# Zinc transporter somatic gene mutations cause primary aldosteronism

**DOI:** 10.1101/2022.07.25.501443

**Authors:** Juilee Rege, Kazutaka Nanba, Sascha Bandulik, Carla Kosmann, Amy R. Blinder, Pankaj Vats, Chandan Kumar-Sinha, Antonio M. Lerario, Tobias Else, Yuto Yamazaki, Fumitoshi Satoh, Hironobu Sasano, Thomas J. Giordano, Tracy Ann Williams, Martin Reincke, Adina F. Turcu, Aaron M. Udager, Richard Warth, William E. Rainey

## Abstract

Primary aldosteronism (PA) is the most common form of endocrine hypertension and effects one in 50 adults. PA is characterized by inappropriately elevated aldosterone production via renin-independent mechanisms. Driver somatic mutations for aldosterone excess have been found in approximately 90% of aldosterone-producing adenomas (APAs). Using next-generation sequencing, we identified recurrent in-frame deletions in *SLC30A1* in five APAs (p.L51_A57del, n=3; p.L49_L55del, n=2). *SLC30A1* encodes the ubiquitous zinc efflux transporter ZnT1 (zinc transporter 1). The identified *SLC30A1* variants are situated in close proximity of the zinc-binding site (H43 and D47) in transmembrane domain II and likely cause abnormal ion transport. PA cases with the unique *SLC30A1* mutations showed male dominance and demonstrated increased aldosterone and 18-oxo-cortisol concentrations. Functional studies of the mutant SLC30A1^51_57del^ variant in a doxycycline-inducible adrenal cell system revealed abnormal Na^+^ conductivity caused by the mutant, which in turn led to the depolarization of the resting membrane potential, and thus to the opening of voltage-gated calcium channels. This resulted in an increase in cytosolic Ca^2+^ activity, which stimulated *CYP11B2* mRNA expression and aldosterone production. Collectively, these data implicate the first-in-field zinc transporter mutations as a dominant driver of aldosterone excess in PA.

## Manuscript Text

Primary Aldosteronism (PA), the most common form of endocrine hypertension, accounts for 5-8% of hypertension cases^1-6^ and for 11-20% of resistant hypertension^7-9^. PA is associated with higher cardiovascular and renal morbidity and mortality compared to essential hypertension of similar severity^10-14^. Biochemically, PA is characterized by inappropriate renin-independent production of aldosterone^15^. Most patients exhibit sporadic forms of PA^16^. The two major subtypes of PA are unilateral aldosterone-producing adenomas (APAs), and bilateral hyperaldosteronism (BHA)^2,17^.

Next-generation sequencing (NGS) has identified somatic mutations in genes encoding ion channels and pumps in APAs. These variants lead to adrenal cell membrane depolarization, facilitate intracellular calcium influx, and promote aldosterone synthase (CYP11B2) expression, which, in turn, drives dysregulated, renin/angiotensin II-independent aldosterone production. Mutated genes described so far include: *KCNJ5* (encoding the Kir3.4 (GIRK4) potassium channel), *ATP1A1* (encoding a Na_+_/K_+_ ATPase alpha subunit), *ATP2B3* (encoding a Ca^2+^ ATPase), *CACNA1D* (encoding a voltage-dependent L-type calcium channel), *CACNA1H* (encoding a voltage-dependent T-type calcium channel alpha-1H subunit), and *CLCN2* (encoding a voltagegated chloride channel ClC-2)^18-25^ In 2018, we developed a CYP11B2-targeted amplicon sequencing method, using immunohistochemistry (IHC) guidance on formalin-fixed paraffin-embedded (FFPE) tissue. Compared to previous genetic studies that utilized gross dissection of fresh frozen APAs, the CYP11B2-guided approach boosted the detection of aldosterone-driver mutations in ∼90% of the APAs^26,27^.

In the present study, whole-exome sequencing (WES) **(Methods secction)** was performed on DNA from two FFPE APAs without known aldosterone-drive mutations and matched adjacent adrenal tissue using our aforementioned CYP11B2 IHC-guided targeted NGS approach^26,27^. An in-frame p.L51_A57del (Leucine, L; A, Alanine) deletion mutation was identified in the *SLC30A1* gene in two APAs **(APA_UM109 and APA_UM110, Supplemental Tables 1 and 2)**. Subsequent targeted NGS **(Online Methods)** using an updated panel and Sanger sequencing identified *SLC30A1* in-frame deletions in three other APAs (APA_S19, APA_S1, and APA_LMU1) that had previously tested negative for known aldosterone-driver mutations **(Supplemental Table 3, Figure 1a,)**; one of these APA harbored the p.L51_A57del variant identified by our WES data, while a novel overlapping deletion (p.L49_L55del) was detected in the other two APAs. *SLC30A1* encodes the zinc efflux transporter ZnT1 (zinc transporter 1), and importantly, no *SLC30A1* mutations have been previously reported in APA-rendering this a first-in-field protein class (zinc transporter) cause of PA. Somatic mutations in *SLC30A1* were identified in 5 of 204 APAs analyzed with targeted NGS, resulting in a prevalence of 2.45%.

**Figure 1.**
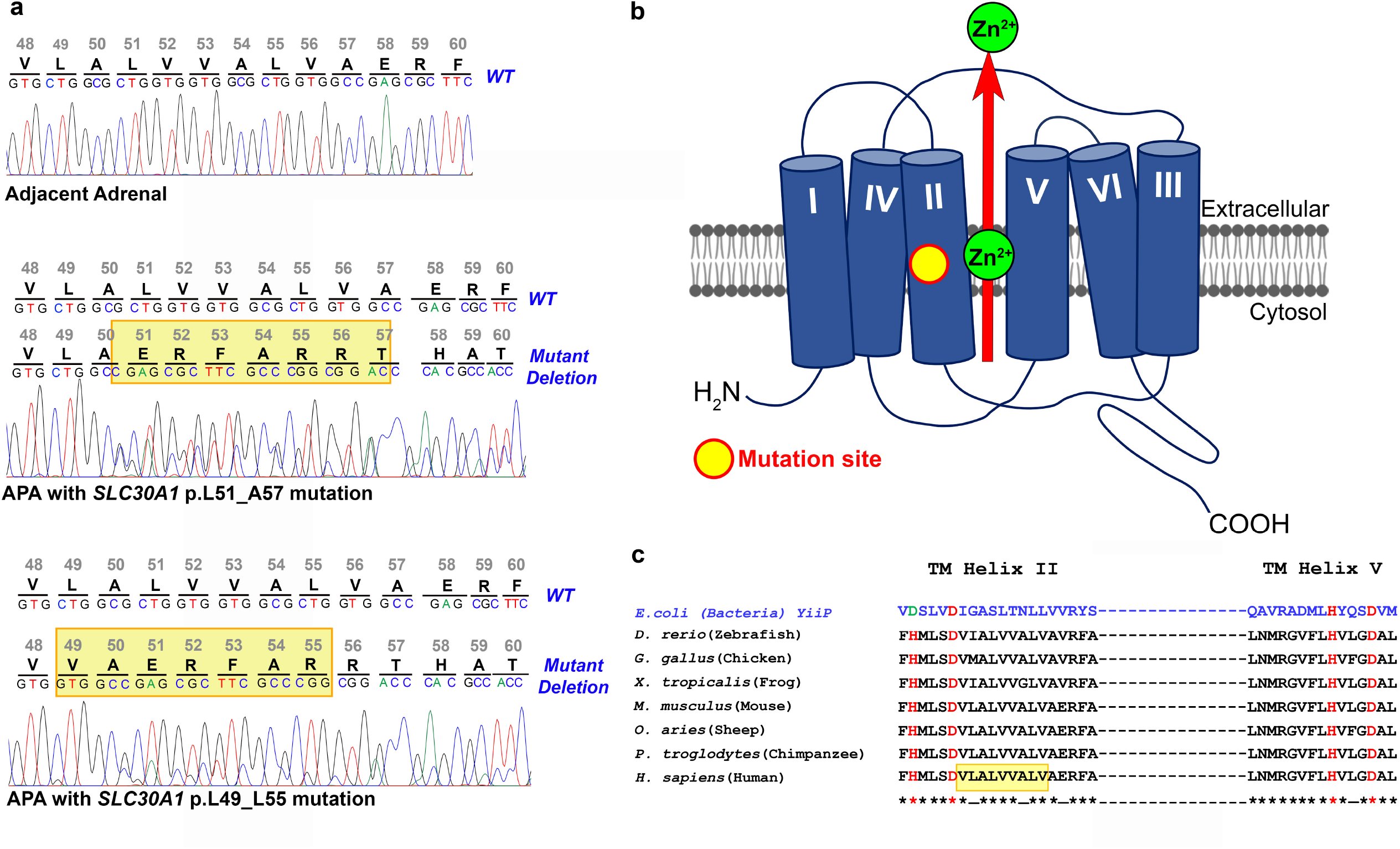
*SLC30A1* mutations in APAs and primary aldosteronism. **(a)** Sanger chromatogram sequences of CYP11 B2-expressing tumor region and adjacent adrenal FFPE genomic DNA for *SLC30A1* codons 48-60 in APAs with the p.L51_A57del and p.L49_L55del mutations. The top row of nucleotides denotes the nucleotide trace in the matched adjacent adrenal tissue obtained by Sanger sequencing. Mutations are present in only the tumor region. **(b)** Schematic structure of ZnT1, the zinc efflux transporter encoded by the *SLC30A1* gene, adapted from Kambe et al^46^. The proposed topology of the transporter is based on the X-ray structure of the *E. coli* homolog, YiiP. The TM helices I, II, IV, and V likely form a compact four-helix bundle. The conserved zinc-binding motif (HD-HD) lies in TM helices II and V. The yellow circle indicates the positions where somatic alterations have been described in APAs, lies in the close vicinity of the zinc-binding site. Zn^2+^, zinc. **(c)** Alignment and conservation of residues encoded by ZnT1 among orthologous species. The conserved residues have been indicated by an asterisk (*). The conserved zinc-binding motif (HD-HD) in TM helices II and V has been indicated in red. The yellow box imposed in the human *H. sapiens* ortholog indicates the region of the APA alterations between codons 49 and 57.

As shown in **Table 1**, APAs harboring somatic *SLC30A1* mutations were all from men, between 59 and 73 years old, with PA diagnosed. All patients had severe clinical and biochemical PA, with resistant hypertension, hypokalemia, and dramatically elevated aldosterone-to-renin ratio [208-1350 (ng/dL)/(ng/mL/h)]. All but one showed a solitary adrenal tumor on cross-sectional imaging studies and adrenal venous sampling lateralization on the side harboring the nodule. Presence of CYP11B2 expression in tumor cells was confirmed by IHC in all cases **(Figure 2)**. The expression of CYP17A1 was observed to vary from scanty to moderate **(Figure 2)**. Owing to abnormal cellular co-expression of CYP17A1 and CYP11B2 in APAs harboring *KCNJ5* mutations, several studies using serum samples from PA patients have demonstrated that along with aldosterone, the hybrid steroids 18-hydroxycortisol (18OHF) and 18-oxocortisol (18oxoF), are a hallmark of these APAs^28-36^. Aldosterone, 18OHF, 18oxoF, and cortisol concentrations were measured in peripheral serum samples from 3/5 patients and 30 age-matched male controls without PA **(Supplemental Table 4)**^37^. Aldosterone,18OHF, and 18oxoF were significantly elevated in the PA patients with *SLC30A1*-mutated APAs vs. controls.

**Table 1.** Clinical characteristics of patients with *SLC30A1*-mutated APA

| Sample ID | APA_UM109 | APA_UM110 | APA_S19 | APA_S21 | APA_LMU1 |
| --- | --- | --- | --- | --- | --- |
| <u>Baseline characteristics</u> |  |  |  |  |  |
| Age at surgery | 66 | 67 | 73 | 62 | 59 |
| Sex | M | M | M | M | M |
| Race | White | White | Asian | Asian | NA |
| *SLC30A1* somatic variant | p.L51_A57del | p.L51_A57del | p.L49_L55del | p.L51_A57del | p.L49_L55del |
| Blood pressure (mmHg) | 137/72 | 185/79 | 138/82 | 145/93 | 162/95 |
| Number of anti-hypertensive medications | 7 | 4 | 1 | 3 | 6 |
| Serum potassium (mEq/L) | 3.9 | 3.1 | 3.8 | 3.0 | 2.5 |
| Potassium supplementation | Yes | Yes | Yes | Yes | Yes |
| PAC (ng/dL) | 37.0 | 135 | 41.6 | 44.4 | 94.3 |
| PRA (ng/mL/h) | 0.1 | 0.1 | 0.2 | 0.1 | NA |
| DRC (mU/L) | NA | NA | NA | NA | 2 |
| <u>Imaging study</u> |  |  |  |  |  |
| Diameter of adrenal tumor (mm) | 13 | 21 | Not detected | 14 | 18 |
| Side of adrenal tumor | L | L | - | L | L |
| Side of aldosterone excess on AVS | L | L | R | L | L |
| Side of adrenalectomy | L | L | R | L | L |
| <u>Post-surgical follow-up data</u> |  |  |  |  |  |
| Follow-up timing | 2 weeks | 1 month | 6 months | 6 months | 6-12 months |
| Blood pressure (mmHg) | 137/63 | 140/65 | 128/81 | 150/91 | 138/81 |
| Number of anti-hypertensive medications | 1 | 3 | 1 | 0 | 0 |
| Serum potassium (mEq/L) | 5.1* | 4.9 | 4.6 | 3.9 | 4.4 |
| PAC (ng/dL) | 7.2 | < 3.0 | 6.7 | 4.4 | 5.5 |
| PRA (ng/mL/h) | 0.3 | < 0.1 | 0.4 | 0.1 | NA |
| DRC (mU/L) | NA | NA | NA | NA | 2 |

PAC, plasma aldosterone concentration; PRA, plasma renin activity; DRC, direct renin concentration; AVS, adrenal venous sampling; M, man; W, woman; \*, potassium supplementation was discontinued at this visit

**Figure 2.**
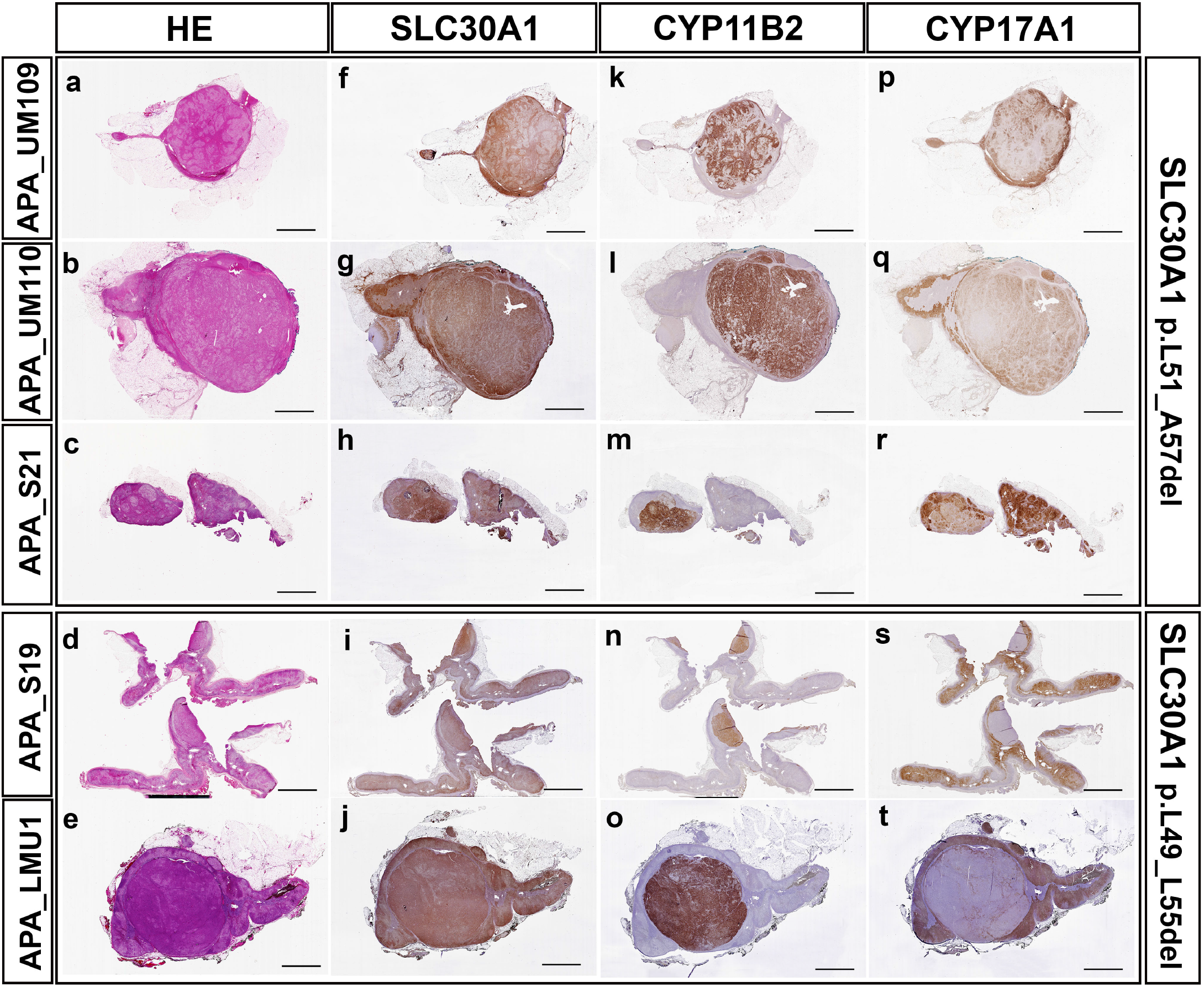
Histopathologic characteristics of aldosterone-producing adenomas (APAs) with somatic *SLC30A1* mutations. Scanned images of the tumors following hematoxylin and eosin staining **(a-e)**, and immunohistochemistry for zinc transporter 1 (SLC30A1/ZnT1) **(f-j)**, aldosterone synthase (CYP11B2) **(k-o)**, and 1?a-hydroxylase/17,20 lyase (CYP17A1) **(p-t)**. Scale: 5 mm.

The *SLC30A1* gene is located on chromosome 1q32.2 (hg19), on the reverse strand. ZnT1, the zinc efflux transporter encoded by *SLC30A1*, is one of ten members of a zinc transporter (ZnT) family, and the most extensively studied. ZnT1 is the most ubiquitously expressed and the sole zinc transporter localized to the cellular plasma membrane^38,39^. ZnT1 functions by exporting cytosolic zinc into the extracellular space, thereby modulating cellular zinc homeostasis, and safeguarding multiple tissues and cell types against zinc toxicity^40,41^. Consistent with its critical role in zinc homeostasis, homozygous *Znt1* knockout mice are embryonically lethal^42^. While no information is available on the structure of any of the ZnT proteins, the crystal structure of YiiP, a bacterial homolog, has been studied extensively^43-45^. The prediction of the structure and function of all the ZnTs is based on the YiiP model. Similar to YiiP, ZnT proteins are predicted to function as metal transporters and demonstrate six transmembrane (TM) helices (except ZnT5), of which TM helices I, II, IV, and V probably form a four-helix bundle harboring a signature metal-binding site^41,46^ **(Figure 1b)**. While the metal-binding site in the TM helices II and V consists of a DD-HD (Aspartic acid, D; Histidine, H) site in the bacterial YiiP, allowing zinc and cadmium transport, the HD-HD motif in the ZnT1-9 confers zinc-specific transport^43,45-47^ **(Figure 1c)**. The ND-HD (Asparagine, N) site in ZnT10 allows for manganese-specific transport. The zinc-specific HD-HD motif has been entirely conserved in proteins encoded by homologs from invertebrates to humans **(Figure 1c)**. Generally, ZnT proteins function as homodimers^18,48-51^, except ZnT5 that forms heterodimers with ZnT6^4,46,48,50^. Another hallmark of ZnT proteins is a histidine-rich loop between TM helices IV and V on the cytosolic side. It is suggested that this loop acts as a sensing mechanism for cytosolic zinc content and could regulate the entry of zinc to the HD-HD site in the four-helix bundle^52-54^. The *SLC30A1* variants identified in the APAs in the present study (p.L51_A57del and p.L49_L55del) are situated in close proximity to the zinc-binding site (H43 and D47)^55-57^ in TM II **(Figure 1c)** suggesting a possible loss of ion transport or cause an abnormal gain-of-function, as described for other PA-associated mutant proteins (e.g. KCNJ5, ATP1A1, ATP2B3).

We investigated the differential expression of *SLC30A1* transcript levels in multiple human tissues. Real time quantitative PCR (qPCR) analysis revealed that the *SLC30A1* mRNA expression is indeed ubiquitous, with the highest transcript levels present in the adrenal cortex **(Supplemental Figure 1a)**. Within the adrenal cortex, unlike CYP11B2, that demonstrated localization to the zona glomerulosa (ZG), *SLC30A1* was abundantly expressed throughout the cortex and did not exhibit zone-specific expression **(Supplemental Figure 1b and 1c)**. We further investigated the histological characteristics of the *SLC30A1*-mutated APAs **(Figure 2)**. All mutant *SLC30A1* APAs demonstrated abundant immunoreactivity for CYP11B2 and SLC30A1 (ZnT1). Unlike CYP11B2, however, SLC30A1 (ZnT1) was also expressed in the adjacent adrenal tissue.

We previously demonstrated the utility of IHC-guided tissue capture followed by targeted next-generation RNA sequencing (RNAseq) to characterized transcriptomic patterns of adrenal zones and adrenal cortical tumors^58,59^. In the current study, we performed targeted RNAseq on the five *SLC30A1*-mutated APAs and compared their gene expression profiles to adrenal cortical zones [zona glomerulosa (ZG), zona fasciculata (ZF), and zona reticularis (ZR)], and other adrenal cortical tumors [APAs with mutations in genes other than *SLC30A1*, cortisol-producing adenoma (CPA) and adrenal cortical carcinoma (ACC)], as described previously^58,59^. Principal component analysis (PCA) and a heatmap of differentially expressed genes (DEGs) among the adrenal cortical zones showed that *SLC30A1*-mutated APAs clustered with the aldosterone-producing ZG (**Supplemental Figure 2a and 2b)**. For example, *SLC30A1*-mutated APAs demonstrated elevated expression of *CYP11B2*, as well as the ZG markers *VSNL1, NOV* and *KCNJ5*, and the WNT pathway-related genes *WNT4, LGR5* and *RSPO3* **(Supplemental Figure 2b)**. Similarly, a PCA plot and DEG heatmap showed that *SLC30A1*-mutated APAs cluster with other APAs of different genotypes and had a distinct gene expression profile from CPA and ACC **(Supplemental Figure 3a and 3b)**. For example, compared to CPA and ACC, *SLC30A1*-mutated APAs demonstrated elevated expression of *CYP11B2, ALDH1A2, RELN, CABP7*, and *VPREB3* **(Supplemental Figure 3b)**.

To characterize the functional impact of the wild type (WT) *SLC30A1*^*WT*^ and the mutant *SLC30A1*^*51*^*-*^*57del*^ transporters on adrenal aldosterone production, we developed human adrenocortical cell lines using HAC15 cells containing a *CYP11B2* promoter-driven secreted Gaussia luciferase reporter (HAC15-B2Luc)^33^,6^0^, with doxycycline (Doxy)-mediated conditional expression of these transporters **(Supplemental Figure 4a)**. Doxy treatment increased *SLC30A1* transcript levels by over 45-fold after 24h **(Supplemental Figure 4b)**. Confocal microscopy confirmed cell membrane localization of *SLC30A1*^*WT*^ and the mutant *SLC30A1*^*51*^*-*^*57del*^ after 24h of Doxy-induction **(Supplemental Figure 4c)**. While Doxy-induction of *SLC30A1*^*WT*^ did not increase either basal *CYP11B2* transcript levels or *CYP11B2* promoter activity, the *SLC30A1*^*51*^*-*^*57del*^ variant increased both over time **(Figure 3a and b)**. In addition, the Doxy induction of mutant protein increased expression of Nurr1 *(NR4A2)*, an upstream regulator of *CYP11B2* transcription^61,62^ **(Figure 3c)**. RNAseq analysis confirmed the augmentative effect of *SLC30A1*^*51*^*-*^*57del*^ on *SLC30A1, CYP11B2*, and *NR4A2* expression levels of the Doxy-induced variant at 24h vs. 0h of treatment **(Supplemental Table 4)**. The transcriptomic analysis of *SLC30A1*^*WT*^ vs. *SLC30A1*^*51*^*-*^*57del*^ cells after 24h Doxy-induction further corroborated the increments in *CYP11B2* and *NR4A2* mRNA expression with the mutant but not wild-type protein **(Figure 3d and e; Supplemental Table 5)**. Moreover, Doxy-activation of the *SLC30A1*^*51*^*-*^*57del*^ gene led to a 14-fold elevation in aldosterone production **(Figure 3f)**.

**Figure 3.**
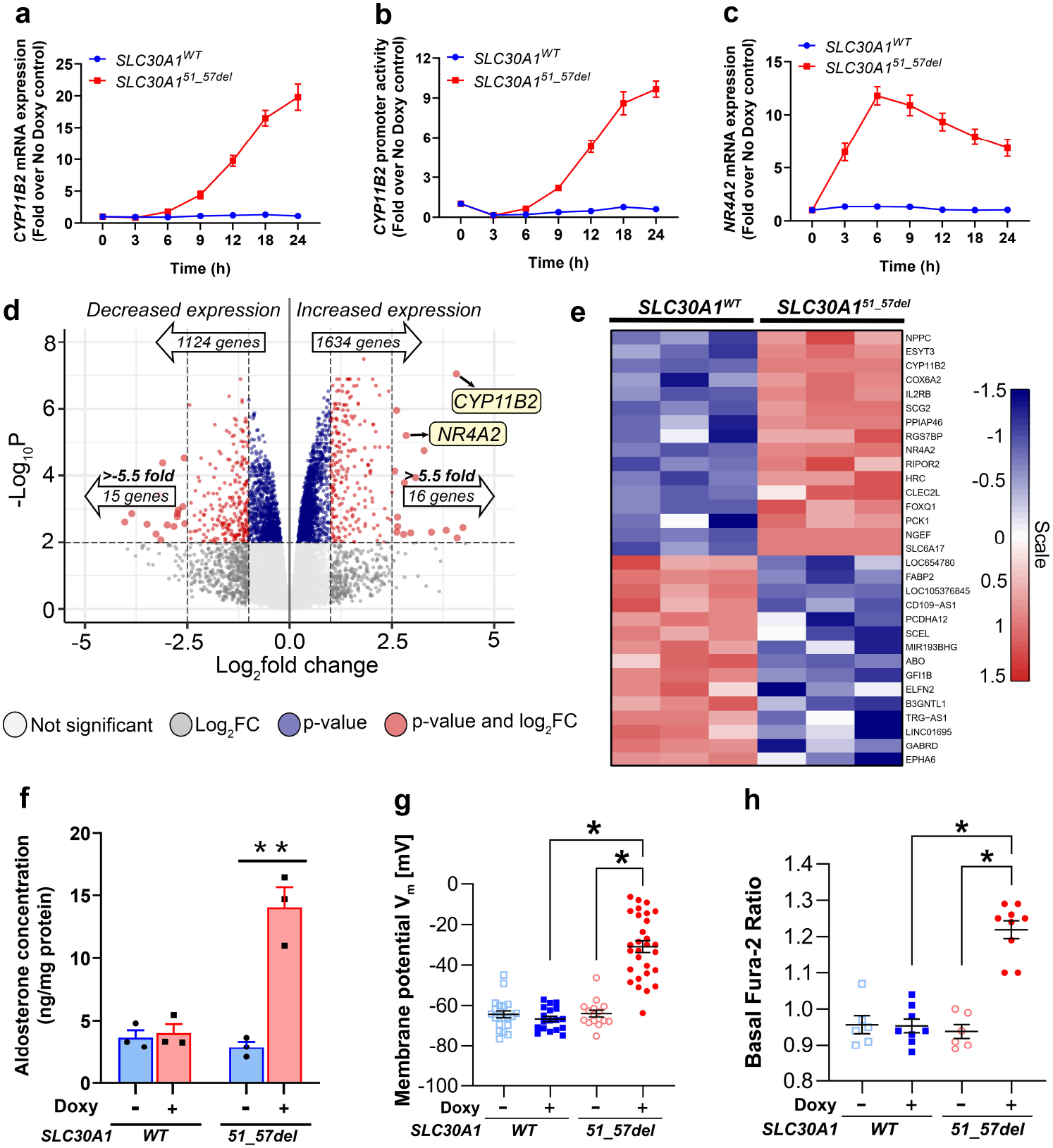
Doxycycline (Doxy)-induced activation of *SLC30A1*^*WT*^ *and SLC30A1*^*51*^*-*^*57del*^ engineered in HAC15-B2Luc cells. Cells were treated with 1 pg/mL Doxy for the indicated times. Doxy causes a time-dependent induction of **(a)** *CYP11B2* mRNA, **(b)** *CYP11B2* promoter activity, and **(c)** *NR4A2* mRNA. Quantitative RT-PCR was used for mRNA transcript detection and gaussian luciferase assay for *CYP11B2* promoter activity, **(d)** Volcano plot and **(e)** Heatmap analyses following RNA sequencing of HAC15-B2Luc cells stably transfected with inducible *SLC30A1*^*m*^ and *SLC30A1*^*51*^*-*^*57del*^, post 24h of Doxy treatment. *CYP11B2* and *NR4A2* show a robust increase in expression in mutant cells vs WT cells. Genes for the volcano plot and heatmap analyses are filtered according to adjusted P < 0.01. Doxy also causes a substantial increase in **(f)** aldosterone production in mutant cells vs WT cells. **, P < 0.005. **(g)** Cell membrane potential of HAC15-B2Luc cells with and without Doxy-induction of *SLC30A1*^*WT*^ and *SLC30A1*^*51*^*-*^*57del*^. Membrane potential was measured by whole-cell patch-clamp 25-33 h after Doxy-induction of *SLC30A1*. Data are presented as mean ±SEM. In addition, single cell data is shown by dots. Overexpression of mutant *SLC30A1*^*51*^*-*^*57del*^ caused a significant depolarization of the resting membrane potential compared to cells overexpressing *SLC30A1*^*WT*^ or to *SLC30A1*^*51*^*-*^*57del*^ cells without Doxy induction. V_m_, membrane potential; *, P < 0.05. (h) Basal cytosolic calcium activity in HAC15-B2Luc cells with and without doxy-induction of *SLC30A1*^*WT*^ and *SLC30A1*^*51*^*-*^*57del*^. The mean of the ratio (± SEM) of the Fura-2 fluorescence intensity at 490-530 nm after excitation at 340 and 380 nm was calculated as a measure of relative cytosolic calcium activity. Data were obtained 25-33 h after Doxy-induction of *SLC30A1*. Overexpression of mutant *SLC30A1*^*51*^*-*^*57de]*^ caused a significant increase of cytosolic calcium compared to cells overexpressing *SLC30A1*^*m*^ or to *SLC30A1*^*51*^*-*^*57del*^ cells without induction. *, P < 0.05.

To investigate the underlying pathological mechanism of mutant SLC30A1, we measured membrane potential and cytosolic Ca^2+^ activity, parameters crucial for the physiological control of aldosterone production in adrenal cells. Similar to native ZG cells, HAC15 cells possess negative resting membrane potential^55,56^. Following Doxy treatment, patch-clamping studies indicated a robust depolarization of the resting membrane potential in the *SLC30A1*^*51*^*-*^57del^-harboring HAC15-B2Luc cells (−31.0 ± 2.9 mV, n=29), as opposed to the *SLC30A1*^*WT*^*-* expressing HAC15-B2Luc cells (−66.3 ± 1.3 mV, n=18) **(Figure 3g)**. The depolarized membrane potential in the *SLC30A1*^*51*^*-*^*57de*^*’* cells was accompanied by increased calcium activity **(Figure 3h)**. Pathogenesis of APAs with mutations in *KCNJ5, ATP2B3* and *CACNA1H* has been attributed to depolarization of the membrane potential due to abnormal Na+ influx^18,63,64^. Therefore, we investigated whether increased Na+ conductance could also play a role in the pathophysiology of mutant *SLC30A1*^*51*^*-*^*57de*^*’*. Overexpression of *SLC30A1*^*51*^*-*^*57de*^*’* caused a significant abnormal inward current (visible at -120 mV up to -60 mV voltage clamp steps), which was absent in control cells **(Figure 4a-d)**. The removal of extracellular Na+, which was replaced by impermeable N-methyl-D-glucamine (NMDG+), largely reduced the voltage-activated transient inward currents in the mutant cells to the level of the control cells **(Figure 4d)**. We also observed that the membrane potential of Doxy-induced *SLC30A1*^*51*^*-*^*57de*^*’* cells, which was depolarized under control conditions (−37.3 ± 2.4 mV), became hyperpolarized under Na_+_ free condition (−66.8 ± 1.9 mV) **(Figure 4e)**. Lastly, flame photometric measurements of the Na_+_ content in cell lysates demonstrated a robust elevation in intracellular Na_+_ in Doxy-induced *SLC30A1*^*51*^*-*^*57de]*^ cells **(Figure 4f)**. This indicates that the abnormal inward current in the mutant cells is causing Na_+_ influx that cannot be compensated by increased Na_+_ export. Treatment with the Na_+_/K_+_-ATPase inhibitor Ouabain also led to an increase in Na_+_ in the control cells by inhibiting Na_+_ export. In the Doxy-induced *SLC30A1*^*51*^*-*^57del^ cells, the increase in intracellular Na+ content was exacerbated with Ouabain, presumably as a result of the abnormal Na+ influx **(Figure 4f)**. These data provide strong evidence for an abnormal Na_+_ conductance caused by the expression of mutant *SLC30A1*^*51*^*-*^57del^. This imbalance in the electrochemical equilibrium causes depolarization of the cell membrane and the activation of voltage-dependent Ca^2+^ channels leading to the eventual downstream transcriptional activation of *CYP11B2* and increased aldosterone production. This hypothesis was further strengthened by verapamil (L-type calcium channel blocker) inhibition of *SLC30A1*^*51*^*-*^57del^-induced adrenal cell *NR4A2* and *CYP11B2* expression, and of aldosterone production **(Supplemental Figure 5)**. Interestingly, several studies have demonstrated that *SLC30A1/ZnT1* can downregulate L-type calcium channel activity in multiple cell types, by decreasing the trafficking of its a1-subunit to the cell surface membrane^38,39,55,65-67^. Recently, neuronal *SLC30A1/ZnT1* was shown to function as a zinc/calcium exchanger^68^, and a zinc/H_+_ exchanger in the HEK293 kidney cells^57^. The functional versatility of *SLC30A1*, in conjunction with our current observations, provides ample scope for studying the role of *SLC30A1* in adrenal aldosterone production.

**Figure 4.**
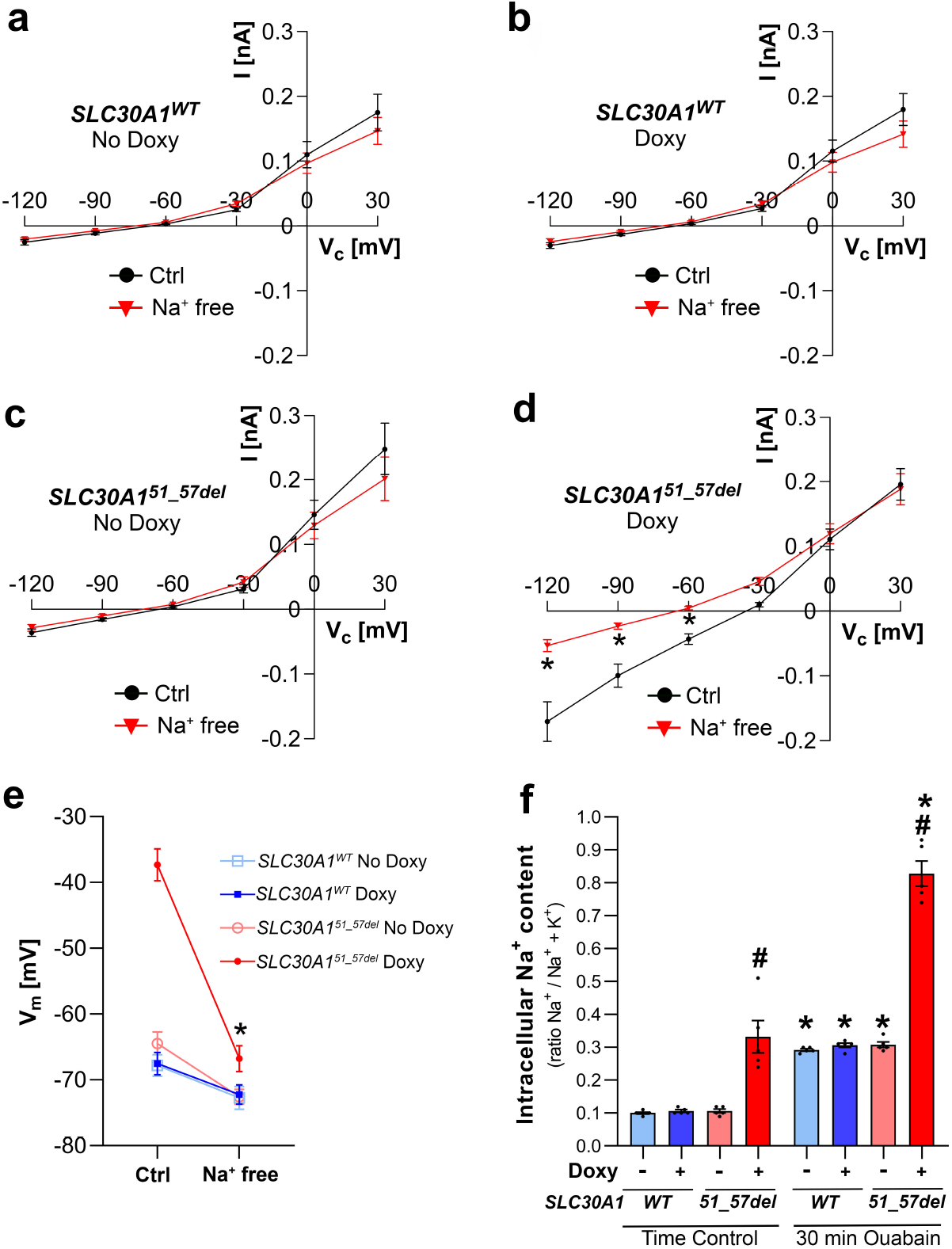
Effect of extracellular Na^+^ removal on the ion currents and the cell membrane potential after Doxy-induction of *SLC30A1*^*WT*^ and *SLC30A1*^*s1*^*-*^*57del*^ in HAC15-B2Luc cells, and intracellular Na^+^ content at basal condition and upon inhibition of the Na7K^+^-ATPase using Ouabain. **(a-d)** Ion currents and membrane potential were measured by whole-cell patch-clamp 25-33 h after induction of *SLC30A1*^*WT*^ *and SLC30A1*^*51*^*-*^*57del*^ with 1 pg/mL doxycycline. Data are presented as mean ±SEM. Overexpression of mutant *SLC30A1*^*51*^*-*^*57del*^ caused an abnormal inward current, which was absent in cells overexpressing *SLC30A1*^*WT*^ or *SI_C30A1*^*51*^*-*^*57del*^ cells without Doxy induction. This inward current was largely reduced to the level of control cells upon replacement of extracellular Na^+^ by impermeable N-methyl-D-glucamine (Na+ free), **(e)** The membrane potential of induced *SLC30A1*^*51*^*-*^*57del*^ cells, which was depolarized under control condition, became hyperpolarized under Na^+^ free condition. *, P < 0.05, Control (Ctrl) vs Na^+^ free, **(f)** Intracellular Na^+^ content of cell lysates was measured using flame photometry and is presented here as mean ±SEM of ratio comparing Na+ concentration to the sum of Na^+^ and potassium (K^+^) concentration. Doxy-induced mutant *SI_C30A1*^*51*^*-*^*57del*^ cells exhibited increased intracellular Na^+^ content, vs. Doxy-induced *SLC30A1*^*WT*^cells and *SLC30A1*^*51*^*-*^*57del*^ cells without Doxy-induction. Inhibition of the sodium pump Na7K^+^-ATPase with Ouabain (10 pM for 30 min) resulted in increased intracellular Na+ concentration in all groups compared to conditions without Ouabain. However, the elevation in intracellular Na+ concentration was even more pronounced in Doxy-induced mutant *SLC30A1*^*51*^*-*^*57del*^ cells. #, P < 0.05, Doxy-treated *SLC30A1*^*51*^*-*^*57del*^ cells vs. Doxy-treated *SLC30A1*^*WT*^ cells and cells without Doxy-induction. *, P < 0.05, No Ouabain vs. Ouabain-treated.

In summary, we identified recurrent somatic mutations in a zinc transporter in a subset of APAs. The mutant *SLC30A1* (ZnT1) causes abnormal Na_+_ conductance, which leads to depolarization of the cell membrane and an increase in cytosolic Ca^2+^ activity, triggering autonomous aldosterone production. **(Figure 5)**. The discovery of a first-in-field protein class (zinc transporter) causing PA gives insight into the structure-function relationship of this class of proteins and extends our understanding of the molecular pathogenesis of APAs.

**Figure 5.**
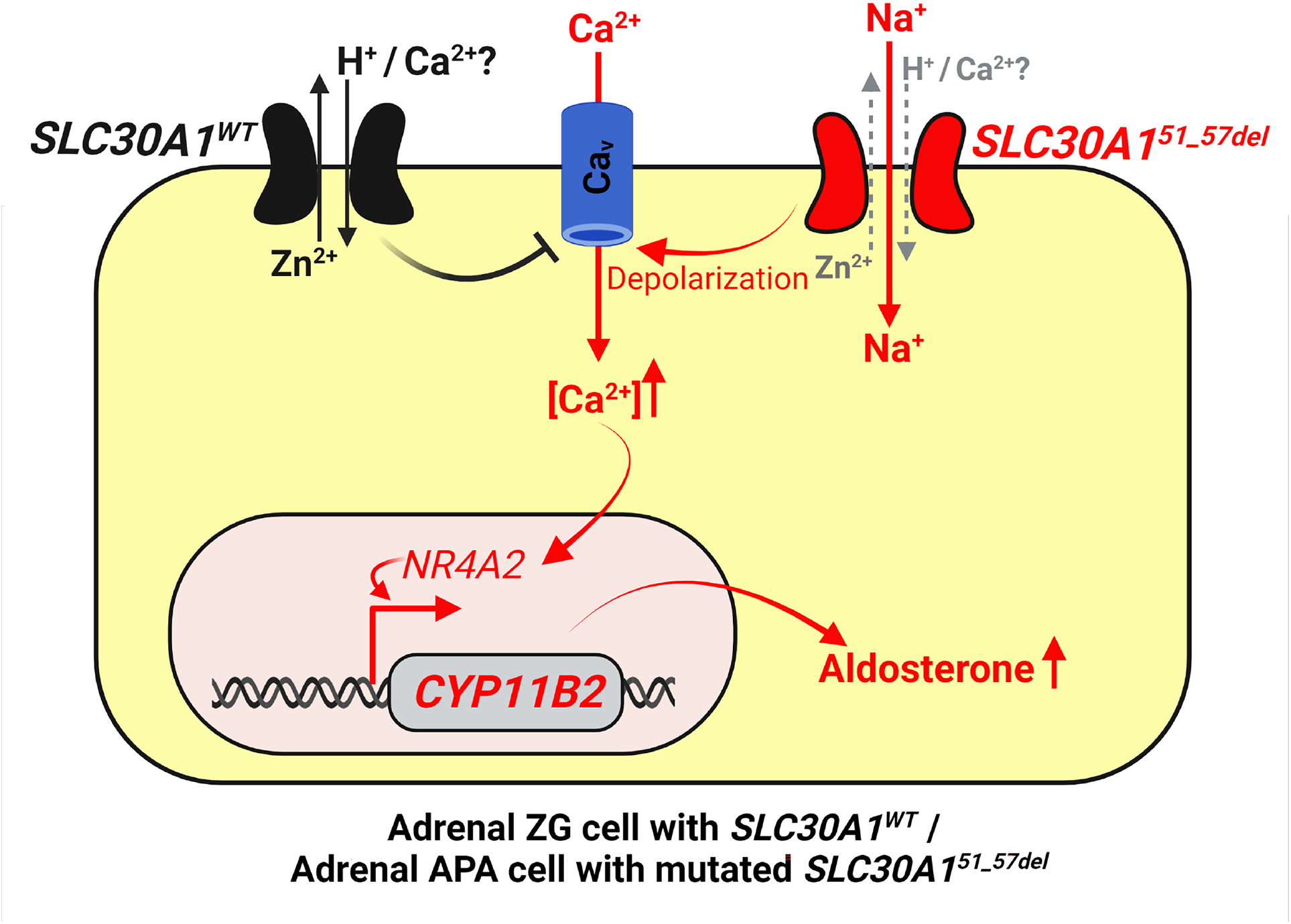
Proposed model for autonomous aldosterone secretion in adrenal zona glomerulosa cells with the *SLC30A1*^*51*^*-*^*57del*^ mutant. The *SLC30A1*^*WT*^ has been shown to (1) downregulate voltage-gated calcium channel activity, (2) function as a zinc/calcium exchanger, and/or (3) function as a zinc/H^+^exchanger in multiple cell types. The *SLC30A1*^*51*^*-*^*57del*^ variant causes abnormal Na^+^ influx which leads to cell depolarization which in turn, activates the voltage-gated calcium channels stimulating calcium influx. Increased calcium signal leads to elevated expression of *NR4A2* and *CYP11B2*, and eventually aldosterone production. [Ca^2+^], intracellular calcium concentration; Na^+^, sodium ion; Ca_v_, voltage-gated calcium channels. Created with BioRender.com.

## Methods

Methods, including statements of data availability and associated references, are available in the online version of the paper.

## Acknowledgements

We thank Michelle Vinco and Farah Keyoumarsi at the University of Michigan for assistance in slide preparation. We are also thankful to Elena Klayman, Marcin Cieslik, Fengyun Su, Rui Wang, Xuhong Cao, and Arul M. Chinnaiyan at the University of Michigan for support with whole-exome sequencing and Chia-Jen Liu at the University of Michigan for technical support with the targeted NGS. We would also like to acknowledge Dr. Richard J. Auchus at the University of Michigan for support with the LC-MS/MS facilities, and Florian Gürtler and Eva Wacker at the University of Regensburg for their technical assistance with the electrophysiological studies.

This work was supported by grants from NIDDK (R01DK106618 and R01DK043140 to W.E.R), NCATS/ Michigan Institute for Clinical and Health Research (UL1TR002240 to J.R.), NHLBI (1R01HL130106 to T.E, and R01HL155834 to A.F.T.), the Doris Duke Charitable Foundation (2019087 to A.F.T), the American Heart Association (17SDG33660447 to K.N. and 20CDA35320016 to J.R.), the Ministry of Health, Labour, and Welfare, Japan (20FC1020 to H.S.). and the Deutsche Forschungsgemeinschaft (WA 1275/6 to R.W., BA 4436/2-1 to S. B., CRC/TRR 205 “The Adrenal Gland”, RE 752/31-1 and WI 5359/2-1 “Project 444776998” to M.R. and T. A.W). M.R. also received funding from the Else Kröner-Fresenius Stiftung (2012_A103, 2015_A228, and 2019_A104 “Else-Kröner Hyperaldosteronismus-German Conn Registry”), and the European Research Council (Grant # 694913). The lentiviral preparation was performed at the University of Michigan Vector Core and was supported by a grant from NIDDK to the University of Michigan Center for Gastrointestinal Research (5P30DK034933).

## Author contributions

J.R., K.N., S.B. and W.E.R designed experiments. J.R., K.N., S.B. and W.E.R. wrote the manuscript. P.V. and C.K-S performed and analyzed whole-exome sequencing. A.R.B performed IHC studies and tissue DNA/RNA extraction. A.M.U. analyzed targeted NGS data from tissue samples. J.R., A.R.B. and W.E.R performed and analyzed the results of *in vitro* studies on HAC15-B2Luc cells. A.M.L. analyzed RNAseq data in the engineered HAC15-B2Luc cells. S.B., C.K. and R.W. performed electrophysiological studies, calcium measurements, IF staining and flame photometry measurements of cells. A.F.T and J.R. performed LC-MS/MS studies and analyzed the results. A.F.T., T.E., Y.Y., F.S., H.S., T.J.G., T.A.W., M.R. assisted with patient recruitment, medical care, specimen collection, and clinical data acquisition. All authors reviewed and revised the manuscript draft.

## Methods

### Patients

Patients diagnosed with primary aldosteronism (PA), who underwent unilateral adrenalectomy at the University of Michigan, Tohoku University and Ludwig Maximilian University of Munich, were studied. The diagnosis of PA was established as per the Endocrine Society’s clinical practice guideline^1^ or the institutional consensus at that time. The availability of archival formalin-fixed paraffin-embedded (FFPE) blocks of resected tumors determined the inclusion of the patients in the study. Normal adrenal glands were obtained from kidney transplantation donors at the University of Michigan. This study was approved by the institutional review boards of the University of Michigan (HUM00083056 and HUM00069665), Tohoku University (2021-1-482) and Ludwig Maximilian University of Munich (379-10). Baseline clinical characteristics of these individuals are summarized in **Table 1**. The approval allows for the NGS analysis of somatic variants on archival tissue. Therefore, we will share limited NGS data on request.

### Immunohistochemistry (IHC) and immunofluorescence (IF) of adrenal FFPE tissue

Hematoxylin and eosin (H&E) staining, CYP11B2 and 17α-hydroxylase/17,20 lyase (CYP17A1) IHC were performed as previously described^2,3^. 5 µm APA FFPE sections were deparaffinized and epitope retrieval was performed by heating samples for 15 min in a pH 9 buffer (Vector Laboratories Inc., Burlingame, CA). Following peroxidase (10 min) and serum blocking [10% goat serum in Triton X-100 phosphate buffered saline (TBS) for 1 hour), FFPE sections were incubated at room temperature for 1 hour with a mouse monoclonal antibody against CYP11B2 (clone 41-17B; diluted 1:1250; Millipore Sigma; Cat # MABS1251)^4^ and a rabbit polyclonal antibody against CYP17A1 (diluted 1:2000; LifeSpan Biosciences, Cat # LS-B14227)^5^; and overnight at 4°C with a rabbit polyclonal antibody against SLC30A1/ZnT1 (diluted 1:250; LifeSpan Biosciences, Cat # LS-B14285)^6^. The Polink-2 HRP Plus Mouse DAB System (GBI Labs, Bothell, WA) was used for detection. Slides were counterstained with Harris hematoxylin for 10-20 seconds followed by dehydration and coverslipping. After confirmation of appropriate staining under a light microscope, slides were electronically scanned by the PathScan Enabler IV (Meyer Instruments, Houston, TX).

Dual immunofluorescence was performed for CYP11B2 (mouse monoclonal antibody, Millipore Sigma) and SLC30A1/ZnT1 (rabbit polyclonal antibody,). After blocking (1:10 goat serum in TBS for 1 hour), FFPE tissue sections were incubated with the primary antibody solutions overnight (1:1250 dilution for both CYP11B2 and SLC30A1/ZnT1 antibodies), washed with TBS, and subsequently incubated with secondary fluorescent antibodies for 1 hour (Alexa fluor 488-conjugated goat anti-mouse and Alexa fluor 594 – conjugated goat antirabbit, both diluted 1:100, Jackson Immunoresearch, West Grove, PA). Prolong Gold mounting medium with DAPI (Life Technologies, Carlsbad, CA) was used to visualize the cell nucleus. Immunofluorescence was viewed under an inverted microscope (Leica, Wetzlar, Germany).

### gDNA and RNA isolation from adrenal FFPE tissue

5 µm serial sections were prepared from the APA and normal adrenal FFPE blocks. For each APA case, slides 1, 2, and 3 were used for H&E staining, CYP11B2 IHC, and CYP17A1 IHC, respectively as described previously^7,8^. Using a sterile scalpel, the subsequent 8-9 unstained FFPE slides were dissected based on the CYP11B2 IHC results for CYP11B2-positive tumor regions under an Olympus SZ-40 microscope (Olympus, Tokyo, Japan). The adrenal tissues adjacent to the tumor region was also separately dissected.

For the normal adrenal, CYP11B2, VSNL1 (visinin-like 1), HSD3B2 (3²-hydroxysteroid dehydrogenase type 2), CYB5A (cytochrome b5A), and TH (tyrosine hydroxylase) IHC were used to guide capture of zona glomerulosa (ZG), zona fasciculata (ZF) and zona reticularis (ZR), respectively as previously described^9^.

gDNA and RNA were isolated from the captured FFPE tissue using the AllPrep DNA/RNA FFPE kit (Qiagen, Germantown, MD) as described previously^7,8^.

### Whole Exome Sequencing (WES) on gDNA From FFPE tissue

WES was performed^10,11^ on gDNA from two APAs that were found to be CYP11B2-positive but negative for aldosterone-driver mutations based on targeted next-generation sequencing (NGS) analysis^7,8^. The matched germline DNA from the adjacent adrenal tissue of the aforementioned APA was also included in the WES analysis. WES was performed using standard protocols in our Clinical Laboratory Improvement Amendments (CLIA)-compliant sequencing lab^12,13^. 500 ng of gDNA was sheared using a Covaris S2 ultrasonicator to a peak target size of 250 base pairs (bp). Concentration of the fragmented DNA was performed with AMPure beads, followed by end-repair, A-base addition, ligation of the Illumina indexed adapters, and size selection on 3% Nusieve agarose gels (Lonza, Basel, Switzerland). Illumina index primers and AMPure beads were used to amplify and purify fragments between 300 to 350 bp. 1 µg of the library was hybridized to the Agilent SureSelect Human All Exon v.4. The targeted exon fragments were captured and enriched following the manufacturer’s protocol (Agilent, Santa Clara, CA). Analysis of the paired-end whole-exome libraries was performed by the Agilent 2100 Bioanalyzer and DNA 1000 reagents and sequencing was performed with the Illumina HiSeq 2500 sequencing system (Illumina, San Diego, CA). The primary base call files were converted into FASTQ sequence files using the bcl2fastq converter tool bcl2fastq-1.8.4 in the CASAVA 1.8 pipeline.

### Ion Torrent-based NGS on gDNA From FFPE tissue

Validation of the *SLC30A1* variant identified by WES was performed by targeted NGS of using a custom AmpliSeq DNA panel and the Ion Torrent NGS System (Thermo Fisher Scientific, Waltham, MA). Targeted regions included the complete coding sequences of *KCNJ5, CACNA1D, CACNA1H, ATP1A1, ATP2B3, CLCN2, SLC30A1*, and *CTNNB1*. NGS library preparation, sequencing, and identification of somatic variants was performed as described previously^2,7,14^.

### Sanger sequencing

Further validation of the *SLC30A1* mutation status was done by direct bidirectional Sanger sequencing of the PCR-amplified gDNA from the APAs identified with an *SLC30A1* alteration by WES or targeted NGS. Sanger sequencing was also carried out on the matched adjacent adrenal tissue. gDNA was PCR-amplified using the Promega GoTaq Flexi DNA Polymerase following the manufacturer’s protocol (Promega, Madison, WI). An annealing temperature of 63.9°C was applied for 35 cycles. The gDNA in APA-adjacent adrenal tissue pairs was sequenced following PCR amplification using specific *SLC30A1* primers. Forward: 5’-GCGCTGACCTTCATGTTCATGGTG-3’; Reverse: 5’-GCGAAACAGAGGCCAGTCAGGAAG-3’.

### Targeted next-generation RNA sequencing (RNAseq)

Targeted RNAseq was performed as previously described^9,15^. Briefly, up to 15 ng of RNA was reverse transcribed using SuperScript VILO (Thermo Fisher Scientific, Waltham, MA), and NGS libraries were generated for each sample from the resulting cDNA using the Ion AmpliSeq Library Kit Plus (Thermo Fisher Scientific) and a custom adrenal-specific AmpliSeq RNA panel targeting 194 genes^9,15^. Differential gene expression was visualized using heatmaps constructed with Cluster 3.0 and TreeView 1.1.6r4, and principal component analysis was carried out using normalized gene expression values and the prcomp function of the stats package in R (version 4.1.2) prior to visualization in Microsoft Excel (Redmond, WA).

### Serum steroid quantification by liquid chromatography-tandem mass spectrometry (LC-MS/MS)

LC-MS/MS was utilized to quantify aldosterone, cortisol, 18OH-cortisol and 18oxo-cortisol in a single assay. Steroid extraction and LC-MS/MS quantitation was performed as previously described^16,17^.

### Generation of vectors and lentiviruses

The human wild type (WT) *SLC30A1* gene cDNA ORF in the pcDNA3.1-C-(k)DYK backbone (OHu13086; NM_021194.3) was purchased from GenScript (Piscataway, NJ). GenScript also performed site-directed deletion on the WT clone to obtain the *SLC30A1* variant p.L51_A57del in pcDNA3.1-C-(k)DYK. The in-frame deletion of the nucleotide sequence 5’-CTGGTGGTGGCGCTGGTGGCC-3’ resulted in the deletion of the amino acids L51-V52-V53-A54-A55-V56-A57 in the *SLC30A1* (p.L51_A57del) clone. GenScript also provided services to sub-clone WT and the mutant *SLC30A1* amplicons into the shuttle vector pENTR1A-GFP-N2 (FR1) (Addgene plasmid # 19364)^18,19^. The resultant *SLC30A1* shuttle vectors were then inserted into the doxycycline (Doxy)-inducible lentiviral vector pCLX-pTF-R1-DEST-R2-EBR65 (Addgene plasmid # 45952)^20,21^ to generate the lentivectors pCLX-pTF-Blast-*SLC30A1*^*WT*^ and pCLX-pTF-Blast-*SLC30A1*^*51*^*-*^*57del*^ respectively. Lentiviruses (∼1 × 10^6^ TU/mL) were generated at the University of Michigan Biomedical Vector Core and used for transducing adrenal cells.

### Lentiviral transduction and cell experiments

The human adrenocortical carcinoma cell line HAC15 containing a *CYP11B2* promoter-driven secreted Gaussia luciferase reporter (HAC15-B2Luc)^22,23^, was used as parental cells (provided by Dr. Celso Gomez-Sanchez, University of Mississippi Medical Center). The HAC15-B2Luc cells were grown in DMEM-F12 containing 10% cosmic calf serum (CCS), 1% insulin-transferrin-selenium, and antibiotics (penicillin, streptomycin and gentamicin) and maintained in a 37°C humidified atmosphere (5% CO_2_), as previously described^23^. These cells were transduced^10,23,24^ with the *SLC30A1*^*WT*^ and *SLC30A1*^*51*^*-*^*57del*^ lentiviruses to generate the cell lines HAC15B2Luc-Doxy-*SLC30A1*^*WT*^ and HAC15B2Luc-Doxy-*SLC30A1*^*51*^*-*^*57del*^ respectively, followed by antibiotic selection with geneticin (Gaussia luciferase selection marker, 1 mg/mL) and blasticidin *(SLC30A1* selection marker, 2 µg/mL). A mixed population of the respective geneticin/blasticidin-selected HAC15B2Luc cells were used to analyze the functional effects of the WT gene and the variant.

For experiments, cells were plated at a density of 75,000 cells/well in a 48-well dish for 48 hours. After incubation in a low serum medium (DMEM/F-12 containing 0.1% CCS and antibiotics) for 24 hours, expression of *SLC30A1*^*WT*^ and *SLC30A1*^*51*^*-*^*57del*^ in the cells was initiated by addition of 1 µg/mL Doxy (Sigma Aldrich, St. Louis, MO) for the indicated times. For the inhibitor studies, cells were incubated with 10 µM verapamil (EMD Millipore, Temecula, CA), for 30 min before, and during the course of Doxy treatment. At the end of each treatment, the medium was collected from each well and stored at -20°C until aldosterone quantitation and gaussia luciferase assay, whereas the cells were frozen at -80°C for or RNA isolation for cDNA generation, quantitative real-time RT-PCR (qPCR) and/or RNAseq, as well as protein assay. All experiments were conducted in at least triplicate.

### Adrenal cell RNA isolation, RNAseq and qPCR

Total RNA was extracted from cells using RNeasy plus mini kit (Qiagen, Valencia, CA). The purity and integrity of the RNA were checked spectroscopically using a Nano Drop spectrometer (Nano Drop Technologies). Total RNA was applied to downstream processes such as quantitative RT-PCR (qPCR) and/or RNAseq (LC Sciences, Houston, TX).

100 ng of RNA was reverse transcribed using the High-capacity cDNA Archive kit (Life Technologies). For qPCR, 5 ng of cDNA was mixed with Taqman Fast Universal Master Mix (Life Technologies) following manufacturer’s recommendations. qPCR was performed using a StepOne Plus Fast Real-Time PCR system (Applied Biosystems). The primer-probe sets for *CYP11B2* were designed in-house and purchased from Integrated DNA Technologies^25,26^. The primer-probe sets for Peptidylprolyl isomerase A *(PPIA*, Cyclophilin A), *SLC30A1* and Nurr1 *(NR4A1)* were purchased from Life Technologies^26^. Quantitative normalization of cDNA in each tissue-derived sample was performed using the expression of *PPIA* as an internal control. Relative quantification was determined using the comparative threshold cycle method. Values of all experiments are presented as the Mean±SEM of three independent experiments performed in experimental triplicate for each condition.

For cellular RNAseq, sequencing metrics such as quality scores and number of reads were assessed with *fastQC* (https://www.bioinformatics.babraham.ac.uk/projects/fastqc/). Read adapters were trimmed with the *bbduk* tool from *bbtools* (https://sourceforge.net/projects/bbmap/). Paired-end reads were aligned to the human transcriptome reference sequence (Release 40, obtained from gencodegenes.org) using *kallisto*^*27*^. Downstream analyses were performed in R using Bioconductor packages as briefly described: Expression values were summarized at the gene level using the lengthScaledTPM method from *tximport*^*28*^. Inter-experiment gene level expression data was scaled to library size with *edgeR* using the TMM method^29^. Non-expressed genes were filtered out with *filterByExpr*function from *edgeR*. Unwanted and hidden sources of variation were removed from the data using *sva*^30^. Differential gene expression analysis was performed using *limma*^*31*^. Heatmaps and volcano plots were built using *pheatmap* and *EnhancedVolcano*, respectively [Raivo Kolde (2019). pheatmap: Pretty Heatmaps. R package version 1.0.12. https://CRAN.R-project.org/package=pheatmap; Kevin Blighe, Sharmila Rana and Myles Lewis (2021). EnhancedVolcano: Publication-ready volcano plots with enhanced colouring and labeling. R package version 1.12.0. https://github.com/kevinblighe/EnhancedVolcano].

### Aldosterone measurements

The aldosterone content of the cell culture medium was analyzed using an inhouse competitive ELISA kit^32,33^. Briefly, high-binding 96-well plates were coated with 2µg/100µl goat anti-mouse IgG (Sigma Aldrich, Cat # M2650-1ML)^34^ and incubated overnight at 4°C. Post washing with PBS and Tween, the wells were coated with an aldosterone antibody (Aldo A2E11-6G1, kindly provided by Dr. Celso Gomez-Sanchez)^35^. This was followed by the addition of aldosterone standards and cell media samples in the respective wells. The wells were then incubated with aldosterone conjugated to horseradish peroxidase and the plate was shaken at room temperature for 2 hours. The contents of the wells were discarded, and the wells were washed multiple times with PBS and Tween. The substrate solution containing tetramethylbenzidine (TMB) was the added to all wells and the plate was incubated on a shaker for 30 min. Finally, the reaction was stopped with 1N sulfuric acid. The absorbance of each well was measured at 450 nm within 15 minutes of stopping the reaction. A standard curve was then constructed by plotting the mean absorbance obtained from each standard against its concentration with absorbance value on the vertical (Y) axis and concentration on the horizontal (X) axis. The concentration of aldosterone in the samples was determined by extrapolating the mean absorbance value for each sample against the corresponding concentrations obtained from the calibration curve. Values of all experiments are presented as the Mean±SEM of three independent experiments performed in experimental triplicate for each condition. Each dot (n=3) represents the Mean or Average of each individual experiment performed in triplicate.

### Protein extraction and protein assay

Cells were lysed in 100 µL mammalian protein extraction reagent (Pierce Chemical Co), and the protein content was estimated by the bicinchoninic acid protein assay following manufacturer’s protocol (Thermo Scientific).

### Gaussia luciferase assay for *CYP11B2* promoter activity

For the gaussia luciferase luminescence bioassay, 25 µL of medium was mixed with 50 µL of 0.01 mg/mL coelenterazine (GoldBio, St. Louis, MO). The coelenterazine solution was prepared in an assay buffer comprising of 50 mM Tris HCl (pH 7.5) and 150 mM of NaCl. Luminescence was then measured by the Tecan Spark Microplate Reader (Tecan, Männedorf, Switzerland).

### Immunocytochemistry

Doxy-inducible HAC15B2Luc-Doxy-*SLC30A1*^*WT*^ and HAC15B2Luc-Doxy-*SLC30A1*^*51*^*-*^*57del*^ cells were seeded onto fibronectin/collagen-coated glass cover slips in 35 mm cell culture dishes in growth medium. After 2-3 days, growth medium was replaced with a low serum medium. After an additional day, expression of wild type or mutant *SLC30A1* was induced by 1 µg/mL doxycycline in low serum medium. For control (No Doxy) cells, low serum medium without doxycycline was used. After 24 hours, cells were washed twice with Ringer solution and then fixed for 15 min with a solution containing 3% paraformaldehyde, 3.4% sucrose, 90 mM NaCl, 15 mM K_2_HPO_4_, 1 mM EGTA and 2 mM MgCl_2_, pH 7.4. Between the following steps, cells were washed each time for 5 min in PBS. Cells were treated for 5 min with 0.1% SDS-containing PBS to permeabilize the cells and unmask epitopes. Non-specific binding sites were blocked by incubation for 15 min in PBS containing 5% bovine serum albumin (BSA) and 0.04% Triton-X 100. Primary antibody (rabbit polyclonal SLC30A1/ZnT1, Novus Biologicals, Centennial, Colorado, USA, Cat # NBP1-86825)^36^ was applied for 1 hour at room temperature diluted 1:400 in PBS with 1% BSA and 0.008% Triton-X 100. Secondary antibody (donkey-anti-rabbit Alexa 488 nm, Life Technologies GmbH, Darmstadt, Germany) was applied for 1 hour at room temperature diluted 1:400 in PBS with 1% BSA and 0.008% Triton-X 100 together with HOE33258 (1.25 µM, for staining of cell nuclei). Finally, the glass coverslips with the cells were mounted on slides with fluorescent-free glycergel mounting medium (DakoCytomation, Hamburg, Germany). The staining was then analyzed with a confocal microscope (LSM 710; Zeiss, Jena, Germany) using sequential scanning (Plan Apochromat 63x/1.3 water objective) with an optical slice thickness of 1 µm.

### Electrophysiology/Patch-clamp measurements

Doxy-inducible HAC15B2Luc-Doxy-*SLC30A1*^*WT*^ and HAC15B2Luc-Doxy-*SLC30A1*^*51*^*-*^*57de/*^ cells were seeded onto fibronectin/collagen-coated glass cover slips in 35 mm cell culture dishes in growth medium. After 2-3 days, growth medium was replaced by medium with low serum. After an additional day, expression of wild type or mutant *SLC30A1* was induced by addition of 1 µg/mL doxycycline in low serum medium. For control (No Doxy) cells, low serum medium without doxycycline was used. 25-33 hours after induction of *SLC30A1* expression, cell membrane potential was measured by patch-clamp technique. Whole cell patch recordings were performed at room temperature using an EPC 10 amplifier (Heka, Lambrecht, Germany) and a Powerlab Data Acquisition System (ADInstruments GmbH, Spechbach, Germany). The PatchMaster v2×50 software (Heka, Lambrecht, Germany) and the LabChartPro v7 software (ADInstruments GmbH, Spechbach, Germany) were used for data acquisition and analysis. Cell membrane potential was measured in current clamp mode with subtraction of 10 mV liquid junction potential. Inward and outward currents were measured in voltage clamp mode by a series of clamp steps (from -120 to +30 mV incremented in 30 mV steps). Patch pipettes with 5-10 MΩ were used for the recordings. The patch pipette solution contained (mM): 95 K-gluconate, 30 KCl, 4.8 Na_2_HPO_4_, 1.2 NaH_2_PO_4_, 5 glucose, 2.38 MgCl_2_, 0.726 CaCl_2_, 1 EGTA, 3 ATP, pH 7.2. The extracellular Ringer-type control solution contained (mM): 145 NaCl, 0.4 KH_2_ PO_4_, 1.6 K_2_HPO_4_, 5 glucose, 1 MgCl_2_, 1.3 CaCl_2_, 5 HEPES, pH 7.4. For Na^+^-free solution, Na^+^ was replaced by cell impermeable N-methyl-D-glucamine (NMDG^+^). The measurements were carried out at different days and from different cell preparations using different cell passages to ensure the reproducibility of the experiments. each cell represents an independent experiment because the stimulation protocol for individual cells was started separately.

### Na^+^ and K^+^ content measurements in cell lysates using flame photometry

Doxy-inducible *SLC30A1*^*WT*^ and *SLC30A1*^*51*^*-*^*57de/*^ HAC15-B2Luc cells were seeded in 12-well cell culture plates in serum (10%) containing medium (8,000,000 cells/well). After 2-3 days, growth medium was replaced by medium with low serum. After an additional day, expression of wild type or mutant *SLC30A1* was induced by addition of 1 µg/mL doxycycline in low serum medium. For control (No Doxy) cells, low serum medium without doxycycline was used. The next day, half of the wells was supplemented with 10 µM Ouabain to inhibit Na+/K+ ATPase. After 30 min, the plates were placed on ice, cell culture medium was removed, and the cells were quickly washed 3 times with Na^+^- and K^+^-free 300 mM Mannitol solution to remove Na^+^ and K^+^ that may have left from the extracellular medium. Cell swelling and rupture was then induced by incubation in 300 µL deionized water (Milli-Q) on ice for 2 hours, followed by placing the plates in an ultrasonic water bath for 1 min. Subsequently, cell lysates were further homogenized by freezing and thawing. Finally, the cell lysates were clarified from cell debris by centrifugation at 14,800 rpm for 10 min. The supernatant (250 µL) was diluted 1:17 in deionized water to measure the Na^+^ and K^+^ concentration with flame photometry (BWB-XP Technologies, UK). The relative cellular Na^+^ content was calculated as the ratio of the Na^+^ concentration divided by the sum of the Na^+^ and K^+^ concentrations. The measurements were carried out at different days and from different cell preparations using different cell passages to ensure the reproducibility of the experiments.

### Calcium measurements

Cytosolic free calcium activity was measured at 37°C using the ratiometric fluorescent Ca^2+^ sensitive dye Fura-2-AM (Life Technologies GmbH, Darmstadt, Germany). Cells were loaded at room temperature for 60 min with 3 µM Fura-2-AM in the presence of 1X Power Load permeabilizing reagent (Life Technologies GmbH, Darmstadt, Germany). The extracellular Ringer-type control solution contained (mM): 145 NaCl, 0.4 KH_2_PO_4_, 1.6 K_2_HPO_4_, 5 glucose, 1 MgCl_2_, 1.3 CaCl_2_, 5 HEPES, pH 7.4. Fluorescence ratios of emission at 490-530 nm after excitation at 340 nm and 380 nm were calculated for single cells after subtraction of the background signal using the Axiovision software (Zeiss, Jena, Germany). Mean fluorescence ratios of 5-18 cells per dish were then calculated from a measurement period of 5 min. The measurements were carried out at different days and from different cell preparations using different cell passages to ensure the reproducibility of the experiments. The averaged signals per dish represent the number of experiments since several cells per dish were measured at the same time (n=5 to 18).

### Statistical Analysis

For cell culture experiments, data were analyzed in Prism software (GraphPad, San Diego, CA) using one-way ANOVA or two-way ANOVA, as appropriate, plus Tukey post-test to calculate the level of significance. Differences between the groups were considered significant for p<0.05.

### URLs

FastQC (https://www.bioinformatics.babraham.ac.uk/projects/fastqc/); Read adapters trimming with the *bbduk* tool from *bbtools* (https://sourceforge.net/projects/bbmap/). pheatmap: Pretty Heatmaps. R package version 1.0.12. https://CRAN.R-project.org/package=pheatmap; EnhancedVolcano: Publication-ready volcano plots with enhanced colouring and labeling. R package version 1.12.0. https://github.com/kevinblighe/EnhancedVolcano.

### Data availability

The data that support the findings of this study are available from the authors upon reasonable request. NGS data are available upon request within a scientific cooperation.

