## Supplemental Material for "Zinc transporter somatic gene mutations cause primary aldosteronism"

Supplemental Figure 1

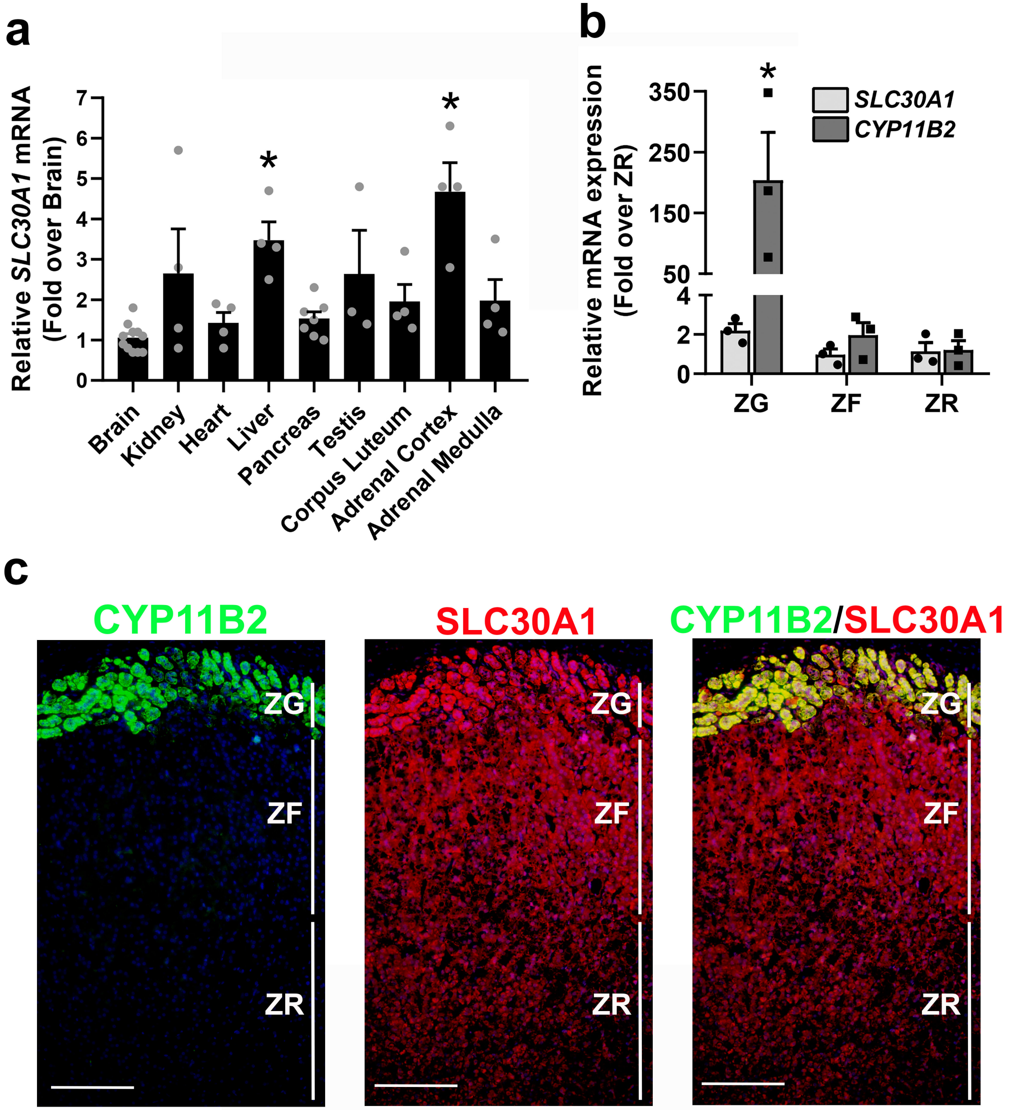

**Supplemental Figure 1. SLC30A1 expression in multiple tissues. (a)** Quantitative RT-PCR analyses of *SLC30A1* in adrenal cortex and medulla in comparison with various tissues. *, P < 0.005 vs. brain. Peptidylprolyl isomerase A (PPIA, Cyclophilin A) was used as the housekeeping gene. Each dot represents individual tissue samples. Comparison of *SLC30A1* and *CYP11B2* **(b)** transcript levels and **(c)** protein expression in the adrenal ZG, ZF and ZR. While CYP11B2 shows a distinct increased expression in the adrenal ZG, SLC30A1 remains uniformly distributed throughout the adrenal cortex. Immunofluorescence scale: 200 µm

Supplemental Figure 2

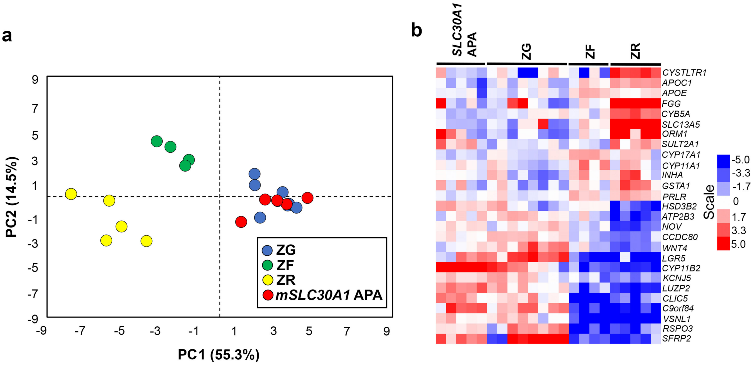

**Supplemental Figure 2. Targeted RNAseq demonstrates clustering of mutant *SLC30A1* APAs with zona glomerulosa (ZG). (a)** PCA plot of F1 and F2 reveals distinct clustering of ZG (n=8), zona fasciculata (ZF, n=4), and zona reticularis (ZR, n=5) samples. It also demonstrates grouping of mutant SLC30A1 APAs (n=5) with the aldosterone-producing ZG. **(b)** Hierarchical clustering of differentially expressed genes shows sample-level grouping, with mutant *SLC30A1* APAs aligning with ZG. Heatmap depicts median-centered gene expression values (red = high and blue = low). The *SLC30A1*-mutated APAs exhibit elevated expression of *CYP11B2*, the ZG markers *VSNL1*, *NOV* and *KCNJ5*, and several WNT pathway-related genes. *mSLC30A1* APA, mutant SLC30A1 APA

Supplemental Figure 3

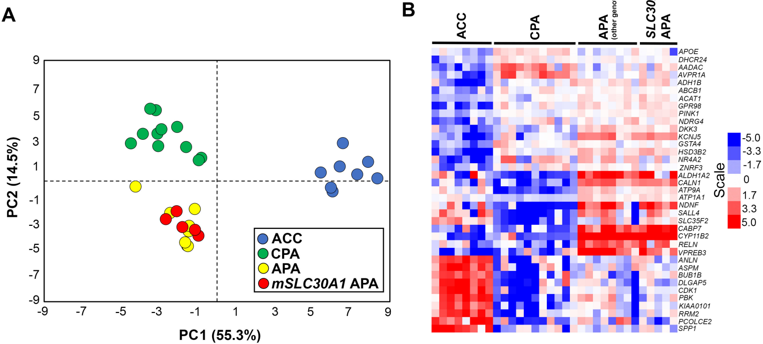

**Supplemental Figure 3. Targeted RNAseq demonstrates clustering of mutant *SLC30A1* APAs with APAs of other genotypes. (a)** PCA plot of F1 and F2 reveals distinct clustering of ACCs (n=8), CPAs (n=11), and APAs of genotypes other than *SLC30A1* [total n=8 (mutated *CACNA1D*, n=2, mutated *KCNJ5*, n=3, mutated *ATP1A1*, n=2; mutated *ATP2B3*, n=1) samples. It also demonstrates grouping of mutant *SLC30A1* APAs (n=5) with other APAs. **(b)** Hierarchical clustering of differentially expressed genes shows sample-level grouping with mutant *SLC30A1* adenomas aligning with APAs harboring other mutations. Heatmap depicts median-centered gene expression values (red = high and blue = low). The *SLC30A1*-mutated APAs demonstrated increased expression of *CYP11B2*, *ALDH1A2*, *RELN*, *CABP7* and *VPREB3*; consistent with the transcriptomic signature of APAs of other genotypes. ACCs, adrenocortical carcinomas; CPAs, cortisol-producing adenomas; *mSLC30A1* APA, mutant *SLC30A1* APA

Supplemental Figure 4

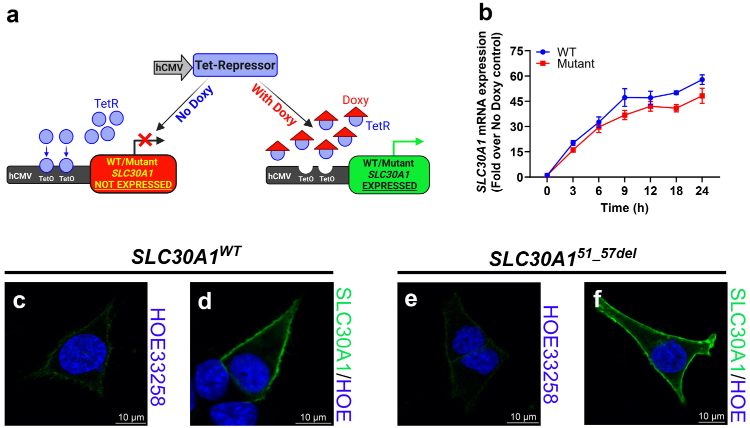

**Supplemental Figure 4. (a)** Overview of the doxycycline (Doxy)-inducible adrenal cell system. **(b)** *SLC30A1*$^{WT}$ and *SLC30A1*$^{51\_57del}$ mRNA and **(c)** protein expression selectively increases in the stably transduced HAC15-B2Luc cell lines in the presence of Doxy. Immunofluorescence scale: 10 µm

Supplemental Figure 5

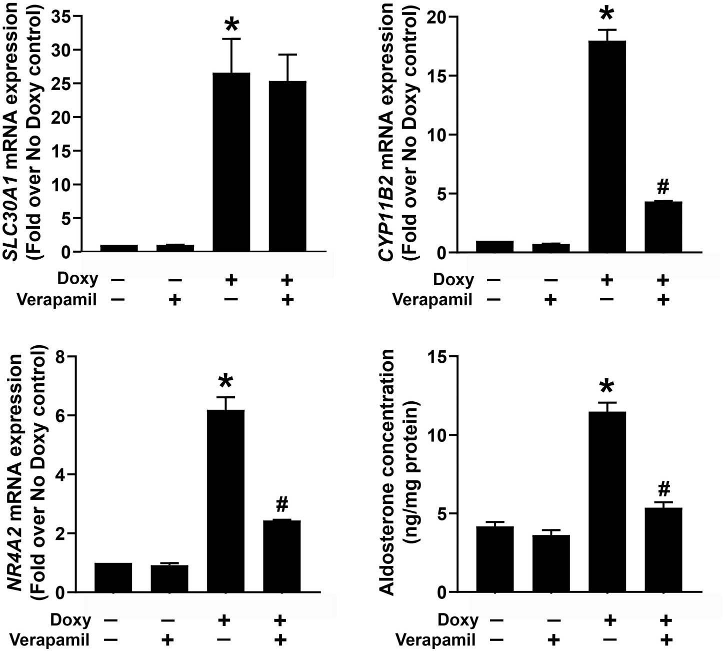

**Supplemental Figure 5.** ***SLC30A1***$^{51\_57del}$ **increases *CYP11B2* mRNA expression and aldosterone production through calcium-mediated pathway.** While pre-incubation with the L-type calcium channel blocker, verapamil (10 μM), did not cause an abrogation in doxy-induced **(a)** *SLC30A1* mRNA, inhibition a dose-dependent inhibition was observed in SLC30A1$^{51_57del}$ -stimulated **(b)** *CYP11B2* mRNA, **(c)** *NR4A2* mRNA and **(d)** aldosterone production. *, P < 0.005, No Doxy vs. Doxy-treated; #, P < 0.005, Doxy-treated vs. Verapamil plus Doxy-treated.

**Supplemental Table 1. List of somatic variants prioritized by whole-exome sequencing in two APAs**

| Sample ID | Gene | Position | Amino acid change | Mutation type | VAF (Tumor) | VAF (Adjacent adrenal tissue) |
|---|---|---|---|---|---|---|
| APA_UM109 | *ARHGAP5* | chr14:32563216 | p.Asp1114Gly | missense | 27/268(10%) | 0/245(0%) |
| APA_UM109 | *FSIP2* | chr2:186662123 | p.Lys3420Asn | missense | 43/329(13%) | 0/268(0%) |
| APA_UM109 | *GPATCH8* | chr17:42478580 | p.Asn289Asp | missense | 12/179(7%) | 0/169(0%) |
| APA_UM109 | *MADCAM1* | chr19:501762 | p.Thr259_Asp266dup | inframe_insertion | 7/40(18%) | 0/25(0%) |
| APA_UM109 | *PHACTR3* | chr20:58318183 | p.Pro47Leu | missense | 31/109(28%) | 0/76(0%) |
| APA_UM109 | *RAB10* | chr2:26357810 | p.Pro175Ala | missense | 73/259(28%) | 0/220(0%) |
| **APA_UM109** | ***SLC30A1*** | chr1:211751784 | **p.Leu51_Ala57del** | **inframe_del** | **117/461(25%)** | **0/402(0%)** |
| APA_UM109 | *TMEM72* | chr10:45430359 | p.Pro202His | missense | 123/428(29%) | 1/325(0%) |
| APA_UM110 | *AP1G2* | chr14:24033862 | p.Thr277Ile | missense | 22/233(9%) | 0/205(0%) |
| APA_UM110 | *ATRN* | chr20:3553565 | p.Glu687Lys | missense | 44/193(23%) | 0/122(0%) |
| APA_UM110 | *C11orf42* | chr11:6231729 | p.Pro242fs | frameshift | 40/345(12%) | 0/265(0%) |
| APA_UM110 | *DCAF1* | chr3:51440672 | p.Ser1404Cys | missense | 33/327(10%) | 1/209(0%) |
| APA_UM110 | *DSPP* | chr4:88535088 | p.Gly425Glu | missense | 28/282(10%) | 0/201(0%) |
| APA_UM110 | *FRAS1* | chr4:79366883 | p.Val1958Ala | missense | 15/297(5%) | 0/192(0%) |
| APA_UM110 | *PI4K2A* | chr10:99433426 | p.Ser456Phe | missense | 38/225(17%) | 0/164(0%) |
| APA_UM110 | *RGS2* | chr1:192780132 | p.Ala99Val | missense | 8/103(8%) | 0/72(0%) |
| **APA_UM110** | ***SLC30A1*** | chr1:211751784 | **p.Leu51_Ala57del** | **inframe_del** | **151/567(27%)** | **0/353(0%)** |
| APA_UM110 | *TACC2* | chr10:123844724 | p.Ser903Arg | missense | 63/521(12%) | 0/413(0%) |
| APA_UM110 | *TMED6* | chr16:69377315 | p.Cys240Ser | missense | 20/245(8%) | 0/136(0%) |

Whole-exome sequencing was performed on two FFPE mutation-negative APAs and their matched germline DNA from the adjacent adrenal tissue. Two of the APAs demonstrated the SLC30A1 (p.L51_A57del) somatic variant. VAF, variant allele frequency.

**Supplemental Table 2. Results of whole exome sequencing.**

| Sample ID | Tissue Type | Total Reads | Mean Coverage per Base | % Alignment to hg19 reference genome |
|---|---|---|---|---|
| APA_UM109 | Tumor | 105940791 | 245.41 | 92.12 |
| APA_UM110 | Adjacent | 82673747 | 195.23 | 92.93 |
| APA_S19 | Tumor | 115255724 | 263.31 | 93.11 |
| APA_S21 | Adjacent | 74028633 | 179.48 | 93.49 |

**Supplemental Table 3. Results of targeted next-generation sequencing.**

| Sample ID | Gene | Nucleotide change | Amino Acid Change | FDP | VAF (%) |
|---|---|---|---|---|---|
| APA_UM109 | *SLC30A1* | c.150_170del | p.L51_A57del | 705 | 69 |
| APA_UM110 | *SLC30A1* | c.150_170del | p.L51_A57del | 638 | 65 |
| APA_S19 | *SLC30A1* | c.145_165del | p.L49_L55del | 547 | 63 |
| APA_S21 | *SLC30A1* | c.150_170del | p.L51_A57del | 481 | 79 |
| APA_LMU1 | *SLC30A1* | c.145_165del | p.L49_L55del | 544 | 60 |

FDP, flow-corrected read depth; VAF, variant allele frequency

**Supplemental Table 4. Serum steroids measured by LC-MS/MS**

| Steroids (pg/mL) | Individual Patients | | | PA patients with *SLC30A1* mutations Median (IQ range) | Age-matched male controls Median (IQ range) |
|---|---|---|---|---|---|
| | APA_UM109 | APA_UM110 | APA_LMU1 | | |
| Aldosterone | 204.9 | 1929.5 | 699.7 | 699.7 (204.9-1930)* | 0.0 (0.0-65.7) |
| Cortisol | 42546 | 122929 | 227751 | 122929 (42546-227751) | 114687 (93770-139119) |
| 18OH-cortisol | 5627 | 10907 | 2104 | 5627 (2104-10907)* | 633.1 (511.8-819.2) |
| 18oxo-cortisol | 62.9 | 1004 | 116.4 | 116.4 (62.9-1004)* | 0.0 (0.0-65.7) |

Steroids were measured by LC-MS/MS. The steroid concentrations (pg/mL) for the individual patients are expressed in the first three data columns. Data are also expressed as median with corresponding interquartile range for the three PA patients with mutant *SLC30A1* APAs. Steroid concentrations were not available for APA_S19 and APA_S21. Age-matched male subjects (n=30) were utilized as controls for statistical comparison. Statistical significance was determined by nonparametric Mann–Whitney U test. *, P<0.005. Note: Aldosterone and 18oxo-cortisol were not detected in 19 and 28 controls respectively.

**Supplemental Table 5.** RNAseq analysis of *SLC30A1* $^{51\_57del}$ at 24 hours vs. 0 hours of Doxy induction

| Entrez ID | Symbol | Ensembl | *SLC30A1* $^{51\_57del}$ Doxy 24h | *SLC30A1* $^{51\_57del}$ Doxy 0h | 24h Doxy vs. 0h Doxy *Log FC* | 24h Doxy vs. 0h Doxy *FC* | P.Value | adj.P.Value (Significance P < 0.01) |
|---|---|---|---|---|---|---|---|---|
| 337 | *APOA4* | ENSG00000110244.7 | -5.35 | -0.12 | 5.22 | 37.35 | 6.11185E-07 | 2.47926E-05 |
| 92749 | *DRC1* | ENSG00000157856.12 | -5.30 | -0.46 | 4.83 | 28.51 | 0.000122591 | 0.001131911 |
| 359 | *AQP2* | ENSG00000167580.8 | 0.10 | 4.90 | 4.80 | 27.91 | 5.09059E-09 | 8.79125E-07 |
| 1585 | *CYP11B2* | ENSG00000179142.2 | 2.90 | 7.66 | 4.76 | 27.01 | 2.2312E-12 | 6.69761E-09 |
| 3560 | *IL2RB* | ENSG00000100385.14 | -4.36 | 0.34 | 4.71 | 26.24 | 5.68813E-05 | 0.000647747 |
| 7779 | *SLC30A1* | ENSG00000170385.10 | 6.75 | 11.39 | 4.64 | 24.91 | 5.69755E-16 | 8.55146E-12 |
| 1339 | *COX6A2* | ENSG00000156885.6 | -4.61 | -0.24 | 4.37 | 20.68 | 0.00020565 | 0.00167477 |
| 4581 | *MTX1P1* | ENSG00000236675.1 | -3.86 | 0.44 | 4.31 | 19.89 | 0.001755668 | 0.008763159 |
| 103611157 | *TGFB2-OT1* | ENSG00000281453.1 | -1.60 | 2.71 | 4.31 | 19.77 | 0.000252971 | 0.001969315 |
| 107984450 | *LOC107984450* | ENSG00000256001.3 | -3.20 | 0.73 | 3.92 | 15.13 | 0.002074028 | 0.009977273 |
| 4880 | *NPPC* | ENSG00000163273.4 | -3.88 | -0.07 | 3.79 | 13.81 | 0.001063958 | 0.005983121 |
| 7857 | *SCG2* | ENSG00000171951.5 | -0.52 | 2.90 | 3.42 | 10.68 | 1.70779E-07 | 1.05483E-05 |
| 25791 | *NGEF* | ENSG00000066248.15 | 4.62 | 7.96 | 3.34 | 10.10 | 2.7716E-10 | 1.38663E-07 |
| 284021 | *MILR1* | ENSG00000271605.6 | -3.15 | 0.11 | 3.25 | 9.49 | 0.000408811 | 0.002866271 |
| 11211 | *FZD10* | ENSG00000111432.5 | -1.49 | 1.70 | 3.19 | 9.10 | 1.6099E-06 | 4.75972E-05 |
| 3400 | *ID4* | ENSG00000172201.12 | -3.18 | -0.10 | 3.09 | 8.49 | 0.001411783 | 0.007432287 |
| 55824 | *PAG1* | ENSG00000076641.4 | -3.63 | -0.57 | 3.08 | 8.45 | 0.000824543 | 0.004879955 |
| 4929 | *NR4A2* | ENSG00000153234.15 | 1.46 | 4.50 | 3.04 | 8.23 | 2.90306E-08 | 3.02584E-06 |
| 3822 | *KLRC2* | ENSG00000205809.11 | -2.31 | 0.74 | 3.03 | 8.19 | 0.001483816 | 0.007727689 |
| 729739 | *PPIAP46* | ENSG00000260266.1 | 0.10 | 2.91 | 2.82 | 7.04 | 9.83472E-06 | 0.000181785 |
| 6863 | *TAC1* | ENSG00000006128.12 | -1.26 | 1.49 | 2.76 | 6.75 | 1.67357E-05 | 0.000265216 |
| 2353 | *FOS* | ENSG00000170345.10 | -0.64 | 2.10 | 2.75 | 6.71 | 0.000403636 | 0.002836223 |
| 5970 | *RELA* | ENSG00000173039.20 | 7.59 | 10.32 | 2.74 | 6.66 | 1.32175E-14 | 9.91905E-11 |
| 26207 | *PITPNC1* | ENSG00000154217.16 | 4.34 | 6.94 | 2.60 | 6.05 | 1.94426E-10 | 1.25652E-07 |
| 1244 | *ABCC2* | ENSG00000023839.12 | -2.40 | 0.16 | 2.58 | 5.96 | 0.001081436 | 0.006054184 |
| 84623 | *KIRREL3* | ENSG00000149571.12 | 1.77 | 4.34 | 2.56 | 5.90 | 9.41269E-08 | 6.82488E-06 |
| 2827 | *GPR3* | ENSG00000181773.7 | 0.08 | 2.62 | 2.55 | 5.84 | 2.02771E-05 | 0.000305254 |
| 26018 | *LRIG1* | ENSG00000144749.14 | -0.02 | 2.50 | 2.52 | 5.74 | 0.000182107 | 0.00151951 |
| 5105 | *PCK1* | ENSG00000124253.11 | -0.93 | 1.58 | 2.51 | 5.71 | 0.000943823 | 0.00543377 |
| 8736 | *MYOM1* | ENSG00000101605.14 | 4.89 | 7.38 | 2.49 | 5.61 | 2.27634E-09 | 5.31861E-07 |
| 94234 | *FOXQ1* | ENSG00000164379.7 | -1.96 | 0.50 | 2.46 | 5.52 | 0.00030048 | 0.002260603 |
| 973 | *CD79A* | ENSG00000105369.10 | 3.66 | 6.11 | 2.45 | 5.45 | 8.55628E-12 | 1.83459E-08 |
| 9750 | *RIPOR2* | ENSG00000111913.20 | -0.11 | 2.33 | 2.43 | 5.39 | 2.60124E-05 | 0.000364538 |
| 11069 | *RAPGEF4* | ENSG00000091428.18 | 4.91 | 7.32 | 2.42 | 5.35 | 8.83752E-09 | 1.27541E-06 |
| 100133941 | *CD24* | ENSG00000272398.6 | -0.72 | 1.69 | 2.40 | 5.29 | 0.000140516 | 0.001257741 |
| 81606 | *LBH* | ENSG00000213626.13 | 3.04 | 5.40 | 2.37 | 5.16 | 4.7168E-07 | 2.10424E-05 |
| 28513 | *CDH19* | ENSG00000071991.9 | -3.50 | -1.13 | 2.36 | 5.13 | 0.00150416 | 0.007795559 |
| 23349 | *PHF24* | ENSG00000122733.12 | -1.80 | 0.53 | 2.33 | 5.02 | 0.001593075 | 0.008136445 |
| 109729181 | *ZNF710-AS1* | ENSG00000259291.2 | 2.06 | 4.38 | 2.32 | 4.98 | 2.75541E-07 | 1.47628E-05 |
| 2303 | *FOXC2* | ENSG00000176692.8 | -0.24 | 2.07 | 2.31 | 4.95 | 6.01714E-07 | 2.47926E-05 |
| 344838 | *PAQR9* | ENSG00000188582.10 | 0.65 | 2.95 | 2.30 | 4.92 | 4.72468E-07 | 2.10424E-05 |
| 10221 | *TRIB1* | ENSG00000173334.4 | 3.35 | 5.62 | 2.27 | 4.83 | 7.65319E-08 | 6.04562E-06 |
| 161253 | *REM2* | ENSG00000139890.10 | -2.89 | -0.66 | 2.22 | 4.66 | 0.001996268 | 0.009696438 |
| 80144 | *FRAS1* | ENSG00000138759.20 | 5.87 | 8.07 | 2.20 | 4.61 | 6.00606E-10 | 2.25362E-07 |
| 3624 | *INHBA* | ENSG00000122641.11 | 2.34 | 4.54 | 2.20 | 4.60 | 8.44223E-09 | 1.24225E-06 |
| 9586 | *CREB5* | ENSG00000146592.17 | 1.61 | 3.81 | 2.20 | 4.59 | 0.000731607 | 0.00446007 |
| 2118 | *ETV4* | ENSG00000175832.13 | 1.13 | 3.33 | 2.19 | 4.57 | 4.46679E-06 | 0.000101272 |
| 4158 | *MC2R* | ENSG00000185231.5 | 3.27 | 5.46 | 2.18 | 4.55 | 7.96547E-07 | 2.95476E-05 |
| 6546 | *SLC8A1* | ENSG00000183023.18 | 1.48 | 3.65 | 2.18 | 4.52 | 0.000135055 | 0.001218311 |
| 1584 | *CYP11B1* | ENSG00000160882.13 | -0.36 | 1.80 | 2.16 | 4.47 | 1.57072E-05 | 0.000254315 |
| 10930 | *APOBEC2* | ENSG00000124701.6 | -2.48 | -0.32 | 2.16 | 4.47 | 0.001853461 | 0.009147843 |
| 6622 | *SNCA* | ENSG00000145335.17 | 3.20 | 5.35 | 2.16 | 4.46 | 1.25613E-06 | 4.076E-05 |
| 3491 | *CCN1* | ENSG00000142871.18 | 4.24 | 6.39 | 2.15 | 4.44 | 1.7731E-09 | 4.8302E-07 |
| 6539 | *SLC6A12* | ENSG00000111181.13 | -1.27 | 0.85 | 2.13 | 4.38 | 0.000300266 | 0.002260126 |
| 57419 | *SLC24A3* | ENSG00000185052.13 | 2.95 | 5.06 | 2.11 | 4.30 | 1.33125E-10 | 1.13323E-07 |
| 23704 | *KCNE4* | ENSG00000152049.7 | 4.87 | 6.94 | 2.07 | 4.20 | 5.78854E-12 | 1.448E-08 |
| 388662 | *SLC6A17* | ENSG00000197106.7 | 2.65 | 4.71 | 2.06 | 4.17 | 1.75054E-05 | 0.000271705 |
| 390937 | *ERFL* | ENSG00000268041.3 | -0.32 | 1.73 | 2.06 | 4.16 | 0.000629783 | 0.00399173 |
| 729475 | *RAD51AP2* | ENSG00000214842.6 | -2.25 | -0.20 | 2.05 | 4.15 | 0.001041546 | 0.005881327 |
| 2239 | *GPC4* | ENSG00000076716.9 | 6.98 | 9.03 | 2.04 | 4.12 | 5.91551E-13 | 2.21965E-09 |
| 359845 | *RFLNB* | ENSG00000183688.4 | 4.33 | 6.37 | 2.04 | 4.10 | 2.73187E-10 | 1.38663E-07 |
| 23462 | *HEY1* | ENSG00000164683.18 | -0.81 | 1.21 | 2.03 | 4.08 | 6.09283E-05 | 0.000682952 |
| 4883 | *NPR3* | ENSG00000113389.16 | 1.57 | 3.56 | 1.99 | 3.98 | 0.000120099 | 0.001116144 |
| 25816 | *TNFAIP8* | ENSG00000145779.8 | 4.15 | 6.13 | 1.98 | 3.95 | 6.01064E-09 | 9.49618E-07 |
| 627 | *BDNF* | ENSG00000176697.20 | 1.25 | 3.24 | 1.98 | 3.95 | 8.63466E-07 | 3.11861E-05 |
| 51209 | *RAB9B* | ENSG00000288597.1 | -1.15 | 0.84 | 1.98 | 3.94 | 0.00104779 | 0.005907696 |
| 330 | *BIRC3* | ENSG00000023445.16 | -1.08 | 0.87 | 1.95 | 3.87 | 2.54111E-05 | 0.000359467 |
| 51330 | *TNFRSF12A* | ENSG00000006327.14 | 2.81 | 4.73 | 1.92 | 3.78 | 1.09625E-07 | 7.76114E-06 |
| 387763 | *C11orf96* | ENSG00000187479.8 | 2.33 | 4.24 | 1.90 | 3.73 | 3.19998E-07 | 1.60095E-05 |
| 222546 | *RFX6* | ENSG00000185002.10 | 1.71 | 3.60 | 1.90 | 3.72 | 1.18836E-06 | 3.91783E-05 |
| 64577 | *ALDH8A1* | ENSG00000118514.14 | 0.01 | 1.91 | 1.89 | 3.72 | 0.000108394 | 0.001042501 |
| 7042 | *TGFB2* | ENSG00000092969.12 | 4.94 | 6.80 | 1.86 | 3.62 | 6.33682E-10 | 2.31974E-07 |
| 148641 | *SLC35F3* | ENSG00000183780.13 | 0.47 | 2.28 | 1.81 | 3.51 | 3.36205E-05 | 0.000439556 |
| 619518 | *SSBP3-AS1* | ENSG00000198711.5 | -1.42 | 0.40 | 1.80 | 3.49 | 0.000747942 | 0.004546714 |
| 81792 | *ADAMTS12* | ENSG00000151388.11 | 1.35 | 3.15 | 1.79 | 3.47 | 2.01955E-06 | 5.65513E-05 |
| 27098 | *CLUL1* | ENSG00000079101.17 | 3.22 | 5.00 | 1.79 | 3.45 | 8.04689E-08 | 6.18901E-06 |
| 688 | *KLF5* | ENSG00000102554.14 | 2.17 | 3.93 | 1.76 | 3.39 | 0.000131875 | 0.001200312 |
| 6324 | *SCN1B* | ENSG00000105711.14 | 0.05 | 1.81 | 1.76 | 3.38 | 0.001674848 | 0.008441166 |
| 554282 | *FAM72C* | ENSG00000263513.6 | 3.85 | 5.60 | 1.75 | 3.36 | 2.37816E-09 | 5.3792E-07 |
| 894 | *CCND2* | ENSG00000118971.9 | 5.73 | 7.48 | 1.75 | 3.36 | 9.56691E-11 | 9.68773E-08 |
| 384 | *ARG2* | ENSG00000081181.8 | 4.95 | 6.69 | 1.74 | 3.33 | 1.53518E-10 | 1.18336E-07 |
| 81786 | *TRIM7* | ENSG00000146054.18 | 0.11 | 1.83 | 1.72 | 3.29 | 0.000547388 | 0.003584535 |

| | | | | | | | | |
|---|---|---|---|---|---|---|---|---|
| 1490 | *CCN2* | ENSG00000118523.6 | -0.07 | 1.65 | 1.71 | 3.26 | 0.001302593 | 0.007009902 |
| 83604 | *TMEM47* | ENSG00000147027.4 | 8.76 | 10.47 | 1.70 | 3.26 | 5.64932E-13 | 2.21965E-09 |
| 23764 | *MAFF* | ENSG00000185022.12 | 1.57 | 3.24 | 1.68 | 3.21 | 8.97495E-06 | 0.000170513 |
| 25780 | *RASGRP3* | ENSG00000152689.18 | 2.13 | 3.81 | 1.68 | 3.21 | 2.66711E-06 | 6.93772E-05 |
| 4783 | *NFIL3* | ENSG00000165030.4 | 4.92 | 6.60 | 1.68 | 3.20 | 1.57686E-10 | 1.18336E-07 |
| 2669 | *GEM* | ENSG00000164949.8 | 2.93 | 4.61 | 1.67 | 3.19 | 3.46409E-07 | 1.6611E-05 |
| 100506827 | *LINC02475* | ENSG00000251350.2 | 3.26 | 4.92 | 1.66 | 3.17 | 8.09501E-07 | 2.96337E-05 |
| 219539 | *YPEL4* | ENSG00000166793.13 | -0.63 | 1.02 | 1.66 | 3.16 | 0.000222505 | 0.001786824 |
| 101928721 | *LOC101928721* | ENSG00000247810.8 | 0.23 | 1.88 | 1.65 | 3.14 | 1.89897E-05 | 0.000289356 |
| 245806 | *VGLL2* | ENSG00000170162.14 | 1.11 | 2.72 | 1.62 | 3.07 | 0.000124801 | 0.00114776 |
| 84570 | *COL25A1* | ENSG00000188517.17 | 6.42 | 8.04 | 1.62 | 3.07 | 3.11064E-10 | 1.45065E-07 |
| 283078 | *MKX* | ENSG00000150051.14 | -0.32 | 1.30 | 1.61 | 3.05 | 0.000103597 | 0.001009015 |
| 57574 | *MARCHF4* | ENSG00000144583.5 | 1.54 | 3.14 | 1.60 | 3.03 | 1.16332E-05 | 0.000205416 |
| 64359 | *NXN* | ENSG00000167693.17 | 4.08 | 5.67 | 1.59 | 3.01 | 4.59915E-05 | 0.000553558 |
| 2119 | *ETV5* | ENSG00000244405.8 | 5.70 | 7.28 | 1.57 | 2.98 | 7.23692E-09 | 1.08619E-06 |
| 6506 | *SLC1A2* | ENSG00000110436.13 | 1.83 | 3.40 | 1.57 | 2.97 | 4.58216E-05 | 0.000552399 |
| 26050 | *SLITRK5* | ENSG00000165300.8 | 0.80 | 2.38 | 1.57 | 2.97 | 4.10434E-05 | 0.000509916 |
| 481 | *ATP1B1* | ENSG00000143153.14 | 7.47 | 9.04 | 1.56 | 2.96 | 2.96623E-11 | 4.49291E-08 |
| 55117 | *SLC6A15* | ENSG00000072041.18 | 1.52 | 3.08 | 1.56 | 2.95 | 0.00053298 | 0.003510091 |
| 27129 | *HSPB7* | ENSG00000173641.18 | 6.91 | 8.47 | 1.56 | 2.94 | 6.50371E-10 | 2.32415E-07 |
| 3397 | *ID1* | ENSG00000125968.9 | 0.47 | 2.02 | 1.55 | 2.94 | 0.000577952 | 0.00374546 |
| 53342 | *IL17D* | ENSG00000172458.5 | 1.65 | 3.20 | 1.55 | 2.93 | 7.77551E-07 | 2.90305E-05 |
| 1604 | *CD55* | ENSG00000196352.17 | 6.25 | 7.80 | 1.55 | 2.93 | 2.54316E-10 | 1.38663E-07 |
| 375387 | *NRROS* | ENSG00000174004.6 | -1.88 | -0.34 | 1.54 | 2.91 | 0.001556578 | 0.008009145 |
| 222545 | *GPRC6A* | ENSG00000173612.10 | 0.23 | 1.76 | 1.53 | 2.90 | 0.001502637 | 0.007795559 |
| 196294 | *IMMP1L* | ENSG00000148950.11 | 1.25 | 2.78 | 1.53 | 2.90 | 0.000169298 | 0.001444569 |
| 55244 | *SLC47A1* | ENSG00000142494.13 | 6.78 | 8.30 | 1.53 | 2.88 | 2.09294E-10 | 1.25652E-07 |
| 58476 | *TP53INP2* | ENSG00000078804.13 | 5.48 | 7.00 | 1.52 | 2.86 | 1.90289E-09 | 4.84076E-07 |
| 2355 | *FOSL2* | ENSG00000075426.12 | 5.47 | 6.97 | 1.51 | 2.84 | 7.39576E-10 | 2.58147E-07 |
| 3815 | *KIT* | ENSG00000157404.17 | 0.83 | 2.33 | 1.50 | 2.84 | 0.000628905 | 0.003987845 |
| 3751 | *KCND2* | ENSG00000184408.10 | 1.80 | 3.30 | 1.50 | 2.84 | 0.000109044 | 0.001046442 |
| 145270 | *PRIMA1* | ENSG00000175785.13 | 2.19 | 3.69 | 1.50 | 2.83 | 1.49186E-06 | 4.51344E-05 |
| 8553 | *BHLHE40* | ENSG00000134107.5 | 3.99 | 5.49 | 1.50 | 2.83 | 3.25738E-07 | 1.61135E-05 |
| 3418 | *IDH2* | ENSG00000182054.10 | 6.53 | 8.03 | 1.50 | 2.82 | 4.50548E-11 | 5.7363E-08 |
| 23554 | *TSPAN12* | ENSG00000106025.9 | 6.80 | 8.29 | 1.50 | 2.82 | 2.44894E-11 | 4.49291E-08 |
| 3953 | *LEPR* | ENSG00000116678.20 | 3.01 | 4.49 | 1.49 | 2.80 | 3.03792E-07 | 1.57074E-05 |
| 6303 | *SAT1* | ENSG00000130066.17 | 4.56 | 6.04 | 1.48 | 2.79 | 2.40127E-09 | 5.3792E-07 |
| 100302736 | *TMED7-TICAM2* | ENSG00000251201.8 | 3.76 | 5.24 | 1.48 | 2.79 | 3.32962E-07 | 1.61986E-05 |
| 3516 | *RBPJ* | ENSG00000168214.22 | 7.71 | 9.19 | 1.48 | 2.78 | 3.22374E-10 | 1.45065E-07 |
| 105375666 | *VPS13B-DT* | ENSG00000253948.3 | -0.81 | 0.66 | 1.47 | 2.78 | 0.001357325 | 0.007215973 |
| 687 | *KLF9* | ENSG00000119138.5 | 1.72 | 3.19 | 1.47 | 2.77 | 1.52189E-05 | 0.000248013 |
| 115827 | *RAB3C* | ENSG00000152932.8 | 6.39 | 7.86 | 1.47 | 2.76 | 5.87436E-07 | 2.44233E-05 |
| 9248 | *GPR50* | ENSG00000102195.10 | 1.42 | 2.89 | 1.47 | 2.76 | 0.00021408 | 0.001731214 |
| 22837 | *COBLL1* | ENSG00000082438.18 | 1.14 | 2.60 | 1.46 | 2.75 | 0.001797348 | 0.008935541 |
| 56521 | *DNAJC12* | ENSG00000108176.15 | 4.47 | 5.93 | 1.46 | 2.75 | 3.89942E-09 | 7.60084E-07 |
| 6533 | *SLC6A6* | ENSG00000131389.18 | 7.36 | 8.82 | 1.46 | 2.74 | 4.58629E-11 | 5.7363E-08 |
| 9497 | *SLC4A7* | ENSG00000033867.17 | 4.33 | 5.78 | 1.45 | 2.74 | 7.53269E-08 | 6.02913E-06 |
| 54805 | *CNNM2* | ENSG00000148842.18 | 6.09 | 7.53 | 1.44 | 2.72 | 3.66631E-09 | 7.33296E-07 |
| 55450 | *CAMK2N1* | ENSG00000162545.6 | 6.75 | 8.19 | 1.44 | 2.71 | 1.69541E-10 | 1.21174E-07 |
| 128209 | *KLF17* | ENSG00000171872.5 | 2.51 | 3.94 | 1.44 | 2.71 | 1.82893E-06 | 5.25212E-05 |
| 6422 | *SFRP1* | ENSG00000104332.12 | 6.30 | 7.73 | 1.44 | 2.70 | 9.68193E-11 | 9.68773E-08 |
| 9021 | *SOCS3* | ENSG00000184557.4 | 2.57 | 4.00 | 1.43 | 2.70 | 5.56824E-08 | 4.8503E-06 |
| 283209 | *PGM2L1* | ENSG00000165434.8 | 6.06 | 7.49 | 1.43 | 2.70 | 1.2961E-10 | 1.13323E-07 |
| 22808 | *MRAS* | ENSG00000158186.13 | 6.89 | 8.32 | 1.43 | 2.70 | 7.81043E-08 | 6.13101E-06 |
| 729533 | *FAM72A* | ENSG00000196550.11 | 4.13 | 5.56 | 1.43 | 2.70 | 4.36983E-07 | 1.98147E-05 |
| 753 | *LDLRAD4* | ENSG00000168675.19 | 2.82 | 4.25 | 1.43 | 2.69 | 5.57817E-08 | 4.8503E-06 |
| 378805 | *LINC-PINT* | ENSG00000231721.7 | 1.88 | 3.29 | 1.41 | 2.66 | 0.000258704 | 0.00200252 |
| 3955 | *LFNG* | ENSG00000106003.14 | 0.81 | 2.22 | 1.41 | 2.66 | 0.000140091 | 0.00125605 |
| 1028 | *CDKN1C* | ENSG00000129757.15 | 0.89 | 2.30 | 1.41 | 2.65 | 0.000462893 | 0.003146542 |
| 54855 | *TENT5C* | ENSG00000183508.5 | 4.49 | 5.90 | 1.41 | 2.65 | 8.78958E-09 | 1.27541E-06 |
| 29970 | *SCHIP1* | ENSG00000283154.3 | 1.62 | 3.03 | 1.41 | 2.65 | 1.48322E-05 | 0.000242766 |
| 6277 | *S100A6* | ENSG00000197956.10 | 6.37 | 7.78 | 1.40 | 2.65 | 5.55182E-08 | 4.8503E-06 |
| 22795 | *NID2* | ENSG00000087303.19 | 3.94 | 5.35 | 1.40 | 2.64 | 2.9266E-06 | 7.39484E-05 |
| 65055 | *REEP1* | ENSG00000068615.20 | 2.46 | 3.86 | 1.40 | 2.64 | 7.03343E-06 | 0.000140006 |
| 57480 | *PLEKHG1* | ENSG00000120278.16 | 2.87 | 4.26 | 1.40 | 2.64 | 0.000161583 | 0.001390984 |
| 1647 | *GADD45A* | ENSG00000116717.13 | 3.23 | 4.62 | 1.40 | 2.64 | 3.44386E-07 | 1.6567E-05 |
| 1263 | *PLK3* | ENSG00000173846.13 | 1.56 | 2.94 | 1.39 | 2.62 | 4.51829E-05 | 0.000546015 |
| 55203 | *LGI2* | ENSG00000153012.12 | 2.37 | 3.76 | 1.39 | 2.61 | 7.01369E-05 | 0.000759523 |
| 56895 | *AGPAT4* | ENSG00000026652.15 | 4.06 | 5.42 | 1.37 | 2.58 | 7.23004E-08 | 5.86571E-06 |
| 353322 | *ANKRD37* | ENSG00000186352.9 | -0.33 | 1.04 | 1.37 | 2.58 | 0.001467642 | 0.007667192 |
| 22801 | *ITGA11* | ENSG00000137809.17 | 1.59 | 2.95 | 1.36 | 2.56 | 0.001162612 | 0.006405892 |
| 56246 | *MRAP* | ENSG00000170262.13 | 2.38 | 3.73 | 1.35 | 2.55 | 1.13206E-05 | 0.000201555 |
| 4430 | *MYO1B* | ENSG00000128641.19 | 3.64 | 4.99 | 1.35 | 2.55 | 1.88045E-07 | 1.12445E-05 |
| 653820 | *FAM72B* | ENSG00000188610.12 | 4.48 | 5.83 | 1.35 | 2.55 | 4.50194E-08 | 4.22311E-06 |
| 1846 | *DUSP4* | ENSG00000120875.9 | 0.89 | 2.24 | 1.35 | 2.55 | 1.76587E-05 | 0.000273386 |
| 220441 | *RNF152* | ENSG00000176641.11 | 3.28 | 4.64 | 1.35 | 2.54 | 1.19388E-06 | 3.921E-05 |
| 91373 | *UAP1L1* | ENSG00000197355.11 | 3.82 | 5.16 | 1.35 | 2.54 | 2.01984E-08 | 2.29665E-06 |
| 23302 | *WSCD1* | ENSG00000179314.16 | 4.13 | 5.48 | 1.34 | 2.54 | 2.22461E-07 | 1.25523E-05 |
| 654 | *BMP6* | ENSG00000153162.9 | 4.37 | 5.71 | 1.34 | 2.54 | 1.93696E-08 | 2.2363E-06 |
| 5033 | *P4HA1* | ENSG00000122884.13 | 5.99 | 7.33 | 1.34 | 2.53 | 1.86656E-09 | 4.8302E-07 |
| 8828 | *NRP2* | ENSG00000118257.17 | 6.81 | 8.14 | 1.33 | 2.52 | 9.11221E-08 | 6.70417E-06 |
| 114134 | *SLC2A13* | ENSG00000151229.13 | 6.03 | 7.36 | 1.33 | 2.51 | 8.51909E-10 | 2.78965E-07 |
| 29785 | *CYP2S1* | ENSG00000167600.14 | 2.20 | 3.52 | 1.33 | 2.51 | 0.000305915 | 0.002291156 |
| 9367 | *RAB9A* | ENSG00000123595.8 | 4.35 | 5.67 | 1.32 | 2.50 | 8.04052E-07 | 2.95476E-05 |
| 253559 | *CADM2* | ENSG00000175161.14 | 4.38 | 5.70 | 1.32 | 2.50 | 3.51025E-09 | 7.11964E-07 |
| 441061 | *MARCHF11* | ENSG00000183654.9 | 0.63 | 1.95 | 1.31 | 2.49 | 5.09814E-05 | 0.0005964 |
| 4241 | *MELTF* | ENSG00000163975.13 | 2.55 | 3.86 | 1.31 | 2.49 | 2.77581E-05 | 0.000382573 |

| | | | | | | | | |
|---|---|---|---|---|---|---|---|---|
| 81029 | *WNT5B* | ENSG00000111186.13 | 0.31 | 1.62 | 1.31 | 2.49 | 0.000794675 | 0.004761391 |
| 3651 | *PDX1* | ENSG00000139515.6 | 0.65 | 1.96 | 1.31 | 2.48 | 1.20996E-05 | 0.000211165 |
| 4811 | *NID1* | ENSG00000116962.15 | 7.48 | 8.79 | 1.31 | 2.48 | 4.11196E-10 | 1.76333E-07 |
| 728833 | *FAM72D* | ENSG00000215784.6 | 4.66 | 5.97 | 1.31 | 2.48 | 5.65529E-08 | 4.8503E-06 |
| 970 | *CD70* | ENSG00000125726.11 | 1.07 | 2.38 | 1.31 | 2.47 | 0.000481481 | 0.003244969 |
| 83875 | *BCO2* | ENSG00000197580.13 | 2.39 | 3.69 | 1.30 | 2.47 | 0.000576665 | 0.003738733 |
| 127544 | *RNF19B* | ENSG00000116514.17 | 5.22 | 6.52 | 1.30 | 2.47 | 5.28689E-09 | 8.91584E-07 |
| 345275 | *HSD17B13* | ENSG00000170509.12 | -0.63 | 0.66 | 1.29 | 2.44 | 0.001360513 | 0.007223184 |
| 57520 | *HECW2* | ENSG00000138411.13 | 1.75 | 3.03 | 1.28 | 2.43 | 0.000677351 | 0.004195775 |
| 84302 | *PGAP4* | ENSG00000165152.9 | 7.31 | 8.59 | 1.28 | 2.43 | 1.35906E-10 | 1.13323E-07 |
| 84418 | *CYSTM1* | ENSG00000120306.11 | 5.13 | 6.41 | 1.28 | 2.42 | 1.14512E-09 | 3.58066E-07 |
| 57526 | *PCDH19* | ENSG00000165194.15 | 1.58 | 2.85 | 1.28 | 2.42 | 0.000565915 | 0.003684954 |
| 8870 | *IER3* | ENSG00000137331.12 | -0.30 | 0.97 | 1.27 | 2.42 | 0.000522978 | 0.003454834 |
| 9510 | *ADAMTS1* | ENSG00000154734.16 | 1.69 | 2.96 | 1.26 | 2.40 | 0.000778841 | 0.004694629 |
| 57103 | *TIGAR* | ENSG00000078237.7 | 3.67 | 4.93 | 1.26 | 2.40 | 0.000133427 | 0.001210765 |
| 150094 | *SIK1* | ENSG00000142178.9 | 3.95 | 5.20 | 1.26 | 2.40 | 0.000316727 | 0.002349854 |
| 4121 | *MAN1A1* | ENSG00000111885.7 | 4.22 | 5.48 | 1.26 | 2.39 | 7.33226E-08 | 5.91666E-06 |
| 114990 | *VASN* | ENSG00000168140.5 | 1.81 | 3.07 | 1.26 | 2.39 | 4.97907E-06 | 0.000109096 |
| 10560 | *SLC19A2* | ENSG00000117479.15 | 5.68 | 6.93 | 1.26 | 2.39 | 5.15444E-09 | 8.79125E-07 |
| 2571 | *GAD1* | ENSG00000128683.14 | 2.02 | 3.27 | 1.25 | 2.38 | 0.000571762 | 0.003714968 |
| 26278 | *SACS* | ENSG00000151835.17 | 7.38 | 8.63 | 1.25 | 2.37 | 3.10688E-07 | 1.57538E-05 |
| 643201 | *LOC643201* | ENSG00000248596.8 | 2.93 | 4.17 | 1.24 | 2.36 | 1.38144E-05 | 0.000233491 |
| 25861 | *WHRN* | ENSG00000095397.16 | 7.62 | 8.85 | 1.24 | 2.36 | 5.61049E-09 | 9.25361E-07 |
| 25984 | *KRT23* | ENSG00000108244.17 | 2.86 | 4.09 | 1.23 | 2.35 | 6.35482E-06 | 0.000130699 |
| 3164 | *NR4A1* | ENSG00000123358.20 | 6.61 | 7.84 | 1.23 | 2.34 | 4.99993E-10 | 2.02821E-07 |
| 11015 | *KDELR3* | ENSG00000100196.11 | 3.10 | 4.33 | 1.23 | 2.34 | 1.80147E-06 | 5.20968E-05 |
| 5592 | *PRKG1* | ENSG00000185532.20 | 2.32 | 3.54 | 1.23 | 2.34 | 0.001568296 | 0.008048318 |
| 8447 | *DOC2B* | ENSG00000272636.4 | 5.28 | 6.50 | 1.22 | 2.33 | 6.13936E-08 | 5.1478E-06 |
| 5764 | *PTN* | ENSG00000105894.12 | 6.61 | 7.83 | 1.22 | 2.33 | 2.71661E-10 | 1.38663E-07 |
| 4646 | *MYO6* | ENSG00000196586.17 | 6.30 | 7.52 | 1.22 | 2.32 | 6.10823E-09 | 9.54983E-07 |
| 27071 | *DAPP1* | ENSG00000070190.13 | 1.09 | 2.29 | 1.21 | 2.31 | 0.00037525 | 0.002677517 |
| 54583 | *EGLN1* | ENSG00000135766.9 | 6.78 | 7.99 | 1.21 | 2.31 | 1.06265E-08 | 1.44994E-06 |
| 5169 | *ENPP3* | ENSG00000154269.15 | 1.92 | 3.12 | 1.20 | 2.30 | 1.21453E-05 | 0.000211472 |
| 160851 | *DGKH* | ENSG00000102780.17 | 4.06 | 5.26 | 1.20 | 2.30 | 6.28999E-06 | 0.000129858 |
| 27309 | *ZNF330* | ENSG00000109445.11 | 6.22 | 7.41 | 1.19 | 2.29 | 2.30335E-09 | 5.31861E-07 |
| 390 | *RND3* | ENSG00000115963.14 | 3.19 | 4.38 | 1.19 | 2.28 | 1.21409E-05 | 0.000211472 |
| 340485 | *ACER2* | ENSG00000177076.6 | 1.65 | 2.84 | 1.18 | 2.27 | 1.48963E-05 | 0.000243549 |
| 7421 | *VDR* | ENSG00000111424.12 | 6.53 | 7.71 | 1.18 | 2.27 | 2.00506E-09 | 4.87725E-07 |
| 93010 | *B3GNT7* | ENSG00000156966.7 | 2.95 | 4.13 | 1.18 | 2.26 | 5.14424E-06 | 0.000111898 |
| 114789 | *SLC25A25* | ENSG00000148339.13 | 5.32 | 6.49 | 1.18 | 2.26 | 8.21269E-08 | 6.22436E-06 |
| 7078 | *TIMP3* | ENSG00000100234.12 | 9.99 | 11.16 | 1.17 | 2.26 | 5.69525E-11 | 6.57538E-08 |
| 26503 | *SLC17A5* | ENSG00000119899.13 | 5.07 | 6.24 | 1.17 | 2.26 | 2.82326E-07 | 1.50264E-05 |
| 23493 | *HEY2* | ENSG00000135547.9 | 2.76 | 3.93 | 1.17 | 2.25 | 0.000413177 | 0.002889738 |
| 6482 | *ST3GAL1* | ENSG00000008513.16 | 5.68 | 6.84 | 1.16 | 2.24 | 5.64553E-08 | 4.8503E-06 |
| 84624 | *FNDC1* | ENSG00000164694.17 | 1.91 | 3.07 | 1.16 | 2.24 | 0.001779439 | 0.008860623 |
| 195827 | *PRXL2C* | ENSG00000158122.12 | 3.60 | 4.77 | 1.16 | 2.24 | 9.21223E-07 | 3.261E-05 |
| 728215 | *NALF1* | ENSG00000204442.4 | 5.57 | 6.73 | 1.16 | 2.24 | 5.13986E-09 | 8.79125E-07 |
| 9242 | *MSC* | ENSG00000178860.9 | 0.05 | 1.21 | 1.16 | 2.23 | 0.001463697 | 0.007649242 |
| 415116 | *PIM3* | ENSG00000198355.5 | 4.03 | 5.19 | 1.16 | 2.23 | 1.3054E-07 | 8.62284E-06 |
| 51030 | *TVP23B* | ENSG00000171928.14 | 5.66 | 6.82 | 1.16 | 2.23 | 4.39595E-09 | 8.24735E-07 |
| 9200 | *HACD1* | ENSG00000165996.14 | 1.93 | 3.09 | 1.15 | 2.23 | 0.000470714 | 0.003182406 |
| 8518 | *ELP1* | ENSG00000070061.16 | 9.08 | 10.24 | 1.15 | 2.23 | 1.70658E-09 | 4.8302E-07 |
| 23645 | *PPP1R15A* | ENSG00000087074.8 | 4.78 | 5.93 | 1.15 | 2.22 | 1.40692E-08 | 1.7597E-06 |
| 7803 | *PTP4A1* | ENSG00000112245.13 | 7.36 | 8.52 | 1.15 | 2.22 | 4.72612E-08 | 4.37867E-06 |
| 140469 | *MYO3B* | ENSG00000071909.19 | 0.44 | 1.59 | 1.15 | 2.22 | 0.000174764 | 0.001483612 |
| 1843 | *DUSP1* | ENSG00000120129.6 | 3.79 | 4.94 | 1.15 | 2.21 | 3.8138E-08 | 3.74126E-06 |
| 4747 | *NEFL* | ENSG00000277586.4 | 7.00 | 8.15 | 1.15 | 2.21 | 1.85336E-10 | 1.25652E-07 |
| 302 | *ANXA2* | ENSG00000182718.18 | 5.87 | 7.02 | 1.14 | 2.21 | 5.11522E-08 | 4.59726E-06 |
| 51026 | *GOLT1B* | ENSG00000111711.10 | 7.27 | 8.41 | 1.14 | 2.20 | 1.85865E-09 | 4.8302E-07 |
| 6696 | *SPP1* | ENSG00000118785.15 | 6.26 | 7.40 | 1.14 | 2.20 | 1.04048E-08 | 1.43271E-06 |
| 100506710 | *EBLN3P* | ENSG00000281649.3 | 7.57 | 8.70 | 1.14 | 2.20 | 4.32605E-09 | 8.21894E-07 |
| 8611 | *PLPP1* | ENSG00000067113.17 | 5.61 | 6.75 | 1.13 | 2.20 | 1.55916E-09 | 4.58852E-07 |
| 23086 | *EXPH5* | ENSG00000110723.12 | 4.75 | 5.88 | 1.13 | 2.19 | 1.52358E-07 | 9.64867E-06 |
| 1525 | *CXADR* | ENSG00000154639.19 | 5.59 | 6.72 | 1.13 | 2.19 | 4.63767E-08 | 4.3234E-06 |
| 883 | *KYAT1* | ENSG00000171097.14 | 2.38 | 3.51 | 1.13 | 2.18 | 8.45465E-05 | 0.000868835 |
| 9709 | *HERPUD1* | ENSG00000051108.15 | 7.63 | 8.76 | 1.13 | 2.18 | 8.44725E-10 | 2.78965E-07 |
| 10253 | *SPRY2* | ENSG00000136158.12 | 2.14 | 3.27 | 1.13 | 2.18 | 2.27604E-05 | 0.000331018 |
| 167127 | *UGT3A2* | ENSG00000168671.10 | 1.26 | 2.39 | 1.13 | 2.18 | 0.000153082 | 0.001338934 |
| 26524 | *LATS2* | ENSG00000150457.9 | 4.71 | 5.84 | 1.13 | 2.18 | 1.08647E-08 | 1.46908E-06 |
| 6275 | *S100A4* | ENSG00000196154.12 | 2.81 | 3.94 | 1.12 | 2.18 | 2.08015E-05 | 0.0003108 |
| 51765 | *STK26* | ENSG00000134602.16 | 5.34 | 6.46 | 1.12 | 2.17 | 2.0552E-09 | 4.89628E-07 |
| 2274 | *FHL2* | ENSG00000115641.19 | 5.93 | 7.05 | 1.12 | 2.17 | 4.77083E-09 | 8.50382E-07 |
| 9762 | *LZTS3* | ENSG00000088899.16 | 3.24 | 4.35 | 1.11 | 2.16 | 4.13458E-06 | 9.54707E-05 |
| 10150 | *MBNL2* | ENSG00000139793.20 | 7.42 | 8.53 | 1.11 | 2.16 | 1.75746E-09 | 4.8302E-07 |
| 3632 | *INPP5A* | ENSG00000068383.19 | 6.19 | 7.30 | 1.11 | 2.16 | 6.76319E-08 | 5.5774E-06 |
| 644815 | *FAM83G* | ENSG00000188522.15 | 2.76 | 3.86 | 1.11 | 2.15 | 0.000223381 | 0.001790989 |
| 105378853 | *LINC02609* | ENSG00000233593.11 | 1.79 | 2.89 | 1.11 | 2.15 | 0.000176527 | 0.001490993 |
| 23252 | *OTUD3* | ENSG00000169914.6 | 4.02 | 5.13 | 1.11 | 2.15 | 0.001912106 | 0.009369505 |
| 5165 | *PDK3* | ENSG00000067992.17 | 6.33 | 7.44 | 1.10 | 2.15 | 2.9657E-07 | 1.55095E-05 |
| 6386 | *SDCBP* | ENSG00000137575.12 | 7.88 | 8.98 | 1.10 | 2.14 | 5.32982E-10 | 2.05116E-07 |
| 5187 | *PER1* | ENSG00000179094.16 | 6.37 | 7.47 | 1.10 | 2.14 | 1.16674E-05 | 0.000205777 |
| 56254 | *RNF20* | ENSG00000155827.12 | 7.78 | 8.88 | 1.10 | 2.14 | 2.72718E-07 | 1.4671E-05 |
| 9314 | *KLF4* | ENSG00000136826.15 | 2.27 | 3.36 | 1.10 | 2.14 | 1.05822E-05 | 0.000192519 |
| 253512 | *SLC25A30* | ENSG00000174032.17 | 4.57 | 5.67 | 1.10 | 2.14 | 1.62255E-05 | 0.000259902 |
| 101410533 | *YTHDF3-DT* | ENSG00000270673.1 | -0.57 | 0.53 | 1.10 | 2.14 | 0.001932615 | 0.009442259 |
| 23710 | *GABARAPL1* | ENSG00000139112.11 | 7.29 | 8.39 | 1.10 | 2.14 | 9.14533E-10 | 2.92047E-07 |
| 9499 | *MYOT* | ENSG00000120729.10 | 9.03 | 10.13 | 1.09 | 2.14 | 2.08246E-10 | 1.25652E-07 |

| | | | | | | | | |
|---|---|---|---|---|---|---|---|---|
| 1305 | *COL13A1* | ENSG00000197467.17 | 2.34 | 3.43 | 1.09 | 2.13 | 6.02089E-05 | 0.000677046 |
| 87 | *ACTN1* | ENSG00000072110.16 | 5.64 | 6.73 | 1.09 | 2.13 | 2.50218E-08 | 2.71859E-06 |
| 79850 | *TLCD3A* | ENSG00000167695.15 | 4.46 | 5.55 | 1.09 | 2.13 | 7.88965E-08 | 6.13553E-06 |
| 91608 | *RASL10B* | ENSG00000270885.2 | 5.66 | 6.74 | 1.09 | 2.12 | 4.51557E-09 | 8.36719E-07 |
| 27022 | *FOXD3* | ENSG00000187140.6 | 1.25 | 2.33 | 1.09 | 2.12 | 0.000149441 | 0.001317834 |
| 58515 | *SELENOK* | ENSG00000113811.12 | 5.62 | 6.71 | 1.09 | 2.12 | 6.59702E-09 | 1.00015E-06 |
| 55816 | *DOK5* | ENSG00000101134.12 | 2.92 | 4.00 | 1.08 | 2.12 | 0.00018723 | 0.00155686 |
| 3486 | *IGFBP3* | ENSG00000146674.16 | 3.71 | 4.79 | 1.08 | 2.12 | 6.03186E-07 | 2.47926E-05 |
| 91768 | *CABLES1* | ENSG00000134508.13 | 3.59 | 4.67 | 1.08 | 2.11 | 1.70282E-05 | 0.00026762 |
| 491 | *ATP2B2* | ENSG00000157087.20 | 4.34 | 5.41 | 1.08 | 2.11 | 8.09448E-06 | 0.000156358 |
| 84957 | *RELT* | ENSG00000054967.13 | 2.89 | 3.97 | 1.08 | 2.11 | 9.02712E-06 | 0.000170796 |
| 2027 | *ENO3* | ENSG00000108515.18 | 4.71 | 5.79 | 1.07 | 2.10 | 4.89359E-07 | 2.14705E-05 |
| 6517 | *SLC2A4* | ENSG00000181856.15 | 4.41 | 5.48 | 1.07 | 2.10 | 7.22287E-08 | 5.86571E-06 |
| 727910 | *TLCD2* | ENSG00000185561.10 | 2.59 | 3.66 | 1.07 | 2.10 | 0.000197935 | 0.001629625 |
| 1039 | *CDR2* | ENSG00000140743.8 | 5.49 | 6.56 | 1.07 | 2.09 | 8.21462E-08 | 6.22436E-06 |
| 9871 | *SEC24D* | ENSG00000150961.15 | 5.60 | 6.67 | 1.06 | 2.09 | 5.44644E-08 | 4.8503E-06 |
| 132864 | *CPEB2* | ENSG00000137449.17 | 3.21 | 4.28 | 1.06 | 2.09 | 0.000182131 | 0.00151951 |
| 5292 | *PIM1* | ENSG00000137193.14 | 5.21 | 6.27 | 1.06 | 2.09 | 8.51925E-07 | 3.08854E-05 |
| 55062 | *WIPI1* | ENSG00000070540.13 | 4.65 | 5.71 | 1.06 | 2.08 | 6.67902E-07 | 2.62046E-05 |
| 538 | *ATP7A* | ENSG00000165240.22 | 6.32 | 7.37 | 1.06 | 2.08 | 2.23453E-08 | 2.4843E-06 |
| 1903 | *S1PR3* | ENSG00000213694.6 | 8.45 | 9.50 | 1.06 | 2.08 | 3.28617E-10 | 1.45065E-07 |
| 9745 | *ZNF536* | ENSG00000198597.9 | 4.20 | 5.25 | 1.05 | 2.08 | 6.72215E-07 | 2.62742E-05 |
| 5596 | *MAPK4* | ENSG00000141639.12 | 5.54 | 6.59 | 1.05 | 2.07 | 5.97421E-09 | 9.49618E-07 |
| 8970 | *H2BC11* | ENSG00000124635.9 | 1.57 | 2.63 | 1.05 | 2.07 | 0.000207198 | 0.001684633 |
| 55214 | *P3H2* | ENSG00000090530.10 | 7.24 | 8.29 | 1.05 | 2.07 | 1.45297E-06 | 4.45054E-05 |
| 144501 | *KRT80* | ENSG00000167767.14 | 3.04 | 4.09 | 1.05 | 2.06 | 1.5422E-07 | 9.72558E-06 |
| 100505812 | *CARD8-AS1* | ENSG00000268001.1 | 1.59 | 2.64 | 1.04 | 2.06 | 0.000579347 | 0.00375288 |
| 8577 | *TMEFF1* | ENSG00000241697.5 | 1.58 | 2.62 | 1.04 | 2.06 | 0.000375779 | 0.002678135 |
| 8651 | *SOCS1* | ENSG00000185338.7 | 2.95 | 3.99 | 1.04 | 2.05 | 2.33123E-05 | 0.000336761 |
| 8912 | *CACNA1H* | ENSG00000196557.13 | 5.42 | 6.45 | 1.03 | 2.05 | 5.35082E-07 | 2.28155E-05 |
| 65009 | *NDRG4* | ENSG00000103034.15 | 4.59 | 5.62 | 1.03 | 2.05 | 5.61393E-06 | 0.000119348 |
| 667 | *DST* | ENSG00000151914.22 | 7.42 | 8.45 | 1.03 | 2.04 | 2.25182E-06 | 6.12274E-05 |
| 1803 | *DPP4* | ENSG00000197635.11 | 4.80 | 5.83 | 1.03 | 2.04 | 2.10783E-06 | 5.82622E-05 |
| 84628 | *NTNG2* | ENSG00000196358.12 | 3.51 | 4.54 | 1.03 | 2.04 | 0.001822631 | 0.009028342 |
| 57205 | *ATP10D* | ENSG00000145246.14 | 6.39 | 7.41 | 1.03 | 2.04 | 2.6213E-07 | 1.43966E-05 |
| 10950 | *BTG3* | ENSG00000154640.15 | 5.58 | 6.61 | 1.02 | 2.03 | 1.30363E-08 | 1.65815E-06 |
| 5621 | *PRNP* | ENSG00000171867.18 | 5.67 | 6.69 | 1.02 | 2.03 | 1.18991E-07 | 8.04473E-06 |
| 8987 | *STBD1* | ENSG00000118804.9 | 5.50 | 6.52 | 1.02 | 2.03 | 1.56156E-08 | 1.89012E-06 |
| 5901 | *RAN* | ENSG00000132341.12 | 8.86 | 9.88 | 1.02 | 2.03 | 3.46788E-09 | 7.11964E-07 |
| 7280 | *TUBB2A* | ENSG00000137267.7 | 4.75 | 5.77 | 1.02 | 2.02 | 2.52132E-08 | 2.71859E-06 |
| 9223 | *MAGI1* | ENSG00000151276.24 | 5.93 | 6.95 | 1.01 | 2.02 | 2.43013E-06 | 6.51235E-05 |
| 101928464 | *ZNF433-AS1* | ENSG00000219665.8 | 1.41 | 2.42 | 1.01 | 2.02 | 0.001147782 | 0.006354507 |
| 57795 | *BRINP2* | ENSG00000198797.7 | 8.13 | 9.15 | 1.01 | 2.02 | 4.78896E-10 | 1.9966E-07 |
| 54842 | *MFSD6* | ENSG00000151690.15 | 5.38 | 6.39 | 1.01 | 2.01 | 1.59554E-05 | 0.000256122 |
| 3612 | *IMPA1* | ENSG00000133731.10 | 4.17 | 5.17 | 1.00 | 2.01 | 0.000114249 | 0.001080507 |
| 7301 | *TYRO3* | ENSG00000092445.12 | 6.17 | 7.17 | 1.00 | 2.01 | 3.54303E-08 | 3.54516E-06 |
| 101927809 | *COSMOC* | ENSG00000260552.2 | 0.89 | 1.89 | 1.00 | 2.00 | 0.000553697 | 0.003616376 |
| 6907 | *TBL1X* | ENSG00000101849.18 | 5.40 | 6.40 | 1.00 | 2.00 | 6.13424E-07 | 2.48164E-05 |
| 775 | *CACNA1C* | ENSG00000151067.23 | 5.88 | 6.88 | 1.00 | 2.00 | 5.31673E-05 | 0.000616173 |
| 65991 | *FUNDC2* | ENSG00000165775.18 | 6.57 | 7.57 | 1.00 | 2.00 | 6.04146E-08 | 5.09417E-06 |
| 170850 | *KCNG3* | ENSG00000171126.8 | 1.02 | 2.02 | 1.00 | 2.00 | 0.00036544 | 0.002615591 |
| 157378 | *TMEM65* | ENSG00000164983.8 | 4.22 | 5.21 | 0.99 | 1.99 | 8.73362E-08 | 6.45728E-06 |
| 10855 | *HPSE* | ENSG00000173083.16 | 2.45 | 3.44 | 0.99 | 1.99 | 8.11164E-06 | 0.000156488 |
| 30844 | *EHD4* | ENSG00000103966.11 | 4.84 | 5.83 | 0.99 | 1.99 | 4.61858E-07 | 2.06926E-05 |
| 283624 | *LINC00641* | ENSG00000258441.1 | 2.44 | 3.43 | 0.99 | 1.98 | 0.00066134 | 0.004123828 |
| 57559 | *STAMBPL1* | ENSG00000138134.12 | 2.66 | 3.64 | 0.98 | 1.98 | 1.58375E-05 | 0.000255507 |
| 84740 | *AFAP1-AS1* | ENSG00000272620.2 | 1.75 | 2.74 | 0.98 | 1.98 | 0.00020099 | 0.00164755 |
| 9854 | *C2CD2L* | ENSG00000172375.14 | 3.95 | 4.92 | 0.98 | 1.97 | 0.001762422 | 0.008793946 |
| 25907 | *TMEM158* | ENSG00000249992.2 | 3.91 | 4.89 | 0.98 | 1.97 | 4.39121E-05 | 0.000534098 |
| 8974 | *P4HA2* | ENSG00000072682.20 | 2.83 | 3.81 | 0.98 | 1.97 | 0.000210376 | 0.001705854 |
| 5238 | *PGM3* | ENSG00000013375.16 | 5.84 | 6.81 | 0.97 | 1.97 | 4.33896E-07 | 1.97344E-05 |
| 3628 | *INPP1* | ENSG00000151689.13 | 3.20 | 4.16 | 0.97 | 1.96 | 0.000329526 | 0.002412421 |
| 84803 | *GPAT3* | ENSG00000138678.11 | 2.62 | 3.59 | 0.97 | 1.96 | 1.44001E-05 | 0.000239613 |
| 678 | *ZFP36L2* | ENSG00000152518.8 | 7.83 | 8.80 | 0.97 | 1.96 | 1.28598E-09 | 3.86025E-07 |
| 80031 | *SEMA6D* | ENSG00000137872.17 | 4.52 | 5.49 | 0.97 | 1.96 | 0.000815048 | 0.004835973 |
| 2977 | *GUCY1A2* | ENSG00000152402.11 | 3.64 | 4.61 | 0.97 | 1.96 | 0.00010467 | 0.00101551 |
| 8013 | *NR4A3* | ENSG00000119508.18 | 6.44 | 7.41 | 0.97 | 1.95 | 1.94465E-09 | 4.86454E-07 |
| 388677 | *NOTCH2NLA* | ENSG00000264343.7 | 4.91 | 5.88 | 0.97 | 1.95 | 1.86628E-07 | 1.12315E-05 |
| 57620 | *STIM2* | ENSG00000109689.19 | 5.44 | 6.40 | 0.96 | 1.95 | 1.25349E-05 | 0.000217019 |
| 10493 | *VAT1* | ENSG00000108828.16 | 8.32 | 9.28 | 0.96 | 1.95 | 4.06309E-09 | 7.81832E-07 |
| 57030 | *SLC17A7* | ENSG00000104888.10 | 1.59 | 2.55 | 0.96 | 1.94 | 0.001076809 | 0.006031344 |
| 3488 | *IGFBP5* | ENSG00000115461.5 | 2.53 | 3.48 | 0.96 | 1.94 | 0.000305563 | 0.002290805 |
| 127733 | *UBXN10* | ENSG00000162543.6 | 3.26 | 4.21 | 0.95 | 1.94 | 1.0515E-05 | 0.000191529 |
| 109729137 | *MEF2C-AS2* | ENSG00000245864.4 | 3.05 | 4.00 | 0.95 | 1.94 | 0.000298062 | 0.002246919 |
| 10627 | *MYL12A* | ENSG00000101608.13 | 7.22 | 8.18 | 0.95 | 1.93 | 4.78086E-09 | 8.50382E-07 |
| 1164 | *CKS2* | ENSG00000123975.5 | 7.51 | 8.46 | 0.95 | 1.93 | 2.3892E-08 | 2.63673E-06 |
| 79645 | *EFCAB1* | ENSG00000034239.11 | 3.44 | 4.39 | 0.95 | 1.93 | 0.000410768 | 0.002876909 |
| 6367 | *CCL22* | ENSG00000102962.5 | 4.49 | 5.44 | 0.95 | 1.93 | 1.41398E-06 | 4.38481E-05 |
| 51673 | *TPPP3* | ENSG00000159713.11 | 2.93 | 3.88 | 0.95 | 1.93 | 3.04591E-05 | 0.00040818 |
| 5166 | *PDK4* | ENSG00000004799.8 | 10.32 | 11.27 | 0.95 | 1.93 | 2.43606E-10 | 1.38663E-07 |
| 135154 | *SDHAF4* | ENSG00000154079.6 | 2.36 | 3.30 | 0.94 | 1.92 | 0.00013507 | 0.001218311 |
| 91404 | *SESTD1* | ENSG00000187231.14 | 6.69 | 7.63 | 0.94 | 1.92 | 5.18816E-07 | 2.24268E-05 |
| 1839 | *HBEGF* | ENSG00000113070.8 | 1.11 | 2.05 | 0.94 | 1.92 | 0.000283609 | 0.00215747 |
| 57484 | *RNF150* | ENSG00000170153.11 | 4.95 | 5.89 | 0.94 | 1.92 | 6.05294E-07 | 2.47926E-05 |
| 440145 | *MZT1* | ENSG00000204899.6 | 4.96 | 5.89 | 0.94 | 1.91 | 8.82898E-07 | 3.16263E-05 |
| 6322 | *SCML1* | ENSG00000047634.15 | 5.11 | 6.05 | 0.94 | 1.91 | 4.80626E-08 | 4.39861E-06 |
| 7277 | *TUBA4A* | ENSG00000127824.15 | 4.01 | 4.95 | 0.93 | 1.91 | 2.29574E-05 | 0.000332915 |

| | | | | | | | | |
|---|---|---|---|---|---|---|---|---|
| 51661 | *FKBP7* | ENSG00000079150.19 | 4.56 | 5.50 | 0.93 | 1.91 | 1.45326E-05 | 0.000240484 |
| 6781 | *STC1* | ENSG00000159167.12 | 3.44 | 4.38 | 0.93 | 1.90 | 0.000355617 | 0.002571032 |
| 8218 | *CLTCL1* | ENSG00000070371.16 | 4.56 | 5.49 | 0.92 | 1.90 | 1.4818E-06 | 4.49301E-05 |
| 1847 | *DUSP5* | ENSG00000138166.6 | 4.11 | 5.04 | 0.92 | 1.90 | 2.16827E-06 | 5.94946E-05 |
| 100506311 | *HOTAIRM1* | ENSG00000233429.9 | 2.21 | 3.13 | 0.92 | 1.89 | 0.000496734 | 0.003322409 |
| 11120 | *BTN2A1* | ENSG00000112763.17 | 3.91 | 4.83 | 0.92 | 1.89 | 0.000181055 | 0.001513061 |
| 57605 | *PITPNM2* | ENSG00000090975.14 | 4.95 | 5.86 | 0.92 | 1.89 | 0.000397173 | 0.002802619 |
| 10529 | *NEBL* | ENSG00000078114.19 | 3.76 | 4.68 | 0.92 | 1.89 | 1.26016E-05 | 0.000217649 |
| 58527 | *ABRACL* | ENSG00000146386.8 | 5.04 | 5.96 | 0.92 | 1.89 | 4.11723E-07 | 1.89557E-05 |
| 1306 | *COL15A1* | ENSG00000204291.11 | 8.67 | 9.59 | 0.92 | 1.89 | 5.20888E-10 | 2.05116E-07 |
| 9456 | *HOMER1* | ENSG00000152413.15 | 6.41 | 7.33 | 0.91 | 1.89 | 0.000244109 | 0.00191323 |
| 8556 | *CDC14A* | ENSG00000079335.20 | 1.95 | 2.87 | 0.91 | 1.88 | 0.000129515 | 0.001183851 |
| 81539 | *SLC38A1* | ENSG00000111371.16 | 8.34 | 9.25 | 0.91 | 1.88 | 2.01472E-09 | 4.87725E-07 |
| 81575 | *APOLD1* | ENSG00000178878.13 | 4.91 | 5.82 | 0.91 | 1.88 | 7.55278E-07 | 2.8451E-05 |
| 4881 | *NPR1* | ENSG00000169418.10 | 2.17 | 3.08 | 0.91 | 1.88 | 0.000719927 | 0.004404839 |
| 114793 | *FMNL2* | ENSG00000157827.20 | 5.12 | 6.03 | 0.91 | 1.88 | 1.14987E-07 | 7.9414E-06 |
| 7494 | *XBP1* | ENSG00000100219.17 | 6.54 | 7.45 | 0.91 | 1.88 | 1.7375E-08 | 2.06241E-06 |
| 400073 | *C12orf76* | ENSG00000174456.16 | 3.74 | 4.65 | 0.91 | 1.88 | 7.31117E-05 | 0.000782559 |
| 23389 | *MED13L* | ENSG00000123066.9 | 6.67 | 7.57 | 0.91 | 1.88 | 6.22667E-07 | 2.50553E-05 |
| 160760 | *PPTC7* | ENSG00000196850.6 | 6.68 | 7.58 | 0.90 | 1.87 | 8.37987E-09 | 1.24225E-06 |
| 5831 | *PYCR1* | ENSG00000183010.17 | 4.86 | 5.76 | 0.90 | 1.87 | 4.00518E-06 | 9.31997E-05 |
| 3033 | *HADH* | ENSG00000138796.18 | 6.37 | 7.27 | 0.90 | 1.87 | 5.61007E-09 | 9.25361E-07 |
| 3202 | *HOXA5* | ENSG00000106004.5 | 4.80 | 5.71 | 0.90 | 1.86 | 2.2273E-06 | 6.06707E-05 |
| 7769 | *ZNF226* | ENSG00000167380.17 | 3.94 | 4.84 | 0.90 | 1.86 | 9.74467E-07 | 3.41048E-05 |
| 101929704 | *TTC23L-AS1* | ENSG00000272323.1 | 1.94 | 2.84 | 0.90 | 1.86 | 0.000182329 | 0.001520322 |
| 488 | *ATP2A2* | ENSG00000174437.18 | 8.56 | 9.46 | 0.90 | 1.86 | 5.94089E-09 | 9.49618E-07 |
| 4824 | *NKX3-1* | ENSG00000167034.10 | 2.69 | 3.59 | 0.90 | 1.86 | 0.001154916 | 0.006375186 |
| 143686 | *SESN3* | ENSG00000149212.12 | 5.59 | 6.49 | 0.89 | 1.86 | 2.76373E-08 | 2.92119E-06 |
| 64651 | *CSRNP1* | ENSG00000144655.15 | 3.91 | 4.80 | 0.89 | 1.86 | 1.87502E-06 | 5.32996E-05 |
| 64762 | *GAREM1* | ENSG00000141441.16 | 4.66 | 5.56 | 0.89 | 1.86 | 0.000466325 | 0.003161282 |
| 22987 | *SV2C* | ENSG00000122012.14 | 7.55 | 8.44 | 0.89 | 1.86 | 1.5601E-08 | 1.89012E-06 |
| 90874 | *ZNF697* | ENSG00000143067.5 | 3.63 | 4.52 | 0.89 | 1.85 | 2.11026E-05 | 0.000313705 |
| 55304 | *SPTLC3* | ENSG00000172296.13 | 2.97 | 3.86 | 0.89 | 1.85 | 0.000806627 | 0.004808046 |
| 55254 | *TMEM39A* | ENSG00000176142.13 | 5.90 | 6.79 | 0.89 | 1.85 | 1.41159E-06 | 4.38481E-05 |
| 1750 | *DLX6* | ENSG00000006377.11 | 1.95 | 2.84 | 0.89 | 1.85 | 0.000114606 | 0.001082516 |
| 341640 | *FREM2* | ENSG00000150893.11 | 3.69 | 4.58 | 0.89 | 1.85 | 0.000455359 | 0.003112244 |
| 79026 | *AHNAK* | ENSG00000124942.14 | 4.58 | 5.47 | 0.89 | 1.85 | 8.75817E-05 | 0.000889989 |
| 492311 | *IGIP* | ENSG00000182700.5 | 3.21 | 4.10 | 0.89 | 1.85 | 0.000381683 | 0.002711159 |
| 26277 | *TINF2* | ENSG00000092330.18 | 5.63 | 6.52 | 0.89 | 1.85 | 4.87783E-07 | 2.14696E-05 |
| 5029 | *P2RY2* | ENSG00000175591.12 | 4.03 | 4.91 | 0.88 | 1.84 | 1.72898E-05 | 0.000269754 |
| 8744 | *TNFSF9* | ENSG00000125657.5 | 1.25 | 2.13 | 0.88 | 1.84 | 0.001852207 | 0.009145533 |
| 54602 | *NDFIP2* | ENSG00000102471.15 | 5.72 | 6.60 | 0.88 | 1.84 | 2.04668E-07 | 1.18148E-05 |
| 55353 | *LAPTM4B* | ENSG00000104341.17 | 7.34 | 8.21 | 0.88 | 1.84 | 5.58281E-07 | 2.34218E-05 |
| 81550 | *TDRD3* | ENSG00000083544.16 | 4.25 | 5.13 | 0.88 | 1.84 | 5.2695E-05 | 0.000613101 |
| 3512 | *JCHAIN* | ENSG00000132465.12 | 2.25 | 3.12 | 0.88 | 1.84 | 0.001471886 | 0.00767867 |
| 81615 | *TMEM163* | ENSG00000152128.13 | 2.59 | 3.46 | 0.87 | 1.83 | 2.86658E-05 | 0.000390422 |
| 22927 | *HABP4* | ENSG00000130956.14 | 4.77 | 5.64 | 0.87 | 1.83 | 1.96698E-07 | 1.15721E-05 |
| 8573 | *CASK* | ENSG00000147044.23 | 6.34 | 7.21 | 0.87 | 1.82 | 1.1903E-06 | 3.91783E-05 |
| 604 | *BCL6* | ENSG00000113916.18 | 5.12 | 5.98 | 0.87 | 1.82 | 5.11448E-07 | 2.21859E-05 |
| 345757 | *FAM174A* | ENSG00000174132.9 | 4.05 | 4.91 | 0.86 | 1.82 | 4.26634E-06 | 9.78418E-05 |
| 8821 | *INPP4B* | ENSG00000109452.13 | 5.08 | 5.94 | 0.86 | 1.82 | 4.67993E-05 | 0.000561929 |
| 54814 | *QPCTL* | ENSG00000011478.13 | 3.27 | 4.13 | 0.86 | 1.81 | 0.000165939 | 0.001421511 |
| 154141 | *MBOAT1* | ENSG00000172197.11 | 4.39 | 5.25 | 0.86 | 1.81 | 1.55979E-06 | 4.6595E-05 |
| 2057 | *EPOR* | ENSG00000187266.14 | 3.86 | 4.72 | 0.86 | 1.81 | 2.1414E-05 | 0.000316965 |
| 83999 | *KREMEN1* | ENSG00000183762.13 | 6.59 | 7.45 | 0.86 | 1.81 | 6.11108E-07 | 2.47926E-05 |
| 80210 | *ARMC9* | ENSG00000135931.19 | 4.07 | 4.93 | 0.86 | 1.81 | 5.3372E-05 | 0.000617626 |
| 5139 | *PDE3A* | ENSG00000172572.7 | 4.82 | 5.68 | 0.86 | 1.81 | 4.54693E-05 | 0.000548593 |
| 81035 | *COLEC12* | ENSG00000158270.12 | 4.92 | 5.78 | 0.86 | 1.81 | 3.01367E-07 | 1.57056E-05 |
| 128553 | *TSHZ2* | ENSG00000182463.16 | 4.61 | 5.46 | 0.86 | 1.81 | 0.000117243 | 0.001099126 |
| 2673 | *GFPT1* | ENSG00000198380.13 | 7.33 | 8.18 | 0.85 | 1.81 | 1.6109E-07 | 1.00323E-05 |
| 5428 | *POLG* | ENSG00000140521.17 | 5.58 | 6.43 | 0.85 | 1.81 | 0.000118839 | 0.001107861 |
| 404203 | *SPINK6* | ENSG00000178172.7 | 9.04 | 9.89 | 0.85 | 1.80 | 2.36813E-07 | 1.32131E-05 |
| 160518 | *DENND5B* | ENSG00000170456.16 | 6.81 | 7.66 | 0.85 | 1.80 | 1.77701E-07 | 1.08667E-05 |
| 79974 | *CPED1* | ENSG00000106034.18 | 5.79 | 6.65 | 0.85 | 1.80 | 4.91023E-06 | 0.00010833 |
| 2054 | *STX2* | ENSG00000111450.14 | 5.66 | 6.51 | 0.85 | 1.80 | 1.11164E-07 | 7.79656E-06 |
| 1265 | *CNN2* | ENSG00000064666.15 | 3.26 | 4.11 | 0.85 | 1.80 | 0.000808816 | 0.004811541 |
| 6809 | *STX3* | ENSG00000166900.17 | 5.45 | 6.30 | 0.85 | 1.80 | 3.26607E-07 | 1.61135E-05 |
| 8324 | *FZD7* | ENSG00000155760.3 | 6.62 | 7.47 | 0.85 | 1.80 | 2.79311E-08 | 2.9316E-06 |
| 760 | *CA2* | ENSG00000104267.10 | 1.80 | 2.65 | 0.85 | 1.80 | 0.000848929 | 0.004998657 |
| 10209 | *EIF1* | ENSG00000173812.11 | 8.60 | 9.44 | 0.85 | 1.80 | 1.72841E-08 | 2.06241E-06 |
| 58516 | *SINHCAF* | ENSG00000139146.14 | 7.88 | 8.72 | 0.85 | 1.80 | 1.74513E-08 | 2.06241E-06 |
| 5971 | *RELB* | ENSG00000104856.14 | 1.90 | 2.74 | 0.85 | 1.80 | 0.00054604 | 0.003578784 |
| 5062 | *PAK2* | ENSG00000180370.10 | 7.72 | 8.57 | 0.84 | 1.79 | 1.15875E-07 | 7.9414E-06 |
| 5507 | *PPP1R3C* | ENSG00000119938.9 | 3.65 | 4.49 | 0.84 | 1.79 | 1.41529E-05 | 0.000236548 |
| 56975 | *FAM20C* | ENSG00000177706.9 | 4.80 | 5.64 | 0.84 | 1.79 | 0.00014394 | 0.001280026 |
| 121457 | *IKBIP* | ENSG00000166130.15 | 4.03 | 4.88 | 0.84 | 1.79 | 4.58517E-07 | 2.06045E-05 |
| 3099 | *HK2* | ENSG00000159399.10 | 3.54 | 4.38 | 0.84 | 1.79 | 2.12923E-05 | 0.000315475 |
| 6016 | *RIT1* | ENSG00000143622.11 | 6.31 | 7.15 | 0.84 | 1.79 | 3.0555E-07 | 1.57074E-05 |
| 6659 | *SOX4* | ENSG00000124766.7 | 6.09 | 6.93 | 0.84 | 1.79 | 3.67106E-08 | 3.64893E-06 |
| 4189 | *DNAJB9* | ENSG00000128590.5 | 5.88 | 6.71 | 0.84 | 1.79 | 1.1583E-06 | 3.85971E-05 |
| 5805 | *PTS* | ENSG00000150787.8 | 5.10 | 5.94 | 0.84 | 1.79 | 2.47714E-07 | 1.36689E-05 |
| 8639 | *AOC3* | ENSG00000131471.7 | 2.75 | 3.59 | 0.84 | 1.78 | 3.48356E-05 | 0.000451509 |
| 100507436 | *MICA* | ENSG00000204520.14 | 4.44 | 5.27 | 0.84 | 1.78 | 8.72011E-07 | 3.13404E-05 |
| 117154 | *DACH2* | ENSG00000126733.22 | 5.41 | 6.25 | 0.83 | 1.78 | 1.09546E-07 | 7.76114E-06 |
| 2791 | *GNG11* | ENSG00000127920.6 | 6.36 | 7.20 | 0.83 | 1.78 | 2.76391E-07 | 1.47628E-05 |
| 817 | *CAMK2D* | ENSG00000145349.18 | 4.24 | 5.08 | 0.83 | 1.78 | 1.3854E-06 | 4.3501E-05 |
| 25897 | *RNF19A* | ENSG00000034677.13 | 6.01 | 6.84 | 0.83 | 1.78 | 1.26194E-07 | 8.45557E-06 |

| | | | | | | | | |
|---|---|---|---|---|---|---|---|---|
| 254251 | *LCORL* | ENSG00000178177.16 | 6.01 | 6.84 | 0.83 | 1.78 | 4.08142E-07 | 1.88486E-05 |
| 2026 | *ENO2* | ENSG00000111674.9 | 3.97 | 4.80 | 0.83 | 1.78 | 2.23957E-05 | 0.000327939 |
| 813 | *CALU* | ENSG00000128595.17 | 9.44 | 10.27 | 0.83 | 1.78 | 2.58386E-09 | 5.46214E-07 |
| 79693 | *YRDC* | ENSG00000196449.4 | 3.99 | 4.82 | 0.83 | 1.78 | 6.97494E-06 | 0.000139454 |
| 7376 | *NR1H2* | ENSG00000131408.15 | 6.02 | 6.86 | 0.83 | 1.78 | 9.74813E-07 | 3.41048E-05 |
| 60682 | *SMAP1* | ENSG00000112305.15 | 6.30 | 7.13 | 0.83 | 1.78 | 1.22257E-08 | 1.61332E-06 |
| 55075 | *UACA* | ENSG00000137831.15 | 6.92 | 7.75 | 0.83 | 1.78 | 3.08493E-07 | 1.57489E-05 |
| 6890 | *TAP1* | ENSG00000168394.12 | 5.52 | 6.35 | 0.83 | 1.78 | 3.1456E-07 | 1.58731E-05 |
| 1573 | *CYP2J2* | ENSG00000134716.11 | 2.95 | 3.78 | 0.83 | 1.78 | 6.2254E-05 | 0.000692925 |
| 84295 | *PHF6* | ENSG00000156531.18 | 6.58 | 7.41 | 0.83 | 1.77 | 4.84525E-08 | 4.40742E-06 |
| 80019 | *UBTD1* | ENSG00000165886.5 | 1.70 | 2.53 | 0.83 | 1.77 | 0.000208095 | 0.001690198 |
| 7253 | *TSHR* | ENSG00000165409.18 | 4.71 | 5.53 | 0.83 | 1.77 | 1.27412E-05 | 0.000219303 |
| 153241 | *CEP120* | ENSG00000168944.17 | 5.24 | 6.06 | 0.82 | 1.77 | 8.2527E-08 | 6.22436E-06 |
| 483 | *ATP1B3* | ENSG00000069849.11 | 6.55 | 7.37 | 0.82 | 1.77 | 2.64157E-07 | 1.43966E-05 |
| 2296 | *FOXC1* | ENSG00000054598.9 | 3.05 | 3.87 | 0.82 | 1.77 | 0.000282409 | 0.002149431 |
| 25843 | *MOB4* | ENSG00000115540.15 | 8.64 | 9.46 | 0.82 | 1.77 | 4.81594E-09 | 8.50382E-07 |
| 54541 | *DDIT4* | ENSG00000168209.6 | 3.96 | 4.78 | 0.82 | 1.77 | 0.001492421 | 0.00775883 |
| 84343 | *HPS3* | ENSG00000163755.9 | 6.49 | 7.30 | 0.82 | 1.76 | 5.08305E-08 | 4.59587E-06 |
| 9415 | *FADS2* | ENSG00000134824.14 | 3.51 | 4.33 | 0.82 | 1.76 | 3.03992E-05 | 0.00040774 |
| 3133 | *HLA-E* | ENSG00000204592.9 | 5.96 | 6.78 | 0.82 | 1.76 | 2.42996E-07 | 1.35079E-05 |
| 7702 | *ZNF143* | ENSG00000166478.10 | 3.58 | 4.40 | 0.82 | 1.76 | 5.78126E-05 | 0.000657287 |
| 5252 | *PHF1* | ENSG00000112511.18 | 5.38 | 6.20 | 0.81 | 1.76 | 3.58391E-06 | 8.57908E-05 |
| 26502 | *NARF* | ENSG00000141562.19 | 4.88 | 5.70 | 0.81 | 1.76 | 3.97812E-07 | 1.84853E-05 |
| 647135 | *SRGAP2B* | ENSG00000196369.12 | 6.54 | 7.35 | 0.81 | 1.76 | 4.32448E-07 | 1.97283E-05 |
| 9143 | *SYNGR3* | ENSG00000127561.16 | 2.17 | 2.98 | 0.81 | 1.75 | 0.001617762 | 0.008236429 |
| 388 | *RHOB* | ENSG00000143878.10 | 7.48 | 8.29 | 0.81 | 1.75 | 2.01167E-07 | 1.17028E-05 |
| 10769 | *PLK2* | ENSG00000145632.15 | 5.62 | 6.43 | 0.81 | 1.75 | 2.66969E-05 | 0.000371013 |
| 285704 | *RGMB* | ENSG00000174136.13 | 5.00 | 5.81 | 0.81 | 1.75 | 1.26009E-06 | 4.076E-05 |
| 84163 | *GTF2IRD2* | ENSG00000196275.16 | 2.65 | 3.45 | 0.81 | 1.75 | 0.001792816 | 0.008915962 |
| 4000 | *LMNA* | ENSG00000160789.24 | 8.08 | 8.89 | 0.80 | 1.75 | 3.75625E-08 | 3.70905E-06 |
| 203228 | *C9orf72* | ENSG00000147894.17 | 3.86 | 4.66 | 0.80 | 1.74 | 0.000440398 | 0.003028617 |
| 80315 | *CPEB4* | ENSG00000113742.14 | 6.72 | 7.52 | 0.80 | 1.74 | 4.75794E-08 | 4.3811E-06 |
| 285590 | *SH3PXD2B* | ENSG00000174705.13 | 6.11 | 6.91 | 0.80 | 1.74 | 2.64764E-07 | 1.43966E-05 |
| 23603 | *CORO1C* | ENSG00000110880.11 | 8.01 | 8.81 | 0.80 | 1.74 | 6.01013E-08 | 5.09417E-06 |
| 4126 | *MANBA* | ENSG00000109323.11 | 4.59 | 5.39 | 0.80 | 1.74 | 3.6265E-05 | 0.000468418 |
| 221914 | *GPC2* | ENSG00000213420.8 | 3.11 | 3.91 | 0.80 | 1.74 | 0.000348169 | 0.002525695 |
| 6385 | *SDC4* | ENSG00000124145.6 | 6.03 | 6.82 | 0.80 | 1.74 | 1.83968E-07 | 1.11338E-05 |
| 6627 | *SNRPA1* | ENSG00000131876.18 | 5.21 | 6.01 | 0.80 | 1.74 | 1.36364E-05 | 0.000231526 |
| 2801 | *GOLGA2* | ENSG00000167110.19 | 6.38 | 7.17 | 0.80 | 1.74 | 1.96243E-06 | 5.5365E-05 |
| 29948 | *OSGIN1* | ENSG00000140961.14 | 5.21 | 6.01 | 0.80 | 1.74 | 1.91527E-07 | 1.13174E-05 |
| 81579 | *PLA2G12A* | ENSG00000123739.11 | 5.98 | 6.78 | 0.79 | 1.73 | 6.47131E-07 | 2.57951E-05 |
| 55151 | *TMEM38B* | ENSG00000095209.12 | 6.12 | 6.92 | 0.79 | 1.73 | 3.87492E-07 | 1.82456E-05 |
| 6782 | *HSPA13* | ENSG00000155304.6 | 7.63 | 8.42 | 0.79 | 1.73 | 3.03007E-07 | 1.57074E-05 |
| 901 | *CCNG2* | ENSG00000138764.15 | 4.15 | 4.95 | 0.79 | 1.73 | 0.000260692 | 0.002016869 |
| 23204 | *ARL6IP1* | ENSG00000170540.15 | 9.28 | 10.07 | 0.79 | 1.73 | 1.22539E-08 | 1.61332E-06 |
| 128239 | *IQGAP3* | ENSG00000183856.11 | 6.06 | 6.84 | 0.79 | 1.73 | 2.94407E-07 | 1.55095E-05 |
| 928 | *CD9* | ENSG00000010278.15 | 1.73 | 2.51 | 0.79 | 1.73 | 0.001724805 | 0.008632076 |
| 54206 | *ERRFI1* | ENSG00000116285.13 | 5.17 | 5.96 | 0.79 | 1.72 | 2.16603E-07 | 1.24084E-05 |
| 7791 | *ZYX* | ENSG00000159840.16 | 4.76 | 5.54 | 0.79 | 1.72 | 2.56235E-06 | 6.75892E-05 |
| 9232 | *PTTG1* | ENSG00000164611.13 | 6.94 | 7.73 | 0.78 | 1.72 | 1.11853E-07 | 7.80836E-06 |
| 11237 | *RNF24* | ENSG00000101236.17 | 4.14 | 4.93 | 0.78 | 1.72 | 0.00014881 | 0.001313957 |
| 2170 | *FABP3* | ENSG00000121769.8 | 5.85 | 6.64 | 0.78 | 1.72 | 3.72828E-06 | 8.80888E-05 |
| 11004 | *KIF2C* | ENSG00000142945.13 | 6.26 | 7.04 | 0.78 | 1.72 | 4.13831E-07 | 1.89945E-05 |
| 11014 | *KDELR2* | ENSG00000136240.10 | 7.70 | 8.48 | 0.78 | 1.72 | 3.17336E-08 | 3.26093E-06 |
| 7424 | *VEGFC* | ENSG00000150630.4 | 2.13 | 2.90 | 0.78 | 1.72 | 0.000544577 | 0.003570801 |
| 127396 | *ZNF684* | ENSG00000117010.17 | 2.89 | 3.67 | 0.78 | 1.72 | 0.001121635 | 0.006248932 |
| 6100 | *RP9* | ENSG00000164610.10 | 3.39 | 4.17 | 0.78 | 1.72 | 6.18726E-05 | 0.00069 |
| 2114 | *ETS2* | ENSG00000157557.13 | 3.74 | 4.51 | 0.78 | 1.71 | 2.52938E-05 | 0.000358823 |
| 65018 | *PINK1* | ENSG00000158828.8 | 7.58 | 8.36 | 0.77 | 1.71 | 1.97441E-08 | 2.26214E-06 |
| 5986 | *RFNG* | ENSG00000169733.12 | 3.56 | 4.33 | 0.77 | 1.71 | 0.000396887 | 0.002802619 |
| 5864 | *RAB3A* | ENSG00000105649.10 | 2.82 | 3.59 | 0.77 | 1.71 | 5.61104E-05 | 0.000640914 |
| 23352 | *UBR4* | ENSG00000127481.15 | 6.87 | 7.65 | 0.77 | 1.71 | 0.000808646 | 0.004811541 |
| 8459 | *TPST2* | ENSG00000128294.16 | 5.07 | 5.84 | 0.77 | 1.70 | 7.01877E-07 | 2.69424E-05 |
| 7538 | *ZFP36* | ENSG00000128016.7 | 2.79 | 3.56 | 0.77 | 1.70 | 0.001331398 | 0.007111368 |
| 91750 | *LIN52* | ENSG00000205659.12 | 3.46 | 4.22 | 0.77 | 1.70 | 3.79088E-05 | 0.000482999 |
| 1058 | *CENPA* | ENSG00000115163.15 | 4.76 | 5.53 | 0.77 | 1.70 | 7.10084E-07 | 2.71879E-05 |
| 9167 | *COX7A2L* | ENSG00000115944.15 | 6.43 | 7.20 | 0.77 | 1.70 | 1.45993E-06 | 4.46274E-05 |
| 23433 | *RHOQ* | ENSG00000119729.12 | 5.94 | 6.70 | 0.77 | 1.70 | 5.41792E-07 | 2.29064E-05 |
| 9553 | *MRPL33* | ENSG00000243147.8 | 5.60 | 6.37 | 0.77 | 1.70 | 2.82713E-06 | 7.21639E-05 |
| 4509 | *ATP8* | ENSG00000228253.1 | 7.86 | 8.62 | 0.77 | 1.70 | 0.000518353 | 0.003434234 |
| 89927 | *BMERB1* | ENSG00000166780.11 | 4.23 | 4.99 | 0.76 | 1.70 | 0.000117806 | 0.001101647 |
| 2697 | *GJA1* | ENSG00000152661.9 | 5.35 | 6.11 | 0.76 | 1.70 | 1.05133E-05 | 0.000191529 |
| 4758 | *NEU1* | ENSG00000204386.12 | 5.98 | 6.75 | 0.76 | 1.70 | 1.49282E-07 | 9.49398E-06 |
| 6747 | *SSR3* | ENSG00000114850.7 | 7.14 | 7.91 | 0.76 | 1.70 | 1.02165E-06 | 3.54368E-05 |
| 2195 | *FAT1* | ENSG00000083857.15 | 7.86 | 8.62 | 0.76 | 1.70 | 7.96647E-05 | 0.000834395 |
| 5066 | *PAM* | ENSG00000145730.21 | 9.09 | 9.85 | 0.76 | 1.69 | 1.46466E-08 | 1.81679E-06 |
| 10783 | *NEK6* | ENSG00000119408.17 | 5.37 | 6.13 | 0.76 | 1.69 | 3.797E-06 | 8.89066E-05 |
| 57186 | *RALGAPA2* | ENSG00000188559.16 | 6.97 | 7.73 | 0.76 | 1.69 | 3.7328E-05 | 0.000476814 |
| 152185 | *SPICE1* | ENSG00000163611.11 | 3.81 | 4.56 | 0.76 | 1.69 | 6.63606E-05 | 0.00072914 |
| 6478 | *SIAH2* | ENSG00000181788.4 | 9.34 | 10.10 | 0.76 | 1.69 | 5.70299E-08 | 4.86342E-06 |
| 79956 | *ERMP1* | ENSG00000099219.15 | 3.95 | 4.71 | 0.76 | 1.69 | 2.36836E-05 | 0.000340812 |
| 1163 | *CKS1B* | ENSG00000173207.13 | 7.09 | 7.85 | 0.75 | 1.69 | 7.19492E-08 | 5.86571E-06 |
| 102157402 | *AK6* | ENSG00000085231.14 | 5.20 | 5.95 | 0.75 | 1.68 | 0.000196848 | 0.001622454 |
| 55829 | *SELENOS* | ENSG00000131871.15 | 5.45 | 6.20 | 0.75 | 1.68 | 7.16525E-07 | 2.72952E-05 |
| 23314 | *SATB2* | ENSG00000119042.17 | 2.79 | 3.53 | 0.75 | 1.68 | 0.000530958 | 0.003501382 |
| 59 | *ACTA2* | ENSG00000107796.13 | 4.68 | 5.42 | 0.75 | 1.68 | 1.19863E-05 | 0.000209798 |
| 54407 | *SLC38A2* | ENSG00000134294.14 | 8.53 | 9.27 | 0.75 | 1.68 | 8.56362E-08 | 6.3946E-06 |

| | | | | | | | | |
|---|---|---|---|---|---|---|---|---|
| 6376 | *CX3CL1* | ENSG00000006210.7 | 6.71 | 7.46 | 0.75 | 1.68 | 2.45833E-08 | 2.69322E-06 |
| 23162 | *MAPK8IP3* | ENSG00000138834.15 | 4.66 | 5.40 | 0.74 | 1.67 | 0.000245795 | 0.001923429 |
| 8462 | *KLF11* | ENSG00000172059.11 | 5.08 | 5.83 | 0.74 | 1.67 | 7.34511E-07 | 2.78391E-05 |
| 23016 | *EXOSC7* | ENSG00000075914.13 | 4.23 | 4.97 | 0.74 | 1.67 | 1.3107E-05 | 0.000224827 |
| 5997 | *RGS2* | ENSG00000116741.8 | 4.46 | 5.20 | 0.74 | 1.67 | 1.55588E-06 | 4.6595E-05 |
| 151887 | *CCDC80* | ENSG00000091986.16 | 4.95 | 5.68 | 0.74 | 1.67 | 0.000989289 | 0.005639286 |
| 100507602 | *TRIM52-AS1* | ENSG00000248275.3 | 2.57 | 3.31 | 0.74 | 1.67 | 0.00097484 | 0.005573856 |
| 91748 | *MIDEAS* | ENSG00000156030.14 | 4.11 | 4.85 | 0.74 | 1.67 | 4.77004E-05 | 0.000570013 |
| 3858 | *KRT10* | ENSG00000186395.9 | 4.19 | 4.93 | 0.74 | 1.67 | 6.95051E-07 | 2.68866E-05 |
| 6790 | *AURKA* | ENSG00000087586.18 | 6.17 | 6.90 | 0.74 | 1.66 | 5.07497E-07 | 2.20783E-05 |
| 224 | *ALDH3A2* | ENSG00000072210.19 | 8.31 | 9.04 | 0.73 | 1.66 | 2.55394E-08 | 2.71859E-06 |
| 201895 | *SMIM14* | ENSG00000163683.12 | 6.54 | 7.27 | 0.73 | 1.66 | 3.24765E-06 | 8.0171E-05 |
| 3998 | *LMAN1* | ENSG00000074695.6 | 8.67 | 9.40 | 0.73 | 1.66 | 5.86439E-09 | 9.49618E-07 |
| 7456 | *WIPF1* | ENSG00000115935.18 | 4.66 | 5.39 | 0.73 | 1.66 | 4.17564E-05 | 0.000515396 |
| 11142 | *PKIG* | ENSG00000168734.14 | 5.31 | 6.04 | 0.73 | 1.66 | 6.54012E-06 | 0.00013326 |
| 5564 | *PRKAB1* | ENSG00000111725.11 | 5.59 | 6.32 | 0.73 | 1.66 | 4.38366E-06 | 9.95376E-05 |
| 9754 | *STARD8* | ENSG00000130052.14 | 5.44 | 6.17 | 0.73 | 1.66 | 2.31439E-06 | 6.25887E-05 |
| 9572 | *NR1D1* | ENSG00000126368.6 | 4.02 | 4.75 | 0.73 | 1.66 | 5.42652E-05 | 0.000626748 |
| 3764 | *KCNJ8* | ENSG00000121361.5 | 5.30 | 6.03 | 0.73 | 1.66 | 8.25082E-06 | 0.000158765 |
| 10447 | *FAM3C* | ENSG00000196937.11 | 7.97 | 8.70 | 0.73 | 1.66 | 1.81056E-07 | 1.10019E-05 |
| 201164 | *PLD6* | ENSG00000179598.6 | 4.44 | 5.17 | 0.73 | 1.66 | 2.3824E-06 | 6.41965E-05 |
| 8575 | *PRKRA* | ENSG00000180228.14 | 6.15 | 6.88 | 0.73 | 1.66 | 3.87664E-06 | 9.04891E-05 |
| 65249 | *ZSWIM4* | ENSG00000132003.10 | 2.96 | 3.69 | 0.73 | 1.66 | 2.20161E-05 | 0.000323644 |
| 55355 | *HJURP* | ENSG00000123485.12 | 4.87 | 5.59 | 0.73 | 1.65 | 0.000291486 | 0.002206207 |
| 148523 | *CIART* | ENSG00000159208.16 | 3.62 | 4.35 | 0.72 | 1.65 | 0.000784702 | 0.004722104 |
| 26993 | *AKAP8L* | ENSG00000011243.19 | 5.92 | 6.65 | 0.72 | 1.65 | 6.08922E-07 | 2.47926E-05 |
| 7402 | *UTRN* | ENSG00000152818.19 | 6.06 | 6.78 | 0.72 | 1.65 | 0.001635983 | 0.008307498 |
| 57820 | *CCNB1IP1* | ENSG00000100814.18 | 4.83 | 5.55 | 0.72 | 1.65 | 7.81486E-06 | 0.000152527 |
| 7922 | *SLC39A7* | ENSG00000112473.18 | 7.60 | 8.32 | 0.72 | 1.65 | 3.1938E-08 | 3.26093E-06 |
| 3306 | *HSPA2* | ENSG00000126803.10 | 7.32 | 8.04 | 0.72 | 1.65 | 1.06952E-07 | 7.64405E-06 |
| 5569 | *PKIA* | ENSG00000171033.13 | 6.01 | 6.73 | 0.72 | 1.65 | 9.07389E-07 | 3.21962E-05 |
| 6505 | *SLC1A1* | ENSG00000106688.12 | 4.74 | 5.46 | 0.72 | 1.65 | 2.88083E-05 | 0.000391652 |
| 170685 | *NUDT10* | ENSG00000122824.11 | 4.14 | 4.86 | 0.72 | 1.65 | 1.23951E-06 | 4.04431E-05 |
| 55132 | *LARP1B* | ENSG00000138709.19 | 3.80 | 4.52 | 0.72 | 1.65 | 1.32182E-05 | 0.000225985 |
| 222068 | *TMED4* | ENSG00000158604.15 | 7.57 | 8.29 | 0.72 | 1.65 | 1.25517E-08 | 1.62404E-06 |
| 6443 | *SGCB* | ENSG00000163069.13 | 6.87 | 7.59 | 0.72 | 1.64 | 2.5438E-08 | 2.71859E-06 |
| 3297 | *HSF1* | ENSG00000185122.11 | 4.60 | 5.32 | 0.72 | 1.64 | 0.000290903 | 0.002202911 |
| 3399 | *ID3* | ENSG00000117318.9 | 2.98 | 3.70 | 0.72 | 1.64 | 6.66767E-05 | 0.000730846 |
| 6652 | *SORD* | ENSG00000140263.15 | 6.99 | 7.70 | 0.72 | 1.64 | 5.0634E-07 | 2.20783E-05 |
| 55802 | *DCP1A* | ENSG00000272886.6 | 5.77 | 6.48 | 0.72 | 1.64 | 6.60209E-06 | 0.000134269 |
| 142891 | *SAMD8* | ENSG00000156671.15 | 5.61 | 6.33 | 0.71 | 1.64 | 4.82908E-07 | 2.13175E-05 |
| 27338 | *UBE2S* | ENSG00000108106.14 | 7.48 | 8.19 | 0.71 | 1.64 | 2.94672E-05 | 0.000398086 |
| 5604 | *MAP2K1* | ENSG00000169032.11 | 7.05 | 7.77 | 0.71 | 1.64 | 5.48179E-08 | 4.8503E-06 |
| 100129550 | *LINC02035* | ENSG00000273033.2 | 4.39 | 5.10 | 0.71 | 1.64 | 8.74671E-05 | 0.000889989 |
| 151648 | *SGO1* | ENSG00000129810.15 | 4.92 | 5.62 | 0.71 | 1.63 | 6.03124E-05 | 0.000677059 |
| 2992 | *GYG1* | ENSG00000163754.18 | 4.98 | 5.69 | 0.71 | 1.63 | 1.02233E-06 | 3.54368E-05 |
| 80014 | *WWC2* | ENSG00000151718.16 | 6.22 | 6.93 | 0.71 | 1.63 | 6.36611E-06 | 0.000130699 |
| 25963 | *TMEM87A* | ENSG00000103978.16 | 7.19 | 7.90 | 0.71 | 1.63 | 9.30739E-07 | 3.28693E-05 |
| 23530 | *NNT* | ENSG00000112992.18 | 8.45 | 9.15 | 0.70 | 1.63 | 0.000175562 | 0.001487864 |
| 401097 | *C3orf80* | ENSG00000180044.6 | 3.13 | 3.83 | 0.70 | 1.63 | 0.000904333 | 0.005254797 |
| 9976 | *CLEC2B* | ENSG00000110852.5 | 5.10 | 5.80 | 0.70 | 1.63 | 1.38673E-05 | 0.000233652 |
| 2585 | *GALK2* | ENSG00000156958.15 | 5.29 | 6.00 | 0.70 | 1.63 | 1.41115E-05 | 0.00023612 |
| 10472 | *ZBTB18* | ENSG00000179456.10 | 3.72 | 4.43 | 0.70 | 1.63 | 0.000271311 | 0.002080793 |
| 8470 | *SORBS2* | ENSG00000154556.18 | 7.44 | 8.15 | 0.70 | 1.63 | 1.09257E-06 | 3.71844E-05 |
| 1316 | *KLF6* | ENSG00000067082.15 | 5.92 | 6.63 | 0.70 | 1.63 | 7.4658E-07 | 2.82252E-05 |
| 221491 | *SMIM29* | ENSG00000186577.14 | 3.20 | 3.90 | 0.70 | 1.63 | 5.50734E-05 | 0.000633407 |
| 10327 | *AKR1A1* | ENSG00000117448.14 | 5.45 | 6.15 | 0.70 | 1.63 | 3.43322E-07 | 1.6567E-05 |
| 6812 | *STXBP1* | ENSG00000136854.24 | 6.04 | 6.74 | 0.70 | 1.62 | 1.6907E-06 | 4.93692E-05 |
| 5121 | *PCP4* | ENSG00000183036.11 | 9.88 | 10.58 | 0.70 | 1.62 | 9.40616E-08 | 6.82488E-06 |
| 699 | *BUB1* | ENSG00000169679.15 | 7.53 | 8.23 | 0.70 | 1.62 | 1.40572E-07 | 9.01643E-06 |
| 3706 | *ITPKA* | ENSG00000137825.11 | 3.41 | 4.11 | 0.70 | 1.62 | 8.95888E-06 | 0.000170423 |
| 9026 | *HIP1R* | ENSG00000130787.14 | 4.95 | 5.64 | 0.70 | 1.62 | 0.000703862 | 0.004333168 |
| 254170 | *FBXO33* | ENSG00000165355.8 | 4.71 | 5.40 | 0.70 | 1.62 | 5.98154E-06 | 0.000124345 |
| 92597 | *MOB1B* | ENSG00000173542.9 | 5.14 | 5.84 | 0.70 | 1.62 | 1.44268E-06 | 4.45054E-05 |
| 51076 | *CUTC* | ENSG00000119929.13 | 3.04 | 3.73 | 0.70 | 1.62 | 0.001582149 | 0.008096311 |
| 54704 | *PDP1* | ENSG00000164951.16 | 5.78 | 6.48 | 0.69 | 1.62 | 2.7586E-06 | 7.10565E-05 |
| 4541 | *ND6* | ENSG00000198695.2 | 5.15 | 5.84 | 0.69 | 1.61 | 0.000256828 | 0.001991078 |
| 22820 | *COPG1* | ENSG00000181789.14 | 7.70 | 8.39 | 0.69 | 1.61 | 4.18795E-07 | 1.91637E-05 |
| 55635 | *DEPDC1* | ENSG00000024526.17 | 6.66 | 7.35 | 0.69 | 1.61 | 4.86184E-06 | 0.000107691 |
| 9567 | *GTPBP1* | ENSG00000100226.16 | 6.43 | 7.12 | 0.69 | 1.61 | 0.000125491 | 0.001152612 |
| 3106 | *HLA-B* | ENSG00000234745.13 | 6.58 | 7.26 | 0.69 | 1.61 | 7.74142E-07 | 2.89753E-05 |
| 11138 | *TBC1D8* | ENSG00000204634.13 | 3.26 | 3.95 | 0.69 | 1.61 | 0.001923417 | 0.009415708 |
| 11177 | *BAZ1A* | ENSG00000198604.11 | 4.54 | 5.23 | 0.69 | 1.61 | 1.70487E-05 | 0.000267661 |
| 7692 | *ZNF133* | ENSG00000125846.16 | 3.86 | 4.55 | 0.68 | 1.61 | 0.000122776 | 0.001131911 |
| 6396 | *SEC13* | ENSG00000157020.18 | 6.67 | 7.36 | 0.68 | 1.61 | 2.96377E-07 | 1.55095E-05 |
| 11186 | *RASSF1* | ENSG00000068028.18 | 3.19 | 3.88 | 0.68 | 1.61 | 3.83111E-05 | 0.000486473 |
| 2191 | *FAP* | ENSG00000078098.15 | 5.76 | 6.44 | 0.68 | 1.60 | 6.76194E-07 | 2.6361E-05 |
| 3309 | *HSPA5* | ENSG00000044574.9 | 9.61 | 10.29 | 0.68 | 1.60 | 7.56068E-08 | 6.02913E-06 |
| 4008 | *LMO7* | ENSG00000136153.20 | 5.66 | 6.34 | 0.68 | 1.60 | 1.38706E-05 | 0.000233652 |
| 10184 | *LHFPL2* | ENSG00000145685.14 | 5.17 | 5.86 | 0.68 | 1.60 | 1.56155E-06 | 4.6595E-05 |
| 22824 | *HSPA4L* | ENSG00000164070.12 | 5.71 | 6.39 | 0.68 | 1.60 | 1.00299E-06 | 3.49277E-05 |
| 653381 | *SORD2P* | ENSG00000259479.6 | 3.92 | 4.59 | 0.68 | 1.60 | 0.000152673 | 0.001336255 |
| 51023 | *MRPS18C* | ENSG00000163319.11 | 4.23 | 4.91 | 0.68 | 1.60 | 0.000108494 | 0.001042501 |
| 169200 | *TMEM64* | ENSG00000180694.14 | 6.31 | 6.99 | 0.68 | 1.60 | 4.80014E-06 | 0.000106892 |
| 8660 | *IRS2* | ENSG00000185950.9 | 8.66 | 9.33 | 0.68 | 1.60 | 3.99988E-07 | 1.85291E-05 |
| 9451 | *EIF2AK3* | ENSG00000172071.15 | 5.38 | 6.06 | 0.67 | 1.60 | 2.07548E-06 | 5.76867E-05 |
| 9788 | *MTSS1* | ENSG00000170873.19 | 7.02 | 7.70 | 0.67 | 1.60 | 8.64377E-07 | 3.11861E-05 |

| | | | | | | | | |
|---|---|---|---|---|---|---|---|---|
| 360023 | *ZBTB41* | ENSG00000177888.8 | 6.08 | 6.75 | 0.67 | 1.60 | 6.01202E-06 | 0.000124805 |
| 4300 | *MLLT3* | ENSG00000171843.17 | 5.45 | 6.12 | 0.67 | 1.60 | 9.97704E-05 | 0.000982581 |
| 23380 | *SRGAP2* | ENSG00000266028.8 | 6.40 | 7.07 | 0.67 | 1.59 | 3.05587E-07 | 1.57074E-05 |
| 85363 | *TRIM5* | ENSG00000132256.20 | 2.85 | 3.53 | 0.67 | 1.59 | 0.001157893 | 0.006386922 |
| 57026 | *PDXP* | ENSG00000241360.2 | 5.88 | 6.55 | 0.67 | 1.59 | 5.7895E-06 | 0.000122043 |
| 5597 | *MAPK6* | ENSG00000069956.14 | 6.58 | 7.25 | 0.67 | 1.59 | 1.42964E-06 | 4.42423E-05 |
| 11248 | *NXPH3* | ENSG00000182575.8 | 4.86 | 5.53 | 0.67 | 1.59 | 5.81539E-06 | 0.000122074 |
| 50640 | *PNPLA8* | ENSG00000135241.17 | 6.14 | 6.81 | 0.67 | 1.59 | 1.40374E-05 | 0.000235142 |
| 221830 | *POLR1F* | ENSG00000105849.6 | 5.77 | 6.44 | 0.67 | 1.59 | 4.79814E-07 | 2.13063E-05 |
| 79752 | *ZFAND1* | ENSG00000104231.11 | 5.26 | 5.93 | 0.67 | 1.59 | 1.03152E-05 | 0.000188805 |
| 23787 | *MTCH1* | ENSG00000137409.20 | 7.90 | 8.57 | 0.67 | 1.59 | 1.28012E-07 | 8.50149E-06 |
| 9049 | *AIP* | ENSG00000110711.11 | 5.33 | 6.00 | 0.67 | 1.59 | 3.15576E-06 | 7.85486E-05 |
| 375 | *ARF1* | ENSG00000143761.16 | 8.56 | 9.22 | 0.67 | 1.59 | 1.15367E-07 | 7.9414E-06 |
| 5866 | *RAB3IL1* | ENSG00000167994.13 | 3.29 | 3.96 | 0.67 | 1.59 | 0.001835482 | 0.00907702 |
| 64645 | *MFSD14A* | ENSG00000156875.14 | 6.33 | 6.99 | 0.67 | 1.59 | 4.52919E-06 | 0.000102293 |
| 590 | *BCHE* | ENSG00000114200.10 | 7.86 | 8.52 | 0.67 | 1.59 | 3.73273E-06 | 8.80888E-05 |
| 5916 | *RARG* | ENSG00000172819.17 | 5.26 | 5.92 | 0.67 | 1.59 | 0.000263017 | 0.002031717 |
| 4124 | *MAN2A1* | ENSG00000112893.10 | 7.68 | 8.35 | 0.67 | 1.59 | 1.37155E-07 | 8.87311E-06 |
| 57154 | *SMURF1* | ENSG00000198742.10 | 6.42 | 7.08 | 0.66 | 1.59 | 9.81583E-07 | 3.42618E-05 |
| 200734 | *SPRED2* | ENSG00000198369.10 | 3.75 | 4.41 | 0.66 | 1.58 | 0.000635901 | 0.004016935 |
| 51569 | *UFM1* | ENSG00000120686.12 | 5.03 | 5.69 | 0.66 | 1.58 | 1.4665E-06 | 4.46466E-05 |
| 25930 | *PTPN23* | ENSG00000076201.16 | 5.98 | 6.64 | 0.66 | 1.58 | 7.78128E-06 | 0.000152069 |
| 79581 | *SLC52A2* | ENSG00000185803.12 | 4.10 | 4.77 | 0.66 | 1.58 | 4.07215E-05 | 0.000508054 |
| 26354 | *GNL3* | ENSG00000163938.17 | 6.40 | 7.06 | 0.66 | 1.58 | 2.61597E-06 | 6.84626E-05 |
| 3107 | *HLA-C* | ENSG00000204525.18 | 5.27 | 5.94 | 0.66 | 1.58 | 1.39556E-06 | 4.37286E-05 |
| 55198 | *APPL2* | ENSG00000136044.12 | 5.68 | 6.34 | 0.66 | 1.58 | 2.78305E-05 | 0.000383218 |
| 79699 | *ZYG11B* | ENSG00000162378.13 | 6.55 | 7.21 | 0.66 | 1.58 | 6.68689E-07 | 2.62046E-05 |
| 5194 | *PEX13* | ENSG00000162928.9 | 5.06 | 5.72 | 0.66 | 1.58 | 7.34181E-05 | 0.000784852 |
| 5611 | *DNAJC3* | ENSG00000102580.15 | 7.81 | 8.47 | 0.66 | 1.58 | 2.94784E-07 | 1.55095E-05 |
| 56900 | *TMEM167B* | ENSG00000215717.7 | 6.19 | 6.85 | 0.66 | 1.58 | 1.56889E-06 | 4.67212E-05 |
| 25798 | *BRI3* | ENSG00000164713.10 | 4.28 | 4.94 | 0.66 | 1.58 | 3.90159E-05 | 0.000493607 |
| 23558 | *WBP2* | ENSG00000132471.12 | 6.92 | 7.58 | 0.66 | 1.58 | 2.90864E-06 | 7.36186E-05 |
| 5091 | *PC* | ENSG00000173599.15 | 5.54 | 6.19 | 0.66 | 1.58 | 1.12478E-06 | 3.78517E-05 |
| 10484 | *SEC23A* | ENSG00000100934.15 | 7.42 | 8.08 | 0.66 | 1.58 | 7.84299E-08 | 6.13101E-06 |
| 284129 | *SLC26A11* | ENSG00000181045.15 | 5.29 | 5.95 | 0.66 | 1.58 | 0.00011389 | 0.001078483 |
| 23275 | *POFUT2* | ENSG00000186866.17 | 6.36 | 7.02 | 0.66 | 1.58 | 0.000148784 | 0.001313957 |
| 10114 | *HIPK3* | ENSG00000110422.12 | 5.64 | 6.30 | 0.66 | 1.58 | 3.72024E-06 | 8.80711E-05 |
| 10014 | *HDAC5* | ENSG00000108840.16 | 6.39 | 7.05 | 0.66 | 1.58 | 2.61826E-06 | 6.84626E-05 |
| 4440 | *MSI1* | ENSG00000135097.7 | 3.94 | 4.60 | 0.66 | 1.58 | 0.00016093 | 0.001388156 |
| 53916 | *RAB4B* | ENSG00000167578.18 | 2.76 | 3.42 | 0.65 | 1.57 | 0.001307108 | 0.007026643 |
| 10020 | *GNE* | ENSG00000159921.20 | 6.34 | 6.99 | 0.65 | 1.57 | 2.34141E-07 | 1.31128E-05 |
| 78992 | *YIPF2* | ENSG00000130733.11 | 5.49 | 6.14 | 0.65 | 1.57 | 0.000195176 | 0.001611326 |
| 5930 | *RBBP6* | ENSG00000122257.20 | 6.01 | 6.66 | 0.65 | 1.57 | 0.000243157 | 0.001910678 |
| 4864 | *NPC1* | ENSG00000141458.13 | 6.33 | 6.98 | 0.65 | 1.57 | 2.1488E-05 | 0.000317746 |
| 3739 | *KCNA4* | ENSG00000182255.7 | 5.16 | 5.81 | 0.65 | 1.57 | 0.00010674 | 0.001031774 |
| 23231 | *SEL1L3* | ENSG00000091490.11 | 5.29 | 5.94 | 0.65 | 1.57 | 1.254E-05 | 0.000217019 |
| 63982 | *ANO3* | ENSG00000134343.14 | 3.41 | 4.06 | 0.65 | 1.57 | 0.000503223 | 0.003359821 |
| 126731 | *CCSAP* | ENSG00000154429.11 | 5.48 | 6.13 | 0.65 | 1.57 | 5.35964E-06 | 0.000114918 |
| 1875 | *E2F5* | ENSG00000133740.11 | 3.10 | 3.75 | 0.65 | 1.57 | 0.000133618 | 0.001211679 |
| 3598 | *IL13RA2* | ENSG00000123496.8 | 6.52 | 7.17 | 0.65 | 1.57 | 3.15156E-07 | 1.58731E-05 |
| 7006 | *TEC* | ENSG00000135605.13 | 5.26 | 5.91 | 0.65 | 1.57 | 6.11005E-05 | 0.000684371 |
| 51278 | *IER5* | ENSG00000162783.11 | 3.89 | 4.54 | 0.65 | 1.57 | 4.79803E-05 | 0.000572446 |
| 283991 | *UBALD2* | ENSG00000185262.9 | 4.54 | 5.19 | 0.65 | 1.57 | 1.71497E-05 | 0.000268684 |
| 27244 | *SESN1* | ENSG00000080546.14 | 5.06 | 5.71 | 0.65 | 1.57 | 1.4743E-05 | 0.000241834 |
| 55924 | *INKA2* | ENSG00000197852.12 | 3.47 | 4.12 | 0.65 | 1.57 | 0.001286302 | 0.006932173 |
| 23345 | *SYNE1* | ENSG00000131018.25 | 7.81 | 8.46 | 0.65 | 1.57 | 1.21654E-06 | 3.97801E-05 |
| 11018 | *TMED1* | ENSG00000099203.7 | 4.99 | 5.64 | 0.65 | 1.57 | 2.41895E-05 | 0.000346101 |
| 9659 | *PDE4DIP* | ENSG00000178104.19 | 5.74 | 6.39 | 0.65 | 1.57 | 3.70157E-06 | 8.77675E-05 |
| 1845 | *DUSP3* | ENSG00000108861.9 | 7.06 | 7.70 | 0.65 | 1.57 | 1.18428E-06 | 3.91756E-05 |
| 4084 | *MXD1* | ENSG00000059728.11 | 3.82 | 4.46 | 0.65 | 1.57 | 0.000572116 | 0.003715659 |
| 6397 | *SEC14L1* | ENSG00000129657.16 | 7.78 | 8.43 | 0.65 | 1.57 | 1.87079E-07 | 1.12315E-05 |
| 123811 | *CEP20* | ENSG00000133393.13 | 5.84 | 6.48 | 0.65 | 1.57 | 2.72912E-05 | 0.000378222 |
| 7975 | *MAFK* | ENSG00000198517.10 | 3.82 | 4.47 | 0.65 | 1.57 | 0.000937554 | 0.005403899 |
| 25976 | *TIPARP* | ENSG00000163659.13 | 3.93 | 4.58 | 0.65 | 1.57 | 5.61051E-05 | 0.000640914 |
| 3838 | *KPNA2* | ENSG00000182481.10 | 8.56 | 9.20 | 0.65 | 1.57 | 1.31051E-07 | 8.62284E-06 |
| 81537 | *SGPP1* | ENSG00000126821.8 | 6.35 | 6.99 | 0.65 | 1.57 | 1.45264E-06 | 4.45054E-05 |
| 92737 | *DNER* | ENSG00000187957.8 | 4.39 | 5.04 | 0.65 | 1.56 | 0.000130921 | 0.001194526 |
| 100996763 | *NOTCH2NLB* | ENSG00000286019.1 | 2.51 | 3.16 | 0.64 | 1.56 | 0.000843662 | 0.004971545 |
| 355 | *FAS* | ENSG00000026103.23 | 4.41 | 5.05 | 0.64 | 1.56 | 3.96269E-05 | 0.000499799 |
| 27248 | *ERLEC1* | ENSG00000068912.15 | 7.01 | 7.66 | 0.64 | 1.56 | 9.16634E-08 | 6.7111E-06 |
| 10438 | *C1D* | ENSG00000197223.12 | 4.09 | 4.73 | 0.64 | 1.56 | 1.73236E-05 | 0.000269999 |
| 64783 | *RBM15* | ENSG00000162775.17 | 4.60 | 5.24 | 0.64 | 1.56 | 4.83739E-06 | 0.000107562 |
| 55270 | *NUDT15* | ENSG00000136159.4 | 5.06 | 5.70 | 0.64 | 1.56 | 7.00675E-07 | 2.69424E-05 |
| 79139 | *DERL1* | ENSG00000136986.10 | 7.42 | 8.06 | 0.64 | 1.56 | 3.99363E-05 | 0.000502671 |
| 1534 | *CYB561* | ENSG00000008283.16 | 6.74 | 7.38 | 0.64 | 1.56 | 2.64246E-06 | 6.89433E-05 |
| 63925 | *ZNF335* | ENSG00000198026.8 | 3.90 | 4.54 | 0.64 | 1.56 | 0.00017368 | 0.001476921 |
| 151126 | *ZNF385B* | ENSG00000144331.20 | 2.52 | 3.15 | 0.64 | 1.56 | 0.000481829 | 0.003245859 |
| 146760 | *RTN4RL1* | ENSG00000185924.7 | 3.37 | 4.01 | 0.64 | 1.56 | 0.000377685 | 0.002689123 |
| 9262 | *STK17B* | ENSG00000081320.11 | 6.08 | 6.72 | 0.64 | 1.55 | 2.67879E-06 | 6.95469E-05 |
| 4085 | *MAD2L1* | ENSG00000164109.14 | 6.73 | 7.37 | 0.64 | 1.55 | 3.54708E-05 | 0.00045895 |
| 84679 | *SLC9A7* | ENSG00000065923.10 | 4.66 | 5.29 | 0.64 | 1.55 | 0.001899359 | 0.009322262 |
| 51533 | *PHF7* | ENSG00000010318.22 | 3.00 | 3.64 | 0.64 | 1.55 | 0.000180474 | 0.001510727 |
| 84542 | *SANBR* | ENSG00000162929.14 | 3.12 | 3.76 | 0.63 | 1.55 | 0.000119554 | 0.001111762 |
| 8214 | *DGCR6* | ENSG00000183628.14 | 3.95 | 4.58 | 0.63 | 1.55 | 2.05469E-05 | 0.000308081 |
| 100303755 | *PET117* | ENSG00000232838.4 | 3.25 | 3.89 | 0.63 | 1.55 | 0.000797271 | 0.004769214 |
| 10957 | *PNRC1* | ENSG00000146278.11 | 6.37 | 7.00 | 0.63 | 1.55 | 9.9031E-06 | 0.000182599 |
| 143888 | *POGLUT3* | ENSG00000178202.13 | 6.25 | 6.88 | 0.63 | 1.55 | 3.30687E-07 | 1.61986E-05 |

| | | | | | | | | |
|---|---|---|---|---|---|---|---|---|
| 5144 | *PDE4D* | ENSG00000113448.20 | 5.87 | 6.50 | 0.63 | 1.55 | 9.43912E-05 | 0.000941967 |
| 348487 | *FAM131C* | ENSG00000185519.9 | 2.33 | 2.96 | 0.63 | 1.55 | 0.001755022 | 0.008762852 |
| 81691 | *REXO5* | ENSG00000005189.20 | 3.75 | 4.38 | 0.63 | 1.55 | 3.96746E-05 | 0.00049998 |
| 283643 | *TEDC1* | ENSG00000185347.18 | 3.12 | 3.76 | 0.63 | 1.55 | 0.00062028 | 0.003951521 |
| 159013 | *CXorf38* | ENSG00000185753.13 | 4.35 | 4.98 | 0.63 | 1.55 | 0.000313601 | 0.002333585 |
| 291 | *SLC25A4* | ENSG00000151729.11 | 4.76 | 5.39 | 0.63 | 1.55 | 1.9111E-05 | 0.000290027 |
| 5007 | *OSBP* | ENSG00000110048.12 | 7.76 | 8.39 | 0.63 | 1.55 | 1.37E-07 | 8.87311E-06 |
| 1974 | *EIF4A2* | ENSG00000156976.17 | 8.44 | 9.07 | 0.63 | 1.55 | 4.08677E-08 | 3.9069E-06 |
| 53358 | *SHC3* | ENSG00000148082.10 | 3.77 | 4.40 | 0.63 | 1.54 | 0.001369132 | 0.007260152 |
| 64746 | *ACBD3* | ENSG00000182827.9 | 6.44 | 7.06 | 0.63 | 1.54 | 6.99095E-07 | 2.69424E-05 |
| 23321 | *TRIM2* | ENSG00000109654.16 | 6.09 | 6.72 | 0.63 | 1.54 | 7.73662E-06 | 0.000151591 |
| 56005 | *MYDGF* | ENSG00000074842.8 | 5.98 | 6.61 | 0.63 | 1.54 | 5.58665E-07 | 2.34218E-05 |
| 10678 | *B3GNT2* | ENSG00000170340.11 | 3.86 | 4.49 | 0.62 | 1.54 | 0.000116876 | 0.001096367 |
| 55785 | *FGD6* | ENSG00000180263.14 | 4.75 | 5.38 | 0.62 | 1.54 | 8.77743E-05 | 0.000891342 |
| 50854 | *SNHG32* | ENSG00000204387.14 | 5.89 | 6.51 | 0.62 | 1.54 | 1.03525E-06 | 3.57196E-05 |
| 10383 | *TUBB4B* | ENSG00000188229.6 | 9.03 | 9.65 | 0.62 | 1.54 | 1.17221E-07 | 7.97942E-06 |
| 30850 | *CDR2L* | ENSG00000109089.7 | 3.05 | 3.67 | 0.62 | 1.54 | 0.000580074 | 0.003755968 |
| 80021 | *TMEM62* | ENSG00000137842.7 | 4.23 | 4.85 | 0.62 | 1.54 | 0.000184533 | 0.001536993 |
| 55738 | *ARFGAP1* | ENSG00000101199.13 | 4.42 | 5.04 | 0.62 | 1.54 | 0.001045434 | 0.005898842 |
| 51729 | *WBP11* | ENSG00000084463.8 | 7.39 | 8.01 | 0.62 | 1.54 | 0.00047805 | 0.003226196 |
| 79801 | *SHCBP1* | ENSG00000171241.9 | 6.63 | 7.24 | 0.62 | 1.53 | 6.14224E-05 | 0.000686952 |
| 58505 | *OSTC* | ENSG00000198856.13 | 6.34 | 6.96 | 0.62 | 1.53 | 4.85228E-06 | 0.000107691 |
| 51439 | *FAM8A1* | ENSG00000137414.7 | 5.63 | 6.24 | 0.62 | 1.53 | 2.99147E-06 | 7.52076E-05 |
| 10063 | *COX17* | ENSG00000138495.7 | 5.34 | 5.96 | 0.62 | 1.53 | 3.11905E-05 | 0.000415138 |
| 7111 | *TMOD1* | ENSG00000136842.14 | 5.30 | 5.91 | 0.62 | 1.53 | 2.22378E-06 | 6.06707E-05 |
| 10724 | *OGA* | ENSG00000198408.14 | 6.58 | 7.20 | 0.62 | 1.53 | 3.21251E-06 | 7.95653E-05 |
| 2309 | *FOXO3* | ENSG00000118689.15 | 5.52 | 6.14 | 0.62 | 1.53 | 3.66955E-06 | 8.74226E-05 |
| 9949 | *AMMECR1* | ENSG00000101935.12 | 4.15 | 4.77 | 0.62 | 1.53 | 0.000124576 | 0.001146393 |
| 96459 | *FNIP1* | ENSG00000217128.13 | 5.40 | 6.02 | 0.61 | 1.53 | 3.50801E-06 | 8.44894E-05 |
| 347902 | *AMIGO2* | ENSG00000139211.7 | 5.34 | 5.95 | 0.61 | 1.53 | 6.40867E-06 | 0.000131046 |
| 4976 | *OPA1* | ENSG00000198836.11 | 6.79 | 7.40 | 0.61 | 1.53 | 1.5885E-05 | 0.000255813 |
| 84138 | *SLC7A6OS* | ENSG00000103061.13 | 4.11 | 4.72 | 0.61 | 1.53 | 0.001311792 | 0.007044031 |
| 1212 | *CLTB* | ENSG00000175416.16 | 5.08 | 5.70 | 0.61 | 1.53 | 5.56428E-05 | 0.000637531 |
| 10979 | *FERMT2* | ENSG00000073712.15 | 5.59 | 6.20 | 0.61 | 1.53 | 2.30937E-05 | 0.000334569 |
| 55972 | *SLC25A40* | ENSG00000075303.13 | 6.12 | 6.73 | 0.61 | 1.53 | 0.000131994 | 0.00120067 |
| 147657 | *ZNF480* | ENSG00000198464.14 | 3.82 | 4.44 | 0.61 | 1.53 | 0.001194799 | 0.006547185 |
| 134359 | *POC5* | ENSG00000152359.15 | 3.92 | 4.53 | 0.61 | 1.53 | 7.21293E-05 | 0.000774384 |
| 734 | *OSGIN2* | ENSG00000164823.11 | 5.12 | 5.73 | 0.61 | 1.53 | 9.11271E-06 | 0.000172028 |
| 2805 | *GOT1* | ENSG00000120053.12 | 6.59 | 7.20 | 0.61 | 1.52 | 2.09394E-06 | 5.80923E-05 |
| 7423 | *VEGFB* | ENSG00000173511.10 | 5.60 | 6.21 | 0.61 | 1.52 | 5.85608E-05 | 0.00066335 |
| 10026 | *PIGK* | ENSG00000142892.15 | 5.66 | 6.27 | 0.61 | 1.52 | 9.56055E-05 | 0.000950293 |
| 80219 | *COQ10B* | ENSG00000115520.9 | 4.18 | 4.78 | 0.61 | 1.52 | 0.000118072 | 0.001103449 |
| 100526842 | *RPL17-C18orf32* | ENSG00000215472.10 | 5.05 | 5.65 | 0.61 | 1.52 | 3.09053E-05 | 0.000412318 |
| 25840 | *METTL7A* | ENSG00000185432.12 | 7.15 | 7.76 | 0.61 | 1.52 | 4.60604E-06 | 0.000103646 |
| 8613 | *PLPP3* | ENSG00000162407.9 | 4.40 | 5.00 | 0.61 | 1.52 | 6.66314E-05 | 0.000730846 |
| 8702 | *B4GALT4* | ENSG00000121578.13 | 6.20 | 6.80 | 0.60 | 1.52 | 2.53347E-05 | 0.000359046 |
| 5167 | *ENPP1* | ENSG00000197594.14 | 6.43 | 7.04 | 0.60 | 1.52 | 1.03058E-06 | 3.56406E-05 |
| 57563 | *KLHL8* | ENSG00000145332.14 | 4.81 | 5.41 | 0.60 | 1.52 | 0.000672728 | 0.004177482 |
| 9516 | *LITAF* | ENSG00000189067.14 | 5.36 | 5.96 | 0.60 | 1.52 | 0.000666893 | 0.004151553 |
| 1486 | *CTBS* | ENSG00000117151.13 | 4.80 | 5.40 | 0.60 | 1.52 | 2.06852E-05 | 0.000309535 |
| 55379 | *LRRC59* | ENSG00000108829.10 | 7.99 | 8.59 | 0.60 | 1.52 | 1.10158E-07 | 7.76229E-06 |
| 91272 | *BOD1* | ENSG00000145919.11 | 5.60 | 6.20 | 0.60 | 1.52 | 3.72869E-05 | 0.00047678 |
| 51299 | *NRN1* | ENSG00000124785.9 | 3.99 | 4.59 | 0.60 | 1.52 | 0.000143959 | 0.001280026 |
| 7461 | *CLIP2* | ENSG00000106665.16 | 5.04 | 5.64 | 0.60 | 1.52 | 0.00115177 | 0.006367795 |
| 54918 | *CMTM6* | ENSG00000091317.8 | 7.27 | 7.87 | 0.60 | 1.52 | 3.43602E-07 | 1.6567E-05 |
| 158399 | *ZNF483* | ENSG00000173258.13 | 4.35 | 4.95 | 0.60 | 1.52 | 0.001829722 | 0.009060474 |
| 6836 | *SURF4* | ENSG00000148248.14 | 8.73 | 9.33 | 0.60 | 1.52 | 2.85285E-05 | 0.000389369 |
| 221079 | *ARL5B* | ENSG00000165997.5 | 6.70 | 7.30 | 0.60 | 1.52 | 1.04518E-06 | 3.59797E-05 |
| 25777 | *SUN2* | ENSG00000100242.16 | 7.51 | 8.11 | 0.60 | 1.51 | 1.59705E-06 | 4.74655E-05 |
| 2744 | *GLS* | ENSG00000115419.13 | 5.65 | 6.25 | 0.60 | 1.51 | 0.000551347 | 0.003604164 |
| 79101 | *TAF1D* | ENSG00000166012.17 | 4.93 | 5.52 | 0.60 | 1.51 | 2.74319E-05 | 0.000379535 |
| 757 | *TMEM50B* | ENSG00000142188.17 | 5.41 | 6.01 | 0.60 | 1.51 | 2.34336E-06 | 6.32582E-05 |
| 54187 | *NANS* | ENSG00000095380.11 | 7.00 | 7.59 | 0.60 | 1.51 | 2.71691E-07 | 1.46684E-05 |
| 5716 | *PSMD10* | ENSG00000101843.19 | 6.25 | 6.85 | 0.60 | 1.51 | 1.41846E-05 | 0.000236816 |
| 79786 | *KLHL36* | ENSG00000135686.13 | 6.36 | 6.96 | 0.60 | 1.51 | 1.76343E-06 | 5.11941E-05 |
| 221424 | *LRRC73* | ENSG00000204052.4 | 3.69 | 4.28 | 0.59 | 1.51 | 0.000504505 | 0.003365383 |
| 85027 | *SMIM3* | ENSG00000256235.3 | 2.77 | 3.36 | 0.59 | 1.51 | 0.000462063 | 0.003146063 |
| 51302 | *CYP39A1* | ENSG00000146233.8 | 4.14 | 4.73 | 0.59 | 1.51 | 6.34129E-05 | 0.000703968 |
| 65986 | *ZBTB10* | ENSG00000205189.12 | 5.26 | 5.86 | 0.59 | 1.51 | 9.33091E-06 | 0.000174841 |
| 100526694 | *ISANTD3-TMEFF* | ENSG00000251349.3 | 4.24 | 4.83 | 0.59 | 1.51 | 0.000426375 | 0.00296546 |
| 120103 | *SLC36A4* | ENSG00000180773.15 | 6.05 | 6.65 | 0.59 | 1.51 | 2.50421E-06 | 6.64642E-05 |
| 54681 | *P4HTM* | ENSG00000178467.18 | 4.33 | 4.92 | 0.59 | 1.51 | 7.52718E-05 | 0.000798978 |
| 51646 | *YPEL5* | ENSG00000119801.13 | 6.20 | 6.79 | 0.59 | 1.51 | 8.2451E-06 | 0.000158765 |
| 130502 | *TTC32* | ENSG00000183891.6 | 2.88 | 3.46 | 0.59 | 1.51 | 0.001543464 | 0.007960773 |
| 84919 | *PPP1R15B* | ENSG00000158615.10 | 7.07 | 7.66 | 0.59 | 1.51 | 3.09755E-07 | 1.57538E-05 |
| 29803 | *REPIN1* | ENSG00000214022.12 | 7.25 | 7.84 | 0.59 | 1.51 | 7.56342E-07 | 2.8451E-05 |
| 1901 | *S1PR1* | ENSG00000170989.10 | 2.96 | 3.55 | 0.59 | 1.51 | 0.0009509 | 0.005461946 |
| 2803 | *GOLGA4* | ENSG00000144674.17 | 6.69 | 7.28 | 0.59 | 1.51 | 0.000435428 | 0.003007525 |
| 64225 | *ATL2* | ENSG00000119787.14 | 6.35 | 6.95 | 0.59 | 1.51 | 1.183E-05 | 0.00020779 |
| 332 | *BIRC5* | ENSG00000089685.15 | 6.65 | 7.24 | 0.59 | 1.51 | 2.89463E-05 | 0.000392726 |
| 152926 | *PPM1K* | ENSG00000163644.15 | 4.67 | 5.26 | 0.59 | 1.50 | 4.00221E-05 | 0.000502671 |
| 1183 | *CLCN4* | ENSG00000073464.13 | 6.28 | 6.87 | 0.59 | 1.50 | 2.18726E-06 | 5.99061E-05 |
| 135112 | *NCOA7* | ENSG00000111912.20 | 5.40 | 5.99 | 0.59 | 1.50 | 0.0002526 | 0.001968472 |
| 7272 | *TTK* | ENSG00000112742.10 | 6.40 | 6.99 | 0.59 | 1.50 | 5.60875E-06 | 0.000119348 |
| 220213 | *OTUD1* | ENSG00000165312.7 | 3.79 | 4.38 | 0.59 | 1.50 | 9.24036E-05 | 0.000925825 |
| 728769 | *SCAMP1-AS1* | ENSG00000245556.4 | 3.05 | 3.64 | 0.59 | 1.50 | 0.00023474 | 0.001864136 |
| 6541 | *SLC7A1* | ENSG00000139514.13 | 6.42 | 7.01 | 0.59 | 1.50 | 3.78524E-06 | 8.89066E-05 |

| | | | | | | | | |
|---|---|---|---|---|---|---|---|---|
| 81567 | *TXNDC5* | ENSG00000239264.9 | 8.10 | 8.69 | 0.59 | 1.50 | 1.72971E-07 | 1.06398E-05 |
| 10289 | *EIF1B* | ENSG00000114784.4 | 6.23 | 6.82 | 0.59 | 1.50 | 1.85195E-06 | 5.29447E-05 |
| 3920 | *LAMP2* | ENSG00000005893.16 | 7.69 | 8.28 | 0.59 | 1.50 | 3.27444E-07 | 1.61135E-05 |
| 6925 | *TCF4* | ENSG00000196628.20 | 7.91 | 8.50 | 0.59 | 1.50 | 1.6126E-06 | 4.75972E-05 |
| 6884 | *TAF13* | ENSG00000197780.11 | 4.64 | 5.23 | 0.59 | 1.50 | 0.000102828 | 0.001006037 |
| 9819 | *TSC22D2* | ENSG00000196428.13 | 5.41 | 6.00 | 0.59 | 1.50 | 0.000197208 | 0.001624532 |
| 283635 | *FAM177A1* | ENSG00000151327.13 | 4.53 | 5.12 | 0.59 | 1.50 | 3.72046E-05 | 0.00047678 |
| 162962 | *ZNF836* | ENSG00000196267.14 | 5.64 | 6.22 | 0.58 | 1.50 | 9.31987E-05 | 0.000932547 |
| 2531 | *KDSR* | ENSG00000119537.18 | 6.31 | 6.90 | 0.58 | 1.50 | 4.92078E-05 | 0.000582921 |
| 8407 | *TAGLN2* | ENSG00000158710.15 | 7.77 | 8.35 | 0.58 | 1.50 | 7.19489E-07 | 2.73388E-05 |
| 55366 | *LGR4* | ENSG00000205213.14 | 5.09 | 5.67 | 0.58 | 1.50 | 0.001502858 | 0.007795559 |
| 55504 | *TNFRSF19* | ENSG00000127863.16 | 7.74 | 8.32 | 0.58 | 1.50 | 2.17593E-07 | 1.24177E-05 |
| 51171 | *HSD17B14* | ENSG00000087076.9 | 3.25 | 3.83 | 0.58 | 1.50 | 0.000140614 | 0.001257741 |
| 196 | *AHR* | ENSG00000106546.14 | 5.15 | 5.73 | 0.58 | 1.50 | 0.001314276 | 0.007049971 |
| 153339 | *TMEM167A* | ENSG00000174695.10 | 7.51 | 8.09 | 0.58 | 1.50 | 2.16657E-05 | 0.000319744 |
| 80347 | *COASY* | ENSG00000068120.15 | 6.47 | 7.05 | 0.58 | 1.49 | 2.77485E-06 | 7.11927E-05 |
| 57720 | *GPR107* | ENSG00000148358.20 | 7.86 | 8.43 | 0.58 | 1.49 | 7.87713E-07 | 2.93369E-05 |
| 56929 | *FEM1C* | ENSG00000145780.8 | 5.62 | 6.20 | 0.58 | 1.49 | 1.83014E-06 | 5.25212E-05 |
| 110599564 | *EEF1AKMT4* | ENSG00000284753.2 | 2.76 | 3.34 | 0.58 | 1.49 | 0.00114312 | 0.006342732 |
| 56650 | *CLDND1* | ENSG00000080822.17 | 6.85 | 7.43 | 0.58 | 1.49 | 3.18907E-06 | 7.91273E-05 |
| 8440 | *NCK2* | ENSG00000071051.14 | 6.01 | 6.58 | 0.58 | 1.49 | 2.26964E-05 | 0.000330728 |
| 4888 | *NPY6R* | ENSG00000226306.7 | 8.07 | 8.65 | 0.58 | 1.49 | 2.03765E-07 | 1.18081E-05 |
| 23024 | *PDZRN3* | ENSG00000121440.15 | 7.75 | 8.32 | 0.58 | 1.49 | 6.5926E-07 | 2.60391E-05 |
| 2131 | *EXT1* | ENSG00000182197.13 | 5.49 | 6.07 | 0.58 | 1.49 | 4.36682E-05 | 0.000531562 |
| 7430 | *EZR* | ENSG00000092820.19 | 6.25 | 6.83 | 0.58 | 1.49 | 5.48765E-07 | 2.3136E-05 |
| 143662 | *MUC15* | ENSG00000169550.14 | 6.66 | 7.24 | 0.57 | 1.49 | 1.08869E-06 | 3.71368E-05 |
| 4245 | *MGAT1* | ENSG00000131446.17 | 6.26 | 6.83 | 0.57 | 1.49 | 2.93685E-05 | 0.000397507 |
| 26051 | *PPP1R16B* | ENSG00000101445.10 | 3.96 | 4.53 | 0.57 | 1.49 | 0.000714998 | 0.004383745 |
| 81566 | *CSRNP2* | ENSG00000110925.7 | 6.28 | 6.85 | 0.57 | 1.49 | 5.81487E-06 | 0.000122074 |
| 4508 | *ATP6* | ENSG00000198899.2 | 9.70 | 10.27 | 0.57 | 1.49 | 0.00103503 | 0.005851376 |
| 399818 | *EEF1AKMT2* | ENSG00000203791.15 | 4.13 | 4.70 | 0.57 | 1.49 | 0.000154855 | 0.001349253 |
| 55088 | *CCDC186* | ENSG00000165813.20 | 5.45 | 6.02 | 0.57 | 1.49 | 7.39912E-05 | 0.000788731 |
| 4751 | *NEK2* | ENSG00000117650.13 | 6.05 | 6.62 | 0.57 | 1.48 | 3.43207E-06 | 8.30429E-05 |
| 4602 | *MYB* | ENSG00000118513.20 | 4.38 | 4.94 | 0.57 | 1.48 | 0.000104211 | 0.001013674 |
| 137964 | *GPAT4* | ENSG00000158669.11 | 6.95 | 7.52 | 0.57 | 1.48 | 9.17949E-05 | 0.000923425 |
| 280636 | *SELENOH* | ENSG00000211450.11 | 4.50 | 5.07 | 0.57 | 1.48 | 6.59866E-05 | 0.000725562 |
| 55841 | *WWC3* | ENSG00000047644.20 | 7.08 | 7.65 | 0.57 | 1.48 | 0.000228169 | 0.001819651 |
| 54947 | *LPCAT2* | ENSG00000087253.13 | 6.01 | 6.58 | 0.57 | 1.48 | 2.18817E-05 | 0.000322299 |
| 6009 | *RHEB* | ENSG00000106615.10 | 7.93 | 8.50 | 0.57 | 1.48 | 1.97378E-07 | 1.15721E-05 |
| 2 | *A2M* | ENSG00000175899.15 | 9.31 | 9.88 | 0.57 | 1.48 | 2.47525E-06 | 6.58706E-05 |
| 10263 | *CDK2AP2* | ENSG00000167797.8 | 5.13 | 5.70 | 0.57 | 1.48 | 8.85894E-05 | 0.000897798 |
| 387103 | *CENPW* | ENSG00000203760.9 | 4.49 | 5.05 | 0.57 | 1.48 | 0.000152528 | 0.001336255 |
| 57182 | *ANKRD50* | ENSG00000151458.12 | 5.42 | 5.98 | 0.57 | 1.48 | 2.25062E-05 | 0.000329151 |
| 146909 | *KIF18B* | ENSG00000186185.14 | 6.07 | 6.64 | 0.57 | 1.48 | 8.11548E-05 | 0.00084587 |
| 51101 | *ZC2HC1A* | ENSG00000104427.12 | 4.17 | 4.73 | 0.57 | 1.48 | 0.001651149 | 0.008353426 |
| 10567 | *RABAC1* | ENSG00000105404.11 | 5.56 | 6.12 | 0.57 | 1.48 | 2.60038E-05 | 0.000364538 |
| 127262 | *TPRG1L* | ENSG00000158109.15 | 4.67 | 5.24 | 0.56 | 1.48 | 3.4163E-05 | 0.000445135 |
| 6734 | *SRPRA* | ENSG00000182934.12 | 7.84 | 8.40 | 0.56 | 1.48 | 3.71203E-07 | 1.76869E-05 |
| 79845 | *RNF122* | ENSG00000133874.2 | 4.00 | 4.56 | 0.56 | 1.48 | 0.000506259 | 0.003372587 |
| 23052 | *ENDOD1* | ENSG00000149218.5 | 5.67 | 6.24 | 0.56 | 1.48 | 4.62926E-05 | 0.000556662 |
| 10112 | *KIF20A* | ENSG00000112984.12 | 6.99 | 7.55 | 0.56 | 1.47 | 1.09931E-06 | 3.73292E-05 |
| 23576 | *DDAH1* | ENSG00000153904.21 | 6.77 | 7.33 | 0.56 | 1.47 | 1.26909E-06 | 4.09384E-05 |
| 27230 | *SERP1* | ENSG00000120742.11 | 8.04 | 8.60 | 0.56 | 1.47 | 6.51365E-07 | 2.57951E-05 |
| 54819 | *ZCCHC10* | ENSG00000155329.12 | 3.83 | 4.38 | 0.56 | 1.47 | 0.0011567 | 0.006382689 |
| 148266 | *ZNF569* | ENSG00000196437.12 | 3.71 | 4.27 | 0.56 | 1.47 | 0.001664091 | 0.008403885 |
| 4363 | *ABCC1* | ENSG00000103222.20 | 6.17 | 6.73 | 0.56 | 1.47 | 4.91524E-06 | 0.00010833 |
| 144717 | *PHETA1* | ENSG00000198324.14 | 3.81 | 4.37 | 0.56 | 1.47 | 0.00010309 | 0.001006037 |
| 25957 | *PNISR* | ENSG00000132424.17 | 5.59 | 6.15 | 0.56 | 1.47 | 1.91608E-05 | 0.00029049 |
| 8703 | *B4GALT3* | ENSG00000158850.15 | 5.39 | 5.95 | 0.56 | 1.47 | 0.000422452 | 0.002944997 |
| 160897 | *GPR180* | ENSG00000152749.8 | 5.64 | 6.20 | 0.56 | 1.47 | 2.39933E-05 | 0.000344279 |
| 80223 | *RAB11FIP1* | ENSG00000156675.16 | 6.24 | 6.79 | 0.56 | 1.47 | 4.50474E-05 | 0.00054552 |
| 5876 | *RABGGTB* | ENSG00000137955.16 | 5.26 | 5.81 | 0.56 | 1.47 | 0.000208587 | 0.001692258 |
| 100506144 | *TMEM35B* | ENSG00000243749.1 | 3.24 | 3.80 | 0.56 | 1.47 | 0.000120522 | 0.001117996 |
| 54813 | *KLHL28* | ENSG00000179454.14 | 4.96 | 5.51 | 0.56 | 1.47 | 0.000143324 | 0.001276647 |
| 1997 | *ELF1* | ENSG00000120690.16 | 5.37 | 5.92 | 0.56 | 1.47 | 4.87188E-06 | 0.000107691 |
| 27089 | *UQCRQ* | ENSG00000164405.11 | 6.73 | 7.29 | 0.56 | 1.47 | 3.45281E-05 | 0.000448686 |
| 5214 | *PFKP* | ENSG00000067057.18 | 6.17 | 6.73 | 0.56 | 1.47 | 4.03277E-05 | 0.000505169 |
| 22911 | *WDR47* | ENSG00000085433.17 | 4.50 | 5.06 | 0.56 | 1.47 | 0.000314294 | 0.00233642 |
| 23731 | *TMEM245* | ENSG00000106771.13 | 9.19 | 9.75 | 0.56 | 1.47 | 6.17275E-07 | 2.49051E-05 |
| 7296 | *TXNRD1* | ENSG00000198431.17 | 8.82 | 9.37 | 0.55 | 1.47 | 3.19427E-07 | 1.60095E-05 |
| 23413 | *NCS1* | ENSG00000107130.10 | 6.65 | 7.20 | 0.55 | 1.47 | 5.18956E-05 | 0.000604737 |
| 29887 | *SNX10* | ENSG00000086300.16 | 6.58 | 7.13 | 0.55 | 1.47 | 1.27986E-06 | 4.11109E-05 |
| 2037 | *EPB41L2* | ENSG00000079819.19 | 7.18 | 7.73 | 0.55 | 1.47 | 3.42896E-06 | 8.30429E-05 |
| 22872 | *SEC31A* | ENSG00000138674.17 | 7.41 | 7.97 | 0.55 | 1.47 | 3.41599E-05 | 0.000445135 |
| 51109 | *RDH11* | ENSG00000072042.13 | 6.80 | 7.35 | 0.55 | 1.46 | 1.39939E-06 | 4.37392E-05 |
| 9439 | *MED23* | ENSG00000112282.18 | 6.32 | 6.87 | 0.55 | 1.46 | 7.97414E-06 | 0.00015463 |
| 56994 | *CHPT1* | ENSG00000111666.11 | 5.26 | 5.81 | 0.55 | 1.46 | 3.74936E-06 | 8.82039E-05 |
| 60313 | *GPBP1L1* | ENSG00000159592.11 | 6.41 | 6.96 | 0.55 | 1.46 | 4.75333E-05 | 0.000568468 |
| 8905 | *AP1S2* | ENSG00000182287.15 | 4.49 | 5.04 | 0.55 | 1.46 | 0.000706728 | 0.004343688 |
| 57552 | *NCEH1* | ENSG00000144959.11 | 6.09 | 6.64 | 0.55 | 1.46 | 0.000237091 | 0.001879819 |
| 1649 | *DDIT3* | ENSG00000175197.13 | 3.82 | 4.37 | 0.55 | 1.46 | 0.001560248 | 0.008018927 |
| 80854 | *SETD7* | ENSG00000145391.14 | 7.46 | 8.01 | 0.55 | 1.46 | 1.89039E-06 | 5.36349E-05 |
| 65078 | *RTN4R* | ENSG00000040608.14 | 3.87 | 4.42 | 0.55 | 1.46 | 0.001105159 | 0.006175475 |
| 8349 | *H2BC21* | ENSG00000184678.11 | 5.96 | 6.51 | 0.55 | 1.46 | 4.08147E-06 | 9.46813E-05 |
| 3159 | *HMGA1* | ENSG00000137309.20 | 3.38 | 3.93 | 0.55 | 1.46 | 0.000785802 | 0.004725204 |
| 134111 | *UBE2QL1* | ENSG00000215218.4 | 3.06 | 3.60 | 0.55 | 1.46 | 0.00098815 | 0.00563638 |
| 5190 | *PEX6* | ENSG00000124587.14 | 4.83 | 5.38 | 0.55 | 1.46 | 7.35105E-05 | 0.000785281 |

| | | | | | | | | |
|---|---|---|---|---|---|---|---|---|
| 84897 | *TBRG1* | ENSG00000154144.13 | 5.14 | 5.68 | 0.55 | 1.46 | 0.000163747 | 0.001404388 |
| 9534 | *ZNF254* | ENSG00000213096.11 | 3.36 | 3.91 | 0.55 | 1.46 | 0.001994816 | 0.00969252 |
| 9824 | *ARHGAP11A* | ENSG00000198826.11 | 7.61 | 8.16 | 0.55 | 1.46 | 5.57024E-06 | 0.000119094 |
| 50808 | *AK3* | ENSG00000147853.17 | 5.75 | 6.30 | 0.55 | 1.46 | 3.72233E-05 | 0.00047678 |
| 221178 | *SPATA13* | ENSG00000182957.16 | 5.61 | 6.15 | 0.55 | 1.46 | 0.001917502 | 0.009389815 |
| 23232 | *TBC1D12* | ENSG00000108239.9 | 4.02 | 4.57 | 0.55 | 1.46 | 0.000256822 | 0.001991078 |
| 51399 | *TRAPPC4* | ENSG00000196655.12 | 5.25 | 5.80 | 0.54 | 1.46 | 0.000293111 | 0.002216272 |
| 8312 | *AXIN1* | ENSG00000103126.15 | 4.93 | 5.47 | 0.54 | 1.46 | 0.000119223 | 0.001109368 |
| 890 | *CCNA2* | ENSG00000145386.11 | 7.09 | 7.63 | 0.54 | 1.46 | 1.19016E-05 | 0.00020868 |
| 55276 | *PGM2* | ENSG00000169299.14 | 5.53 | 6.07 | 0.54 | 1.46 | 0.000350886 | 0.002540498 |
| 1317 | *SLC31A1* | ENSG00000136868.11 | 7.77 | 8.32 | 0.54 | 1.46 | 2.0005E-06 | 5.63329E-05 |
| 8543 | *LMO4* | ENSG00000143013.13 | 4.12 | 4.67 | 0.54 | 1.46 | 0.000415901 | 0.002906082 |
| 5794 | *PTPRH* | ENSG00000080031.10 | 6.54 | 7.08 | 0.54 | 1.46 | 8.22508E-05 | 0.000853736 |
| 11065 | *UBE2C* | ENSG00000175063.17 | 6.83 | 7.37 | 0.54 | 1.46 | 3.63927E-06 | 8.68392E-05 |
| 25924 | *MYRIP* | ENSG00000170011.14 | 5.59 | 6.13 | 0.54 | 1.46 | 7.5994E-06 | 0.000149097 |
| 284403 | *WDR62* | ENSG00000075702.19 | 6.28 | 6.82 | 0.54 | 1.45 | 2.55616E-05 | 0.000360577 |
| 983 | *CDK1* | ENSG00000170312.17 | 7.87 | 8.41 | 0.54 | 1.45 | 3.24161E-07 | 1.61104E-05 |
| 5142 | *PDE4B* | ENSG00000184588.18 | 4.34 | 4.88 | 0.54 | 1.45 | 2.16384E-05 | 0.000319656 |
| 7532 | *YWHAG* | ENSG00000170027.7 | 9.04 | 9.58 | 0.54 | 1.45 | 2.0061E-07 | 1.17028E-05 |
| 85416 | *ZIC5* | ENSG00000139800.9 | 3.10 | 3.64 | 0.54 | 1.45 | 0.00041268 | 0.002887608 |
| 55857 | *KIZ* | ENSG00000088970.16 | 5.10 | 5.64 | 0.54 | 1.45 | 2.66801E-05 | 0.000371013 |
| 26353 | *HSPB8* | ENSG00000152137.8 | 4.35 | 4.89 | 0.54 | 1.45 | 0.001462388 | 0.007645067 |
| 57491 | *AHRR* | ENSG00000063438.20 | 5.00 | 5.53 | 0.54 | 1.45 | 0.000249066 | 0.001946998 |
| 55113 | *XKR8* | ENSG00000158156.9 | 3.66 | 4.20 | 0.54 | 1.45 | 0.000528396 | 0.003486176 |
| 967 | *CD63* | ENSG00000135404.12 | 9.42 | 9.95 | 0.54 | 1.45 | 1.84732E-06 | 5.2913E-05 |
| 55275 | *VPS53* | ENSG00000141252.21 | 5.36 | 5.90 | 0.54 | 1.45 | 0.00017804 | 0.001497004 |
| 891 | *CCNB1* | ENSG00000134057.15 | 8.07 | 8.61 | 0.54 | 1.45 | 4.44239E-07 | 2.00228E-05 |
| 3422 | *IDI1* | ENSG00000067064.12 | 7.06 | 7.59 | 0.54 | 1.45 | 9.76458E-06 | 0.000180713 |
| 167691 | *LCA5* | ENSG00000135338.14 | 4.13 | 4.67 | 0.54 | 1.45 | 0.000238281 | 0.001883285 |
| 25948 | *KBTBD2* | ENSG00000170852.12 | 6.03 | 6.57 | 0.53 | 1.45 | 4.86785E-06 | 0.000107691 |
| 3148 | *HMGB2* | ENSG00000164104.12 | 6.94 | 7.48 | 0.53 | 1.45 | 1.27106E-06 | 4.09384E-05 |
| 6429 | *SRSF4* | ENSG00000116350.18 | 6.84 | 7.37 | 0.53 | 1.45 | 1.0709E-05 | 0.00019412 |
| 5998 | *RGS3* | ENSG00000138835.22 | 7.04 | 7.57 | 0.53 | 1.45 | 2.57815E-05 | 0.000363337 |
| 1831 | *TSC22D3* | ENSG00000157514.18 | 6.91 | 7.44 | 0.53 | 1.45 | 6.508E-07 | 2.57951E-05 |
| 28970 | *C11orf54* | ENSG00000182919.15 | 4.95 | 5.48 | 0.53 | 1.45 | 0.000342257 | 0.002490032 |
| 995 | *CDC25C* | ENSG00000158402.21 | 4.15 | 4.68 | 0.53 | 1.45 | 0.000483051 | 0.003252632 |
| 1508 | *CTSB* | ENSG00000164733.22 | 8.16 | 8.69 | 0.53 | 1.45 | 1.52642E-06 | 4.58201E-05 |
| 529 | *ATP6V1E1* | ENSG00000131100.13 | 7.71 | 8.24 | 0.53 | 1.45 | 7.68734E-07 | 2.88448E-05 |
| 11240 | *PADI2* | ENSG00000117115.13 | 3.16 | 3.69 | 0.53 | 1.44 | 0.000504885 | 0.003366425 |
| 56180 | *MOSPD1* | ENSG00000101928.13 | 4.54 | 5.07 | 0.53 | 1.44 | 0.000401341 | 0.002824063 |
| 51309 | *ARMCX1* | ENSG00000126947.13 | 5.09 | 5.62 | 0.53 | 1.44 | 5.7445E-05 | 0.000653671 |
| 10926 | *DBF4* | ENSG00000006634.8 | 5.89 | 6.42 | 0.53 | 1.44 | 0.000103034 | 0.001006037 |
| 1955 | *MEGF9* | ENSG00000106780.9 | 6.34 | 6.87 | 0.53 | 1.44 | 4.37947E-06 | 9.95376E-05 |
| 9394 | *HS6ST1* | ENSG00000136720.7 | 6.13 | 6.65 | 0.53 | 1.44 | 4.7972E-06 | 0.000106892 |
| 9266 | *CYTH2* | ENSG00000105443.16 | 6.41 | 6.94 | 0.53 | 1.44 | 4.90064E-05 | 0.000581452 |
| 85415 | *RHPN2* | ENSG00000131941.8 | 4.69 | 5.21 | 0.53 | 1.44 | 0.00102311 | 0.00579525 |
| 54675 | *CRLS1* | ENSG00000088766.12 | 5.10 | 5.62 | 0.53 | 1.44 | 0.000466806 | 0.003163111 |
| 6520 | *SLC3A2* | ENSG00000168003.18 | 6.83 | 7.36 | 0.53 | 1.44 | 1.84526E-05 | 0.000282015 |
| 653464 | *SRGAP2C* | ENSG00000171943.12 | 5.82 | 6.35 | 0.52 | 1.44 | 0.000393234 | 0.002783988 |
| 164832 | *LONRF2* | ENSG00000170500.13 | 6.65 | 7.17 | 0.52 | 1.44 | 7.15237E-06 | 0.000141562 |
| 5264 | *PHYH* | ENSG00000107537.14 | 5.29 | 5.82 | 0.52 | 1.44 | 6.10704E-06 | 0.000126428 |
| 5202 | *PFDN2* | ENSG00000143256.5 | 5.22 | 5.74 | 0.52 | 1.44 | 7.50127E-05 | 0.000796892 |
| 5255 | *PHKA1* | ENSG00000067177.15 | 6.08 | 6.60 | 0.52 | 1.44 | 2.42722E-05 | 0.000346954 |
| 1315 | *COPB1* | ENSG00000129083.13 | 6.97 | 7.49 | 0.52 | 1.44 | 4.81594E-07 | 2.13175E-05 |
| 9552 | *SPAG7* | ENSG00000091640.8 | 5.05 | 5.58 | 0.52 | 1.44 | 8.63497E-05 | 0.00088045 |
| 2101 | *ESRRA* | ENSG00000173153.17 | 4.87 | 5.39 | 0.52 | 1.44 | 0.000128538 | 0.001176354 |
| 11072 | *DUSP14* | ENSG00000276023.5 | 5.34 | 5.87 | 0.52 | 1.44 | 6.10598E-06 | 0.000126428 |
| 427 | *ASAH1* | ENSG00000104763.20 | 5.58 | 6.10 | 0.52 | 1.44 | 5.86565E-05 | 0.000663933 |
| 51272 | *BET1L* | ENSG00000177951.18 | 4.58 | 5.10 | 0.52 | 1.44 | 0.000145365 | 0.001291756 |
| 1347 | *COX7A2* | ENSG00000112695.13 | 6.55 | 7.07 | 0.52 | 1.44 | 3.68803E-05 | 0.000474732 |
| 7153 | *TOP2A* | ENSG00000131747.15 | 9.63 | 10.15 | 0.52 | 1.43 | 8.99335E-07 | 3.2062E-05 |
| 3105 | *HLA-A* | ENSG00000206503.13 | 7.30 | 7.82 | 0.52 | 1.43 | 1.52524E-06 | 4.58201E-05 |
| 5110 | *PCMT1* | ENSG00000120265.19 | 6.82 | 7.34 | 0.52 | 1.43 | 6.51232E-07 | 2.57951E-05 |
| 83857 | *TMTC1* | ENSG00000133687.16 | 8.10 | 8.62 | 0.52 | 1.43 | 5.34938E-06 | 0.000114908 |
| 388789 | *SMIM26* | ENSG00000232388.5 | 4.42 | 4.94 | 0.52 | 1.43 | 0.000541588 | 0.003556806 |
| 2770 | *GNAI1* | ENSG00000127955.17 | 6.20 | 6.72 | 0.52 | 1.43 | 1.78284E-06 | 5.16575E-05 |
| 2643 | *GCH1* | ENSG00000131979.20 | 3.69 | 4.21 | 0.52 | 1.43 | 0.001638763 | 0.008315143 |
| 10282 | *BET1* | ENSG00000105829.13 | 5.35 | 5.87 | 0.52 | 1.43 | 1.84703E-05 | 0.000282015 |
| 163859 | *SDE2* | ENSG00000143751.10 | 5.52 | 6.03 | 0.52 | 1.43 | 0.000136835 | 0.00123201 |
| 84926 | *SPRYD3* | ENSG00000167778.9 | 6.62 | 7.14 | 0.52 | 1.43 | 9.22273E-06 | 0.000173681 |
| 154791 | *FMC1* | ENSG00000164898.13 | 4.04 | 4.56 | 0.52 | 1.43 | 0.00031505 | 0.002339724 |
| 3344 | *FOXN2* | ENSG00000170802.17 | 4.48 | 5.00 | 0.52 | 1.43 | 0.001703537 | 0.008551302 |
| 5480 | *PPIC* | ENSG00000168938.6 | 4.24 | 4.76 | 0.52 | 1.43 | 0.000166028 | 0.001421511 |
| 11040 | *PIM2* | ENSG00000102096.9 | 4.28 | 4.80 | 0.52 | 1.43 | 5.04559E-05 | 0.000593025 |
| 79589 | *RNF128* | ENSG00000133135.14 | 6.72 | 7.24 | 0.52 | 1.43 | 3.28063E-06 | 8.08523E-05 |
| 219654 | *ZCCHC24* | ENSG00000165424.7 | 4.75 | 5.26 | 0.51 | 1.43 | 7.96224E-05 | 0.000834395 |
| 8531 | *YBX3* | ENSG00000060138.13 | 7.96 | 8.48 | 0.51 | 1.43 | 1.33642E-05 | 0.000227936 |
| 9069 | *CLDN12* | ENSG00000157224.16 | 6.57 | 7.09 | 0.51 | 1.43 | 3.80893E-06 | 8.90471E-05 |
| 4097 | *MAFG* | ENSG00000197063.11 | 5.76 | 6.28 | 0.51 | 1.43 | 3.37629E-06 | 8.2398E-05 |
| 7326 | *UBE2G1* | ENSG00000132388.13 | 6.52 | 7.04 | 0.51 | 1.43 | 9.35566E-05 | 0.000934881 |
| 23551 | *RASD2* | ENSG00000100302.7 | 5.79 | 6.31 | 0.51 | 1.43 | 9.76468E-06 | 0.000180713 |
| 83931 | *STK40* | ENSG00000196182.11 | 5.26 | 5.77 | 0.51 | 1.43 | 0.000272973 | 0.002091401 |
| 201475 | *RAB12* | ENSG00000206418.5 | 5.86 | 6.37 | 0.51 | 1.43 | 5.84043E-06 | 0.000122363 |
| 2115 | *ETV1* | ENSG00000006468.14 | 6.60 | 7.11 | 0.51 | 1.43 | 0.000658764 | 0.004114604 |
| 9764 | *KIAA0513* | ENSG00000135709.14 | 5.34 | 5.86 | 0.51 | 1.43 | 0.000270567 | 0.002076148 |
| 6839 | *SUV39H1* | ENSG00000101945.17 | 4.63 | 5.14 | 0.51 | 1.43 | 0.000719322 | 0.004403975 |
| 54205 | *CYCS* | ENSG00000172115.9 | 7.76 | 8.27 | 0.51 | 1.43 | 5.75594E-07 | 2.40643E-05 |

| | | | | | | | | |
|---|---|---|---|---|---|---|---|---|
| 7048 | *TGFBR2* | ENSG00000163513.19 | 5.78 | 6.29 | 0.51 | 1.43 | 2.32795E-05 | 0.00033661 |
| 22903 | *BTBD3* | ENSG00000132640.15 | 5.91 | 6.42 | 0.51 | 1.42 | 7.06009E-05 | 0.000762886 |
| 23139 | *MAST2* | ENSG00000086015.23 | 6.40 | 6.91 | 0.51 | 1.42 | 0.000243388 | 0.001910678 |
| 80314 | *EPC1* | ENSG00000120616.16 | 4.83 | 5.34 | 0.51 | 1.42 | 0.000204595 | 0.001668937 |
| 113402 | *SFT2D1* | ENSG00000198818.10 | 4.92 | 5.43 | 0.51 | 1.42 | 1.82704E-05 | 0.000280103 |
| 6487 | *ST3GAL3* | ENSG00000126091.21 | 4.71 | 5.22 | 0.51 | 1.42 | 0.000107568 | 0.001036918 |
| 91663 | *MYADM* | ENSG00000179820.16 | 6.38 | 6.89 | 0.51 | 1.42 | 2.31796E-05 | 0.000335489 |
| 115294 | *PCMTD1* | ENSG00000168300.14 | 5.89 | 6.39 | 0.51 | 1.42 | 2.76199E-06 | 7.10565E-05 |
| 83941 | *TM2D1* | ENSG00000162604.12 | 4.20 | 4.71 | 0.51 | 1.42 | 0.000469766 | 0.003177429 |
| 1040 | *CDS1* | ENSG00000163624.6 | 4.61 | 5.11 | 0.51 | 1.42 | 0.000101603 | 0.000997359 |
| 56271 | *BEX4* | ENSG00000102409.10 | 3.76 | 4.27 | 0.51 | 1.42 | 0.000381355 | 0.002711159 |
| 64764 | *CREB3L2* | ENSG00000182158.16 | 7.23 | 7.74 | 0.51 | 1.42 | 3.31025E-05 | 0.000434297 |
| 9513 | *FXR2* | ENSG00000129245.12 | 6.49 | 6.99 | 0.51 | 1.42 | 2.05212E-06 | 5.71433E-05 |
| 6617 | *SNAPC1* | ENSG00000023608.5 | 4.24 | 4.75 | 0.51 | 1.42 | 0.000277159 | 0.002116988 |
| 26175 | *TMEM251* | ENSG00000153485.6 | 4.14 | 4.65 | 0.51 | 1.42 | 0.000178597 | 0.001499197 |
| 9781 | *RNF144A* | ENSG00000151692.15 | 6.11 | 6.61 | 0.51 | 1.42 | 8.05928E-06 | 0.000155879 |
| 23484 | *LEPROTL1* | ENSG00000104660.19 | 5.14 | 5.64 | 0.50 | 1.42 | 1.65631E-05 | 0.000263343 |
| 10802 | *SEC24A* | ENSG00000113615.13 | 6.59 | 7.09 | 0.50 | 1.42 | 1.10232E-05 | 0.00019814 |
| 991 | *CDC20* | ENSG00000117399.14 | 6.87 | 7.37 | 0.50 | 1.42 | 1.46757E-05 | 0.000241521 |
| 92259 | *MRPS36* | ENSG00000134056.12 | 5.25 | 5.76 | 0.50 | 1.42 | 3.53131E-05 | 0.000457303 |
| 7743 | *ZNF189* | ENSG00000136870.11 | 7.21 | 7.72 | 0.50 | 1.42 | 1.52137E-06 | 4.58201E-05 |
| 4957 | *ODF2* | ENSG00000136811.17 | 6.29 | 6.80 | 0.50 | 1.42 | 1.74348E-05 | 0.00027117 |
| 79960 | *JADE1* | ENSG00000077684.16 | 6.30 | 6.81 | 0.50 | 1.42 | 1.6647E-06 | 4.87998E-05 |
| 6560 | *SLC12A4* | ENSG00000124067.17 | 5.03 | 5.53 | 0.50 | 1.42 | 0.001363565 | 0.007234271 |
| 255743 | *NPNT* | ENSG00000168743.13 | 6.08 | 6.58 | 0.50 | 1.42 | 1.56604E-05 | 0.000253886 |
| 84886 | *C1orf198* | ENSG00000119280.17 | 5.33 | 5.83 | 0.50 | 1.42 | 1.79593E-05 | 0.00027618 |
| 64951 | *MRPS24* | ENSG00000062582.14 | 6.42 | 6.92 | 0.50 | 1.42 | 3.28751E-06 | 8.08664E-05 |
| 439921 | *MXRA7* | ENSG00000182534.14 | 6.06 | 6.56 | 0.50 | 1.41 | 3.20631E-05 | 0.000424524 |
| 273 | *AMPH* | ENSG00000078053.17 | 5.20 | 5.70 | 0.50 | 1.41 | 4.08717E-05 | 0.00050932 |
| 5347 | *PLK1* | ENSG00000166851.15 | 6.75 | 7.24 | 0.50 | 1.41 | 6.5436E-06 | 0.00013326 |
| 137695 | *TMEM68* | ENSG00000167904.16 | 4.61 | 5.11 | 0.50 | 1.41 | 0.000729255 | 0.004449347 |
| 121536 | *AEBP2* | ENSG00000139154.16 | 7.86 | 8.36 | 0.50 | 1.41 | 2.89439E-06 | 7.36186E-05 |
| 4792 | *NFKBIA* | ENSG00000100906.11 | 5.50 | 6.00 | 0.50 | 1.41 | 4.22234E-05 | 0.000518177 |
| 55033 | *FKBP14* | ENSG00000106080.11 | 5.79 | 6.28 | 0.50 | 1.41 | 0.000321175 | 0.0023723 |
| 79666 | *PLEKHF2* | ENSG00000175895.4 | 4.29 | 4.79 | 0.50 | 1.41 | 0.000303862 | 0.002281475 |
| 55186 | *SLC25A36* | ENSG00000114120.14 | 7.05 | 7.55 | 0.50 | 1.41 | 6.37202E-06 | 0.000130699 |
| 133746 | *JMY* | ENSG00000152409.9 | 6.32 | 6.82 | 0.50 | 1.41 | 1.6801E-05 | 0.000265216 |
| 9353 | *SLIT2* | ENSG00000145147.20 | 5.10 | 5.60 | 0.50 | 1.41 | 0.000161206 | 0.001389745 |
| 55334 | *SLC39A9* | ENSG00000029364.12 | 6.41 | 6.91 | 0.50 | 1.41 | 6.94829E-06 | 0.000139421 |
| 10389 | *SCML2* | ENSG00000102098.19 | 5.30 | 5.79 | 0.50 | 1.41 | 5.45198E-05 | 0.000627521 |
| 8904 | *CPNE1* | ENSG00000214078.13 | 6.39 | 6.88 | 0.49 | 1.41 | 0.000151072 | 0.001328319 |
| 80267 | *EDEM3* | ENSG00000116406.20 | 6.65 | 7.15 | 0.49 | 1.41 | 1.46182E-05 | 0.000241104 |
| 54532 | *USP53* | ENSG00000145390.12 | 4.37 | 4.86 | 0.49 | 1.41 | 0.001411042 | 0.007430992 |
| 10519 | *CIB1* | ENSG00000185043.12 | 5.77 | 6.26 | 0.49 | 1.41 | 7.35822E-05 | 0.000785487 |
| 3638 | *INSIG1* | ENSG00000186480.13 | 7.41 | 7.90 | 0.49 | 1.41 | 1.20475E-05 | 0.000210502 |
| 23306 | *NEMP1* | ENSG00000166881.10 | 7.39 | 7.88 | 0.49 | 1.41 | 1.24696E-06 | 4.05979E-05 |
| 30000 | *TNPO2* | ENSG00000105576.16 | 6.87 | 7.37 | 0.49 | 1.41 | 2.48076E-05 | 0.000352926 |
| 2273 | *FHL1* | ENSG00000022267.19 | 5.23 | 5.73 | 0.49 | 1.41 | 0.00061429 | 0.003926728 |
| 23173 | *METAP1* | ENSG00000164024.12 | 5.70 | 6.19 | 0.49 | 1.40 | 2.03548E-05 | 0.000306117 |
| 53616 | *ADAM22* | ENSG00000008277.15 | 6.07 | 6.56 | 0.49 | 1.40 | 0.000152484 | 0.001336255 |
| 2889 | *RAPGEF1* | ENSG00000107263.19 | 6.98 | 7.47 | 0.49 | 1.40 | 2.02122E-05 | 0.000304583 |
| 5819 | *NECTIN2* | ENSG00000130202.10 | 6.48 | 6.97 | 0.49 | 1.40 | 3.06473E-05 | 0.000409969 |
| 128077 | *LIX1L* | ENSG00000271601.4 | 6.88 | 7.37 | 0.49 | 1.40 | 1.05001E-06 | 3.60476E-05 |
| 54853 | *WDR55* | ENSG00000120314.19 | 4.52 | 5.00 | 0.49 | 1.40 | 0.000204701 | 0.001668937 |
| 51478 | *HSD17B7* | ENSG00000132196.16 | 4.64 | 5.12 | 0.49 | 1.40 | 0.00020342 | 0.001663831 |
| 2717 | *GLA* | ENSG00000102393.14 | 5.38 | 5.87 | 0.49 | 1.40 | 1.77478E-05 | 0.000274332 |
| 56605 | *ERO1B* | ENSG00000086619.15 | 5.48 | 5.97 | 0.49 | 1.40 | 9.52874E-05 | 0.000948387 |
| 84892 | *POMGNT2* | ENSG00000144647.7 | 6.52 | 7.01 | 0.49 | 1.40 | 2.51399E-06 | 6.65475E-05 |
| 53335 | *BCL11A* | ENSG00000119866.22 | 4.44 | 4.92 | 0.49 | 1.40 | 0.001004826 | 0.005710498 |
| 8565 | *YARS1* | ENSG00000134684.12 | 6.63 | 7.12 | 0.49 | 1.40 | 1.0229E-05 | 0.000187786 |
| 80346 | *REEP4* | ENSG00000168476.12 | 4.26 | 4.74 | 0.48 | 1.40 | 0.000742828 | 0.004519297 |
| 57553 | *MICAL3* | ENSG00000243156.9 | 6.46 | 6.95 | 0.48 | 1.40 | 0.000861285 | 0.005039779 |
| 1654 | *DDX3X* | ENSG00000215301.11 | 8.95 | 9.43 | 0.48 | 1.40 | 7.4912E-06 | 0.000147167 |
| 27238 | *GPKOW* | ENSG00000068394.11 | 4.96 | 5.44 | 0.48 | 1.40 | 0.000491235 | 0.003294435 |
| 23348 | *DOCK9* | ENSG00000088387.21 | 6.10 | 6.58 | 0.48 | 1.40 | 0.000336384 | 0.002450864 |
| 204851 | *HIPK1* | ENSG00000163349.22 | 7.27 | 7.75 | 0.48 | 1.40 | 9.0354E-06 | 0.000170796 |
| 84182 | *MINDY4* | ENSG00000106125.14 | 4.18 | 4.66 | 0.48 | 1.40 | 0.000359962 | 0.002591209 |
| 93058 | *COQ10A* | ENSG00000135469.14 | 4.49 | 4.97 | 0.48 | 1.40 | 0.000433108 | 0.002999777 |
| 6814 | *STXBP3* | ENSG00000116266.11 | 5.63 | 6.12 | 0.48 | 1.40 | 6.79088E-05 | 0.000742347 |
| 201595 | *STT3B* | ENSG00000163527.10 | 8.10 | 8.59 | 0.48 | 1.40 | 9.25668E-06 | 0.000173938 |
| 6770 | *STAR* | ENSG00000147465.12 | 10.05 | 10.54 | 0.48 | 1.40 | 1.61218E-06 | 4.75972E-05 |
| 4286 | *MITF* | ENSG00000187098.18 | 6.11 | 6.59 | 0.48 | 1.40 | 2.10707E-05 | 0.000313705 |
| 30001 | *ERO1A* | ENSG00000197930.13 | 6.48 | 6.96 | 0.48 | 1.40 | 5.14784E-05 | 0.000601276 |
| 6868 | *ADAM17* | ENSG00000151694.14 | 6.20 | 6.68 | 0.48 | 1.39 | 5.30355E-05 | 0.00061563 |
| 5261 | *PHKG2* | ENSG00000156873.16 | 4.36 | 4.84 | 0.48 | 1.39 | 0.000189952 | 0.001573392 |
| 55252 | *ASXL2* | ENSG00000143970.18 | 7.09 | 7.57 | 0.48 | 1.39 | 6.84801E-06 | 0.000137962 |
| 54867 | *TMEM214* | ENSG00000119777.20 | 6.70 | 7.18 | 0.48 | 1.39 | 2.22405E-05 | 0.000326303 |
| 1459 | *CSNK2A2* | ENSG00000070770.10 | 6.92 | 7.40 | 0.48 | 1.39 | 4.04318E-05 | 0.000505701 |
| 26258 | *BLOC1S6* | ENSG00000104164.12 | 7.18 | 7.66 | 0.48 | 1.39 | 4.96293E-05 | 0.000585602 |
| 55352 | *COPRS* | ENSG00000172301.11 | 5.64 | 6.11 | 0.48 | 1.39 | 9.75381E-06 | 0.000180713 |
| 23158 | *TBC1D9* | ENSG00000109436.8 | 6.02 | 6.50 | 0.48 | 1.39 | 7.47689E-05 | 0.000795891 |
| 100506844 | *GIHCG* | ENSG00000257698.3 | 4.00 | 4.48 | 0.48 | 1.39 | 0.001747368 | 0.008730441 |
| 9777 | *TM9SF4* | ENSG00000101337.16 | 7.26 | 7.73 | 0.48 | 1.39 | 4.19528E-06 | 9.65751E-05 |
| 253430 | *IPMK* | ENSG00000151151.6 | 4.59 | 5.06 | 0.48 | 1.39 | 0.001033751 | 0.005848311 |
| 84790 | *TUBA1C* | ENSG00000167553.17 | 8.85 | 9.33 | 0.48 | 1.39 | 3.41713E-06 | 8.30429E-05 |
| 6645 | *SNTB2* | ENSG00000168807.17 | 5.52 | 5.99 | 0.47 | 1.39 | 4.19317E-05 | 0.000516286 |
| 4215 | *MAP3K3* | ENSG00000198909.8 | 3.84 | 4.32 | 0.47 | 1.39 | 0.000650468 | 0.004083178 |

| | | | | | | | | |
|---|---|---|---|---|---|---|---|---|
| 10015 | *PDCD6IP* | ENSG00000170248.15 | 7.39 | 7.86 | 0.47 | 1.39 | 0.000225514 | 0.001804249 |
| 961 | *CD47* | ENSG00000196776.17 | 7.20 | 7.67 | 0.47 | 1.39 | 2.39072E-06 | 6.43052E-05 |
| 10954 | *PDIA5* | ENSG00000065485.20 | 5.00 | 5.48 | 0.47 | 1.39 | 0.001200386 | 0.006568206 |
| 6609 | *SMPD1* | ENSG00000166311.10 | 5.26 | 5.74 | 0.47 | 1.39 | 0.000462795 | 0.003146542 |
| 55905 | *RNF114* | ENSG00000124226.11 | 5.75 | 6.22 | 0.47 | 1.39 | 0.001863229 | 0.009177949 |
| 2512 | *FTL* | ENSG00000087086.15 | 9.16 | 9.64 | 0.47 | 1.39 | 1.39174E-05 | 0.000234177 |
| 9212 | *AURKB* | ENSG00000178999.13 | 4.93 | 5.41 | 0.47 | 1.39 | 0.000128081 | 0.001172887 |
| 2787 | *GNG5* | ENSG00000174021.12 | 5.82 | 6.29 | 0.47 | 1.39 | 6.74942E-06 | 0.00013708 |
| 10681 | *GNB5* | ENSG00000069966.19 | 5.88 | 6.35 | 0.47 | 1.39 | 2.18036E-05 | 0.000321464 |
| 9751 | *SNPH* | ENSG00000101298.15 | 4.46 | 4.94 | 0.47 | 1.39 | 0.000401749 | 0.00282561 |
| 63915 | *BLOC1S5* | ENSG00000188428.20 | 5.11 | 5.58 | 0.47 | 1.39 | 8.7326E-05 | 0.000889196 |
| 211 | *ALAS1* | ENSG00000023330.15 | 6.85 | 7.32 | 0.47 | 1.39 | 4.11847E-06 | 9.53036E-05 |
| 26064 | *RAI14* | ENSG00000039560.14 | 7.11 | 7.59 | 0.47 | 1.39 | 1.72362E-05 | 0.000269478 |
| 64946 | *CENPH* | ENSG00000153044.10 | 6.03 | 6.50 | 0.47 | 1.39 | 3.93959E-05 | 0.000497303 |
| 51594 | *NBAS* | ENSG00000151779.13 | 5.56 | 6.03 | 0.47 | 1.39 | 0.000412161 | 0.00288532 |
| 80146 | *UXS1* | ENSG00000115652.15 | 5.62 | 6.09 | 0.47 | 1.39 | 0.000101493 | 0.000996925 |
| 84268 | *RPAIN* | ENSG00000129197.15 | 4.51 | 4.98 | 0.47 | 1.39 | 0.000147214 | 0.001302788 |
| 54820 | *NDE1* | ENSG00000072864.16 | 5.57 | 6.04 | 0.47 | 1.39 | 0.000536375 | 0.003527803 |
| 11345 | *GABARAPL2* | ENSG00000034713.8 | 6.44 | 6.91 | 0.47 | 1.39 | 5.03136E-06 | 0.000109761 |
| 84662 | *GLIS2* | ENSG00000126603.9 | 4.03 | 4.50 | 0.47 | 1.39 | 0.001875178 | 0.009227719 |
| 10043 | *TOM1* | ENSG00000100284.22 | 5.61 | 6.08 | 0.47 | 1.39 | 0.001748462 | 0.008733001 |
| 9875 | *URB1* | ENSG00000142207.7 | 6.30 | 6.77 | 0.47 | 1.39 | 2.53574E-05 | 0.000359046 |
| 23556 | *PIGN* | ENSG00000197563.11 | 5.97 | 6.44 | 0.47 | 1.38 | 5.53179E-05 | 0.000635246 |
| 6515 | *SLC2A3* | ENSG00000059804.16 | 5.19 | 5.66 | 0.47 | 1.38 | 8.14002E-05 | 0.000846402 |
| 143384 | *CACUL1* | ENSG00000151893.15 | 6.71 | 7.18 | 0.47 | 1.38 | 0.000206636 | 0.001681891 |
| 4209 | *MEF2D* | ENSG00000116604.19 | 6.28 | 6.74 | 0.47 | 1.38 | 1.84251E-05 | 0.000281899 |
| 9261 | *MAPKAPK2* | ENSG00000162889.11 | 5.60 | 6.07 | 0.47 | 1.38 | 0.000639815 | 0.004033175 |
| 28998 | *MRPL13* | ENSG00000172172.8 | 5.58 | 6.04 | 0.47 | 1.38 | 7.85855E-05 | 0.000827131 |
| 84144 | *SYDE2* | ENSG00000097096.9 | 3.69 | 4.16 | 0.47 | 1.38 | 0.000852058 | 0.005009827 |
| 57462 | *MYORG* | ENSG00000164976.9 | 5.99 | 6.45 | 0.47 | 1.38 | 0.000161721 | 0.001390984 |
| 2879 | *GPX4* | ENSG00000167468.19 | 6.86 | 7.32 | 0.47 | 1.38 | 4.57946E-06 | 0.000103203 |
| 4179 | *CD46* | ENSG00000117335.20 | 7.44 | 7.90 | 0.47 | 1.38 | 1.11357E-05 | 0.000199684 |
| 5573 | *PRKAR1A* | ENSG00000108946.16 | 9.50 | 9.97 | 0.47 | 1.38 | 2.90316E-06 | 7.36186E-05 |
| 51430 | *SUCO* | ENSG00000094975.14 | 7.58 | 8.04 | 0.47 | 1.38 | 5.23549E-06 | 0.000112902 |
| 4591 | *TRIM37* | ENSG00000108395.14 | 6.90 | 7.37 | 0.46 | 1.38 | 2.58128E-05 | 0.000363438 |
| 3688 | *ITGB1* | ENSG00000150093.20 | 9.08 | 9.54 | 0.46 | 1.38 | 9.05512E-07 | 3.21962E-05 |
| 55761 | *TTC17* | ENSG00000052841.15 | 6.61 | 7.08 | 0.46 | 1.38 | 1.4252E-05 | 0.000237677 |
| 8615 | *USO1* | ENSG00000138768.15 | 6.90 | 7.36 | 0.46 | 1.38 | 5.81363E-06 | 0.000122074 |
| 80381 | *CD276* | ENSG00000103855.18 | 6.69 | 7.16 | 0.46 | 1.38 | 0.000268097 | 0.002064581 |
| 23471 | *TRAM1* | ENSG00000067167.8 | 8.63 | 9.10 | 0.46 | 1.38 | 2.76481E-06 | 7.10565E-05 |
| 64771 | *ILRUN* | ENSG00000196821.10 | 7.29 | 7.75 | 0.46 | 1.38 | 0.000216755 | 0.001749075 |
| 8516 | *ITGA8* | ENSG00000077943.8 | 4.41 | 4.87 | 0.46 | 1.38 | 0.000641896 | 0.004041456 |
| 6048 | *RNF5* | ENSG00000204308.8 | 6.55 | 7.02 | 0.46 | 1.38 | 0.001608183 | 0.008196001 |
| 55143 | *CDCA8* | ENSG00000134690.11 | 6.38 | 6.85 | 0.46 | 1.38 | 4.2449E-06 | 9.75677E-05 |
| 9643 | *MORF4L2* | ENSG00000123562.18 | 8.55 | 9.01 | 0.46 | 1.38 | 1.11099E-05 | 0.00019946 |
| 10550 | *ARL6IP5* | ENSG00000144746.7 | 7.69 | 8.15 | 0.46 | 1.38 | 1.70887E-06 | 4.98026E-05 |
| 9846 | *GAB2* | ENSG00000033327.13 | 5.36 | 5.82 | 0.46 | 1.38 | 1.76684E-05 | 0.000273386 |
| 23463 | *ICMT* | ENSG00000116237.16 | 7.18 | 7.64 | 0.46 | 1.38 | 2.5616E-06 | 6.75892E-05 |
| 9470 | *EIF4E2* | ENSG00000135930.15 | 5.48 | 5.94 | 0.46 | 1.38 | 0.000162677 | 0.001396809 |
| 220965 | *FAM13C* | ENSG00000148541.13 | 4.05 | 4.51 | 0.46 | 1.38 | 0.000124145 | 0.001143829 |
| 23170 | *TTLL12* | ENSG00000100304.13 | 5.48 | 5.94 | 0.46 | 1.38 | 0.00014933 | 0.001317636 |
| 7184 | *HSP90B1* | ENSG00000166598.16 | 10.29 | 10.75 | 0.46 | 1.38 | 5.21484E-07 | 2.24268E-05 |
| 7841 | *MOGS* | ENSG00000115275.15 | 5.97 | 6.43 | 0.46 | 1.37 | 0.000304203 | 0.002282889 |
| 118813 | *ZFYVE27* | ENSG00000155256.18 | 5.21 | 5.67 | 0.46 | 1.37 | 7.94793E-05 | 0.000833617 |
| 7386 | *UQCRFS1* | ENSG00000169021.6 | 6.96 | 7.41 | 0.46 | 1.37 | 1.30302E-05 | 0.000223765 |
| 55787 | *TXLNG* | ENSG00000086712.13 | 5.29 | 5.75 | 0.46 | 1.37 | 0.000131773 | 0.001200114 |
| 55788 | *LMBRD1* | ENSG00000168216.13 | 5.94 | 6.40 | 0.46 | 1.37 | 5.82674E-05 | 0.000661524 |
| 3155 | *HMGCL* | ENSG00000117305.15 | 4.62 | 5.08 | 0.46 | 1.37 | 0.000743867 | 0.004523787 |
| 1054 | *CEBPG* | ENSG00000153879.9 | 6.37 | 6.83 | 0.46 | 1.37 | 1.45945E-05 | 0.000240978 |
| 51776 | *MAP3K20* | ENSG00000091436.17 | 6.14 | 6.60 | 0.46 | 1.37 | 0.000160531 | 0.001385511 |
| 9585 | *KIF20B* | ENSG00000138182.15 | 6.52 | 6.97 | 0.46 | 1.37 | 8.59552E-05 | 0.00087762 |
| 2050 | *EPHB4* | ENSG00000196411.10 | 7.78 | 8.24 | 0.46 | 1.37 | 5.62548E-06 | 0.000119424 |
| 65061 | *CDK15* | ENSG00000138395.17 | 4.49 | 4.95 | 0.46 | 1.37 | 0.000691884 | 0.004273452 |
| 10133 | *OPTN* | ENSG00000123240.17 | 6.73 | 7.18 | 0.46 | 1.37 | 2.43448E-05 | 0.000347661 |
| 285753 | *CEP57L1* | ENSG00000183137.15 | 4.64 | 5.10 | 0.46 | 1.37 | 0.001430573 | 0.007511974 |
| 3727 | *JUND* | ENSG00000130522.6 | 6.32 | 6.78 | 0.46 | 1.37 | 0.000399073 | 0.002810737 |
| 3752 | *KCND3* | ENSG00000171385.10 | 5.93 | 6.39 | 0.46 | 1.37 | 1.939E-05 | 0.000293565 |
| 1032 | *CDKN2D* | ENSG00000129355.7 | 3.85 | 4.31 | 0.46 | 1.37 | 0.000954339 | 0.005471226 |
| 1200 | *TPP1* | ENSG00000166340.18 | 6.16 | 6.62 | 0.46 | 1.37 | 8.55721E-05 | 0.000874899 |
| 8826 | *IQGAP1* | ENSG00000140575.13 | 8.81 | 9.26 | 0.45 | 1.37 | 3.14827E-05 | 0.000418163 |
| 5168 | *ENPP2* | ENSG00000136960.13 | 8.72 | 9.17 | 0.45 | 1.37 | 6.78378E-06 | 0.000137406 |
| 8428 | *STK24* | ENSG00000102572.15 | 7.48 | 7.93 | 0.45 | 1.37 | 0.000573971 | 0.003726097 |
| 359948 | *IRF2BP2* | ENSG00000168264.11 | 6.96 | 7.41 | 0.45 | 1.37 | 0.000716139 | 0.004388947 |
| 10329 | *RXYLT1* | ENSG00000118600.13 | 5.49 | 5.94 | 0.45 | 1.37 | 0.000175112 | 0.001485731 |
| 23008 | *KLHDC10* | ENSG00000128607.14 | 7.55 | 8.00 | 0.45 | 1.37 | 0.000651445 | 0.004086804 |
| 51107 | *APH1A* | ENSG00000117362.13 | 8.11 | 8.56 | 0.45 | 1.37 | 9.88809E-06 | 0.000182547 |
| 93081 | *TEX30* | ENSG00000151287.17 | 4.67 | 5.13 | 0.45 | 1.37 | 0.000880427 | 0.00513577 |
| 79586 | *CHPF* | ENSG00000123989.15 | 5.23 | 5.68 | 0.45 | 1.37 | 0.000113637 | 0.001077431 |
| 161742 | *SPRED1* | ENSG00000166068.13 | 4.47 | 4.92 | 0.45 | 1.37 | 0.000789345 | 0.004742704 |
| 3679 | *ITGA7* | ENSG00000135424.19 | 8.13 | 8.58 | 0.45 | 1.37 | 8.3409E-06 | 0.000160088 |
| 8566 | *PDXK* | ENSG00000160209.19 | 7.06 | 7.51 | 0.45 | 1.37 | 0.000324822 | 0.002388656 |
| 23411 | *SIRT1* | ENSG00000096717.12 | 5.48 | 5.93 | 0.45 | 1.37 | 0.0001521 | 0.001335728 |
| 2217 | *FCGRT* | ENSG00000104870.13 | 5.42 | 5.87 | 0.45 | 1.37 | 0.000773396 | 0.004665557 |
| 9276 | *COPB2* | ENSG00000184432.11 | 8.33 | 8.78 | 0.45 | 1.37 | 4.31593E-06 | 9.84466E-05 |
| 1112 | *FOXN3* | ENSG00000053254.16 | 7.71 | 8.16 | 0.45 | 1.37 | 4.27687E-05 | 0.000523585 |
| 7360 | *UGP2* | ENSG00000169764.16 | 7.14 | 7.59 | 0.45 | 1.37 | 1.3952E-05 | 0.000234496 |
| 10695 | *CNPY3* | ENSG00000137161.18 | 5.95 | 6.40 | 0.45 | 1.37 | 4.72648E-06 | 0.00010588 |

| | | | | | | | | |
|---|---|---|---|---|---|---|---|---|
| 7204 | *TRIO* | ENSG00000038382.21 | 5.15 | 5.60 | 0.45 | 1.36 | 0.000829128 | 0.004903223 |
| 196527 | *ANO6* | ENSG00000177119.17 | 7.53 | 7.98 | 0.45 | 1.36 | 6.3743E-06 | 0.000130699 |
| 441951 | *ZFAS1* | ENSG00000177410.13 | 4.96 | 5.41 | 0.45 | 1.36 | 0.000670902 | 0.004167869 |
| 25912 | *C1orf43* | ENSG00000143612.21 | 8.01 | 8.46 | 0.45 | 1.36 | 1.3632E-05 | 0.000231526 |
| 84513 | *PLPP5* | ENSG00000147535.17 | 3.93 | 4.38 | 0.45 | 1.36 | 0.000592419 | 0.003822707 |
| 79875 | *THSD4* | ENSG00000187720.15 | 7.16 | 7.60 | 0.45 | 1.36 | 5.52249E-05 | 0.000634663 |
| 25987 | *TSKU* | ENSG00000182704.8 | 6.46 | 6.91 | 0.45 | 1.36 | 1.32197E-05 | 0.000225985 |
| 64083 | *GOLPH3* | ENSG00000113384.14 | 6.56 | 7.01 | 0.45 | 1.36 | 1.10172E-05 | 0.00019814 |
| 25797 | *QPCT* | ENSG00000115828.17 | 9.72 | 10.16 | 0.45 | 1.36 | 1.35241E-05 | 0.00023014 |
| 641638 | *SNHG6* | ENSG00000245910.9 | 4.89 | 5.34 | 0.45 | 1.36 | 0.000314558 | 0.002337227 |
| 9278 | *ZBTB22* | ENSG00000236104.3 | 4.41 | 4.85 | 0.45 | 1.36 | 0.000255016 | 0.001983178 |
| 9528 | *TMEM59* | ENSG00000116209.12 | 7.57 | 8.01 | 0.45 | 1.36 | 5.68992E-06 | 0.000120351 |
| 64755 | *RUSF1* | ENSG00000140688.17 | 6.18 | 6.63 | 0.45 | 1.36 | 4.20443E-05 | 0.000516824 |
| 9521 | *EEF1E1* | ENSG00000124802.12 | 4.98 | 5.42 | 0.44 | 1.36 | 0.000668905 | 0.004160626 |
| 400 | *ARL1* | ENSG00000120805.14 | 7.16 | 7.60 | 0.44 | 1.36 | 0.000166551 | 0.001425181 |
| 27333 | *GOLIM4* | ENSG00000173905.9 | 6.62 | 7.06 | 0.44 | 1.36 | 9.50616E-06 | 0.00017702 |
| 1603 | *DAD1* | ENSG00000129562.11 | 6.83 | 7.27 | 0.44 | 1.36 | 0.000278122 | 0.00212326 |
| 6238 | *RRBP1* | ENSG00000125844.17 | 7.19 | 7.64 | 0.44 | 1.36 | 5.44483E-05 | 0.00062718 |
| 58528 | *RRAGD* | ENSG00000025039.15 | 6.43 | 6.88 | 0.44 | 1.36 | 5.09519E-05 | 0.0005964 |
| 83930 | *STARD3NL* | ENSG00000010270.14 | 5.37 | 5.81 | 0.44 | 1.36 | 0.000381592 | 0.002711159 |
| 51069 | *MRPL2* | ENSG00000112651.12 | 4.53 | 4.97 | 0.44 | 1.36 | 0.001108979 | 0.006194517 |
| 57515 | *SERINC1* | ENSG00000111897.7 | 8.67 | 9.11 | 0.44 | 1.36 | 1.26728E-05 | 0.000218376 |
| 58478 | *ENOPH1* | ENSG00000145293.17 | 6.36 | 6.80 | 0.44 | 1.36 | 1.57708E-05 | 0.000254795 |
| 25822 | *DNAJB5* | ENSG00000137094.16 | 5.64 | 6.08 | 0.44 | 1.36 | 7.26149E-05 | 0.00077904 |
| 9522 | *SCAMP1* | ENSG00000085365.18 | 7.22 | 7.66 | 0.44 | 1.36 | 2.86876E-06 | 7.31023E-05 |
| 26234 | *FBXL5* | ENSG00000118564.15 | 6.54 | 6.98 | 0.44 | 1.36 | 4.75317E-05 | 0.000568468 |
| 114984 | *FLYWCH2* | ENSG00000162076.13 | 4.51 | 4.96 | 0.44 | 1.36 | 0.001104911 | 0.006175475 |
| 9265 | *CYTH3* | ENSG00000008256.16 | 6.21 | 6.65 | 0.44 | 1.36 | 0.000390917 | 0.002768889 |
| 91452 | *ACBD5* | ENSG00000107897.20 | 6.40 | 6.84 | 0.44 | 1.36 | 0.000224989 | 0.001801951 |
| 124446 | *TMEM219* | ENSG00000149932.17 | 4.53 | 4.97 | 0.44 | 1.36 | 0.00115188 | 0.006367795 |
| 23200 | *ATP11B* | ENSG00000058063.16 | 6.17 | 6.60 | 0.44 | 1.36 | 9.58943E-05 | 0.000951765 |
| 25831 | *HECTD1* | ENSG00000092148.14 | 8.00 | 8.44 | 0.44 | 1.36 | 0.000122652 | 0.001131911 |
| 5495 | *PPM1B* | ENSG00000138032.21 | 6.22 | 6.66 | 0.44 | 1.36 | 0.000134939 | 0.001218311 |
| 51124 | *IER3IP1* | ENSG00000134049.6 | 5.89 | 6.33 | 0.44 | 1.36 | 0.000133767 | 0.001211679 |
| 337867 | *UBAC2* | ENSG00000134882.16 | 7.05 | 7.49 | 0.44 | 1.36 | 1.90869E-05 | 0.000289954 |
| 29015 | *SLC43A3* | ENSG00000134802.18 | 5.06 | 5.50 | 0.44 | 1.36 | 0.000619763 | 0.003951521 |
| 6184 | *RPN1* | ENSG00000163902.12 | 8.66 | 9.09 | 0.44 | 1.36 | 2.36396E-05 | 0.000340506 |
| 10615 | *SPAG5* | ENSG00000076382.17 | 7.03 | 7.47 | 0.44 | 1.36 | 0.000110133 | 0.001053527 |
| 29098 | *RANGRF* | ENSG00000108961.14 | 4.30 | 4.73 | 0.44 | 1.35 | 0.001013758 | 0.005752547 |
| 25978 | *CHMP2B* | ENSG00000083937.10 | 6.09 | 6.52 | 0.44 | 1.35 | 6.51586E-05 | 0.000719092 |
| 3799 | *KIF5B* | ENSG00000170759.11 | 8.51 | 8.94 | 0.44 | 1.35 | 7.09659E-06 | 0.000141077 |
| 1810 | *DR1* | ENSG00000117505.13 | 7.06 | 7.50 | 0.44 | 1.35 | 8.4874E-06 | 0.000162691 |
| 2878 | *GPX3* | ENSG00000211445.13 | 4.85 | 5.28 | 0.43 | 1.35 | 0.001695924 | 0.008524489 |
| 23042 | *PDXDC1* | ENSG00000179889.20 | 7.47 | 7.90 | 0.43 | 1.35 | 1.44581E-05 | 0.000240046 |
| 55041 | *PLEKHB2* | ENSG00000115762.17 | 7.67 | 8.10 | 0.43 | 1.35 | 4.78986E-06 | 0.000106892 |
| 29127 | *RACGAP1* | ENSG00000161800.13 | 7.87 | 8.30 | 0.43 | 1.35 | 0.000255504 | 0.001984917 |
| 23181 | *DIP2A* | ENSG00000160305.18 | 5.84 | 6.27 | 0.43 | 1.35 | 7.12592E-05 | 0.000767788 |
| 11273 | *ATXN2L* | ENSG00000168488.19 | 7.27 | 7.71 | 0.43 | 1.35 | 7.83039E-06 | 0.000152632 |
| 81855 | *SFXN3* | ENSG00000107819.14 | 4.40 | 4.83 | 0.43 | 1.35 | 0.001574658 | 0.008066225 |
| 4061 | *LY6E* | ENSG00000160932.11 | 8.18 | 8.61 | 0.43 | 1.35 | 2.80362E-05 | 0.000384963 |
| 7105 | *TSPAN6* | ENSG00000000003.15 | 6.45 | 6.88 | 0.43 | 1.35 | 0.000646143 | 0.004064525 |
| 79102 | *RNF26* | ENSG00000173456.5 | 6.22 | 6.65 | 0.43 | 1.35 | 5.17206E-05 | 0.000603165 |
| 6525 | *SMTN* | ENSG00000183963.19 | 5.26 | 5.69 | 0.43 | 1.35 | 0.001402194 | 0.007397376 |
| 29089 | *UBE2T* | ENSG00000077152.11 | 5.64 | 6.08 | 0.43 | 1.35 | 0.000793515 | 0.004758235 |
| 5774 | *PTPN3* | ENSG00000070159.14 | 7.28 | 7.71 | 0.43 | 1.35 | 1.63521E-05 | 0.000261094 |
| 219738 | *FAM241B* | ENSG00000171224.9 | 3.64 | 4.07 | 0.43 | 1.35 | 0.001020862 | 0.005788487 |
| 23353 | *SUN1* | ENSG00000164828.18 | 6.97 | 7.40 | 0.43 | 1.35 | 0.000346495 | 0.00251599 |
| 55771 | *PRR11* | ENSG00000068489.13 | 6.37 | 6.80 | 0.43 | 1.35 | 0.001146519 | 0.006352197 |
| 64319 | *FBRS* | ENSG00000156860.16 | 6.12 | 6.55 | 0.43 | 1.35 | 0.000250328 | 0.001953243 |
| 8729 | *GBF1* | ENSG00000107862.7 | 7.17 | 7.60 | 0.43 | 1.35 | 0.00084252 | 0.004966766 |
| 57142 | *RTN4* | ENSG00000115310.19 | 7.74 | 8.18 | 0.43 | 1.35 | 5.64334E-06 | 0.000119634 |
| 10746 | *MAP3K2* | ENSG00000169967.17 | 6.65 | 7.08 | 0.43 | 1.35 | 0.000114092 | 0.001079697 |
| 103910 | *MYL12B* | ENSG00000118680.14 | 8.22 | 8.65 | 0.43 | 1.35 | 4.95887E-06 | 0.000109096 |
| 51622 | *CCZ1* | ENSG00000122674.12 | 5.41 | 5.84 | 0.43 | 1.35 | 0.000765996 | 0.004636593 |
| 23157 | *SEPTIN6* | ENSG00000125354.24 | 7.57 | 8.00 | 0.43 | 1.35 | 3.30276E-06 | 8.08664E-05 |
| 7555 | *CNBP* | ENSG00000169714.17 | 8.34 | 8.77 | 0.43 | 1.35 | 1.67642E-05 | 0.000265216 |
| 23585 | *TMEM50A* | ENSG00000183726.11 | 7.04 | 7.47 | 0.43 | 1.35 | 5.84242E-05 | 0.000662302 |
| 4502 | *MT2A* | ENSG00000125148.7 | 7.82 | 8.25 | 0.43 | 1.35 | 0.002063549 | 0.009945986 |
| 23210 | *JMJD6* | ENSG00000070495.15 | 4.79 | 5.22 | 0.43 | 1.35 | 8.38681E-05 | 0.000863549 |
| 3303 | *HSPA1A* | ENSG00000204389.10 | 6.46 | 6.89 | 0.43 | 1.35 | 0.000399905 | 0.002815277 |
| 1063 | *CENPF* | ENSG00000117724.14 | 8.42 | 8.85 | 0.43 | 1.35 | 0.001521489 | 0.007874491 |
| 54764 | *ZRANB1* | ENSG00000019995.6 | 6.46 | 6.89 | 0.43 | 1.35 | 7.16818E-06 | 0.000141562 |
| 1894 | *ECT2* | ENSG00000114346.14 | 7.99 | 8.42 | 0.43 | 1.35 | 1.78353E-05 | 0.000275118 |
| 6788 | *STK3* | ENSG00000104375.17 | 5.11 | 5.54 | 0.43 | 1.35 | 0.001583113 | 0.008098481 |
| 10149 | *ADGRG2* | ENSG00000173698.18 | 7.35 | 7.78 | 0.43 | 1.35 | 2.93979E-05 | 0.000397507 |
| 11005 | *SPINK5* | ENSG00000133710.16 | 7.62 | 8.05 | 0.43 | 1.35 | 0.000116218 | 0.001092246 |
| 25901 | *CCDC28A* | ENSG00000024862.18 | 4.87 | 5.29 | 0.43 | 1.35 | 5.78503E-05 | 0.000657287 |
| 9550 | *ATP6V1G1* | ENSG00000136888.8 | 7.24 | 7.67 | 0.43 | 1.35 | 2.27563E-05 | 0.000331018 |
| 2339 | *FNTA* | ENSG00000168522.13 | 5.48 | 5.91 | 0.43 | 1.34 | 0.000234202 | 0.001861833 |
| 55322 | *C5orf22* | ENSG00000082213.18 | 5.49 | 5.91 | 0.43 | 1.34 | 8.86883E-05 | 0.000898193 |
| 50999 | *TMED5* | ENSG00000117500.13 | 8.15 | 8.58 | 0.43 | 1.34 | 2.95719E-05 | 0.000398782 |
| 8754 | *ADAM9* | ENSG00000168615.13 | 7.86 | 8.29 | 0.43 | 1.34 | 2.12838E-06 | 5.86527E-05 |
| 80321 | *CEP70* | ENSG00000114107.9 | 5.33 | 5.75 | 0.43 | 1.34 | 8.24784E-05 | 0.000855507 |
| 57534 | *MIB1* | ENSG00000101752.12 | 7.08 | 7.51 | 0.43 | 1.34 | 3.24422E-05 | 0.000427127 |
| 144699 | *FBXL14* | ENSG00000171823.7 | 4.03 | 4.46 | 0.43 | 1.34 | 0.002017117 | 0.009775562 |
| 286451 | *YIPF6* | ENSG00000181704.12 | 5.99 | 6.41 | 0.43 | 1.34 | 0.000149853 | 0.001319922 |
| 9117 | *SEC22C* | ENSG00000093183.14 | 6.61 | 7.03 | 0.43 | 1.34 | 1.34635E-05 | 0.000229369 |

| | | | | | | | | |
|---|---|---|---|---|---|---|---|---|
| 54928 | *BPNT2* | ENSG00000104331.9 | 8.11 | 8.53 | 0.43 | 1.34 | 1.11794E-05 | 0.000199991 |
| 5832 | *ALDH18A1* | ENSG00000059573.9 | 6.75 | 7.17 | 0.43 | 1.34 | 9.96502E-05 | 0.000982042 |
| 7046 | *TGFBR1* | ENSG00000106799.13 | 7.73 | 8.15 | 0.43 | 1.34 | 0.000116037 | 0.00109123 |
| 59345 | *GNB4* | ENSG00000114450.11 | 4.80 | 5.22 | 0.43 | 1.34 | 0.001343881 | 0.007165296 |
| 5905 | *RANGAP1* | ENSG00000100401.20 | 6.55 | 6.97 | 0.42 | 1.34 | 0.000653102 | 0.004091156 |
| 51094 | *ADIPOR1* | ENSG00000159346.13 | 7.02 | 7.44 | 0.42 | 1.34 | 1.38299E-05 | 0.000233491 |
| 23011 | *RAB21* | ENSG00000080371.6 | 6.45 | 6.87 | 0.42 | 1.34 | 0.000172154 | 0.001465604 |
| 10140 | *TOB1* | ENSG00000141232.5 | 7.30 | 7.72 | 0.42 | 1.34 | 4.71276E-06 | 0.000105731 |
| 54890 | *ALKBH5* | ENSG00000091542.9 | 7.28 | 7.71 | 0.42 | 1.34 | 2.61202E-05 | 0.000365157 |
| 55970 | *GNG12* | ENSG00000172380.6 | 8.12 | 8.54 | 0.42 | 1.34 | 2.81619E-06 | 7.2007E-05 |
| 9950 | *GOLGA5* | ENSG00000066455.13 | 5.88 | 6.30 | 0.42 | 1.34 | 0.000152687 | 0.001336255 |
| 10120 | *ACTR1B* | ENSG00000115073.8 | 5.74 | 6.16 | 0.42 | 1.34 | 6.20624E-05 | 0.000691533 |
| 10776 | *ARPP19* | ENSG00000128989.11 | 8.64 | 9.06 | 0.42 | 1.34 | 1.63095E-05 | 0.000260692 |
| 3685 | *ITGAV* | ENSG00000138448.12 | 7.31 | 7.73 | 0.42 | 1.34 | 9.32701E-06 | 0.000174841 |
| 9052 | *GPRC5A* | ENSG00000013588.9 | 5.13 | 5.55 | 0.42 | 1.34 | 0.000138862 | 0.001247266 |
| 219743 | *TYSND1* | ENSG00000156521.14 | 4.13 | 4.55 | 0.42 | 1.34 | 0.00014708 | 0.001302788 |
| 10159 | *ATP6AP2* | ENSG00000182220.15 | 7.39 | 7.81 | 0.42 | 1.34 | 2.05926E-05 | 0.000308457 |
| 83606 | *GUCD1* | ENSG00000138867.17 | 6.75 | 7.17 | 0.42 | 1.34 | 0.000459614 | 0.003132761 |
| 2590 | *GALNT2* | ENSG00000143641.10 | 7.63 | 8.05 | 0.42 | 1.34 | 7.01477E-06 | 0.00013982 |
| 6785 | *ELOVL4* | ENSG00000118402.6 | 5.72 | 6.14 | 0.42 | 1.34 | 0.000120256 | 0.001116908 |
| 22993 | *HMGXB3* | ENSG00000113716.13 | 6.10 | 6.52 | 0.42 | 1.34 | 2.55341E-05 | 0.000360527 |
| 55105 | *GPATCH2* | ENSG00000092978.11 | 5.26 | 5.68 | 0.42 | 1.34 | 0.00032822 | 0.002407747 |
| 81533 | *ITFG1* | ENSG00000129636.13 | 7.32 | 7.74 | 0.42 | 1.34 | 9.25953E-06 | 0.000173938 |
| 147179 | *WIPF2* | ENSG00000171475.14 | 6.16 | 6.58 | 0.42 | 1.34 | 0.000500581 | 0.003345155 |
| 28962 | *OSTM1* | ENSG00000081087.15 | 6.32 | 6.74 | 0.42 | 1.34 | 0.000469071 | 0.00317416 |
| 387882 | *C12orf75* | ENSG00000235162.9 | 4.67 | 5.08 | 0.42 | 1.34 | 0.001073472 | 0.006018583 |
| 286410 | *ATP11C* | ENSG00000101974.15 | 5.66 | 6.08 | 0.42 | 1.34 | 0.000310033 | 0.002314296 |
| 5606 | *MAP2K3* | ENSG00000034152.19 | 4.66 | 5.07 | 0.42 | 1.34 | 0.000495061 | 0.003312694 |
| 21 | *ABCA3* | ENSG00000167972.14 | 5.92 | 6.34 | 0.42 | 1.34 | 4.10552E-05 | 0.000509916 |
| 57157 | *PHTF2* | ENSG00000006576.16 | 6.40 | 6.81 | 0.42 | 1.34 | 3.78181E-05 | 0.000482253 |
| 10475 | *TRIM38* | ENSG00000112343.11 | 4.46 | 4.88 | 0.42 | 1.34 | 0.000352315 | 0.00254961 |
| 51335 | *NGRN* | ENSG00000182768.9 | 7.25 | 7.67 | 0.42 | 1.33 | 1.25292E-05 | 0.000217019 |
| 246175 | *CNOT6L* | ENSG00000138767.14 | 7.05 | 7.47 | 0.42 | 1.33 | 0.000440369 | 0.003028617 |
| 201562 | *HACD2* | ENSG00000206527.10 | 7.04 | 7.45 | 0.42 | 1.33 | 1.19932E-05 | 0.000209798 |
| 55323 | *LARP6* | ENSG00000166173.11 | 5.57 | 5.98 | 0.42 | 1.33 | 0.000215484 | 0.001740686 |
| 122416 | *ANKRD9* | ENSG00000156381.9 | 5.16 | 5.57 | 0.42 | 1.33 | 0.001966783 | 0.009581127 |
| 9493 | *KIF23* | ENSG00000137807.16 | 7.88 | 8.29 | 0.41 | 1.33 | 6.8008E-06 | 0.000137565 |
| 57326 | *PBXIP1* | ENSG00000163346.17 | 8.42 | 8.84 | 0.41 | 1.33 | 3.43591E-06 | 8.30429E-05 |
| 29919 | *RMC1* | ENSG00000141452.10 | 4.85 | 5.27 | 0.41 | 1.33 | 0.000667531 | 0.004153803 |
| 8411 | *EEA1* | ENSG00000102189.17 | 6.35 | 6.77 | 0.41 | 1.33 | 0.000658738 | 0.004114604 |
| 10116 | *FEM1B* | ENSG00000169018.6 | 7.94 | 8.36 | 0.41 | 1.33 | 1.8072E-05 | 0.000277628 |
| 5250 | *SLC25A3* | ENSG00000075415.15 | 8.77 | 9.19 | 0.41 | 1.33 | 2.8062E-05 | 0.000384963 |
| 7417 | *VDAC2* | ENSG00000165637.14 | 6.92 | 7.34 | 0.41 | 1.33 | 4.15249E-05 | 0.000512961 |
| 7873 | *MANF* | ENSG00000145050.19 | 5.79 | 6.21 | 0.41 | 1.33 | 5.43736E-05 | 0.000626847 |
| 64981 | *MRPL34* | ENSG00000130312.7 | 4.76 | 5.17 | 0.41 | 1.33 | 0.000347647 | 0.002523129 |
| 90231 | *KIAA2013* | ENSG00000116685.16 | 5.80 | 6.21 | 0.41 | 1.33 | 0.000133812 | 0.001211679 |
| 64395 | *GMCL1* | ENSG00000087338.5 | 5.30 | 5.72 | 0.41 | 1.33 | 0.000263846 | 0.002037074 |
| 64116 | *SLC39A8* | ENSG00000138821.14 | 7.03 | 7.44 | 0.41 | 1.33 | 1.18369E-05 | 0.00020779 |
| 27107 | *ZBTB11* | ENSG00000066422.6 | 5.79 | 6.21 | 0.41 | 1.33 | 4.93168E-05 | 0.000583291 |
| 372 | *ARCN1* | ENSG00000095139.15 | 8.13 | 8.54 | 0.41 | 1.33 | 1.64479E-05 | 0.0002618 |
| 8812 | *CCNK* | ENSG00000090061.17 | 5.69 | 6.10 | 0.41 | 1.33 | 0.001021471 | 0.00578975 |
| 57645 | *POGK* | ENSG00000143157.12 | 6.76 | 7.17 | 0.41 | 1.33 | 6.97952E-06 | 0.000139454 |
| 51510 | *CHMP5* | ENSG00000086065.14 | 6.19 | 6.60 | 0.41 | 1.33 | 3.41659E-05 | 0.000445135 |
| 205327 | *C2orf69* | ENSG00000178074.6 | 4.28 | 4.69 | 0.41 | 1.33 | 0.000648358 | 0.004073705 |
| 92370 | *PXYLP1* | ENSG00000155893.13 | 7.82 | 8.23 | 0.41 | 1.33 | 1.12458E-05 | 0.000200699 |
| 3939 | *LDHA* | ENSG00000134333.14 | 8.34 | 8.75 | 0.41 | 1.33 | 0.00062819 | 0.003986835 |
| 10390 | *CEPT1* | ENSG00000134255.15 | 5.18 | 5.59 | 0.41 | 1.33 | 0.001382083 | 0.007311836 |
| 3029 | *HAGH* | ENSG00000063854.13 | 4.94 | 5.35 | 0.41 | 1.33 | 0.000649712 | 0.004080135 |
| 112483 | *SAT2* | ENSG00000141504.12 | 5.46 | 5.87 | 0.41 | 1.33 | 0.000135032 | 0.001218311 |
| 54344 | *DPM3* | ENSG00000179085.8 | 4.32 | 4.73 | 0.41 | 1.33 | 0.001281642 | 0.006914507 |
| 51669 | *SARAF* | ENSG00000133872.14 | 7.33 | 7.75 | 0.41 | 1.33 | 9.12348E-06 | 0.000172028 |
| 3091 | *HIF1A* | ENSG00000100644.17 | 9.27 | 9.68 | 0.41 | 1.33 | 5.86765E-06 | 0.000122472 |
| 5516 | *PPP2CB* | ENSG00000104695.13 | 5.97 | 6.38 | 0.41 | 1.33 | 0.000142824 | 0.001273418 |
| 55839 | *CENPN* | ENSG00000166451.14 | 6.82 | 7.23 | 0.41 | 1.33 | 1.59453E-05 | 0.000256122 |
| 64969 | *MRPS5* | ENSG00000144029.12 | 5.33 | 5.74 | 0.41 | 1.33 | 0.000227851 | 0.001818876 |
| 55343 | *SLC35C1* | ENSG00000181830.9 | 4.80 | 5.21 | 0.41 | 1.33 | 0.000125559 | 0.001152612 |
| 10652 | *YKT6* | ENSG00000106636.9 | 8.18 | 8.59 | 0.41 | 1.33 | 1.12007E-05 | 0.000200133 |
| 9337 | *CNOT8* | ENSG00000155508.14 | 6.28 | 6.69 | 0.41 | 1.33 | 4.33362E-05 | 0.000529237 |
| 5471 | *PPAT* | ENSG00000128059.8 | 5.99 | 6.40 | 0.41 | 1.33 | 0.000397954 | 0.002805491 |
| 10440 | *TIMM17A* | ENSG00000134375.11 | 6.40 | 6.81 | 0.41 | 1.33 | 7.19644E-05 | 0.000774384 |
| 84925 | *SLC49A4* | ENSG00000138463.9 | 5.50 | 5.91 | 0.41 | 1.33 | 0.000177747 | 0.001496243 |
| 54926 | *UBE2R2* | ENSG00000107341.5 | 6.78 | 7.19 | 0.41 | 1.33 | 9.58809E-06 | 0.000178324 |
| 53838 | *C11orf24* | ENSG00000171067.11 | 5.29 | 5.70 | 0.41 | 1.33 | 9.19166E-05 | 0.00092403 |
| 6166 | *RPL36AL* | ENSG00000165502.7 | 6.70 | 7.11 | 0.41 | 1.33 | 2.59516E-05 | 0.000364507 |
| 56942 | *CMC2* | ENSG00000103121.9 | 5.55 | 5.95 | 0.41 | 1.33 | 0.00011633 | 0.00109261 |
| 25820 | *ARIH1* | ENSG00000166233.15 | 7.26 | 7.67 | 0.41 | 1.33 | 0.00120377 | 0.006581927 |
| 3775 | *KCNK1* | ENSG00000135750.15 | 4.92 | 5.33 | 0.41 | 1.33 | 0.000214979 | 0.001737542 |
| 51187 | *RSL24D1* | ENSG00000137876.11 | 6.97 | 7.38 | 0.41 | 1.33 | 7.05294E-05 | 0.000762663 |
| 7385 | *UQCRC2* | ENSG00000140740.11 | 6.98 | 7.39 | 0.41 | 1.33 | 7.92495E-05 | 0.000832369 |
| 124842 | *TMEM132E* | ENSG00000181291.8 | 6.81 | 7.21 | 0.41 | 1.33 | 0.000481167 | 0.003244311 |
| 9853 | *RUSC2* | ENSG00000198853.12 | 6.08 | 6.49 | 0.41 | 1.32 | 0.000170219 | 0.001451604 |
| 5868 | *RAB5A* | ENSG00000144566.11 | 6.11 | 6.52 | 0.41 | 1.32 | 0.000104458 | 0.001014771 |
| 29927 | *SEC61A1* | ENSG00000058262.10 | 9.04 | 9.45 | 0.41 | 1.32 | 1.17165E-05 | 0.0002064 |
| 85450 | *ITPRIP* | ENSG00000148841.17 | 4.36 | 4.77 | 0.40 | 1.32 | 0.001282527 | 0.006915324 |
| 5861 | *RAB1A* | ENSG00000138069.18 | 7.04 | 7.44 | 0.40 | 1.32 | 1.68624E-05 | 0.000265848 |
| 54749 | *EPDR1* | ENSG00000086289.12 | 6.39 | 6.80 | 0.40 | 1.32 | 0.000173911 | 0.001478043 |
| 54861 | *SNRK* | ENSG00000163788.14 | 5.22 | 5.62 | 0.40 | 1.32 | 0.000686228 | 0.004243756 |

| | | | | | | | | |
|---|---|---|---|---|---|---|---|---|
| 27131 | *SNX5* | ENSG00000089006.17 | 6.92 | 7.33 | 0.40 | 1.32 | 0.000269192 | 0.002069828 |
| 345778 | *MTX3* | ENSG00000177034.17 | 6.39 | 6.79 | 0.40 | 1.32 | 7.08765E-05 | 0.000765313 |
| 55744 | *COA1* | ENSG00000106603.20 | 7.24 | 7.64 | 0.40 | 1.32 | 1.23925E-05 | 0.000215526 |
| 7965 | *AIMP2* | ENSG00000106305.10 | 4.85 | 5.26 | 0.40 | 1.32 | 0.00014352 | 0.001277637 |
| 259266 | *ASPM* | ENSG00000066279.19 | 7.35 | 7.76 | 0.40 | 1.32 | 5.88411E-05 | 0.00066492 |
| 8086 | *AAAS* | ENSG00000094914.14 | 6.23 | 6.64 | 0.40 | 1.32 | 7.77515E-05 | 0.000821233 |
| 30834 | *POLR1H* | ENSG00000066379.15 | 3.58 | 3.99 | 0.40 | 1.32 | 0.001952037 | 0.009527844 |
| 90488 | *TMEM263* | ENSG00000151135.10 | 7.13 | 7.53 | 0.40 | 1.32 | 1.08808E-05 | 0.000196535 |
| 23673 | *STX12* | ENSG00000117758.14 | 6.01 | 6.41 | 0.40 | 1.32 | 0.000110218 | 0.00105367 |
| 3703 | *STT3A* | ENSG00000134910.14 | 7.96 | 8.37 | 0.40 | 1.32 | 0.000142877 | 0.001273418 |
| 84333 | *PCGF5* | ENSG00000180628.15 | 6.75 | 7.15 | 0.40 | 1.32 | 0.000236951 | 0.001879703 |
| 10725 | *NFAT5* | ENSG00000102908.22 | 6.05 | 6.45 | 0.40 | 1.32 | 0.000856376 | 0.005022179 |
| 3660 | *IRF2* | ENSG00000168310.11 | 4.88 | 5.28 | 0.40 | 1.32 | 0.001862265 | 0.009176211 |
| 51447 | *IP6K2* | ENSG00000068745.15 | 5.45 | 5.85 | 0.40 | 1.32 | 0.000244461 | 0.001914987 |
| 482 | *ATP1B2* | ENSG00000129244.9 | 5.95 | 6.35 | 0.40 | 1.32 | 0.000302613 | 0.002274367 |
| 51310 | *SLC22A17* | ENSG00000092096.19 | 5.51 | 5.91 | 0.40 | 1.32 | 0.001322542 | 0.007079184 |
| 267 | *AMFR* | ENSG00000159461.15 | 7.61 | 8.01 | 0.40 | 1.32 | 1.68046E-05 | 0.000265216 |
| 80727 | *TTYH3* | ENSG00000136295.15 | 6.74 | 7.14 | 0.40 | 1.32 | 3.71268E-05 | 0.00047678 |
| 284612 | *SYPL2* | ENSG00000143028.9 | 3.79 | 4.19 | 0.40 | 1.32 | 0.000436962 | 0.003013954 |
| 994 | *CDC25B* | ENSG00000101224.18 | 6.42 | 6.82 | 0.40 | 1.32 | 0.000118558 | 0.001106613 |
| 80830 | *APOL6* | ENSG00000221963.6 | 5.75 | 6.15 | 0.40 | 1.32 | 0.000231033 | 0.001839559 |
| 25847 | *ANAPC13* | ENSG00000129055.13 | 6.43 | 6.83 | 0.40 | 1.32 | 8.03477E-05 | 0.000839206 |
| 5087 | *PBX1* | ENSG00000185630.19 | 8.67 | 9.07 | 0.40 | 1.32 | 1.73891E-05 | 0.00027074 |
| 93664 | *CADPS2* | ENSG00000081803.17 | 5.04 | 5.43 | 0.40 | 1.32 | 0.001168826 | 0.006428329 |
| 94032 | *CAMK2N2* | ENSG00000163888.4 | 5.02 | 5.42 | 0.40 | 1.32 | 0.001430306 | 0.007511974 |
| 26049 | *FAM169A* | ENSG00000198780.13 | 6.78 | 7.18 | 0.40 | 1.32 | 0.000451361 | 0.003090548 |
| 54793 | *KCTD9* | ENSG00000104756.16 | 4.68 | 5.07 | 0.40 | 1.32 | 0.001542776 | 0.007959958 |
| 23467 | *NPTXR* | ENSG00000221890.6 | 5.77 | 6.16 | 0.40 | 1.32 | 5.83831E-05 | 0.000662302 |
| 10525 | *HYOU1* | ENSG00000149428.20 | 8.34 | 8.74 | 0.40 | 1.32 | 1.07932E-05 | 0.00019541 |
| 6628 | *SNRPB* | ENSG00000125835.20 | 7.47 | 7.87 | 0.40 | 1.32 | 6.17031E-05 | 0.000689064 |
| 51 | *ACOX1* | ENSG00000161533.12 | 5.82 | 6.22 | 0.39 | 1.31 | 0.001141257 | 0.006334968 |
| 2720 | *GLB1* | ENSG00000170266.16 | 7.09 | 7.48 | 0.39 | 1.31 | 0.000284576 | 0.002162632 |
| 200576 | *PIKFYVE* | ENSG00000115020.17 | 5.38 | 5.78 | 0.39 | 1.31 | 0.000236293 | 0.001875474 |
| 6235 | *RPS29* | ENSG00000213741.11 | 6.48 | 6.87 | 0.39 | 1.31 | 0.002015452 | 0.009770646 |
| 677 | *ZFP36L1* | ENSG00000185650.10 | 5.04 | 5.43 | 0.39 | 1.31 | 0.001220421 | 0.006651164 |
| 8933 | *RTL8C* | ENSG00000134590.14 | 6.21 | 6.60 | 0.39 | 1.31 | 3.46883E-05 | 0.000449988 |
| 6176 | *RPLP1* | ENSG00000137818.12 | 9.40 | 9.80 | 0.39 | 1.31 | 0.000107515 | 0.001036918 |
| 3052 | *HCCS* | ENSG00000004961.15 | 5.37 | 5.76 | 0.39 | 1.31 | 0.000139596 | 0.001252362 |
| 900 | *CCNG1* | ENSG00000113328.19 | 6.78 | 7.17 | 0.39 | 1.31 | 2.11101E-05 | 0.000313705 |
| 9474 | *ATG5* | ENSG00000057663.16 | 5.82 | 6.21 | 0.39 | 1.31 | 0.000628214 | 0.003986835 |
| 57617 | *VPS18* | ENSG00000104142.11 | 5.85 | 6.24 | 0.39 | 1.31 | 0.000847977 | 0.004995008 |
| 56889 | *TM9SF3* | ENSG00000077147.16 | 8.52 | 8.91 | 0.39 | 1.31 | 2.802E-05 | 0.000384963 |
| 221937 | *FOXK1* | ENSG00000164916.11 | 6.36 | 6.75 | 0.39 | 1.31 | 0.000134964 | 0.001218311 |
| 10124 | *ARL4A* | ENSG00000122644.13 | 5.31 | 5.70 | 0.39 | 1.31 | 0.000311738 | 0.002322022 |
| 84336 | *TMEM101* | ENSG00000091947.10 | 4.45 | 4.84 | 0.39 | 1.31 | 0.001980724 | 0.009630282 |
| 54732 | *TMED9* | ENSG00000184840.12 | 7.10 | 7.49 | 0.39 | 1.31 | 0.000189192 | 0.001570563 |
| 23074 | *UHRF1BP1L* | ENSG00000111647.13 | 6.22 | 6.61 | 0.39 | 1.31 | 0.00014963 | 0.001318726 |
| 7988 | *ZNF212* | ENSG00000170260.9 | 4.91 | 5.29 | 0.39 | 1.31 | 0.000768501 | 0.004647236 |
| 51727 | *CMPK1* | ENSG00000162368.14 | 7.70 | 8.09 | 0.39 | 1.31 | 3.30053E-05 | 0.0004334 |
| 6892 | *TAPBP* | ENSG00000231925.13 | 7.26 | 7.65 | 0.39 | 1.31 | 6.15816E-05 | 0.000688219 |
| 2935 | *GSPT1* | ENSG00000103342.13 | 8.14 | 8.53 | 0.39 | 1.31 | 7.10893E-06 | 0.000141135 |
| 8665 | *EIF3F* | ENSG00000175390.15 | 6.13 | 6.52 | 0.39 | 1.31 | 0.000204816 | 0.001668937 |
| 85403 | *EAF1* | ENSG00000144597.14 | 6.19 | 6.57 | 0.39 | 1.31 | 0.000109785 | 0.001052213 |
| 170954 | *PPP1R18* | ENSG00000146112.12 | 5.55 | 5.94 | 0.39 | 1.31 | 0.000222223 | 0.001785517 |
| 80723 | *SLC35G2* | ENSG00000168917.9 | 4.61 | 5.00 | 0.39 | 1.31 | 0.000860979 | 0.005039779 |
| 55872 | *PBK* | ENSG00000168078.10 | 6.00 | 6.39 | 0.39 | 1.31 | 8.45738E-05 | 0.000868835 |
| 54504 | *CPVL* | ENSG00000106066.15 | 6.07 | 6.45 | 0.39 | 1.31 | 2.85626E-05 | 0.000389369 |
| 23432 | *GPR161* | ENSG00000143147.15 | 5.22 | 5.60 | 0.39 | 1.31 | 0.001504046 | 0.007795559 |
| 51512 | *GTSE1* | ENSG00000075218.19 | 5.69 | 6.08 | 0.39 | 1.31 | 0.000212622 | 0.001720347 |
| 4698 | *NDUFA5* | ENSG00000128609.16 | 8.00 | 8.38 | 0.39 | 1.31 | 3.05152E-05 | 0.000408566 |
| 10509 | *SEMA4B* | ENSG00000185033.15 | 5.99 | 6.37 | 0.39 | 1.31 | 0.000140699 | 0.001257741 |
| 23423 | *TMED3* | ENSG00000166557.14 | 5.56 | 5.94 | 0.39 | 1.31 | 0.000105391 | 0.00102118 |
| 8237 | *USP11* | ENSG00000102226.10 | 7.29 | 7.67 | 0.39 | 1.31 | 0.000349085 | 0.002530651 |
| 22931 | *RAB18* | ENSG00000099246.18 | 7.48 | 7.87 | 0.39 | 1.31 | 5.56443E-05 | 0.000637531 |
| 9055 | *PRC1* | ENSG00000198901.14 | 8.42 | 8.80 | 0.38 | 1.31 | 1.02667E-05 | 0.000188148 |
| 26046 | *LTN1* | ENSG00000198862.14 | 6.22 | 6.60 | 0.38 | 1.31 | 9.52548E-05 | 0.000948387 |
| 28972 | *SPCS1* | ENSG00000114902.14 | 7.04 | 7.43 | 0.38 | 1.31 | 6.67106E-05 | 0.000730846 |
| 25994 | *HIGD1A* | ENSG00000181061.14 | 7.03 | 7.41 | 0.38 | 1.30 | 0.000494301 | 0.003310557 |
| 9445 | *ITM2B* | ENSG00000136156.15 | 8.31 | 8.69 | 0.38 | 1.30 | 1.09012E-05 | 0.000196617 |
| 221035 | *REEP3* | ENSG00000165476.14 | 5.43 | 5.82 | 0.38 | 1.30 | 0.000948088 | 0.005452052 |
| 112942 | *CFAP36* | ENSG00000163001.12 | 4.83 | 5.21 | 0.38 | 1.30 | 0.000861805 | 0.005040855 |
| 28989 | *NTMT1* | ENSG00000148335.15 | 5.74 | 6.12 | 0.38 | 1.30 | 0.000521087 | 0.00344689 |
| 55165 | *CEP55* | ENSG00000138180.16 | 6.33 | 6.71 | 0.38 | 1.30 | 2.40414E-05 | 0.000344639 |
| 4052 | *LTBP1* | ENSG00000049323.16 | 8.11 | 8.49 | 0.38 | 1.30 | 0.001246132 | 0.00676182 |
| 10730 | *YME1L1* | ENSG00000136758.20 | 8.02 | 8.40 | 0.38 | 1.30 | 8.28704E-05 | 0.000858692 |
| 11041 | *B4GAT1* | ENSG00000174684.7 | 5.85 | 6.23 | 0.38 | 1.30 | 0.000122197 | 0.001130735 |
| 23207 | *PLEKHM2* | ENSG00000116786.13 | 6.17 | 6.55 | 0.38 | 1.30 | 8.59048E-05 | 0.00087762 |
| 6272 | *SORT1* | ENSG00000134243.12 | 7.82 | 8.20 | 0.38 | 1.30 | 4.14593E-05 | 0.000512848 |
| 51504 | *TRMT112* | ENSG00000173113.7 | 6.53 | 6.91 | 0.38 | 1.30 | 3.19052E-05 | 0.000423026 |
| 167153 | *TENT2* | ENSG00000164329.14 | 6.17 | 6.55 | 0.38 | 1.30 | 0.000755298 | 0.00458216 |
| 200916 | *RPL22L1* | ENSG00000163584.18 | 5.17 | 5.55 | 0.38 | 1.30 | 0.000355855 | 0.002571509 |
| 1198 | *CLK3* | ENSG00000179335.20 | 5.71 | 6.09 | 0.38 | 1.30 | 0.001279633 | 0.006906154 |
| 57544 | *TXNDC16* | ENSG00000087301.9 | 6.28 | 6.66 | 0.38 | 1.30 | 5.97771E-05 | 0.000673064 |
| 7525 | *YES1* | ENSG00000176105.14 | 7.29 | 7.67 | 0.38 | 1.30 | 1.62586E-05 | 0.000260156 |
| 23582 | *CCNDBP1* | ENSG00000166946.14 | 5.36 | 5.74 | 0.38 | 1.30 | 0.000319323 | 0.002364443 |
| 348801 | *LNP1* | ENSG00000206535.8 | 4.23 | 4.62 | 0.38 | 1.30 | 0.000327985 | 0.002407204 |
| 10983 | *CCNI* | ENSG00000118816.11 | 8.07 | 8.45 | 0.38 | 1.30 | 1.45213E-05 | 0.000240484 |

| | | | | | | | | |
|---|---|---|---|---|---|---|---|---|
| 2734 | *GLG1* | ENSG00000090863.12 | 8.35 | 8.73 | 0.38 | 1.30 | 2.68201E-05 | 0.000372036 |
| 133619 | *PRRC1* | ENSG00000164244.21 | 6.77 | 7.15 | 0.38 | 1.30 | 0.001047447 | 0.005907696 |
| 54965 | *PIGX* | ENSG00000163964.18 | 5.57 | 5.95 | 0.38 | 1.30 | 0.000117765 | 0.001101647 |
| 8727 | *CTNNAL1* | ENSG00000119326.15 | 8.87 | 9.25 | 0.38 | 1.30 | 1.55202E-05 | 0.000252471 |
| 154810 | *AMOTL1* | ENSG00000166025.18 | 7.07 | 7.45 | 0.38 | 1.30 | 0.000108437 | 0.001042501 |
| 1390 | *CREM* | ENSG00000095794.20 | 6.85 | 7.22 | 0.38 | 1.30 | 5.32054E-05 | 0.000616173 |
| 5372 | *PMM1* | ENSG00000100417.12 | 4.48 | 4.85 | 0.38 | 1.30 | 0.002050417 | 0.009895405 |
| 4217 | *MAP3K5* | ENSG00000197442.10 | 6.14 | 6.52 | 0.38 | 1.30 | 0.000464193 | 0.003153952 |
| 5792 | *PTPRF* | ENSG00000142949.17 | 7.49 | 7.87 | 0.38 | 1.30 | 0.00048476 | 0.003262676 |
| 3835 | *KIF22* | ENSG00000079616.14 | 6.68 | 7.06 | 0.38 | 1.30 | 5.06784E-05 | 0.000595174 |
| 2132 | *EXT2* | ENSG00000151348.16 | 7.62 | 8.00 | 0.38 | 1.30 | 0.000238815 | 0.001885519 |
| 57291 | *DANCR* | ENSG00000226950.8 | 5.07 | 5.45 | 0.38 | 1.30 | 0.001188427 | 0.006517024 |
| 5045 | *FURIN* | ENSG00000140564.13 | 6.02 | 6.39 | 0.38 | 1.30 | 0.000396807 | 0.002802619 |
| 9296 | *ATP6V1F* | ENSG00000128524.5 | 6.61 | 6.99 | 0.38 | 1.30 | 0.000112999 | 0.001074781 |
| 6480 | *ST6GAL1* | ENSG00000073849.16 | 6.56 | 6.93 | 0.38 | 1.30 | 0.000282093 | 0.002148114 |
| 5584 | *PRKCI* | ENSG00000163558.13 | 6.73 | 7.10 | 0.37 | 1.30 | 5.95875E-05 | 0.000671434 |
| 26224 | *FBXL3* | ENSG00000005812.11 | 5.56 | 5.93 | 0.37 | 1.30 | 4.00008E-05 | 0.000502671 |
| 10010 | *TANK* | ENSG00000136560.14 | 5.62 | 5.99 | 0.37 | 1.30 | 0.000189037 | 0.001570146 |
| 28996 | *HIPK2* | ENSG00000064393.16 | 7.64 | 8.01 | 0.37 | 1.29 | 8.8524E-05 | 0.000897741 |
| 805 | *CALM2* | ENSG00000143933.20 | 8.93 | 9.31 | 0.37 | 1.29 | 1.61206E-05 | 0.000258497 |
| 51074 | *APIP* | ENSG00000149089.13 | 4.57 | 4.94 | 0.37 | 1.29 | 0.001404037 | 0.007404492 |
| 64065 | *PERP* | ENSG00000112378.12 | 4.47 | 4.84 | 0.37 | 1.29 | 0.00052842 | 0.003486176 |
| 55770 | *EXOC2* | ENSG00000112685.14 | 6.09 | 6.46 | 0.37 | 1.29 | 3.35373E-05 | 0.00043885 |
| 57120 | *GOPC* | ENSG00000047932.14 | 6.29 | 6.66 | 0.37 | 1.29 | 0.000179072 | 0.001501506 |
| 3619 | *INCENP* | ENSG00000149503.13 | 7.14 | 7.51 | 0.37 | 1.29 | 0.001841251 | 0.009099549 |
| 8303 | *SNN* | ENSG00000184602.6 | 6.49 | 6.86 | 0.37 | 1.29 | 3.21783E-05 | 0.00042477 |
| 4253 | *MIA2* | ENSG00000150527.18 | 8.62 | 8.99 | 0.37 | 1.29 | 6.55637E-05 | 0.000722693 |
| 8099 | *CDK2AP1* | ENSG00000111328.7 | 5.89 | 6.26 | 0.37 | 1.29 | 0.000594023 | 0.003828393 |
| 401548 | *SNX30* | ENSG00000148158.17 | 6.07 | 6.44 | 0.37 | 1.29 | 9.30765E-05 | 0.000931945 |
| 339287 | *MSL1* | ENSG00000188895.12 | 6.60 | 6.96 | 0.37 | 1.29 | 0.000543386 | 0.003564548 |
| 10403 | *NDC80* | ENSG00000080986.13 | 5.95 | 6.32 | 0.37 | 1.29 | 0.000648416 | 0.004073705 |
| 5269 | *SERPINB6* | ENSG00000124570.21 | 6.05 | 6.42 | 0.37 | 1.29 | 0.000218687 | 0.001759934 |
| 5527 | *PPP2R5C* | ENSG00000078304.19 | 7.64 | 8.01 | 0.37 | 1.29 | 2.96502E-05 | 0.000399479 |
| 57117 | *INTS12* | ENSG00000138785.16 | 4.16 | 4.53 | 0.37 | 1.29 | 0.001711228 | 0.008581407 |
| 6867 | *TACC1* | ENSG00000147526.20 | 7.24 | 7.61 | 0.37 | 1.29 | 0.000160526 | 0.001385511 |
| 54386 | *TERF2IP* | ENSG00000166848.7 | 6.75 | 7.12 | 0.37 | 1.29 | 6.02661E-05 | 0.000677046 |
| 55632 | *G2E3* | ENSG00000092140.16 | 5.46 | 5.83 | 0.37 | 1.29 | 0.001976581 | 0.009616371 |
| 6558 | *SLC12A2* | ENSG00000064651.14 | 6.04 | 6.41 | 0.37 | 1.29 | 0.001817256 | 0.009010639 |
| 64131 | *XYLT1* | ENSG00000103489.12 | 7.10 | 7.47 | 0.37 | 1.29 | 0.000138234 | 0.001242593 |
| 10424 | *PGRMC2* | ENSG00000164040.18 | 6.43 | 6.80 | 0.37 | 1.29 | 0.001007895 | 0.005723823 |
| 54536 | *EXOC6* | ENSG00000138190.17 | 5.89 | 6.26 | 0.37 | 1.29 | 9.17633E-05 | 0.000923425 |
| 2990 | *GUSB* | ENSG00000169919.17 | 5.80 | 6.17 | 0.36 | 1.29 | 0.000994162 | 0.005660049 |
| 4240 | *MFGE8* | ENSG00000140545.15 | 6.96 | 7.32 | 0.36 | 1.29 | 2.66742E-05 | 0.000371013 |
| 83658 | *DYNLRB1* | ENSG00000125971.17 | 6.53 | 6.90 | 0.36 | 1.29 | 0.000238992 | 0.001885924 |
| 6655 | *SOS2* | ENSG00000100485.12 | 6.60 | 6.97 | 0.36 | 1.29 | 4.18309E-05 | 0.000515892 |
| 116068 | *LYSMD3* | ENSG00000176018.13 | 6.60 | 6.97 | 0.36 | 1.29 | 0.000111604 | 0.001063535 |
| 283431 | *GAS2L3* | ENSG00000139354.11 | 6.25 | 6.62 | 0.36 | 1.29 | 0.001924205 | 0.009416498 |
| 10723 | *SLC12A7* | ENSG00000113504.21 | 5.62 | 5.99 | 0.36 | 1.29 | 0.000429187 | 0.002980871 |
| 4105 | *MAGEA6* | ENSG00000197172.11 | 7.17 | 7.53 | 0.36 | 1.29 | 0.000118917 | 0.001107896 |
| 84918 | *LRP11* | ENSG00000120256.11 | 7.07 | 7.43 | 0.36 | 1.29 | 0.000380468 | 0.002706375 |
| 7009 | *TMBIM6* | ENSG00000139644.13 | 10.84 | 11.20 | 0.36 | 1.28 | 3.23564E-05 | 0.000426746 |
| 7040 | *TGFB1* | ENSG00000105329.11 | 5.76 | 6.13 | 0.36 | 1.28 | 0.000509203 | 0.00338769 |
| 8804 | *CREG1* | ENSG00000143162.9 | 6.20 | 6.56 | 0.36 | 1.28 | 8.3049E-05 | 0.000859051 |
| 83667 | *SESN2* | ENSG00000130766.5 | 4.52 | 4.88 | 0.36 | 1.28 | 0.001449319 | 0.007589961 |
| 4190 | *MDH1* | ENSG00000014641.21 | 7.95 | 8.31 | 0.36 | 1.28 | 6.8091E-05 | 0.000743257 |
| 8545 | *CGGBP1* | ENSG00000163320.12 | 7.57 | 7.93 | 0.36 | 1.28 | 0.000162071 | 0.001393195 |
| 780 | *DDR1* | ENSG00000204580.14 | 8.90 | 9.26 | 0.36 | 1.28 | 0.000138259 | 0.001242593 |
| 27125 | *AFF4* | ENSG00000072364.13 | 7.79 | 8.15 | 0.36 | 1.28 | 0.000460542 | 0.003137666 |
| 10959 | *TMED2* | ENSG00000086598.11 | 8.16 | 8.52 | 0.36 | 1.28 | 0.000125441 | 0.001152612 |
| 3460 | *IFNGR2* | ENSG00000159128.15 | 5.63 | 5.99 | 0.36 | 1.28 | 0.000630336 | 0.00399355 |
| 8140 | *SLC7A5* | ENSG00000103257.9 | 6.01 | 6.36 | 0.36 | 1.28 | 0.000634959 | 0.004012674 |
| 11252 | *PACSIN2* | ENSG00000100266.19 | 5.58 | 5.93 | 0.36 | 1.28 | 0.00191008 | 0.009368757 |
| 9144 | *SYNGR2* | ENSG00000108639.8 | 5.58 | 5.94 | 0.36 | 1.28 | 0.000506543 | 0.003372983 |
| 3074 | *HEXB* | ENSG00000049860.14 | 6.82 | 7.18 | 0.36 | 1.28 | 7.12181E-05 | 0.000767788 |
| 5236 | *PGM1* | ENSG00000079739.17 | 7.24 | 7.60 | 0.36 | 1.28 | 0.000112963 | 0.001074781 |
| 9919 | *SEC16A* | ENSG00000148396.19 | 7.43 | 7.79 | 0.36 | 1.28 | 0.000310768 | 0.002318247 |
| 23268 | *DNMBP* | ENSG00000107554.17 | 5.42 | 5.78 | 0.36 | 1.28 | 0.001461236 | 0.007641706 |
| 7110 | *TMF1* | ENSG00000144747.17 | 6.26 | 6.62 | 0.36 | 1.28 | 0.000281916 | 0.002147853 |
| 6160 | *RPL31* | ENSG00000071082.11 | 8.02 | 8.38 | 0.36 | 1.28 | 0.001308209 | 0.007030043 |
| 7295 | *TXN* | ENSG00000136810.13 | 7.86 | 8.22 | 0.36 | 1.28 | 8.62274E-05 | 0.000879801 |
| 84221 | *SPATC1L* | ENSG00000160284.15 | 4.03 | 4.38 | 0.36 | 1.28 | 0.001500537 | 0.007792925 |
| 11321 | *GPN1* | ENSG00000198522.14 | 5.42 | 5.78 | 0.36 | 1.28 | 0.000111313 | 0.00106278 |
| 10608 | *MXD4* | ENSG00000123933.17 | 5.57 | 5.92 | 0.36 | 1.28 | 0.00150137 | 0.007794558 |
| 7533 | *YWHAH* | ENSG00000128245.15 | 8.12 | 8.47 | 0.36 | 1.28 | 7.20902E-05 | 0.000774384 |
| 9928 | *KIF14* | ENSG00000118193.12 | 5.61 | 5.97 | 0.36 | 1.28 | 0.000637703 | 0.004024931 |
| 84896 | *ATAD1* | ENSG00000138138.14 | 6.53 | 6.89 | 0.35 | 1.28 | 0.000308974 | 0.002309249 |
| 38 | *ACAT1* | ENSG00000075239.14 | 7.24 | 7.60 | 0.35 | 1.28 | 5.67872E-05 | 0.000647167 |
| 51290 | *ERGIC2* | ENSG00000087502.19 | 7.06 | 7.41 | 0.35 | 1.28 | 0.000675037 | 0.004186624 |
| 10092 | *ARPC5* | ENSG00000162704.16 | 8.38 | 8.74 | 0.35 | 1.28 | 4.3579E-05 | 0.000530907 |
| 90417 | *KNSTRN* | ENSG00000128944.14 | 6.22 | 6.57 | 0.35 | 1.28 | 0.000161676 | 0.001390984 |
| 56851 | *EMC7* | ENSG00000134153.10 | 6.58 | 6.93 | 0.35 | 1.28 | 0.000152182 | 0.001335728 |
| 25940 | *FAM98A* | ENSG00000119812.20 | 6.36 | 6.71 | 0.35 | 1.28 | 0.000153507 | 0.001341087 |
| 4925 | *NUCB2* | ENSG00000070081.18 | 5.74 | 6.09 | 0.35 | 1.28 | 0.00019327 | 0.001597351 |
| 22974 | *TPX2* | ENSG00000088325.16 | 8.36 | 8.72 | 0.35 | 1.28 | 1.09122E-05 | 0.000196617 |
| 11231 | *SEC63* | ENSG00000025796.14 | 7.71 | 8.06 | 0.35 | 1.28 | 0.001274359 | 0.006890079 |
| 3608 | *ILF2* | ENSG00000143621.17 | 8.77 | 9.12 | 0.35 | 1.28 | 1.26719E-05 | 0.000218376 |
| 10376 | *TUBA1B* | ENSG00000123416.15 | 11.56 | 11.92 | 0.35 | 1.28 | 8.33125E-05 | 0.00086059 |

| | | | | | | | | |
|---|---|---|---|---|---|---|---|---|
| 5376 | *PMP22* | ENSG00000109099.16 | 5.21 | 5.56 | 0.35 | 1.28 | 0.000531341 | 0.003501882 |
| 5290 | *PIK3CA* | ENSG00000121879.6 | 5.62 | 5.97 | 0.35 | 1.28 | 0.000852643 | 0.005009827 |
| 56681 | *SAR1A* | ENSG00000079332.15 | 7.13 | 7.48 | 0.35 | 1.28 | 5.95059E-05 | 0.000671017 |
| 729082 | *OIP5-AS1* | ENSG00000247556.6 | 7.99 | 8.34 | 0.35 | 1.28 | 0.000189584 | 0.001572077 |
| 55633 | *TBC1D22B* | ENSG00000065491.8 | 5.02 | 5.37 | 0.35 | 1.28 | 0.000748243 | 0.004546714 |
| 65125 | *WNK1* | ENSG00000060237.19 | 9.19 | 9.54 | 0.35 | 1.28 | 0.000372312 | 0.002660969 |
| 222484 | *LNX2* | ENSG00000139517.9 | 5.25 | 5.60 | 0.35 | 1.27 | 0.000462848 | 0.003146542 |
| 200933 | *FBXO45* | ENSG00000174013.8 | 5.84 | 6.19 | 0.35 | 1.27 | 0.000323133 | 0.002383096 |
| 3485 | *IGFBP2* | ENSG00000115457.10 | 7.95 | 8.30 | 0.35 | 1.27 | 0.000297036 | 0.002241431 |
| 9787 | *DLGAP5* | ENSG00000126787.13 | 6.91 | 7.26 | 0.35 | 1.27 | 0.000178378 | 0.001498194 |
| 10542 | *LAMTOR5* | ENSG00000134248.14 | 5.94 | 6.29 | 0.35 | 1.27 | 0.000267116 | 0.002059142 |
| 55161 | *TMEM33* | ENSG00000109133.13 | 7.78 | 8.13 | 0.35 | 1.27 | 3.89914E-05 | 0.000493607 |
| 10483 | *SEC23B* | ENSG00000101310.17 | 7.13 | 7.48 | 0.35 | 1.27 | 0.000136756 | 0.00123201 |
| 790 | *CAD* | ENSG00000084774.14 | 6.88 | 7.23 | 0.35 | 1.27 | 0.001694076 | 0.008518052 |
| 9867 | *PJA2* | ENSG00000198961.10 | 7.77 | 8.12 | 0.35 | 1.27 | 3.99566E-05 | 0.000502671 |
| 7114 | *TMSB4X* | ENSG00000205542.11 | 6.52 | 6.87 | 0.35 | 1.27 | 0.001306698 | 0.007026643 |
| 4154 | *MBNL1* | ENSG00000152601.18 | 7.23 | 7.58 | 0.35 | 1.27 | 8.28999E-05 | 0.000858692 |
| 23198 | *PSME4* | ENSG00000068878.15 | 6.96 | 7.31 | 0.35 | 1.27 | 0.000590447 | 0.003813264 |
| 9802 | *DAZAP2* | ENSG00000183283.16 | 8.86 | 9.20 | 0.35 | 1.27 | 3.91682E-05 | 0.000494845 |
| 6845 | *VAMP7* | ENSG00000124333.16 | 6.04 | 6.38 | 0.35 | 1.27 | 0.000159943 | 0.001382027 |
| 6845 | *VAMP7* | SG00000124333.16_PAR | 6.04 | 6.38 | 0.35 | 1.27 | 0.000159943 | 0.001382027 |
| 23216 | *TBC1D1* | ENSG00000065882.17 | 6.21 | 6.56 | 0.35 | 1.27 | 0.001383636 | 0.007317475 |
| 54887 | *UHRF1BP1* | ENSG00000065060.18 | 6.07 | 6.42 | 0.35 | 1.27 | 9.80773E-05 | 0.000968448 |
| 348235 | *SKA2* | ENSG00000182628.13 | 6.94 | 7.29 | 0.35 | 1.27 | 0.000122297 | 0.001130967 |
| 26001 | *RNF167* | ENSG00000108523.16 | 6.68 | 7.02 | 0.35 | 1.27 | 0.000158687 | 0.001377521 |
| 369 | *ARAF* | ENSG00000078061.14 | 5.43 | 5.77 | 0.35 | 1.27 | 0.001439773 | 0.007545236 |
| 9792 | *SERTAD2* | ENSG00000179833.4 | 5.44 | 5.78 | 0.35 | 1.27 | 0.001834315 | 0.009076854 |
| 55884 | *WSB2* | ENSG00000176871.9 | 6.96 | 7.30 | 0.35 | 1.27 | 0.000571493 | 0.003714831 |
| 9444 | *QKI* | ENSG00000112531.17 | 7.96 | 8.30 | 0.35 | 1.27 | 0.000535311 | 0.003522352 |
| 10944 | *C11orf58* | ENSG00000110696.10 | 6.81 | 7.16 | 0.35 | 1.27 | 0.001151187 | 0.006367795 |
| 9921 | *RNF10* | ENSG00000022840.17 | 7.61 | 7.95 | 0.34 | 1.27 | 0.000949614 | 0.005458736 |
| 8087 | *FXR1* | ENSG00000114416.18 | 7.49 | 7.83 | 0.34 | 1.27 | 4.99661E-05 | 0.000588189 |
| 6745 | *SSR1* | ENSG00000124783.14 | 8.20 | 8.55 | 0.34 | 1.27 | 4.33168E-05 | 0.000529237 |
| 84858 | *ZNF503* | ENSG00000165655.17 | 6.13 | 6.47 | 0.34 | 1.27 | 0.000201247 | 0.001648753 |
| 5048 | *PAFAH1B1* | ENSG00000007168.14 | 7.90 | 8.24 | 0.34 | 1.27 | 0.001258529 | 0.006816765 |
| 9655 | *SOCS5* | ENSG00000171150.9 | 5.08 | 5.42 | 0.34 | 1.27 | 0.000621528 | 0.003956112 |
| 4170 | *MCL1* | ENSG00000143384.14 | 8.50 | 8.84 | 0.34 | 1.27 | 1.63705E-05 | 0.00026111 |
| 2109 | *ETFB* | ENSG00000105379.10 | 5.88 | 6.22 | 0.34 | 1.27 | 0.001321898 | 0.007078263 |
| 6049 | *RNF6* | ENSG00000127870.17 | 5.76 | 6.10 | 0.34 | 1.27 | 0.00120978 | 0.006602758 |
| 4478 | *MSN* | ENSG00000147065.17 | 7.86 | 8.20 | 0.34 | 1.27 | 0.000325912 | 0.002394327 |
| 5885 | *RAD21* | ENSG00000164754.16 | 9.10 | 9.44 | 0.34 | 1.27 | 4.06428E-05 | 0.000507494 |
| 5781 | *PTPN11* | ENSG00000179295.19 | 7.85 | 8.19 | 0.34 | 1.27 | 4.39836E-05 | 0.000534534 |
| 64282 | *TENT4B* | ENSG00000121274.14 | 6.37 | 6.71 | 0.34 | 1.27 | 0.001679167 | 0.008457056 |
| 1027 | *CDKN1B* | ENSG00000111276.12 | 6.93 | 7.27 | 0.34 | 1.27 | 0.000164016 | 0.001405895 |
| 1453 | *CSNK1D* | ENSG00000141551.15 | 7.03 | 7.37 | 0.34 | 1.27 | 0.001975256 | 0.009614733 |
| 148867 | *SLC30A7* | ENSG00000162695.12 | 5.76 | 6.10 | 0.34 | 1.27 | 0.00170164 | 0.008547493 |
| 10169 | *SERF2* | ENSG00000140264.21 | 8.46 | 8.80 | 0.34 | 1.27 | 0.000199993 | 0.001642843 |
| 81610 | *FAM83D* | ENSG00000101447.15 | 6.55 | 6.89 | 0.34 | 1.26 | 0.000784972 | 0.004722104 |
| 81563 | *C1orf21* | ENSG00000116667.15 | 5.95 | 6.29 | 0.34 | 1.26 | 0.001810202 | 0.008978626 |
| 113178 | *SCAMP4* | ENSG00000227500.10 | 5.90 | 6.23 | 0.34 | 1.26 | 0.000660748 | 0.004121845 |
| 1398 | *CRK* | ENSG00000167193.8 | 6.62 | 6.96 | 0.34 | 1.26 | 0.00010283 | 0.001006037 |
| 4247 | *MGAT2* | ENSG00000168282.6 | 6.66 | 7.00 | 0.34 | 1.26 | 8.88801E-05 | 0.000899529 |
| 3842 | *TNPO1* | ENSG00000083312.19 | 8.19 | 8.53 | 0.34 | 1.26 | 9.66355E-05 | 0.00095673 |
| 53834 | *FGFRL1* | ENSG00000127418.15 | 7.25 | 7.59 | 0.34 | 1.26 | 0.000122004 | 0.001129647 |
| 26097 | *CHTOP* | ENSG00000160679.13 | 6.96 | 7.30 | 0.34 | 1.26 | 0.000541732 | 0.003556806 |
| 10079 | *ATP9A* | ENSG00000054793.14 | 6.98 | 7.32 | 0.34 | 1.26 | 0.000255179 | 0.001983419 |
| 4102 | *MAGEA3* | ENSG00000221867.9 | 8.33 | 8.67 | 0.34 | 1.26 | 0.000107358 | 0.001036226 |
| 55754 | *TMEM30A* | ENSG00000112697.17 | 8.43 | 8.77 | 0.34 | 1.26 | 0.000145889 | 0.001294118 |
| 60592 | *SCOC* | ENSG00000153130.18 | 7.89 | 8.23 | 0.34 | 1.26 | 7.82539E-05 | 0.000824219 |
| 158405 | *KIAA1958* | ENSG00000165185.15 | 6.01 | 6.35 | 0.34 | 1.26 | 0.001651317 | 0.008353426 |
| 5518 | *PPP2R1A* | ENSG00000105568.18 | 9.49 | 9.83 | 0.34 | 1.26 | 4.94566E-05 | 0.000584023 |
| 5204 | *PFDN5* | ENSG00000123349.15 | 6.94 | 7.27 | 0.34 | 1.26 | 0.00043474 | 0.003005534 |
| 3915 | *LAMC1* | ENSG00000135862.6 | 9.18 | 9.51 | 0.34 | 1.26 | 0.001119705 | 0.006242812 |
| 871 | *SERPINH1* | ENSG00000149257.16 | 7.02 | 7.36 | 0.34 | 1.26 | 0.000304925 | 0.002287165 |
| 81608 | *FIP1L1* | ENSG00000145216.16 | 6.00 | 6.33 | 0.34 | 1.26 | 0.000222669 | 0.001787183 |
| 283638 | *CEP170B* | ENSG00000099814.17 | 5.85 | 6.19 | 0.34 | 1.26 | 0.000728115 | 0.004444193 |
| 9263 | *STK17A* | ENSG00000164543.7 | 5.60 | 5.94 | 0.34 | 1.26 | 0.000298981 | 0.00225158 |
| 2632 | *GBE1* | ENSG00000114480.13 | 7.54 | 7.88 | 0.33 | 1.26 | 9.22052E-05 | 0.000924662 |
| 51454 | *GULP1* | ENSG00000144366.16 | 5.34 | 5.67 | 0.33 | 1.26 | 0.000359084 | 0.002587371 |
| 1176 | *AP3S1* | ENSG00000177879.17 | 5.45 | 5.79 | 0.33 | 1.26 | 0.00090906 | 0.005274096 |
| 7171 | *TPM4* | ENSG00000167460.17 | 8.76 | 9.09 | 0.33 | 1.26 | 1.64486E-05 | 0.0002618 |
| 57231 | *SNX14* | ENSG00000135317.14 | 6.41 | 6.75 | 0.33 | 1.26 | 0.000851157 | 0.005007844 |
| 9341 | *VAMP3* | ENSG00000049245.13 | 6.03 | 6.37 | 0.33 | 1.26 | 0.000449332 | 0.003082277 |
| 9246 | *UBE2L6* | ENSG00000156587.16 | 6.07 | 6.40 | 0.33 | 1.26 | 0.00059631 | 0.003837918 |
| 6944 | *VPS72* | ENSG00000163159.15 | 6.20 | 6.53 | 0.33 | 1.26 | 7.80503E-05 | 0.000822653 |
| 55773 | *TBC1D23* | ENSG00000036054.13 | 6.13 | 6.46 | 0.33 | 1.26 | 0.001560616 | 0.008018927 |
| 51119 | *SBDS* | ENSG00000126524.11 | 6.48 | 6.81 | 0.33 | 1.26 | 0.000340017 | 0.002474937 |
| 51696 | *HECA* | ENSG00000112406.5 | 5.60 | 5.93 | 0.33 | 1.26 | 0.000618085 | 0.003945914 |
| 9133 | *CCNB2* | ENSG00000157456.8 | 7.07 | 7.40 | 0.33 | 1.26 | 0.000124545 | 0.001146393 |
| 7905 | *REEP5* | ENSG00000129625.13 | 6.72 | 7.05 | 0.33 | 1.26 | 0.000199511 | 0.001640797 |
| 10491 | *CRTAP* | ENSG00000170275.15 | 8.14 | 8.47 | 0.33 | 1.26 | 0.000176503 | 0.001490993 |
| 83540 | *NUF2* | ENSG00000143228.13 | 6.18 | 6.51 | 0.33 | 1.26 | 0.000233661 | 0.001859497 |
| 10057 | *ABCC5* | ENSG00000114770.17 | 6.47 | 6.80 | 0.33 | 1.26 | 0.001175529 | 0.006460458 |
| 80204 | *FBXO11* | ENSG00000138081.22 | 6.19 | 6.52 | 0.33 | 1.26 | 9.37883E-05 | 0.000936573 |
| 9040 | *UBE2M* | ENSG00000130725.8 | 6.24 | 6.57 | 0.33 | 1.26 | 0.000674665 | 0.004186624 |
| 55030 | *FBXO34* | ENSG00000178974.11 | 5.76 | 6.09 | 0.33 | 1.26 | 0.00180913 | 0.008978626 |
| 6223 | *RPS19* | ENSG00000105372.8 | 7.79 | 8.12 | 0.33 | 1.26 | 0.000639083 | 0.004031943 |

| | | | | | | | | |
|---|---|---|---|---|---|---|---|---|
| 7086 | *TKT* | ENSG00000163931.17 | 8.81 | 9.14 | 0.33 | 1.26 | 0.000153457 | 0.001341087 |
| 387 | *RHOA* | ENSG00000067560.14 | 9.17 | 9.50 | 0.33 | 1.26 | 3.4083E-05 | 0.000445135 |
| 10613 | *ERLIN1* | ENSG00000107566.14 | 6.54 | 6.87 | 0.33 | 1.26 | 0.000492436 | 0.003301017 |
| 4893 | *NRAS* | ENSG00000213281.5 | 7.26 | 7.59 | 0.33 | 1.26 | 0.00031116 | 0.002320019 |
| 523 | *ATP6V1A* | ENSG00000114573.10 | 8.15 | 8.48 | 0.33 | 1.26 | 6.22797E-05 | 0.000692925 |
| 83638 | *C11orf68* | ENSG00000175573.7 | 4.65 | 4.97 | 0.33 | 1.26 | 0.002068891 | 0.009959759 |
| 113201 | *GOLM2* | ENSG00000166734.20 | 8.46 | 8.79 | 0.33 | 1.26 | 6.02557E-05 | 0.000677046 |
| 1540 | *CYLD* | ENSG00000083799.18 | 6.83 | 7.15 | 0.33 | 1.25 | 0.001321736 | 0.007078263 |
| 5547 | *PRCP* | ENSG00000137509.12 | 7.98 | 8.31 | 0.33 | 1.25 | 0.000161588 | 0.001390984 |
| 9403 | *SELENOF* | ENSG00000183291.18 | 7.80 | 8.13 | 0.33 | 1.25 | 0.000109138 | 0.00104668 |
| 22818 | *COPZ1* | ENSG00000111481.10 | 7.80 | 8.12 | 0.33 | 1.25 | 0.000603286 | 0.003871193 |
| 23338 | *JADE2* | ENSG00000043143.22 | 6.23 | 6.55 | 0.33 | 1.25 | 0.001484286 | 0.007727689 |
| 2286 | *FKBP2* | ENSG00000173486.14 | 5.33 | 5.65 | 0.33 | 1.25 | 0.000721234 | 0.004409371 |
| 219333 | *USP12* | ENSG00000152484.14 | 5.76 | 6.09 | 0.32 | 1.25 | 0.001132635 | 0.006300862 |
| 1314 | *COPA* | ENSG00000122218.16 | 8.10 | 8.43 | 0.32 | 1.25 | 0.001067904 | 0.005998567 |
| 55969 | *RAB5IF* | ENSG00000101084.19 | 6.57 | 6.90 | 0.32 | 1.25 | 0.000575968 | 0.003735828 |
| 10444 | *ZER1* | ENSG00000160445.11 | 7.35 | 7.67 | 0.32 | 1.25 | 6.94194E-05 | 0.000753374 |
| 55830 | *GLT8D1* | ENSG00000016864.19 | 7.18 | 7.51 | 0.32 | 1.25 | 0.000363935 | 0.002609792 |
| 23291 | *FBXW11* | ENSG00000072803.18 | 6.81 | 7.14 | 0.32 | 1.25 | 0.000318763 | 0.002361457 |
| 60559 | *SPCS3* | ENSG00000129128.13 | 6.37 | 6.69 | 0.32 | 1.25 | 0.001115261 | 0.006224972 |
| 10024 | *TROAP* | ENSG00000135451.13 | 5.99 | 6.32 | 0.32 | 1.25 | 0.001785539 | 0.008882717 |
| 49854 | *ZBTB21* | ENSG00000173276.14 | 6.01 | 6.33 | 0.32 | 1.25 | 0.001369895 | 0.007260152 |
| 3490 | *IGFBP7* | ENSG00000163453.11 | 5.63 | 5.95 | 0.32 | 1.25 | 0.001282714 | 0.006915324 |
| 2040 | *STOM* | ENSG00000148175.13 | 7.62 | 7.94 | 0.32 | 1.25 | 4.08909E-05 | 0.00050932 |
| 23171 | *GPD1L* | ENSG00000152642.11 | 7.07 | 7.39 | 0.32 | 1.25 | 0.000115113 | 0.00108594 |
| 10430 | *TMEM147* | ENSG00000105677.12 | 6.46 | 6.78 | 0.32 | 1.25 | 0.00065159 | 0.004086804 |
| 54492 | *NEURL1B* | ENSG00000214357.9 | 7.58 | 7.90 | 0.32 | 1.25 | 0.000114541 | 0.001082516 |
| 80790 | *CMIP* | ENSG00000153815.18 | 6.15 | 6.47 | 0.32 | 1.25 | 0.000906211 | 0.005261621 |
| 10473 | *HMGN4* | ENSG00000182952.5 | 6.93 | 7.25 | 0.32 | 1.25 | 0.001348299 | 0.007181705 |
| 2888 | *GRB14* | ENSG00000115290.10 | 6.35 | 6.67 | 0.32 | 1.25 | 0.00205238 | 0.009901694 |
| 83461 | *CDCA3* | ENSG00000111665.12 | 5.81 | 6.13 | 0.32 | 1.25 | 0.000425176 | 0.002959866 |
| 7844 | *RNF103* | ENSG00000239305.7 | 5.65 | 5.98 | 0.32 | 1.25 | 0.001503387 | 0.007795559 |
| 65979 | *PHACTR4* | ENSG00000204138.13 | 6.31 | 6.63 | 0.32 | 1.25 | 0.000924788 | 0.005346744 |
| 64793 | *CEP85* | ENSG00000130695.16 | 5.73 | 6.05 | 0.32 | 1.25 | 0.001978428 | 0.009622238 |
| 25949 | *SYF2* | ENSG00000117614.10 | 5.96 | 6.28 | 0.32 | 1.25 | 0.001152988 | 0.006369229 |
| 54502 | *RBM47* | ENSG00000163694.15 | 7.67 | 7.99 | 0.32 | 1.25 | 5.35216E-05 | 0.000618879 |
| 51324 | *SPG21* | ENSG00000090487.11 | 6.50 | 6.82 | 0.32 | 1.25 | 0.00024764 | 0.001936859 |
| 26959 | *HBP1* | ENSG00000105856.14 | 6.64 | 6.96 | 0.32 | 1.25 | 0.000237841 | 0.001881789 |
| 9255 | *AIMP1* | ENSG00000164022.18 | 5.71 | 6.03 | 0.32 | 1.25 | 0.000814074 | 0.004833244 |
| 537 | *ATP6AP1* | ENSG00000071553.18 | 8.42 | 8.74 | 0.32 | 1.25 | 6.89722E-05 | 0.000749109 |
| 80273 | *GRPEL1* | ENSG00000109519.13 | 6.25 | 6.56 | 0.32 | 1.25 | 0.000171117 | 0.001457603 |
| 4134 | *MAP4* | ENSG00000047849.22 | 7.72 | 8.04 | 0.32 | 1.25 | 0.000115459 | 0.001088524 |
| 26127 | *FGFR1OP2* | ENSG00000111790.14 | 6.10 | 6.41 | 0.32 | 1.24 | 0.0007202 | 0.004404839 |
| 7088 | *TLE1* | ENSG00000196781.16 | 7.58 | 7.89 | 0.32 | 1.24 | 0.000122597 | 0.001131911 |
| 26470 | *SEZ6L2* | ENSG00000174938.15 | 6.96 | 7.28 | 0.32 | 1.24 | 0.000217387 | 0.001752286 |
| 91369 | *ANKRD40* | ENSG00000154945.7 | 6.23 | 6.55 | 0.31 | 1.24 | 0.000152179 | 0.001335728 |
| 26586 | *CKAP2* | ENSG00000136108.15 | 7.19 | 7.50 | 0.31 | 1.24 | 5.53939E-05 | 0.000635632 |
| 2319 | *FLOT2* | ENSG00000132589.16 | 6.62 | 6.94 | 0.31 | 1.24 | 0.001358674 | 0.007215973 |
| 139322 | *APOOL* | ENSG00000155008.16 | 6.35 | 6.67 | 0.31 | 1.24 | 0.000384027 | 0.002725229 |
| 64782 | *AEN* | ENSG00000181026.15 | 5.09 | 5.41 | 0.31 | 1.24 | 0.001255459 | 0.006802595 |
| 6856 | *SYPL1* | ENSG00000008282.9 | 7.72 | 8.03 | 0.31 | 1.24 | 0.000145728 | 0.001293456 |
| 51726 | *DNAJB11* | ENSG00000090520.12 | 6.03 | 6.35 | 0.31 | 1.24 | 0.000549153 | 0.003592952 |
| 25932 | *CLIC4* | ENSG00000169504.15 | 8.08 | 8.39 | 0.31 | 1.24 | 0.000208341 | 0.001691178 |
| 253782 | *CERS6* | ENSG00000172292.15 | 6.29 | 6.61 | 0.31 | 1.24 | 0.000267778 | 0.002063184 |
| 29956 | *CERS2* | ENSG00000143418.20 | 8.04 | 8.35 | 0.31 | 1.24 | 2.83707E-05 | 0.000388164 |
| 2631 | *NIPSNAP2* | ENSG00000146729.10 | 6.92 | 7.23 | 0.31 | 1.24 | 0.00011328 | 0.001076765 |
| 51203 | *NUSAP1* | ENSG00000137804.14 | 8.49 | 8.80 | 0.31 | 1.24 | 0.00066921 | 0.004160802 |
| 567 | *B2M* | ENSG00000166710.22 | 10.17 | 10.48 | 0.31 | 1.24 | 0.000241949 | 0.001904254 |
| 3073 | *HEXA* | ENSG00000213614.11 | 7.43 | 7.74 | 0.31 | 1.24 | 0.000339792 | 0.002474499 |
| 1859 | *DYRK1A* | ENSG00000157540.22 | 6.54 | 6.85 | 0.31 | 1.24 | 0.001243197 | 0.006748335 |
| 5243 | *ABCB1* | ENSG00000085563.15 | 9.29 | 9.60 | 0.31 | 1.24 | 0.000211592 | 0.001713861 |
| 342371 | *ATXN1L* | ENSG00000224470.9 | 6.74 | 7.05 | 0.31 | 1.24 | 0.001377505 | 0.007292761 |
| 10787 | *NCKAP1* | ENSG00000061676.15 | 8.32 | 8.63 | 0.31 | 1.24 | 0.000150275 | 0.001322086 |
| 10527 | *IPO7* | ENSG00000205339.10 | 7.57 | 7.88 | 0.31 | 1.24 | 0.000217572 | 0.001752841 |
| 4694 | *NDUFA1* | ENSG00000125356.7 | 5.22 | 5.53 | 0.31 | 1.24 | 0.001531332 | 0.007917247 |
| 57092 | *PCNP* | ENSG00000081154.12 | 7.95 | 8.26 | 0.31 | 1.24 | 0.00010263 | 0.00100547 |
| 7076 | *TIMP1* | ENSG00000102265.12 | 6.30 | 6.61 | 0.31 | 1.24 | 0.002069061 | 0.009959759 |
| 10594 | *PRPF8* | ENSG00000174231.17 | 9.46 | 9.77 | 0.31 | 1.24 | 0.001438282 | 0.007540056 |
| 64853 | *AIDA* | ENSG00000186063.13 | 6.99 | 7.30 | 0.31 | 1.24 | 0.000134463 | 0.001216484 |
| 27069 | *GHITM* | ENSG00000165678.22 | 8.52 | 8.83 | 0.31 | 1.24 | 0.000133851 | 0.001211679 |
| 10434 | *LYPLA1* | ENSG00000120992.18 | 6.67 | 6.98 | 0.31 | 1.24 | 0.000765886 | 0.004636593 |
| 10682 | *EBP* | ENSG00000147155.11 | 6.73 | 7.04 | 0.31 | 1.24 | 0.000809793 | 0.004815443 |
| 51586 | *MED15* | ENSG00000099917.17 | 7.11 | 7.42 | 0.31 | 1.24 | 0.000859531 | 0.00503344 |
| 3836 | *KPNA1* | ENSG00000114030.13 | 7.59 | 7.90 | 0.31 | 1.24 | 0.001511929 | 0.007828082 |
| 57019 | *CIAPIN1* | ENSG00000005194.15 | 6.33 | 6.64 | 0.31 | 1.24 | 0.001975604 | 0.009614733 |
| 8887 | *TAX1BP1* | ENSG00000106052.14 | 7.98 | 8.29 | 0.31 | 1.24 | 0.000113574 | 0.001077431 |
| 60343 | *FAM3A* | ENSG00000071889.17 | 5.61 | 5.92 | 0.31 | 1.24 | 0.001719455 | 0.008608172 |
| 10130 | *PDIA6* | ENSG00000143870.13 | 8.20 | 8.50 | 0.30 | 1.24 | 0.000136056 | 0.001226462 |
| 3313 | *HSPA9* | ENSG00000113013.16 | 9.24 | 9.55 | 0.30 | 1.24 | 6.8477E-05 | 0.000746385 |
| 2885 | *GRB2* | ENSG00000177885.15 | 7.11 | 7.42 | 0.30 | 1.24 | 0.000329821 | 0.002412421 |
| 79646 | *PANK3* | ENSG00000120137.7 | 7.64 | 7.94 | 0.30 | 1.23 | 0.000397778 | 0.002805491 |
| 1725 | *DHPS* | ENSG00000095059.17 | 5.54 | 5.85 | 0.30 | 1.23 | 0.001278204 | 0.006903407 |
| 84726 | *PRRC2B* | ENSG00000288701.1 | 8.81 | 9.11 | 0.30 | 1.23 | 0.000950247 | 0.005460283 |
| 83636 | *C19orf12* | ENSG00000131943.20 | 5.50 | 5.80 | 0.30 | 1.23 | 0.00078129 | 0.00470561 |
| 146223 | *CMTM4* | ENSG00000183723.13 | 7.20 | 7.50 | 0.30 | 1.23 | 0.000147162 | 0.001302788 |
| 26123 | *TCTN3* | ENSG00000119977.22 | 6.17 | 6.47 | 0.30 | 1.23 | 0.000588342 | 0.003804576 |
| 81624 | *DIAPH3* | ENSG00000139734.19 | 5.97 | 6.27 | 0.30 | 1.23 | 0.000494846 | 0.003312694 |

| | | | | | | | | |
|---|---|---|---|---|---|---|---|---|
| 6233 | *RPS27A* | ENSG00000143947.15 | 8.08 | 8.39 | 0.30 | 1.23 | 0.001457403 | 0.007624315 |
| 10632 | *ATP5MG* | ENSG00000167283.9 | 7.00 | 7.31 | 0.30 | 1.23 | 0.000343817 | 0.002498959 |
| 5869 | *RAB5B* | ENSG00000111540.16 | 8.42 | 8.72 | 0.30 | 1.23 | 0.000612301 | 0.003920659 |
| 5104 | *SERPINA5* | ENSG00000188488.14 | 7.95 | 8.25 | 0.30 | 1.23 | 0.000159016 | 0.001378778 |
| 11171 | *STRAP* | ENSG00000023734.11 | 7.70 | 8.01 | 0.30 | 1.23 | 0.000111404 | 0.001062846 |
| 7534 | *YWHAZ* | ENSG00000164924.18 | 9.25 | 9.55 | 0.30 | 1.23 | 7.49856E-05 | 0.000796892 |
| 8731 | *RNMT* | ENSG00000101654.18 | 6.30 | 6.60 | 0.30 | 1.23 | 0.000798884 | 0.004773271 |
| 9686 | *VGLL4* | ENSG00000144560.16 | 6.69 | 6.99 | 0.30 | 1.23 | 0.001144061 | 0.006344188 |
| 283820 | *NOMO2* | ENSG00000185164.15 | 8.26 | 8.56 | 0.30 | 1.23 | 7.44477E-05 | 0.000793035 |
| 6416 | *MAP2K4* | ENSG00000065559.15 | 6.00 | 6.30 | 0.30 | 1.23 | 0.001429742 | 0.007511974 |
| 11228 | *RASSF8* | ENSG00000123094.16 | 6.64 | 6.94 | 0.30 | 1.23 | 0.000857274 | 0.005022179 |
| 682 | *BSG* | ENSG00000172270.22 | 8.93 | 9.23 | 0.30 | 1.23 | 0.00048707 | 0.003273815 |
| 11135 | *CDC42EP1* | ENSG00000128283.7 | 5.50 | 5.80 | 0.30 | 1.23 | 0.001842848 | 0.009104444 |
| 48 | *ACO1* | ENSG00000122729.19 | 6.50 | 6.80 | 0.30 | 1.23 | 0.000896443 | 0.005215002 |
| 138050 | *HGSNAT* | ENSG00000165102.15 | 6.10 | 6.40 | 0.30 | 1.23 | 0.001277331 | 0.006901175 |
| 27173 | *SLC39A1* | ENSG00000143570.19 | 6.83 | 7.12 | 0.30 | 1.23 | 0.000147098 | 0.001302788 |
| 2923 | *PDIA3* | ENSG00000167004.14 | 9.72 | 10.02 | 0.30 | 1.23 | 4.85252E-05 | 0.000577111 |
| 51142 | *CHCHD2* | ENSG00000106153.13 | 8.46 | 8.76 | 0.30 | 1.23 | 0.00041879 | 0.002921964 |
| 29889 | *GNL2* | ENSG00000134697.13 | 6.63 | 6.93 | 0.30 | 1.23 | 0.000729685 | 0.004450156 |
| 8803 | *SUCLA2* | ENSG00000136143.16 | 5.90 | 6.19 | 0.30 | 1.23 | 0.000322644 | 0.002380807 |
| 57179 | *KIAA1191* | ENSG00000122203.15 | 7.41 | 7.71 | 0.29 | 1.23 | 9.5542E-05 | 0.000950291 |
| 166647 | *ADGRA3* | ENSG00000152990.14 | 6.90 | 7.19 | 0.29 | 1.23 | 0.001889146 | 0.009284893 |
| 55500 | *ETNK1* | ENSG00000139163.16 | 7.96 | 8.25 | 0.29 | 1.23 | 0.002004874 | 0.009728791 |
| 9230 | *RAB11B* | ENSG00000185236.12 | 6.44 | 6.73 | 0.29 | 1.23 | 0.000657821 | 0.004112132 |
| 6730 | *SRP68* | ENSG00000167881.15 | 7.14 | 7.43 | 0.29 | 1.22 | 0.000510609 | 0.003395538 |
| 4055 | *LTBR* | ENSG00000111321.11 | 6.13 | 6.42 | 0.29 | 1.22 | 0.000991508 | 0.005648336 |
| 6169 | *RPL38* | ENSG00000172809.13 | 7.00 | 7.30 | 0.29 | 1.22 | 0.001663629 | 0.008403885 |
| 56947 | *MFF* | ENSG00000168958.20 | 6.06 | 6.35 | 0.29 | 1.22 | 0.000874601 | 0.005103769 |
| 7763 | *ZFAND5* | ENSG00000107372.13 | 8.45 | 8.74 | 0.29 | 1.22 | 0.00029703 | 0.002241431 |
| 2799 | *GNS* | ENSG00000135677.11 | 9.77 | 10.06 | 0.29 | 1.22 | 0.000118152 | 0.001103515 |
| 51526 | *OSER1* | ENSG00000132823.11 | 5.75 | 6.04 | 0.29 | 1.22 | 0.00073202 | 0.004460777 |
| 26092 | *TOR1AIP1* | ENSG00000143337.19 | 7.72 | 8.01 | 0.29 | 1.22 | 0.000518487 | 0.003434234 |
| 55967 | *NDUFA12* | ENSG00000184752.14 | 5.82 | 6.11 | 0.29 | 1.22 | 0.000792578 | 0.004754518 |
| 27075 | *TSPAN13* | ENSG00000106537.8 | 8.57 | 8.86 | 0.29 | 1.22 | 0.001023213 | 0.00579525 |
| 51246 | *SHISA5* | ENSG00000164054.17 | 6.40 | 6.69 | 0.29 | 1.22 | 0.001625752 | 0.008268692 |
| 23376 | *UFL1* | ENSG00000014123.10 | 6.43 | 6.72 | 0.29 | 1.22 | 0.000450833 | 0.003088341 |
| 9554 | *SEC22B* | ENSG00000265808.4 | 7.07 | 7.36 | 0.29 | 1.22 | 0.000599113 | 0.003852653 |
| 5440 | *POLR2K* | ENSG00000147669.11 | 6.22 | 6.50 | 0.29 | 1.22 | 0.001596654 | 0.008151083 |
| 8836 | *GGH* | ENSG00000137563.13 | 6.56 | 6.85 | 0.29 | 1.22 | 0.000688228 | 0.004254374 |
| 51719 | *CAB39* | ENSG00000135932.12 | 6.12 | 6.40 | 0.29 | 1.22 | 0.000541982 | 0.003556888 |
| 11057 | *ABHD2* | ENSG00000140526.18 | 9.50 | 9.79 | 0.29 | 1.22 | 0.000250365 | 0.001953243 |
| 90411 | *MCFD2* | ENSG00000180398.13 | 8.62 | 8.91 | 0.29 | 1.22 | 0.00019144 | 0.001583968 |
| 27257 | *LSM1* | ENSG00000175324.10 | 4.93 | 5.22 | 0.28 | 1.22 | 0.001871608 | 0.009213175 |
| 63905 | *MANBAL* | ENSG00000101363.13 | 5.60 | 5.89 | 0.28 | 1.22 | 0.001640113 | 0.008319181 |
| 23588 | *KLHDC2* | ENSG00000165516.11 | 5.97 | 6.26 | 0.28 | 1.22 | 0.001456747 | 0.007623542 |
| 25793 | *FBXO7* | ENSG00000100225.18 | 7.30 | 7.58 | 0.28 | 1.22 | 0.000309005 | 0.002309249 |
| 644 | *BLVRA* | ENSG00000106605.11 | 6.66 | 6.95 | 0.28 | 1.22 | 0.001542447 | 0.007959958 |
| 51247 | *PAIP2* | ENSG00000120727.13 | 7.37 | 7.66 | 0.28 | 1.22 | 0.000312331 | 0.002325286 |
| 444 | *ASPH* | ENSG00000198363.18 | 8.92 | 9.20 | 0.28 | 1.22 | 0.000186131 | 0.001548585 |
| 26035 | *GLCE* | ENSG00000138604.10 | 7.28 | 7.57 | 0.28 | 1.22 | 0.000226678 | 0.00181065 |
| 10857 | *PGRMC1* | ENSG00000101856.10 | 7.85 | 8.13 | 0.28 | 1.22 | 0.000795439 | 0.004764066 |
| 5660 | *PSAP* | ENSG00000197746.15 | 10.67 | 10.95 | 0.28 | 1.22 | 0.000714389 | 0.004383589 |
| 490 | *ATP2B1* | ENSG00000070961.16 | 6.75 | 7.04 | 0.28 | 1.22 | 0.000961579 | 0.005502228 |
| 5217 | *PFN2* | ENSG00000070087.15 | 7.53 | 7.81 | 0.28 | 1.22 | 0.000323483 | 0.002383096 |
| 81631 | *MAP1LC3B* | ENSG00000140941.14 | 6.78 | 7.06 | 0.28 | 1.21 | 0.000344487 | 0.002502619 |
| 387066 | *SNHG5* | ENSG00000203875.13 | 5.85 | 6.13 | 0.28 | 1.21 | 0.001810102 | 0.008978626 |
| 6625 | *SNRNP70* | ENSG00000104852.15 | 7.06 | 7.34 | 0.28 | 1.21 | 0.001593245 | 0.008136445 |
| 152006 | *RNF38* | ENSG00000137075.18 | 7.59 | 7.87 | 0.28 | 1.21 | 0.000906073 | 0.005261621 |
| 80213 | *TM2D3* | ENSG00000184277.13 | 4.73 | 5.01 | 0.28 | 1.21 | 0.002078835 | 0.009993992 |
| 51449 | *PCYOX1* | ENSG00000116005.12 | 7.43 | 7.71 | 0.28 | 1.21 | 0.000237489 | 0.001880985 |
| 27043 | *PELP1* | ENSG00000141456.16 | 6.70 | 6.97 | 0.28 | 1.21 | 0.001041291 | 0.005881327 |
| 5528 | *PPP2R5D* | ENSG00000112640.16 | 6.78 | 7.06 | 0.28 | 1.21 | 0.000435629 | 0.003007525 |
| 2230 | *FDX1* | ENSG00000137714.3 | 7.22 | 7.50 | 0.28 | 1.21 | 0.000465752 | 0.003158824 |
| 5886 | *RAD23A* | ENSG00000179262.10 | 7.03 | 7.31 | 0.28 | 1.21 | 0.000443943 | 0.003049494 |
| 57414 | *RHBDD2* | ENSG00000005486.17 | 7.24 | 7.52 | 0.28 | 1.21 | 0.001647614 | 0.008342111 |
| 10906 | *TRAFD1* | ENSG00000135148.12 | 5.82 | 6.10 | 0.27 | 1.21 | 0.001579972 | 0.008087925 |
| 55361 | *PI4K2A* | ENSG00000155252.13 | 6.35 | 6.62 | 0.27 | 1.21 | 0.001004739 | 0.005710498 |
| 3658 | *IREB2* | ENSG00000136381.13 | 7.71 | 7.98 | 0.27 | 1.21 | 0.001144815 | 0.006345098 |
| 10605 | *PAIP1* | ENSG00000172239.14 | 7.38 | 7.66 | 0.27 | 1.21 | 0.000423169 | 0.002947687 |
| 9530 | *BAG4* | ENSG00000156735.11 | 5.71 | 5.98 | 0.27 | 1.21 | 0.001137808 | 0.006320266 |
| 10564 | *ARFGEF2* | ENSG00000124198.10 | 6.89 | 7.16 | 0.27 | 1.21 | 0.000837198 | 0.004940133 |
| 3423 | *IDS* | ENSG00000010404.18 | 8.41 | 8.68 | 0.27 | 1.21 | 0.000984653 | 0.005625681 |
| 3094 | *HINT1* | ENSG00000169567.13 | 7.37 | 7.64 | 0.27 | 1.21 | 0.000258277 | 0.002000247 |
| 10302 | *SNAPC5* | ENSG00000174446.13 | 5.41 | 5.68 | 0.27 | 1.21 | 0.000790626 | 0.0047466 |
| 9975 | *NR1D2* | ENSG00000174738.13 | 7.98 | 8.25 | 0.27 | 1.21 | 0.00119926 | 0.006564442 |
| 54878 | *DPP8* | ENSG00000074603.19 | 6.71 | 6.98 | 0.27 | 1.21 | 0.001899275 | 0.009322262 |
| 6139 | *RPL17* | ENSG00000265681.8 | 8.27 | 8.54 | 0.27 | 1.21 | 0.001141299 | 0.006334968 |
| 754 | *PTTG1IP* | ENSG00000183255.12 | 8.47 | 8.74 | 0.27 | 1.20 | 0.000517508 | 0.00343381 |
| 57223 | *PPP4R3B* | ENSG00000275052.5 | 7.15 | 7.42 | 0.27 | 1.20 | 0.000675867 | 0.004188309 |
| 5894 | *RAF1* | ENSG00000132155.14 | 7.68 | 7.95 | 0.27 | 1.20 | 0.000363591 | 0.002608576 |
| 5034 | *P4HB* | ENSG00000185624.16 | 9.38 | 9.65 | 0.27 | 1.20 | 0.000342601 | 0.002491327 |
| 541578 | *EOLA2* | ENSG00000197021.9 | 5.04 | 5.31 | 0.27 | 1.20 | 0.001687762 | 0.008491994 |
| 9741 | *LAPTM4A* | ENSG00000068697.7 | 8.47 | 8.74 | 0.27 | 1.20 | 0.001704432 | 0.008552933 |
| 57459 | *GATAD2B* | ENSG00000143614.10 | 6.83 | 7.09 | 0.27 | 1.20 | 0.00058626 | 0.003792745 |
| 10768 | *AHCYL1* | ENSG00000168710.18 | 7.72 | 7.98 | 0.27 | 1.20 | 0.001158613 | 0.006388548 |
| 7416 | *VDAC1* | ENSG00000213585.11 | 7.91 | 8.17 | 0.26 | 1.20 | 0.000489861 | 0.003288157 |
| 134553 | *C5orf24* | ENSG00000181904.9 | 7.28 | 7.54 | 0.26 | 1.20 | 0.001167047 | 0.006420899 |

| | | | | | | | | |
|---|---|---|---|---|---|---|---|---|
| 25 | *ABL1* | ENSG00000097007.19 | 7.78 | 8.04 | 0.26 | 1.20 | 0.001733393 | 0.00866833 |
| 10899 | *JTB* | ENSG00000143543.15 | 6.26 | 6.53 | 0.26 | 1.20 | 0.000912274 | 0.005288655 |
| 63971 | *KIF13A* | ENSG00000137177.20 | 5.09 | 5.35 | 0.26 | 1.20 | 0.001857251 | 0.009160043 |
| 65056 | *GPBP1* | ENSG00000062194.16 | 7.50 | 7.76 | 0.26 | 1.20 | 0.000613603 | 0.003925649 |
| 6059 | *ABCE1* | ENSG00000164163.11 | 7.40 | 7.66 | 0.26 | 1.20 | 0.000991629 | 0.005648336 |
| 1362 | *CPD* | ENSG00000108582.12 | 7.55 | 7.81 | 0.26 | 1.20 | 0.001534441 | 0.007930585 |
| 5756 | *TWF1* | ENSG00000151239.14 | 8.43 | 8.69 | 0.26 | 1.20 | 0.000597908 | 0.003846552 |
| 51465 | *UBE2J1* | ENSG00000198833.7 | 6.47 | 6.73 | 0.26 | 1.20 | 0.000704995 | 0.004336583 |
| 203547 | *VMA21* | ENSG00000160131.14 | 7.76 | 8.02 | 0.26 | 1.20 | 0.000272857 | 0.002091401 |
| 155066 | *ATP6V0E2* | ENSG00000171130.19 | 6.96 | 7.22 | 0.26 | 1.20 | 0.000867021 | 0.005066443 |
| 114882 | *OSBPL8* | ENSG00000091039.17 | 7.29 | 7.55 | 0.26 | 1.20 | 0.000929694 | 0.005373035 |
| 53339 | *BTBD1* | ENSG00000064726.10 | 6.89 | 7.15 | 0.26 | 1.20 | 0.00063322 | 0.004005055 |
| 25829 | *TMEM184B* | ENSG00000198792.13 | 6.31 | 6.57 | 0.26 | 1.20 | 0.001702834 | 0.008550632 |
| 130074 | *FAM168B* | ENSG00000152102.18 | 8.04 | 8.30 | 0.26 | 1.20 | 0.000322254 | 0.002379099 |
| 51096 | *UTP18* | ENSG00000011260.14 | 5.69 | 5.94 | 0.26 | 1.20 | 0.001057529 | 0.005951427 |
| 60485 | *SAV1* | ENSG00000151748.15 | 6.20 | 6.45 | 0.26 | 1.20 | 0.001247332 | 0.006765888 |
| 7879 | *RAB7A* | ENSG00000075785.14 | 8.04 | 8.29 | 0.26 | 1.19 | 0.000620217 | 0.003951521 |
| 6612 | *SUMO3* | ENSG00000184900.16 | 7.55 | 7.81 | 0.26 | 1.19 | 0.001544467 | 0.007963212 |
| 3191 | *HNRNPL* | ENSG00000104824.18 | 8.61 | 8.87 | 0.26 | 1.19 | 0.00056529 | 0.00368248 |
| 6678 | *SPARC* | ENSG00000113140.11 | 10.84 | 11.10 | 0.26 | 1.19 | 0.001668421 | 0.008420085 |
| 29035 | *C16orf72* | ENSG00000182831.12 | 6.96 | 7.21 | 0.26 | 1.19 | 0.001197072 | 0.006557245 |
| 647979 | *NORAD* | ENSG00000260032.2 | 9.26 | 9.51 | 0.25 | 1.19 | 0.000625976 | 0.00397768 |
| 10313 | *RTN3* | ENSG00000133318.14 | 7.88 | 8.14 | 0.25 | 1.19 | 0.000613038 | 0.003923707 |
| 1386 | *ATF2* | ENSG00000115966.18 | 6.66 | 6.91 | 0.25 | 1.19 | 0.000522104 | 0.003451899 |
| 506 | *ATP5F1B* | ENSG00000110955.9 | 10.26 | 10.52 | 0.25 | 1.19 | 0.000361717 | 0.002598856 |
| 2664 | *GDI1* | ENSG00000203879.12 | 7.93 | 8.19 | 0.25 | 1.19 | 0.000356798 | 0.00257472 |
| 136319 | *MTPN* | ENSG00000105887.11 | 9.11 | 9.36 | 0.25 | 1.19 | 0.001617372 | 0.008236429 |
| 253461 | *ZBTB38* | ENSG00000177311.12 | 6.89 | 7.15 | 0.25 | 1.19 | 0.001765254 | 0.008803251 |
| 3183 | *HNRNPC* | ENSG00000092199.19 | 9.29 | 9.54 | 0.25 | 1.19 | 0.000306913 | 0.002297483 |
| 3607 | *FOXK2* | ENSG00000141568.21 | 6.82 | 7.07 | 0.25 | 1.19 | 0.002029378 | 0.009825465 |
| 5763 | *PTMS* | ENSG00000159335.18 | 8.76 | 9.01 | 0.25 | 1.19 | 0.000946268 | 0.00544367 |
| 133686 | *NADK2* | ENSG00000152620.13 | 7.48 | 7.73 | 0.25 | 1.19 | 0.000614434 | 0.003926728 |
| 3021 | *H3-3B* | ENSG00000132475.10 | 8.47 | 8.72 | 0.25 | 1.19 | 0.001993481 | 0.00968917 |
| 29959 | *NRBP1* | ENSG00000115216.14 | 7.49 | 7.74 | 0.25 | 1.19 | 0.000362485 | 0.002601885 |
| 23071 | *ERP44* | ENSG00000023318.9 | 6.97 | 7.22 | 0.25 | 1.19 | 0.000750294 | 0.004556688 |
| 6729 | *SRP54* | ENSG00000100883.13 | 7.10 | 7.35 | 0.25 | 1.19 | 0.001957412 | 0.009550973 |
| 583 | *BBS2* | ENSG00000125124.14 | 7.27 | 7.52 | 0.25 | 1.19 | 0.000805567 | 0.004805544 |
| 5577 | *PRKAR2B* | ENSG00000005249.13 | 8.36 | 8.61 | 0.25 | 1.19 | 0.00068101 | 0.004214962 |
| 8242 | *KDM5C* | ENSG00000126012.13 | 7.10 | 7.35 | 0.25 | 1.19 | 0.001012849 | 0.005749566 |
| 84193 | *SETD3* | ENSG00000183576.13 | 5.96 | 6.20 | 0.25 | 1.19 | 0.002043656 | 0.009869123 |
| 80351 | *TNKS2* | ENSG00000107854.6 | 6.94 | 7.18 | 0.24 | 1.18 | 0.000599834 | 0.003854177 |
| 93663 | *ARHGAP18* | ENSG00000146376.11 | 7.84 | 8.08 | 0.24 | 1.18 | 0.000951473 | 0.005462685 |
| 140809 | *SRXN1* | ENSG00000271303.2 | 7.14 | 7.38 | 0.24 | 1.18 | 0.00054796 | 0.003586715 |
| 51150 | *SDF4* | ENSG00000078808.19 | 6.13 | 6.38 | 0.24 | 1.18 | 0.001116578 | 0.00623001 |
| 6647 | *SOD1* | ENSG00000142168.15 | 8.02 | 8.26 | 0.24 | 1.18 | 0.001961904 | 0.009566672 |
| 116150 | *NUS1* | ENSG00000153989.8 | 6.86 | 7.10 | 0.24 | 1.18 | 0.000914418 | 0.005299035 |
| 57132 | *CHMP1B* | ENSG00000255112.3 | 6.33 | 6.57 | 0.24 | 1.18 | 0.001378869 | 0.007297406 |
| 5934 | *RBL2* | ENSG00000103479.17 | 6.85 | 7.08 | 0.24 | 1.18 | 0.001886183 | 0.009275791 |
| 351 | *APP* | ENSG00000142192.21 | 8.67 | 8.90 | 0.24 | 1.18 | 0.000490776 | 0.003292825 |
| 134429 | *STARD4* | ENSG00000164211.13 | 6.02 | 6.25 | 0.24 | 1.18 | 0.001926044 | 0.009419353 |
| 8763 | *CD164* | ENSG00000135535.17 | 8.43 | 8.67 | 0.24 | 1.18 | 0.00141445 | 0.007443716 |
| 8266 | *UBL4A* | ENSG00000102178.13 | 6.82 | 7.05 | 0.23 | 1.18 | 0.001972304 | 0.009604905 |
| 80829 | *ZFP91* | ENSG00000186660.15 | 7.96 | 8.19 | 0.23 | 1.18 | 0.000758048 | 0.004595132 |
| 5987 | *TRIM27* | ENSG00000204713.11 | 6.11 | 6.34 | 0.23 | 1.18 | 0.001410931 | 0.007430992 |
| 22883 | *CLSTN1* | ENSG00000171603.18 | 8.03 | 8.27 | 0.23 | 1.18 | 0.001528816 | 0.007906958 |
| 476 | *ATP1A1* | ENSG00000163399.16 | 9.14 | 9.38 | 0.23 | 1.18 | 0.002041979 | 0.009867141 |
| 10134 | *BCAP31* | ENSG00000185825.17 | 8.31 | 8.55 | 0.23 | 1.18 | 0.001083053 | 0.006060978 |
| 8788 | *DLK1* | ENSG00000185559.16 | 10.59 | 10.83 | 0.23 | 1.18 | 0.001428543 | 0.007511974 |
| 2052 | *EPHX1* | ENSG00000143819.13 | 9.44 | 9.68 | 0.23 | 1.17 | 0.001386815 | 0.007326542 |
| 6574 | *SLC20A1* | ENSG00000144136.11 | 7.20 | 7.43 | 0.23 | 1.17 | 0.000998891 | 0.00568323 |
| 10935 | *PRDX3* | ENSG00000165672.7 | 8.19 | 8.42 | 0.23 | 1.17 | 0.001287181 | 0.006934424 |
| 7414 | *VCL* | ENSG00000035403.18 | 7.85 | 8.08 | 0.23 | 1.17 | 0.001484371 | 0.007727689 |
| 29978 | *UBQLN2* | ENSG00000188021.9 | 6.42 | 6.65 | 0.23 | 1.17 | 0.001597843 | 0.008151607 |
| 23568 | *ARL2BP* | ENSG00000102931.8 | 6.74 | 6.97 | 0.23 | 1.17 | 0.001640939 | 0.008320557 |
| 382 | *ARF6* | ENSG00000165527.8 | 7.41 | 7.63 | 0.22 | 1.17 | 0.001773363 | 0.008839722 |
| 10099 | *TSPAN3* | ENSG00000140391.15 | 8.37 | 8.59 | 0.22 | 1.16 | 0.001857763 | 0.009160043 |
| 5479 | *PPIB* | ENSG00000166794.6 | 8.49 | 8.70 | 0.22 | 1.16 | 0.001323105 | 0.007079671 |
| 1736 | *DKC1* | ENSG00000130826.18 | 7.62 | 7.84 | 0.22 | 1.16 | 0.000773366 | 0.004665557 |
| 10211 | *FLOT1* | ENSG00000137312.15 | 7.05 | 7.26 | 0.22 | 1.16 | 0.001659458 | 0.008386128 |
| 6926 | *TBX3* | ENSG00000135111.16 | 9.55 | 9.76 | 0.21 | 1.16 | 0.001240211 | 0.006739436 |
| 219988 | *PATL1* | ENSG00000166889.14 | 7.05 | 7.26 | 0.21 | 1.16 | 0.002055964 | 0.009915796 |
| 9833 | *MELK* | ENSG00000165304.8 | 7.13 | 7.34 | 0.21 | 1.16 | 0.001778059 | 0.00885725 |
| 1975 | *EIF4B* | ENSG00000063046.18 | 9.78 | 9.98 | 0.21 | 1.15 | 0.00099444 | 0.005660049 |
| 26112 | *CCDC69* | ENSG00000198624.13 | 8.31 | 8.51 | 0.20 | 1.15 | 0.001684244 | 0.008477136 |
| 3329 | *HSPD1* | ENSG00000144381.18 | 11.33 | 11.52 | 0.19 | 1.14 | 0.001471241 | 0.00767867 |
| 821 | *CANX* | ENSG00000127022.16 | 10.39 | 10.58 | 0.19 | 1.14 | 0.001799419 | 0.008942875 |
| 10492 | *SYNCRIP* | ENSG00000135316.19 | 8.46 | 8.27 | -0.20 | 0.87 | 0.001832636 | 0.009071911 |
| 51280 | *GOLM1* | ENSG00000135052.16 | 8.33 | 8.13 | -0.21 | 0.87 | 0.001712949 | 0.008581407 |
| 3178 | *HNRNPA1* | ENSG00000135486.19 | 10.80 | 10.59 | -0.21 | 0.86 | 0.001644059 | 0.008333566 |
| 10280 | *SIGMAR1* | ENSG00000147955.18 | 6.69 | 6.47 | -0.22 | 0.86 | 0.002024284 | 0.009803963 |
| 22803 | *XRN2* | ENSG00000088930.8 | 7.47 | 7.25 | -0.22 | 0.86 | 0.001548991 | 0.007976906 |
| 2539 | *G6PD* | ENSG00000160211.19 | 7.53 | 7.31 | -0.23 | 0.86 | 0.000808652 | 0.004811541 |
| 149603 | *RNF187* | ENSG00000168159.15 | 7.96 | 7.73 | -0.23 | 0.85 | 0.001545562 | 0.007966118 |
| 6950 | *TCP1* | ENSG00000120438.12 | 7.90 | 7.67 | -0.23 | 0.85 | 0.000796378 | 0.004767785 |
| 222658 | *KCTD20* | ENSG00000112078.14 | 7.58 | 7.35 | -0.23 | 0.85 | 0.001779917 | 0.008860623 |
| 1653 | *DDX1* | ENSG00000079785.16 | 7.17 | 6.93 | -0.23 | 0.85 | 0.001181143 | 0.006486563 |
| 54839 | *LRRC49* | ENSG00000137821.12 | 6.92 | 6.69 | -0.24 | 0.85 | 0.001836104 | 0.009077104 |

| | | | | | | | | |
|---|---|---|---|---|---|---|---|---|
| 2058 | *EPRS1* | ENSG00000136628.18 | 11.43 | 11.19 | -0.24 | 0.85 | 0.000362247 | 0.00260142 |
| 142 | *PARP1* | ENSG00000143799.14 | 8.64 | 8.40 | -0.24 | 0.85 | 0.000594576 | 0.003828393 |
| 3190 | *HNRNPK* | ENSG00000165119.21 | 10.13 | 9.89 | -0.24 | 0.85 | 0.00043088 | 0.002987101 |
| 5425 | *POLD2* | ENSG00000106628.12 | 7.96 | 7.71 | -0.24 | 0.85 | 0.000574435 | 0.003727492 |
| 6830 | *SUPT6H* | ENSG00000109111.15 | 7.62 | 7.38 | -0.24 | 0.85 | 0.001254585 | 0.006800312 |
| 23516 | *SLC39A14* | ENSG00000104635.15 | 6.64 | 6.40 | -0.24 | 0.85 | 0.001227924 | 0.006687193 |
| 80010 | *RMI1* | ENSG00000178966.17 | 6.91 | 6.67 | -0.24 | 0.85 | 0.001711867 | 0.008581407 |
| 85406 | *DNAJC14* | ENSG00000135392.19 | 6.71 | 6.47 | -0.24 | 0.84 | 0.001319074 | 0.007068185 |
| 5095 | *PCCA* | ENSG00000175198.17 | 7.10 | 6.85 | -0.24 | 0.84 | 0.001052102 | 0.00592755 |
| 84168 | *ANTXR1* | ENSG00000169604.21 | 7.54 | 7.30 | -0.25 | 0.84 | 0.000854046 | 0.005015014 |
| 3609 | *ILF3* | ENSG00000129351.17 | 8.88 | 8.63 | -0.25 | 0.84 | 0.001110077 | 0.006198344 |
| 23270 | *TSPYL4* | ENSG00000187189.11 | 7.36 | 7.11 | -0.25 | 0.84 | 0.000632708 | 0.004003506 |
| 7410 | *VAV2* | ENSG00000160293.17 | 7.45 | 7.20 | -0.25 | 0.84 | 0.000736353 | 0.00448536 |
| 23636 | *NUP62* | ENSG00000213024.12 | 7.08 | 6.83 | -0.26 | 0.84 | 0.001585315 | 0.008101817 |
| 9318 | *COPS2* | ENSG00000166200.15 | 7.33 | 7.07 | -0.26 | 0.84 | 0.00055112 | 0.003604164 |
| 23607 | *CD2AP* | ENSG00000198087.7 | 7.00 | 6.74 | -0.26 | 0.84 | 0.001054333 | 0.005937892 |
| 9373 | *PLAA* | ENSG00000137055.15 | 6.14 | 5.88 | -0.26 | 0.84 | 0.001241109 | 0.006741874 |
| 9203 | *ZMYM3* | ENSG00000147130.14 | 7.05 | 6.80 | -0.26 | 0.84 | 0.000631994 | 0.004000674 |
| 219 | *ALDH1B1* | ENSG00000137124.8 | 6.90 | 6.64 | -0.26 | 0.84 | 0.001260722 | 0.00682618 |
| 988 | *CDC5L* | ENSG00000096401.8 | 7.13 | 6.87 | -0.26 | 0.84 | 0.000468472 | 0.003171535 |
| 8030 | *CCDC6* | ENSG00000108091.11 | 6.31 | 6.05 | -0.26 | 0.83 | 0.001385138 | 0.007322838 |
| 50512 | *PODXL2* | ENSG00000114631.11 | 5.63 | 5.37 | -0.26 | 0.83 | 0.001574115 | 0.008066196 |
| 824 | *CAPN2* | ENSG00000162909.18 | 7.16 | 6.90 | -0.26 | 0.83 | 0.001136831 | 0.00631746 |
| 26985 | *AP3M1* | ENSG00000185009.13 | 6.98 | 6.71 | -0.26 | 0.83 | 0.000921692 | 0.005330894 |
| 9898 | *UBAP2L* | ENSG00000143569.19 | 8.59 | 8.33 | -0.26 | 0.83 | 0.000438027 | 0.003018936 |
| 120 | *ADD3* | ENSG00000148700.15 | 7.62 | 7.36 | -0.26 | 0.83 | 0.001313567 | 0.007048742 |
| 9791 | *PTDSS1* | ENSG00000156471.13 | 6.79 | 6.53 | -0.27 | 0.83 | 0.001636019 | 0.008307498 |
| 219699 | *UNC5B* | ENSG00000107731.12 | 7.34 | 7.08 | -0.27 | 0.83 | 0.001147256 | 0.006353935 |
| 1967 | *EIF2B1* | ENSG00000111361.13 | 5.89 | 5.62 | -0.27 | 0.83 | 0.001959689 | 0.009558979 |
| 7019 | *TFAM* | ENSG00000108064.11 | 6.77 | 6.50 | -0.27 | 0.83 | 0.000599864 | 0.003854177 |
| 7529 | *YWHAB* | ENSG00000166913.13 | 8.55 | 8.28 | -0.27 | 0.83 | 0.000313991 | 0.002335327 |
| 328 | *APEX1* | ENSG00000100823.12 | 7.75 | 7.47 | -0.27 | 0.83 | 0.001454533 | 0.00761461 |
| 178 | *AGL* | ENSG00000162688.17 | 6.41 | 6.14 | -0.27 | 0.83 | 0.000434587 | 0.003005534 |
| 9782 | *MATR3* | ENSG00000015479.20 | 9.04 | 8.76 | -0.27 | 0.83 | 0.00023465 | 0.001864136 |
| 5705 | *PSMC5* | ENSG00000087191.13 | 6.67 | 6.40 | -0.27 | 0.83 | 0.000615118 | 0.003928643 |
| 7398 | *USP1* | ENSG00000162607.13 | 7.50 | 7.23 | -0.27 | 0.83 | 0.000354574 | 0.002564725 |
| 11331 | *PHB2* | ENSG00000215021.10 | 7.33 | 7.06 | -0.28 | 0.83 | 0.000706154 | 0.004341938 |
| 3151 | *HMGN2* | ENSG00000198830.11 | 9.40 | 9.12 | -0.28 | 0.83 | 0.000427917 | 0.002973428 |
| 54476 | *RNF216* | ENSG00000011275.19 | 6.30 | 6.02 | -0.28 | 0.82 | 0.000462194 | 0.003146063 |
| 10772 | *SRSF10* | ENSG00000188529.15 | 7.26 | 6.98 | -0.28 | 0.82 | 0.001855823 | 0.009156494 |
| 79590 | *MRPL24* | ENSG00000143314.12 | 6.19 | 5.91 | -0.28 | 0.82 | 0.001737744 | 0.008685248 |
| 3434 | *IFIT1* | ENSG00000185745.10 | 5.05 | 4.77 | -0.28 | 0.82 | 0.001350444 | 0.007187521 |
| 64756 | *ATPAF1* | ENSG00000123472.12 | 7.04 | 6.76 | -0.28 | 0.82 | 0.000368888 | 0.002637754 |
| 9126 | *SMC3* | ENSG00000108055.11 | 7.66 | 7.38 | -0.28 | 0.82 | 0.000628512 | 0.003987042 |
| 39 | *ACAT2* | ENSG00000120437.9 | 6.76 | 6.48 | -0.28 | 0.82 | 0.000769964 | 0.004654204 |
| 23511 | *NUP188* | ENSG00000095319.14 | 7.84 | 7.56 | -0.28 | 0.82 | 0.000179359 | 0.001502234 |
| 1434 | *CSE1L* | ENSG00000124207.17 | 7.81 | 7.53 | -0.28 | 0.82 | 0.0003091 | 0.002309249 |
| 231 | *AKR1B1* | ENSG00000085662.14 | 6.63 | 6.35 | -0.28 | 0.82 | 0.001069176 | 0.006003464 |
| 5902 | *RANBP1* | ENSG00000099901.17 | 7.41 | 7.13 | -0.28 | 0.82 | 0.000819166 | 0.004857712 |
| 6602 | *SMARCD1* | ENSG00000066117.15 | 7.75 | 7.47 | -0.28 | 0.82 | 0.000288741 | 0.002188741 |
| 10928 | *RALBP1* | ENSG00000017797.13 | 8.15 | 7.87 | -0.28 | 0.82 | 0.000365009 | 0.002614243 |
| 23339 | *VPS39* | ENSG00000166887.16 | 7.13 | 6.84 | -0.28 | 0.82 | 0.000464561 | 0.003154616 |
| 835 | *CASP2* | ENSG00000106144.20 | 7.09 | 6.81 | -0.28 | 0.82 | 0.001948123 | 0.009514929 |
| 84915 | *FAM222A* | ENSG00000139438.6 | 6.55 | 6.27 | -0.28 | 0.82 | 0.000774365 | 0.004669524 |
| 1836 | *SLC26A2* | ENSG00000155850.9 | 6.00 | 5.72 | -0.28 | 0.82 | 0.001273724 | 0.006889127 |
| 2289 | *FKBP5* | ENSG00000096060.15 | 6.14 | 5.86 | -0.28 | 0.82 | 0.000891546 | 0.005189691 |
| 2029 | *ENSA* | ENSG00000143420.19 | 7.73 | 7.45 | -0.29 | 0.82 | 0.000375341 | 0.002677517 |
| 114932 | *MRFAP1L1* | ENSG00000178988.11 | 5.81 | 5.52 | -0.29 | 0.82 | 0.001036805 | 0.005858961 |
| 23168 | *RTF1* | ENSG00000137815.15 | 8.02 | 7.73 | -0.29 | 0.82 | 0.001480226 | 0.007714135 |
| 10105 | *PPIF* | ENSG00000108179.14 | 9.17 | 8.88 | -0.29 | 0.82 | 0.000229723 | 0.00183107 |
| 9879 | *DDX46* | ENSG00000145833.16 | 7.60 | 7.31 | -0.29 | 0.82 | 0.000559949 | 0.003649273 |
| 6252 | *RTN1* | ENSG00000139970.18 | 6.01 | 5.72 | -0.29 | 0.82 | 0.001507518 | 0.007810265 |
| 8241 | *RBM10* | ENSG00000182872.16 | 6.29 | 6.00 | -0.29 | 0.82 | 0.001166593 | 0.006420752 |
| 646719 | *NIPBL-DT* | ENSG00000285967.1 | 5.77 | 5.48 | -0.29 | 0.82 | 0.000609033 | 0.003904729 |
| 91775 | *NXPE3* | ENSG00000144815.17 | 7.19 | 6.89 | -0.29 | 0.82 | 0.000403451 | 0.002836223 |
| 1499 | *CTNNB1* | ENSG00000168036.18 | 9.33 | 9.03 | -0.29 | 0.82 | 6.68553E-05 | 0.000731898 |
| 57190 | *SELENON* | ENSG00000162430.18 | 6.82 | 6.52 | -0.29 | 0.82 | 0.000663887 | 0.004137991 |
| 54881 | *TEX10* | ENSG00000136891.14 | 6.24 | 5.94 | -0.30 | 0.81 | 0.00192936 | 0.009432497 |
| 55837 | *EAPP* | ENSG00000129518.10 | 5.35 | 5.05 | -0.30 | 0.81 | 0.002033308 | 0.009838141 |
| 7389 | *UROD* | ENSG00000126088.14 | 6.65 | 6.35 | -0.30 | 0.81 | 0.001659085 | 0.008386128 |
| 1639 | *DCTN1* | ENSG00000204843.13 | 7.81 | 7.51 | -0.30 | 0.81 | 0.000181205 | 0.001513466 |
| 84823 | *LMNB2* | ENSG00000176619.13 | 7.66 | 7.36 | -0.30 | 0.81 | 0.000104044 | 0.00101271 |
| 55917 | *CTTNBP2NL* | ENSG00000143079.15 | 5.80 | 5.49 | -0.30 | 0.81 | 0.001561297 | 0.008019682 |
| 6421 | *SFPQ* | ENSG00000116560.11 | 8.68 | 8.38 | -0.30 | 0.81 | 0.000108463 | 0.001042501 |
| 112770 | *GLMP* | ENSG00000198715.13 | 5.90 | 5.60 | -0.30 | 0.81 | 0.000939292 | 0.005411835 |
| 8655 | *DYNLL1* | ENSG00000088986.11 | 6.95 | 6.65 | -0.30 | 0.81 | 0.001669831 | 0.008422769 |
| 54480 | *CHPF2* | ENSG00000033100.17 | 6.58 | 6.28 | -0.30 | 0.81 | 0.001044942 | 0.005898284 |
| 2091 | *FBL* | ENSG00000105202.9 | 7.41 | 7.11 | -0.30 | 0.81 | 0.000335277 | 0.00244399 |
| 221895 | *JAZF1* | ENSG00000153814.13 | 6.37 | 6.06 | -0.30 | 0.81 | 0.000328724 | 0.002410266 |
| 50650 | *ARHGEF3* | ENSG00000163947.12 | 6.93 | 6.62 | -0.30 | 0.81 | 0.000178702 | 0.001499236 |
| 5239 | *PGM5* | ENSG00000154330.13 | 7.16 | 6.85 | -0.31 | 0.81 | 0.001242806 | 0.006748335 |
| 2771 | *GNAI2* | ENSG00000114353.17 | 8.08 | 7.78 | -0.31 | 0.81 | 0.000510954 | 0.003395822 |
| 55588 | *MED29* | ENSG00000063322.15 | 6.40 | 6.09 | -0.31 | 0.81 | 0.001336195 | 0.007129381 |
| 11130 | *ZWINT* | ENSG00000122952.17 | 6.74 | 6.43 | -0.31 | 0.81 | 0.000168087 | 0.001435047 |
| 23392 | *ECPAS* | ENSG00000136813.15 | 7.74 | 7.43 | -0.31 | 0.81 | 0.000458356 | 0.003127524 |
| 56181 | *MTFR1L* | ENSG00000117640.18 | 6.58 | 6.27 | -0.31 | 0.81 | 0.001186734 | 0.006510122 |
| 29 | *ABR* | ENSG00000159842.16 | 7.82 | 7.50 | -0.31 | 0.81 | 0.000223946 | 0.00179456 |

| | | | | | | | | |
|---|---|---|---|---|---|---|---|---|
| 5550 | *PREP* | ENSG00000085377.15 | 5.50 | 5.19 | -0.31 | 0.81 | 0.001357768 | 0.007215973 |
| 684 | *BST2* | ENSG00000130303.14 | 7.04 | 6.73 | -0.31 | 0.81 | 0.000669877 | 0.004163225 |
| 9697 | *TRAM2* | ENSG00000065308.5 | 6.02 | 5.71 | -0.31 | 0.81 | 0.00164484 | 0.008334707 |
| 4522 | *MTHFD1* | ENSG00000100714.17 | 7.52 | 7.20 | -0.31 | 0.81 | 0.000477617 | 0.003224719 |
| 9459 | *ARHGEF6* | ENSG00000129675.16 | 7.28 | 6.97 | -0.31 | 0.80 | 6.58633E-05 | 0.00072527 |
| 118 | *ADD1* | ENSG00000087274.19 | 7.92 | 7.61 | -0.31 | 0.80 | 0.000242272 | 0.001905785 |
| 51535 | *PPHLN1* | ENSG00000134283.18 | 7.28 | 6.97 | -0.31 | 0.80 | 0.0004084 | 0.002865671 |
| 55827 | *DCAF6* | ENSG00000143164.16 | 6.01 | 5.70 | -0.32 | 0.80 | 0.000935931 | 0.005396612 |
| 7919 | *DDX39B* | ENSG00000198563.14 | 7.56 | 7.25 | -0.32 | 0.80 | 0.000555043 | 0.003623593 |
| 10606 | *PAICS* | ENSG00000128050.9 | 7.54 | 7.23 | -0.32 | 0.80 | 0.000611433 | 0.003916774 |
| 55819 | *RNF130* | ENSG00000113269.14 | 5.70 | 5.38 | -0.32 | 0.80 | 0.001003353 | 0.005706449 |
| 283464 | *GXYLT1* | ENSG00000151233.11 | 6.62 | 6.30 | -0.32 | 0.80 | 0.001335688 | 0.007129213 |
| 9352 | *TXNL1* | ENSG00000091164.13 | 6.49 | 6.17 | -0.32 | 0.80 | 0.000141877 | 0.001266761 |
| 112574 | *SNX18* | ENSG00000178996.14 | 7.14 | 6.82 | -0.32 | 0.80 | 0.000333478 | 0.002432057 |
| 5701 | *PSMC2* | ENSG00000161057.13 | 7.10 | 6.78 | -0.32 | 0.80 | 0.000891343 | 0.005189691 |
| 7013 | *TERF1* | ENSG00000147601.15 | 5.68 | 5.36 | -0.32 | 0.80 | 0.000520128 | 0.003442065 |
| 220988 | *HNRNPA3* | ENSG00000170144.21 | 9.69 | 9.37 | -0.32 | 0.80 | 0.000180859 | 0.001512264 |
| 6414 | *SELENOP* | ENSG00000250722.6 | 6.27 | 5.95 | -0.32 | 0.80 | 0.00069419 | 0.00428241 |
| 8910 | *SGCE* | ENSG00000127990.19 | 6.18 | 5.86 | -0.32 | 0.80 | 0.000589278 | 0.003808991 |
| 79966 | *SCD5* | ENSG00000145284.12 | 7.22 | 6.90 | -0.32 | 0.80 | 0.000317901 | 0.002356237 |
| 23598 | *PATZ1* | ENSG00000100105.18 | 5.29 | 4.97 | -0.32 | 0.80 | 0.001207818 | 0.006596184 |
| 2781 | *GNAZ* | ENSG00000128266.9 | 5.88 | 5.56 | -0.32 | 0.80 | 0.000653941 | 0.004094704 |
| 148789 | *B3GALNT2* | ENSG00000162885.14 | 5.42 | 5.10 | -0.32 | 0.80 | 0.00059226 | 0.003822707 |
| 79048 | *SECISBP2* | ENSG00000187742.15 | 6.13 | 5.80 | -0.32 | 0.80 | 0.000652889 | 0.004091156 |
| 3337 | *DNAJB1* | ENSG00000132002.9 | 7.50 | 7.18 | -0.32 | 0.80 | 0.000814038 | 0.004833244 |
| 1588 | *CYP19A1* | ENSG00000137869.16 | 5.75 | 5.42 | -0.32 | 0.80 | 0.001523078 | 0.007879998 |
| 124245 | *ZC3H18* | ENSG00000158545.16 | 6.84 | 6.52 | -0.32 | 0.80 | 0.00080865 | 0.004811541 |
| 11078 | *TRIOBP* | ENSG00000100106.22 | 5.99 | 5.67 | -0.32 | 0.80 | 0.000454047 | 0.003106302 |
| 6434 | *TRA2B* | ENSG00000136527.19 | 7.58 | 7.26 | -0.32 | 0.80 | 7.54013E-05 | 0.000799786 |
| 3192 | *HNRNPU* | ENSG00000153187.20 | 9.55 | 9.23 | -0.33 | 0.80 | 5.64125E-05 | 0.000643875 |
| 10144 | *FAM13A* | ENSG00000138640.15 | 5.02 | 4.69 | -0.33 | 0.80 | 0.001190275 | 0.006524774 |
| 5723 | *PSPH* | ENSG00000146733.14 | 6.66 | 6.34 | -0.33 | 0.80 | 0.000824286 | 0.004879955 |
| 6599 | *SMARCC1* | ENSG00000173473.11 | 7.73 | 7.40 | -0.33 | 0.80 | 0.000167384 | 0.001430673 |
| 51339 | *DACT1* | ENSG00000165617.15 | 6.93 | 6.60 | -0.33 | 0.80 | 0.00085693 | 0.005022179 |
| 11161 | *ERG28* | ENSG00000133935.7 | 5.51 | 5.18 | -0.33 | 0.80 | 0.001346803 | 0.007178328 |
| 8220 | *ESS2* | ENSG00000100056.12 | 5.31 | 4.98 | -0.33 | 0.80 | 0.000558497 | 0.003641392 |
| 79109 | *MAPKAP1* | ENSG00000119487.17 | 7.28 | 6.95 | -0.33 | 0.80 | 0.000238709 | 0.001885519 |
| 23384 | *SPECC1L* | ENSG00000100014.20 | 7.03 | 6.69 | -0.33 | 0.79 | 0.000476493 | 0.003218579 |
| 8106 | *PABPN1* | ENSG00000100836.11 | 6.55 | 6.22 | -0.33 | 0.79 | 0.00055819 | 0.003640969 |
| 64219 | *PJA1* | ENSG00000181191.12 | 6.52 | 6.19 | -0.33 | 0.79 | 0.000906559 | 0.005261621 |
| 11168 | *PSIP1* | ENSG00000164985.15 | 8.29 | 7.96 | -0.33 | 0.79 | 0.000159521 | 0.001381561 |
| 9923 | *ZBTB40* | ENSG00000184677.18 | 6.34 | 6.01 | -0.33 | 0.79 | 0.000538692 | 0.003539943 |
| 1020 | *CDK5* | ENSG00000164885.13 | 5.54 | 5.21 | -0.33 | 0.79 | 0.001583955 | 0.008100025 |
| 1114 | *CHGB* | ENSG00000089199.10 | 6.48 | 6.14 | -0.33 | 0.79 | 0.000204823 | 0.001668937 |
| 4940 | *OAS3* | ENSG00000111331.14 | 6.35 | 6.01 | -0.34 | 0.79 | 0.000177358 | 0.001495197 |
| 51088 | *KLHL5* | ENSG00000109790.18 | 6.88 | 6.54 | -0.34 | 0.79 | 0.000356332 | 0.002573718 |
| 23466 | *CBX6* | ENSG00000183741.12 | 7.27 | 6.93 | -0.34 | 0.79 | 3.87836E-05 | 0.000491641 |
| 64388 | *GREM2* | ENSG00000180875.5 | 6.37 | 6.03 | -0.34 | 0.79 | 0.00083691 | 0.004940133 |
| 81875 | *ISG20L2* | ENSG00000143319.17 | 6.57 | 6.23 | -0.34 | 0.79 | 0.001679691 | 0.008457056 |
| 89891 | *DYNC2I2* | ENSG00000119333.12 | 5.93 | 5.59 | -0.34 | 0.79 | 0.000305831 | 0.002291156 |
| 9343 | *EFTUD2* | ENSG00000108883.13 | 7.87 | 7.53 | -0.34 | 0.79 | 0.00031742 | 0.002353829 |
| 11180 | *WDR6* | ENSG00000178252.19 | 7.63 | 7.29 | -0.34 | 0.79 | 3.33775E-05 | 0.00043714 |
| 8872 | *CDC123* | ENSG00000151465.14 | 6.34 | 6.00 | -0.34 | 0.79 | 0.000719384 | 0.004403975 |
| 5297 | *PI4KA* | ENSG00000241973.11 | 8.23 | 7.89 | -0.34 | 0.79 | 0.000175964 | 0.001489588 |
| 5394 | *EXOSC10* | ENSG00000171824.14 | 5.95 | 5.61 | -0.34 | 0.79 | 0.000456717 | 0.003120099 |
| 79903 | *NAA60* | ENSG00000122390.19 | 6.07 | 5.73 | -0.34 | 0.79 | 0.002048288 | 0.00988831 |
| 25828 | *TXN2* | ENSG00000100348.10 | 6.13 | 5.79 | -0.34 | 0.79 | 0.00162138 | 0.008252047 |
| 80020 | *FOXRED2* | ENSG00000100350.16 | 7.39 | 7.05 | -0.34 | 0.79 | 5.30905E-05 | 0.000615793 |
| 149951 | *COMMD7* | ENSG00000149600.12 | 4.80 | 4.46 | -0.34 | 0.79 | 0.001645676 | 0.008336131 |
| 25776 | *CBY1* | ENSG00000100211.11 | 5.31 | 4.97 | -0.34 | 0.79 | 0.000644247 | 0.004054302 |
| 22858 | *CILK1* | ENSG00000112144.16 | 6.60 | 6.26 | -0.34 | 0.79 | 0.000104459 | 0.001014771 |
| 57178 | *ZMIZ1* | ENSG00000108175.19 | 6.73 | 6.38 | -0.34 | 0.79 | 0.001800606 | 0.008945813 |
| 126299 | *ZNF428* | ENSG00000131116.12 | 5.14 | 4.80 | -0.34 | 0.79 | 0.000822023 | 0.004870805 |
| 55000 | *TUG1* | ENSG00000253352.10 | 8.54 | 8.20 | -0.34 | 0.79 | 0.000116016 | 0.00109123 |
| 51287 | *COA4* | ENSG00000181924.7 | 5.49 | 5.15 | -0.34 | 0.79 | 0.001498306 | 0.007784034 |
| 1376 | *CPT2* | ENSG00000157184.7 | 5.84 | 5.49 | -0.34 | 0.79 | 0.000374995 | 0.002677517 |
| 85458 | *DIXDC1* | ENSG00000150764.14 | 5.88 | 5.54 | -0.34 | 0.79 | 0.000242398 | 0.001905785 |
| 51668 | *HSPB11* | ENSG00000081870.12 | 5.00 | 4.65 | -0.34 | 0.79 | 0.001436358 | 0.007532599 |
| 10576 | *CCT2* | ENSG00000166226.13 | 8.23 | 7.89 | -0.35 | 0.79 | 0.000253168 | 0.00196983 |
| 29914 | *UBIAD1* | ENSG00000120942.14 | 5.15 | 4.81 | -0.35 | 0.79 | 0.000654613 | 0.004097054 |
| 78988 | *MRPL57* | ENSG00000173141.5 | 5.74 | 5.39 | -0.35 | 0.79 | 0.000170998 | 0.00145742 |
| 1952 | *CELSR2* | ENSG00000143126.8 | 5.59 | 5.24 | -0.35 | 0.79 | 0.001965926 | 0.009580058 |
| 10969 | *EBNA1BP2* | ENSG00000117395.13 | 6.21 | 5.87 | -0.35 | 0.79 | 0.00039634 | 0.002802013 |
| 399668 | *SMIM10L2A* | ENSG00000178947.9 | 8.38 | 8.03 | -0.35 | 0.79 | 2.47738E-05 | 0.000352779 |
| 162494 | *RHBDL3* | ENSG00000141314.13 | 6.71 | 6.37 | -0.35 | 0.79 | 2.95717E-05 | 0.000398782 |
| 11112 | *HIBADH* | ENSG00000106049.9 | 7.10 | 6.75 | -0.35 | 0.79 | 8.5389E-05 | 0.000874218 |
| 2629 | *GBA* | ENSG00000177628.16 | 7.12 | 6.77 | -0.35 | 0.79 | 0.000320319 | 0.002369178 |
| 10745 | *PHTF1* | ENSG00000116793.16 | 5.88 | 5.53 | -0.35 | 0.79 | 0.000200475 | 0.001644221 |
| 9358 | *ITGBL1* | ENSG00000198542.15 | 5.69 | 5.33 | -0.35 | 0.78 | 0.00048002 | 0.00323803 |
| 10922 | *FASTK* | ENSG00000164896.21 | 5.69 | 5.34 | -0.35 | 0.78 | 0.000604004 | 0.003874144 |
| 54799 | *MBTD1* | ENSG00000011258.16 | 4.82 | 4.47 | -0.35 | 0.78 | 0.001865293 | 0.009185099 |
| 81889 | *FAHD1* | ENSG00000180185.12 | 4.89 | 4.54 | -0.35 | 0.78 | 0.00069251 | 0.004274114 |
| 80145 | *THOC7* | ENSG00000163634.12 | 5.32 | 4.97 | -0.35 | 0.78 | 0.001329326 | 0.007107894 |
| 55631 | *LRRC40* | ENSG00000066557.6 | 5.03 | 4.68 | -0.35 | 0.78 | 0.000589698 | 0.003810064 |
| 4046 | *LSP1* | ENSG00000130592.17 | 8.64 | 8.29 | -0.35 | 0.78 | 6.55812E-05 | 0.000722693 |
| 10528 | *NOP56* | ENSG00000101361.17 | 6.61 | 6.25 | -0.35 | 0.78 | 0.000360745 | 0.002594358 |
| 9156 | *EXO1* | ENSG00000174371.17 | 5.83 | 5.47 | -0.36 | 0.78 | 0.001597284 | 0.008151525 |

| | | | | | | | | |
|---|---|---|---|---|---|---|---|---|
| 55608 | *ANKRD10* | ENSG00000088448.14 | 6.55 | 6.20 | -0.36 | 0.78 | 0.000273589 | 0.002094181 |
| 2516 | *NR5A1* | ENSG00000136931.10 | 8.81 | 8.45 | -0.36 | 0.78 | 0.000139105 | 0.001248699 |
| 55243 | *KIRREL1* | ENSG00000183853.18 | 7.09 | 6.73 | -0.36 | 0.78 | 8.14315E-05 | 0.000846402 |
| 8125 | *ANP32A* | ENSG00000140350.15 | 7.86 | 7.50 | -0.36 | 0.78 | 8.65594E-05 | 0.000881989 |
| 22906 | *TRAK1* | ENSG00000182606.17 | 5.93 | 5.57 | -0.36 | 0.78 | 0.000946143 | 0.00544367 |
| 54888 | *NSUN2* | ENSG00000037474.15 | 6.36 | 6.00 | -0.36 | 0.78 | 4.14816E-05 | 0.000512848 |
| 79711 | *IPO4* | ENSG00000196497.17 | 5.96 | 5.60 | -0.36 | 0.78 | 0.000387047 | 0.002745363 |
| 221154 | *MICU2* | ENSG00000165487.14 | 5.92 | 5.55 | -0.36 | 0.78 | 0.000505814 | 0.003371119 |
| 339324 | *ZNF260* | ENSG00000254004.7 | 6.02 | 5.66 | -0.36 | 0.78 | 0.00182018 | 0.009019177 |
| 23379 | *ICE1* | ENSG00000164151.12 | 6.57 | 6.20 | -0.36 | 0.78 | 0.002061459 | 0.009939106 |
| 55794 | *DDX28* | ENSG00000182810.7 | 4.21 | 3.85 | -0.36 | 0.78 | 0.001251634 | 0.006786767 |
| 2189 | *FANCG* | ENSG00000221829.10 | 6.19 | 5.82 | -0.36 | 0.78 | 0.001546575 | 0.007968602 |
| 27246 | *RNF115* | ENSG00000265491.5 | 6.15 | 5.79 | -0.36 | 0.78 | 9.21514E-05 | 0.000924662 |
| 4174 | *MCM5* | ENSG00000100297.16 | 6.90 | 6.53 | -0.36 | 0.78 | 0.000954331 | 0.005471226 |
| 60492 | *CCDC90B* | ENSG00000137500.10 | 5.67 | 5.31 | -0.36 | 0.78 | 0.001185117 | 0.006506007 |
| 6573 | *SLC19A1* | ENSG00000173638.19 | 5.10 | 4.74 | -0.36 | 0.78 | 0.001891549 | 0.009293048 |
| 25946 | *ZNF385A* | ENSG00000161642.18 | 7.84 | 7.47 | -0.37 | 0.78 | 2.11381E-05 | 0.000313809 |
| 79710 | *MORC4* | ENSG00000133131.15 | 5.32 | 4.96 | -0.37 | 0.78 | 0.00032334 | 0.002383096 |
| 9683 | *N4BP1* | ENSG00000102921.8 | 6.11 | 5.74 | -0.37 | 0.78 | 0.000917964 | 0.005311381 |
| 55086 | *RADX* | ENSG00000147231.14 | 5.97 | 5.60 | -0.37 | 0.78 | 0.001176255 | 0.006462086 |
| 8528 | *DDO* | ENSG00000203797.12 | 4.30 | 3.94 | -0.37 | 0.78 | 0.000697245 | 0.004297723 |
| 9727 | *RAB11FIP3* | ENSG00000090565.17 | 5.95 | 5.59 | -0.37 | 0.78 | 0.001071739 | 0.006013357 |
| 29086 | *BABAM1* | ENSG00000105393.16 | 5.86 | 5.50 | -0.37 | 0.77 | 0.000855554 | 0.005021902 |
| 92399 | *MRRF* | ENSG00000148187.18 | 5.07 | 4.70 | -0.37 | 0.77 | 0.000917524 | 0.005310882 |
| 7559 | *ZNF12* | ENSG00000164631.19 | 5.55 | 5.18 | -0.37 | 0.77 | 0.000241409 | 0.001901004 |
| 225 | *ABCD2* | ENSG00000173208.4 | 4.77 | 4.40 | -0.37 | 0.77 | 0.000797362 | 0.004769214 |
| 9962 | *SLC23A2* | ENSG00000089057.15 | 8.96 | 8.59 | -0.37 | 0.77 | 1.12851E-05 | 0.000201161 |
| 2580 | *GAK* | ENSG00000178950.18 | 6.99 | 6.62 | -0.37 | 0.77 | 0.00045074 | 0.003088341 |
| 11169 | *WDHD1* | ENSG00000198554.12 | 5.74 | 5.37 | -0.37 | 0.77 | 0.000418761 | 0.002921964 |
| 375748 | *ERCC6L2* | ENSG00000182150.19 | 5.81 | 5.43 | -0.37 | 0.77 | 0.001349401 | 0.007184522 |
| 51701 | *NLK* | ENSG00000087095.13 | 7.78 | 7.41 | -0.37 | 0.77 | 4.7528E-05 | 0.000568468 |
| 140885 | *SIRPA* | ENSG00000198053.12 | 6.98 | 6.60 | -0.37 | 0.77 | 2.74366E-05 | 0.000379535 |
| 56904 | *SH3GLB2* | ENSG00000148341.18 | 5.51 | 5.14 | -0.37 | 0.77 | 0.001352892 | 0.007195449 |
| 79665 | *DHX40* | ENSG00000108406.10 | 6.95 | 6.57 | -0.37 | 0.77 | 6.18789E-05 | 0.00069 |
| 6742 | *SSBP1* | ENSG00000106028.11 | 7.18 | 6.81 | -0.37 | 0.77 | 8.91937E-05 | 0.000902094 |
| 10109 | *ARPC2* | ENSG00000163466.16 | 6.69 | 6.31 | -0.38 | 0.77 | 0.000117716 | 0.001101647 |
| 1616 | *DAXX* | ENSG00000204209.13 | 6.50 | 6.13 | -0.38 | 0.77 | 0.000366225 | 0.002619958 |
| 4082 | *MARCKS* | ENSG00000277443.3 | 6.91 | 6.53 | -0.38 | 0.77 | 6.49721E-05 | 0.000718369 |
| 2941 | *GSTA4* | ENSG00000170899.11 | 6.70 | 6.32 | -0.38 | 0.77 | 9.51325E-05 | 0.000948103 |
| 222553 | *SLC35F1* | ENSG00000196376.11 | 6.23 | 5.86 | -0.38 | 0.77 | 5.64733E-05 | 0.000644079 |
| 6988 | *TCTA* | ENSG00000145022.5 | 4.96 | 4.58 | -0.38 | 0.77 | 0.001385959 | 0.007324599 |
| 51435 | *SCARA3* | ENSG00000168077.14 | 5.24 | 4.86 | -0.38 | 0.77 | 0.000330075 | 0.002413103 |
| 6259 | *RYK* | ENSG00000163785.13 | 5.71 | 5.33 | -0.38 | 0.77 | 0.001880014 | 0.009248485 |
| 26750 | *RPS6KC1* | ENSG00000136643.12 | 6.20 | 5.82 | -0.38 | 0.77 | 0.000278502 | 0.002124001 |
| 63979 | *FIGNL1* | ENSG00000132436.12 | 6.96 | 6.57 | -0.38 | 0.77 | 7.80024E-05 | 0.000822653 |
| 85476 | *GFM1* | ENSG00000168827.15 | 6.56 | 6.18 | -0.38 | 0.77 | 0.000725882 | 0.004434171 |
| 9765 | *ZFYVE16* | ENSG00000039319.17 | 7.03 | 6.65 | -0.38 | 0.77 | 1.37324E-05 | 0.000232892 |
| 54939 | *COMMD4* | ENSG00000140365.16 | 5.35 | 4.96 | -0.38 | 0.77 | 9.87197E-05 | 0.000974151 |
| 79605 | *PGBD5* | ENSG00000177614.11 | 6.59 | 6.20 | -0.38 | 0.77 | 0.000204213 | 0.001668169 |
| 9672 | *SDC3* | ENSG00000162512.16 | 7.28 | 6.89 | -0.38 | 0.77 | 0.000103552 | 0.001009015 |
| 79142 | *PHF23* | ENSG00000040633.13 | 6.34 | 5.95 | -0.39 | 0.77 | 0.000113423 | 0.001076765 |
| 79738 | *BBS10* | ENSG00000179941.9 | 4.92 | 4.54 | -0.39 | 0.77 | 0.000291927 | 0.002208432 |
| 3028 | *HSD17B10* | ENSG00000072506.14 | 6.05 | 5.67 | -0.39 | 0.77 | 0.001264195 | 0.006842516 |
| 5358 | *PLS3* | ENSG00000102024.19 | 7.81 | 7.42 | -0.39 | 0.77 | 0.000805914 | 0.004805707 |
| 23012 | *STK38L* | ENSG00000211455.8 | 6.15 | 5.76 | -0.39 | 0.77 | 0.001032536 | 0.005843639 |
| 9749 | *PHACTR2* | ENSG00000112419.15 | 8.04 | 7.65 | -0.39 | 0.77 | 0.001186034 | 0.006508662 |
| 1432 | *MAPK14* | ENSG00000112062.11 | 7.58 | 7.19 | -0.39 | 0.76 | 0.000177563 | 0.001495535 |
| 11346 | *SYNPO* | ENSG00000171992.13 | 7.21 | 6.82 | -0.39 | 0.76 | 3.72936E-05 | 0.00047678 |
| 3996 | *LLGL1* | ENSG00000131899.12 | 6.36 | 5.98 | -0.39 | 0.76 | 8.4987E-05 | 0.00087185 |
| 9141 | *PDCD5* | ENSG00000105185.12 | 5.95 | 5.56 | -0.39 | 0.76 | 0.000174394 | 0.001481313 |
| 81605 | *URM1* | ENSG00000167118.11 | 6.23 | 5.84 | -0.39 | 0.76 | 0.000269125 | 0.002069828 |
| 23654 | *PLXNB2* | ENSG00000196576.16 | 6.48 | 6.09 | -0.39 | 0.76 | 0.000295737 | 0.00223388 |
| 6817 | *SULT1A1* | ENSG00000196502.13 | 5.61 | 5.22 | -0.39 | 0.76 | 0.00160558 | 0.008185514 |
| 83594 | *NUDT12* | ENSG00000112874.10 | 4.69 | 4.30 | -0.39 | 0.76 | 0.001229454 | 0.006690671 |
| 24145 | *PANX1* | ENSG00000110218.9 | 4.65 | 4.26 | -0.39 | 0.76 | 0.001432071 | 0.007515367 |
| 55131 | *RBM28* | ENSG00000106344.8 | 6.07 | 5.67 | -0.39 | 0.76 | 0.000471063 | 0.003183337 |
| 8540 | *AGPS* | ENSG00000018510.18 | 7.78 | 7.39 | -0.39 | 0.76 | 4.0097E-05 | 0.00050319 |
| 159 | *ADSS2* | ENSG00000035687.10 | 6.20 | 5.81 | -0.39 | 0.76 | 3.82022E-05 | 0.000485913 |
| 80851 | *SH3BP5L* | ENSG00000175137.11 | 5.81 | 5.41 | -0.39 | 0.76 | 4.03556E-05 | 0.000505169 |
| 1808 | *DPYSL2* | ENSG00000092964.18 | 5.32 | 4.93 | -0.39 | 0.76 | 0.000501457 | 0.003349519 |
| 26153 | *KIF26A* | ENSG00000066735.15 | 5.74 | 5.35 | -0.39 | 0.76 | 0.000320436 | 0.002369178 |
| 51222 | *ZNF219* | ENSG00000165804.16 | 4.87 | 4.47 | -0.39 | 0.76 | 0.00176546 | 0.008803251 |
| 9324 | *HMGN3* | ENSG00000118418.14 | 6.68 | 6.29 | -0.39 | 0.76 | 1.27927E-05 | 0.000219938 |
| 7027 | *TFDP1* | ENSG00000198176.13 | 8.40 | 8.01 | -0.39 | 0.76 | 6.35158E-06 | 0.000130699 |
| 79762 | *C1orf115* | ENSG00000162817.7 | 4.79 | 4.39 | -0.40 | 0.76 | 0.00032552 | 0.002392621 |
| 266655 | *BRD3OS* | ENSG00000235106.10 | 5.75 | 5.35 | -0.40 | 0.76 | 0.000464776 | 0.003154616 |
| 23557 | *SNAPIN* | ENSG00000143553.11 | 5.36 | 4.96 | -0.40 | 0.76 | 0.000454076 | 0.003106302 |
| 10154 | *PLXNC1* | ENSG00000136040.9 | 7.09 | 6.69 | -0.40 | 0.76 | 0.000357264 | 0.002576731 |
| 4130 | *MAP1A* | ENSG00000166963.13 | 5.08 | 4.68 | -0.40 | 0.76 | 0.000387547 | 0.002747611 |
| 8078 | *USP5* | ENSG00000111667.14 | 8.39 | 7.99 | -0.40 | 0.76 | 1.9783E-05 | 0.000299016 |
| 9517 | *SPTLC2* | ENSG00000100596.8 | 5.95 | 5.55 | -0.40 | 0.76 | 0.001560433 | 0.008018927 |
| 25929 | *GEMIN5* | ENSG00000082516.9 | 5.97 | 5.57 | -0.40 | 0.76 | 6.66817E-05 | 0.000730846 |
| 6241 | *RRM2* | ENSG00000171848.16 | 8.05 | 7.65 | -0.40 | 0.76 | 3.26408E-05 | 0.000429365 |
| 25960 | *ADGRA2* | ENSG00000020181.18 | 4.41 | 4.01 | -0.40 | 0.76 | 0.001441851 | 0.007553489 |
| 51520 | *LARS1* | ENSG00000133706.19 | 7.13 | 6.73 | -0.40 | 0.76 | 0.000117354 | 0.001099482 |
| 6821 | *SUOX* | ENSG00000139531.13 | 5.77 | 5.37 | -0.40 | 0.76 | 0.001558059 | 0.008014017 |
| 10190 | *TXNDC9* | ENSG00000115514.12 | 4.55 | 4.15 | -0.41 | 0.76 | 0.000397074 | 0.002802619 |

| | | | | | | | | |
|---|---|---|---|---|---|---|---|---|
| 23224 | *SYNE2* | ENSG00000054654.20 | 6.92 | 6.52 | -0.41 | 0.75 | 0.000196445 | 0.001620027 |
| 55508 | *SLC35E3* | ENSG00000175782.11 | 5.31 | 4.90 | -0.41 | 0.75 | 0.001027576 | 0.00581776 |
| 5356 | *PLRG1* | ENSG00000171566.12 | 5.87 | 5.47 | -0.41 | 0.75 | 0.00106697 | 0.00599781 |
| 22985 | *ACIN1* | ENSG00000100813.15 | 7.88 | 7.48 | -0.41 | 0.75 | 0.000103001 | 0.001006037 |
| 134147 | *CMBL* | ENSG00000164237.9 | 6.18 | 5.77 | -0.41 | 0.75 | 0.00042323 | 0.002947687 |
| 27090 | *ST6GALNAC4* | ENSG00000136840.19 | 3.80 | 3.40 | -0.41 | 0.75 | 0.001429333 | 0.007511974 |
| 56160 | *NSMCE3* | ENSG00000185115.6 | 5.74 | 5.33 | -0.41 | 0.75 | 0.000326873 | 0.00240021 |
| 64785 | *GINS3* | ENSG00000181938.14 | 6.16 | 5.75 | -0.41 | 0.75 | 1.67182E-05 | 0.000265216 |
| 7145 | *TNS1* | ENSG00000079308.20 | 6.63 | 6.22 | -0.41 | 0.75 | 0.001216215 | 0.006630646 |
| 79600 | *TCTN1* | ENSG00000204852.17 | 4.80 | 4.40 | -0.41 | 0.75 | 0.001573762 | 0.008066196 |
| 7324 | *UBE2E1* | ENSG00000170142.12 | 5.70 | 5.29 | -0.41 | 0.75 | 0.000901695 | 0.005241494 |
| 2035 | *EPB41* | ENSG00000159023.22 | 7.67 | 7.26 | -0.41 | 0.75 | 0.000408867 | 0.002866271 |
| 4343 | *MOV10* | ENSG00000155363.19 | 6.64 | 6.23 | -0.41 | 0.75 | 8.04225E-05 | 0.000839403 |
| 56892 | *TCIM* | ENSG00000176907.5 | 4.08 | 3.67 | -0.41 | 0.75 | 0.001677436 | 0.008451372 |
| 7077 | *TIMP2* | ENSG00000035862.12 | 7.21 | 6.80 | -0.41 | 0.75 | 4.24462E-05 | 0.000520487 |
| 6837 | *MED22* | ENSG00000148297.16 | 5.85 | 5.43 | -0.41 | 0.75 | 0.00085717 | 0.005022179 |
| 79954 | *NOL10* | ENSG00000115761.16 | 5.61 | 5.19 | -0.41 | 0.75 | 4.80898E-05 | 0.000573237 |
| 57446 | *NDRG3* | ENSG00000101079.21 | 6.31 | 5.89 | -0.42 | 0.75 | 2.48685E-05 | 0.000353458 |
| 5610 | *EIF2AK2* | ENSG00000055332.19 | 7.52 | 7.10 | -0.42 | 0.75 | 5.11406E-05 | 0.000597795 |
| 8263 | *F8A1* | ENSG00000288722.1 | 6.22 | 5.81 | -0.42 | 0.75 | 2.87341E-05 | 0.000390997 |
| 8899 | *PRPF4B* | ENSG00000112739.17 | 6.70 | 6.28 | -0.42 | 0.75 | 4.50693E-05 | 0.00054552 |
| 54916 | *TMEM260* | ENSG00000070269.14 | 4.33 | 3.91 | -0.42 | 0.75 | 0.001569877 | 0.008052729 |
| 84861 | *KLHL22* | ENSG00000099910.17 | 5.81 | 5.39 | -0.42 | 0.75 | 0.000438278 | 0.003018936 |
| 116224 | *PABIR1* | ENSG00000187866.10 | 6.37 | 5.95 | -0.42 | 0.75 | 2.79041E-05 | 0.00038388 |
| 9937 | *DCLRE1A* | ENSG00000198924.8 | 5.08 | 4.66 | -0.42 | 0.75 | 8.00778E-05 | 0.000838137 |
| 116461 | *TSEN15* | ENSG00000198860.14 | 4.82 | 4.40 | -0.42 | 0.75 | 0.001421489 | 0.007478137 |
| 8437 | *RASAL1* | ENSG00000111344.12 | 5.19 | 4.77 | -0.42 | 0.75 | 0.000488373 | 0.003279636 |
| 114991 | *ZNF618* | ENSG00000157657.15 | 8.22 | 7.80 | -0.42 | 0.75 | 6.05081E-05 | 0.000678749 |
| 203 | *AK1* | ENSG00000106992.19 | 5.58 | 5.16 | -0.42 | 0.75 | 0.00116308 | 0.006406115 |
| 9183 | *ZW10* | ENSG00000086827.9 | 5.40 | 4.98 | -0.42 | 0.75 | 0.000654863 | 0.004097054 |
| 4605 | *MYBL2* | ENSG00000101057.16 | 6.57 | 6.15 | -0.42 | 0.75 | 2.08111E-05 | 0.0003108 |
| 55964 | *SEPTIN3* | ENSG00000100167.21 | 4.84 | 4.41 | -0.42 | 0.75 | 0.000155666 | 0.001353642 |
| 25936 | *NSL1* | ENSG00000117697.15 | 5.11 | 4.68 | -0.43 | 0.74 | 0.000708752 | 0.00435434 |
| 55809 | *TRERF1* | ENSG00000124496.12 | 5.07 | 4.64 | -0.43 | 0.74 | 0.000262332 | 0.002028508 |
| 5715 | *PSMD9* | ENSG00000110801.14 | 5.09 | 4.66 | -0.43 | 0.74 | 0.000819662 | 0.004858733 |
| 3187 | *HNRNPH1* | ENSG00000169045.17 | 9.04 | 8.62 | -0.43 | 0.74 | 7.85548E-06 | 0.000152922 |
| 79085 | *SLC25A23* | ENSG00000125648.15 | 6.35 | 5.92 | -0.43 | 0.74 | 0.000931642 | 0.00538015 |
| 114805 | *GALNT13* | ENSG00000144278.15 | 5.48 | 5.05 | -0.43 | 0.74 | 0.000594563 | 0.003828393 |
| 439994 | *LINC00863* | ENSG00000224914.4 | 4.68 | 4.25 | -0.43 | 0.74 | 0.001136882 | 0.00631746 |
| 51616 | *TAF9B* | ENSG00000187325.5 | 5.58 | 5.15 | -0.43 | 0.74 | 6.59682E-05 | 0.000725562 |
| 9989 | *PPP4R1* | ENSG00000154845.16 | 6.15 | 5.72 | -0.43 | 0.74 | 0.00015004 | 0.00132079 |
| 23491 | *CES3* | ENSG00000172828.13 | 3.47 | 3.04 | -0.43 | 0.74 | 0.001634133 | 0.00830569 |
| 2067 | *ERCC1* | ENSG00000012061.16 | 5.74 | 5.30 | -0.43 | 0.74 | 8.17252E-05 | 0.000848867 |
| 210 | *ALAD* | ENSG00000148218.16 | 6.24 | 5.81 | -0.43 | 0.74 | 4.59776E-05 | 0.000553558 |
| 1119 | *CHKA* | ENSG00000110721.12 | 6.48 | 6.04 | -0.43 | 0.74 | 0.001129949 | 0.006288247 |
| 57325 | *KAT14* | ENSG00000149474.15 | 5.53 | 5.09 | -0.43 | 0.74 | 0.000153619 | 0.001341285 |
| 138162 | *C9orf116* | ENSG00000160345.13 | 3.71 | 3.27 | -0.44 | 0.74 | 0.001487523 | 0.007741412 |
| 116442 | *RAB39B* | ENSG00000155961.5 | 4.80 | 4.36 | -0.44 | 0.74 | 0.000331731 | 0.002420492 |
| 5445 | *PON2* | ENSG00000105854.13 | 7.23 | 6.79 | -0.44 | 0.74 | 1.25506E-05 | 0.000217019 |
| 154796 | *AMOT* | ENSG00000126016.17 | 6.46 | 6.02 | -0.44 | 0.74 | 0.000148826 | 0.001313957 |
| 339290 | *LINC00667* | ENSG00000263753.9 | 4.98 | 4.54 | -0.44 | 0.74 | 0.000487685 | 0.003276482 |
| 165055 | *CCDC138* | ENSG00000163006.12 | 3.33 | 2.89 | -0.44 | 0.74 | 0.00189642 | 0.009313931 |
| 27153 | *ZNF777* | ENSG00000196453.8 | 5.28 | 4.84 | -0.44 | 0.74 | 0.000137055 | 0.001233252 |
| 5982 | *RFC2* | ENSG00000049541.11 | 5.45 | 5.01 | -0.44 | 0.74 | 0.00053468 | 0.003519743 |
| 23277 | *CLUH* | ENSG00000132361.18 | 6.78 | 6.34 | -0.44 | 0.74 | 0.001159291 | 0.00638994 |
| 63967 | *CLSPN* | ENSG00000092853.14 | 6.00 | 5.55 | -0.44 | 0.74 | 0.000237237 | 0.001879987 |
| 100128252 | *ZNF667-AS1* | ENSG00000166770.12 | 5.39 | 4.95 | -0.44 | 0.74 | 0.000942031 | 0.005425532 |
| 9315 | *NREP* | ENSG00000134986.14 | 8.53 | 8.09 | -0.44 | 0.74 | 9.95358E-05 | 0.000981559 |
| 3092 | *HIP1* | ENSG00000127946.17 | 5.97 | 5.52 | -0.44 | 0.73 | 0.001208136 | 0.006596184 |
| 27000 | *DNAJC2* | ENSG00000105821.15 | 4.93 | 4.48 | -0.44 | 0.73 | 0.000713893 | 0.004382336 |
| 6642 | *SNX1* | ENSG00000028528.15 | 7.47 | 7.02 | -0.45 | 0.73 | 0.001276149 | 0.006897269 |
| 5980 | *REV3L* | ENSG00000009413.16 | 6.45 | 6.01 | -0.45 | 0.73 | 0.000674813 | 0.004186624 |
| 54865 | *GPATCH4* | ENSG00000160818.17 | 5.20 | 4.75 | -0.45 | 0.73 | 0.001549249 | 0.007976906 |
| 10040 | *TOM1L1* | ENSG00000141198.16 | 5.10 | 4.66 | -0.45 | 0.73 | 0.000180805 | 0.001512264 |
| 4841 | *NONO* | ENSG00000147140.17 | 9.12 | 8.67 | -0.45 | 0.73 | 4.28218E-06 | 9.79744E-05 |
| 8805 | *TRIM24* | ENSG00000122779.18 | 7.88 | 7.43 | -0.45 | 0.73 | 3.14483E-06 | 7.85015E-05 |
| 11338 | *U2AF2* | ENSG00000063244.13 | 7.86 | 7.41 | -0.45 | 0.73 | 1.39725E-05 | 0.000234579 |
| 9134 | *CCNE2* | ENSG00000175305.18 | 5.93 | 5.48 | -0.45 | 0.73 | 1.8206E-05 | 0.0002794 |
| 10382 | *TUBB4A* | ENSG00000104833.12 | 6.82 | 6.37 | -0.45 | 0.73 | 3.7142E-05 | 0.00047678 |
| 220972 | *MARCHF8* | ENSG00000165406.16 | 6.14 | 5.69 | -0.45 | 0.73 | 0.000126071 | 0.001156599 |
| 5337 | *PLD1* | ENSG00000075651.17 | 8.11 | 7.66 | -0.45 | 0.73 | 1.04073E-05 | 0.000190259 |
| 7913 | *DEK* | ENSG00000124795.17 | 8.49 | 8.04 | -0.45 | 0.73 | 4.29395E-06 | 9.80943E-05 |
| 3376 | *IARS1* | ENSG00000196305.19 | 8.61 | 8.15 | -0.45 | 0.73 | 4.53228E-06 | 0.000102293 |
| 9931 | *HELZ* | ENSG00000198265.12 | 6.75 | 6.29 | -0.45 | 0.73 | 0.000675494 | 0.004187728 |
| 29087 | *THYN1* | ENSG00000151500.15 | 5.01 | 4.56 | -0.46 | 0.73 | 0.000870585 | 0.005082305 |
| 7342 | *UBP1* | ENSG00000153560.12 | 6.53 | 6.07 | -0.46 | 0.73 | 0.001615466 | 0.008230321 |
| 6941 | *TCF19* | ENSG00000137310.12 | 6.90 | 6.44 | -0.46 | 0.73 | 7.15091E-06 | 0.000141562 |
| 2800 | *GOLGA1* | ENSG00000136935.14 | 5.43 | 4.97 | -0.46 | 0.73 | 0.000865175 | 0.005058595 |
| 54809 | *SAMD9* | ENSG00000205413.8 | 6.51 | 6.05 | -0.46 | 0.73 | 2.0131E-05 | 0.000303664 |
| 10885 | *WDR3* | ENSG00000065183.16 | 6.65 | 6.19 | -0.46 | 0.73 | 4.20004E-05 | 0.000516708 |
| 27350 | *APOBEC3C* | ENSG00000244509.4 | 6.18 | 5.72 | -0.46 | 0.73 | 4.18972E-05 | 0.000516285 |
| 3480 | *IGF1R* | ENSG00000140443.15 | 7.00 | 6.54 | -0.46 | 0.73 | 0.000440747 | 0.003028927 |
| 84851 | *TRIM52* | ENSG00000183718.6 | 4.51 | 4.05 | -0.46 | 0.73 | 0.000429847 | 0.00298276 |
| 9379 | *NRXN2* | ENSG00000110076.20 | 6.62 | 6.17 | -0.46 | 0.73 | 0.001564344 | 0.00803258 |
| 90861 | *JPT2* | ENSG00000206053.13 | 7.50 | 7.04 | -0.46 | 0.73 | 0.000250516 | 0.001953243 |
| 26133 | *TRPC4AP* | ENSG00000100991.12 | 6.51 | 6.05 | -0.46 | 0.73 | 2.44826E-06 | 6.52681E-05 |
| 7756 | *ZNF207* | ENSG00000010244.19 | 7.94 | 7.48 | -0.46 | 0.73 | 1.34798E-06 | 4.2864E-05 |

| | | | | | | | | |
|---|---|---|---|---|---|---|---|---|
| 3184 | *HNRNPD* | ENSG00000138668.19 | 8.66 | 8.20 | -0.46 | 0.73 | 1.12451E-06 | 3.78517E-05 |
| 25989 | *ULK3* | ENSG00000140474.14 | 5.99 | 5.53 | -0.46 | 0.73 | 0.000726223 | 0.004434449 |
| 340481 | *ZDHHC21* | ENSG00000175893.12 | 4.86 | 4.40 | -0.46 | 0.73 | 0.001782259 | 0.008869337 |
| 26523 | *AGO1* | ENSG00000092847.13 | 8.23 | 7.77 | -0.46 | 0.73 | 2.28127E-06 | 6.18043E-05 |
| 26273 | *FBXO3* | ENSG00000110429.14 | 4.93 | 4.47 | -0.46 | 0.73 | 0.001215658 | 0.006630016 |
| 256471 | *MFSD8* | ENSG00000164073.11 | 4.35 | 3.89 | -0.46 | 0.73 | 0.001351543 | 0.007190824 |
| 23657 | *SLC7A11* | ENSG00000151012.13 | 5.57 | 5.11 | -0.46 | 0.73 | 0.001070343 | 0.006007768 |
| 1741 | *DLG3* | ENSG00000082458.12 | 5.94 | 5.47 | -0.46 | 0.73 | 0.001647969 | 0.008342111 |
| 57664 | *PLEKHA4* | ENSG00000105559.12 | 4.14 | 3.68 | -0.46 | 0.72 | 0.000773299 | 0.004665557 |
| 51367 | *POP5* | ENSG00000167272.11 | 4.37 | 3.91 | -0.46 | 0.72 | 0.000713334 | 0.0043807 |
| 7447 | *VSNL1* | ENSG00000163032.12 | 6.01 | 5.54 | -0.47 | 0.72 | 0.000162517 | 0.001396231 |
| 152007 | *GLIPR2* | ENSG00000122694.16 | 7.40 | 6.94 | -0.47 | 0.72 | 2.03672E-06 | 5.69257E-05 |
| 3762 | *KCNJ5* | ENSG00000120457.12 | 8.15 | 7.68 | -0.47 | 0.72 | 1.59227E-05 | 0.000256122 |
| 9568 | *GABBR2* | ENSG00000136928.7 | 7.54 | 7.07 | -0.47 | 0.72 | 7.19813E-05 | 0.000774384 |
| 51747 | *LUC7L3* | ENSG00000108848.16 | 6.21 | 5.74 | -0.47 | 0.72 | 0.0006836 | 0.00422925 |
| 55035 | *NOL8* | ENSG00000198000.12 | 6.15 | 5.68 | -0.47 | 0.72 | 0.000365078 | 0.002614243 |
| 23398 | *PPWD1* | ENSG00000113593.12 | 5.59 | 5.12 | -0.47 | 0.72 | 0.002035535 | 0.009842573 |
| 79912 | *PYROXD1* | ENSG00000121350.16 | 4.98 | 4.51 | -0.47 | 0.72 | 0.001348394 | 0.007181705 |
| 26019 | *UPF2* | ENSG00000151461.20 | 5.66 | 5.19 | -0.47 | 0.72 | 8.97759E-05 | 0.000907182 |
| 3204 | *HOXA7* | ENSG00000122592.8 | 5.29 | 4.82 | -0.47 | 0.72 | 0.000167614 | 0.001431825 |
| 10587 | *TXNRD2* | ENSG00000184470.21 | 6.06 | 5.58 | -0.47 | 0.72 | 7.36418E-05 | 0.000785564 |
| 64759 | *TNS3* | ENSG00000136205.17 | 7.52 | 7.05 | -0.47 | 0.72 | 2.25443E-05 | 0.000329151 |
| 338761 | *C1QL4* | ENSG00000186897.5 | 4.24 | 3.77 | -0.47 | 0.72 | 0.000837231 | 0.004940133 |
| 90324 | *CCDC97* | ENSG00000142039.4 | 5.25 | 4.78 | -0.47 | 0.72 | 2.85362E-05 | 0.000389369 |
| 4858 | *NOVA2* | ENSG00000104967.8 | 5.43 | 4.95 | -0.47 | 0.72 | 0.000120421 | 0.00111775 |
| 64792 | *IFT22* | ENSG00000128581.17 | 6.03 | 5.55 | -0.47 | 0.72 | 0.000177382 | 0.001495197 |
| 51763 | *INPP5K* | ENSG00000132376.20 | 5.09 | 4.61 | -0.48 | 0.72 | 0.00133232 | 0.007113762 |
| 11140 | *CDC37* | ENSG00000105401.9 | 6.93 | 6.45 | -0.48 | 0.72 | 3.12525E-06 | 7.81782E-05 |
| 57542 | *KLHL42* | ENSG00000087448.11 | 7.07 | 6.59 | -0.48 | 0.72 | 2.00654E-06 | 5.63973E-05 |
| 56886 | *UGGT1* | ENSG00000136731.13 | 7.47 | 7.00 | -0.48 | 0.72 | 0.000237804 | 0.001881789 |
| 3837 | *KPNB1* | ENSG00000108424.11 | 8.76 | 8.28 | -0.48 | 0.72 | 8.66757E-06 | 0.000165722 |
| 65108 | *MARCKSL1* | ENSG00000175130.7 | 7.22 | 6.74 | -0.48 | 0.72 | 4.07619E-06 | 9.46813E-05 |
| 90557 | *CCDC74A* | ENSG00000163040.15 | 4.40 | 3.92 | -0.48 | 0.72 | 0.000700341 | 0.004315035 |
| 8881 | *CDC16* | ENSG00000130177.16 | 7.58 | 7.10 | -0.48 | 0.72 | 4.67857E-06 | 0.000105121 |
| 9987 | *HNRNPDL* | ENSG00000152795.18 | 8.06 | 7.58 | -0.48 | 0.72 | 1.8194E-06 | 5.24132E-05 |
| 4686 | *NCBP1* | ENSG00000136937.13 | 7.36 | 6.87 | -0.48 | 0.72 | 2.96645E-06 | 7.47038E-05 |
| 9844 | *ELMO1* | ENSG00000155849.16 | 4.52 | 4.04 | -0.48 | 0.72 | 0.001278953 | 0.006904964 |
| 5702 | *PSMC3* | ENSG00000165916.9 | 7.69 | 7.20 | -0.48 | 0.72 | 0.000275433 | 0.002104878 |
| 23167 | *EFR3A* | ENSG00000132294.15 | 5.30 | 4.82 | -0.48 | 0.72 | 0.001712965 | 0.008581407 |
| 493 | *ATP2B4* | ENSG00000058668.15 | 7.83 | 7.35 | -0.48 | 0.72 | 8.02548E-05 | 0.000839206 |
| 5050 | *PAFAH1B3* | ENSG00000079462.8 | 5.30 | 4.82 | -0.48 | 0.72 | 0.002014388 | 0.009768645 |
| 2260 | *FGFR1* | ENSG00000077782.22 | 7.67 | 7.18 | -0.48 | 0.72 | 6.89765E-05 | 0.000749109 |
| 9063 | *PIAS2* | ENSG00000078043.16 | 6.00 | 5.51 | -0.48 | 0.72 | 2.76069E-05 | 0.000381188 |
| 169981 | *SPIN3* | ENSG00000204271.13 | 3.72 | 3.24 | -0.48 | 0.71 | 0.000641937 | 0.004041456 |
| 145957 | *NRG4* | ENSG00000169752.17 | 4.09 | 3.60 | -0.48 | 0.71 | 0.001315424 | 0.007051143 |
| 79989 | *TTC26* | ENSG00000105948.13 | 5.84 | 5.36 | -0.49 | 0.71 | 0.000794604 | 0.004761391 |
| 1107 | *CHD3* | ENSG00000170004.18 | 8.02 | 7.53 | -0.49 | 0.71 | 0.000198139 | 0.001630406 |
| 80233 | *FAAP100* | ENSG00000185504.17 | 4.92 | 4.44 | -0.49 | 0.71 | 0.000958657 | 0.005489695 |
| 2618 | *GART* | ENSG00000159131.17 | 7.06 | 6.58 | -0.49 | 0.71 | 0.000526411 | 0.003475979 |
| 1601 | *DAB2* | ENSG00000153071.15 | 9.77 | 9.28 | -0.49 | 0.71 | 2.50641E-06 | 6.64642E-05 |
| 404217 | *CTXN1* | ENSG00000178531.6 | 4.97 | 4.49 | -0.49 | 0.71 | 0.000364658 | 0.002613734 |
| 7596 | *ZNF45* | ENSG00000124459.12 | 5.30 | 4.81 | -0.49 | 0.71 | 0.001913722 | 0.009374362 |
| 1075 | *CTSC* | ENSG00000109861.17 | 7.56 | 7.07 | -0.49 | 0.71 | 1.24378E-05 | 0.000216064 |
| 837 | *CASP4* | ENSG00000196954.14 | 4.22 | 3.73 | -0.49 | 0.71 | 9.07369E-05 | 0.000914621 |
| 23506 | *BICRAL* | ENSG00000112624.13 | 4.98 | 4.49 | -0.49 | 0.71 | 0.000593023 | 0.003824959 |
| 56062 | *KLHL4* | ENSG00000102271.14 | 5.43 | 4.94 | -0.49 | 0.71 | 1.94028E-05 | 0.000293565 |
| 65258 | *MPPE1* | ENSG00000154889.17 | 4.45 | 3.96 | -0.49 | 0.71 | 0.001670075 | 0.008422769 |
| 9486 | *CHST10* | ENSG00000115526.11 | 5.41 | 4.92 | -0.49 | 0.71 | 0.000273615 | 0.002094181 |
| 7915 | *ALDH5A1* | ENSG00000112294.14 | 5.42 | 4.93 | -0.49 | 0.71 | 0.000486454 | 0.00327114 |
| 7353 | *UFD1* | ENSG00000070010.19 | 6.95 | 6.46 | -0.49 | 0.71 | 4.10746E-05 | 0.000509916 |
| 494115 | *RBMXL1* | ENSG00000213516.10 | 5.46 | 4.97 | -0.49 | 0.71 | 0.000331696 | 0.002420492 |
| 7580 | *ZNF32* | ENSG00000169740.14 | 3.91 | 3.42 | -0.49 | 0.71 | 0.001144228 | 0.006344188 |
| 715 | *C1R* | ENSG00000159403.18 | 5.63 | 5.14 | -0.49 | 0.71 | 7.00055E-05 | 0.000759185 |
| 342926 | *ZNF677* | ENSG00000197928.11 | 4.51 | 4.01 | -0.50 | 0.71 | 0.000664829 | 0.004142144 |
| 1983 | *EIF5* | ENSG00000100664.11 | 7.41 | 6.91 | -0.50 | 0.71 | 0.000132449 | 0.001204075 |
| 3437 | *IFIT3* | ENSG00000119917.15 | 4.41 | 3.92 | -0.50 | 0.71 | 0.000119038 | 0.001108336 |
| 2627 | *GATA6* | ENSG00000141448.11 | 4.46 | 3.96 | -0.50 | 0.71 | 0.001818574 | 0.009014195 |
| 55860 | *ACTR10* | ENSG00000131966.14 | 5.97 | 5.47 | -0.50 | 0.71 | 9.68932E-05 | 0.000958649 |
| 54962 | *TIPIN* | ENSG00000075131.10 | 3.73 | 3.23 | -0.50 | 0.71 | 0.000752357 | 0.004566167 |
| 3312 | *HSPA8* | ENSG00000109971.14 | 10.28 | 9.78 | -0.50 | 0.71 | 4.90665E-07 | 2.14705E-05 |
| 123920 | *CMTM3* | ENSG00000140931.20 | 4.26 | 3.76 | -0.50 | 0.71 | 0.001072443 | 0.006015057 |
| 27291 | *R3HCC1L* | ENSG00000166024.14 | 4.23 | 3.73 | -0.50 | 0.71 | 0.000516946 | 0.003431595 |
| 57458 | *TMCC3* | ENSG00000057704.13 | 5.00 | 4.50 | -0.50 | 0.71 | 0.001542108 | 0.007959958 |
| 388021 | *TMEM179* | ENSG00000258986.7 | 6.74 | 6.24 | -0.50 | 0.71 | 5.87837E-05 | 0.000664872 |
| 9380 | *GRHPR* | ENSG00000137106.18 | 7.31 | 6.81 | -0.50 | 0.71 | 3.49174E-06 | 8.42564E-05 |
| 83759 | *RBM4B* | ENSG00000173914.12 | 4.88 | 4.38 | -0.50 | 0.71 | 9.99977E-05 | 0.000984174 |
| 51092 | *SIDT2* | ENSG00000149577.16 | 4.62 | 4.12 | -0.50 | 0.71 | 0.001238864 | 0.006734556 |
| 64327 | *LMBR1* | ENSG00000105983.23 | 7.50 | 7.00 | -0.50 | 0.71 | 7.55103E-05 | 0.000800377 |
| 79096 | *C11orf49* | ENSG00000149179.14 | 5.28 | 4.78 | -0.50 | 0.71 | 0.000286349 | 0.00217281 |
| 348262 | *MCRIP1* | ENSG00000225663.8 | 5.10 | 4.60 | -0.50 | 0.71 | 3.72593E-05 | 0.00047678 |
| 672 | *BRCA1* | ENSG00000012048.23 | 6.17 | 5.67 | -0.50 | 0.71 | 6.89739E-05 | 0.000749109 |
| 5073 | *PARN* | ENSG00000140694.17 | 6.53 | 6.02 | -0.50 | 0.71 | 7.94498E-06 | 0.000154264 |
| 124222 | *PAQR4* | ENSG00000162073.14 | 6.63 | 6.13 | -0.50 | 0.71 | 1.45285E-06 | 4.45054E-05 |
| 1788 | *DNMT3A* | ENSG00000119772.19 | 5.65 | 5.15 | -0.50 | 0.71 | 5.09774E-05 | 0.0005964 |
| 6646 | *SOAT1* | ENSG00000057252.13 | 9.98 | 9.47 | -0.51 | 0.70 | 7.98191E-07 | 2.95476E-05 |
| 84187 | *TMEM164* | ENSG00000157600.12 | 6.76 | 6.26 | -0.51 | 0.70 | 0.00015651 | 0.001359407 |
| 57405 | *SPC25* | ENSG00000152253.9 | 4.52 | 4.02 | -0.51 | 0.70 | 0.000957727 | 0.005486461 |

| | | | | | | | | |
|---|---|---|---|---|---|---|---|---|
| 7343 | *UBTF* | ENSG00000108312.15 | 7.43 | 6.93 | -0.51 | 0.70 | 1.28189E-06 | 4.11109E-05 |
| 23233 | *EXOC6B* | ENSG00000144036.16 | 5.01 | 4.50 | -0.51 | 0.70 | 0.000891746 | 0.005189691 |
| 126792 | *B3GALT6* | ENSG00000176022.7 | 4.36 | 3.85 | -0.51 | 0.70 | 0.000799215 | 0.004773347 |
| 7903 | *ST8SIA4* | ENSG00000113532.13 | 5.48 | 4.98 | -0.51 | 0.70 | 0.000624255 | 0.003968424 |
| 1793 | *DOCK1* | ENSG00000150760.13 | 5.83 | 5.32 | -0.51 | 0.70 | 1.67734E-05 | 0.000265216 |
| 4171 | *MCM2* | ENSG00000073111.14 | 7.58 | 7.07 | -0.51 | 0.70 | 1.13261E-06 | 3.803E-05 |
| 5001 | *ORC5* | ENSG00000164815.11 | 5.09 | 4.59 | -0.51 | 0.70 | 2.61471E-05 | 0.000365157 |
| 10400 | *PEMT* | ENSG00000133027.18 | 3.52 | 3.01 | -0.51 | 0.70 | 0.00195106 | 0.009526176 |
| 91057 | *CCDC34* | ENSG00000109881.17 | 4.12 | 3.61 | -0.51 | 0.70 | 0.000985691 | 0.005629468 |
| 9947 | *MAGEC1* | ENSG00000155495.9 | 6.00 | 5.49 | -0.51 | 0.70 | 1.11619E-05 | 0.000199916 |
| 23635 | *SSBP2* | ENSG00000145687.17 | 5.09 | 4.58 | -0.51 | 0.70 | 0.000783305 | 0.004715854 |
| 11267 | *SNF8* | ENSG00000159210.10 | 5.73 | 5.22 | -0.51 | 0.70 | 8.34042E-05 | 0.000860945 |
| 100289097 | *FRG1CP* | ENSG00000282826.3 | 4.09 | 3.58 | -0.51 | 0.70 | 0.000218178 | 0.001756774 |
| 80208 | *SPG11* | ENSG00000104133.16 | 6.23 | 5.72 | -0.52 | 0.70 | 9.46808E-05 | 0.000944229 |
| 388969 | *C2orf68* | ENSG00000168887.11 | 5.43 | 4.91 | -0.52 | 0.70 | 8.4669E-05 | 0.000869218 |
| 65110 | *UPF3A* | ENSG00000169062.15 | 7.27 | 6.75 | -0.52 | 0.70 | 5.1781E-06 | 0.000112455 |
| 56848 | *SPHK2* | ENSG00000063176.16 | 4.77 | 4.25 | -0.52 | 0.70 | 6.39842E-05 | 0.000709785 |
| 373156 | *GSTK1* | ENSG00000197448.14 | 6.92 | 6.40 | -0.52 | 0.70 | 2.10528E-06 | 5.82622E-05 |
| 9577 | *BABAM2* | ENSG00000158019.21 | 4.76 | 4.24 | -0.52 | 0.70 | 9.64662E-05 | 0.000955684 |
| 5793 | *PTPRG* | ENSG00000144724.20 | 4.25 | 3.74 | -0.52 | 0.70 | 0.000771901 | 0.004664036 |
| 25941 | *TPGS2* | ENSG00000134779.15 | 6.62 | 6.11 | -0.52 | 0.70 | 3.21313E-05 | 0.000424524 |
| 27065 | *NSG1* | ENSG00000168824.15 | 5.39 | 4.87 | -0.52 | 0.70 | 0.000518356 | 0.003434234 |
| 1466 | *CSRP2* | ENSG00000175183.10 | 5.72 | 5.20 | -0.52 | 0.70 | 1.43085E-05 | 0.000238353 |
| 84629 | *TNRC18* | ENSG00000182095.15 | 6.13 | 5.61 | -0.52 | 0.70 | 0.000109883 | 0.001052482 |
| 83666 | *PARP9* | ENSG00000138496.17 | 5.87 | 5.35 | -0.52 | 0.70 | 8.37193E-05 | 0.000863549 |
| 585 | *BBS4* | ENSG00000140463.14 | 4.95 | 4.43 | -0.52 | 0.70 | 0.000131361 | 0.001197088 |
| 83989 | *FAM172A* | ENSG00000113391.19 | 4.53 | 4.01 | -0.52 | 0.70 | 0.001711916 | 0.008581407 |
| 9158 | *FIBP* | ENSG00000172500.13 | 5.41 | 4.89 | -0.52 | 0.70 | 0.000200127 | 0.001642843 |
| 22980 | *TCF25* | ENSG00000141002.20 | 6.10 | 5.58 | -0.52 | 0.70 | 0.00017274 | 0.001469761 |
| 7321 | *UBE2D1* | ENSG00000072401.15 | 4.94 | 4.41 | -0.52 | 0.70 | 0.000988404 | 0.00563638 |
| 84305 | *PYM1* | ENSG00000170473.17 | 5.16 | 4.64 | -0.53 | 0.69 | 0.000121001 | 0.001121748 |
| 51514 | *DTL* | ENSG00000143476.18 | 6.98 | 6.45 | -0.53 | 0.69 | 0.00079757 | 0.004769214 |
| 9202 | *ZMYM4* | ENSG00000146463.12 | 7.00 | 6.48 | -0.53 | 0.69 | 4.98859E-05 | 0.000588168 |
| 54361 | *WNT4* | ENSG00000162552.15 | 6.12 | 5.59 | -0.53 | 0.69 | 4.34175E-05 | 0.0005298 |
| 26121 | *PRPF31* | ENSG00000105618.14 | 6.08 | 5.55 | -0.53 | 0.69 | 1.57444E-05 | 0.000254642 |
| 5305 | *PIP4K2A* | ENSG00000150867.14 | 6.59 | 6.06 | -0.53 | 0.69 | 2.50441E-05 | 0.000355617 |
| 5591 | *PRKDC* | ENSG00000253729.8 | 8.58 | 8.05 | -0.53 | 0.69 | 0.000193059 | 0.001596482 |
| 4324 | *MMP15* | ENSG00000102996.5 | 5.17 | 4.64 | -0.53 | 0.69 | 0.000464922 | 0.003154616 |
| 375346 | *STIMATE* | ENSG00000213533.13 | 5.98 | 5.46 | -0.53 | 0.69 | 0.001652742 | 0.00835782 |
| 7626 | *ZNF75D* | ENSG00000186376.15 | 5.97 | 5.44 | -0.53 | 0.69 | 4.40236E-05 | 0.000534588 |
| 55703 | *POLR3B* | ENSG00000013503.10 | 5.26 | 4.73 | -0.53 | 0.69 | 0.001228463 | 0.006687702 |
| 728661 | *SLC35E2B* | ENSG00000189339.12 | 5.39 | 4.86 | -0.53 | 0.69 | 0.000133327 | 0.001210588 |
| 4897 | *NRCAM* | ENSG00000091129.22 | 7.05 | 6.52 | -0.53 | 0.69 | 0.000665113 | 0.00414219 |
| 54942 | *ABITRAM* | ENSG00000119328.12 | 4.55 | 4.02 | -0.53 | 0.69 | 0.001237704 | 0.006730685 |
| 1968 | *EIF2S3* | ENSG00000130741.11 | 7.12 | 6.58 | -0.54 | 0.69 | 3.57804E-06 | 8.57872E-05 |
| 643988 | *FNDC10* | ENSG00000228594.4 | 5.15 | 4.61 | -0.54 | 0.69 | 3.75163E-05 | 0.000478811 |
| 8209 | *GATD3* | ENSG00000160221.18 | 5.76 | 5.23 | -0.54 | 0.69 | 0.001469536 | 0.007674416 |
| 54931 | *TRMT10C* | ENSG00000174173.7 | 6.25 | 5.71 | -0.54 | 0.69 | 6.49974E-05 | 0.000718369 |
| 23229 | *ARHGEF9* | ENSG00000131089.17 | 4.79 | 4.25 | -0.54 | 0.69 | 0.000986551 | 0.005632233 |
| 728819 | *C1GALT1C1L* | ENSG00000223658.8 | 3.39 | 2.85 | -0.54 | 0.69 | 0.000511104 | 0.003395822 |
| 9851 | *KIAA0753* | ENSG00000198920.11 | 4.11 | 3.57 | -0.54 | 0.69 | 0.001809749 | 0.008978626 |
| 9897 | *WASHC5* | ENSG00000164961.16 | 6.83 | 6.29 | -0.54 | 0.69 | 3.59254E-06 | 8.58604E-05 |
| 80142 | *PTGES2* | ENSG00000148334.16 | 5.18 | 4.64 | -0.54 | 0.69 | 0.002037488 | 0.009848845 |
| 124637 | *CYB5D1* | ENSG00000182224.12 | 4.68 | 4.14 | -0.54 | 0.69 | 1.90223E-05 | 0.00028956 |
| 80742 | *PRR3* | ENSG00000204576.12 | 5.40 | 4.86 | -0.54 | 0.69 | 2.22031E-05 | 0.000326073 |
| 2937 | *GSS* | ENSG00000100983.12 | 5.63 | 5.09 | -0.54 | 0.69 | 2.89486E-05 | 0.000392726 |
| 23089 | *PEG10* | ENSG00000242265.6 | 10.80 | 10.27 | -0.54 | 0.69 | 2.45461E-05 | 0.00034987 |
| 9830 | *TRIM14* | ENSG00000106785.15 | 5.94 | 5.40 | -0.54 | 0.69 | 5.13333E-06 | 0.000111823 |
| 2593 | *GAMT* | ENSG00000130005.13 | 4.78 | 4.23 | -0.54 | 0.69 | 0.000270188 | 0.0020743 |
| 23452 | *ANGPTL2* | ENSG00000136859.10 | 5.91 | 5.36 | -0.54 | 0.69 | 0.00027505 | 0.002103019 |
| 55298 | *RNF121* | ENSG00000137522.18 | 5.28 | 4.74 | -0.54 | 0.69 | 0.001058791 | 0.005956296 |
| 7425 | *VGF* | ENSG00000128564.8 | 6.35 | 5.81 | -0.54 | 0.69 | 5.43777E-05 | 0.000626847 |
| 6603 | *SMARCD2* | ENSG00000108604.17 | 5.61 | 5.07 | -0.54 | 0.69 | 0.000121921 | 0.001129578 |
| 55833 | *UBAP2* | ENSG00000137073.24 | 6.33 | 5.79 | -0.54 | 0.69 | 5.59402E-05 | 0.000640432 |
| 26999 | *CYFIP2* | ENSG00000055163.20 | 6.28 | 5.74 | -0.54 | 0.69 | 0.00022795 | 0.001818876 |
| 24148 | *PRPF6* | ENSG00000101161.8 | 5.79 | 5.25 | -0.54 | 0.69 | 2.90785E-06 | 7.36186E-05 |
| 55572 | *FOXRED1* | ENSG00000110074.12 | 4.67 | 4.12 | -0.54 | 0.69 | 1.15562E-05 | 0.000204296 |
| 93210 | *PGAP3* | ENSG00000161395.14 | 4.94 | 4.40 | -0.54 | 0.69 | 0.000225516 | 0.001804249 |
| 51157 | *ZNF580* | ENSG00000213015.9 | 3.90 | 3.36 | -0.54 | 0.69 | 0.000930195 | 0.005373865 |
| 56950 | *SMYD2* | ENSG00000143499.14 | 4.79 | 4.25 | -0.54 | 0.69 | 0.000467054 | 0.003163361 |
| 7629 | *ZNF76* | ENSG00000065029.15 | 4.97 | 4.42 | -0.55 | 0.69 | 8.55711E-05 | 0.000874899 |
| 10518 | *CIB2* | ENSG00000136425.14 | 3.49 | 2.95 | -0.55 | 0.68 | 0.000704804 | 0.004336583 |
| 5933 | *RBL1* | ENSG00000080839.12 | 6.06 | 5.52 | -0.55 | 0.68 | 0.001997364 | 0.009698618 |
| 776 | *CACNA1D* | ENSG00000157388.20 | 6.91 | 6.36 | -0.55 | 0.68 | 6.12741E-05 | 0.000685804 |
| 644873 | *LINC01184* | ENSG00000245937.9 | 4.15 | 3.60 | -0.55 | 0.68 | 0.000375785 | 0.002678135 |
| 108783654 | *HSD17B1-AS1* | ENSG00000266962.2 | 3.71 | 3.17 | -0.55 | 0.68 | 0.000867811 | 0.005068085 |
| 9836 | *LCMT2* | ENSG00000168806.8 | 4.73 | 4.18 | -0.55 | 0.68 | 8.13725E-05 | 0.000846402 |
| 11164 | *NUDT5* | ENSG00000165609.13 | 5.68 | 5.13 | -0.55 | 0.68 | 1.5849E-05 | 0.000255507 |
| 51659 | *GINS2* | ENSG00000131153.9 | 5.89 | 5.34 | -0.55 | 0.68 | 2.41553E-05 | 0.000345941 |
| 203522 | *INTS6L* | ENSG00000165359.16 | 4.20 | 3.66 | -0.55 | 0.68 | 0.000320169 | 0.002369178 |
| 53 | *ACP2* | ENSG00000134575.13 | 6.07 | 5.52 | -0.55 | 0.68 | 0.000104928 | 0.001017354 |
| 339229 | *OXLD1* | ENSG00000204237.5 | 3.62 | 3.07 | -0.55 | 0.68 | 0.000293906 | 0.002221166 |
| 55269 | *PSPC1* | ENSG00000121390.19 | 5.82 | 5.27 | -0.55 | 0.68 | 9.42116E-06 | 0.000176092 |
| 57608 | *JCAD* | ENSG00000165757.9 | 4.09 | 3.54 | -0.55 | 0.68 | 0.000105551 | 0.001022073 |
| 9555 | *MACROH2A1* | ENSG00000113648.17 | 7.94 | 7.39 | -0.55 | 0.68 | 1.46568E-06 | 4.46466E-05 |
| 80152 | *CENPT* | ENSG00000102901.13 | 4.25 | 3.70 | -0.55 | 0.68 | 9.7582E-05 | 0.000964193 |
| 2521 | *FUS* | ENSG00000089280.19 | 9.19 | 8.64 | -0.55 | 0.68 | 1.36323E-06 | 4.32572E-05 |

| | | | | | | | | |
|---|---|---|---|---|---|---|---|---|
| 55796 | *MBNL3* | ENSG00000076770.15 | 6.99 | 6.44 | -0.55 | 0.68 | 8.75728E-05 | 0.000889989 |
| 3551 | *IKBKB* | ENSG00000104365.16 | 5.12 | 4.56 | -0.55 | 0.68 | 7.36413E-06 | 0.00014505 |
| 6904 | *TBCD* | ENSG00000141556.22 | 6.20 | 5.65 | -0.55 | 0.68 | 2.10959E-05 | 0.000313705 |
| 113419 | *TEX261* | ENSG00000144043.12 | 6.48 | 5.92 | -0.56 | 0.68 | 8.03282E-05 | 0.000839206 |
| 93183 | *PIGM* | ENSG00000143315.8 | 6.26 | 5.70 | -0.56 | 0.68 | 6.96332E-06 | 0.000139454 |
| 503637 | *DUXAP8* | ENSG00000206195.11 | 6.19 | 5.63 | -0.56 | 0.68 | 2.25254E-05 | 0.000329151 |
| 88745 | *RRP36* | ENSG00000124541.7 | 5.32 | 4.76 | -0.56 | 0.68 | 1.37693E-05 | 0.000233254 |
| 83637 | *ZMIZ2* | ENSG00000122515.16 | 6.81 | 6.25 | -0.56 | 0.68 | 4.91072E-05 | 0.000582188 |
| 51209 | *RAB9B* | ENSG00000123570.4 | 3.86 | 3.30 | -0.56 | 0.68 | 0.000660605 | 0.004121845 |
| 348110 | *ARPIN* | ENSG00000242498.8 | 5.65 | 5.08 | -0.56 | 0.68 | 0.001121384 | 0.006248932 |
| 412 | *STS* | ENSG00000101846.9 | 4.13 | 3.57 | -0.56 | 0.68 | 0.000393429 | 0.002784052 |
| 159371 | *SLC35G1* | ENSG00000176273.15 | 3.92 | 3.35 | -0.56 | 0.68 | 0.000867193 | 0.005066443 |
| 100506668 | *NRAV* | ENSG00000248008.4 | 3.97 | 3.41 | -0.56 | 0.68 | 4.8999E-05 | 0.000581452 |
| 83990 | *BRIP1* | ENSG00000136492.10 | 5.64 | 5.08 | -0.56 | 0.68 | 3.11237E-05 | 0.000414863 |
| 10717 | *AP4B1* | ENSG00000134262.13 | 4.07 | 3.50 | -0.56 | 0.68 | 9.73395E-05 | 0.000962429 |
| 1870 | *E2F2* | ENSG00000007968.7 | 5.12 | 4.56 | -0.56 | 0.68 | 0.000303064 | 0.002276621 |
| 1718 | *DHCR24* | ENSG00000116133.13 | 9.44 | 8.87 | -0.57 | 0.68 | 3.90221E-07 | 1.82456E-05 |
| 79084 | *WDR77* | ENSG00000116455.14 | 5.71 | 5.14 | -0.57 | 0.68 | 8.31111E-05 | 0.000859101 |
| 57794 | *SUGP1* | ENSG00000105705.16 | 5.18 | 4.62 | -0.57 | 0.68 | 0.001089991 | 0.006097533 |
| 2639 | *GCDH* | ENSG00000105607.13 | 4.89 | 4.32 | -0.57 | 0.68 | 0.000454662 | 0.003108895 |
| 435 | *ASL* | ENSG00000126522.17 | 4.00 | 3.43 | -0.57 | 0.67 | 0.001054797 | 0.005938276 |
| 55506 | *MACROH2A2* | ENSG00000099284.15 | 5.81 | 5.24 | -0.57 | 0.67 | 0.00024011 | 0.001892761 |
| 130535 | *KCTD18* | ENSG00000155729.13 | 3.42 | 2.85 | -0.57 | 0.67 | 0.000719471 | 0.004403975 |
| 283131 | *NEAT1* | ENSG00000245532.10 | 5.35 | 4.78 | -0.57 | 0.67 | 0.001118119 | 0.006236289 |
| 54556 | *ING3* | ENSG00000071243.16 | 4.98 | 4.40 | -0.57 | 0.67 | 1.51962E-05 | 0.000247913 |
| 2235 | *FECH* | ENSG00000066926.13 | 6.42 | 5.84 | -0.57 | 0.67 | 4.48767E-05 | 0.000544067 |
| 56731 | *SLC2A4RG* | ENSG00000125520.14 | 5.02 | 4.45 | -0.57 | 0.67 | 0.000297289 | 0.002242219 |
| 286077 | *FAM83H* | ENSG00000180921.7 | 4.94 | 4.37 | -0.57 | 0.67 | 1.15512E-05 | 0.000204296 |
| 9899 | *SV2B* | ENSG00000185518.12 | 6.54 | 5.97 | -0.57 | 0.67 | 3.42381E-06 | 8.30429E-05 |
| 445 | *ASS1* | ENSG00000130707.18 | 5.03 | 4.45 | -0.58 | 0.67 | 0.000856519 | 0.005022179 |
| 81928 | *CABLES2* | ENSG00000149679.12 | 3.95 | 3.37 | -0.58 | 0.67 | 0.000823048 | 0.004874949 |
| 374618 | *TEX9* | ENSG00000151575.14 | 3.65 | 3.07 | -0.58 | 0.67 | 0.000274398 | 0.002099101 |
| 121642 | *ALKBH2* | ENSG00000189046.11 | 3.32 | 2.74 | -0.58 | 0.67 | 0.001221088 | 0.006652379 |
| 114785 | *MBD6* | ENSG00000166987.15 | 6.21 | 5.63 | -0.58 | 0.67 | 1.17707E-05 | 0.000207111 |
| 64976 | *MRPL40* | ENSG00000185608.9 | 5.45 | 4.88 | -0.58 | 0.67 | 2.61539E-05 | 0.000365157 |
| 987 | *LRBA* | ENSG00000198589.14 | 6.62 | 6.05 | -0.58 | 0.67 | 0.000178136 | 0.001497004 |
| 55709 | *KBTBD4* | ENSG00000123444.14 | 5.28 | 4.70 | -0.58 | 0.67 | 7.31515E-05 | 0.000782559 |
| 79622 | *SNRNP25* | ENSG00000161981.11 | 5.56 | 4.98 | -0.58 | 0.67 | 7.16436E-06 | 0.000141562 |
| 9076 | *CLDN1* | ENSG00000163347.6 | 9.04 | 8.47 | -0.58 | 0.67 | 3.91822E-07 | 1.82635E-05 |
| 92017 | *SNX29* | ENSG00000048471.14 | 5.27 | 4.69 | -0.58 | 0.67 | 0.000381869 | 0.002711197 |
| 440465 | *BAIAP2-DT* | ENSG00000226137.5 | 5.16 | 4.58 | -0.58 | 0.67 | 4.54567E-05 | 0.000548593 |
| 2176 | *FANCC* | ENSG00000158169.13 | 5.30 | 4.72 | -0.58 | 0.67 | 0.001173981 | 0.006454315 |
| 79643 | *CHMP6* | ENSG00000176108.9 | 4.31 | 3.73 | -0.58 | 0.67 | 0.000130207 | 0.00118873 |
| 5588 | *PRKCQ* | ENSG00000065675.16 | 3.90 | 3.32 | -0.58 | 0.67 | 7.88918E-05 | 0.000829193 |
| 113455421 | *DERPC* | ENSG00000286140.2 | 5.58 | 5.00 | -0.58 | 0.67 | 0.001889271 | 0.009284893 |
| 112487 | *DTD2* | ENSG00000129480.13 | 4.32 | 3.74 | -0.58 | 0.67 | 0.000329648 | 0.002412421 |
| 3936 | *LCP1* | ENSG00000136167.15 | 5.43 | 4.85 | -0.58 | 0.67 | 7.04229E-05 | 0.00076206 |
| 1719 | *DHFR* | ENSG00000228716.7 | 8.06 | 7.48 | -0.58 | 0.67 | 1.61642E-06 | 4.75972E-05 |
| 7174 | *TPP2* | ENSG00000134900.12 | 6.86 | 6.27 | -0.59 | 0.67 | 2.75081E-06 | 7.10565E-05 |
| 9344 | *TAOK2* | ENSG00000149930.18 | 6.49 | 5.90 | -0.59 | 0.67 | 1.56586E-05 | 0.000253886 |
| 84263 | *HSDL2* | ENSG00000119471.15 | 7.74 | 7.16 | -0.59 | 0.67 | 5.20641E-07 | 2.24268E-05 |
| 57469 | *PNMA8B* | ENSG00000204851.7 | 5.17 | 4.58 | -0.59 | 0.67 | 4.13655E-05 | 0.000512256 |
| 79184 | *BRCC3* | ENSG00000185515.16 | 6.14 | 5.55 | -0.59 | 0.66 | 5.60278E-06 | 0.000119348 |
| 27143 | *PALD1* | ENSG00000107719.9 | 3.89 | 3.30 | -0.59 | 0.66 | 0.000175403 | 0.001487357 |
| 78991 | *PCYOX1L* | ENSG00000145882.11 | 4.74 | 4.15 | -0.59 | 0.66 | 0.000128812 | 0.001178148 |
| 5331 | *PLCB3* | ENSG00000149782.13 | 5.05 | 4.45 | -0.59 | 0.66 | 0.001478814 | 0.007709457 |
| 80823 | *BHLHB9* | ENSG00000198908.12 | 4.02 | 3.42 | -0.59 | 0.66 | 4.94442E-05 | 0.000584023 |
| 6822 | *SULT2A1* | ENSG00000105398.4 | 7.66 | 7.07 | -0.59 | 0.66 | 5.30311E-07 | 2.26764E-05 |
| 84456 | *L3MBTL3* | ENSG00000198945.8 | 5.93 | 5.33 | -0.60 | 0.66 | 8.55455E-06 | 0.00016377 |
| 5325 | *PLAGL1* | ENSG00000118495.21 | 8.22 | 7.62 | -0.60 | 0.66 | 3.5216E-06 | 8.45691E-05 |
| 66008 | *TRAK2* | ENSG00000115993.13 | 6.19 | 5.59 | -0.60 | 0.66 | 4.69158E-05 | 0.000562877 |
| 79968 | *WDR76* | ENSG00000092470.12 | 6.30 | 5.70 | -0.60 | 0.66 | 3.60544E-05 | 0.000466099 |
| 51062 | *ATL1* | ENSG00000198513.14 | 4.74 | 4.14 | -0.60 | 0.66 | 0.000204284 | 0.001668169 |
| 26873 | *OPLAH* | ENSG00000178814.17 | 3.81 | 3.21 | -0.60 | 0.66 | 0.001067539 | 0.005998567 |
| 54621 | *VSIG10* | ENSG00000176834.14 | 5.40 | 4.79 | -0.60 | 0.66 | 9.15807E-05 | 0.000922507 |
| 6920 | *TCEA3* | ENSG00000204219.11 | 6.21 | 5.60 | -0.60 | 0.66 | 1.14816E-05 | 0.000203457 |
| 219854 | *TMEM218* | ENSG00000150433.10 | 3.80 | 3.20 | -0.61 | 0.66 | 0.000609509 | 0.003906116 |
| 22859 | *ADGRL1* | ENSG00000072071.17 | 6.41 | 5.80 | -0.61 | 0.66 | 0.000183557 | 0.001529707 |
| 6426 | *SRSF1* | ENSG00000136450.13 | 8.55 | 7.94 | -0.61 | 0.66 | 2.23932E-07 | 1.2588E-05 |
| 2850 | *GPR27* | ENSG00000170837.3 | 3.61 | 3.01 | -0.61 | 0.66 | 0.000688723 | 0.004255678 |
| 90957 | *DHX57* | ENSG00000163214.21 | 5.99 | 5.38 | -0.61 | 0.66 | 1.74953E-05 | 0.000271705 |
| 645323 | *LINC00461* | ENSG00000245526.12 | 6.09 | 5.49 | -0.61 | 0.66 | 0.000147581 | 0.00130527 |
| 79862 | *ZNF669* | ENSG00000188295.15 | 3.18 | 2.57 | -0.61 | 0.66 | 0.001535996 | 0.007933991 |
| 4144 | *MAT2A* | ENSG00000168906.13 | 7.23 | 6.62 | -0.61 | 0.66 | 3.65324E-05 | 0.000471466 |
| 79183 | *TTPAL* | ENSG00000124120.11 | 5.14 | 4.53 | -0.61 | 0.66 | 8.41476E-05 | 0.000865642 |
| 55644 | *OSGEP* | ENSG00000092094.11 | 3.99 | 3.38 | -0.61 | 0.65 | 0.000852829 | 0.005009827 |
| 79624 | *ARMT1* | ENSG00000146476.11 | 7.16 | 6.55 | -0.61 | 0.65 | 6.30384E-07 | 2.52979E-05 |
| 6522 | *SLC4A2* | ENSG00000164889.15 | 7.39 | 6.78 | -0.61 | 0.65 | 5.85498E-06 | 0.000122392 |
| 64847 | *SPATA20* | ENSG00000006282.21 | 5.78 | 5.17 | -0.61 | 0.65 | 0.000103027 | 0.001006037 |
| 402665 | *IGLON5* | ENSG00000142549.10 | 5.89 | 5.28 | -0.61 | 0.65 | 8.02251E-07 | 2.95476E-05 |
| 11282 | *MGAT4B* | ENSG00000161013.17 | 6.10 | 5.49 | -0.62 | 0.65 | 7.72036E-05 | 0.00081602 |
| 9208 | *LRRFIP1* | ENSG00000124831.19 | 4.43 | 3.82 | -0.62 | 0.65 | 5.09815E-05 | 0.0005964 |
| 10654 | *PMVK* | ENSG00000163344.6 | 4.51 | 3.90 | -0.62 | 0.65 | 8.98176E-05 | 0.000907182 |
| 4670 | *HNRNPM* | ENSG00000099783.12 | 8.38 | 7.76 | -0.62 | 0.65 | 6.09182E-07 | 2.47926E-05 |
| 9774 | *BCLAF1* | ENSG00000029363.17 | 8.59 | 7.97 | -0.62 | 0.65 | 8.72829E-07 | 3.13404E-05 |
| 57610 | *RANBP10* | ENSG00000141084.12 | 4.75 | 4.13 | -0.62 | 0.65 | 8.77286E-06 | 0.000167522 |
| 23169 | *SLC35D1* | ENSG00000116704.8 | 7.13 | 6.51 | -0.62 | 0.65 | 3.29922E-06 | 8.08664E-05 |

| | | | | | | | | |
|---|---|---|---|---|---|---|---|---|
| 7733 | *ZNF180* | ENSG00000167384.11 | 3.53 | 2.91 | -0.62 | 0.65 | 0.000238067 | 0.001882584 |
| 55384 | *MEG3* | ENSG00000214548.18 | 7.70 | 7.08 | -0.62 | 0.65 | 0.000647806 | 0.004073281 |
| 151613 | *TTC14* | ENSG00000163728.11 | 4.59 | 3.97 | -0.62 | 0.65 | 0.000195348 | 0.001611863 |
| 65123 | *INTS3* | ENSG00000143624.14 | 6.63 | 6.00 | -0.62 | 0.65 | 1.34078E-06 | 4.28027E-05 |
| 63920 | *ZBED8* | ENSG00000221886.4 | 3.80 | 3.17 | -0.62 | 0.65 | 0.000129971 | 0.0011873 |
| 23542 | *MAPK8IP2* | ENSG00000008735.14 | 3.76 | 3.13 | -0.63 | 0.65 | 0.000349391 | 0.002530893 |
| 7390 | *UROS* | ENSG00000188690.15 | 5.42 | 4.79 | -0.63 | 0.65 | 8.95859E-06 | 0.000170423 |
| 4931 | *NVL* | ENSG00000143748.18 | 4.76 | 4.13 | -0.63 | 0.65 | 0.001372793 | 0.007272942 |
| 4311 | *MME* | ENSG00000196549.13 | 9.35 | 8.73 | -0.63 | 0.65 | 1.15979E-06 | 3.85971E-05 |
| 79813 | *EHMT1* | ENSG00000181090.21 | 7.08 | 6.46 | -0.63 | 0.65 | 0.000189571 | 0.001572077 |
| 55532 | *SLC30A10* | ENSG00000196660.11 | 8.00 | 7.37 | -0.63 | 0.65 | 1.05196E-06 | 3.60476E-05 |
| 57447 | *NDRG2* | ENSG00000165795.25 | 6.65 | 6.03 | -0.63 | 0.65 | 1.47332E-06 | 4.47632E-05 |
| 221883 | *HOXA11-AS* | ENSG00000240990.10 | 5.21 | 4.59 | -0.63 | 0.65 | 2.93742E-05 | 0.000397507 |
| 55027 | *HEATR3* | ENSG00000155393.14 | 5.10 | 4.47 | -0.63 | 0.65 | 0.000218906 | 0.001760749 |
| 55005 | *RMND1* | ENSG00000155906.20 | 4.60 | 3.97 | -0.63 | 0.65 | 0.000107901 | 0.001039465 |
| 101930085 | *HERPUD2-AS1* | ENSG00000271122.1 | 3.74 | 3.11 | -0.63 | 0.65 | 6.41372E-05 | 0.000710957 |
| 9894 | *TELO2* | ENSG00000100726.15 | 5.32 | 4.69 | -0.63 | 0.65 | 0.000190795 | 0.001579502 |
| 2934 | *GSN* | ENSG00000148180.21 | 5.73 | 5.10 | -0.63 | 0.65 | 1.38081E-05 | 0.000233491 |
| 875 | *CBS* | ENSG00000160200.18 | 4.03 | 3.39 | -0.63 | 0.65 | 0.000250445 | 0.001953243 |
| 10274 | *STAG1* | ENSG00000118007.13 | 6.54 | 5.91 | -0.63 | 0.65 | 3.3349E-07 | 1.61986E-05 |
| 8504 | *PEX3* | ENSG00000034693.15 | 4.76 | 4.13 | -0.64 | 0.64 | 4.14613E-06 | 9.55903E-05 |
| 10240 | *MRPS31* | ENSG00000102738.8 | 3.68 | 3.04 | -0.64 | 0.64 | 2.59616E-05 | 0.000364507 |
| 4200 | *ME2* | ENSG00000082212.13 | 5.43 | 4.80 | -0.64 | 0.64 | 6.84359E-06 | 0.000137962 |
| 54872 | *PIGG* | ENSG00000174227.16 | 6.51 | 5.87 | -0.64 | 0.64 | 3.21789E-06 | 7.95672E-05 |
| 84993 | *UBL7* | ENSG00000138629.16 | 5.95 | 5.31 | -0.64 | 0.64 | 2.82509E-05 | 0.000386878 |
| 100132074 | *FOXO6* | ENSG00000204060.8 | 3.60 | 2.96 | -0.64 | 0.64 | 0.000772233 | 0.004664162 |
| 2135 | *EXTL2* | ENSG00000162694.14 | 6.28 | 5.64 | -0.64 | 0.64 | 3.30054E-06 | 8.08664E-05 |
| 81790 | *RNF170* | ENSG00000120925.16 | 4.15 | 3.51 | -0.64 | 0.64 | 0.001154048 | 0.006372741 |
| 6695 | *SPOCK1* | ENSG00000152377.14 | 7.02 | 6.38 | -0.64 | 0.64 | 1.41074E-06 | 4.38481E-05 |
| 284613 | *CYB561D1* | ENSG00000174151.15 | 4.49 | 3.84 | -0.65 | 0.64 | 0.001203557 | 0.006581927 |
| 10752 | *CHL1* | ENSG00000134121.10 | 5.56 | 4.91 | -0.65 | 0.64 | 0.000692561 | 0.004274114 |
| 55701 | *ARHGEF40* | ENSG00000165801.10 | 6.43 | 5.78 | -0.65 | 0.64 | 3.66856E-05 | 0.000473036 |
| 23187 | *PHLDB1* | ENSG00000019144.20 | 6.26 | 5.61 | -0.65 | 0.64 | 4.21534E-05 | 0.000517741 |
| 726 | *CAPN5* | ENSG00000149260.18 | 4.83 | 4.18 | -0.65 | 0.64 | 2.44726E-05 | 0.000349154 |
| 5892 | *RAD51D* | ENSG00000185379.21 | 5.06 | 4.42 | -0.65 | 0.64 | 3.42705E-05 | 0.00044611 |
| 26149 | *ZNF658* | ENSG00000274349.5 | 3.94 | 3.29 | -0.65 | 0.64 | 0.000244081 | 0.00191323 |
| 23228 | *PLCL2* | ENSG00000154822.18 | 7.91 | 7.26 | -0.65 | 0.64 | 5.59721E-08 | 4.8503E-06 |
| 147947 | *ZNF542P* | ENSG00000240225.10 | 4.03 | 3.38 | -0.65 | 0.64 | 0.000509152 | 0.00338769 |
| 57628 | *DPP10* | ENSG00000175497.17 | 7.08 | 6.43 | -0.65 | 0.64 | 2.19412E-06 | 5.99847E-05 |
| 124961 | *ZFP3* | ENSG00000180787.6 | 4.64 | 3.99 | -0.65 | 0.64 | 0.000163435 | 0.001402511 |
| 55084 | *SOBP* | ENSG00000112320.12 | 4.23 | 3.58 | -0.65 | 0.64 | 0.000324543 | 0.002387777 |
| 90806 | *ANGEL2* | ENSG00000174606.14 | 5.53 | 4.89 | -0.65 | 0.64 | 3.13243E-05 | 0.000416427 |
| 9609 | *RAB36* | ENSG00000100228.14 | 6.30 | 5.65 | -0.65 | 0.64 | 1.06326E-05 | 0.000192969 |
| 139285 | *AMER1* | ENSG00000184675.11 | 6.09 | 5.44 | -0.65 | 0.64 | 1.40173E-06 | 4.37392E-05 |
| 2798 | *GNRHR* | ENSG00000109163.7 | 5.60 | 4.95 | -0.65 | 0.64 | 1.1444E-06 | 3.83399E-05 |
| 26521 | *TIMM8B* | ENSG00000150779.12 | 5.52 | 4.87 | -0.65 | 0.64 | 4.26986E-06 | 9.78418E-05 |
| 80095 | *ZNF606* | ENSG00000166704.13 | 3.25 | 2.59 | -0.65 | 0.64 | 0.000320947 | 0.002371784 |
| 1945 | *EFNA4* | ENSG00000243364.8 | 5.56 | 4.90 | -0.65 | 0.64 | 5.72852E-06 | 0.000120927 |
| 1660 | *DHX9* | ENSG00000135829.17 | 8.82 | 8.17 | -0.65 | 0.64 | 6.69934E-08 | 5.55527E-06 |
| 58985 | *IL22RA1* | ENSG00000142677.4 | 3.00 | 2.35 | -0.65 | 0.64 | 0.001834844 | 0.009076854 |
| 64789 | *EXO5* | ENSG00000164002.12 | 3.64 | 2.99 | -0.65 | 0.64 | 0.000177894 | 0.001496644 |
| 83543 | *AIF1L* | ENSG00000126878.13 | 8.15 | 7.49 | -0.65 | 0.64 | 2.96244E-06 | 7.47038E-05 |
| 285362 | *SUMF1* | ENSG00000144455.14 | 4.12 | 3.47 | -0.65 | 0.64 | 7.6775E-05 | 0.000812062 |
| 9886 | *RHOBTB1* | ENSG00000072422.17 | 6.38 | 5.72 | -0.66 | 0.63 | 0.000166853 | 0.001426953 |
| 114132 | *SIGLEC11* | ENSG00000161640.15 | 6.63 | 5.98 | -0.66 | 0.63 | 4.96916E-06 | 0.000109096 |
| 4820 | *NKTR* | ENSG00000114857.19 | 5.84 | 5.18 | -0.66 | 0.63 | 2.89658E-05 | 0.000392726 |
| 340526 | *RTL5* | ENSG00000242732.4 | 5.61 | 4.95 | -0.66 | 0.63 | 9.48723E-06 | 0.000176887 |
| 29882 | *ANAPC2* | ENSG00000176248.9 | 5.33 | 4.67 | -0.66 | 0.63 | 3.43749E-05 | 0.000447082 |
| 1468 | *SLC25A10* | ENSG00000183048.12 | 4.80 | 4.14 | -0.66 | 0.63 | 4.01621E-05 | 0.000503586 |
| 3065 | *HDAC1* | ENSG00000116478.12 | 6.97 | 6.31 | -0.66 | 0.63 | 1.38165E-06 | 4.3501E-05 |
| 3482 | *IGF2R* | ENSG00000197081.16 | 8.29 | 7.63 | -0.66 | 0.63 | 9.33485E-05 | 0.000933423 |
| 55971 | *BAIAP2L1* | ENSG00000006453.14 | 6.28 | 5.62 | -0.66 | 0.63 | 3.6869E-06 | 8.75581E-05 |
| 3623 | *INHA* | ENSG00000123999.5 | 2.72 | 2.06 | -0.66 | 0.63 | 0.001123862 | 0.006259015 |
| 129642 | *MBOAT2* | ENSG00000143797.12 | 4.52 | 3.85 | -0.66 | 0.63 | 0.000827218 | 0.004893859 |
| 23613 | *ZMYND8* | ENSG00000101040.20 | 6.99 | 6.32 | -0.66 | 0.63 | 4.26151E-05 | 0.00052213 |
| 200895 | *DHFR2* | ENSG00000178700.9 | 4.13 | 3.47 | -0.66 | 0.63 | 4.81878E-05 | 0.000573553 |
| 2157 | *F8* | ENSG00000185010.15 | 4.31 | 3.65 | -0.66 | 0.63 | 0.000933824 | 0.005388604 |
| 5046 | *PCSK6* | ENSG00000140479.18 | 5.70 | 5.03 | -0.66 | 0.63 | 1.31223E-05 | 0.000224831 |
| 113675 | *SDSL* | ENSG00000139410.15 | 3.70 | 3.03 | -0.67 | 0.63 | 7.31155E-05 | 0.000782559 |
| 5257 | *PHKB* | ENSG00000102893.16 | 7.45 | 6.78 | -0.67 | 0.63 | 2.61694E-06 | 6.84626E-05 |
| 138428 | *PTRH1* | ENSG00000187024.15 | 3.19 | 2.53 | -0.67 | 0.63 | 0.000807331 | 0.004810331 |
| 56896 | *DPYSL5* | ENSG00000157851.17 | 5.54 | 4.88 | -0.67 | 0.63 | 0.001101286 | 0.006158423 |
| 25927 | *CNRIP1* | ENSG00000119865.9 | 2.90 | 2.23 | -0.67 | 0.63 | 0.001903026 | 0.009337205 |
| 3428 | *IFI16* | ENSG00000163565.20 | 5.85 | 5.18 | -0.67 | 0.63 | 5.58517E-06 | 0.000119243 |
| 10964 | *IFI44L* | ENSG00000137959.17 | 4.21 | 3.54 | -0.67 | 0.63 | 0.000115656 | 0.00108969 |
| 91975 | *ZNF300* | ENSG00000145908.13 | 3.78 | 3.10 | -0.68 | 0.63 | 1.72854E-05 | 0.000269754 |
| 755 | *CFAP410* | ENSG00000160226.16 | 4.44 | 3.76 | -0.68 | 0.63 | 0.000360323 | 0.002592561 |
| 9308 | *CD83* | ENSG00000112149.10 | 7.99 | 7.32 | -0.68 | 0.63 | 4.06765E-08 | 3.9069E-06 |
| 9557 | *CHD1L* | ENSG00000131778.20 | 5.63 | 4.95 | -0.68 | 0.63 | 2.99825E-06 | 7.52521E-05 |
| 286101 | *ZNF252P* | ENSG00000196922.11 | 4.48 | 3.80 | -0.68 | 0.62 | 0.000518083 | 0.003434234 |
| 79587 | *CARS2* | ENSG00000134905.17 | 5.44 | 4.76 | -0.68 | 0.62 | 4.97524E-06 | 0.000109096 |
| 1666 | *DECR1* | ENSG00000104325.7 | 6.10 | 5.41 | -0.68 | 0.62 | 1.86173E-06 | 5.31231E-05 |
| 51313 | *GASK1B* | ENSG00000164125.16 | 6.34 | 5.66 | -0.68 | 0.62 | 1.55261E-05 | 0.000252471 |
| 5836 | *PYGL* | ENSG00000100504.17 | 5.22 | 4.53 | -0.69 | 0.62 | 1.15048E-06 | 3.84578E-05 |
| 10165 | *SLC25A13* | ENSG00000004864.14 | 6.22 | 5.53 | -0.69 | 0.62 | 3.99367E-06 | 9.3076E-05 |
| 11022 | *TDRKH* | ENSG00000182134.17 | 4.20 | 3.51 | -0.69 | 0.62 | 0.00023052 | 0.001836448 |
| 2874 | *GPS2* | ENSG00000132522.16 | 5.45 | 4.76 | -0.69 | 0.62 | 9.02858E-06 | 0.000170796 |

| | | | | | | | | |
|---|---|---|---|---|---|---|---|---|
| 56926 | *NCLN* | ENSG00000125912.11 | 6.87 | 6.18 | -0.69 | 0.62 | 8.18523E-07 | 2.9891E-05 |
| 670 | *BPHL* | ENSG00000137274.13 | 3.67 | 2.98 | -0.69 | 0.62 | 0.000427424 | 0.002971379 |
| 64848 | *YTHDC2* | ENSG00000047188.16 | 5.04 | 4.35 | -0.69 | 0.62 | 0.000264532 | 0.002040265 |
| 25999 | *CLIP3* | ENSG00000105270.15 | 6.25 | 5.57 | -0.69 | 0.62 | 0.000307401 | 0.002299989 |
| 5314 | *PKHD1* | ENSG00000170927.15 | 2.81 | 2.12 | -0.69 | 0.62 | 0.001911569 | 0.009369505 |
| 401337 | *LINC01446* | ENSG00000205628.5 | 2.55 | 1.86 | -0.69 | 0.62 | 0.001693248 | 0.008516744 |
| 22881 | *ANKRD6* | ENSG00000135299.17 | 5.08 | 4.38 | -0.69 | 0.62 | 0.000849989 | 0.005002938 |
| 5587 | *PRKD1* | ENSG00000184304.17 | 4.48 | 3.79 | -0.69 | 0.62 | 0.000798333 | 0.004771877 |
| 8482 | *SEMA7A* | ENSG00000138623.10 | 6.59 | 5.89 | -0.69 | 0.62 | 3.33286E-07 | 1.61986E-05 |
| 10277 | *UBE4B* | ENSG00000130939.20 | 6.10 | 5.40 | -0.70 | 0.62 | 5.00199E-05 | 0.000588361 |
| 150726 | *FBXO41* | ENSG00000163013.12 | 4.44 | 3.74 | -0.70 | 0.62 | 0.000284099 | 0.002160101 |
| 6936 | *GCFC2* | ENSG00000005436.14 | 2.71 | 2.01 | -0.70 | 0.62 | 0.000233961 | 0.001860903 |
| 84859 | *LRCH3* | ENSG00000186001.14 | 5.36 | 4.67 | -0.70 | 0.62 | 0.00027888 | 0.002125808 |
| 388341 | *LRRC75A* | ENSG00000181350.12 | 4.89 | 4.19 | -0.70 | 0.62 | 1.79064E-05 | 0.000275649 |
| 50937 | *CDON* | ENSG00000064309.16 | 5.52 | 4.82 | -0.70 | 0.62 | 1.45508E-05 | 0.000240521 |
| 27148 | *STK36* | ENSG00000163482.12 | 3.93 | 3.23 | -0.70 | 0.62 | 0.000417147 | 0.002913429 |
| 29903 | *CCDC106* | ENSG00000173581.8 | 4.03 | 3.32 | -0.70 | 0.61 | 0.000152363 | 0.001336255 |
| 10785 | *WDR4* | ENSG00000160193.12 | 3.26 | 2.55 | -0.70 | 0.61 | 0.00058355 | 0.003776844 |
| 64718 | *UNKL* | ENSG00000059145.19 | 5.62 | 4.92 | -0.70 | 0.61 | 0.001075969 | 0.006030326 |
| 57728 | *WDR19* | ENSG00000157796.18 | 5.66 | 4.96 | -0.70 | 0.61 | 0.000202418 | 0.001657444 |
| 9479 | *MAPK8IP1* | ENSG00000121653.11 | 5.80 | 5.10 | -0.70 | 0.61 | 1.0196E-07 | 7.35729E-06 |
| 79634 | *SCRN3* | ENSG00000144306.15 | 4.21 | 3.50 | -0.71 | 0.61 | 0.000208108 | 0.001690198 |
| 5164 | *PDK2* | ENSG00000005882.12 | 5.79 | 5.08 | -0.71 | 0.61 | 0.000176068 | 0.001489629 |
| 27077 | *B9D1* | ENSG00000108641.20 | 4.69 | 3.99 | -0.71 | 0.61 | 4.63236E-05 | 0.000556662 |
| 10778 | *ZNF271P* | ENSG00000257267.4 | 5.24 | 4.53 | -0.71 | 0.61 | 2.80416E-06 | 7.18218E-05 |
| 56922 | *MCCC1* | ENSG00000078070.14 | 5.22 | 4.52 | -0.71 | 0.61 | 1.91325E-06 | 5.41812E-05 |
| 340348 | *TSPAN33* | ENSG00000158457.6 | 3.79 | 3.08 | -0.71 | 0.61 | 0.000679291 | 0.004206056 |
| 7507 | *XPA* | ENSG00000136936.11 | 4.01 | 3.30 | -0.71 | 0.61 | 0.000107128 | 0.001034675 |
| 84674 | *CARD6* | ENSG00000132357.14 | 5.21 | 4.50 | -0.71 | 0.61 | 9.36254E-06 | 0.000175215 |
| 5575 | *PRKAR1B* | ENSG00000188191.15 | 4.84 | 4.13 | -0.71 | 0.61 | 2.40381E-06 | 6.45417E-05 |
| 64427 | *TTC31* | ENSG00000115282.21 | 3.48 | 2.77 | -0.71 | 0.61 | 0.001369594 | 0.007260152 |
| 9681 | *DEPDC5* | ENSG00000100150.20 | 5.21 | 4.51 | -0.71 | 0.61 | 0.000176298 | 0.001490735 |
| 5152 | *PDE9A* | ENSG00000160191.18 | 4.74 | 4.03 | -0.71 | 0.61 | 3.24156E-05 | 0.000427127 |
| 134 | *ADORA1* | ENSG00000163485.17 | 3.06 | 2.34 | -0.71 | 0.61 | 0.001493586 | 0.007762198 |
| 7627 | *ZNF75A* | ENSG00000162086.16 | 4.10 | 3.39 | -0.71 | 0.61 | 0.001375308 | 0.007283695 |
| 55561 | *CDC42BPG* | ENSG00000171219.9 | 5.28 | 4.57 | -0.71 | 0.61 | 0.000405395 | 0.002846276 |
| 4292 | *MLH1* | ENSG00000076242.16 | 5.45 | 4.73 | -0.71 | 0.61 | 5.28699E-06 | 0.000113849 |
| 8558 | *CDK10* | ENSG00000185324.22 | 4.33 | 3.61 | -0.72 | 0.61 | 0.000410629 | 0.002876909 |
| 6604 | *SMARCD3* | ENSG00000082014.17 | 6.04 | 5.32 | -0.72 | 0.61 | 2.11837E-07 | 1.21819E-05 |
| 2184 | *FAH* | ENSG00000103876.14 | 4.50 | 3.79 | -0.72 | 0.61 | 8.08864E-05 | 0.000843658 |
| 11170 | *FAM107A* | ENSG00000168309.18 | 5.51 | 4.79 | -0.72 | 0.61 | 1.3432E-06 | 4.28027E-05 |
| 5936 | *RBM4* | ENSG00000173933.21 | 5.91 | 5.19 | -0.72 | 0.61 | 3.11997E-05 | 0.000415138 |
| 133121 | *ENPP6* | ENSG00000164303.11 | 3.28 | 2.56 | -0.72 | 0.61 | 0.000323589 | 0.002383096 |
| 2766 | *GMPR* | ENSG00000137198.10 | 4.00 | 3.28 | -0.72 | 0.61 | 5.94199E-05 | 0.000670551 |
| 25925 | *ZNF521* | ENSG00000198795.11 | 6.32 | 5.60 | -0.72 | 0.61 | 0.000154229 | 0.001345045 |
| 286527 | *TMSB15B* | ENSG00000269226.7 | 2.65 | 1.93 | -0.72 | 0.61 | 0.001405295 | 0.007408524 |
| 23649 | *POLA2* | ENSG00000014138.9 | 6.04 | 5.32 | -0.72 | 0.61 | 0.000429856 | 0.00298276 |
| 4810 | *NHS* | ENSG00000188158.17 | 5.32 | 4.60 | -0.72 | 0.61 | 5.69322E-06 | 0.000120351 |
| 1174 | *AP1S1* | ENSG00000106367.15 | 7.97 | 7.24 | -0.72 | 0.61 | 6.69799E-08 | 5.55527E-06 |
| 747 | *DAGLA* | ENSG00000134780.10 | 3.74 | 3.02 | -0.72 | 0.61 | 0.000298725 | 0.002250787 |
| 11146 | *GLMN* | ENSG00000174842.17 | 4.05 | 3.33 | -0.73 | 0.60 | 0.002065441 | 0.009948719 |
| 57727 | *NCOA5* | ENSG00000124160.12 | 6.46 | 5.74 | -0.73 | 0.60 | 2.44258E-06 | 6.52324E-05 |
| 5311 | *PKD2* | ENSG00000118762.8 | 5.21 | 4.49 | -0.73 | 0.60 | 1.00149E-05 | 0.000184434 |
| 2150 | *F2RL1* | ENSG00000164251.5 | 2.93 | 2.21 | -0.73 | 0.60 | 0.001401895 | 0.007397376 |
| 79810 | *PTCD2* | ENSG00000049883.15 | 4.77 | 4.04 | -0.73 | 0.60 | 4.7128E-05 | 0.000564971 |
| 1944 | *EFNA3* | ENSG00000143590.14 | 6.02 | 5.29 | -0.73 | 0.60 | 1.14236E-07 | 7.93779E-06 |
| 192683 | *SCAMP5* | ENSG00000198794.12 | 6.30 | 5.57 | -0.73 | 0.60 | 2.20118E-05 | 0.000323644 |
| 7412 | *VCAM1* | ENSG00000162692.12 | 7.49 | 6.76 | -0.73 | 0.60 | 1.65618E-07 | 1.02717E-05 |
| 25894 | *PLEKHG4* | ENSG00000196155.13 | 5.32 | 4.59 | -0.73 | 0.60 | 0.00043448 | 0.003005534 |
| 283104 | *SBF2-AS1* | ENSG00000246273.8 | 2.73 | 2.01 | -0.73 | 0.60 | 0.000546272 | 0.003578784 |
| 26268 | *FBXO9* | ENSG00000112146.17 | 6.33 | 5.60 | -0.73 | 0.60 | 2.01509E-06 | 5.65317E-05 |
| 5092 | *PCBD1* | ENSG00000166228.9 | 4.22 | 3.49 | -0.73 | 0.60 | 4.51501E-05 | 0.000546015 |
| 79641 | *ROGDI* | ENSG00000067836.13 | 4.03 | 3.30 | -0.73 | 0.60 | 0.000243402 | 0.001910678 |
| 91947 | *ARRDC4* | ENSG00000140450.9 | 5.45 | 4.72 | -0.73 | 0.60 | 9.34394E-07 | 3.29209E-05 |
| 29082 | *CHMP4A* | ENSG00000254505.11 | 4.03 | 3.29 | -0.73 | 0.60 | 5.09371E-05 | 0.0005964 |
| 10795 | *ZNF268* | ENSG00000090612.22 | 5.20 | 4.46 | -0.74 | 0.60 | 4.05138E-05 | 0.000506305 |
| 8677 | *STX10* | ENSG00000104915.15 | 3.93 | 3.20 | -0.74 | 0.60 | 0.000110001 | 0.001052936 |
| 1780 | *DYNC1I1* | ENSG00000158560.14 | 4.19 | 3.45 | -0.74 | 0.60 | 0.000722896 | 0.004417728 |
| 4065 | *LY75* | ENSG00000054219.11 | 5.50 | 4.76 | -0.74 | 0.60 | 1.78737E-05 | 0.000275428 |
| 79640 | *C22orf46* | ENSG00000184208.12 | 3.83 | 3.09 | -0.74 | 0.60 | 4.29602E-05 | 0.000525501 |
| 162282 | *ANKFN1* | ENSG00000153930.13 | 7.23 | 6.49 | -0.74 | 0.60 | 2.94327E-08 | 3.04659E-06 |
| 344595 | *DUBR* | ENSG00000243701.8 | 5.80 | 5.05 | -0.74 | 0.60 | 8.78524E-05 | 0.000891533 |
| 55622 | *TTC27* | ENSG00000018699.13 | 3.92 | 3.17 | -0.75 | 0.60 | 0.000986986 | 0.005632574 |
| 79736 | *TEFM* | ENSG00000172171.11 | 3.01 | 2.27 | -0.75 | 0.60 | 0.000189848 | 0.001573392 |
| 3070 | *HELLS* | ENSG00000119969.15 | 6.24 | 5.49 | -0.75 | 0.60 | 3.88628E-07 | 1.82456E-05 |
| 8850 | *KAT2B* | ENSG00000114166.8 | 4.43 | 3.68 | -0.75 | 0.60 | 7.20373E-05 | 0.000774384 |
| 29800 | *ZDHHC1* | ENSG00000159714.12 | 3.25 | 2.49 | -0.75 | 0.59 | 0.000601608 | 0.003863729 |
| 2845 | *GPR22* | ENSG00000172209.6 | 5.29 | 4.54 | -0.75 | 0.59 | 7.8682E-05 | 0.000827567 |
| 196740 | *VSTM4* | ENSG00000165633.13 | 6.62 | 5.87 | -0.75 | 0.59 | 7.59215E-08 | 6.02913E-06 |
| 65094 | *JMJD4* | ENSG00000081692.13 | 3.82 | 3.07 | -0.75 | 0.59 | 9.59438E-05 | 0.000951765 |
| 64943 | *NT5DC2* | ENSG00000168268.11 | 6.06 | 5.30 | -0.75 | 0.59 | 1.1396E-05 | 0.00020228 |
| 9941 | *EXOG* | ENSG00000157036.13 | 4.52 | 3.77 | -0.75 | 0.59 | 0.00131222 | 0.007044031 |
| 64925 | *CCDC71* | ENSG00000177352.10 | 4.31 | 3.56 | -0.75 | 0.59 | 5.42515E-06 | 0.000116157 |
| 10801 | *SEPTIN9* | ENSG00000184640.20 | 7.80 | 7.05 | -0.75 | 0.59 | 6.68488E-07 | 2.62046E-05 |
| 10439 | *OLFM1* | ENSG00000130558.20 | 5.12 | 4.37 | -0.75 | 0.59 | 7.13043E-07 | 2.72317E-05 |
| 27340 | *UTP20* | ENSG00000120800.5 | 6.23 | 5.47 | -0.76 | 0.59 | 2.38515E-05 | 0.000342899 |
| 375323 | *LHFPL4* | ENSG00000156959.9 | 3.96 | 3.21 | -0.76 | 0.59 | 0.000177423 | 0.001495197 |

| | | | | | | | | |
|---|---|---|---|---|---|---|---|---|
| 23532 | *PRAME* | ENSG00000185686.18 | 5.47 | 4.71 | -0.76 | 0.59 | 8.88722E-06 | 0.00016949 |
| 23313 | *KIAA0930* | ENSG00000100364.19 | 4.76 | 4.00 | -0.76 | 0.59 | 0.000132836 | 0.001206858 |
| 23708 | *GSPT2* | ENSG00000189369.9 | 2.57 | 1.81 | -0.76 | 0.59 | 0.000952121 | 0.005462685 |
| 9852 | *EPM2AIP1* | ENSG00000178567.8 | 6.77 | 6.01 | -0.76 | 0.59 | 9.44917E-06 | 0.000176396 |
| 378708 | *CENPS* | ENSG00000175279.22 | 2.87 | 2.11 | -0.76 | 0.59 | 0.00021714 | 0.001751241 |
| 129831 | *RBM45* | ENSG00000155636.15 | 3.80 | 3.04 | -0.76 | 0.59 | 0.000285709 | 0.002169049 |
| 222183 | *SRRM3* | ENSG00000177679.16 | 6.22 | 5.46 | -0.76 | 0.59 | 5.80976E-07 | 2.42219E-05 |
| 113763 | *ZBED6CL* | ENSG00000188707.6 | 3.72 | 2.96 | -0.76 | 0.59 | 0.00116559 | 0.006417587 |
| 80017 | *DGLUCY* | ENSG00000133943.21 | 6.15 | 5.38 | -0.76 | 0.59 | 2.76708E-05 | 0.00038172 |
| 8100 | *IFT88* | ENSG00000032742.18 | 4.21 | 3.44 | -0.76 | 0.59 | 0.00020695 | 0.001683527 |
| 5565 | *PRKAB2* | ENSG00000131791.8 | 5.06 | 4.29 | -0.76 | 0.59 | 0.000329397 | 0.002412421 |
| 101928601 | *MEI4* | ENSG00000269964.4 | 4.31 | 3.54 | -0.76 | 0.59 | 8.38393E-05 | 0.000863549 |
| 54734 | *RAB39A* | ENSG00000179331.3 | 2.27 | 1.51 | -0.76 | 0.59 | 0.001585385 | 0.008101817 |
| 23331 | *TTC28* | ENSG00000100154.15 | 6.03 | 5.27 | -0.77 | 0.59 | 0.000932513 | 0.005383113 |
| 4609 | *MYC* | ENSG00000136997.21 | 5.40 | 4.63 | -0.77 | 0.59 | 2.36048E-05 | 0.00034033 |
| 255928 | *SYT14* | ENSG00000143469.20 | 3.44 | 2.67 | -0.77 | 0.59 | 0.00083497 | 0.004935829 |
| 8913 | *CACNA1G* | ENSG00000006283.18 | 4.28 | 3.51 | -0.77 | 0.59 | 3.79503E-06 | 8.89066E-05 |
| 4163 | *MCC* | ENSG00000171444.18 | 3.26 | 2.50 | -0.77 | 0.59 | 0.001303851 | 0.007014157 |
| 6660 | *SOX5* | ENSG00000134532.19 | 3.72 | 2.95 | -0.77 | 0.59 | 0.000714975 | 0.004383745 |
| 389023 | *DPP10-AS1* | ENSG00000235026.7 | 4.05 | 3.28 | -0.77 | 0.59 | 2.62856E-05 | 0.000366655 |
| 6932 | *TCF7* | ENSG00000081059.20 | 5.16 | 4.39 | -0.77 | 0.59 | 1.6205E-06 | 4.75972E-05 |
| 146857 | *SLFN13* | ENSG00000154760.14 | 4.24 | 3.47 | -0.77 | 0.58 | 0.000623761 | 0.003968424 |
| 11092 | *SPACA9* | ENSG00000165698.16 | 3.68 | 2.90 | -0.77 | 0.58 | 6.48667E-05 | 0.000718369 |
| 84455 | *EFCAB7* | ENSG00000203965.13 | 3.17 | 2.40 | -0.77 | 0.58 | 0.001387378 | 0.007326939 |
| 284358 | *MAMSTR* | ENSG00000176909.12 | 4.23 | 3.46 | -0.78 | 0.58 | 0.000961264 | 0.005502228 |
| 5422 | *POLA1* | ENSG00000101868.13 | 5.58 | 4.80 | -0.78 | 0.58 | 2.61422E-06 | 6.84626E-05 |
| 284339 | *TMEM145* | ENSG00000167619.13 | 4.77 | 3.99 | -0.78 | 0.58 | 0.000330354 | 0.002413962 |
| 2938 | *GSTA1* | ENSG00000243955.6 | 9.63 | 8.85 | -0.78 | 0.58 | 1.17493E-07 | 7.97942E-06 |
| 548596 | *CKMT1A* | ENSG00000223572.10 | 4.24 | 3.46 | -0.78 | 0.58 | 0.000567232 | 0.003691927 |
| 8777 | *MPDZ* | ENSG00000107186.17 | 5.10 | 4.32 | -0.78 | 0.58 | 0.000552491 | 0.003610074 |
| 80199 | *FUZ* | ENSG00000010361.14 | 4.39 | 3.61 | -0.78 | 0.58 | 9.21976E-05 | 0.000924662 |
| 55154 | *MSTO1* | ENSG00000125459.17 | 5.34 | 4.55 | -0.78 | 0.58 | 1.89789E-05 | 0.000289356 |
| 10628 | *TXNIP* | ENSG00000265972.6 | 8.28 | 7.50 | -0.79 | 0.58 | 4.43901E-08 | 4.19026E-06 |
| 92806 | *CENPBD1P* | ENSG00000177946.7 | 3.85 | 3.06 | -0.79 | 0.58 | 6.81578E-05 | 0.000743445 |
| 253039 | *CUTALP* | ENSG00000226752.10 | 5.72 | 4.93 | -0.79 | 0.58 | 4.52057E-06 | 0.000102293 |
| 57570 | *TRMT5* | ENSG00000126814.7 | 4.40 | 3.61 | -0.79 | 0.58 | 0.000446602 | 0.003066355 |
| 3321 | *IGSF3* | ENSG00000143061.18 | 5.73 | 4.94 | -0.79 | 0.58 | 3.68395E-06 | 8.75581E-05 |
| 7161 | *TP73* | ENSG00000078900.15 | 4.29 | 3.50 | -0.79 | 0.58 | 0.000154125 | 0.001344917 |
| 110806273 | *SLC5A4-AS1* | ENSG00000242082.2 | 3.41 | 2.62 | -0.80 | 0.58 | 0.000108657 | 0.001043402 |
| 54948 | *MRPL16* | ENSG00000166902.5 | 4.75 | 3.96 | -0.80 | 0.58 | 1.02344E-05 | 0.000187786 |
| 3430 | *IFI35* | ENSG00000068079.8 | 3.73 | 2.93 | -0.80 | 0.57 | 0.001199174 | 0.006564442 |
| 4306 | *NR3C2* | ENSG00000151623.15 | 3.93 | 3.13 | -0.80 | 0.57 | 1.45186E-05 | 0.000240484 |
| 728130 | *NUTM2D* | ENSG00000214562.15 | 3.42 | 2.62 | -0.80 | 0.57 | 0.000141768 | 0.001266547 |
| 55359 | *STYK1* | ENSG00000060140.9 | 3.53 | 2.72 | -0.80 | 0.57 | 6.88667E-05 | 0.000749109 |
| 83401 | *ELOVL3* | ENSG00000119915.5 | 3.89 | 3.09 | -0.80 | 0.57 | 2.26951E-06 | 6.15968E-05 |
| 64425 | *POLR1E* | ENSG00000137054.16 | 6.41 | 5.61 | -0.81 | 0.57 | 9.3689E-09 | 1.31419E-06 |
| 84277 | *DNAJC30* | ENSG00000176410.8 | 4.23 | 3.43 | -0.81 | 0.57 | 3.27268E-05 | 0.000430119 |
| 3909 | *LAMA3* | ENSG00000053747.17 | 3.13 | 2.33 | -0.81 | 0.57 | 0.000255923 | 0.001987145 |
| 4651 | *MYO10* | ENSG00000145555.15 | 8.32 | 7.51 | -0.81 | 0.57 | 2.22044E-07 | 1.25523E-05 |
| 10069 | *RWDD2B* | ENSG00000156253.7 | 3.97 | 3.16 | -0.81 | 0.57 | 0.000110611 | 0.001056754 |
| 284695 | *ZNF326* | ENSG00000162664.17 | 6.07 | 5.26 | -0.81 | 0.57 | 6.84854E-07 | 2.66295E-05 |
| 29945 | *ANAPC4* | ENSG00000053900.11 | 4.69 | 3.88 | -0.81 | 0.57 | 6.98712E-06 | 0.000139454 |
| 400954 | *EML6* | ENSG00000214595.13 | 4.64 | 3.83 | -0.81 | 0.57 | 0.000210646 | 0.001707123 |
| 10533 | *ATG7* | ENSG00000197548.13 | 4.54 | 3.72 | -0.81 | 0.57 | 0.000240767 | 0.00189694 |
| 9829 | *DNAJC6* | ENSG00000116675.16 | 5.68 | 4.87 | -0.81 | 0.57 | 2.3957E-05 | 0.000344087 |
| 22994 | *CEP131* | ENSG00000141577.14 | 4.65 | 3.84 | -0.81 | 0.57 | 3.46881E-05 | 0.000449988 |
| 26267 | *FBXO10* | ENSG00000147912.13 | 7.05 | 6.24 | -0.81 | 0.57 | 2.28864E-05 | 0.000332206 |
| 4886 | *NPY1R* | ENSG00000164128.7 | 4.04 | 3.22 | -0.81 | 0.57 | 4.35225E-05 | 0.000530649 |
| 30832 | *ZNF354C* | ENSG00000177932.7 | 2.59 | 1.77 | -0.81 | 0.57 | 0.000916518 | 0.00530855 |
| 1159 | *CKMT1B* | ENSG00000237289.10 | 5.45 | 4.64 | -0.82 | 0.57 | 1.00968E-05 | 0.000185714 |
| 730101 | *LOC730101* | ENSG00000216775.4 | 5.53 | 4.71 | -0.82 | 0.57 | 8.30528E-07 | 3.01825E-05 |
| 57476 | *GRAMD1B* | ENSG00000023171.20 | 8.92 | 8.10 | -0.82 | 0.57 | 1.90103E-07 | 1.13174E-05 |
| 2648 | *KAT2A* | ENSG00000108773.11 | 4.78 | 3.96 | -0.82 | 0.57 | 1.7771E-05 | 0.000274408 |
| 80070 | *ADAMTS20* | ENSG00000173157.18 | 1.84 | 1.02 | -0.82 | 0.57 | 0.000951848 | 0.005462685 |
| 134728 | *IRAK1BP1* | ENSG00000146243.14 | 3.15 | 2.34 | -0.82 | 0.57 | 0.001536157 | 0.007933991 |
| 2690 | *GHR* | ENSG00000112964.14 | 3.18 | 2.36 | -0.82 | 0.57 | 0.000515426 | 0.003423023 |
| 51236 | *HGH1* | ENSG00000235173.7 | 3.11 | 2.30 | -0.82 | 0.57 | 0.000101307 | 0.000995757 |
| 1153 | *CIRBP* | ENSG00000099622.14 | 7.47 | 6.65 | -0.82 | 0.57 | 3.05652E-06 | 7.65865E-05 |
| 80298 | *MTERF2* | ENSG00000120832.10 | 4.22 | 3.40 | -0.82 | 0.57 | 9.0315E-05 | 0.000911592 |
| 4968 | *OGG1* | ENSG00000114026.22 | 4.06 | 3.24 | -0.82 | 0.57 | 3.74488E-06 | 8.82039E-05 |
| 81704 | *DOCK8* | ENSG00000107099.18 | 7.71 | 6.88 | -0.82 | 0.56 | 1.25494E-08 | 1.62404E-06 |
| 79774 | *GRTP1* | ENSG00000139835.14 | 3.58 | 2.76 | -0.82 | 0.56 | 0.000792401 | 0.004754518 |
| 23154 | *NCDN* | ENSG00000020129.16 | 5.08 | 4.26 | -0.83 | 0.56 | 6.38881E-06 | 0.000130818 |
| 54970 | *TTC12* | ENSG00000149292.17 | 3.83 | 3.00 | -0.83 | 0.56 | 0.00120782 | 0.006596184 |
| 399726 | *MIR1915HG* | ENSG00000204682.8 | 2.94 | 2.11 | -0.83 | 0.56 | 0.000310084 | 0.002314296 |
| 150290 | *DUSP18* | ENSG00000167065.14 | 2.59 | 1.76 | -0.83 | 0.56 | 0.001399962 | 0.007390796 |
| 92104 | *TTC30A* | ENSG00000197557.7 | 5.38 | 4.55 | -0.83 | 0.56 | 3.89066E-07 | 1.82456E-05 |
| 55170 | *PRMT6* | ENSG00000198890.9 | 5.28 | 4.45 | -0.83 | 0.56 | 1.05807E-06 | 3.61743E-05 |
| 440104 | *TMEM198B* | ENSG00000182796.15 | 4.55 | 3.72 | -0.83 | 0.56 | 0.000194034 | 0.001602783 |
| 10481 | *HOXB13* | ENSG00000159184.8 | 4.13 | 3.29 | -0.84 | 0.56 | 1.46633E-05 | 0.000241521 |
| 862 | *RUNX1T1* | ENSG00000079102.16 | 3.76 | 2.92 | -0.84 | 0.56 | 0.001624046 | 0.008262816 |
| 7098 | *TLR3* | ENSG00000164342.13 | 3.52 | 2.68 | -0.84 | 0.56 | 0.000883589 | 0.005152211 |
| 23211 | *ZC3H4* | ENSG00000130749.10 | 5.94 | 5.11 | -0.84 | 0.56 | 8.2266E-07 | 2.99692E-05 |
| 79414 | *LRFN3* | ENSG00000126243.9 | 5.27 | 4.43 | -0.84 | 0.56 | 0.000504468 | 0.003365383 |
| 374868 | *ATP9B* | ENSG00000166377.21 | 4.61 | 3.78 | -0.84 | 0.56 | 4.92525E-05 | 0.00058299 |
| 54957 | *TXNL4B* | ENSG00000140830.9 | 3.91 | 3.07 | -0.84 | 0.56 | 0.000179298 | 0.001502234 |
| 255082 | *CASC2* | ENSG00000177640.16 | 2.78 | 1.93 | -0.84 | 0.56 | 0.000458557 | 0.003127524 |

| | | | | | | | | |
|---|---|---|---|---|---|---|---|---|
| 2121 | *EVC* | ENSG00000072840.13 | 5.13 | 4.28 | -0.84 | 0.56 | 2.23738E-05 | 0.000327938 |
| 158572 | *USP27X-DT* | ENSG00000234390.4 | 3.18 | 2.34 | -0.84 | 0.56 | 0.000114711 | 0.001082827 |
| 26022 | *TMEM98* | ENSG00000006042.12 | 4.60 | 3.76 | -0.85 | 0.56 | 7.27788E-05 | 0.000780241 |
| 256364 | *EML3* | ENSG00000149499.12 | 4.33 | 3.49 | -0.85 | 0.56 | 0.001636703 | 0.008307498 |
| 463 | *ZFHX3* | ENSG00000140836.17 | 4.77 | 3.92 | -0.85 | 0.55 | 0.000311388 | 0.002320567 |
| 11178 | *LZTS1* | ENSG00000061337.15 | 3.32 | 2.47 | -0.85 | 0.55 | 9.22259E-05 | 0.000924662 |
| 90203 | *SNX21* | ENSG00000124104.19 | 4.80 | 3.94 | -0.85 | 0.55 | 1.38471E-06 | 4.3501E-05 |
| 119392 | *SFR1* | ENSG00000156384.15 | 2.57 | 1.72 | -0.86 | 0.55 | 0.000159155 | 0.001379187 |
| 92170 | *MTG1* | ENSG00000148824.19 | 4.40 | 3.54 | -0.86 | 0.55 | 0.001568481 | 0.008048318 |
| 93611 | *FBXO44* | ENSG00000132879.14 | 4.11 | 3.25 | -0.86 | 0.55 | 3.67374E-05 | 0.000473297 |
| 7587 | *ZNF37A* | ENSG00000075407.19 | 5.12 | 4.26 | -0.86 | 0.55 | 8.87228E-07 | 3.17057E-05 |
| 140738 | *TMEM37* | ENSG00000171227.7 | 2.53 | 1.67 | -0.86 | 0.55 | 0.000425489 | 0.00296067 |
| 79818 | *ZNF552* | ENSG00000178935.5 | 2.53 | 1.67 | -0.86 | 0.55 | 0.000203736 | 0.001665509 |
| 23428 | *SLC7A8* | ENSG00000092068.21 | 7.39 | 6.53 | -0.86 | 0.55 | 3.26383E-08 | 3.30569E-06 |
| 79979 | *TRMT2B* | ENSG00000188917.15 | 4.60 | 3.73 | -0.86 | 0.55 | 4.09972E-05 | 0.000509916 |
| 57118 | *CAMK1D* | ENSG00000183049.14 | 3.39 | 2.53 | -0.86 | 0.55 | 0.000203258 | 0.001663417 |
| 55607 | *PPP1R9A* | ENSG00000158528.12 | 4.88 | 4.02 | -0.86 | 0.55 | 0.000431642 | 0.002991001 |
| 55157 | *DARS2* | ENSG00000117593.12 | 6.71 | 5.84 | -0.87 | 0.55 | 1.39747E-08 | 1.7597E-06 |
| 9856 | *KIAA0319* | ENSG00000137261.15 | 1.62 | 0.75 | -0.87 | 0.55 | 0.000374237 | 0.002673452 |
| 10799 | *RPP40* | ENSG00000124787.14 | 2.75 | 1.88 | -0.87 | 0.55 | 0.000185424 | 0.001543558 |
| 201255 | *LRRC45* | ENSG00000169683.8 | 3.55 | 2.68 | -0.87 | 0.55 | 2.04812E-05 | 0.000307402 |
| 10940 | *POP1* | ENSG00000104356.11 | 5.08 | 4.21 | -0.87 | 0.55 | 5.02834E-06 | 0.000109761 |
| 92840 | *REEP6* | ENSG00000115255.12 | 5.34 | 4.47 | -0.87 | 0.55 | 1.59433E-07 | 9.97055E-06 |
| 50944 | *SHANK1* | ENSG00000161681.17 | 7.07 | 6.20 | -0.88 | 0.54 | 1.19943E-06 | 3.93063E-05 |
| 10810 | *WASF3* | ENSG00000132970.14 | 6.41 | 5.54 | -0.88 | 0.54 | 7.62431E-05 | 0.000807005 |
| 2898 | *GRIK2* | ENSG00000164418.22 | 5.61 | 4.73 | -0.88 | 0.54 | 4.121E-06 | 9.53036E-05 |
| 10806 | *SDCCAG8* | ENSG00000054282.16 | 4.57 | 3.69 | -0.88 | 0.54 | 0.000887705 | 0.005174197 |
| 374393 | *FAM111B* | ENSG00000189057.11 | 6.04 | 5.16 | -0.88 | 0.54 | 1.33868E-06 | 4.28027E-05 |
| 9704 | *DHX34* | ENSG00000134815.19 | 4.03 | 3.15 | -0.88 | 0.54 | 2.08776E-05 | 0.000311483 |
| 259230 | *SGMS1* | ENSG00000198964.14 | 5.61 | 4.73 | -0.88 | 0.54 | 0.000702264 | 0.004325107 |
| 84152 | *PPP1R1B* | ENSG00000131771.14 | 3.64 | 2.75 | -0.88 | 0.54 | 2.59586E-05 | 0.000364507 |
| 56952 | *PRTFDC1* | ENSG00000099256.19 | 4.28 | 3.40 | -0.88 | 0.54 | 2.60763E-05 | 0.000365092 |
| 51276 | *ZNF571* | ENSG00000180479.14 | 3.47 | 2.58 | -0.89 | 0.54 | 0.000226031 | 0.001807409 |
| 90673 | *PPP1R3E* | ENSG00000235194.9 | 4.34 | 3.45 | -0.89 | 0.54 | 0.000356814 | 0.00257472 |
| 145773 | *FAM81A* | ENSG00000157470.12 | 3.80 | 2.92 | -0.89 | 0.54 | 0.00025703 | 0.001991614 |
| 729013 | *ZBED5-AS1* | ENSG00000247271.8 | 1.79 | 0.91 | -0.89 | 0.54 | 0.000448725 | 0.003079521 |
| 114801 | *TMEM200A* | ENSG00000164484.12 | 6.06 | 5.18 | -0.89 | 0.54 | 1.31563E-07 | 8.62284E-06 |
| 80310 | *PDGFD* | ENSG00000170962.13 | 7.13 | 6.24 | -0.89 | 0.54 | 3.28168E-08 | 3.30569E-06 |
| 100302692 | *FTX* | ENSG00000230590.12 | 3.66 | 2.77 | -0.89 | 0.54 | 0.000652171 | 0.004088733 |
| 5557 | *PRIM1* | ENSG00000198056.15 | 4.80 | 3.90 | -0.89 | 0.54 | 8.38866E-05 | 0.000863549 |
| 170960 | *ZNF721* | ENSG00000182903.16 | 3.07 | 2.18 | -0.90 | 0.54 | 0.000636991 | 0.004022125 |
| 112268269 | *LOC112268269* | ENSG00000268858.2 | 2.90 | 2.00 | -0.90 | 0.54 | 5.42856E-05 | 0.000626748 |
| 2048 | *EPHB2* | ENSG00000133216.17 | 5.75 | 4.85 | -0.90 | 0.54 | 1.10829E-06 | 3.75381E-05 |
| 54841 | *BIVM* | ENSG00000134897.14 | 4.86 | 3.96 | -0.90 | 0.54 | 6.91245E-06 | 0.000138888 |
| 116987 | *AGAP1* | ENSG00000157985.19 | 4.68 | 3.77 | -0.90 | 0.54 | 0.000269947 | 0.00207351 |
| 9677 | *PPIP5K1* | ENSG00000168781.23 | 5.77 | 4.87 | -0.90 | 0.54 | 0.000115882 | 0.00109114 |
| 79695 | *GALNT12* | ENSG00000119514.7 | 3.02 | 2.12 | -0.90 | 0.53 | 0.000434496 | 0.003005534 |
| 6431 | *SRSF6* | ENSG00000124193.16 | 6.84 | 5.93 | -0.90 | 0.53 | 6.04293E-07 | 2.47926E-05 |
| 8908 | *GYG2* | ENSG00000056998.20 | 3.87 | 2.97 | -0.90 | 0.53 | 1.37255E-06 | 4.33696E-05 |
| 9227 | *LRAT* | ENSG00000121207.12 | 4.65 | 3.74 | -0.90 | 0.53 | 1.75641E-05 | 0.000272334 |
| 81033 | *KCNH6* | ENSG00000173826.15 | 2.43 | 1.53 | -0.90 | 0.53 | 0.000430152 | 0.002983431 |
| 390927 | *ZNF793* | ENSG00000188227.14 | 2.54 | 1.63 | -0.91 | 0.53 | 0.001362431 | 0.007230807 |
| 9187 | *SLC24A1* | ENSG00000074621.14 | 3.57 | 2.65 | -0.91 | 0.53 | 0.000916767 | 0.00530855 |
| 57102 | *C12orf4* | ENSG00000047621.12 | 4.36 | 3.45 | -0.91 | 0.53 | 0.000113891 | 0.001078483 |
| 4773 | *NFATC2* | ENSG00000101096.20 | 3.11 | 2.20 | -0.91 | 0.53 | 0.000118815 | 0.001107861 |
| 54982 | *CLN6* | ENSG00000128973.13 | 6.14 | 5.23 | -0.91 | 0.53 | 5.219E-06 | 0.000112708 |
| 115207 | *KCTD12* | ENSG00000178695.6 | 3.08 | 2.17 | -0.91 | 0.53 | 2.64757E-05 | 0.000368964 |
| 80217 | *CFAP43* | ENSG00000197748.13 | 1.99 | 1.07 | -0.91 | 0.53 | 0.002035458 | 0.009842573 |
| 197335 | *WDR90* | ENSG00000161996.19 | 4.31 | 3.40 | -0.91 | 0.53 | 0.000750492 | 0.004556688 |
| 594 | *BCKDHB* | ENSG00000083123.15 | 4.28 | 3.37 | -0.91 | 0.53 | 6.25059E-05 | 0.000694927 |
| 10402 | *ST3GAL6* | ENSG00000064225.13 | 2.95 | 2.04 | -0.91 | 0.53 | 0.000602836 | 0.00386996 |
| 136227 | *COL26A1* | ENSG00000160963.14 | 4.74 | 3.83 | -0.91 | 0.53 | 0.000324062 | 0.002385409 |
| 147949 | *ZNF583* | ENSG00000198440.9 | 3.12 | 2.21 | -0.92 | 0.53 | 0.000316538 | 0.002349617 |
| 745 | *MYRF* | ENSG00000124920.14 | 3.33 | 2.41 | -0.92 | 0.53 | 0.00027835 | 0.002123924 |
| 150737 | *TTC30B* | ENSG00000196659.10 | 4.85 | 3.93 | -0.92 | 0.53 | 2.44539E-07 | 1.35435E-05 |
| 118738 | *ZNF488* | ENSG00000265763.4 | 4.12 | 3.20 | -0.92 | 0.53 | 1.47003E-05 | 0.000241661 |
| 59283 | *CACNG8* | ENSG00000142408.7 | 2.87 | 1.95 | -0.92 | 0.53 | 0.001716473 | 0.008596111 |
| 125113 | *KRT222* | ENSG00000213424.9 | 4.04 | 3.12 | -0.93 | 0.53 | 1.831E-05 | 0.000280424 |
| 440173 | *LINC02893* | ENSG00000269994.3 | 4.26 | 3.34 | -0.93 | 0.53 | 5.1848E-06 | 0.000112455 |
| 135932 | *TMEM139* | ENSG00000178826.11 | 1.07 | 0.13 | -0.93 | 0.52 | 0.000624193 | 0.003968424 |
| 89795 | *NAV3* | ENSG00000067798.16 | 4.78 | 3.85 | -0.93 | 0.52 | 0.002031043 | 0.009830353 |
| 80206 | *FHOD3* | ENSG00000134775.16 | 4.74 | 3.80 | -0.93 | 0.52 | 2.85599E-05 | 0.000389369 |
| 3992 | *FADS1* | ENSG00000149485.19 | 5.95 | 5.01 | -0.94 | 0.52 | 2.89561E-07 | 1.5357E-05 |
| 201232 | *SLC16A13* | ENSG00000174327.7 | 3.08 | 2.14 | -0.94 | 0.52 | 0.000436461 | 0.003011882 |
| 27034 | *ACAD8* | ENSG00000151498.12 | 4.73 | 3.80 | -0.94 | 0.52 | 3.35352E-06 | 8.19754E-05 |
| 144571 | *A2M-AS1* | ENSG00000245105.4 | 2.89 | 1.95 | -0.94 | 0.52 | 3.07825E-05 | 0.000411044 |
| 282679 | *AQP11* | ENSG00000178301.4 | 3.28 | 2.33 | -0.94 | 0.52 | 1.51344E-05 | 0.000247173 |
| 100129461 | *LYRM4-AS1* | ENSG00000272142.4 | 3.37 | 2.42 | -0.94 | 0.52 | 9.7353E-06 | 0.000180713 |
| 5639 | *PRRG2* | ENSG00000126460.11 | 2.95 | 1.99 | -0.94 | 0.52 | 0.001488761 | 0.007743795 |
| 92659 | *MAFG-DT* | ENSG00000265688.2 | 1.52 | 0.57 | -0.95 | 0.52 | 0.001325995 | 0.007092607 |
| 59269 | *HIVEP3* | ENSG00000127124.16 | 3.59 | 2.65 | -0.95 | 0.52 | 1.44518E-05 | 0.000240046 |
| 100288842 | *B3GALT9* | ENSG00000214654.9 | 2.72 | 1.77 | -0.95 | 0.52 | 9.04685E-05 | 0.000912528 |
| 389840 | *MAP3K15* | ENSG00000180815.15 | 3.61 | 2.66 | -0.95 | 0.52 | 2.12977E-06 | 5.86527E-05 |
| 219931 | *TPCN2* | ENSG00000162341.18 | 2.68 | 1.72 | -0.95 | 0.52 | 0.00065537 | 0.004098518 |
| 56154 | *TEX15* | ENSG00000133863.9 | 4.20 | 3.25 | -0.95 | 0.52 | 5.84543E-06 | 0.000122363 |
| 60529 | *ALX4* | ENSG00000052850.8 | 3.25 | 2.29 | -0.95 | 0.52 | 5.89295E-06 | 0.000122673 |
| 51196 | *PLCE1* | ENSG00000138193.17 | 3.73 | 2.78 | -0.95 | 0.52 | 0.000302273 | 0.002272954 |

| | | | | | | | | |
|---|---|---|---|---|---|---|---|---|
| 64429 | *ZDHHC6* | ENSG00000023041.12 | 5.54 | 4.58 | -0.96 | 0.52 | 2.65698E-07 | 1.43966E-05 |
| 22885 | *ABLIM3* | ENSG00000173210.20 | 3.51 | 2.55 | -0.96 | 0.51 | 4.79311E-06 | 0.000106892 |
| 149233 | *IL23R* | ENSG00000162594.16 | 3.19 | 2.23 | -0.96 | 0.51 | 0.000159942 | 0.001382027 |
| 285382 | *C3orf70* | ENSG00000187068.3 | 3.27 | 2.31 | -0.96 | 0.51 | 3.3265E-05 | 0.000436048 |
| 114787 | *GPRIN1* | ENSG00000169258.7 | 5.10 | 4.13 | -0.96 | 0.51 | 9.48462E-07 | 3.33383E-05 |
| 93517 | *SDR42E1* | ENSG00000184860.10 | 3.49 | 2.52 | -0.96 | 0.51 | 0.001331357 | 0.007111368 |
| 114987 | *WDR31* | ENSG00000148225.16 | 3.75 | 2.79 | -0.97 | 0.51 | 1.08815E-05 | 0.000196535 |
| 10046 | *MAMLD1* | ENSG00000013619.15 | 3.82 | 2.85 | -0.97 | 0.51 | 6.88902E-05 | 0.000749109 |
| 6508 | *SLC4A3* | ENSG00000114923.17 | 2.89 | 1.92 | -0.97 | 0.51 | 0.000126804 | 0.001162611 |
| 36 | *ACADSB* | ENSG00000196177.13 | 5.77 | 4.79 | -0.98 | 0.51 | 8.68767E-08 | 6.45511E-06 |
| 128710 | *SLX4IP* | ENSG00000149346.15 | 3.74 | 2.77 | -0.98 | 0.51 | 3.83932E-05 | 0.000487103 |
| 84852 | *ATP1A1-AS1* | ENSG00000203865.12 | 2.80 | 1.82 | -0.98 | 0.51 | 0.001860732 | 0.009171668 |
| 64881 | *PCDH20* | ENSG00000280165.1 | 4.04 | 3.06 | -0.98 | 0.51 | 1.185E-06 | 3.91756E-05 |
| 83478 | *ARHGAP24* | ENSG00000138639.18 | 4.15 | 3.17 | -0.98 | 0.51 | 0.000176827 | 0.001492684 |
| 6469 | *SHH* | ENSG00000164690.8 | 2.78 | 1.79 | -0.98 | 0.51 | 0.001570677 | 0.00805408 |
| 170712 | *COX7B2* | ENSG00000170516.17 | 1.49 | 0.51 | -0.98 | 0.51 | 0.000449649 | 0.003083041 |
| 653689 | *GSTT2B* | ENSG00000133433.11 | 6.81 | 5.82 | -0.99 | 0.50 | 4.01856E-08 | 3.9069E-06 |
| 100131369 | *DUX4L27* | ENSG00000258794.3 | 1.83 | 0.85 | -0.99 | 0.50 | 0.001152411 | 0.006368388 |
| 55930 | *MYO5C* | ENSG00000128833.13 | 4.82 | 3.84 | -0.99 | 0.50 | 9.58461E-05 | 0.000951765 |
| 9619 | *ABCG1* | ENSG00000160179.19 | 2.68 | 1.68 | -0.99 | 0.50 | 0.000243898 | 0.00191323 |
| 100289635 | *ZNF605* | ENSG00000196458.11 | 4.08 | 3.09 | -0.99 | 0.50 | 0.001726874 | 0.00863955 |
| 60401 | *EDA2R* | ENSG00000131080.15 | 3.69 | 2.70 | -0.99 | 0.50 | 8.28419E-06 | 0.000159203 |
| 2395 | *FXN* | ENSG00000165060.15 | 4.53 | 3.54 | -0.99 | 0.50 | 1.06141E-05 | 0.000192866 |
| 10999 | *SLC27A4* | ENSG00000167114.13 | 6.83 | 5.83 | -0.99 | 0.50 | 2.88768E-09 | 6.01961E-07 |
| 1909 | *EDNRA* | ENSG00000151617.17 | 2.69 | 1.70 | -1.00 | 0.50 | 0.002065208 | 0.009948719 |
| 6427 | *SRSF2* | ENSG00000161547.17 | 7.82 | 6.82 | -1.00 | 0.50 | 1.726E-09 | 4.8302E-07 |
| 83879 | *CDCA7* | ENSG00000144354.14 | 4.65 | 3.65 | -1.00 | 0.50 | 3.23151E-07 | 1.61104E-05 |
| 10813 | *UTP14A* | ENSG00000156697.13 | 4.70 | 3.69 | -1.00 | 0.50 | 3.78243E-07 | 1.79087E-05 |
| 359809 | *PEG13* | ENSG00000282164.3 | 2.72 | 1.71 | -1.00 | 0.50 | 0.001512 | 0.007828082 |
| 4693 | *NDP* | ENSG00000124479.11 | 2.32 | 1.32 | -1.00 | 0.50 | 6.80225E-05 | 0.00074305 |
| 105370333 | *PCCA-DT* | ENSG00000274605.3 | 3.46 | 2.45 | -1.01 | 0.50 | 3.17146E-05 | 0.000420871 |
| 729262 | *NUTM2B* | ENSG00000188199.11 | 1.81 | 0.81 | -1.01 | 0.50 | 0.000159913 | 0.001382027 |
| 55672 | *NBPF1* | ENSG00000219481.11 | 7.95 | 6.94 | -1.01 | 0.50 | 5.00663E-06 | 0.00010954 |
| 286016 | *TPI1P2* | ENSG00000230359.5 | 1.69 | 0.68 | -1.01 | 0.50 | 0.000494086 | 0.003310557 |
| 8527 | *DGKD* | ENSG00000077044.11 | 5.07 | 4.06 | -1.01 | 0.50 | 0.000522304 | 0.003451899 |
| 54487 | *DGCR8* | ENSG00000128191.16 | 6.06 | 5.05 | -1.01 | 0.50 | 0.000104635 | 0.00101551 |
| 29075 | *LINC00652* | ENSG00000179935.10 | 1.54 | 0.53 | -1.01 | 0.50 | 0.000974591 | 0.005573856 |
| 100129518 | *SOD2-OT1* | ENSG00000285427.1 | 1.88 | 0.87 | -1.01 | 0.50 | 0.000694608 | 0.004283223 |
| 147495 | *APCDD1* | ENSG00000154856.13 | 7.78 | 6.77 | -1.01 | 0.50 | 3.66085E-07 | 1.74986E-05 |
| 54843 | *SYTL2* | ENSG00000137501.18 | 8.94 | 7.92 | -1.02 | 0.49 | 8.5498E-10 | 2.78965E-07 |
| 2264 | *FGFR4* | ENSG00000160867.15 | 5.33 | 4.31 | -1.02 | 0.49 | 1.73008E-06 | 5.03232E-05 |
| 285368 | *PRRT3* | ENSG00000163704.12 | 4.86 | 3.85 | -1.02 | 0.49 | 4.12366E-05 | 0.000511504 |
| 4684 | *NCAM1* | ENSG00000149294.17 | 7.03 | 6.01 | -1.02 | 0.49 | 1.91215E-07 | 1.13174E-05 |
| 9478 | *CABP1* | ENSG00000157782.10 | 2.96 | 1.94 | -1.02 | 0.49 | 0.001125483 | 0.006265717 |
| 100101467 | *ZSCAN30* | ENSG00000186814.14 | 4.33 | 3.30 | -1.02 | 0.49 | 1.4806E-05 | 0.000242602 |
| 4919 | *ROR1* | ENSG00000185483.13 | 3.86 | 2.83 | -1.02 | 0.49 | 2.12469E-05 | 0.000315114 |
| 285908 | *LINC00174* | ENSG00000179406.9 | 3.32 | 2.30 | -1.03 | 0.49 | 1.70829E-05 | 0.000267918 |
| 197342 | *EME2* | ENSG00000197774.14 | 2.31 | 1.28 | -1.03 | 0.49 | 0.000627288 | 0.003984328 |
| 2304 | *FOXE1* | ENSG00000178919.9 | 2.89 | 1.86 | -1.03 | 0.49 | 8.30366E-05 | 0.000859051 |
| 9622 | *KLK4* | ENSG00000167749.12 | 2.67 | 1.64 | -1.03 | 0.49 | 0.000935459 | 0.005395966 |
| 50617 | *ATP6V0A4* | ENSG00000105929.16 | 1.51 | 0.48 | -1.03 | 0.49 | 8.13139E-05 | 0.000846402 |
| 8187 | *ZNF239* | ENSG00000196793.14 | 1.81 | 0.78 | -1.03 | 0.49 | 0.000458501 | 0.003127524 |
| 1122 | *CHML* | ENSG00000203668.3 | 5.78 | 4.74 | -1.03 | 0.49 | 1.11046E-06 | 3.75381E-05 |
| 124152 | *IQCK* | ENSG00000174628.16 | 2.79 | 1.75 | -1.03 | 0.49 | 5.2023E-05 | 0.000605751 |
| 26053 | *AUTS2* | ENSG00000158321.18 | 4.39 | 3.36 | -1.04 | 0.49 | 0.000639749 | 0.004033175 |
| 8740 | *TNFSF14* | ENSG00000125735.11 | 1.81 | 0.77 | -1.04 | 0.49 | 0.000285167 | 0.002166027 |
| 7004 | *TEAD4* | ENSG00000197905.10 | 2.45 | 1.41 | -1.04 | 0.49 | 0.000485063 | 0.00326325 |
| 1589 | *CYP21A2* | ENSG00000231852.9 | 8.32 | 7.28 | -1.04 | 0.49 | 1.28081E-08 | 1.64306E-06 |
| 1010 | *CDH12* | ENSG00000154162.15 | 5.03 | 3.99 | -1.04 | 0.49 | 8.01287E-06 | 0.000155181 |
| 7771 | *ZNF112* | ENSG00000062370.17 | 3.53 | 2.49 | -1.05 | 0.48 | 3.21274E-05 | 0.000424524 |
| 54661 | *PCDHB17P* | ENSG00000255622.3 | 1.43 | 0.38 | -1.05 | 0.48 | 0.001076954 | 0.006031344 |
| 147660 | *ZNF578* | ENSG00000258405.11 | 3.16 | 2.11 | -1.05 | 0.48 | 0.000741903 | 0.004515503 |
| 54549 | *SDK2* | ENSG00000069188.17 | 3.94 | 2.89 | -1.05 | 0.48 | 2.33449E-05 | 0.000336908 |
| 28988 | *DBNL* | ENSG00000136279.21 | 7.80 | 6.75 | -1.05 | 0.48 | 2.21585E-08 | 2.48192E-06 |
| 100329135 | *TRPC5OS* | ENSG00000204025.8 | 5.26 | 4.21 | -1.05 | 0.48 | 6.88405E-07 | 2.66984E-05 |
| 6855 | *SYP* | ENSG00000102003.12 | 5.69 | 4.64 | -1.06 | 0.48 | 8.53127E-08 | 6.3946E-06 |
| 9814 | *SFI1* | ENSG00000198089.16 | 3.60 | 2.54 | -1.07 | 0.48 | 0.000740785 | 0.004510523 |
| 2939 | *GSTA2* | ENSG00000244067.3 | 4.10 | 3.03 | -1.07 | 0.48 | 3.51265E-06 | 8.44894E-05 |
| 56124 | *PCDHB12* | ENSG00000120328.6 | 1.72 | 0.65 | -1.07 | 0.48 | 0.000756024 | 0.004584713 |
| 101927021 | *ZFAND2A-DT* | ENSG00000229043.3 | 1.10 | 0.03 | -1.07 | 0.47 | 0.000619017 | 0.003950182 |
| 339039 | *PKD1P3* | ENSG00000183458.14 | 3.79 | 2.71 | -1.08 | 0.47 | 0.000976897 | 0.00558349 |
| 185 | *AGTR1* | ENSG00000144891.19 | 6.25 | 5.18 | -1.08 | 0.47 | 2.48528E-09 | 5.40603E-07 |
| 100507437 | *LOC100507437* | ENSG00000251602.7 | 2.46 | 1.38 | -1.08 | 0.47 | 0.000113351 | 0.001076765 |
| 9515 | *STXBP5L* | ENSG00000145087.13 | 5.51 | 4.42 | -1.08 | 0.47 | 7.44692E-06 | 0.000146489 |
| 4053 | *LTBP2* | ENSG00000119681.12 | 3.19 | 2.11 | -1.08 | 0.47 | 4.99646E-05 | 0.000588189 |
| 25939 | *SAMHD1* | ENSG00000101347.11 | 7.62 | 6.53 | -1.09 | 0.47 | 9.27108E-09 | 1.31273E-06 |
| 176 | *ACAN* | ENSG00000157766.19 | 3.03 | 1.93 | -1.10 | 0.47 | 1.39892E-05 | 0.000234596 |
| 55520 | *ELAC1* | ENSG00000141642.9 | 2.87 | 1.77 | -1.10 | 0.47 | 0.000250253 | 0.001953243 |
| 55329 | *MNS1* | ENSG00000138587.6 | 2.63 | 1.53 | -1.10 | 0.47 | 3.06885E-05 | 0.000410155 |
| 202243 | *CCDC125* | ENSG00000183323.13 | 3.66 | 2.56 | -1.10 | 0.47 | 0.000571223 | 0.003714681 |
| 50863 | *NTM* | ENSG00000182667.15 | 4.27 | 3.17 | -1.10 | 0.47 | 1.13876E-05 | 0.00020228 |
| 399947 | *C11orf87* | ENSG00000185742.7 | 1.08 | -0.02 | -1.10 | 0.47 | 0.001925492 | 0.009419353 |
| 582 | *BBS1* | ENSG00000174483.20 | 5.31 | 4.20 | -1.10 | 0.46 | 1.2716E-07 | 8.48239E-06 |
| 100134713 | *NDUFB2-AS1* | ENSG00000240889.1 | 0.80 | -0.31 | -1.11 | 0.46 | 0.001007933 | 0.005723823 |
| 387640 | *SKIDA1* | ENSG00000180592.17 | 2.42 | 1.31 | -1.11 | 0.46 | 0.000175784 | 0.001488907 |
| 10544 | *PROCR* | ENSG00000101000.6 | 2.76 | 1.65 | -1.11 | 0.46 | 0.000252962 | 0.001969315 |
| 399948 | *COLCA1* | ENSG00000196167.10 | 4.04 | 2.92 | -1.11 | 0.46 | 1.81747E-06 | 5.24132E-05 |

| | | | | | | | | |
|---|---|---|---|---|---|---|---|---|
| 5198 | *PFAS* | ENSG00000178921.14 | 5.20 | 4.09 | -1.11 | 0.46 | 3.20828E-05 | 0.000424524 |
| 245812 | *CNPY4* | ENSG00000166997.8 | 4.08 | 2.97 | -1.11 | 0.46 | 6.34886E-07 | 2.54107E-05 |
| 5935 | *RBM3* | ENSG00000102317.18 | 6.75 | 5.64 | -1.11 | 0.46 | 1.78108E-07 | 1.08667E-05 |
| 55259 | *DNAI7* | ENSG00000118307.20 | 1.75 | 0.63 | -1.12 | 0.46 | 0.001213613 | 0.006621272 |
| 50861 | *STMN3* | ENSG00000197457.10 | 5.63 | 4.51 | -1.12 | 0.46 | 9.73793E-09 | 1.3533E-06 |
| 57821 | *CCDC181* | ENSG00000117477.13 | 2.38 | 1.27 | -1.12 | 0.46 | 0.000268743 | 0.002068491 |
| 151647 | *TAFA4* | ENSG00000163377.16 | 7.75 | 6.63 | -1.12 | 0.46 | 6.28802E-09 | 9.63029E-07 |
| 57545 | *CC2D2A* | ENSG00000048342.18 | 4.38 | 3.26 | -1.12 | 0.46 | 1.32858E-05 | 0.000226856 |
| 105369364 | *LINC02701* | ENSG00000250508.1 | 4.43 | 3.30 | -1.12 | 0.46 | 3.76075E-07 | 1.78624E-05 |
| 1577 | *CYP3A5* | ENSG00000106258.15 | 3.86 | 2.74 | -1.13 | 0.46 | 8.52327E-05 | 0.000873214 |
| 4674 | *NAP1L2* | ENSG00000186462.9 | 4.47 | 3.34 | -1.13 | 0.46 | 5.87511E-06 | 0.000122472 |
| 5939 | *RBMS2* | ENSG00000076067.13 | 5.23 | 4.09 | -1.13 | 0.46 | 0.0005315 | 0.003501882 |
| 7691 | *ZNF132* | ENSG00000131849.13 | 2.29 | 1.15 | -1.13 | 0.46 | 0.001489013 | 0.007743795 |
| 441046 | *GUSBP5* | ENSG00000236296.8 | 2.02 | 0.88 | -1.14 | 0.46 | 0.000106282 | 0.001028491 |
| 157983 | *DOCK8-AS1* | ENSG00000183784.7 | 0.12 | -1.01 | -1.14 | 0.45 | 0.001309951 | 0.007036885 |
| 9079 | *LDB2* | ENSG00000169744.13 | 5.37 | 4.24 | -1.14 | 0.45 | 2.04148E-06 | 5.69526E-05 |
| 161582 | *DNAAF4* | ENSG00000256061.7 | 2.79 | 1.65 | -1.14 | 0.45 | 0.001852383 | 0.009145533 |
| 220963 | *SLC16A9* | ENSG00000165449.12 | 2.19 | 1.05 | -1.14 | 0.45 | 3.79995E-05 | 0.000483744 |
| 84457 | *PHYHIPL* | ENSG00000165443.12 | 4.87 | 3.73 | -1.14 | 0.45 | 3.07053E-07 | 1.57289E-05 |
| 19 | *ABCA1* | ENSG00000165029.17 | 4.94 | 3.79 | -1.15 | 0.45 | 3.39705E-06 | 8.277E-05 |
| 100506930 | *LINC00665* | ENSG00000232677.10 | 5.24 | 4.10 | -1.15 | 0.45 | 4.08555E-08 | 3.9069E-06 |
| 120071 | *LARGE2* | ENSG00000165905.18 | 3.63 | 2.48 | -1.15 | 0.45 | 0.000761355 | 0.004613315 |
| 4900 | *NRGN* | ENSG00000154146.13 | 3.19 | 2.04 | -1.15 | 0.45 | 5.20834E-06 | 0.000112708 |
| 11182 | *SLC2A6* | ENSG00000160326.14 | 2.45 | 1.29 | -1.15 | 0.45 | 0.000435466 | 0.003007525 |
| 79159 | *NOL12* | ENSG00000273899.5 | 3.69 | 2.54 | -1.16 | 0.45 | 4.87381E-05 | 0.000579185 |
| 18 | *ABAT* | ENSG00000183044.12 | 4.06 | 2.90 | -1.16 | 0.45 | 2.64584E-06 | 6.89433E-05 |
| 100009676 | *ZBTB11-AS1* | ENSG00000256628.3 | 3.01 | 1.85 | -1.16 | 0.45 | 1.25783E-06 | 4.076E-05 |
| 26220 | *DGCR5* | ENSG00000273032.3 | 4.54 | 3.38 | -1.16 | 0.45 | 5.25557E-07 | 2.25374E-05 |
| 51306 | *FAM13B* | ENSG00000031003.11 | 7.09 | 5.93 | -1.16 | 0.45 | 1.39626E-07 | 8.99417E-06 |
| 51167 | *CYB5R4* | ENSG00000065615.14 | 3.89 | 2.74 | -1.16 | 0.45 | 0.000767264 | 0.004641624 |
| 124930 | *ANKRD13B* | ENSG00000198720.13 | 3.91 | 2.75 | -1.17 | 0.45 | 1.6189E-06 | 4.75972E-05 |
| 374378 | *GALNT18* | ENSG00000110328.6 | 2.29 | 1.12 | -1.17 | 0.45 | 6.23628E-06 | 0.000128926 |
| 163131 | *ZNF780B* | ENSG00000128000.17 | 3.86 | 2.69 | -1.17 | 0.45 | 0.001314736 | 0.007049971 |
| 84253 | *GARNL3* | ENSG00000136895.19 | 3.38 | 2.21 | -1.17 | 0.44 | 0.000111461 | 0.001062846 |
| 83758 | *RBP5* | ENSG00000139194.8 | 1.87 | 0.70 | -1.17 | 0.44 | 0.000418953 | 0.002921964 |
| 84532 | *ACSS1* | ENSG00000154930.15 | 7.33 | 6.16 | -1.17 | 0.44 | 1.2543E-09 | 3.84201E-07 |
| 9540 | *TP53I3* | ENSG00000115129.14 | 2.59 | 1.41 | -1.17 | 0.44 | 0.000288228 | 0.002185957 |
| 101927761 | *TH2LCRR* | ENSG00000223442.1 | 5.19 | 4.01 | -1.17 | 0.44 | 1.7652E-08 | 2.06984E-06 |
| 55511 | *SAGE1* | ENSG00000181433.11 | 2.56 | 1.38 | -1.17 | 0.44 | 2.75189E-05 | 0.000380323 |
| 51286 | *CEND1* | ENSG00000184524.6 | 0.82 | -0.35 | -1.17 | 0.44 | 0.0004405 | 0.003028617 |
| 5026 | *P2RX5* | ENSG00000083454.22 | 2.88 | 1.71 | -1.18 | 0.44 | 0.000156341 | 0.001358727 |
| 51725 | *FBXO40* | ENSG00000163833.8 | 5.02 | 3.84 | -1.18 | 0.44 | 4.42386E-08 | 4.19026E-06 |
| 220992 | *ZNF485* | ENSG00000198298.13 | 1.17 | -0.01 | -1.18 | 0.44 | 0.000835946 | 0.004939653 |
| 10900 | *RUNDC3A* | ENSG00000108309.14 | 3.02 | 1.83 | -1.18 | 0.44 | 6.83618E-06 | 0.000137962 |
| 285600 | *KIAA0825* | ENSG00000185261.15 | 1.49 | 0.30 | -1.19 | 0.44 | 0.000331614 | 0.002420492 |
| 404281 | *YY2* | ENSG00000230797.3 | 2.45 | 1.26 | -1.19 | 0.44 | 4.44755E-05 | 0.000539638 |
| 57002 | *YAE1* | ENSG00000241127.8 | 2.83 | 1.64 | -1.19 | 0.44 | 3.90374E-05 | 0.000493607 |
| 23245 | *ASTN2* | ENSG00000148219.18 | 2.75 | 1.56 | -1.19 | 0.44 | 3.00544E-05 | 0.000403476 |
| 393 | *ARHGAP4* | ENSG00000089820.16 | 1.66 | 0.47 | -1.20 | 0.44 | 0.00126907 | 0.006866429 |
| 400591 | *TMEM132E-DT* | ENSG00000197322.4 | 0.85 | -0.35 | -1.20 | 0.44 | 0.00063375 | 0.00400672 |
| 79847 | *MFSD13A* | ENSG00000138111.14 | 3.23 | 2.02 | -1.20 | 0.44 | 0.00177417 | 0.008840809 |
| 101927417 | *FHAD1-AS1* | ENSG00000233485.1 | 4.09 | 2.89 | -1.20 | 0.44 | 2.13671E-06 | 5.87361E-05 |
| 84622 | *ZNF594* | ENSG00000180626.10 | 3.09 | 1.88 | -1.20 | 0.43 | 1.67292E-05 | 0.000265216 |
| 101929780 | *NBPF25P* | ENSG00000272150.5 | 1.39 | 0.18 | -1.20 | 0.43 | 0.000555566 | 0.003625431 |
| 23236 | *PLCB1* | ENSG00000182621.18 | 4.60 | 3.40 | -1.20 | 0.43 | 2.80855E-05 | 0.000384963 |
| 51385 | *ZNF589* | ENSG00000164048.14 | 3.35 | 2.13 | -1.21 | 0.43 | 0.00012752 | 0.001168464 |
| 129804 | *FBLN7* | ENSG00000144152.13 | 2.32 | 1.11 | -1.21 | 0.43 | 0.000379536 | 0.002701019 |
| 64220 | *STRA6* | ENSG00000137868.19 | 3.26 | 2.06 | -1.21 | 0.43 | 9.02527E-06 | 0.000170796 |
| 56967 | *C14orf132* | ENSG00000227051.7 | 3.13 | 1.91 | -1.21 | 0.43 | 0.001051192 | 0.005924647 |
| 100996307 | *LIPE-AS1* | ENSG00000213904.10 | 2.07 | 0.86 | -1.21 | 0.43 | 0.001674329 | 0.008441166 |
| 80320 | *SP6* | ENSG00000189120.5 | 5.36 | 4.14 | -1.22 | 0.43 | 4.77385E-09 | 8.50382E-07 |
| 728411 | *GUSBP1* | ENSG00000215190.9 | 1.75 | 0.52 | -1.22 | 0.43 | 0.001471591 | 0.00767867 |
| 6262 | *RYR2* | ENSG00000198626.18 | 4.63 | 3.41 | -1.22 | 0.43 | 6.50749E-05 | 0.000718697 |
| 283481 | *FGF14-AS2* | ENSG00000272143.1 | 1.96 | 0.73 | -1.22 | 0.43 | 0.00028989 | 0.002196343 |
| 8745 | *ADAM23* | ENSG00000114948.13 | 4.51 | 3.29 | -1.22 | 0.43 | 2.73257E-06 | 7.07124E-05 |
| 100507516 | *LOC100507516* | ENSG00000237807.4 | 2.31 | 1.08 | -1.22 | 0.43 | 2.98407E-05 | 0.000401656 |
| 25903 | *OLFML2B* | ENSG00000162745.11 | 3.67 | 2.44 | -1.23 | 0.43 | 5.41347E-07 | 2.29064E-05 |
| 8941 | *CDK5R2* | ENSG00000171450.6 | 2.26 | 1.03 | -1.23 | 0.43 | 2.67256E-05 | 0.000371068 |
| 201625 | *DNAH12* | ENSG00000174844.15 | 3.00 | 1.77 | -1.23 | 0.43 | 0.000262721 | 0.002030475 |
| 92369 | *SPSB4* | ENSG00000175093.5 | 3.09 | 1.85 | -1.24 | 0.42 | 3.18955E-06 | 7.91273E-05 |
| 158401 | *SHOC1* | ENSG00000165181.17 | 4.99 | 3.75 | -1.24 | 0.42 | 1.04914E-05 | 0.000191529 |
| 79839 | *CCDC102B* | ENSG00000150636.17 | 1.22 | -0.04 | -1.25 | 0.42 | 0.000619824 | 0.003951521 |
| 8536 | *CAMK1* | ENSG00000134072.11 | 4.54 | 3.29 | -1.25 | 0.42 | 1.36858E-06 | 4.33353E-05 |
| 51351 | *ZNF117* | ENSG00000152926.16 | 1.95 | 0.68 | -1.25 | 0.42 | 0.001998361 | 0.009700325 |
| 120379 | *PIH1D2* | ENSG00000150773.12 | 1.32 | 0.05 | -1.26 | 0.42 | 0.001667488 | 0.008418204 |
| 85376 | *RIMBP3* | ENSG00000275793.1 | 2.75 | 1.49 | -1.26 | 0.42 | 8.50415E-05 | 0.00087185 |
| 29896 | *TRA2A* | ENSG00000164548.12 | 6.29 | 5.03 | -1.26 | 0.42 | 2.09317E-08 | 2.36214E-06 |
| 1947 | *EFNB1* | ENSG00000090776.6 | 3.75 | 2.49 | -1.26 | 0.42 | 4.39568E-07 | 1.98719E-05 |
| 150 | *ADRA2A* | ENSG00000150594.7 | 3.78 | 2.51 | -1.26 | 0.42 | 2.28191E-05 | 0.00033155 |
| 134145 | *ATPSCKMT* | ENSG00000150756.14 | 3.15 | 1.88 | -1.27 | 0.42 | 2.2684E-05 | 0.000330728 |
| 596 | *BCL2* | ENSG00000171791.14 | 2.39 | 1.12 | -1.27 | 0.42 | 0.000100677 | 0.000990208 |
| 9068 | *ANGPTL1* | ENSG00000116194.13 | 4.06 | 2.79 | -1.27 | 0.41 | 5.2788E-05 | 0.000613706 |
| 57549 | *IGSF9* | ENSG00000085552.18 | 1.58 | 0.31 | -1.27 | 0.41 | 0.000394846 | 0.002792761 |
| 10332 | *CLEC4M* | ENSG00000104938.18 | 1.78 | 0.51 | -1.28 | 0.41 | 6.49265E-05 | 0.000718369 |
| 4744 | *NEFH* | ENSG00000100285.10 | 3.24 | 1.96 | -1.28 | 0.41 | 1.98193E-05 | 0.000299263 |
| 5608 | *MAP2K6* | ENSG00000108984.15 | 4.37 | 3.09 | -1.28 | 0.41 | 6.90265E-06 | 0.000138877 |
| 100288332 | *NPIPA5* | ENSG00000183793.14 | 2.63 | 1.34 | -1.29 | 0.41 | 0.000405446 | 0.002846276 |

| | | | | | | | | |
|---|---|---|---|---|---|---|---|---|
| 26052 | *DNM3* | ENSG00000197959.15 | 7.25 | 5.95 | -1.29 | 0.41 | 9.1373E-09 | 1.30611E-06 |
| 3995 | *FADS3* | ENSG00000221968.9 | 5.20 | 3.91 | -1.30 | 0.41 | 1.34486E-07 | 8.77609E-06 |
| 442213 | *PTCHD4* | ENSG00000244694.8 | 4.31 | 3.01 | -1.30 | 0.41 | 1.02873E-07 | 7.38767E-06 |
| 284443 | *ZNF493* | ENSG00000196268.12 | 2.50 | 1.20 | -1.31 | 0.40 | 0.00201385 | 0.009768645 |
| 79187 | *FSD1* | ENSG00000105255.11 | 2.82 | 1.52 | -1.31 | 0.40 | 2.04234E-05 | 0.000306841 |
| 4168 | *MCF2* | ENSG00000101977.21 | 4.84 | 3.53 | -1.31 | 0.40 | 6.07439E-07 | 2.47926E-05 |
| 9580 | *SOX13* | ENSG00000143842.15 | 5.82 | 4.50 | -1.32 | 0.40 | 2.44623E-09 | 5.39932E-07 |
| 4099 | *MAG* | ENSG00000105695.15 | 1.28 | -0.04 | -1.32 | 0.40 | 0.001911083 | 0.009369505 |
| 203238 | *CCDC171* | ENSG00000164989.17 | 1.65 | 0.32 | -1.32 | 0.40 | 0.000239732 | 0.001890773 |
| 643276 | *NOC2LP1* | ENSG00000213225.7 | 3.32 | 1.99 | -1.32 | 0.40 | 2.64888E-07 | 1.43966E-05 |
| 129049 | *SGSM1* | ENSG00000167037.19 | 2.62 | 1.30 | -1.32 | 0.40 | 0.001432632 | 0.007515684 |
| 401207 | *C5orf63* | ENSG00000164241.14 | 1.67 | 0.35 | -1.32 | 0.40 | 0.001407469 | 0.007417379 |
| 389799 | *CFAP77* | ENSG00000188523.9 | 1.74 | 0.41 | -1.32 | 0.40 | 0.000155069 | 0.001350017 |
| 4856 | *CCN3* | ENSG00000136999.5 | 8.51 | 7.18 | -1.33 | 0.40 | 2.99348E-11 | 4.49291E-08 |
| 642852 | *LINC00205* | ENSG00000223768.3 | 5.74 | 4.40 | -1.35 | 0.39 | 0.000438288 | 0.003018936 |
| 503569 | *RGMB-AS1* | ENSG00000246763.7 | 0.18 | -1.17 | -1.35 | 0.39 | 0.000569387 | 0.003704344 |
| 64757 | *MTARC1* | ENSG00000186205.13 | 2.69 | 1.33 | -1.36 | 0.39 | 0.000222939 | 0.001788398 |
| 6753 | *SSTR3* | ENSG00000278195.2 | 1.78 | 0.41 | -1.37 | 0.39 | 0.00024537 | 0.001921103 |
| 285084 | *LINC01305* | ENSG00000231453.1 | 0.28 | -1.08 | -1.37 | 0.39 | 0.001932414 | 0.009442259 |
| 728723 | *ZBED3-AS1* | ENSG00000250802.7 | 2.16 | 0.80 | -1.37 | 0.39 | 0.000116729 | 0.001095679 |
| 257396 | *MOCS2-DT* | ENSG00000247796.3 | 3.00 | 1.63 | -1.37 | 0.39 | 2.6829E-06 | 6.95469E-05 |
| 133022 | *TRAM1L1* | ENSG00000174599.5 | 1.61 | 0.23 | -1.38 | 0.39 | 0.001204717 | 0.006584703 |
| 389602 | *LOC389602* | ENSG00000204876.5 | 3.19 | 1.81 | -1.38 | 0.39 | 1.17253E-06 | 3.89348E-05 |
| 100507421 | *TMEM178B* | ENSG00000261115.7 | 1.37 | -0.01 | -1.38 | 0.38 | 0.001341142 | 0.007153236 |
| 55959 | *SULF2* | ENSG00000196562.15 | 5.20 | 3.81 | -1.38 | 0.38 | 7.93506E-08 | 6.13903E-06 |
| 114898 | *C1QTNF2* | ENSG00000145861.9 | 1.42 | 0.03 | -1.39 | 0.38 | 0.000155207 | 0.001350435 |
| 9942 | *XYLB* | ENSG00000093217.11 | 1.98 | 0.58 | -1.39 | 0.38 | 0.000812964 | 0.004830474 |
| 340562 | *SATL1* | ENSG00000184788.14 | 2.20 | 0.81 | -1.40 | 0.38 | 0.000264144 | 0.002038322 |
| 137209 | *ZNF572* | ENSG00000180938.6 | 2.27 | 0.87 | -1.40 | 0.38 | 0.000200197 | 0.001642843 |
| 85301 | *COL27A1* | ENSG00000196739.15 | 3.13 | 1.74 | -1.40 | 0.38 | 0.000614557 | 0.003926728 |
| 29951 | *PDZRN4* | ENSG00000165966.16 | 0.99 | -0.41 | -1.41 | 0.38 | 0.000838803 | 0.004946798 |
| 1748 | *DLX4* | ENSG00000108813.11 | 1.97 | 0.55 | -1.41 | 0.38 | 0.000216191 | 0.001745457 |
| 105369147 | *LOC105369147* | ENSG00000247081.8 | 1.91 | 0.50 | -1.41 | 0.38 | 0.000154891 | 0.001349253 |
| 283008 | *NUTM2E* | ENSG00000228570.8 | 1.98 | 0.57 | -1.42 | 0.37 | 3.82828E-05 | 0.000486473 |
| 90668 | *CARMIL3* | ENSG00000186648.15 | 1.13 | -0.28 | -1.42 | 0.37 | 0.000140669 | 0.001257741 |
| 256933 | *NPB* | ENSG00000183979.8 | 0.77 | -0.64 | -1.42 | 0.37 | 0.000789694 | 0.004742901 |
| 83987 | *CCDC8* | ENSG00000169515.8 | 0.08 | -1.34 | -1.42 | 0.37 | 0.001733781 | 0.00866833 |
| 101927671 | *MIR202HG* | ENSG00000166917.11 | 0.40 | -1.02 | -1.43 | 0.37 | 0.000659675 | 0.004118581 |
| 283726 | *SAXO2* | ENSG00000188659.10 | 0.82 | -0.60 | -1.43 | 0.37 | 0.000802297 | 0.004789846 |
| 340533 | *NEXMIF* | ENSG00000050030.16 | 2.41 | 0.97 | -1.44 | 0.37 | 5.88766E-05 | 0.00066492 |
| 23284 | *ADGRL3* | ENSG00000150471.17 | 4.17 | 2.72 | -1.44 | 0.37 | 8.08213E-08 | 6.18901E-06 |
| 91409 | *CCDC74B* | ENSG00000152076.19 | 1.78 | 0.33 | -1.45 | 0.37 | 0.000620593 | 0.003951835 |
| 9641 | *IKBKE* | ENSG00000263528.8 | 2.92 | 1.48 | -1.45 | 0.37 | 0.000802866 | 0.00479134 |
| 83873 | *GPR61* | ENSG00000156097.13 | 4.12 | 2.66 | -1.46 | 0.36 | 5.35152E-06 | 0.000114908 |
| 100129129 | *TDH-AS1* | ENSG00000255020.1 | 2.16 | 0.70 | -1.46 | 0.36 | 0.000594258 | 0.003828393 |
| 92960 | *PEX11G* | ENSG00000104883.8 | 0.75 | -0.71 | -1.46 | 0.36 | 0.000359625 | 0.002590022 |
| 54332 | *GDAP1* | ENSG00000104381.14 | 5.74 | 4.27 | -1.47 | 0.36 | 3.71314E-09 | 7.33296E-07 |
| 3284 | *HSD3B2* | ENSG00000203859.10 | 5.27 | 3.80 | -1.47 | 0.36 | 6.27769E-09 | 9.63029E-07 |
| 256643 | *BCLAF3* | ENSG00000173681.17 | 3.49 | 2.02 | -1.47 | 0.36 | 1.44859E-06 | 4.45054E-05 |
| 60680 | *CELF5* | ENSG00000161082.13 | 2.81 | 1.33 | -1.47 | 0.36 | 7.09617E-05 | 0.000765683 |
| 54959 | *ODAM* | ENSG00000109205.17 | 0.26 | -1.21 | -1.48 | 0.36 | 0.001474208 | 0.007688114 |
| 1837 | *DTNA* | ENSG00000134769.23 | 5.67 | 4.18 | -1.50 | 0.35 | 1.23635E-07 | 8.32122E-06 |
| 285848 | *PNPLA1* | ENSG00000180316.13 | 3.76 | 2.25 | -1.50 | 0.35 | 2.98653E-05 | 0.000401656 |
| 100128881 | *VPS9D1-AS1* | ENSG00000261373.1 | 2.58 | 1.08 | -1.50 | 0.35 | 0.000256559 | 0.001991052 |
| 55231 | *CCDC87* | ENSG00000182791.5 | 0.38 | -1.13 | -1.50 | 0.35 | 0.000631045 | 0.003996351 |
| 23732 | *FRRS1L* | ENSG00000260230.5 | 1.44 | -0.06 | -1.51 | 0.35 | 0.001636512 | 0.008307498 |
| 113 | *ADCY7* | ENSG00000121281.13 | 2.41 | 0.90 | -1.51 | 0.35 | 7.76264E-06 | 0.000151903 |
| 174 | *AFP* | ENSG00000081051.8 | 1.95 | 0.44 | -1.52 | 0.35 | 0.000458637 | 0.003127524 |
| 7103 | *TSPAN8* | ENSG00000127324.9 | 1.71 | 0.18 | -1.52 | 0.35 | 0.002042588 | 0.009867141 |
| 1795 | *DOCK3* | ENSG00000088538.13 | 3.71 | 2.19 | -1.52 | 0.35 | 8.05182E-07 | 2.95476E-05 |
| 66004 | *LYNX1* | ENSG00000180155.20 | 0.98 | -0.55 | -1.53 | 0.35 | 0.002070445 | 0.009963229 |
| 140706 | *CCM2L* | ENSG00000101331.17 | 1.86 | 0.34 | -1.53 | 0.35 | 0.000122722 | 0.001131911 |
| 100506810 | *LINC01132* | ENSG00000227630.4 | 1.69 | 0.16 | -1.53 | 0.35 | 0.000188851 | 0.001569472 |
| 164633 | *CABP7* | ENSG00000100314.4 | 2.38 | 0.85 | -1.53 | 0.35 | 1.71813E-05 | 0.000268899 |
| 1268 | *CNR1* | ENSG00000118432.13 | 4.68 | 3.14 | -1.54 | 0.34 | 8.02975E-07 | 2.95476E-05 |
| 3755 | *KCNG1* | ENSG00000026559.14 | 1.11 | -0.43 | -1.54 | 0.34 | 0.000641453 | 0.004041456 |
| 114044 | *MCM3AP-AS1* | ENSG00000215424.10 | 2.01 | 0.46 | -1.55 | 0.34 | 0.000954906 | 0.00547239 |
| 102724474 | *LOC102724474* | ENSG00000283828.1 | 1.26 | -0.29 | -1.55 | 0.34 | 0.000766124 | 0.004636593 |
| 793 | *CALB1* | ENSG00000104327.7 | 5.98 | 4.43 | -1.55 | 0.34 | 1.8318E-09 | 4.8302E-07 |
| 84276 | *NICN1* | ENSG00000145029.14 | 2.08 | 0.52 | -1.56 | 0.34 | 1.56638E-05 | 0.000253886 |
| 84536 | *LINC01547* | ENSG00000183250.13 | 2.86 | 1.28 | -1.58 | 0.33 | 7.2692E-06 | 0.000143369 |
| 114769 | *CARD16* | ENSG00000204397.10 | 0.43 | -1.16 | -1.59 | 0.33 | 0.000852476 | 0.005009827 |
| 492 | *ATP2B3* | ENSG00000067842.18 | 5.27 | 3.69 | -1.59 | 0.33 | 1.8397E-08 | 2.14047E-06 |
| 140862 | *ISM1* | ENSG00000101230.6 | 5.62 | 4.02 | -1.60 | 0.33 | 3.04186E-10 | 1.45065E-07 |
| 283601 | *LINC00523* | ENSG00000196273.8 | 2.12 | 0.53 | -1.60 | 0.33 | 5.29626E-05 | 0.00061526 |
| 134265 | *AFAP1L1* | ENSG00000157510.14 | 1.52 | -0.08 | -1.62 | 0.33 | 0.000899828 | 0.005232669 |
| 5820 | *PVT1* | ENSG00000249859.13 | 0.86 | -0.75 | -1.62 | 0.33 | 7.50222E-05 | 0.000796892 |
| 4128 | *MAOA* | ENSG00000189221.11 | 1.09 | -0.54 | -1.63 | 0.32 | 0.000329679 | 0.002412421 |
| 9118 | *INA* | ENSG00000148798.11 | -0.05 | -1.69 | -1.63 | 0.32 | 0.000779611 | 0.004697383 |
| 101929074 | *PIK3CD-AS2* | ENSG00000231789.3 | -0.05 | -1.69 | -1.64 | 0.32 | 0.001018233 | 0.005775756 |
| 167681 | *PRSS35* | ENSG00000146250.7 | 1.60 | -0.03 | -1.64 | 0.32 | 7.01379E-05 | 0.000759523 |
| 56135 | *PCDHAC1* | ENSG00000248383.5 | 0.62 | -1.03 | -1.66 | 0.32 | 0.000360975 | 0.002594769 |
| 91646 | *TDRD12* | ENSG00000173809.18 | 1.86 | 0.20 | -1.66 | 0.32 | 0.000349189 | 0.002530651 |
| 340895 | *MALRD1* | ENSG00000204740.11 | -0.31 | -1.97 | -1.66 | 0.32 | 0.000500209 | 0.003344161 |
| 103724390 | *LINC01287* | ENSG00000234722.5 | -0.33 | -1.99 | -1.66 | 0.32 | 0.001149231 | 0.00636018 |
| 643783 | *USP46-DT* | ENSG00000248866.1 | 0.73 | -0.94 | -1.67 | 0.31 | 0.000377092 | 0.002686177 |
| 64857 | *PLEKHG2* | ENSG00000090924.15 | 4.69 | 2.98 | -1.70 | 0.31 | 0.000269895 | 0.00207351 |

| | | | | | | | | |
|---|---|---|---|---|---|---|---|---|
| 100506622 | *LINC00540* | ENSG00000276476.4 | 1.74 | 0.04 | -1.70 | 0.31 | 5.60067E-05 | 0.000640704 |
| 3747 | *KCNC2* | ENSG00000166006.14 | 3.29 | 1.58 | -1.70 | 0.31 | 1.58747E-07 | 9.96918E-06 |
| 26032 | *SUSD5* | ENSG00000173705.9 | 1.59 | -0.11 | -1.71 | 0.31 | 0.001293669 | 0.006966874 |
| 107985164 | *LOC107985164* | ENSG00000267327.2 | 2.26 | 0.55 | -1.71 | 0.31 | 0.000837344 | 0.004940133 |
| 5138 | *PDE2A* | ENSG00000186642.16 | 7.07 | 5.35 | -1.72 | 0.30 | 1.11873E-08 | 1.49919E-06 |
| 7433 | *VIPR1* | ENSG00000114812.13 | 3.87 | 2.13 | -1.74 | 0.30 | 1.46411E-07 | 9.35098E-06 |
| 10202 | *DHRS2* | ENSG00000100867.15 | 2.18 | 0.43 | -1.75 | 0.30 | 6.50315E-06 | 0.000132797 |
| 84894 | *LINGO1* | ENSG00000169783.13 | 4.33 | 2.57 | -1.75 | 0.30 | 4.34475E-06 | 9.89535E-05 |
| 83876 | *MRO* | ENSG00000134042.14 | -0.24 | -2.00 | -1.76 | 0.30 | 0.000440345 | 0.003028617 |
| 57127 | *RHBG* | ENSG00000132677.13 | 2.60 | 0.84 | -1.76 | 0.30 | 2.20345E-07 | 1.25271E-05 |
| 89846 | *FGD3* | ENSG00000127084.19 | 1.48 | -0.29 | -1.76 | 0.29 | 7.58193E-05 | 0.000803085 |
| 6004 | *RGS16* | ENSG00000143333.7 | 1.90 | 0.13 | -1.77 | 0.29 | 0.000101802 | 0.000998657 |
| 94 | *ACVRL1* | ENSG00000139567.13 | 2.62 | 0.83 | -1.78 | 0.29 | 1.87489E-06 | 5.32996E-05 |
| 23416 | *KCNH3* | ENSG00000135519.8 | 2.99 | 1.20 | -1.80 | 0.29 | 1.69276E-05 | 0.000266597 |
| 101928728 | *LOC101928728* | ENSG00000260063.1 | 0.96 | -0.82 | -1.80 | 0.29 | 2.99792E-05 | 0.000402828 |
| 80274 | *SCUBE1* | ENSG00000159307.20 | 0.79 | -1.02 | -1.80 | 0.29 | 0.001297104 | 0.006982865 |
| 283849 | *EXOC3L1* | ENSG00000179044.16 | 0.10 | -1.72 | -1.82 | 0.28 | 0.001801553 | 0.008947557 |
| 283050 | *ZMIZ1-AS1* | ENSG00000224596.9 | 1.25 | -0.57 | -1.82 | 0.28 | 4.78518E-05 | 0.000571367 |
| 126755 | *LRRC38* | ENSG00000162494.6 | 0.74 | -1.09 | -1.83 | 0.28 | 6.784E-05 | 0.000742136 |
| 653427 | *FOXD4L5* | ENSG00000204779.2 | -0.11 | -1.93 | -1.83 | 0.28 | 0.001578729 | 0.008084322 |
| 200931 | *SLC51A* | ENSG00000163959.10 | 1.01 | -0.83 | -1.84 | 0.28 | 0.000142029 | 0.001267369 |
| 11219 | *TREX2* | ENSG00000183479.13 | 1.94 | 0.10 | -1.85 | 0.28 | 0.001430924 | 0.007511974 |
| 3780 | *KCNN1* | ENSG00000105642.16 | 3.72 | 1.87 | -1.85 | 0.28 | 1.68001E-06 | 4.91526E-05 |
| 196872 | *LINC00638* | ENSG00000258701.1 | 1.82 | -0.04 | -1.86 | 0.28 | 2.43416E-06 | 6.51235E-05 |
| 2900 | *GRIK4* | ENSG00000149403.13 | 3.89 | 2.03 | -1.86 | 0.27 | 1.50653E-08 | 1.8534E-06 |
| 105371708 | *LINC01992* | ENSG00000260019.2 | 2.41 | 0.54 | -1.87 | 0.27 | 1.49456E-06 | 4.51344E-05 |
| 5457 | *POU4F1* | ENSG00000152192.8 | 1.75 | -0.14 | -1.89 | 0.27 | 6.27958E-05 | 0.000697633 |
| 100128025 | *WWTR1-AS1* | ENSG00000241313.3 | -0.78 | -2.70 | -1.93 | 0.26 | 0.000414003 | 0.002894161 |
| 11077 | *HSF2BP* | ENSG00000160207.9 | 0.06 | -1.86 | -1.93 | 0.26 | 0.000815178 | 0.004835973 |
| 22844 | *FRMPD1* | ENSG00000070601.10 | 0.55 | -1.39 | -1.94 | 0.26 | 0.001630109 | 0.008288042 |
| 3291 | *HSD11B2* | ENSG00000176387.7 | 1.15 | -0.79 | -1.95 | 0.26 | 0.000212564 | 0.001720347 |
| 347442 | *DCAF8L2* | ENSG00000189186.11 | 2.80 | 0.85 | -1.96 | 0.26 | 7.80355E-05 | 0.000822653 |
| 25854 | *FAM149A* | ENSG00000109794.14 | 1.12 | -0.85 | -1.97 | 0.25 | 0.000145628 | 0.00129333 |
| 286042 | *FAM86B3P* | ENSG00000173295.9 | 0.48 | -1.50 | -1.99 | 0.25 | 0.001556187 | 0.008009145 |
| 8875 | *VNN2* | ENSG00000112303.14 | -0.89 | -2.89 | -2.00 | 0.25 | 0.002018648 | 0.009779823 |
| 375704 | *ENHO* | ENSG00000168913.7 | 1.64 | -0.37 | -2.02 | 0.25 | 6.773E-06 | 0.000137373 |
| 93164 | *HTR7P1* | ENSG00000183935.5 | 1.21 | -0.81 | -2.03 | 0.25 | 0.000811912 | 0.004826131 |
| 5288 | *PIK3C2G* | ENSG00000139144.11 | 2.42 | 0.40 | -2.03 | 0.25 | 3.70684E-05 | 0.000476744 |
| 339929 | *LPP-AS2* | ENSG00000270959.1 | 1.27 | -0.79 | -2.05 | 0.24 | 7.87767E-06 | 0.000153155 |
| 55619 | *DOCK10* | ENSG00000135905.20 | 2.59 | 0.54 | -2.05 | 0.24 | 0.001235833 | 0.006722952 |
| 133690 | *CAPSL* | ENSG00000152611.12 | -0.32 | -2.38 | -2.07 | 0.24 | 0.001701183 | 0.008547493 |
| 2298 | *FOXD4* | ENSG00000170122.6 | -0.34 | -2.42 | -2.09 | 0.23 | 0.000890404 | 0.005187919 |
| 92340 | *PRR29* | ENSG00000224383.8 | 1.93 | -0.24 | -2.16 | 0.22 | 0.000102048 | 0.001000416 |
| 112268389 | *LOC112268389* | ENSG00000255052.5 | 1.26 | -0.92 | -2.18 | 0.22 | 2.54985E-05 | 0.000360364 |
| 101928489 | *CD109-AS1* | ENSG00000231652.2 | -1.24 | -3.41 | -2.18 | 0.22 | 0.000537244 | 0.003531974 |
| 2169 | *FABP2* | ENSG00000145384.4 | 1.58 | -0.62 | -2.19 | 0.22 | 2.5907E-06 | 6.82172E-05 |
| 9971 | *NR1H4* | ENSG00000012504.15 | 2.73 | 0.53 | -2.21 | 0.22 | 9.63215E-05 | 0.000954881 |
| 100131193 | *CCDC183-AS1* | ENSG00000228544.1 | 1.24 | -1.00 | -2.23 | 0.21 | 5.21745E-06 | 0.000112708 |
| 221938 | *MMD2* | ENSG00000136297.15 | 0.86 | -1.35 | -2.23 | 0.21 | 0.000200115 | 0.001642843 |
| 105376845 | *LOC105376845* | ENSG00000285873.1 | -0.64 | -2.90 | -2.25 | 0.21 | 0.00021956 | 0.001765064 |
| 2953 | *GSTT2* | ENSG00000099984.11 | 1.31 | -0.94 | -2.25 | 0.21 | 1.14018E-05 | 0.00020228 |
| 102724064 | *LOC102724064* | ENSG00000255468.7 | 0.82 | -1.42 | -2.25 | 0.21 | 0.001963302 | 0.009570381 |
| 5148 | *PDE6G* | ENSG00000185527.12 | -0.32 | -2.58 | -2.26 | 0.21 | 1.69824E-05 | 0.00026718 |
| 115677 | *NOSTRIN* | ENSG00000163072.16 | 3.01 | 0.75 | -2.27 | 0.21 | 5.3855E-07 | 2.28983E-05 |
| 160728 | *SLC5A8* | ENSG00000256870.3 | -0.23 | -2.52 | -2.28 | 0.21 | 0.000359066 | 0.002587371 |
| 7092 | *TLL1* | ENSG00000038295.8 | 0.37 | -1.93 | -2.30 | 0.20 | 0.000389249 | 0.002758373 |
| 9914 | *ATP2C2* | ENSG00000064270.13 | 0.69 | -1.62 | -2.30 | 0.20 | 0.000158816 | 0.001377839 |
| 25794 | *FSCN2* | ENSG00000186765.12 | -0.22 | -2.54 | -2.32 | 0.20 | 0.001601431 | 0.008167135 |
| 7694 | *ZNF135* | ENSG00000176293.20 | 1.08 | -1.26 | -2.34 | 0.20 | 4.81231E-05 | 0.000573237 |
| 85397 | *RGS8* | ENSG00000135824.12 | 2.09 | -0.27 | -2.36 | 0.20 | 0.000786345 | 0.004726573 |
| 8911 | *CACNA1I* | ENSG00000100346.18 | 0.81 | -1.58 | -2.39 | 0.19 | 3.14864E-06 | 7.85015E-05 |
| 101929567 | *LINC01814* | ENSG00000236008.4 | 3.19 | 0.78 | -2.41 | 0.19 | 2.55628E-09 | 5.46214E-07 |
| 881 | *CCIN* | ENSG00000185972.6 | -0.39 | -2.80 | -2.41 | 0.19 | 0.002076588 | 0.009986385 |
| 256536 | *TCERG1L* | ENSG00000176769.9 | 1.91 | -0.50 | -2.42 | 0.19 | 4.1286E-05 | 0.000511694 |
| 112267871 | *LOC112267871* | ENSG00000228549.5 | 0.20 | -2.23 | -2.44 | 0.18 | 0.000519914 | 0.003442065 |
| 729467 | *HSD52* | ENSG00000224609.8 | -0.04 | -2.67 | -2.64 | 0.16 | 1.90444E-05 | 0.000289602 |
| 56137 | *PCDHA12* | ENSG00000251664.5 | 0.62 | -2.15 | -2.76 | 0.15 | 0.000112773 | 0.001073995 |
| 4624 | *MYH6* | ENSG00000197616.13 | -0.09 | -2.90 | -2.81 | 0.14 | 0.001035072 | 0.005851376 |
| 728039 | *SSR4P1* | ENSG00000235374.2 | 2.04 | -0.77 | -2.83 | 0.14 | 0.000106759 | 0.001031774 |
| 8796 | *SCEL* | ENSG00000136155.17 | 1.86 | -0.97 | -2.83 | 0.14 | 0.000131132 | 0.001195723 |
| 8328 | *GFI1B* | ENSG00000165702.15 | 0.91 | -1.96 | -2.87 | 0.14 | 1.95121E-06 | 5.5152E-05 |
| 100507462 | *CCDC28A-AS1* | ENSG00000279968.3 | -0.05 | -2.97 | -2.92 | 0.13 | 0.000103479 | 0.001009015 |
| 100124700 | *HOTAIR* | ENSG00000228630.5 | 0.20 | -2.77 | -2.99 | 0.13 | 0.001358365 | 0.007215973 |
| 654780 | *LOC654780* | ENSG00000250685.7 | 0.11 | -2.92 | -3.02 | 0.12 | 7.94524E-05 | 0.000833617 |
| 85478 | *CCDC65* | ENSG00000139537.11 | 0.96 | -2.08 | -3.03 | 0.12 | 0.000226495 | 0.001810148 |
| 56145 | *PCDHA3* | ENSG00000255408.4 | 0.29 | -2.80 | -3.08 | 0.12 | 5.16069E-05 | 0.000602308 |
| 6339 | *SCNN1D* | ENSG00000162572.21 | -0.88 | -4.08 | -3.21 | 0.11 | 0.001330699 | 0.007111368 |
| 100129781 | *MIR193BHG* | ENSG00000262454.4 | 0.59 | -2.61 | -3.23 | 0.11 | 0.000140185 | 0.001256144 |
| 146712 | *B3GNTL1* | ENSG00000175711.9 | 1.25 | -2.07 | -3.32 | 0.10 | 1.47429E-05 | 0.000241834 |
| 114794 | *ELFN2* | ENSG00000166897.16 | 0.86 | -2.50 | -3.35 | 0.10 | 0.000909461 | 0.005274382 |
| 101928737 | *ZCCHC14-DT* | ENSG00000288568.1 | -0.84 | -4.32 | -3.46 | 0.09 | 0.00113321 | 0.006301723 |
| 100506776 | *TRG-AS1* | ENSG00000281103.3 | -0.24 | -3.81 | -3.59 | 0.08 | 0.000398144 | 0.002805512 |

**Supplemental Table 6.** RNAseq analysis of *SLC30A1*$^{WT}$ and *SLC30A1*$^{51\_57del}$ at 24 hours of Doxy induction

| Entrez ID | Symbol | Ensembl | *SLC30A1*$^{51\_57del}$ Doxy 24h | *SLC30A1*$^{WT}$ Doxy 24h | *SLC30A1*$^{51\_57del}$ vs. *SLC30A1*$^{WT}$ Log FC | *SLC30A1*$^{51\_57del}$ vs. *SLC30A1*$^{WT}$ FC | P.Value | adj.P.Value (Significance P < 0.01) |
|---|---|---|---|---|---|---|---|---|
| 4880 | *NPPC* | ENSG00000163273.4 | -0.07 | -4.32 | 4.23 | 18.83 | 0.000461453 | 0.0035793 |
| 83850 | *ESYT3* | ENSG00000158220.14 | -0.11 | -4.21 | 4.09 | 17.07 | 0.001208384 | 0.007280868 |
| 1585 | *CYP11B2* | ENSG00000179142.2 | 7.66 | 3.58 | 4.08 | 16.86 | 1.19742E-11 | 8.98603E-08 |
| 1339 | *COX6A2* | ENSG00000156885.6 | -0.24 | -4.06 | 3.83 | 14.20 | 0.00059215 | 0.004301826 |
| 3560 | *IL2RB* | ENSG00000100385.14 | 0.34 | -3.12 | 3.46 | 11.04 | 0.000711537 | 0.004914011 |
| 7857 | *SCG2* | ENSG00000171951.5 | 2.90 | -0.40 | 3.28 | 9.74 | 2.5908E-07 | 1.81707E-05 |
| 729739 | *PPIAP46* | ENSG00000260266.1 | 2.91 | -0.18 | 3.07 | 8.42 | 4.34705E-06 | 0.000117347 |
| 401190 | *RGS7BP* | ENSG00000186479.5 | 1.07 | -1.90 | 2.96 | 7.78 | 0.000761412 | 0.005157052 |
| 4929 | *NR4A2* | ENSG00000153234.15 | 4.50 | 1.66 | 2.85 | 7.20 | 5.79234E-08 | 6.16576E-06 |
| 9750 | *RIPOR2* | ENSG00000111913.20 | 2.33 | -0.49 | 2.80 | 6.99 | 6.99768E-06 | 0.000165139 |
| 3270 | *HRC* | ENSG00000130528.12 | 1.68 | -1.09 | 2.79 | 6.89 | 0.000871069 | 0.005706625 |
| 154790 | *CLEC2L* | ENSG00000236279.7 | 1.23 | -1.41 | 2.64 | 6.23 | 0.000415128 | 0.003319476 |
| 94234 | *FOXQ1* | ENSG00000164379.7 | 0.50 | -2.13 | 2.64 | 6.22 | 0.000174054 | 0.001734646 |
| 5105 | *PCK1* | ENSG00000124253.11 | 1.58 | -1.05 | 2.62 | 6.14 | 0.000706934 | 0.004891828 |
| 25791 | *NGEF* | ENSG00000066248.15 | 7.96 | 5.34 | 2.61 | 6.12 | 3.72208E-09 | 1.10195E-06 |
| 388662 | *SLC6A17* | ENSG00000197106.7 | 4.71 | 2.14 | 2.57 | 5.93 | 2.19809E-06 | 7.44066E-05 |
| 11211 | *FZD10* | ENSG00000111432.5 | 1.70 | -0.80 | 2.50 | 5.66 | 1.64065E-05 | 0.000310132 |
| 26018 | *LRIG1* | ENSG00000144749.14 | 2.50 | 0.02 | 2.47 | 5.53 | 0.000220661 | 0.002074333 |
| 26207 | *PITPNC1* | ENSG00000154217.16 | 6.94 | 4.52 | 2.42 | 5.35 | 4.15558E-10 | 2.97005E-07 |
| 109729181 | *ZNF710-AS1* | ENSG00000259291.2 | 4.38 | 1.97 | 2.41 | 5.31 | 1.88248E-07 | 1.43703E-05 |
| 390937 | *ERFL* | ENSG00000268041.3 | 1.73 | -0.68 | 2.39 | 5.25 | 0.000190753 | 0.001856692 |
| 6863 | *TAC1* | ENSG00000006128.12 | 1.49 | -0.85 | 2.33 | 5.01 | 7.69576E-05 | 0.00097227 |
| 28513 | *CDH19* | ENSG00000071991.9 | -1.13 | -3.44 | 2.31 | 4.95 | 0.001759795 | 0.009667923 |
| 64168 | *NECAB1* | ENSG00000123119.12 | -0.29 | -2.48 | 2.21 | 4.62 | 0.000974293 | 0.006159713 |
| 29970 | *SCHIP1* | ENSG00000283154.3 | 3.03 | 0.83 | 2.20 | 4.60 | 1.88616E-07 | 1.43703E-05 |
| 80144 | *FRAS1* | ENSG00000138759.20 | 8.07 | 5.87 | 2.20 | 4.59 | 6.18943E-10 | 3.87071E-07 |
| 6324 | *SCN1B* | ENSG00000105711.14 | 1.81 | -0.34 | 2.15 | 4.44 | 0.000375867 | 0.003070983 |
| 337 | *APOA4* | ENSG00000110244.7 | -0.12 | -2.26 | 2.14 | 4.41 | 0.001513771 | 0.008648718 |
| 359845 | *RFLNB* | ENSG00000183688.4 | 6.37 | 4.26 | 2.11 | 4.31 | 1.89138E-10 | 2.00923E-07 |
| 11069 | *RAPGEF4* | ENSG00000091428.18 | 7.32 | 5.22 | 2.10 | 4.30 | 3.79807E-08 | 4.75044E-06 |
| 10221 | *TRIB1* | ENSG00000173334.4 | 5.62 | 3.54 | 2.09 | 4.25 | 1.84673E-07 | 1.42142E-05 |
| 100133941 | *CD24* | ENSG00000272398.6 | 1.69 | -0.38 | 2.06 | 4.17 | 0.00049437 | 0.003758869 |
| 2827 | *GPR3* | ENSG00000181773.7 | 2.62 | 0.55 | 2.06 | 4.16 | 0.000134588 | 0.001443907 |
| 8736 | *MYOM1* | ENSG00000101605.14 | 7.38 | 5.33 | 2.05 | 4.13 | 1.78056E-08 | 2.84302E-06 |
| 3624 | *INHBA* | ENSG00000122641.11 | 4.54 | 2.51 | 2.04 | 4.10 | 1.92574E-08 | 2.93222E-06 |
| 973 | *CD79A* | ENSG00000105369.10 | 6.11 | 4.08 | 2.03 | 4.09 | 6.46959E-11 | 1.29661E-07 |
| 688 | *KLF5* | ENSG00000102554.14 | 3.93 | 1.90 | 2.02 | 4.06 | 4.02338E-05 | 0.000611068 |
| 359 | *AQP2* | ENSG00000167580.8 | 4.90 | 2.88 | 2.02 | 4.05 | 2.88065E-05 | 0.000474597 |
| 81792 | *ADAMTS12* | ENSG00000151388.11 | 3.15 | 1.15 | 2.00 | 4.00 | 7.00934E-07 | 3.5422E-05 |
| 344838 | *PAQR9* | ENSG00000188582.10 | 2.95 | 0.97 | 1.98 | 3.95 | 2.10244E-06 | 7.20414E-05 |
| 148641 | *SLC35F3* | ENSG00000183780.13 | 2.28 | 0.31 | 1.98 | 3.93 | 1.5486E-05 | 0.000295711 |
| 2303 | *FOXC2* | ENSG00000176692.8 | 2.07 | 0.10 | 1.97 | 3.93 | 2.86162E-06 | 8.74745E-05 |
| 4158 | *MC2R* | ENSG00000185231.5 | 5.46 | 3.49 | 1.96 | 3.90 | 2.31483E-06 | 7.68656E-05 |
| 687 | *KLF9* | ENSG00000119138.5 | 3.19 | 1.26 | 1.93 | 3.80 | 1.14953E-06 | 4.87381E-05 |
| 627 | *BDNF* | ENSG00000176697.20 | 3.24 | 1.32 | 1.92 | 3.78 | 1.21325E-06 | 5.07234E-05 |
| 222546 | *RFX6* | ENSG00000185002.10 | 3.60 | 1.69 | 1.91 | 3.77 | 1.09861E-06 | 4.73821E-05 |
| 23764 | *MAFF* | ENSG00000185022.12 | 3.24 | 1.39 | 1.86 | 3.62 | 3.54484E-06 | 0.000102316 |
| 64359 | *NXN* | ENSG00000167693.17 | 5.67 | 3.81 | 1.85 | 3.62 | 1.15096E-05 | 0.000238931 |
| 111 | *ADCY5* | ENSG00000173175.15 | 0.90 | -0.97 | 1.85 | 3.61 | 0.000323532 | 0.002748102 |
| 1584 | *CYP11B1* | ENSG00000160882.13 | 1.80 | -0.04 | 1.85 | 3.60 | 6.50485E-05 | 0.000865526 |
| 9466 | *IL27RA* | ENSG00000104998.4 | 0.80 | -1.05 | 1.84 | 3.59 | 0.000407843 | 0.003268185 |
| 9510 | *ADAMTS1* | ENSG00000154734.16 | 2.96 | 1.13 | 1.84 | 3.58 | 3.59991E-05 | 0.00055933 |
| 25816 | *TNFAIP8* | ENSG00000145779.8 | 6.13 | 4.29 | 1.84 | 3.57 | 1.33796E-08 | 2.2935E-06 |
| 22795 | *NID2* | ENSG00000087303.19 | 5.35 | 3.51 | 1.83 | 3.57 | 2.06847E-07 | 1.51442E-05 |
| 2239 | *GPC4* | ENSG00000076716.9 | 9.03 | 7.21 | 1.81 | 3.51 | 2.17414E-12 | 3.26316E-08 |
| 387763 | *C11orf96* | ENSG00000187479.8 | 4.24 | 2.47 | 1.77 | 3.41 | 6.72508E-07 | 3.45161E-05 |
| 3491 | *CCN1* | ENSG00000142871.18 | 6.39 | 4.67 | 1.72 | 3.29 | 1.892E-08 | 2.92753E-06 |
| 222545 | *GPRC6A* | ENSG00000173612.10 | 1.76 | 0.04 | 1.71 | 3.28 | 0.000676995 | 0.004728252 |
| 6622 | *SNCA* | ENSG00000145335.17 | 5.35 | 3.64 | 1.71 | 3.26 | 1.20062E-05 | 0.000246513 |
| 90993 | *CREB3L1* | ENSG00000157613.11 | 0.91 | -0.78 | 1.69 | 3.22 | 0.000215117 | 0.002035749 |
| 23704 | *KCNE4* | ENSG00000152049.7 | 6.94 | 5.25 | 1.69 | 3.22 | 5.24364E-11 | 1.29661E-07 |
| 2571 | *GAD1* | ENSG00000128683.14 | 3.27 | 1.58 | 1.68 | 3.21 | 4.95115E-05 | 0.000709162 |
| 57419 | *SLC24A3* | ENSG00000185052.13 | 5.06 | 3.38 | 1.68 | 3.20 | 1.52777E-09 | 6.7442E-07 |
| 7042 | *TGFB2* | ENSG00000092969.12 | 6.80 | 5.13 | 1.68 | 3.20 | 1.87128E-09 | 7.20156E-07 |
| 4883 | *NPR3* | ENSG00000113389.16 | 3.56 | 1.90 | 1.67 | 3.19 | 0.000504154 | 0.003815856 |
| 84623 | *KIRREL3* | ENSG00000149571.12 | 4.34 | 2.68 | 1.66 | 3.15 | 7.1456E-06 | 0.000167313 |
| 4783 | *NFIL3* | ENSG00000165030.4 | 6.60 | 4.96 | 1.64 | 3.12 | 2.00803E-10 | 2.00923E-07 |
| 57709 | *SLC7A14* | ENSG00000013293.6 | 1.84 | 0.22 | 1.63 | 3.09 | 0.00026947 | 0.002418943 |
| 27098 | *CLUL1* | ENSG00000079101.17 | 5.00 | 3.40 | 1.60 | 3.03 | 2.49019E-07 | 1.77134E-05 |
| 2118 | *ETV4* | ENSG00000175832.13 | 3.33 | 1.74 | 1.59 | 3.01 | 8.47086E-05 | 0.001037025 |
| 554282 | *FAM72C* | ENSG00000263513.6 | 5.60 | 4.01 | 1.59 | 3.00 | 6.73142E-09 | 1.62955E-06 |
| 81606 | *LBH* | ENSG00000213626.13 | 5.40 | 3.82 | 1.59 | 3.00 | 2.22343E-05 | 0.000390194 |
| 643201 | *LOC643201* | ENSG00000248596.8 | 4.17 | 2.59 | 1.58 | 3.00 | 1.35518E-06 | 5.45306E-05 |
| 107986100 | *LINC02018* | ENSG00000272690.6 | -0.29 | -1.88 | 1.58 | 3.00 | 0.001836071 | 0.009991919 |
| 6546 | *SLC8A1* | ENSG00000183023.18 | 3.65 | 2.09 | 1.57 | 2.98 | 0.001671058 | 0.00932849 |
| 1604 | *CD55* | ENSG00000196352.17 | 7.80 | 6.23 | 1.57 | 2.97 | 2.21662E-10 | 2.07933E-07 |
| 55117 | *SLC6A15* | ENSG00000072041.18 | 3.08 | 1.52 | 1.56 | 2.94 | 0.000545623 | 0.004054084 |
| 8553 | *BHLHE40* | ENSG00000134107.5 | 5.49 | 3.94 | 1.55 | 2.92 | 2.40309E-07 | 1.71752E-05 |
| 728833 | *FAM72D* | ENSG00000215784.6 | 5.97 | 4.42 | 1.54 | 2.92 | 1.00366E-08 | 1.93127E-06 |
| 9248 | *GPR50* | ENSG00000102195.10 | 2.89 | 1.34 | 1.54 | 2.91 | 0.000141804 | 0.001501998 |
| 9242 | *MSC* | ENSG00000178860.9 | 1.21 | -0.33 | 1.53 | 2.90 | 0.000169137 | 0.001696906 |
| 57633 | *LRRN1* | ENSG00000175928.6 | 4.38 | 2.87 | 1.52 | 2.87 | 7.3912E-05 | 0.000943862 |

| | | | | | | | | |
|---|---|---|---|---|---|---|---|---|
| 8447 | *DOC2B* | ENSG00000272636.4 | 6.50 | 4.99 | 1.50 | 2.84 | 7.13765E-09 | 1.63338E-06 |
| 57480 | *PLEKHG1* | ENSG00000120278.16 | 4.26 | 2.77 | 1.50 | 2.84 | 9.01262E-05 | 0.00108303 |
| 384 | *ARG2* | ENSG00000081181.8 | 6.69 | 5.19 | 1.50 | 2.83 | 7.31925E-10 | 4.22518E-07 |
| 3953 | *LEPR* | ENSG00000116678.20 | 4.49 | 3.00 | 1.49 | 2.82 | 2.90807E-07 | 1.93958E-05 |
| 5592 | *PRKG1* | ENSG00000185532.20 | 3.54 | 2.05 | 1.48 | 2.79 | 0.000387348 | 0.003134074 |
| 6422 | *SFRP1* | ENSG00000104332.12 | 7.73 | 6.25 | 1.48 | 2.79 | 6.99488E-11 | 1.29661E-07 |
| 84570 | *COL25A1* | ENSG00000188517.17 | 8.04 | 6.57 | 1.47 | 2.77 | 8.71663E-10 | 4.67242E-07 |
| 64577 | *ALDH8A1* | ENSG00000118514.14 | 1.91 | 0.44 | 1.47 | 2.76 | 0.000850251 | 0.00561435 |
| 4430 | *MYO1B* | ENSG00000128641.19 | 4.99 | 3.53 | 1.46 | 2.76 | 8.40325E-08 | 8.23623E-06 |
| 3418 | *IDH2* | ENSG00000182054.10 | 8.03 | 6.57 | 1.45 | 2.74 | 6.06051E-11 | 1.29661E-07 |
| 2119 | *ETV5* | ENSG00000244405.8 | 7.28 | 5.84 | 1.44 | 2.71 | 1.83603E-08 | 2.90074E-06 |
| 1647 | *GADD45A* | ENSG00000116717.13 | 4.62 | 3.19 | 1.44 | 2.71 | 2.65813E-07 | 1.84245E-05 |
| 3516 | *RBPJ* | ENSG00000168214.22 | 9.19 | 7.76 | 1.43 | 2.70 | 4.48675E-10 | 3.06098E-07 |
| 23554 | *TSPAN12* | ENSG00000106025.9 | 8.29 | 6.86 | 1.43 | 2.69 | 3.93222E-11 | 1.29661E-07 |
| 2669 | *GEM* | ENSG00000164949.8 | 4.61 | 3.18 | 1.43 | 2.69 | 1.72999E-06 | 6.34851E-05 |
| 729533 | *FAM72A* | ENSG00000196550.11 | 5.56 | 4.14 | 1.42 | 2.68 | 4.62878E-07 | 2.67474E-05 |
| 1846 | *DUSP4* | ENSG00000120875.9 | 2.24 | 0.82 | 1.42 | 2.68 | 1.11047E-05 | 0.000232132 |
| 6303 | *SAT1* | ENSG00000130066.17 | 6.04 | 4.62 | 1.42 | 2.68 | 3.74437E-09 | 1.10195E-06 |
| 26050 | *SLITRK5* | ENSG00000165300.8 | 2.38 | 0.96 | 1.41 | 2.66 | 0.000103779 | 0.001198166 |
| 481 | *ATP1B1* | ENSG00000143153.14 | 9.04 | 7.62 | 1.41 | 2.66 | 8.85687E-11 | 1.29661E-07 |
| 91373 | *UAP1L1* | ENSG00000197355.11 | 5.16 | 3.77 | 1.40 | 2.64 | 1.33573E-08 | 2.2935E-06 |
| 128209 | *KLF17* | ENSG00000171872.5 | 3.94 | 2.54 | 1.40 | 2.64 | 2.38692E-06 | 7.78812E-05 |
| 114134 | *SLC2A13* | ENSG00000151229.13 | 7.36 | 5.96 | 1.40 | 2.63 | 5.0431E-10 | 3.29095E-07 |
| 27129 | *HSPB7* | ENSG00000173641.18 | 8.47 | 7.07 | 1.39 | 2.63 | 2.06572E-09 | 7.5769E-07 |
| 3751 | *KCND2* | ENSG00000184408.10 | 3.30 | 1.91 | 1.39 | 2.62 | 0.000210789 | 0.00200875 |
| 100506827 | *LINC02475* | ENSG00000251350.2 | 4.92 | 3.53 | 1.39 | 2.61 | 4.87588E-06 | 0.000127405 |
| 340485 | *ACER2* | ENSG00000177076.6 | 2.84 | 1.46 | 1.38 | 2.60 | 3.59719E-06 | 0.000103232 |
| 56521 | *DNAJC12* | ENSG00000108176.15 | 5.93 | 4.55 | 1.38 | 2.60 | 7.07201E-09 | 1.63338E-06 |
| 654 | *BMP6* | ENSG00000153162.9 | 5.71 | 4.34 | 1.37 | 2.59 | 1.55208E-08 | 2.5599E-06 |
| 84624 | *FNDC1* | ENSG00000164694.17 | 3.07 | 1.71 | 1.37 | 2.58 | 0.000557545 | 0.004116179 |
| 883 | *KYAT1* | ENSG00000171097.14 | 3.51 | 2.15 | 1.36 | 2.56 | 1.61971E-05 | 0.00030656 |
| 219539 | *YPEL4* | ENSG00000166793.13 | 1.02 | -0.34 | 1.36 | 2.56 | 0.001060838 | 0.006576669 |
| 58476 | *TP53INP2* | ENSG00000078804.13 | 7.00 | 5.65 | 1.35 | 2.55 | 6.52127E-09 | 1.60455E-06 |
| 26278 | *SACS* | ENSG00000151835.17 | 8.63 | 7.28 | 1.35 | 2.55 | 1.37874E-07 | 1.15621E-05 |
| 4646 | *MYO6* | ENSG00000196586.17 | 7.52 | 6.17 | 1.34 | 2.54 | 2.11552E-09 | 7.5769E-07 |
| 388730 | *TMEM81* | ENSG00000174529.7 | 1.81 | 0.46 | 1.34 | 2.53 | 0.000182718 | 0.001800669 |
| 653820 | *FAM72B* | ENSG00000188610.12 | 5.83 | 4.50 | 1.33 | 2.51 | 5.32325E-08 | 5.87475E-06 |
| 5033 | *P4HA1* | ENSG00000122884.13 | 7.33 | 6.00 | 1.32 | 2.50 | 2.13747E-09 | 7.5769E-07 |
| 9021 | *SOCS3* | ENSG00000184557.4 | 4.00 | 2.69 | 1.31 | 2.49 | 1.37891E-07 | 1.15621E-05 |
| 970 | *CD70* | ENSG00000125726.11 | 2.38 | 1.06 | 1.31 | 2.48 | 0.000472906 | 0.003639923 |
| 63973 | *NEUROG2* | ENSG00000178403.4 | 1.59 | 0.29 | 1.31 | 2.48 | 0.001710507 | 0.009483932 |
| 115827 | *RAB3C* | ENSG00000152932.8 | 7.86 | 6.55 | 1.31 | 2.48 | 1.79316E-06 | 6.51661E-05 |
| 27022 | *FOXD3* | ENSG00000187140.6 | 2.33 | 1.03 | 1.31 | 2.48 | 2.98233E-05 | 0.000488133 |
| 4493 | *MT1E* | ENSG00000169715.15 | 5.93 | 4.63 | 1.30 | 2.47 | 1.3106E-08 | 2.2935E-06 |
| 10560 | *SLC19A2* | ENSG00000117479.15 | 6.93 | 5.63 | 1.30 | 2.46 | 3.54295E-09 | 1.08523E-06 |
| 6533 | *SLC6A6* | ENSG00000131389.18 | 8.82 | 7.53 | 1.29 | 2.45 | 1.65636E-10 | 2.00923E-07 |
| 283209 | *PGM2L1* | ENSG00000165434.8 | 7.49 | 6.21 | 1.28 | 2.44 | 4.1346E-10 | 2.97005E-07 |
| 114990 | *VASN* | ENSG00000168140.5 | 3.07 | 1.78 | 1.28 | 2.43 | 4.13167E-06 | 0.000113575 |
| 170850 | *KCNG3* | ENSG00000171126.8 | 2.02 | 0.74 | 1.28 | 2.42 | 4.69428E-05 | 0.000684043 |
| 11015 | *KDELR3* | ENSG00000100196.11 | 4.33 | 3.06 | 1.27 | 2.41 | 1.33178E-06 | 5.43172E-05 |
| 9497 | *SLC4A7* | ENSG00000033867.17 | 5.78 | 4.52 | 1.26 | 2.40 | 3.10333E-07 | 1.99905E-05 |
| 51330 | *TNFRSF12A* | ENSG00000006327.14 | 4.73 | 3.46 | 1.26 | 2.40 | 6.87365E-06 | 0.000164017 |
| 6506 | *SLC1A2* | ENSG00000110436.13 | 3.40 | 2.14 | 1.26 | 2.40 | 0.00029996 | 0.002615981 |
| 2296 | *FOXC1* | ENSG00000054598.9 | 3.87 | 2.61 | 1.26 | 2.39 | 6.68252E-06 | 0.000160278 |
| 27231 | *NMRK2* | ENSG00000077009.14 | 2.78 | 1.54 | 1.24 | 2.37 | 0.001582371 | 0.008963362 |
| 5029 | *P2RY2* | ENSG00000175591.12 | 4.91 | 3.68 | 1.23 | 2.35 | 7.1213E-07 | 3.55095E-05 |
| 2274 | *FHL2* | ENSG00000115641.19 | 7.05 | 5.82 | 1.22 | 2.34 | 1.78526E-09 | 7.0513E-07 |
| 4121 | *MAN1A1* | ENSG00000111885.7 | 5.48 | 4.26 | 1.22 | 2.33 | 9.91828E-08 | 9.42174E-06 |
| 4241 | *MELTF* | ENSG00000163975.13 | 3.86 | 2.64 | 1.22 | 2.33 | 5.42138E-05 | 0.000754819 |
| 26503 | *SLC17A5* | ENSG00000119899.13 | 6.24 | 5.02 | 1.22 | 2.33 | 1.93513E-07 | 1.46445E-05 |
| 4747 | *NEFL* | ENSG00000277586.4 | 8.15 | 6.93 | 1.22 | 2.33 | 9.50274E-11 | 1.29661E-07 |
| 245806 | *VGLL2* | ENSG00000170162.14 | 2.72 | 1.50 | 1.22 | 2.33 | 0.001178522 | 0.007143958 |
| 8912 | *CACNA1H* | ENSG00000196557.13 | 6.45 | 5.24 | 1.21 | 2.31 | 1.09306E-07 | 1.00649E-05 |
| 29785 | *CYP2S1* | ENSG00000167600.14 | 3.52 | 2.32 | 1.21 | 2.31 | 0.000645157 | 0.004567526 |
| 54855 | *TENT5C* | ENSG00000183508.5 | 5.90 | 4.69 | 1.21 | 2.31 | 4.41339E-08 | 5.18389E-06 |
| 5764 | *PTN* | ENSG00000105894.12 | 7.83 | 6.63 | 1.20 | 2.30 | 3.15009E-10 | 2.78116E-07 |
| 9762 | *LZTS3* | ENSG00000088899.16 | 4.35 | 3.15 | 1.20 | 2.29 | 2.11142E-06 | 7.20414E-05 |
| 3570 | *IL6R* | ENSG00000160712.13 | 3.15 | 1.97 | 1.19 | 2.28 | 0.000479327 | 0.003666779 |
| 1305 | *COL13A1* | ENSG00000197467.17 | 3.43 | 2.24 | 1.19 | 2.28 | 2.87264E-05 | 0.000474597 |
| 8828 | *NRP2* | ENSG00000118257.17 | 8.14 | 6.95 | 1.19 | 2.27 | 2.99469E-07 | 1.96276E-05 |
| 100302736 | *TMED7-TICAI* | ENSG00000251201.8 | 5.24 | 4.04 | 1.18 | 2.27 | 3.00935E-06 | 9.10631E-05 |
| 6482 | *ST3GAL1* | ENSG00000008513.16 | 6.84 | 5.65 | 1.18 | 2.27 | 4.7376E-08 | 5.42799E-06 |
| 51621 | *KLF13* | ENSG00000169926.11 | 3.46 | 2.27 | 1.18 | 2.27 | 0.001152102 | 0.007009285 |
| 55244 | *SLC47A1* | ENSG00000142494.13 | 8.30 | 7.12 | 1.18 | 2.27 | 3.17223E-09 | 9.91918E-07 |
| 23302 | *WSCD1* | ENSG00000179314.16 | 5.48 | 4.30 | 1.17 | 2.26 | 8.65396E-07 | 4.05731E-05 |
| 5165 | *PDK3* | ENSG00000067992.17 | 7.44 | 6.27 | 1.17 | 2.25 | 1.68285E-07 | 1.35069E-05 |
| 23493 | *HEY2* | ENSG00000135547.9 | 3.93 | 2.77 | 1.16 | 2.24 | 0.00042999 | 0.003394906 |
| 27309 | *ZNF330* | ENSG00000109445.11 | 7.41 | 6.25 | 1.16 | 2.24 | 3.07658E-09 | 9.82477E-07 |
| 2355 | *FOSL2* | ENSG00000075426.12 | 6.97 | 5.81 | 1.16 | 2.23 | 1.17042E-08 | 2.1423E-06 |
| 84302 | *PGAP4* | ENSG00000165152.9 | 8.59 | 7.44 | 1.16 | 2.23 | 3.91929E-10 | 2.97005E-07 |
| 3632 | *INPP5A* | ENSG00000068383.19 | 7.30 | 6.14 | 1.16 | 2.23 | 4.37068E-08 | 5.18389E-06 |
| 83604 | *TMEM47* | ENSG00000147027.4 | 10.47 | 9.31 | 1.16 | 2.23 | 3.721E-11 | 1.29661E-07 |
| 441061 | *MARCHF11* | ENSG00000183654.9 | 1.95 | 0.80 | 1.15 | 2.22 | 0.000160469 | 0.001640098 |
| 4811 | *NID1* | ENSG00000116962.15 | 8.79 | 7.65 | 1.15 | 2.21 | 1.66588E-09 | 6.93414E-07 |
| 6517 | *SLC2A4* | ENSG00000181856.15 | 5.48 | 4.34 | 1.14 | 2.21 | 3.62805E-08 | 4.57591E-06 |
| 538 | *ATP7A* | ENSG00000165240.22 | 7.37 | 6.23 | 1.14 | 2.21 | 9.88187E-09 | 1.93127E-06 |
| 51661 | *FKBP7* | ENSG00000079150.19 | 5.50 | 4.36 | 1.14 | 2.20 | 2.22026E-06 | 7.45501E-05 |
| 415116 | *PIM3* | ENSG00000198355.5 | 5.19 | 4.05 | 1.13 | 2.20 | 1.60186E-07 | 1.29959E-05 |
| 23417 | *MLYCD* | ENSG00000103150.7 | 2.73 | 1.60 | 1.13 | 2.19 | 0.00142262 | 0.008253617 |

| | | | | | | | | |
|---|---|---|---|---|---|---|---|---|
| 7301 | *TYRO3* | ENSG00000092445.12 | 7.17 | 6.04 | 1.13 | 2.19 | 1.02007E-08 | 1.938E-06 |
| 10253 | *SPRY2* | ENSG00000136158.12 | 3.27 | 2.13 | 1.13 | 2.19 | 2.22669E-05 | 0.000390194 |
| 54842 | *MFSD6* | ENSG00000151690.15 | 6.39 | 5.26 | 1.13 | 2.19 | 5.56659E-06 | 0.000139481 |
| 145270 | *PRIMA1* | ENSG00000175785.13 | 3.69 | 2.56 | 1.13 | 2.19 | 2.25642E-05 | 0.000394257 |
| 127544 | *RNF19B* | ENSG00000116514.17 | 6.52 | 5.39 | 1.13 | 2.19 | 2.37693E-08 | 3.39766E-06 |
| 22808 | *MRAS* | ENSG00000158186.13 | 8.32 | 7.20 | 1.12 | 2.18 | 9.14889E-07 | 4.224E-05 |
| 3486 | *IGFBP3* | ENSG00000146674.16 | 4.79 | 3.67 | 1.11 | 2.17 | 4.50047E-07 | 2.64893E-05 |
| 378805 | *LINC-PINT* | ENSG00000231721.7 | 3.29 | 2.18 | 1.11 | 2.16 | 0.001606946 | 0.00906035 |
| 51765 | *STK26* | ENSG00000134602.16 | 6.46 | 5.35 | 1.11 | 2.16 | 2.18277E-09 | 7.5769E-07 |
| 105378853 | *LINC02609* | ENSG00000233593.11 | 2.89 | 1.79 | 1.11 | 2.16 | 0.00017365 | 0.00173292 |
| 2027 | *ENO3* | ENSG00000108515.18 | 5.79 | 4.69 | 1.10 | 2.15 | 3.7478E-07 | 2.28662E-05 |
| 4501 | *MT1X* | ENSG00000187193.9 | 5.55 | 4.45 | 1.10 | 2.14 | 5.60428E-06 | 0.000140191 |
| 3164 | *NR4A1* | ENSG00000123358.20 | 7.84 | 6.74 | 1.10 | 2.14 | 1.62557E-09 | 6.93414E-07 |
| 253559 | *CADM2* | ENSG00000175161.14 | 5.70 | 4.60 | 1.10 | 2.14 | 2.45464E-08 | 3.44314E-06 |
| 8651 | *SOCS1* | ENSG00000185338.7 | 3.99 | 2.89 | 1.10 | 2.14 | 1.41141E-05 | 0.00027947 |
| 55450 | *CAMK2N1* | ENSG00000162545.6 | 8.19 | 7.09 | 1.10 | 2.14 | 3.04517E-09 | 9.82477E-07 |
| 9223 | *MAGI1* | ENSG00000151276.24 | 6.95 | 5.85 | 1.09 | 2.13 | 1.15545E-06 | 4.8851E-05 |
| 3955 | *LFNG* | ENSG00000106003.14 | 2.22 | 1.13 | 1.09 | 2.13 | 0.001077479 | 0.006655093 |
| 56246 | *MRAP* | ENSG00000170262.13 | 3.73 | 2.65 | 1.08 | 2.12 | 8.37024E-05 | 0.00103059 |
| 84957 | *RELT* | ENSG00000054967.13 | 3.97 | 2.88 | 1.08 | 2.12 | 8.49362E-06 | 0.00018886 |
| 314 | *AOC2* | ENSG00000131480.9 | 2.16 | 1.08 | 1.08 | 2.12 | 0.00179565 | 0.009814604 |
| 65009 | *NDRG4* | ENSG00000103034.15 | 5.62 | 4.54 | 1.08 | 2.12 | 3.6743E-06 | 0.000104843 |
| 93010 | *B3GNT7* | ENSG00000156966.7 | 4.13 | 3.06 | 1.08 | 2.11 | 1.21737E-05 | 0.000248254 |
| 441478 | *NRARP* | ENSG00000198435.4 | 2.06 | 0.98 | 1.08 | 2.11 | 0.000510407 | 0.003851532 |
| 132864 | *CPEB2* | ENSG00000137449.17 | 4.28 | 3.21 | 1.07 | 2.10 | 0.000171594 | 0.001715826 |
| 5187 | *PER1* | ENSG00000179094.16 | 7.47 | 6.41 | 1.07 | 2.09 | 1.53232E-05 | 0.000293724 |
| 136 | *ADORA2B* | ENSG00000170425.4 | 1.95 | 0.88 | 1.07 | 2.09 | 0.00098066 | 0.00618737 |
| 579 | *NKX3-2* | ENSG00000109705.8 | 1.97 | 0.91 | 1.06 | 2.09 | 0.001220466 | 0.007335991 |
| 25780 | *RASGRP3* | ENSG00000152689.18 | 3.81 | 2.75 | 1.06 | 2.09 | 0.000175323 | 0.001745506 |
| 56254 | *RNF20* | ENSG00000155827.12 | 8.88 | 7.82 | 1.06 | 2.08 | 4.04869E-07 | 2.4156E-05 |
| 7078 | *TIMP3* | ENSG00000100234.12 | 11.16 | 10.11 | 1.05 | 2.07 | 1.83185E-10 | 2.00923E-07 |
| 5621 | *PRNP* | ENSG00000171867.18 | 6.69 | 5.64 | 1.05 | 2.07 | 8.87517E-08 | 8.53894E-06 |
| 491 | *ATP2B2* | ENSG00000157087.20 | 5.41 | 4.36 | 1.05 | 2.07 | 1.02303E-05 | 0.000219249 |
| 51673 | *TPPP3* | ENSG00000159713.11 | 3.88 | 2.83 | 1.05 | 2.07 | 1.17135E-05 | 0.000242493 |
| 55062 | *WIPI1* | ENSG00000070540.13 | 5.71 | 4.66 | 1.05 | 2.07 | 7.44959E-07 | 3.63023E-05 |
| 53342 | *IL17D* | ENSG00000172458.5 | 3.20 | 2.16 | 1.04 | 2.06 | 3.43504E-05 | 0.0005427 |
| 221749 | *PXDC1* | ENSG00000168994.14 | 2.02 | 0.98 | 1.04 | 2.06 | 0.000784072 | 0.005277188 |
| 84418 | *CYSTM1* | ENSG00000120306.11 | 6.41 | 5.37 | 1.04 | 2.06 | 1.00314E-08 | 1.93127E-06 |
| 390 | *RND3* | ENSG00000115963.14 | 4.38 | 3.34 | 1.04 | 2.06 | 4.18659E-05 | 0.000630256 |
| 3612 | *IMPA1* | ENSG00000133731.10 | 5.17 | 4.13 | 1.04 | 2.06 | 8.60261E-05 | 0.001048876 |
| 1525 | *CXADR* | ENSG00000154639.19 | 6.72 | 5.69 | 1.04 | 2.05 | 1.10651E-07 | 1.00854E-05 |
| 80817 | *CEP44* | ENSG00000164118.13 | 4.12 | 3.08 | 1.04 | 2.05 | 7.5416E-05 | 0.000957173 |
| 9314 | *KLF4* | ENSG00000136826.15 | 3.36 | 2.32 | 1.04 | 2.05 | 1.83179E-05 | 0.000336545 |
| 79730 | *NSUN7* | ENSG00000179299.17 | 4.31 | 3.28 | 1.04 | 2.05 | 0.000126077 | 0.001379215 |
| 81035 | *COLEC12* | ENSG00000158270.12 | 5.78 | 4.74 | 1.03 | 2.04 | 4.49001E-08 | 5.18389E-06 |
| 8611 | *PLPP1* | ENSG00000067113.17 | 6.75 | 5.72 | 1.03 | 2.04 | 4.39675E-09 | 1.24511E-06 |
| 4440 | *MSI1* | ENSG00000135097.7 | 4.60 | 3.57 | 1.03 | 2.04 | 2.74637E-06 | 8.54897E-05 |
| 25907 | *TMEM158* | ENSG00000249992.2 | 4.89 | 3.86 | 1.03 | 2.04 | 2.83134E-05 | 0.000468611 |
| 130888 | *FBXO36* | ENSG00000153832.12 | 1.81 | 0.78 | 1.02 | 2.03 | 0.000431757 | 0.003403492 |
| 100506710 | *EBLN3P* | ENSG00000281649.3 | 8.70 | 7.69 | 1.02 | 2.03 | 1.34471E-08 | 2.2935E-06 |
| 27123 | *DKK2* | ENSG00000155011.9 | 4.11 | 3.10 | 1.02 | 2.03 | 0.000278522 | 0.002477972 |
| 1039 | *CDR2* | ENSG00000140743.8 | 6.56 | 5.54 | 1.02 | 2.02 | 1.30847E-07 | 1.12222E-05 |
| 5292 | *PIM1* | ENSG00000137193.14 | 6.27 | 5.25 | 1.02 | 2.02 | 1.29562E-06 | 5.32766E-05 |
| 55816 | *DOK5* | ENSG00000101134.12 | 4.00 | 2.99 | 1.01 | 2.02 | 0.000324696 | 0.002754868 |
| 9745 | *ZNF536* | ENSG00000198597.9 | 5.25 | 4.24 | 1.01 | 2.01 | 1.04106E-06 | 4.62288E-05 |
| 91608 | *RASL10B* | ENSG00000270885.2 | 6.74 | 5.73 | 1.01 | 2.01 | 9.77887E-09 | 1.93127E-06 |
| 9367 | *RAB9A* | ENSG00000123595.8 | 5.67 | 4.66 | 1.01 | 2.01 | 1.09533E-05 | 0.000230572 |
| 55214 | *P3H2* | ENSG00000090530.10 | 8.29 | 7.28 | 1.01 | 2.01 | 2.13057E-06 | 7.23477E-05 |
| 57620 | *STIM2* | ENSG00000109689.19 | 6.40 | 5.39 | 1.00 | 2.00 | 8.68353E-06 | 0.000192797 |
| 195827 | *PRXL2C* | ENSG00000158122.12 | 4.77 | 3.76 | 1.00 | 2.00 | 3.98507E-06 | 0.000111798 |
| 101928721 | *LOC10192872* | ENSG00000247810.8 | 1.88 | 0.87 | 1.00 | 2.00 | 0.001225314 | 0.007359239 |
| 51026 | *GOLT1B* | ENSG00000111711.10 | 8.41 | 7.42 | 0.99 | 1.99 | 7.69967E-09 | 1.69948E-06 |
| 9871 | *SEC24D* | ENSG00000150961.15 | 6.67 | 5.67 | 0.99 | 1.99 | 1.10873E-07 | 1.00854E-05 |
| 25861 | *WHRN* | ENSG00000095397.16 | 8.85 | 7.86 | 0.99 | 1.99 | 5.64412E-08 | 6.12322E-06 |
| 79850 | *TLCD3A* | ENSG00000167695.15 | 5.55 | 4.56 | 0.99 | 1.98 | 2.17444E-07 | 1.58428E-05 |
| 5596 | *MAPK4* | ENSG00000141639.12 | 6.59 | 5.61 | 0.99 | 1.98 | 1.16638E-08 | 2.1423E-06 |
| 7421 | *VDR* | ENSG00000111424.12 | 7.71 | 6.72 | 0.99 | 1.98 | 1.34157E-08 | 2.2935E-06 |
| 6386 | *SDCBP* | ENSG00000137575.12 | 8.98 | 7.99 | 0.99 | 1.98 | 1.7094E-09 | 6.93414E-07 |
| 114789 | *SLC25A25* | ENSG00000148339.13 | 6.49 | 5.51 | 0.98 | 1.98 | 5.01029E-07 | 2.85793E-05 |
| 5901 | *RAN* | ENSG00000132341.12 | 9.88 | 8.90 | 0.98 | 1.98 | 5.0634E-09 | 1.35708E-06 |
| 127733 | *UBXN10* | ENSG00000162543.6 | 4.21 | 3.23 | 0.98 | 1.97 | 8.32008E-06 | 0.000186717 |
| 54805 | *CNNM2* | ENSG00000148842.18 | 7.53 | 6.55 | 0.98 | 1.97 | 2.04569E-07 | 1.50508E-05 |
| 23086 | *EXPH5* | ENSG00000110723.12 | 5.88 | 4.90 | 0.98 | 1.97 | 6.77621E-07 | 3.45161E-05 |
| 79974 | *CPED1* | ENSG00000106034.18 | 6.65 | 5.67 | 0.98 | 1.97 | 1.30083E-06 | 5.33449E-05 |
| 10950 | *BTG3* | ENSG00000154640.15 | 6.61 | 5.63 | 0.97 | 1.97 | 2.16386E-08 | 3.14766E-06 |
| 51030 | *TVP23B* | ENSG00000171928.14 | 6.82 | 5.84 | 0.97 | 1.96 | 2.6418E-08 | 3.63768E-06 |
| 9709 | *HERPUD1* | ENSG00000051108.15 | 8.76 | 7.79 | 0.97 | 1.96 | 4.06615E-09 | 1.17363E-06 |
| 302 | *ANXA2* | ENSG00000182718.18 | 7.02 | 6.04 | 0.97 | 1.96 | 2.79489E-07 | 1.91546E-05 |
| 5169 | *ENPP3* | ENSG00000154269.15 | 3.12 | 2.15 | 0.97 | 1.96 | 8.40661E-05 | 0.001034219 |
| 1803 | *DPP4* | ENSG00000197635.11 | 5.83 | 4.86 | 0.97 | 1.96 | 3.75902E-06 | 0.000106361 |
| 142689 | *ASB12* | ENSG00000198881.10 | 0.65 | -0.31 | 0.97 | 1.95 | 0.001422606 | 0.008253617 |
| 7803 | *PTP4A1* | ENSG00000112245.13 | 8.52 | 7.55 | 0.96 | 1.95 | 2.87236E-07 | 1.93958E-05 |
| 25984 | *KRT23* | ENSG00000108244.17 | 4.09 | 3.13 | 0.96 | 1.95 | 6.12052E-05 | 0.000826849 |
| 400073 | *C12orf76* | ENSG00000174456.16 | 4.65 | 3.69 | 0.96 | 1.95 | 4.43385E-05 | 0.000657586 |
| 109729137 | *MEF2C-AS2* | ENSG00000245864.4 | 4.00 | 3.05 | 0.95 | 1.94 | 0.000304022 | 0.00264219 |
| 7280 | *TUBB2A* | ENSG00000137267.7 | 5.77 | 4.82 | 0.95 | 1.93 | 5.18151E-08 | 5.87475E-06 |
| 57030 | *SLC17A7* | ENSG00000104888.10 | 2.55 | 1.60 | 0.95 | 1.93 | 0.001158767 | 0.007035571 |
| 84740 | *AFAP1-AS1* | ENSG00000272620.2 | 2.74 | 1.79 | 0.95 | 1.93 | 0.00027404 | 0.002448258 |
| 58516 | *SINHCAF* | ENSG00000139146.14 | 8.72 | 7.78 | 0.95 | 1.93 | 5.27271E-09 | 1.38839E-06 |

| | | | | | | | | |
|---|---|---|---|---|---|---|---|---|
| 727910 | *TLCD2* | ENSG00000185561.10 | 3.66 | 2.72 | 0.95 | 1.93 | 0.000526316 | 0.003945794 |
| 8821 | *INPP4B* | ENSG00000109452.13 | 5.94 | 4.99 | 0.95 | 1.93 | 2.05886E-05 | 0.000368313 |
| 1306 | *COL15A1* | ENSG00000204291.11 | 9.59 | 8.64 | 0.94 | 1.92 | 3.73515E-10 | 2.97005E-07 |
| 57205 | *ATP10D* | ENSG00000145246.14 | 7.41 | 6.47 | 0.94 | 1.92 | 6.09712E-07 | 3.23363E-05 |
| 117154 | *DACH2* | ENSG00000126733.22 | 6.25 | 5.31 | 0.94 | 1.92 | 3.17406E-08 | 4.21588E-06 |
| 1839 | *HBEGF* | ENSG00000113070.8 | 2.05 | 1.11 | 0.94 | 1.92 | 0.000285602 | 0.002521533 |
| 9499 | *MYOT* | ENSG00000120729.10 | 10.13 | 9.19 | 0.94 | 1.91 | 1.08366E-09 | 5.41117E-07 |
| 167127 | *UGT3A2* | ENSG00000168671.10 | 2.39 | 1.46 | 0.94 | 1.91 | 0.000683256 | 0.004760906 |
| 5986 | *RFNG* | ENSG00000169733.12 | 4.33 | 3.40 | 0.93 | 1.91 | 8.31299E-05 | 0.001026067 |
| 79026 | *AHNAK* | ENSG00000124942.14 | 5.47 | 4.54 | 0.93 | 1.90 | 5.82605E-05 | 0.00079784 |
| 57559 | *STAMBPL1* | ENSG00000138134.12 | 3.64 | 2.72 | 0.93 | 1.90 | 2.74056E-05 | 0.00045892 |
| 57795 | *BRINP2* | ENSG00000198797.7 | 9.15 | 8.22 | 0.93 | 1.90 | 1.20353E-09 | 5.64494E-07 |
| 153222 | *CREBRF* | ENSG00000164463.12 | 5.37 | 4.45 | 0.93 | 1.90 | 6.85752E-05 | 0.000900477 |
| 154141 | *MBOAT1* | ENSG00000172197.11 | 5.25 | 4.32 | 0.93 | 1.90 | 7.54557E-07 | 3.66509E-05 |
| 58515 | *SELENOK* | ENSG00000113811.12 | 6.71 | 5.78 | 0.92 | 1.90 | 3.57037E-08 | 4.54133E-06 |
| 4300 | *MLLT3* | ENSG00000171843.17 | 6.12 | 5.20 | 0.92 | 1.89 | 6.03204E-06 | 0.000148662 |
| 5864 | *RAB3A* | ENSG00000105649.10 | 3.59 | 2.68 | 0.91 | 1.88 | 1.22258E-05 | 0.000248979 |
| 160760 | *PPTC7* | ENSG00000196850.6 | 7.58 | 6.67 | 0.91 | 1.88 | 7.64022E-09 | 1.69948E-06 |
| 22927 | *HABP4* | ENSG00000130956.14 | 5.64 | 4.73 | 0.91 | 1.88 | 1.2579E-07 | 1.08606E-05 |
| 6812 | *STXBP1* | ENSG00000136854.24 | 6.74 | 5.84 | 0.91 | 1.88 | 1.25908E-07 | 1.08606E-05 |
| 7704 | *ZBTB16* | ENSG00000109906.15 | 3.18 | 2.28 | 0.91 | 1.87 | 0.000865859 | 0.005679932 |
| 157378 | *TMEM65* | ENSG00000164983.8 | 5.21 | 4.30 | 0.91 | 1.87 | 2.29152E-07 | 1.65124E-05 |
| 678 | *ZFP36L2* | ENSG00000152518.8 | 8.80 | 7.89 | 0.91 | 1.87 | 2.68778E-09 | 8.96465E-07 |
| 91404 | *SESTD1* | ENSG00000187231.14 | 7.63 | 6.73 | 0.90 | 1.87 | 7.91205E-07 | 3.77105E-05 |
| 4026 | *LPP* | ENSG00000145012.14 | 4.76 | 3.85 | 0.90 | 1.87 | 0.000515003 | 0.003878411 |
| 25819 | *NOCT* | ENSG00000151014.6 | 3.57 | 2.67 | 0.90 | 1.87 | 0.001294966 | 0.007675962 |
| 667 | *DST* | ENSG00000151914.22 | 8.45 | 7.55 | 0.90 | 1.87 | 8.01361E-06 | 0.000182513 |
| 254251 | *LCORL* | ENSG00000178177.16 | 6.84 | 5.94 | 0.90 | 1.87 | 1.81338E-07 | 1.41021E-05 |
| 8556 | *CDC14A* | ENSG00000079335.20 | 2.87 | 1.97 | 0.90 | 1.87 | 0.000149309 | 0.001559484 |
| 9456 | *HOMER1* | ENSG00000152413.15 | 7.33 | 6.43 | 0.90 | 1.86 | 0.00028038 | 0.002491552 |
| 6627 | *SNRPA1* | ENSG00000131876.18 | 6.01 | 5.11 | 0.90 | 1.86 | 4.38618E-06 | 0.000117979 |
| 30844 | *EHD4* | ENSG00000103966.11 | 5.83 | 4.94 | 0.90 | 1.86 | 1.26268E-06 | 5.23523E-05 |
| 3033 | *HADH* | ENSG00000138796.18 | 7.27 | 6.37 | 0.90 | 1.86 | 5.89523E-09 | 1.48304E-06 |
| 8744 | *TNFSF9* | ENSG00000125657.5 | 2.13 | 1.24 | 0.89 | 1.86 | 0.001735376 | 0.009576773 |
| 6322 | *SCML1* | ENSG00000047634.15 | 6.05 | 5.16 | 0.89 | 1.85 | 7.95776E-08 | 8.07014E-06 |
| 146760 | *RTN4RL1* | ENSG00000185924.7 | 4.01 | 3.12 | 0.89 | 1.85 | 2.22797E-05 | 0.000390194 |
| 58527 | *ABRACL* | ENSG00000146386.8 | 5.96 | 5.06 | 0.89 | 1.85 | 5.55658E-07 | 3.07744E-05 |
| 6907 | *TBL1X* | ENSG00000101849.18 | 6.40 | 5.51 | 0.89 | 1.85 | 1.97494E-06 | 6.9582E-05 |
| 4502 | *MT2A* | ENSG00000125148.7 | 8.25 | 7.36 | 0.89 | 1.85 | 4.73732E-06 | 0.000124741 |
| 54541 | *DDIT4* | ENSG00000168209.6 | 4.78 | 3.89 | 0.89 | 1.85 | 0.000857505 | 0.0056523 |
| 84343 | *HPS3* | ENSG00000163755.9 | 7.30 | 6.42 | 0.89 | 1.85 | 2.18107E-08 | 3.14766E-06 |
| 4189 | *DNAJB9* | ENSG00000128590.5 | 6.71 | 5.83 | 0.89 | 1.85 | 6.71198E-07 | 3.45161E-05 |
| 4881 | *NPR1* | ENSG00000169418.10 | 3.08 | 2.20 | 0.88 | 1.85 | 0.000907137 | 0.005863573 |
| 10472 | *ZBTB18* | ENSG00000179456.10 | 4.43 | 3.54 | 0.88 | 1.84 | 3.92586E-05 | 0.000599423 |
| 253512 | *SLC25A30* | ENSG00000174032.17 | 5.67 | 4.79 | 0.88 | 1.84 | 0.000113435 | 0.001282999 |
| 817 | *CAMK2D* | ENSG00000145349.18 | 5.08 | 4.19 | 0.88 | 1.84 | 8.04989E-07 | 3.82344E-05 |
| 488 | *ATP2A2* | ENSG00000174437.18 | 9.46 | 8.58 | 0.88 | 1.84 | 7.11136E-09 | 1.63338E-06 |
| 29948 | *OSGIN1* | ENSG00000140961.14 | 6.01 | 5.13 | 0.88 | 1.84 | 6.89064E-08 | 7.03548E-06 |
| 10150 | *MBNL2* | ENSG00000139793.20 | 8.53 | 7.65 | 0.87 | 1.83 | 2.14862E-08 | 3.14766E-06 |
| 1164 | *CKS2* | ENSG00000123975.5 | 8.46 | 7.58 | 0.87 | 1.83 | 5.70416E-08 | 6.12322E-06 |
| 55353 | *LAPTM4B* | ENSG00000104341.17 | 8.21 | 7.34 | 0.87 | 1.83 | 5.93902E-07 | 3.18353E-05 |
| 87 | *ACTN1* | ENSG00000072110.16 | 6.73 | 5.86 | 0.87 | 1.83 | 2.57053E-07 | 1.81132E-05 |
| 283624 | *LINC00641* | ENSG00000258441.1 | 3.43 | 2.55 | 0.87 | 1.83 | 0.001688415 | 0.009393709 |
| 55132 | *LARP1B* | ENSG00000138709.19 | 4.52 | 3.66 | 0.86 | 1.82 | 2.4007E-06 | 7.79915E-05 |
| 3202 | *HOXA5* | ENSG00000106004.5 | 5.71 | 4.85 | 0.86 | 1.81 | 3.46755E-06 | 0.000101394 |
| 84295 | *PHF6* | ENSG00000156531.18 | 7.41 | 6.55 | 0.86 | 1.81 | 3.21716E-08 | 4.23565E-06 |
| 3651 | *PDX1* | ENSG00000139515.6 | 1.96 | 1.11 | 0.86 | 1.81 | 0.000475556 | 0.003652825 |
| 55151 | *TMEM38B* | ENSG00000095209.12 | 6.92 | 6.06 | 0.86 | 1.81 | 1.77388E-07 | 1.38667E-05 |
| 728215 | *NALF1* | ENSG00000204442.4 | 6.73 | 5.87 | 0.86 | 1.81 | 1.1997E-07 | 1.06546E-05 |
| 23710 | *GABARAPL1* | ENSG00000139112.11 | 8.39 | 7.53 | 0.86 | 1.81 | 1.22074E-08 | 2.20747E-06 |
| 23645 | *PPP1R15A* | ENSG00000087074.8 | 5.93 | 5.07 | 0.86 | 1.81 | 3.02379E-07 | 1.96468E-05 |
| 89927 | *BMERB1* | ENSG00000166780.11 | 4.99 | 4.14 | 0.85 | 1.80 | 4.59071E-05 | 0.000672215 |
| 348487 | *FAM131C* | ENSG00000185519.9 | 2.96 | 2.11 | 0.85 | 1.80 | 0.000188442 | 0.001841359 |
| 221914 | *GPC2* | ENSG00000213420.8 | 3.91 | 3.06 | 0.85 | 1.80 | 0.000220909 | 0.002074861 |
| 10014 | *HDAC5* | ENSG00000108840.16 | 7.05 | 6.20 | 0.85 | 1.80 | 2.04363E-07 | 1.50508E-05 |
| 6781 | *STC1* | ENSG00000159167.12 | 4.38 | 3.52 | 0.85 | 1.80 | 0.000726688 | 0.004987134 |
| 8462 | *KLF11* | ENSG00000172059.11 | 5.83 | 4.98 | 0.85 | 1.80 | 2.00288E-07 | 1.49558E-05 |
| 11138 | *TBC1D8* | ENSG00000204634.13 | 3.95 | 3.10 | 0.84 | 1.79 | 0.000420438 | 0.003340575 |
| 5252 | *PHF1* | ENSG00000112511.18 | 6.20 | 5.35 | 0.84 | 1.79 | 2.54919E-06 | 8.14059E-05 |
| 5166 | *PDK4* | ENSG00000004799.8 | 11.27 | 10.43 | 0.84 | 1.79 | 8.42591E-10 | 4.67242E-07 |
| 7424 | *VEGFC* | ENSG00000150630.4 | 2.90 | 2.06 | 0.84 | 1.79 | 0.000310959 | 0.00267438 |
| 5831 | *PYCR1* | ENSG00000183010.17 | 5.76 | 4.93 | 0.84 | 1.79 | 8.09863E-06 | 0.00018417 |
| 2673 | *GFPT1* | ENSG00000198380.13 | 8.18 | 7.35 | 0.83 | 1.78 | 2.02187E-07 | 1.50229E-05 |
| 341640 | *FREM2* | ENSG00000150893.11 | 4.58 | 3.74 | 0.83 | 1.78 | 0.000738029 | 0.005046506 |
| 284129 | *SLC26A11* | ENSG00000181045.15 | 5.95 | 5.12 | 0.83 | 1.78 | 1.45974E-05 | 0.000284166 |
| 1903 | *S1PR3* | ENSG00000213694.6 | 9.50 | 8.68 | 0.82 | 1.77 | 4.5258E-09 | 1.25792E-06 |
| 23162 | *MAPK8IP3* | ENSG00000138834.15 | 5.40 | 4.58 | 0.82 | 1.77 | 0.000107355 | 0.001230449 |
| 23389 | *MED13L* | ENSG00000123066.9 | 7.57 | 6.75 | 0.82 | 1.77 | 1.67346E-06 | 6.18645E-05 |
| 81615 | *TMEM163* | ENSG00000152128.13 | 3.46 | 2.64 | 0.82 | 1.77 | 5.08544E-05 | 0.000721473 |
| 492311 | *IGIP* | ENSG00000182700.5 | 4.10 | 3.28 | 0.82 | 1.76 | 0.000710203 | 0.004909921 |
| 65991 | *FUNDC2* | ENSG00000165775.18 | 7.57 | 6.75 | 0.82 | 1.76 | 4.55112E-07 | 2.66225E-05 |
| 54583 | *EGLN1* | ENSG00000135766.9 | 7.99 | 7.17 | 0.82 | 1.76 | 5.81303E-07 | 3.17519E-05 |
| 26277 | *TINF2* | ENSG00000092330.18 | 6.52 | 5.70 | 0.82 | 1.76 | 1.07291E-06 | 4.66761E-05 |
| 7494 | *XBP1* | ENSG00000100219.17 | 7.45 | 6.63 | 0.82 | 1.76 | 5.28346E-08 | 5.87475E-06 |
| 158399 | *ZNF483* | ENSG00000173258.13 | 4.95 | 4.14 | 0.81 | 1.76 | 0.000180979 | 0.001788224 |
| 54814 | *QPCTL* | ENSG00000011478.13 | 4.13 | 3.31 | 0.81 | 1.76 | 0.000263541 | 0.002387141 |
| 647135 | *SRGAP2B* | ENSG00000196369.12 | 7.35 | 6.54 | 0.81 | 1.75 | 4.43172E-07 | 2.61873E-05 |
| 6809 | *STX3* | ENSG00000166900.17 | 6.30 | 5.50 | 0.81 | 1.75 | 5.39231E-07 | 3.00867E-05 |
| 114793 | *FMNL2* | ENSG00000157827.20 | 6.03 | 5.23 | 0.81 | 1.75 | 3.90528E-07 | 2.35399E-05 |

| | | | | | | | | |
|---|---|---|---|---|---|---|---|---|
| 84803 | *GPAT3* | ENSG00000138678.11 | 3.59 | 2.79 | 0.81 | 1.75 | 7.8517E-05 | 0.000984513 |
| 285830 | *HLA-F-AS1* | ENSG00000214922.9 | 3.32 | 2.52 | 0.81 | 1.75 | 0.000917013 | 0.005895808 |
| 4602 | *MYB* | ENSG00000118513.20 | 4.94 | 4.14 | 0.81 | 1.75 | 4.52589E-06 | 0.000120547 |
| 60682 | *SMAP1* | ENSG00000112305.15 | 7.13 | 6.33 | 0.80 | 1.74 | 1.7556E-08 | 2.8333E-06 |
| 1163 | *CKS1B* | ENSG00000173207.13 | 7.85 | 7.05 | 0.80 | 1.74 | 3.86827E-08 | 4.76126E-06 |
| 23380 | *SRGAP2* | ENSG00000266028.8 | 7.07 | 6.28 | 0.80 | 1.74 | 5.31446E-08 | 5.87475E-06 |
| 10627 | *MYL12A* | ENSG00000101608.13 | 8.18 | 7.38 | 0.80 | 1.74 | 3.05348E-08 | 4.1288E-06 |
| 8459 | *TPST2* | ENSG00000128294.16 | 5.84 | 5.04 | 0.80 | 1.74 | 5.04599E-07 | 2.85793E-05 |
| 10715 | *CERS1* | ENSG00000223802.9 | 4.61 | 3.80 | 0.80 | 1.74 | 0.000528839 | 0.003957182 |
| 26524 | *LATS2* | ENSG00000150457.9 | 5.84 | 5.04 | 0.79 | 1.73 | 3.93716E-07 | 2.36372E-05 |
| 83999 | *KREMEN1* | ENSG00000183762.13 | 7.45 | 6.66 | 0.79 | 1.73 | 1.32212E-06 | 5.40699E-05 |
| 9754 | *STARD8* | ENSG00000130052.14 | 6.17 | 5.38 | 0.79 | 1.73 | 1.03775E-06 | 4.62182E-05 |
| 55254 | *TMEM39A* | ENSG00000176142.13 | 6.79 | 6.00 | 0.79 | 1.73 | 4.41975E-06 | 0.000118669 |
| 54602 | *NDFIP2* | ENSG00000102471.15 | 6.60 | 5.81 | 0.79 | 1.73 | 6.17039E-07 | 3.26097E-05 |
| 5966 | *REL* | ENSG00000162924.15 | 4.87 | 4.08 | 0.79 | 1.73 | 0.000370451 | 0.003041632 |
| 440145 | *MZT1* | ENSG00000204899.6 | 5.89 | 5.11 | 0.79 | 1.73 | 4.84324E-06 | 0.000127307 |
| 11120 | *BTN2A1* | ENSG00000112763.17 | 4.83 | 4.04 | 0.79 | 1.73 | 0.000632521 | 0.004501425 |
| 2697 | *GJA1* | ENSG00000152661.9 | 6.11 | 5.33 | 0.78 | 1.72 | 8.14867E-06 | 0.000184748 |
| 7791 | *ZYX* | ENSG00000159840.16 | 5.54 | 4.76 | 0.78 | 1.72 | 2.71126E-06 | 8.49548E-05 |
| 10327 | *AKR1A1* | ENSG00000117448.14 | 6.15 | 5.37 | 0.78 | 1.72 | 1.13988E-07 | 1.03063E-05 |
| 6016 | *RIT1* | ENSG00000143622.11 | 7.15 | 6.37 | 0.78 | 1.72 | 6.41716E-07 | 3.34464E-05 |
| 81575 | *APOLD1* | ENSG00000178878.13 | 5.82 | 5.04 | 0.78 | 1.72 | 3.43232E-06 | 0.000101011 |
| 23016 | *EXOSC7* | ENSG00000075914.13 | 4.97 | 4.19 | 0.78 | 1.71 | 8.47431E-06 | 0.000188845 |
| 4494 | *MT1F* | ENSG00000198417.7 | 5.50 | 4.72 | 0.78 | 1.71 | 2.75706E-06 | 8.54897E-05 |
| 6659 | *SOX4* | ENSG00000124766.7 | 6.93 | 6.16 | 0.78 | 1.71 | 8.08889E-08 | 8.12287E-06 |
| 8575 | *PRKRA* | ENSG00000180228.14 | 6.88 | 6.10 | 0.78 | 1.71 | 2.0318E-06 | 7.07547E-05 |
| 604 | *BCL6* | ENSG00000113916.18 | 5.98 | 5.21 | 0.78 | 1.71 | 1.51049E-06 | 5.82801E-05 |
| 1364 | *CLDN4* | ENSG00000189143.10 | 3.30 | 2.52 | 0.78 | 1.71 | 0.001447744 | 0.008351959 |
| 6478 | *SIAH2* | ENSG00000181788.4 | 10.10 | 9.32 | 0.77 | 1.71 | 4.48913E-08 | 5.18389E-06 |
| 5805 | *PTS* | ENSG00000150787.8 | 5.94 | 5.17 | 0.77 | 1.71 | 5.60687E-07 | 3.09388E-05 |
| 58191 | *CXCL16* | ENSG00000161921.17 | 4.34 | 3.56 | 0.77 | 1.71 | 9.77403E-05 | 0.00115445 |
| 26354 | *GNL3* | ENSG00000163938.17 | 7.06 | 6.29 | 0.77 | 1.71 | 5.87315E-07 | 3.17519E-05 |
| 160851 | *DGKH* | ENSG00000102780.17 | 5.26 | 4.49 | 0.77 | 1.71 | 0.000313303 | 0.002685246 |
| 160518 | *DENND5B* | ENSG00000170456.16 | 7.66 | 6.89 | 0.77 | 1.71 | 4.85073E-07 | 2.78945E-05 |
| 51302 | *CYP39A1* | ENSG00000146233.8 | 4.73 | 3.96 | 0.77 | 1.70 | 6.04257E-06 | 0.000148677 |
| 5238 | *PGM3* | ENSG00000013375.16 | 6.81 | 6.04 | 0.77 | 1.70 | 4.43011E-06 | 0.000118735 |
| 6696 | *SPP1* | ENSG00000118785.15 | 7.40 | 6.63 | 0.77 | 1.70 | 5.85998E-07 | 3.17519E-05 |
| 9572 | *NR1D1* | ENSG00000126368.6 | 4.75 | 3.99 | 0.77 | 1.70 | 3.50445E-05 | 0.000550768 |
| 134111 | *UBE2QL1* | ENSG00000215218.4 | 3.60 | 2.84 | 0.77 | 1.70 | 6.7131E-05 | 0.000886164 |
| 3706 | *ITPKA* | ENSG00000137825.11 | 4.11 | 3.34 | 0.77 | 1.70 | 3.7309E-06 | 0.000106055 |
| 5062 | *PAK2* | ENSG00000180370.10 | 8.57 | 7.80 | 0.77 | 1.70 | 3.08795E-07 | 1.99772E-05 |
| 143686 | *SESN3* | ENSG00000149212.12 | 6.49 | 5.72 | 0.76 | 1.70 | 1.39261E-07 | 1.15766E-05 |
| 2054 | *STX2* | ENSG00000111450.14 | 6.51 | 5.74 | 0.76 | 1.70 | 3.29845E-07 | 2.0801E-05 |
| 6790 | *AURKA* | ENSG00000087586.18 | 6.90 | 6.14 | 0.76 | 1.70 | 3.48477E-07 | 2.17025E-05 |
| 3488 | *IGFBP5* | ENSG00000115461.5 | 3.48 | 2.72 | 0.76 | 1.69 | 0.001708898 | 0.009483932 |
| 4008 | *LMO7* | ENSG00000136153.20 | 6.34 | 5.58 | 0.76 | 1.69 | 4.88094E-06 | 0.000127405 |
| 25897 | *RNF19A* | ENSG00000034677.13 | 6.84 | 6.08 | 0.76 | 1.69 | 3.21323E-07 | 2.04821E-05 |
| 11014 | *KDELR2* | ENSG00000136240.10 | 8.48 | 7.72 | 0.76 | 1.69 | 4.4741E-08 | 5.18389E-06 |
| 8218 | *CLTCL1* | ENSG00000070371.16 | 5.49 | 4.73 | 0.76 | 1.69 | 1.04353E-05 | 0.000221532 |
| 23138 | *N4BP3* | ENSG00000145911.6 | 2.49 | 1.74 | 0.76 | 1.69 | 0.001778601 | 0.009732056 |
| 2195 | *FAT1* | ENSG00000083857.15 | 8.62 | 7.87 | 0.75 | 1.69 | 8.58056E-05 | 0.001047477 |
| 121457 | *IKBIP* | ENSG00000166130.15 | 4.88 | 4.12 | 0.75 | 1.69 | 1.40539E-06 | 5.59509E-05 |
| 11237 | *RNF24* | ENSG00000101236.17 | 4.93 | 4.18 | 0.75 | 1.69 | 0.000209301 | 0.001999617 |
| 57605 | *PITPNM2* | ENSG00000090975.14 | 5.86 | 5.11 | 0.75 | 1.69 | 0.001731315 | 0.00957199 |
| 10493 | *VAT1* | ENSG00000108828.16 | 9.28 | 8.53 | 0.75 | 1.68 | 5.20759E-08 | 5.87475E-06 |
| 26993 | *AKAP8L* | ENSG00000011243.19 | 6.65 | 5.90 | 0.75 | 1.68 | 4.14951E-07 | 2.46166E-05 |
| 151648 | *SGO1* | ENSG00000129810.15 | 5.62 | 4.88 | 0.75 | 1.68 | 3.55853E-05 | 0.000554805 |
| 11142 | *PKIG* | ENSG00000168734.14 | 6.04 | 5.28 | 0.75 | 1.68 | 4.99598E-06 | 0.000129507 |
| 170685 | *NUDT10* | ENSG00000122824.11 | 4.86 | 4.10 | 0.75 | 1.68 | 8.19397E-07 | 3.8674E-05 |
| 100527963 | *PMF1-BGLAP* | ENSG00000260238.6 | 3.04 | 2.30 | 0.75 | 1.68 | 0.000764197 | 0.005168922 |
| 143888 | *POGLUT3* | ENSG00000178202.13 | 6.88 | 6.13 | 0.75 | 1.68 | 5.90028E-08 | 6.23643E-06 |
| 11018 | *TMED1* | ENSG00000099203.7 | 5.64 | 4.89 | 0.75 | 1.68 | 6.29953E-06 | 0.00015349 |
| 775 | *CACNA1C* | ENSG00000151067.23 | 6.88 | 6.13 | 0.75 | 1.68 | 0.000575345 | 0.004214421 |
| 9232 | *PTTG1* | ENSG00000164611.13 | 7.73 | 6.98 | 0.75 | 1.68 | 1.82607E-07 | 1.41276E-05 |
| 7769 | *ZNF226* | ENSG00000167380.17 | 4.84 | 4.09 | 0.75 | 1.68 | 5.92078E-06 | 0.000146884 |
| 84662 | *GLIS2* | ENSG00000126603.9 | 4.50 | 3.76 | 0.75 | 1.68 | 4.95644E-05 | 0.000709162 |
| 100507436 | *MICA* | ENSG00000204520.14 | 5.27 | 4.53 | 0.74 | 1.67 | 2.77169E-06 | 8.55974E-05 |
| 135154 | *SDHAF4* | ENSG00000154079.6 | 3.30 | 2.57 | 0.74 | 1.67 | 0.000925816 | 0.005938276 |
| 3133 | *HLA-E* | ENSG00000204592.9 | 6.78 | 6.04 | 0.74 | 1.67 | 6.41786E-07 | 3.34464E-05 |
| 57026 | *PDXP* | ENSG00000241360.2 | 6.55 | 5.81 | 0.74 | 1.67 | 2.25623E-06 | 7.55887E-05 |
| 8324 | *FZD7* | ENSG00000155760.3 | 7.47 | 6.73 | 0.74 | 1.67 | 1.15956E-07 | 1.04214E-05 |
| 4092 | *SMAD7* | ENSG00000101665.10 | 3.70 | 2.96 | 0.74 | 1.67 | 0.000523477 | 0.003928996 |
| 2657 | *GDF1* | ENSG00000130283.9 | 3.41 | 2.67 | 0.74 | 1.67 | 0.000919787 | 0.005907182 |
| 757 | *TMEM50B* | ENSG00000142188.17 | 6.01 | 5.27 | 0.74 | 1.67 | 3.01341E-07 | 1.96468E-05 |
| 9415 | *FADS2* | ENSG00000134824.14 | 4.33 | 3.59 | 0.74 | 1.67 | 7.75932E-05 | 0.000977402 |
| 81539 | *SLC38A1* | ENSG00000111371.16 | 9.25 | 8.52 | 0.73 | 1.66 | 1.95364E-08 | 2.93222E-06 |
| 5971 | *RELB* | ENSG00000104856.14 | 2.74 | 2.01 | 0.73 | 1.66 | 0.001574814 | 0.008936252 |
| 6396 | *SEC13* | ENSG00000157020.18 | 7.36 | 6.62 | 0.73 | 1.66 | 1.49335E-07 | 1.21814E-05 |
| 2801 | *GOLGA2* | ENSG00000167110.19 | 7.17 | 6.44 | 0.73 | 1.66 | 4.47779E-06 | 0.000119586 |
| 2026 | *ENO2* | ENSG00000111674.9 | 4.80 | 4.07 | 0.73 | 1.66 | 7.13393E-05 | 0.000921456 |
| 151887 | *CCDC80* | ENSG00000091986.16 | 5.68 | 4.96 | 0.73 | 1.66 | 0.001105888 | 0.006794215 |
| 65055 | *REEP1* | ENSG00000068615.20 | 3.86 | 3.14 | 0.72 | 1.65 | 0.001735158 | 0.009576773 |
| 3159 | *HMGA1* | ENSG00000137309.20 | 3.93 | 3.20 | 0.72 | 1.65 | 8.28644E-05 | 0.001023631 |
| 81550 | *TDRD3* | ENSG00000083544.16 | 5.13 | 4.41 | 0.72 | 1.65 | 0.000273075 | 0.002443994 |
| 25930 | *PTPN23* | ENSG00000076201.16 | 6.64 | 5.92 | 0.72 | 1.65 | 3.52E-06 | 0.000101992 |
| 122525 | *C14orf28* | ENSG00000179476.8 | 2.45 | 1.73 | 0.72 | 1.65 | 0.001455685 | 0.008390314 |
| 1974 | *EIF4A2* | ENSG00000156976.17 | 9.07 | 8.35 | 0.72 | 1.65 | 9.76728E-09 | 1.93127E-06 |
| 285590 | *SH3PXD2B* | ENSG00000174705.13 | 6.91 | 6.19 | 0.72 | 1.64 | 8.07547E-07 | 3.82349E-05 |
| 25843 | *MOB4* | ENSG00000115540.15 | 9.46 | 8.75 | 0.72 | 1.64 | 1.94311E-08 | 2.93222E-06 |

| | | | | | | | | |
|---|---|---|---|---|---|---|---|---|
| 91768 | *CABLES1* | ENSG00000134508.13 | 4.67 | 3.95 | 0.72 | 1.64 | 0.000560289 | 0.004126291 |
| 4000 | *LMNA* | ENSG00000160789.24 | 8.89 | 8.17 | 0.72 | 1.64 | 1.23886E-07 | 1.08606E-05 |
| 2585 | *GALK2* | ENSG00000156958.15 | 6.00 | 5.28 | 0.72 | 1.64 | 1.19474E-05 | 0.000246163 |
| 7376 | *NR1H2* | ENSG00000131408.15 | 6.86 | 6.14 | 0.71 | 1.64 | 4.22739E-06 | 0.000114843 |
| 483 | *ATP1B3* | ENSG00000069849.11 | 7.37 | 6.66 | 0.71 | 1.64 | 1.08986E-06 | 4.71403E-05 |
| 401097 | *C3orf80* | ENSG00000180044.6 | 3.83 | 3.12 | 0.71 | 1.64 | 0.000831186 | 0.005517878 |
| 6385 | *SDC4* | ENSG00000124145.6 | 6.82 | 6.11 | 0.71 | 1.63 | 6.01152E-07 | 3.19953E-05 |
| 11156 | *PTP4A3* | ENSG00000184489.13 | 2.43 | 1.73 | 0.71 | 1.63 | 0.001053312 | 0.006540821 |
| 128553 | *TSHZ2* | ENSG00000182463.16 | 5.46 | 4.75 | 0.71 | 1.63 | 0.000548221 | 0.004065341 |
| 26502 | *NARF* | ENSG00000141562.19 | 5.70 | 4.99 | 0.71 | 1.63 | 1.62358E-06 | 6.12271E-05 |
| 7922 | *SLC39A7* | ENSG00000112473.18 | 8.32 | 7.62 | 0.70 | 1.63 | 4.0241E-08 | 4.91038E-06 |
| 153241 | *CEP120* | ENSG00000168944.17 | 6.06 | 5.36 | 0.70 | 1.63 | 4.05578E-07 | 2.4156E-05 |
| 128239 | *IQGAP3* | ENSG00000183856.11 | 6.84 | 6.14 | 0.70 | 1.63 | 9.17465E-07 | 4.224E-05 |
| 28227 | *PPP2R3B* | ENSG00000167393.18 | 3.30 | 2.60 | 0.70 | 1.63 | 0.000375025 | 0.003068684 |
| 28227 | *PPP2R3B* | ENSG00000167393.18_P/ | 3.30 | 2.60 | 0.70 | 1.63 | 0.000375025 | 0.003068684 |
| 7555 | *CNBP* | ENSG00000169714.17 | 8.77 | 8.07 | 0.70 | 1.63 | 1.39608E-07 | 1.15766E-05 |
| 29803 | *REPIN1* | ENSG00000214022.12 | 7.84 | 7.14 | 0.70 | 1.62 | 1.3616E-07 | 1.15459E-05 |
| 813 | *CALU* | ENSG00000128595.17 | 10.27 | 9.57 | 0.70 | 1.62 | 1.57387E-08 | 2.56763E-06 |
| 6782 | *HSPA13* | ENSG00000155304.6 | 8.42 | 7.72 | 0.70 | 1.62 | 1.05209E-06 | 4.63072E-05 |
| 5997 | *RGS2* | ENSG00000116741.8 | 5.20 | 4.51 | 0.70 | 1.62 | 2.7808E-06 | 8.57024E-05 |
| 10209 | *EIF1* | ENSG00000173812.11 | 9.44 | 8.75 | 0.70 | 1.62 | 1.25534E-07 | 1.08606E-05 |
| 3998 | *LMAN1* | ENSG00000074695.6 | 9.40 | 8.70 | 0.70 | 1.62 | 9.42723E-09 | 1.93127E-06 |
| 23345 | *SYNE1* | ENSG00000131018.25 | 8.46 | 7.76 | 0.70 | 1.62 | 5.88118E-07 | 3.17519E-05 |
| 51076 | *CUTC* | ENSG00000119929.13 | 3.73 | 3.04 | 0.70 | 1.62 | 0.001601438 | 0.00903823 |
| 2977 | *GUCY1A2* | ENSG00000152402.11 | 4.61 | 3.91 | 0.70 | 1.62 | 0.001442908 | 0.008335877 |
| 55829 | *SELENOS* | ENSG00000131871.15 | 6.20 | 5.51 | 0.69 | 1.62 | 1.50613E-06 | 5.82618E-05 |
| 11177 | *BAZ1A* | ENSG00000198604.11 | 5.23 | 4.53 | 0.69 | 1.62 | 1.5252E-05 | 0.000293571 |
| 57182 | *ANKRD50* | ENSG00000151458.12 | 5.98 | 5.29 | 0.69 | 1.62 | 3.36158E-06 | 9.95458E-05 |
| 2803 | *GOLGA4* | ENSG00000144674.17 | 7.28 | 6.58 | 0.69 | 1.62 | 0.000118376 | 0.001326892 |
| 6839 | *SUV39H1* | ENSG00000101945.17 | 5.14 | 4.45 | 0.69 | 1.62 | 6.22089E-05 | 0.000837393 |
| 3297 | *HSF1* | ENSG00000185122.11 | 5.32 | 4.63 | 0.69 | 1.62 | 0.000386513 | 0.003129006 |
| 100129550 | *LINC02035* | ENSG00000273033.2 | 5.10 | 4.41 | 0.69 | 1.61 | 0.000111342 | 0.001263134 |
| 91750 | *LIN52* | ENSG00000205659.12 | 4.22 | 3.53 | 0.69 | 1.61 | 9.73829E-05 | 0.001152697 |
| 55075 | *UACA* | ENSG00000137831.15 | 7.75 | 7.06 | 0.69 | 1.61 | 1.90385E-06 | 6.77129E-05 |
| 54407 | *SLC38A2* | ENSG00000134294.14 | 9.27 | 8.58 | 0.69 | 1.61 | 1.94167E-07 | 1.46445E-05 |
| 23603 | *CORO1C* | ENSG00000110880.11 | 8.81 | 8.12 | 0.69 | 1.61 | 2.88102E-07 | 1.93958E-05 |
| 11248 | *NXPH3* | ENSG00000182575.8 | 5.53 | 4.84 | 0.69 | 1.61 | 4.63731E-06 | 0.000122971 |
| 4215 | *MAP3K3* | ENSG00000198909.8 | 4.32 | 3.63 | 0.69 | 1.61 | 3.04092E-05 | 0.0004961 |
| 57186 | *RALGAPA2* | ENSG00000188559.16 | 7.73 | 7.05 | 0.69 | 1.61 | 8.72331E-05 | 0.001061006 |
| 5604 | *MAP2K1* | ENSG00000169032.11 | 7.77 | 7.08 | 0.68 | 1.61 | 8.13735E-08 | 8.12287E-06 |
| 5930 | *RBBP6* | ENSG00000122257.20 | 6.66 | 5.98 | 0.68 | 1.61 | 0.000165531 | 0.001677549 |
| 56980 | *PRDM10* | ENSG00000170325.16 | 4.73 | 4.04 | 0.68 | 1.61 | 0.001821907 | 0.009932802 |
| 55905 | *RNF114* | ENSG00000124226.11 | 6.22 | 5.54 | 0.68 | 1.60 | 0.000110669 | 0.001256448 |
| 10114 | *HIPK3* | ENSG00000110422.12 | 6.30 | 5.61 | 0.68 | 1.60 | 2.61138E-06 | 8.23678E-05 |
| 9167 | *COX7A2L* | ENSG00000115944.15 | 7.20 | 6.52 | 0.68 | 1.60 | 4.72103E-06 | 0.000124531 |
| 790 | *CAD* | ENSG00000084774.14 | 7.23 | 6.55 | 0.68 | 1.60 | 6.78683E-06 | 0.000162203 |
| 80318 | *GKAP1* | ENSG00000165113.13 | 3.67 | 3.00 | 0.68 | 1.60 | 0.000617188 | 0.004425886 |
| 2992 | *GYG1* | ENSG00000163754.18 | 5.69 | 5.01 | 0.68 | 1.60 | 1.63315E-06 | 6.14336E-05 |
| 101929704 | *TTC23L-AS1* | ENSG00000272323.1 | 2.84 | 2.16 | 0.67 | 1.60 | 0.001639089 | 0.009202527 |
| 222068 | *TMED4* | ENSG00000158604.15 | 8.29 | 7.62 | 0.67 | 1.59 | 2.43858E-08 | 3.44314E-06 |
| 6429 | *SRSF4* | ENSG00000116350.18 | 7.37 | 6.70 | 0.67 | 1.59 | 1.14341E-06 | 4.8616E-05 |
| 56895 | *AGPAT4* | ENSG00000026652.15 | 5.42 | 4.75 | 0.67 | 1.59 | 6.89081E-05 | 0.000901693 |
| 284403 | *WDR62* | ENSG00000075702.19 | 6.82 | 6.15 | 0.67 | 1.59 | 3.3018E-06 | 9.81321E-05 |
| 64651 | *CSRNP1* | ENSG00000144655.15 | 4.80 | 4.13 | 0.67 | 1.59 | 2.76999E-05 | 0.000462456 |
| 80315 | *CPEB4* | ENSG00000113742.14 | 7.52 | 6.85 | 0.67 | 1.59 | 2.92054E-07 | 1.93958E-05 |
| 84679 | *SLC9A7* | ENSG00000065923.10 | 5.29 | 4.62 | 0.67 | 1.59 | 0.001299356 | 0.007684019 |
| 23204 | *ARL6IP1* | ENSG00000170540.15 | 10.07 | 9.40 | 0.67 | 1.59 | 6.39719E-08 | 6.62175E-06 |
| 5866 | *RAB3IL1* | ENSG00000167994.13 | 3.96 | 3.28 | 0.67 | 1.59 | 0.001819133 | 0.009922312 |
| 79699 | *ZYG11B* | ENSG00000162378.13 | 7.21 | 6.54 | 0.67 | 1.59 | 5.82001E-07 | 3.17519E-05 |
| 65018 | *PINK1* | ENSG00000158828.8 | 8.36 | 7.69 | 0.67 | 1.59 | 8.6912E-08 | 8.41588E-06 |
| 388677 | *NOTCH2NLA* | ENSG00000264343.7 | 5.88 | 5.21 | 0.67 | 1.59 | 6.9485E-06 | 0.000164494 |
| 221424 | *LRRC73* | ENSG00000204052.4 | 4.28 | 3.61 | 0.67 | 1.59 | 0.000200667 | 0.001935108 |
| 79956 | *ERMP1* | ENSG00000099219.15 | 4.71 | 4.04 | 0.67 | 1.59 | 7.1904E-05 | 0.000926948 |
| 169200 | *TMEM64* | ENSG00000180694.14 | 6.99 | 6.32 | 0.67 | 1.59 | 5.49235E-06 | 0.000138313 |
| 51023 | *MRPS18C* | ENSG00000163319.11 | 4.91 | 4.24 | 0.67 | 1.59 | 0.000125062 | 0.001371089 |
| 54820 | *NDE1* | ENSG00000072864.16 | 6.04 | 5.38 | 0.67 | 1.59 | 2.96223E-05 | 0.000485373 |
| 27338 | *UBE2S* | ENSG00000108106.14 | 8.19 | 7.53 | 0.67 | 1.59 | 5.38898E-05 | 0.000752402 |
| 11004 | *KIF2C* | ENSG00000142945.13 | 7.04 | 6.37 | 0.66 | 1.59 | 2.05077E-06 | 7.10855E-05 |
| 375 | *ARF1* | ENSG00000143761.16 | 9.22 | 8.56 | 0.66 | 1.58 | 1.21993E-07 | 1.07706E-05 |
| 4126 | *MANBA* | ENSG00000109323.11 | 5.39 | 4.72 | 0.66 | 1.58 | 0.000187189 | 0.001832698 |
| 5091 | *PC* | ENSG00000173599.15 | 6.19 | 5.53 | 0.66 | 1.58 | 1.0485E-06 | 4.62853E-05 |
| 159013 | *CXorf38* | ENSG00000185753.13 | 4.98 | 4.32 | 0.66 | 1.58 | 0.000208789 | 0.001996299 |
| 23433 | *RHOQ* | ENSG00000119729.12 | 6.70 | 6.04 | 0.66 | 1.58 | 2.27662E-06 | 7.57644E-05 |
| 152185 | *SPICE1* | ENSG00000163611.11 | 4.56 | 3.90 | 0.66 | 1.58 | 0.000207269 | 0.001983993 |
| 110599564 | *EEF1AKMT4* | ENSG00000284753.2 | 3.34 | 2.68 | 0.66 | 1.58 | 0.000418968 | 0.003334165 |
| 90874 | *ZNF697* | ENSG00000143067.5 | 4.52 | 3.86 | 0.66 | 1.58 | 0.000283196 | 0.002510623 |
| 29902 | *FAM216A* | ENSG00000204856.12 | 3.42 | 2.77 | 0.66 | 1.58 | 0.000339846 | 0.002851173 |
| 2114 | *ETS2* | ENSG00000157557.13 | 4.51 | 3.85 | 0.66 | 1.58 | 0.000111615 | 0.00126528 |
| 55802 | *DCP1A* | ENSG00000272886.6 | 6.48 | 5.83 | 0.66 | 1.58 | 1.45691E-05 | 0.000284166 |
| 8573 | *CASK* | ENSG00000147044.23 | 7.21 | 6.55 | 0.66 | 1.58 | 1.67884E-05 | 0.000315069 |
| 5569 | *PKIA* | ENSG00000171033.13 | 6.73 | 6.08 | 0.65 | 1.57 | 2.33594E-06 | 7.72248E-05 |
| 8613 | *PLPP3* | ENSG00000162407.9 | 5.00 | 4.35 | 0.65 | 1.57 | 3.48396E-05 | 0.000548696 |
| 57154 | *SMURF1* | ENSG00000198742.10 | 7.08 | 6.43 | 0.65 | 1.57 | 1.18063E-06 | 4.96362E-05 |
| 1316 | *KLF6* | ENSG00000067082.15 | 6.63 | 5.97 | 0.65 | 1.57 | 1.52413E-06 | 5.86194E-05 |
| 6397 | *SEC14L1* | ENSG00000129657.16 | 8.43 | 7.78 | 0.65 | 1.57 | 1.72807E-07 | 1.37231E-05 |
| 11240 | *PADI2* | ENSG00000117115.13 | 3.69 | 3.04 | 0.65 | 1.57 | 9.79401E-05 | 0.00115445 |
| 120103 | *SLC36A4* | ENSG00000180773.15 | 6.65 | 6.00 | 0.65 | 1.57 | 1.05624E-06 | 4.63543E-05 |
| 2191 | *FAP* | ENSG00000078098.15 | 6.44 | 5.80 | 0.65 | 1.57 | 1.12886E-06 | 4.82652E-05 |
| 3306 | *HSPA2* | ENSG00000126803.10 | 8.04 | 7.40 | 0.65 | 1.56 | 3.19541E-07 | 2.04821E-05 |

| | | | | | | | | |
|---|---|---|---|---|---|---|---|---|
| 79752 | *ZFAND1* | ENSG00000104231.11 | 5.93 | 5.29 | 0.65 | 1.56 | 1.4323E-05 | 0.000282118 |
| 2309 | *FOXO3* | ENSG00000118689.15 | 6.14 | 5.49 | 0.64 | 1.56 | 2.39907E-06 | 7.79915E-05 |
| 254170 | *FBXO33* | ENSG00000165355.8 | 5.40 | 4.76 | 0.64 | 1.56 | 1.29678E-05 | 0.000260553 |
| 2553 | *GABPB1* | ENSG00000104064.18 | 4.90 | 4.26 | 0.64 | 1.56 | 0.000908076 | 0.005864595 |
| 345757 | *FAM174A* | ENSG00000174132.9 | 4.91 | 4.27 | 0.64 | 1.56 | 6.56658E-05 | 0.000871306 |
| 79693 | *YRDC* | ENSG00000196449.4 | 4.82 | 4.18 | 0.64 | 1.56 | 7.42097E-05 | 0.000946315 |
| 144717 | *PHETA1* | ENSG00000198324.14 | 4.37 | 3.73 | 0.64 | 1.56 | 3.15656E-05 | 0.000509652 |
| 1534 | *CYB561* | ENSG00000008283.16 | 7.38 | 6.74 | 0.64 | 1.56 | 2.67192E-06 | 8.3897E-05 |
| 6376 | *CX3CL1* | ENSG00000006210.7 | 7.46 | 6.82 | 0.64 | 1.56 | 1.18825E-07 | 1.06158E-05 |
| 5167 | *ENPP1* | ENSG00000197594.14 | 7.04 | 6.40 | 0.64 | 1.56 | 5.90232E-07 | 3.17519E-05 |
| 55031 | *USP47* | ENSG00000170242.19 | 6.38 | 5.74 | 0.64 | 1.56 | 0.000120958 | 0.001345942 |
| 84886 | *C1orf198* | ENSG00000119280.17 | 5.83 | 5.19 | 0.64 | 1.56 | 1.85232E-06 | 6.68306E-05 |
| 894 | *CCND2* | ENSG00000118971.9 | 7.48 | 6.84 | 0.64 | 1.56 | 3.24947E-06 | 9.73479E-05 |
| 10447 | *FAM3C* | ENSG00000196937.11 | 8.70 | 8.06 | 0.64 | 1.56 | 6.84449E-07 | 3.47057E-05 |
| 8660 | *IRS2* | ENSG00000185950.9 | 9.33 | 8.70 | 0.64 | 1.55 | 7.3343E-07 | 3.5974E-05 |
| 6487 | *ST3GAL3* | ENSG00000126091.21 | 5.22 | 4.59 | 0.63 | 1.55 | 1.57674E-05 | 0.000300702 |
| 7277 | *TUBA4A* | ENSG00000127824.15 | 4.95 | 4.32 | 0.63 | 1.55 | 0.000610936 | 0.004399973 |
| 220213 | *OTUD1* | ENSG00000165312.7 | 4.38 | 3.75 | 0.63 | 1.55 | 4.88841E-05 | 0.000703453 |
| 2531 | *KDSR* | ENSG00000119537.18 | 6.90 | 6.27 | 0.63 | 1.55 | 2.43705E-05 | 0.00041803 |
| 1843 | *DUSP1* | ENSG00000120129.6 | 4.94 | 4.30 | 0.63 | 1.55 | 1.42642E-05 | 0.000281788 |
| 2057 | *EPOR* | ENSG00000187266.14 | 4.72 | 4.09 | 0.63 | 1.55 | 0.000307526 | 0.002662566 |
| 122416 | *ANKRD9* | ENSG00000156381.9 | 5.57 | 4.94 | 0.63 | 1.55 | 7.67871E-05 | 0.000971751 |
| 5564 | *PRKAB1* | ENSG00000111725.11 | 6.32 | 5.69 | 0.63 | 1.55 | 1.76687E-05 | 0.000327394 |
| 10793 | *ZNF273* | ENSG00000198039.12 | 2.99 | 2.37 | 0.63 | 1.55 | 0.00152379 | 0.008689425 |
| 10769 | *PLK2* | ENSG00000145632.15 | 6.43 | 5.81 | 0.63 | 1.54 | 0.0002422 | 0.002239791 |
| 65249 | *ZSWIM4* | ENSG00000132003.10 | 3.69 | 3.06 | 0.63 | 1.54 | 8.42723E-05 | 0.001035372 |
| 55198 | *APPL2* | ENSG00000136044.12 | 6.34 | 5.71 | 0.63 | 1.54 | 4.57605E-05 | 0.000671378 |
| 54733 | *SLC35F2* | ENSG00000110660.15 | 5.42 | 4.79 | 0.62 | 1.54 | 0.000106548 | 0.001223252 |
| 1058 | *CENPA* | ENSG00000115163.15 | 5.53 | 4.90 | 0.62 | 1.54 | 5.37347E-06 | 0.000135851 |
| 22820 | *COPG1* | ENSG00000181789.14 | 8.39 | 7.76 | 0.62 | 1.54 | 1.11961E-06 | 4.8012E-05 |
| 2805 | *GOT1* | ENSG00000120053.12 | 7.20 | 6.57 | 0.62 | 1.54 | 1.65614E-06 | 6.15274E-05 |
| 1040 | *CDS1* | ENSG00000163624.6 | 5.11 | 4.49 | 0.62 | 1.54 | 1.64502E-05 | 0.000310567 |
| 753 | *LDLRAD4* | ENSG00000168675.19 | 4.25 | 3.62 | 0.62 | 1.54 | 0.000151644 | 0.001579481 |
| 81579 | *PLA2G12A* | ENSG00000123739.11 | 6.78 | 6.15 | 0.62 | 1.54 | 6.95941E-06 | 0.000164494 |
| 9819 | *TSC22D2* | ENSG00000196428.13 | 6.00 | 5.38 | 0.62 | 1.54 | 0.000121242 | 0.001345942 |
| 355 | *FAS* | ENSG00000026103.23 | 5.05 | 4.43 | 0.62 | 1.54 | 5.65868E-05 | 0.000779184 |
| 56929 | *FEM1C* | ENSG00000145780.8 | 6.20 | 5.58 | 0.62 | 1.54 | 9.49584E-07 | 4.33201E-05 |
| 995 | *CDC25C* | ENSG00000158402.21 | 4.68 | 4.06 | 0.62 | 1.54 | 0.000144777 | 0.001527029 |
| 283991 | *UBALD2* | ENSG00000185262.9 | 5.19 | 4.58 | 0.62 | 1.53 | 2.66933E-05 | 0.000450664 |
| 3099 | *HK2* | ENSG00000159399.10 | 4.38 | 3.77 | 0.62 | 1.53 | 0.000309621 | 0.002670744 |
| 11030 | *RBPMS* | ENSG00000157110.16 | 5.79 | 5.18 | 0.62 | 1.53 | 0.001476284 | 0.008479733 |
| 7253 | *TSHR* | ENSG00000165409.18 | 5.53 | 4.92 | 0.61 | 1.53 | 0.000173158 | 0.001729156 |
| 4084 | *MXD1* | ENSG00000059728.11 | 4.46 | 3.85 | 0.61 | 1.53 | 0.000861034 | 0.005665591 |
| 2137 | *EXTL3* | ENSG00000012232.9 | 7.97 | 7.36 | 0.61 | 1.53 | 0.001697524 | 0.009439843 |
| 9451 | *EIF2AK3* | ENSG00000172071.15 | 6.06 | 5.45 | 0.61 | 1.53 | 5.32892E-06 | 0.000135793 |
| 11186 | *RASSF1* | ENSG00000068028.18 | 3.88 | 3.26 | 0.61 | 1.53 | 0.000104647 | 0.001203562 |
| 57563 | *KLHL8* | ENSG00000145332.14 | 5.41 | 4.80 | 0.61 | 1.53 | 0.000625487 | 0.00446832 |
| 653464 | *SRGAP2C* | ENSG00000171943.12 | 6.35 | 5.74 | 0.61 | 1.52 | 0.00011677 | 0.001312814 |
| 7272 | *TTK* | ENSG00000112742.10 | 6.99 | 6.38 | 0.61 | 1.52 | 4.09477E-06 | 0.000113392 |
| 3106 | *HLA-B* | ENSG00000234745.13 | 7.26 | 6.66 | 0.61 | 1.52 | 2.59646E-06 | 8.23678E-05 |
| 54947 | *LPCAT2* | ENSG00000087253.13 | 6.58 | 5.97 | 0.61 | 1.52 | 1.22554E-05 | 0.000249244 |
| 50640 | *PNPLA8* | ENSG00000135241.17 | 6.81 | 6.20 | 0.61 | 1.52 | 3.48867E-05 | 0.000548863 |
| 9746 | *CLSTN3* | ENSG00000139182.15 | 6.91 | 6.31 | 0.61 | 1.52 | 0.001052222 | 0.00653946 |
| 54206 | *ERRFI1* | ENSG00000116285.13 | 5.96 | 5.35 | 0.61 | 1.52 | 2.93088E-06 | 8.88677E-05 |
| 285704 | *RGMB* | ENSG00000174136.13 | 5.81 | 5.21 | 0.60 | 1.52 | 2.03895E-05 | 0.000366059 |
| 5832 | *ALDH18A1* | ENSG00000059573.9 | 7.17 | 6.57 | 0.60 | 1.52 | 4.11397E-06 | 0.000113457 |
| 5428 | *POLG* | ENSG00000140521.17 | 6.43 | 5.83 | 0.60 | 1.52 | 0.001731491 | 0.00957199 |
| 142891 | *SAMD8* | ENSG00000156671.15 | 6.33 | 5.72 | 0.60 | 1.52 | 2.57842E-06 | 8.19904E-05 |
| 9553 | *MRPL33* | ENSG00000243147.8 | 6.37 | 5.76 | 0.60 | 1.52 | 2.66917E-05 | 0.000450664 |
| 699 | *BUB1* | ENSG00000169679.15 | 8.23 | 7.62 | 0.60 | 1.52 | 6.20738E-07 | 3.269E-05 |
| 590 | *BCHE* | ENSG00000114200.10 | 8.52 | 7.92 | 0.60 | 1.52 | 9.52795E-06 | 0.000204878 |
| 6925 | *TCF4* | ENSG00000196628.20 | 8.50 | 7.90 | 0.60 | 1.52 | 1.24699E-06 | 5.18452E-05 |
| 6652 | *SORD* | ENSG00000140263.15 | 7.70 | 7.10 | 0.60 | 1.52 | 2.76251E-06 | 8.54897E-05 |
| 4124 | *MAN2A1* | ENSG00000112893.10 | 8.35 | 7.75 | 0.60 | 1.52 | 3.78492E-07 | 2.29905E-05 |
| 5480 | *PPIC* | ENSG00000168938.6 | 4.76 | 4.15 | 0.60 | 1.52 | 4.49031E-05 | 0.000663338 |
| 388 | *RHOB* | ENSG00000143878.10 | 8.29 | 7.69 | 0.60 | 1.52 | 3.9299E-06 | 0.000110457 |
| 9049 | *AIP* | ENSG00000110711.11 | 6.00 | 5.40 | 0.60 | 1.51 | 9.00021E-06 | 0.000197491 |
| 1875 | *E2F5* | ENSG00000133740.11 | 3.75 | 3.15 | 0.60 | 1.51 | 0.000268117 | 0.002410853 |
| 4957 | *ODF2* | ENSG00000136811.17 | 6.80 | 6.20 | 0.60 | 1.51 | 3.43978E-06 | 0.000101033 |
| 23348 | *DOCK9* | ENSG00000088387.21 | 6.58 | 5.99 | 0.60 | 1.51 | 5.84438E-05 | 0.000798977 |
| 27244 | *SESN1* | ENSG00000080546.14 | 5.71 | 5.11 | 0.60 | 1.51 | 3.15728E-05 | 0.000509652 |
| 26036 | *ZNF451* | ENSG00000112200.17 | 6.39 | 5.79 | 0.60 | 1.51 | 0.001023842 | 0.006400188 |
| 6890 | *TAP1* | ENSG00000168394.12 | 6.35 | 5.76 | 0.60 | 1.51 | 7.94989E-06 | 0.000181337 |
| 63925 | *ZNF335* | ENSG00000198026.8 | 4.54 | 3.94 | 0.60 | 1.51 | 0.000312424 | 0.00267953 |
| 54910 | *SEMA4C* | ENSG00000168758.11 | 4.85 | 4.25 | 0.59 | 1.51 | 0.001738869 | 0.009581016 |
| 30850 | *CDR2L* | ENSG00000109089.7 | 3.67 | 3.08 | 0.59 | 1.51 | 0.000822319 | 0.005478113 |
| 79801 | *SHCBP1* | ENSG00000171241.9 | 7.24 | 6.65 | 0.59 | 1.51 | 8.52752E-05 | 0.00104311 |
| 23173 | *METAP1* | ENSG00000164024.12 | 6.19 | 5.60 | 0.59 | 1.51 | 3.37066E-06 | 9.9587E-05 |
| 5255 | *PHKA1* | ENSG00000067177.15 | 6.60 | 6.01 | 0.59 | 1.51 | 7.57667E-06 | 0.000174951 |
| 5507 | *PPP1R3C* | ENSG00000119938.9 | 4.49 | 3.89 | 0.59 | 1.51 | 0.000306323 | 0.002657575 |
| 55355 | *HJURP* | ENSG00000123485.12 | 5.59 | 5.00 | 0.59 | 1.51 | 0.001370691 | 0.008024314 |
| 91272 | *BOD1* | ENSG00000145919.11 | 6.20 | 5.60 | 0.59 | 1.51 | 4.32382E-05 | 0.000646376 |
| 5190 | *PEX6* | ENSG00000124587.14 | 5.38 | 4.79 | 0.59 | 1.51 | 3.71681E-05 | 0.000574797 |
| 6525 | *SMTN* | ENSG00000183963.19 | 5.69 | 5.10 | 0.59 | 1.51 | 0.000121233 | 0.001345942 |
| 10484 | *SEC23A* | ENSG00000100934.15 | 8.08 | 7.49 | 0.59 | 1.51 | 2.29935E-07 | 1.65124E-05 |
| 22987 | *SV2C* | ENSG00000122012.14 | 8.44 | 7.85 | 0.59 | 1.51 | 1.0213E-06 | 4.58941E-05 |
| 3838 | *KPNA2* | ENSG00000182481.10 | 9.20 | 8.61 | 0.59 | 1.51 | 3.22553E-07 | 2.04821E-05 |
| 4758 | *NEU1* | ENSG00000204386.12 | 6.75 | 6.15 | 0.59 | 1.51 | 1.89851E-06 | 6.76835E-05 |
| 10957 | *PNRC1* | ENSG00000146278.11 | 7.00 | 6.41 | 0.59 | 1.51 | 1.89044E-05 | 0.000344111 |

| | | | | | | | | |
|---|---|---|---|---|---|---|---|---|
| 5121 | *PCP4* | ENSG00000183036.11 | 10.58 | 9.99 | 0.59 | 1.51 | 5.19197E-07 | 2.92956E-05 |
| 5007 | *OSBP* | ENSG00000110048.12 | 8.39 | 7.80 | 0.59 | 1.51 | 2.5527E-07 | 1.80724E-05 |
| 10678 | *B3GNT2* | ENSG00000170340.11 | 4.49 | 3.90 | 0.59 | 1.51 | 0.000190416 | 0.001855817 |
| 80381 | *CD276* | ENSG00000103855.18 | 7.16 | 6.57 | 0.59 | 1.51 | 3.42501E-05 | 0.000541686 |
| 387103 | *CENPW* | ENSG00000203760.9 | 5.05 | 4.46 | 0.59 | 1.51 | 0.000110477 | 0.001255965 |
| 25963 | *TMEM87A* | ENSG00000103978.16 | 7.90 | 7.31 | 0.59 | 1.50 | 5.29267E-06 | 0.000135328 |
| 221830 | *POLR1F* | ENSG00000105849.6 | 6.44 | 5.85 | 0.59 | 1.50 | 1.72028E-06 | 6.32837E-05 |
| 5794 | *PTPRH* | ENSG00000080031.10 | 7.08 | 6.49 | 0.59 | 1.50 | 3.99069E-05 | 0.000607467 |
| 51278 | *IER5* | ENSG00000162783.11 | 4.54 | 3.95 | 0.59 | 1.50 | 0.000114409 | 0.001291963 |
| 4888 | *NPY6R* | ENSG00000226306.7 | 8.65 | 8.06 | 0.58 | 1.50 | 1.72701E-07 | 1.37231E-05 |
| 55379 | *LRRC59* | ENSG00000108829.10 | 8.59 | 8.01 | 0.58 | 1.50 | 1.48792E-07 | 1.21814E-05 |
| 53616 | *ADAM22* | ENSG00000008277.15 | 6.56 | 5.97 | 0.58 | 1.50 | 3.33277E-05 | 0.000532144 |
| 3309 | *HSPA5* | ENSG00000044574.9 | 10.29 | 9.71 | 0.58 | 1.50 | 3.56525E-07 | 2.21119E-05 |
| 9534 | *ZNF254* | ENSG00000213096.11 | 3.91 | 3.33 | 0.58 | 1.50 | 0.001283913 | 0.007640861 |
| 10289 | *EIF1B* | ENSG00000114784.4 | 6.82 | 6.23 | 0.58 | 1.50 | 1.99659E-06 | 6.98526E-05 |
| 360023 | *ZBTB41* | ENSG00000177888.8 | 6.75 | 6.17 | 0.58 | 1.50 | 2.33987E-05 | 0.000405532 |
| 23787 | *MTCH1* | ENSG00000137409.20 | 8.57 | 7.99 | 0.58 | 1.50 | 5.23963E-07 | 2.94538E-05 |
| 92597 | *MOB1B* | ENSG00000173542.9 | 5.84 | 5.26 | 0.58 | 1.50 | 8.3226E-06 | 0.000186717 |
| 10802 | *SEC24A* | ENSG00000113615.13 | 7.09 | 6.51 | 0.58 | 1.50 | 2.9052E-06 | 8.84466E-05 |
| 133746 | *JMY* | ENSG00000152409.9 | 6.82 | 6.24 | 0.58 | 1.49 | 4.04806E-06 | 0.000113142 |
| 126731 | *CCSAP* | ENSG00000154429.11 | 6.13 | 5.55 | 0.58 | 1.49 | 1.60793E-05 | 0.000305037 |
| 4085 | *MAD2L1* | ENSG00000164109.14 | 7.37 | 6.79 | 0.58 | 1.49 | 8.14101E-05 | 0.001010488 |
| 25924 | *MYRIP* | ENSG00000170011.14 | 6.13 | 5.56 | 0.58 | 1.49 | 4.08031E-06 | 0.000113337 |
| 2037 | *EPB41L2* | ENSG00000079819.19 | 7.73 | 7.15 | 0.58 | 1.49 | 2.26753E-06 | 7.56296E-05 |
| 56005 | *MYDGF* | ENSG00000074842.8 | 6.61 | 6.03 | 0.58 | 1.49 | 1.27181E-06 | 5.25857E-05 |
| 7456 | *WIPF1* | ENSG00000115935.18 | 5.39 | 4.81 | 0.58 | 1.49 | 0.000314055 | 0.002687371 |
| 64746 | *ACBD3* | ENSG00000182827.9 | 7.06 | 6.49 | 0.57 | 1.49 | 1.65182E-06 | 6.15274E-05 |
| 10383 | *TUBB4B* | ENSG00000188229.6 | 9.65 | 9.08 | 0.57 | 1.49 | 2.65736E-07 | 1.84245E-05 |
| 58528 | *RRAGD* | ENSG00000025039.15 | 6.88 | 6.30 | 0.57 | 1.49 | 4.65758E-06 | 0.00012329 |
| 4751 | *NEK2* | ENSG00000117650.13 | 6.62 | 6.05 | 0.57 | 1.49 | 3.26844E-06 | 9.7527E-05 |
| 55785 | *FGD6* | ENSG00000180263.14 | 5.38 | 4.80 | 0.57 | 1.49 | 0.000180781 | 0.001787447 |
| 27248 | *ERLEC1* | ENSG00000068912.15 | 7.66 | 7.08 | 0.57 | 1.49 | 2.97188E-07 | 1.95636E-05 |
| 8566 | *PDXK* | ENSG00000160209.19 | 7.51 | 6.94 | 0.57 | 1.49 | 4.26866E-05 | 0.000640683 |
| 22824 | *HSPA4L* | ENSG00000164070.12 | 6.39 | 5.82 | 0.57 | 1.49 | 5.35939E-06 | 0.000135851 |
| 3148 | *HMGB2* | ENSG00000164104.12 | 7.48 | 6.90 | 0.57 | 1.49 | 6.47798E-07 | 3.36429E-05 |
| 3987 | *LIMS1* | ENSG00000169756.16 | 6.83 | 6.26 | 0.57 | 1.49 | 0.000228575 | 0.002129882 |
| 6100 | *RP9* | ENSG00000164610.10 | 4.17 | 3.60 | 0.57 | 1.49 | 0.000786053 | 0.005283417 |
| 57820 | *CCNB1IP1* | ENSG00000100814.18 | 5.55 | 4.98 | 0.57 | 1.49 | 6.52899E-05 | 0.000867968 |
| 1063 | *CENPF* | ENSG00000117724.14 | 8.85 | 8.28 | 0.57 | 1.49 | 0.000165364 | 0.001676993 |
| 2 | *A2M* | ENSG00000175899.15 | 9.88 | 9.31 | 0.57 | 1.48 | 2.35733E-06 | 7.75832E-05 |
| 79733 | *E2F8* | ENSG00000129173.13 | 4.48 | 3.91 | 0.57 | 1.48 | 0.001171462 | 0.007108297 |
| 8531 | *YBX3* | ENSG00000060138.13 | 8.48 | 7.91 | 0.57 | 1.48 | 5.14037E-06 | 0.000132336 |
| 23275 | *POFUT2* | ENSG00000186866.17 | 7.02 | 6.45 | 0.57 | 1.48 | 0.000474369 | 0.003647438 |
| 80014 | *WWC2* | ENSG00000151718.16 | 6.93 | 6.36 | 0.57 | 1.48 | 4.65337E-05 | 0.000680063 |
| 65061 | *CDK15* | ENSG00000138395.17 | 4.95 | 4.38 | 0.57 | 1.48 | 0.000120577 | 0.00134553 |
| 9911 | *TMCC2* | ENSG00000133069.17 | 3.49 | 2.92 | 0.57 | 1.48 | 0.000654042 | 0.004612921 |
| 255743 | *NPNT* | ENSG00000168743.13 | 6.58 | 6.02 | 0.57 | 1.48 | 5.0384E-06 | 0.000130157 |
| 166929 | *SGMS2* | ENSG00000164023.15 | 4.78 | 4.22 | 0.57 | 1.48 | 0.00044616 | 0.003493176 |
| 5066 | *PAM* | ENSG00000145730.21 | 9.85 | 9.28 | 0.57 | 1.48 | 2.96834E-07 | 1.95636E-05 |
| 5611 | *DNAJC3* | ENSG00000102580.15 | 8.47 | 7.91 | 0.57 | 1.48 | 1.35046E-06 | 5.44866E-05 |
| 8518 | *ELP1* | ENSG00000070061.16 | 10.24 | 9.67 | 0.57 | 1.48 | 2.51744E-06 | 8.07355E-05 |
| 55504 | *TNFRSF19* | ENSG00000127863.16 | 8.32 | 7.76 | 0.57 | 1.48 | 2.8814E-07 | 1.93958E-05 |
| 983 | *CDK1* | ENSG00000170312.17 | 8.41 | 7.85 | 0.57 | 1.48 | 1.98973E-07 | 1.49319E-05 |
| 2512 | *FTL* | ENSG00000087086.15 | 9.64 | 9.07 | 0.57 | 1.48 | 2.573E-06 | 8.19904E-05 |
| 9788 | *MTSS1* | ENSG00000170873.19 | 7.70 | 7.13 | 0.56 | 1.48 | 4.87418E-06 | 0.000127405 |
| 3107 | *HLA-C* | ENSG00000204525.18 | 5.94 | 5.37 | 0.56 | 1.48 | 6.50838E-06 | 0.000157302 |
| 152926 | *PPM1K* | ENSG00000163644.15 | 5.26 | 4.69 | 0.56 | 1.48 | 6.066E-05 | 0.000822445 |
| 81567 | *TXNDC5* | ENSG00000239264.9 | 8.69 | 8.13 | 0.56 | 1.48 | 2.66381E-07 | 1.84245E-05 |
| 100526694 | *MSANTD3-TM* | ENSG00000251349.3 | 4.83 | 4.27 | 0.56 | 1.48 | 0.000660096 | 0.004642633 |
| 5597 | *MAPK6* | ENSG00000069956.14 | 7.25 | 6.69 | 0.56 | 1.48 | 7.88741E-06 | 0.00018046 |
| 890 | *CCNA2* | ENSG00000145386.11 | 7.63 | 7.07 | 0.56 | 1.47 | 8.84668E-06 | 0.000195265 |
| 57553 | *MICAL3* | ENSG00000243156.9 | 6.95 | 6.39 | 0.56 | 1.47 | 0.000278371 | 0.002477972 |
| 201895 | *SMIM14* | ENSG00000163683.12 | 7.27 | 6.71 | 0.56 | 1.47 | 3.94076E-05 | 0.000601086 |
| 146909 | *KIF18B* | ENSG00000186185.14 | 6.64 | 6.08 | 0.56 | 1.47 | 9.09149E-05 | 0.001089019 |
| 221079 | *ARL5B* | ENSG00000165997.5 | 7.30 | 6.75 | 0.56 | 1.47 | 2.11675E-06 | 7.20414E-05 |
| 6443 | *SGCB* | ENSG00000163069.13 | 7.59 | 7.03 | 0.56 | 1.47 | 3.37317E-07 | 2.11832E-05 |
| 11065 | *UBE2C* | ENSG00000175063.17 | 7.37 | 6.82 | 0.56 | 1.47 | 2.75853E-06 | 8.54897E-05 |
| 80854 | *SETD7* | ENSG00000145391.14 | 8.01 | 7.46 | 0.56 | 1.47 | 1.6437E-06 | 6.15219E-05 |
| 55143 | *CDCA8* | ENSG00000134690.11 | 6.85 | 6.29 | 0.56 | 1.47 | 7.085E-07 | 3.55095E-05 |
| 7153 | *TOP2A* | ENSG00000131747.15 | 10.15 | 9.59 | 0.56 | 1.47 | 4.62985E-07 | 2.67474E-05 |
| 677 | *ZFP36L1* | ENSG00000185650.10 | 5.43 | 4.88 | 0.56 | 1.47 | 7.9232E-05 | 0.000990319 |
| 164832 | *LONRF2* | ENSG00000170500.13 | 7.17 | 6.62 | 0.56 | 1.47 | 4.11981E-06 | 0.000113457 |
| 5144 | *PDE4D* | ENSG00000113448.20 | 6.50 | 5.94 | 0.55 | 1.47 | 0.000281378 | 0.002497588 |
| 78992 | *YIPF2* | ENSG00000130733.11 | 6.14 | 5.59 | 0.55 | 1.47 | 0.000717215 | 0.004940191 |
| 8312 | *AXIN1* | ENSG00000103126.15 | 5.47 | 4.92 | 0.55 | 1.47 | 0.000102337 | 0.00118828 |
| 728769 | *SCAMP1-AS1* | ENSG00000245556.4 | 3.64 | 3.08 | 0.55 | 1.47 | 0.000384203 | 0.003113659 |
| 5371 | *PML* | ENSG00000140464.20 | 6.00 | 5.44 | 0.55 | 1.47 | 0.000432873 | 0.003410494 |
| 114984 | *FLYWCH2* | ENSG00000162076.13 | 4.96 | 4.40 | 0.55 | 1.47 | 0.000192065 | 0.001867039 |
| 201164 | *PLD6* | ENSG00000179598.6 | 5.17 | 4.62 | 0.55 | 1.47 | 3.15795E-05 | 0.000509652 |
| 5202 | *PFDN2* | ENSG00000143256.5 | 5.74 | 5.19 | 0.55 | 1.47 | 4.75485E-05 | 0.000690501 |
| 91663 | *MYADM* | ENSG00000179820.16 | 6.89 | 6.34 | 0.55 | 1.47 | 1.11666E-05 | 0.000232958 |
| 81566 | *CSRNP2* | ENSG00000110925.7 | 6.85 | 6.30 | 0.55 | 1.46 | 8.28494E-06 | 0.00018643 |
| 143662 | *MUC15* | ENSG00000169550.14 | 7.24 | 6.69 | 0.55 | 1.46 | 1.6145E-06 | 6.10802E-05 |
| 65986 | *ZBTB10* | ENSG00000205189.12 | 5.86 | 5.31 | 0.55 | 1.46 | 1.89801E-05 | 0.000344881 |
| 144699 | *FBXL14* | ENSG00000171823.7 | 4.46 | 3.91 | 0.55 | 1.46 | 0.000307847 | 0.002662566 |
| 162962 | *ZNF836* | ENSG00000196267.14 | 6.22 | 5.67 | 0.55 | 1.46 | 0.00015621 | 0.001611383 |
| 55113 | *XKR8* | ENSG00000158156.9 | 4.20 | 3.65 | 0.55 | 1.46 | 0.000441784 | 0.003469771 |
| 9069 | *CLDN12* | ENSG00000157224.16 | 7.09 | 6.54 | 0.55 | 1.46 | 2.02835E-06 | 7.07547E-05 |
| 6814 | *STXBP3* | ENSG00000116266.11 | 6.12 | 5.57 | 0.55 | 1.46 | 2.1783E-05 | 0.00038509 |

| | | | | | | | | |
|---|---|---|---|---|---|---|---|---|
| 4209 | *MEF2D* | ENSG00000116604.19 | 6.74 | 6.20 | 0.55 | 1.46 | 4.14807E-06 | 0.000113721 |
| 65078 | *RTN4R* | ENSG00000040608.14 | 4.42 | 3.87 | 0.55 | 1.46 | 0.001132847 | 0.006927016 |
| 54681 | *P4HTM* | ENSG00000178467.18 | 4.92 | 4.37 | 0.55 | 1.46 | 0.000149811 | 0.001562552 |
| 54532 | *USP53* | ENSG00000145390.12 | 4.86 | 4.32 | 0.55 | 1.46 | 0.000672073 | 0.004704822 |
| 83667 | *SESN2* | ENSG00000130766.5 | 4.88 | 4.34 | 0.54 | 1.46 | 5.55324E-05 | 0.00076819 |
| 64645 | *MFSD14A* | ENSG00000156875.14 | 6.99 | 6.45 | 0.54 | 1.46 | 2.99646E-05 | 0.000489378 |
| 2149 | *F2R* | ENSG00000181104.7 | 4.04 | 3.49 | 0.54 | 1.46 | 0.000561524 | 0.004129305 |
| 64783 | *RBM15* | ENSG00000162775.17 | 5.24 | 4.70 | 0.54 | 1.46 | 2.3077E-05 | 0.000400882 |
| 10184 | *LHFPL2* | ENSG00000145685.14 | 5.86 | 5.31 | 0.54 | 1.46 | 1.36853E-05 | 0.000272771 |
| 23731 | *TMEM245* | ENSG00000106771.13 | 9.75 | 9.21 | 0.54 | 1.46 | 7.76374E-07 | 3.74266E-05 |
| 55761 | *TTC17* | ENSG00000052841.15 | 7.08 | 6.54 | 0.54 | 1.46 | 3.27689E-06 | 9.75851E-05 |
| 3764 | *KCNJ8* | ENSG00000121361.5 | 6.03 | 5.49 | 0.54 | 1.46 | 0.000117648 | 0.001320697 |
| 83857 | *TMTC1* | ENSG00000133687.16 | 8.62 | 8.08 | 0.54 | 1.45 | 3.61972E-06 | 0.00010368 |
| 10389 | *SCML2* | ENSG00000102098.19 | 5.79 | 5.25 | 0.54 | 1.45 | 2.50764E-05 | 0.000429158 |
| 8470 | *SORBS2* | ENSG00000154556.18 | 8.15 | 7.61 | 0.54 | 1.45 | 1.31585E-05 | 0.00026368 |
| 2791 | *GNG11* | ENSG00000127920.6 | 7.20 | 6.65 | 0.54 | 1.45 | 1.84163E-05 | 0.000337497 |
| 25987 | *TSKU* | ENSG00000182704.8 | 6.91 | 6.37 | 0.54 | 1.45 | 2.20667E-06 | 7.44261E-05 |
| 219654 | *ZCCHC24* | ENSG00000165424.7 | 5.26 | 4.72 | 0.54 | 1.45 | 5.27482E-05 | 0.000738667 |
| 8639 | *AOC3* | ENSG00000131471.7 | 3.59 | 3.05 | 0.54 | 1.45 | 0.001234182 | 0.007394744 |
| 273 | *AMPH* | ENSG00000078053.17 | 5.70 | 5.16 | 0.54 | 1.45 | 2.08575E-05 | 0.000371793 |
| 10725 | *NFAT5* | ENSG00000102908.22 | 6.45 | 5.91 | 0.54 | 1.45 | 8.179E-05 | 0.00101286 |
| 1212 | *CLTB* | ENSG00000175416.16 | 5.70 | 5.16 | 0.54 | 1.45 | 0.000167215 | 0.001685862 |
| 64319 | *FBRS* | ENSG00000156860.16 | 6.55 | 6.01 | 0.54 | 1.45 | 3.78585E-05 | 0.000581594 |
| 57555 | *NLGN2* | ENSG00000169992.10 | 7.12 | 6.59 | 0.54 | 1.45 | 0.00141268 | 0.008211821 |
| 122961 | *ISCA2* | ENSG00000165898.14 | 3.87 | 3.33 | 0.54 | 1.45 | 0.001631458 | 0.009177868 |
| 80727 | *TTYH3* | ENSG00000136295.15 | 7.14 | 6.60 | 0.54 | 1.45 | 2.38096E-06 | 7.78557E-05 |
| 10892 | *MALT1* | ENSG00000172175.15 | 5.20 | 4.67 | 0.54 | 1.45 | 6.87099E-05 | 0.00090067 |
| 54813 | *KLHL28* | ENSG00000179454.14 | 5.51 | 4.98 | 0.53 | 1.45 | 0.000200713 | 0.001935108 |
| 91748 | *MIDEAS* | ENSG00000156030.14 | 4.85 | 4.32 | 0.53 | 1.45 | 0.000691977 | 0.004808279 |
| 283237 | *TTC9C* | ENSG00000162222.14 | 4.92 | 4.39 | 0.53 | 1.45 | 0.000104554 | 0.001203562 |
| 23551 | *RASD2* | ENSG00000100302.7 | 6.31 | 5.77 | 0.53 | 1.45 | 6.92008E-06 | 0.000164341 |
| 1486 | *CTBS* | ENSG00000117151.13 | 5.40 | 4.87 | 0.53 | 1.45 | 6.3011E-05 | 0.000845125 |
| 8411 | *EEA1* | ENSG00000102189.17 | 6.77 | 6.24 | 0.53 | 1.45 | 8.58416E-05 | 0.001047477 |
| 23576 | *DDAH1* | ENSG00000153904.21 | 7.33 | 6.80 | 0.53 | 1.45 | 2.07236E-06 | 7.15034E-05 |
| 6836 | *SURF4* | ENSG00000148248.14 | 9.33 | 8.80 | 0.53 | 1.45 | 8.21311E-05 | 0.001015408 |
| 79990 | *PLEKHH3* | ENSG00000068137.15 | 4.08 | 3.55 | 0.53 | 1.45 | 0.000515858 | 0.003880959 |
| 54853 | *WDR55* | ENSG00000120314.19 | 5.00 | 4.47 | 0.53 | 1.45 | 0.000102726 | 0.001191064 |
| 7111 | *TMOD1* | ENSG00000136842.14 | 5.91 | 5.38 | 0.53 | 1.45 | 9.09647E-06 | 0.000198732 |
| 8702 | *B4GALT4* | ENSG00000121578.13 | 6.80 | 6.27 | 0.53 | 1.45 | 7.94673E-05 | 0.000992284 |
| 54331 | *GNG2* | ENSG00000186469.9 | 3.95 | 3.41 | 0.53 | 1.45 | 0.000684884 | 0.004770035 |
| 4097 | *MAFG* | ENSG00000197063.11 | 6.28 | 5.75 | 0.53 | 1.45 | 2.47362E-06 | 7.98422E-05 |
| 127262 | *TPRG1L* | ENSG00000158109.15 | 5.24 | 4.71 | 0.53 | 1.44 | 5.84501E-05 | 0.000798977 |
| 10112 | *KIF20A* | ENSG00000112984.12 | 7.55 | 7.02 | 0.53 | 1.44 | 1.86217E-06 | 6.70248E-05 |
| 26258 | *BLOC1S6* | ENSG00000104164.12 | 7.66 | 7.13 | 0.53 | 1.44 | 1.90931E-05 | 0.000346226 |
| 7625 | *ZNF74* | ENSG00000185252.19 | 5.54 | 5.01 | 0.53 | 1.44 | 0.000109805 | 0.001250432 |
| 5998 | *RGS3* | ENSG00000138835.22 | 7.57 | 7.04 | 0.53 | 1.44 | 2.68668E-05 | 0.000452574 |
| 5876 | *RABGGTB* | ENSG00000137955.16 | 5.81 | 5.29 | 0.53 | 1.44 | 0.000311111 | 0.00267438 |
| 283638 | *CEP170B* | ENSG00000099814.17 | 6.19 | 5.66 | 0.53 | 1.44 | 1.51539E-05 | 0.000292346 |
| 25798 | *BRI3* | ENSG00000164713.10 | 4.94 | 4.41 | 0.53 | 1.44 | 0.00025582 | 0.002336945 |
| 1831 | *TSC22D3* | ENSG00000157514.18 | 7.44 | 6.92 | 0.53 | 1.44 | 7.24246E-07 | 3.58067E-05 |
| 51309 | *ARMCX1* | ENSG00000126947.13 | 5.62 | 5.10 | 0.53 | 1.44 | 6.02343E-05 | 0.000818892 |
| 147179 | *WIPF2* | ENSG00000171475.14 | 6.58 | 6.05 | 0.53 | 1.44 | 7.52668E-05 | 0.000956545 |
| 6734 | *SRPRA* | ENSG00000182934.12 | 8.40 | 7.87 | 0.53 | 1.44 | 7.28576E-07 | 3.58531E-05 |
| 1315 | *COPB1* | ENSG00000129083.13 | 7.49 | 6.96 | 0.53 | 1.44 | 4.55859E-07 | 2.66225E-05 |
| 55186 | *SLC25A36* | ENSG00000114120.14 | 7.55 | 7.02 | 0.53 | 1.44 | 3.78632E-06 | 0.000106821 |
| 10519 | *CIB1* | ENSG00000185043.12 | 6.26 | 5.74 | 0.52 | 1.44 | 4.32047E-05 | 0.000646376 |
| 59 | *ACTA2* | ENSG00000107796.13 | 5.42 | 4.90 | 0.52 | 1.44 | 0.000261832 | 0.002380279 |
| 6770 | *STAR* | ENSG00000147465.12 | 10.54 | 10.01 | 0.52 | 1.44 | 7.18384E-07 | 3.57027E-05 |
| 84919 | *PPP1R15B* | ENSG00000158615.10 | 7.66 | 7.14 | 0.52 | 1.44 | 1.04817E-06 | 4.62853E-05 |
| 64225 | *ATL2* | ENSG00000119787.14 | 6.95 | 6.42 | 0.52 | 1.44 | 3.59992E-05 | 0.00055933 |
| 84926 | *SPRYD3* | ENSG00000167778.9 | 7.14 | 6.61 | 0.52 | 1.44 | 8.27227E-06 | 0.00018643 |
| 22911 | *WDR47* | ENSG00000085433.17 | 5.06 | 4.54 | 0.52 | 1.44 | 0.000514584 | 0.003877203 |
| 85415 | *RHPN2* | ENSG00000131941.8 | 5.21 | 4.69 | 0.52 | 1.44 | 0.001116109 | 0.006840213 |
| 135112 | *NCOA7* | ENSG00000111912.20 | 5.99 | 5.47 | 0.52 | 1.43 | 0.000658197 | 0.004633619 |
| 85416 | *ZIC5* | ENSG00000139800.9 | 3.64 | 3.12 | 0.52 | 1.43 | 0.000538736 | 0.004014837 |
| 427 | *ASAH1* | ENSG00000104763.20 | 6.10 | 5.58 | 0.52 | 1.43 | 5.90771E-05 | 0.000806503 |
| 5321 | *PLA2G4A* | ENSG00000116711.10 | 3.60 | 3.08 | 0.52 | 1.43 | 0.001682726 | 0.009377352 |
| 3619 | *INCENP* | ENSG00000149503.13 | 7.51 | 6.99 | 0.52 | 1.43 | 0.000136886 | 0.001463335 |
| 353116 | *RILPL1* | ENSG00000188026.13 | 4.22 | 3.70 | 0.52 | 1.43 | 0.001171692 | 0.007108297 |
| 8615 | *USO1* | ENSG00000138768.15 | 7.36 | 6.85 | 0.52 | 1.43 | 1.99208E-06 | 6.98526E-05 |
| 81537 | *SGPP1* | ENSG00000126821.8 | 6.99 | 6.47 | 0.52 | 1.43 | 1.21623E-05 | 0.000248254 |
| 80346 | *REEP4* | ENSG00000168476.12 | 4.74 | 4.23 | 0.52 | 1.43 | 0.000454103 | 0.003535104 |
| 267 | *AMFR* | ENSG00000159461.15 | 8.01 | 7.49 | 0.52 | 1.43 | 1.43086E-06 | 5.6515E-05 |
| 56605 | *ERO1B* | ENSG00000086619.15 | 5.97 | 5.45 | 0.52 | 1.43 | 5.65033E-05 | 0.000778749 |
| 79139 | *DERL1* | ENSG00000136986.10 | 8.06 | 7.54 | 0.52 | 1.43 | 0.000248882 | 0.002288891 |
| 9212 | *AURKB* | ENSG00000178999.13 | 5.41 | 4.89 | 0.52 | 1.43 | 6.04856E-05 | 0.000820821 |
| 57645 | *POGK* | ENSG00000143157.12 | 7.17 | 6.65 | 0.52 | 1.43 | 7.8849E-07 | 3.77105E-05 |
| 1845 | *DUSP3* | ENSG00000108861.9 | 7.70 | 7.19 | 0.52 | 1.43 | 1.03573E-05 | 0.000220501 |
| 9824 | *ARHGAP11A* | ENSG00000198826.11 | 8.16 | 7.64 | 0.52 | 1.43 | 9.46876E-06 | 0.000204191 |
| 5819 | *NECTIN2* | ENSG00000130202.10 | 6.97 | 6.46 | 0.52 | 1.43 | 1.92413E-05 | 0.000348362 |
| 10979 | *FERMT2* | ENSG00000073712.15 | 6.20 | 5.69 | 0.52 | 1.43 | 0.000104578 | 0.001203562 |
| 8214 | *DGCR6* | ENSG00000183628.14 | 4.58 | 4.07 | 0.51 | 1.43 | 0.000129803 | 0.001408658 |
| 1847 | *DUSP5* | ENSG00000138166.6 | 5.04 | 4.52 | 0.51 | 1.43 | 0.000405357 | 0.003251913 |
| 59274 | *TLNRD1* | ENSG00000140406.4 | 4.02 | 3.50 | 0.51 | 1.43 | 0.000634206 | 0.00451128 |
| 6238 | *RRBP1* | ENSG00000125844.17 | 7.64 | 7.12 | 0.51 | 1.43 | 1.45323E-05 | 0.000284166 |
| 25777 | *SUN2* | ENSG00000100242.16 | 8.11 | 7.60 | 0.51 | 1.43 | 7.14834E-06 | 0.000167313 |
| 8086 | *AAAS* | ENSG00000094914.14 | 6.64 | 6.13 | 0.51 | 1.43 | 8.94088E-06 | 0.000196765 |
| 85439 | *STON2* | ENSG00000140022.14 | 4.70 | 4.19 | 0.51 | 1.43 | 0.000330418 | 0.002795513 |
| 22882 | *ZHX2* | ENSG00000178764.8 | 4.97 | 4.46 | 0.51 | 1.43 | 0.000865275 | 0.00567858 |

| | | | | | | | | |
|---|---|---|---|---|---|---|---|---|
| 23192 | *ATG4B* | ENSG00000168397.17 | 4.72 | 4.21 | 0.51 | 1.43 | 0.000911249 | 0.005874974 |
| 29089 | *UBE2T* | ENSG00000077152.11 | 6.08 | 5.56 | 0.51 | 1.42 | 0.000213844 | 0.002027532 |
| 6184 | *RPN1* | ENSG00000163902.12 | 9.09 | 8.58 | 0.51 | 1.42 | 5.65197E-06 | 0.000141149 |
| 253430 | *IPMK* | ENSG00000151151.6 | 5.06 | 4.55 | 0.51 | 1.42 | 0.000615984 | 0.004423755 |
| 79581 | *SLC52A2* | ENSG00000185803.12 | 4.77 | 4.26 | 0.51 | 1.42 | 0.000377629 | 0.003075272 |
| 8904 | *CPNE1* | ENSG00000214078.13 | 6.88 | 6.37 | 0.51 | 1.42 | 0.000115853 | 0.001303481 |
| 7532 | *YWHAG* | ENSG00000170027.7 | 9.58 | 9.07 | 0.51 | 1.42 | 3.60042E-07 | 2.22382E-05 |
| 54867 | *TMEM214* | ENSG00000119777.20 | 7.18 | 6.67 | 0.51 | 1.42 | 1.29933E-05 | 0.000260718 |
| 529 | *ATP6V1E1* | ENSG00000131100.13 | 8.24 | 7.74 | 0.51 | 1.42 | 1.22883E-06 | 5.12318E-05 |
| 51646 | *YPEL5* | ENSG00000119801.13 | 6.79 | 6.28 | 0.51 | 1.42 | 3.47741E-05 | 0.00054824 |
| 3303 | *HSPA1A* | ENSG00000204389.10 | 6.89 | 6.38 | 0.51 | 1.42 | 0.000103959 | 0.001199328 |
| 10142 | *AKAP9* | ENSG00000127914.19 | 6.50 | 5.99 | 0.50 | 1.42 | 0.000822032 | 0.005478113 |
| 128077 | *LIX1L* | ENSG00000271601.4 | 7.37 | 6.87 | 0.50 | 1.42 | 7.78007E-07 | 3.74266E-05 |
| 85236 | *H2BC12* | ENSG00000197903.8 | 5.22 | 4.71 | 0.50 | 1.42 | 0.000875707 | 0.005724516 |
| 9846 | *GAB2* | ENSG00000033327.13 | 5.82 | 5.32 | 0.50 | 1.42 | 7.87168E-06 | 0.000180376 |
| 10724 | *OGA* | ENSG00000198408.14 | 7.20 | 6.69 | 0.50 | 1.42 | 2.10471E-05 | 0.000374728 |
| 10438 | *C1D* | ENSG00000197223.12 | 4.73 | 4.23 | 0.50 | 1.42 | 0.000147587 | 0.001550123 |
| 3920 | *LAMP2* | ENSG00000005893.16 | 8.28 | 7.77 | 0.50 | 1.42 | 1.50604E-06 | 5.82618E-05 |
| 160897 | *GPR180* | ENSG00000152749.8 | 6.20 | 5.69 | 0.50 | 1.42 | 5.98269E-05 | 0.000814091 |
| 9764 | *KIAA0513* | ENSG00000135709.14 | 5.86 | 5.35 | 0.50 | 1.42 | 0.000321297 | 0.002732211 |
| 221491 | *SMIM29* | ENSG00000186577.14 | 3.90 | 3.40 | 0.50 | 1.42 | 0.000848664 | 0.005606334 |
| 8987 | *STBD1* | ENSG00000118804.9 | 6.52 | 6.01 | 0.50 | 1.42 | 1.83195E-05 | 0.000336545 |
| 55771 | *PRR11* | ENSG00000068489.13 | 6.80 | 6.30 | 0.50 | 1.41 | 0.000368922 | 0.003032395 |
| 10263 | *CDK2AP2* | ENSG00000167797.8 | 5.70 | 5.20 | 0.50 | 1.41 | 0.000255278 | 0.002333415 |
| 8440 | *NCK2* | ENSG00000071051.14 | 6.58 | 6.08 | 0.50 | 1.41 | 8.01785E-05 | 0.000997844 |
| 23413 | *NCS1* | ENSG00000107130.10 | 7.20 | 6.70 | 0.50 | 1.41 | 0.000126312 | 0.001379722 |
| 64763 | *ZNF574* | ENSG00000105732.13 | 4.98 | 4.49 | 0.50 | 1.41 | 0.000598371 | 0.004334433 |
| 26064 | *RAI14* | ENSG00000039560.14 | 7.59 | 7.09 | 0.50 | 1.41 | 1.03744E-05 | 0.000220551 |
| 29919 | *RMC1* | ENSG00000141452.10 | 5.27 | 4.77 | 0.50 | 1.41 | 0.000155988 | 0.001610192 |
| 5269 | *SERPINB6* | ENSG00000124570.21 | 6.42 | 5.92 | 0.50 | 1.41 | 1.60963E-05 | 0.000305037 |
| 9949 | *AMMECR1* | ENSG00000101935.12 | 4.77 | 4.27 | 0.50 | 1.41 | 0.000695419 | 0.004821034 |
| 54187 | *NANS* | ENSG00000095380.11 | 7.59 | 7.10 | 0.50 | 1.41 | 1.71336E-06 | 6.3184E-05 |
| 259266 | *ASPM* | ENSG00000066279.19 | 7.76 | 7.26 | 0.50 | 1.41 | 8.76183E-06 | 0.000194249 |
| 9453 | *GGPS1* | ENSG00000152904.11 | 4.89 | 4.39 | 0.50 | 1.41 | 0.001778318 | 0.009732056 |
| 25831 | *HECTD1* | ENSG00000092148.14 | 8.44 | 7.94 | 0.50 | 1.41 | 4.30358E-05 | 0.000644636 |
| 23231 | *SEL1L3* | ENSG00000091490.11 | 5.94 | 5.44 | 0.50 | 1.41 | 0.000140183 | 0.001494042 |
| 6747 | *SSR3* | ENSG00000114850.7 | 7.91 | 7.41 | 0.50 | 1.41 | 5.71096E-05 | 0.000784226 |
| 10695 | *CNPY3* | ENSG00000137161.18 | 6.40 | 5.91 | 0.50 | 1.41 | 1.81368E-06 | 6.57523E-05 |
| 85352 | *SHISAL1* | ENSG00000138944.8 | 4.78 | 4.28 | 0.50 | 1.41 | 0.000169805 | 0.0017002 |
| 9262 | *STK17B* | ENSG00000081320.11 | 6.72 | 6.22 | 0.50 | 1.41 | 2.79588E-05 | 0.000465743 |
| 10390 | *CEPT1* | ENSG00000134255.15 | 5.59 | 5.10 | 0.50 | 1.41 | 0.00034244 | 0.002868123 |
| 57720 | *GPR107* | ENSG00000148358.20 | 8.43 | 7.94 | 0.49 | 1.41 | 3.63108E-06 | 0.000103807 |
| 84790 | *TUBA1C* | ENSG00000167553.17 | 9.33 | 8.83 | 0.49 | 1.41 | 2.32191E-06 | 7.69305E-05 |
| 55635 | *DEPDC1* | ENSG00000024526.17 | 7.35 | 6.85 | 0.49 | 1.41 | 9.82617E-05 | 0.001156713 |
| 25948 | *KBTBD2* | ENSG00000170852.12 | 6.57 | 6.07 | 0.49 | 1.41 | 1.03378E-05 | 0.000220501 |
| 64764 | *CREB3L2* | ENSG00000182158.16 | 7.74 | 7.25 | 0.49 | 1.41 | 4.16302E-05 | 0.000627967 |
| 55972 | *SLC25A40* | ENSG00000075303.13 | 6.73 | 6.24 | 0.49 | 1.41 | 0.000727711 | 0.004989592 |
| 84858 | *ZNF503* | ENSG00000165655.17 | 6.47 | 5.98 | 0.49 | 1.41 | 8.28283E-06 | 0.00018643 |
| 9976 | *CLEC2B* | ENSG00000110852.5 | 5.80 | 5.31 | 0.49 | 1.41 | 0.000304211 | 0.002642304 |
| 79845 | *RNF122* | ENSG00000133874.2 | 4.56 | 4.07 | 0.49 | 1.41 | 0.001383023 | 0.008073822 |
| 137695 | *TMEM68* | ENSG00000167904.16 | 5.11 | 4.61 | 0.49 | 1.41 | 0.000813761 | 0.005433159 |
| 146330 | *FBXL16* | ENSG00000127585.12 | 5.22 | 4.73 | 0.49 | 1.41 | 8.66504E-05 | 0.001054773 |
| 201475 | *RAB12* | ENSG00000206418.5 | 6.37 | 5.88 | 0.49 | 1.41 | 8.83454E-06 | 0.000195265 |
| 51439 | *FAM8A1* | ENSG00000137414.7 | 6.24 | 5.75 | 0.49 | 1.41 | 2.52721E-05 | 0.000431523 |
| 2273 | *FHL1* | ENSG00000022267.19 | 5.73 | 5.23 | 0.49 | 1.41 | 0.000613247 | 0.004412377 |
| 7743 | *ZNF189* | ENSG00000136870.11 | 7.72 | 7.23 | 0.49 | 1.40 | 1.96292E-06 | 6.94566E-05 |
| 80314 | *EPC1* | ENSG00000120616.16 | 5.34 | 4.85 | 0.49 | 1.40 | 0.00028634 | 0.002525075 |
| 728841 | *NBPF8* | ENSG00000270231.4 | 4.61 | 4.12 | 0.49 | 1.40 | 0.00026333 | 0.002387141 |
| 3460 | *IFNGR2* | ENSG00000159128.15 | 5.99 | 5.50 | 0.49 | 1.40 | 4.96311E-05 | 0.000709441 |
| 81839 | *VANGL1* | ENSG00000173218.15 | 5.91 | 5.42 | 0.49 | 1.40 | 0.000541923 | 0.004032585 |
| 6560 | *SLC12A4* | ENSG00000124067.17 | 5.53 | 5.04 | 0.49 | 1.40 | 0.001664573 | 0.009309253 |
| 84892 | *POMGNT2* | ENSG00000144647.7 | 7.01 | 6.52 | 0.49 | 1.40 | 2.35371E-06 | 7.75832E-05 |
| 143098 | *MPP7* | ENSG00000150054.19 | 4.12 | 3.64 | 0.49 | 1.40 | 0.001686696 | 0.009390064 |
| 27230 | *SERP1* | ENSG00000120742.11 | 8.60 | 8.11 | 0.49 | 1.40 | 2.52664E-06 | 8.0858E-05 |
| 9751 | *SNPH* | ENSG00000101298.15 | 4.94 | 4.45 | 0.49 | 1.40 | 0.000318963 | 0.002723125 |
| 55252 | *ASXL2* | ENSG00000143970.18 | 7.57 | 7.08 | 0.49 | 1.40 | 5.77808E-06 | 0.000144059 |
| 115294 | *PCMTD1* | ENSG00000168300.14 | 6.39 | 5.91 | 0.49 | 1.40 | 4.15405E-06 | 0.000113721 |
| 4504 | *MT3* | ENSG00000087250.9 | 6.04 | 5.55 | 0.49 | 1.40 | 9.90844E-05 | 0.00116003 |
| 64946 | *CENPH* | ENSG00000153044.10 | 6.50 | 6.02 | 0.49 | 1.40 | 3.06453E-05 | 0.000499408 |
| 51430 | *SUCO* | ENSG00000094975.14 | 8.04 | 7.56 | 0.48 | 1.40 | 3.50769E-06 | 0.000101832 |
| 23585 | *TMEM50A* | ENSG00000183726.11 | 7.47 | 6.99 | 0.48 | 1.40 | 1.95915E-05 | 0.000353425 |
| 23467 | *NPTXR* | ENSG00000221890.6 | 6.16 | 5.68 | 0.48 | 1.40 | 9.25854E-06 | 0.000201102 |
| 9265 | *CYTH3* | ENSG00000008256.16 | 6.65 | 6.17 | 0.48 | 1.40 | 0.000184029 | 0.001807649 |
| 9777 | *TM9SF4* | ENSG00000101337.16 | 7.73 | 7.25 | 0.48 | 1.40 | 3.58592E-06 | 0.000103105 |
| 23170 | *TTLL12* | ENSG00000100304.13 | 5.94 | 5.46 | 0.48 | 1.40 | 0.000100296 | 0.001168739 |
| 291 | *SLC25A4* | ENSG00000151729.11 | 5.39 | 4.90 | 0.48 | 1.40 | 0.000193921 | 0.001881424 |
| 1317 | *SLC31A1* | ENSG00000136868.11 | 8.32 | 7.83 | 0.48 | 1.40 | 6.23222E-06 | 0.000152097 |
| 55275 | *VPS53* | ENSG00000141252.21 | 5.90 | 5.41 | 0.48 | 1.40 | 0.000419435 | 0.003334374 |
| 29802 | *VPREB3* | ENSG00000128218.8 | 3.79 | 3.31 | 0.48 | 1.40 | 0.000723583 | 0.004974925 |
| 64782 | *AEN* | ENSG00000181026.15 | 5.41 | 4.93 | 0.48 | 1.40 | 4.00228E-05 | 0.000608615 |
| 80219 | *COQ10B* | ENSG00000115520.9 | 4.78 | 4.30 | 0.48 | 1.40 | 0.000760547 | 0.005155852 |
| 57534 | *MIB1* | ENSG00000101752.12 | 7.51 | 7.02 | 0.48 | 1.40 | 1.08143E-05 | 0.000227967 |
| 2170 | *FABP3* | ENSG00000121769.8 | 6.64 | 6.16 | 0.48 | 1.40 | 0.00028936 | 0.002547218 |
| 23306 | *NEMP1* | ENSG00000166881.10 | 7.88 | 7.40 | 0.48 | 1.40 | 1.58051E-06 | 6.00553E-05 |
| 11273 | *ATXN2L* | ENSG00000168488.19 | 7.71 | 7.23 | 0.48 | 1.40 | 2.91937E-06 | 8.86982E-05 |
| 9276 | *COPB2* | ENSG00000184432.11 | 8.78 | 8.30 | 0.48 | 1.39 | 2.2651E-06 | 7.56296E-05 |
| 57128 | *LYRM4* | ENSG00000214113.11 | 4.12 | 3.64 | 0.48 | 1.39 | 0.001678959 | 0.009363134 |
| 3422 | *IDI1* | ENSG00000067064.12 | 7.59 | 7.11 | 0.48 | 1.39 | 2.67816E-05 | 0.000451647 |
| 80195 | *TMEM254* | ENSG00000133678.14 | 5.75 | 5.28 | 0.48 | 1.39 | 0.001573098 | 0.008929888 |

| | | | | | | | | |
|---|---|---|---|---|---|---|---|---|
| 29887 | *SNX10* | ENSG00000086300.16 | 7.13 | 6.65 | 0.48 | 1.39 | 5.13725E-06 | 0.000132336 |
| 167691 | *LCA5* | ENSG00000135338.14 | 4.67 | 4.19 | 0.48 | 1.39 | 0.000577621 | 0.004226966 |
| 30000 | *TNPO2* | ENSG00000105576.16 | 7.37 | 6.89 | 0.48 | 1.39 | 3.19843E-05 | 0.000513975 |
| 51447 | *IP6K2* | ENSG00000068745.15 | 5.85 | 5.37 | 0.48 | 1.39 | 5.54345E-05 | 0.000767543 |
| 8428 | *STK24* | ENSG00000102572.15 | 7.93 | 7.46 | 0.48 | 1.39 | 0.000388343 | 0.00314043 |
| 9394 | *HS6ST1* | ENSG00000136720.7 | 6.65 | 6.18 | 0.48 | 1.39 | 1.27095E-05 | 0.000256836 |
| 6645 | *SNTB2* | ENSG00000168807.17 | 5.99 | 5.52 | 0.48 | 1.39 | 4.1274E-05 | 0.000623848 |
| 4591 | *TRIM37* | ENSG00000108395.14 | 7.37 | 6.89 | 0.48 | 1.39 | 2.07563E-05 | 0.00037043 |
| 10020 | *GNE* | ENSG00000159921.20 | 6.99 | 6.51 | 0.47 | 1.39 | 5.44219E-06 | 0.000137281 |
| 51171 | *HSD17B14* | ENSG00000087076.9 | 3.83 | 3.35 | 0.47 | 1.39 | 0.000712719 | 0.004915994 |
| 123811 | *CEP20* | ENSG00000133393.13 | 6.48 | 6.01 | 0.47 | 1.39 | 0.000379313 | 0.003085431 |
| 891 | *CCNB1* | ENSG00000134057.15 | 8.61 | 8.13 | 0.47 | 1.39 | 1.46801E-06 | 5.76145E-05 |
| 64771 | *ILRUN* | ENSG00000196821.10 | 7.75 | 7.28 | 0.47 | 1.39 | 0.000179962 | 0.001781695 |
| 56180 | *MOSPD1* | ENSG00000101928.13 | 5.07 | 4.60 | 0.47 | 1.39 | 0.000956701 | 0.006076651 |
| 220965 | *FAM13C* | ENSG00000148541.13 | 4.51 | 4.04 | 0.47 | 1.39 | 0.000100239 | 0.001168739 |
| 23232 | *TBC1D12* | ENSG00000108239.9 | 4.57 | 4.09 | 0.47 | 1.39 | 0.000785459 | 0.005283417 |
| 64770 | *CCDC14* | ENSG00000175455.16 | 5.92 | 5.45 | 0.47 | 1.39 | 0.001640133 | 0.009202527 |
| 51533 | *PHF7* | ENSG00000010318.22 | 3.64 | 3.17 | 0.47 | 1.39 | 0.001738007 | 0.009579781 |
| 51299 | *NRN1* | ENSG00000124785.9 | 4.59 | 4.11 | 0.47 | 1.39 | 0.000975084 | 0.006162118 |
| 55276 | *PGM2* | ENSG00000169299.14 | 6.07 | 5.60 | 0.47 | 1.39 | 0.001025206 | 0.006406045 |
| 27131 | *SNX5* | ENSG00000089006.17 | 7.33 | 6.85 | 0.47 | 1.39 | 7.19498E-05 | 0.000926948 |
| 1603 | *DAD1* | ENSG00000129562.11 | 7.27 | 6.80 | 0.47 | 1.39 | 0.000166377 | 0.001683847 |
| 154810 | *AMOTL1* | ENSG00000166025.18 | 7.45 | 6.98 | 0.47 | 1.39 | 1.50484E-05 | 0.000290684 |
| 961 | *CD47* | ENSG00000196776.17 | 7.67 | 7.20 | 0.47 | 1.39 | 2.49269E-06 | 8.02849E-05 |
| 1407 | *CRY1* | ENSG00000008405.12 | 6.53 | 6.06 | 0.47 | 1.39 | 0.000876369 | 0.005726345 |
| 1508 | *CTSB* | ENSG00000164733.22 | 8.69 | 8.22 | 0.47 | 1.39 | 4.85594E-06 | 0.000127405 |
| 30001 | *ERO1A* | ENSG00000197930.13 | 6.96 | 6.49 | 0.47 | 1.39 | 6.17333E-05 | 0.000832484 |
| 121536 | *AEBP2* | ENSG00000139154.16 | 8.36 | 7.89 | 0.47 | 1.39 | 4.91971E-06 | 0.000127972 |
| 23074 | *UHRF1BP1L* | ENSG00000111647.13 | 6.61 | 6.14 | 0.47 | 1.39 | 2.83184E-05 | 0.000468611 |
| 6625 | *SNRNP70* | ENSG00000104852.15 | 7.34 | 6.87 | 0.47 | 1.39 | 2.27527E-05 | 0.000396165 |
| 9282 | *MED14* | ENSG00000180182.11 | 6.65 | 6.18 | 0.47 | 1.39 | 0.001661816 | 0.009299849 |
| 79586 | *CHPF* | ENSG00000123989.15 | 5.68 | 5.21 | 0.47 | 1.39 | 7.96775E-05 | 0.000993255 |
| 84272 | *YIPF4* | ENSG00000119820.11 | 6.78 | 6.32 | 0.47 | 1.39 | 0.000531379 | 0.003971605 |
| 6515 | *SLC2A3* | ENSG00000059804.16 | 5.66 | 5.19 | 0.47 | 1.38 | 8.06431E-05 | 0.001001964 |
| 9552 | *SPAG7* | ENSG00000091640.8 | 5.58 | 5.11 | 0.47 | 1.38 | 0.000214183 | 0.002028816 |
| 23558 | *WBP2* | ENSG00000132471.12 | 7.58 | 7.11 | 0.47 | 1.38 | 6.82139E-05 | 0.000898089 |
| 4792 | *NFKBIA* | ENSG00000100906.11 | 6.00 | 5.53 | 0.47 | 1.38 | 7.45584E-05 | 0.000948346 |
| 9686 | *VGLL4* | ENSG00000144560.16 | 6.99 | 6.53 | 0.47 | 1.38 | 3.12367E-05 | 0.000505752 |
| 7430 | *EZR* | ENSG00000092820.19 | 6.83 | 6.36 | 0.47 | 1.38 | 4.24665E-06 | 0.000114843 |
| 8407 | *TAGLN2* | ENSG00000158710.15 | 8.35 | 7.88 | 0.47 | 1.38 | 6.39529E-06 | 0.000155068 |
| 118813 | *ZFYVE27* | ENSG00000155256.18 | 5.67 | 5.21 | 0.46 | 1.38 | 7.18444E-05 | 0.000926948 |
| 1955 | *MEGF9* | ENSG00000106780.9 | 6.87 | 6.41 | 0.46 | 1.38 | 1.45523E-05 | 0.000284166 |
| 23052 | *ENDOD1* | ENSG00000149218.5 | 6.24 | 5.77 | 0.46 | 1.38 | 0.000234808 | 0.002179493 |
| 80347 | *COASY* | ENSG00000068120.15 | 7.05 | 6.58 | 0.46 | 1.38 | 2.20376E-05 | 0.000388059 |
| 54504 | *CPVL* | ENSG00000106066.15 | 6.45 | 5.99 | 0.46 | 1.38 | 5.15911E-06 | 0.000132591 |
| 1742 | *DLG4* | ENSG00000132535.22 | 4.49 | 4.02 | 0.46 | 1.38 | 0.001442083 | 0.008335877 |
| 56900 | *TMEM167B* | ENSG00000215717.7 | 6.85 | 6.39 | 0.46 | 1.38 | 4.27964E-05 | 0.00064169 |
| 6520 | *SLC3A2* | ENSG00000168003.18 | 7.36 | 6.90 | 0.46 | 1.38 | 5.67961E-05 | 0.000781349 |
| 283635 | *FAM177A1* | ENSG00000151327.13 | 5.12 | 4.66 | 0.46 | 1.38 | 0.00027672 | 0.002466322 |
| 80146 | *UXS1* | ENSG00000115652.15 | 6.09 | 5.63 | 0.46 | 1.38 | 0.00012171 | 0.001348678 |
| 23042 | *PDXDC1* | ENSG00000179889.20 | 7.90 | 7.44 | 0.46 | 1.38 | 8.18918E-06 | 0.000185387 |
| 25957 | *PNISR* | ENSG00000132424.17 | 6.15 | 5.69 | 0.46 | 1.38 | 0.000103141 | 0.00119356 |
| 25822 | *DNAJB5* | ENSG00000137094.16 | 6.08 | 5.62 | 0.46 | 1.38 | 4.97044E-05 | 0.000709813 |
| 5716 | *PSMD10* | ENSG00000101843.19 | 6.85 | 6.39 | 0.46 | 1.38 | 0.000141631 | 0.001501223 |
| 85459 | *CEP295* | ENSG00000166004.15 | 4.94 | 4.48 | 0.46 | 1.38 | 0.000382468 | 0.003102955 |
| 51114 | *ZDHHC9* | ENSG00000188706.13 | 4.99 | 4.53 | 0.46 | 1.38 | 0.000262273 | 0.002381403 |
| 1654 | *DDX3X* | ENSG00000215301.11 | 9.43 | 8.97 | 0.46 | 1.38 | 1.19782E-05 | 0.000246275 |
| 9643 | *MORF4L2* | ENSG00000123562.18 | 9.01 | 8.55 | 0.46 | 1.38 | 1.1625E-05 | 0.000240994 |
| 3105 | *HLA-A* | ENSG00000206503.13 | 7.82 | 7.37 | 0.46 | 1.38 | 5.02747E-06 | 0.000130099 |
| 57099 | *AVEN* | ENSG00000169857.9 | 4.18 | 3.72 | 0.46 | 1.37 | 0.00158168 | 0.008963362 |
| 93058 | *COQ10A* | ENSG00000135469.14 | 4.97 | 4.51 | 0.46 | 1.37 | 0.00064407 | 0.004561986 |
| 6609 | *SMPD1* | ENSG00000166311.10 | 5.74 | 5.28 | 0.46 | 1.37 | 0.000593073 | 0.004304577 |
| 900 | *CCNG1* | ENSG00000113328.19 | 7.17 | 6.72 | 0.46 | 1.37 | 4.89411E-06 | 0.000127527 |
| 51622 | *CCZ1* | ENSG00000122674.12 | 5.84 | 5.38 | 0.46 | 1.37 | 0.000469341 | 0.003621769 |
| 55279 | *ZNF654* | ENSG00000175105.8 | 5.16 | 4.70 | 0.46 | 1.37 | 0.001075388 | 0.006649683 |
| 3858 | *KRT10* | ENSG00000186395.9 | 4.93 | 4.47 | 0.46 | 1.37 | 6.27089E-05 | 0.000842988 |
| 8013 | *NR4A3* | ENSG00000119508.18 | 7.41 | 6.95 | 0.46 | 1.37 | 4.00202E-06 | 0.000112064 |
| 163859 | *SDE2* | ENSG00000143751.10 | 6.03 | 5.58 | 0.46 | 1.37 | 0.000372237 | 0.003052956 |
| 734 | *OSGIN2* | ENSG00000164823.11 | 5.73 | 5.27 | 0.46 | 1.37 | 0.000120695 | 0.00134585 |
| 9439 | *MED23* | ENSG00000112282.18 | 6.87 | 6.42 | 0.46 | 1.37 | 4.45159E-05 | 0.000659565 |
| 49854 | *ZBTB21* | ENSG00000173276.14 | 6.33 | 5.88 | 0.46 | 1.37 | 8.96217E-05 | 0.001078694 |
| 337867 | *UBAC2* | ENSG00000134882.16 | 7.49 | 7.03 | 0.46 | 1.37 | 1.33461E-05 | 0.000266727 |
| 9522 | *SCAMP1* | ENSG00000085365.18 | 7.66 | 7.20 | 0.46 | 1.37 | 2.09804E-06 | 7.20414E-05 |
| 7525 | *YES1* | ENSG00000176105.14 | 7.67 | 7.21 | 0.46 | 1.37 | 2.89283E-06 | 8.82488E-05 |
| 30834 | *POLR1H* | ENSG00000066379.15 | 3.99 | 3.53 | 0.45 | 1.37 | 0.000836143 | 0.005540693 |
| 2050 | *EPHB4* | ENSG00000196411.10 | 8.24 | 7.79 | 0.45 | 1.37 | 5.95227E-06 | 0.000147179 |
| 80267 | *EDEM3* | ENSG00000116406.20 | 7.15 | 6.70 | 0.45 | 1.37 | 3.17415E-05 | 0.00051062 |
| 991 | *CDC20* | ENSG00000117399.14 | 7.37 | 6.92 | 0.45 | 1.37 | 3.85117E-05 | 0.000589819 |
| 50848 | *F11R* | ENSG00000158769.18 | 7.74 | 7.29 | 0.45 | 1.37 | 0.000140356 | 0.001494042 |
| 3029 | *HAGH* | ENSG00000063854.13 | 5.35 | 4.89 | 0.45 | 1.37 | 0.000309532 | 0.002670744 |
| 55633 | *TBC1D22B* | ENSG00000065491.8 | 5.37 | 4.91 | 0.45 | 1.37 | 9.79925E-05 | 0.00115445 |
| 23471 | *TRAM1* | ENSG00000067167.8 | 9.10 | 8.65 | 0.45 | 1.37 | 3.47386E-06 | 0.000101394 |
| 201562 | *HACD2* | ENSG00000206527.10 | 7.45 | 7.00 | 0.45 | 1.37 | 5.37649E-06 | 0.000135851 |
| 7006 | *TEC* | ENSG00000135605.13 | 5.91 | 5.46 | 0.45 | 1.37 | 0.001126878 | 0.006897761 |
| 23411 | *SIRT1* | ENSG00000096717.12 | 5.93 | 5.48 | 0.45 | 1.37 | 0.000148127 | 0.001553329 |
| 55744 | *COA1* | ENSG00000106603.20 | 7.64 | 7.19 | 0.45 | 1.37 | 4.24483E-06 | 0.000114843 |
| 64755 | *RUSF1* | ENSG00000140688.17 | 6.63 | 6.18 | 0.45 | 1.37 | 3.7395E-05 | 0.000576837 |
| 53834 | *FGFRL1* | ENSG00000127418.15 | 7.59 | 7.14 | 0.45 | 1.37 | 9.14631E-06 | 0.000199241 |
| 57799 | *RAB40C* | ENSG00000197562.10 | 4.27 | 3.83 | 0.45 | 1.37 | 0.00077008 | 0.005203945 |

| | | | | | | | | |
|---|---|---|---|---|---|---|---|---|
| 65979 | *PHACTR4* | ENSG00000204138.13 | 6.63 | 6.18 | 0.45 | 1.37 | 5.91081E-05 | 0.000806503 |
| 27238 | *GPKOW* | ENSG00000068394.11 | 5.44 | 4.99 | 0.45 | 1.37 | 0.000852057 | 0.005621332 |
| 10681 | *GNB5* | ENSG00000069966.19 | 6.35 | 5.90 | 0.45 | 1.37 | 3.39932E-05 | 0.000539898 |
| 56907 | *SPIRE1* | ENSG00000134278.16 | 5.98 | 5.53 | 0.45 | 1.37 | 0.000915732 | 0.005891223 |
| 100526842 | *RPL17-C18orf* | ENSG00000215472.10 | 5.65 | 5.20 | 0.45 | 1.37 | 0.00038577 | 0.003124677 |
| 25983 | *NGDN* | ENSG00000129460.16 | 5.38 | 4.93 | 0.45 | 1.37 | 0.000143367 | 0.001515347 |
| 55033 | *FKBP14* | ENSG00000106080.11 | 6.28 | 5.83 | 0.45 | 1.37 | 0.000712614 | 0.004915994 |
| 9875 | *URB1* | ENSG00000142207.7 | 6.77 | 6.32 | 0.45 | 1.37 | 3.77366E-05 | 0.000580316 |
| 7832 | *BTG2* | ENSG00000159388.6 | 4.03 | 3.58 | 0.45 | 1.37 | 0.001671284 | 0.00932849 |
| 25840 | *METTL7A* | ENSG00000185432.12 | 7.76 | 7.31 | 0.45 | 1.37 | 7.11896E-05 | 0.000921456 |
| 65125 | *WNK1* | ENSG00000060237.19 | 9.54 | 9.10 | 0.45 | 1.36 | 4.68129E-05 | 0.000683478 |
| 5573 | *PRKAR1A* | ENSG00000108946.16 | 9.97 | 9.52 | 0.45 | 1.36 | 4.08523E-06 | 0.000113337 |
| 1200 | *TPP1* | ENSG00000166340.18 | 6.62 | 6.17 | 0.45 | 1.36 | 9.79198E-05 | 0.00115445 |
| 81691 | *REXO5* | ENSG00000005189.20 | 4.38 | 3.94 | 0.45 | 1.36 | 0.000692726 | 0.004811259 |
| 280636 | *SELENOH* | ENSG00000211450.11 | 5.07 | 4.62 | 0.45 | 1.36 | 0.000487488 | 0.003721229 |
| 2590 | *GALNT2* | ENSG00000143641.10 | 8.05 | 7.61 | 0.45 | 1.36 | 3.85836E-06 | 0.00010865 |
| 57157 | *PHTF2* | ENSG00000006576.16 | 6.81 | 6.37 | 0.45 | 1.36 | 2.0179E-05 | 0.000363149 |
| 9263 | *STK17A* | ENSG00000164543.7 | 5.94 | 5.49 | 0.45 | 1.36 | 2.55756E-05 | 0.000435715 |
| 10437 | *IFI30* | ENSG00000216490.4 | 4.91 | 4.47 | 0.45 | 1.36 | 0.001221768 | 0.00734088 |
| 59345 | *GNB4* | ENSG00000114450.11 | 5.22 | 4.78 | 0.45 | 1.36 | 0.000953902 | 0.006069147 |
| 204851 | *HIPK1* | ENSG00000163349.22 | 7.75 | 7.30 | 0.45 | 1.36 | 1.93086E-05 | 0.00034874 |
| 128869 | *PIGU* | ENSG00000101464.11 | 4.99 | 4.55 | 0.44 | 1.36 | 0.000665241 | 0.004668056 |
| 4214 | *MAP3K1* | ENSG00000095015.6 | 4.25 | 3.80 | 0.44 | 1.36 | 0.00079749 | 0.005343541 |
| 3485 | *IGFBP2* | ENSG00000115457.10 | 8.30 | 7.85 | 0.44 | 1.36 | 3.85512E-05 | 0.000589822 |
| 23432 | *GPR161* | ENSG00000143147.15 | 5.60 | 5.16 | 0.44 | 1.36 | 0.00052889 | 0.003957182 |
| 8349 | *H2BC21* | ENSG00000184678.11 | 6.51 | 6.06 | 0.44 | 1.36 | 2.8803E-05 | 0.000474597 |
| 7184 | *HSP90B1* | ENSG00000166598.16 | 10.75 | 10.30 | 0.44 | 1.36 | 7.38485E-07 | 3.6104E-05 |
| 9055 | *PRC1* | ENSG00000198901.14 | 8.80 | 8.36 | 0.44 | 1.36 | 2.60732E-06 | 8.23678E-05 |
| 6046 | *BRD2* | ENSG00000204256.14 | 8.59 | 8.14 | 0.44 | 1.36 | 0.000161514 | 0.001646846 |
| 5527 | *PPP2R5C* | ENSG00000078304.19 | 8.01 | 7.56 | 0.44 | 1.36 | 5.34168E-06 | 0.000135851 |
| 6541 | *SLC7A1* | ENSG00000139514.13 | 7.01 | 6.57 | 0.44 | 1.36 | 5.15046E-05 | 0.000727218 |
| 10079 | *ATP9A* | ENSG00000054793.14 | 7.32 | 6.87 | 0.44 | 1.36 | 2.46978E-05 | 0.000423162 |
| 201595 | *STT3B* | ENSG00000163527.10 | 8.59 | 8.14 | 0.44 | 1.36 | 2.04598E-05 | 0.000366445 |
| 96459 | *FNIP1* | ENSG00000217128.13 | 6.02 | 5.57 | 0.44 | 1.36 | 7.12329E-05 | 0.000921456 |
| 79666 | *PLEKHF2* | ENSG00000175895.4 | 4.79 | 4.35 | 0.44 | 1.36 | 0.000785767 | 0.005283417 |
| 51594 | *NBAS* | ENSG00000151779.13 | 6.03 | 5.59 | 0.44 | 1.36 | 0.000696177 | 0.004824062 |
| 91452 | *ACBD5* | ENSG00000107897.20 | 6.84 | 6.40 | 0.44 | 1.36 | 0.00022393 | 0.002099288 |
| 25978 | *CHMP2B* | ENSG00000083937.10 | 6.52 | 6.08 | 0.44 | 1.36 | 6.07792E-05 | 0.000823317 |
| 51109 | *RDH11* | ENSG00000072042.13 | 7.35 | 6.91 | 0.44 | 1.36 | 1.19231E-05 | 0.000246153 |
| 23024 | *PDZRN3* | ENSG00000121440.15 | 8.32 | 7.88 | 0.44 | 1.36 | 8.87623E-06 | 0.000195629 |
| 9278 | *ZBTB22* | ENSG00000236104.3 | 4.85 | 4.41 | 0.44 | 1.36 | 0.000291209 | 0.002557495 |
| 4179 | *CD46* | ENSG00000117335.20 | 7.90 | 7.46 | 0.44 | 1.36 | 1.89148E-05 | 0.000344111 |
| 339287 | *MSL1* | ENSG00000188895.12 | 6.96 | 6.53 | 0.44 | 1.36 | 0.000133332 | 0.001433505 |
| 51696 | *HECA* | ENSG00000112406.5 | 5.93 | 5.49 | 0.44 | 1.36 | 6.32149E-05 | 0.00084638 |
| 56650 | *CLDND1* | ENSG00000080822.17 | 7.43 | 6.99 | 0.44 | 1.35 | 4.14675E-05 | 0.000626142 |
| 4245 | *MGAT1* | ENSG00000131446.17 | 6.83 | 6.39 | 0.44 | 1.35 | 0.000290555 | 0.002553692 |
| 54764 | *ZRANB1* | ENSG00000019995.6 | 6.89 | 6.46 | 0.44 | 1.35 | 5.84456E-06 | 0.000145233 |
| 55105 | *GPATCH2* | ENSG00000092978.11 | 5.68 | 5.24 | 0.44 | 1.35 | 0.000236655 | 0.002192567 |
| 2717 | *GLA* | ENSG00000102393.14 | 5.87 | 5.43 | 0.44 | 1.35 | 4.75173E-05 | 0.000690501 |
| 9513 | *FXR2* | ENSG00000129245.12 | 6.99 | 6.56 | 0.44 | 1.35 | 8.4242E-06 | 0.000188433 |
| 54965 | *PIGX* | ENSG00000163964.18 | 5.95 | 5.52 | 0.44 | 1.35 | 3.4113E-05 | 0.000540657 |
| 54443 | *ANLN* | ENSG00000011426.11 | 8.31 | 7.87 | 0.44 | 1.35 | 3.35419E-05 | 0.000534427 |
| 137964 | *GPAT4* | ENSG00000158669.11 | 7.52 | 7.09 | 0.44 | 1.35 | 0.000776426 | 0.00523747 |
| 25976 | *TIPARP* | ENSG00000163659.13 | 4.58 | 4.14 | 0.44 | 1.35 | 0.001323678 | 0.007797129 |
| 3598 | *IL13RA2* | ENSG00000123496.8 | 7.17 | 6.73 | 0.44 | 1.35 | 1.49363E-05 | 0.000289636 |
| 2101 | *ESRRA* | ENSG00000173153.17 | 5.39 | 4.96 | 0.44 | 1.35 | 0.000551863 | 0.004084275 |
| 23463 | *ICMT* | ENSG00000116237.16 | 7.64 | 7.21 | 0.44 | 1.35 | 4.47054E-06 | 0.000119586 |
| 1060 | *CENPC* | ENSG00000145241.11 | 5.23 | 4.80 | 0.44 | 1.35 | 0.000528869 | 0.003957182 |
| 58478 | *ENOPH1* | ENSG00000145293.17 | 6.80 | 6.37 | 0.44 | 1.35 | 1.80901E-05 | 0.000333966 |
| 10015 | *PDCD6IP* | ENSG00000170248.15 | 7.86 | 7.43 | 0.43 | 1.35 | 0.000451737 | 0.003525804 |
| 11345 | *GABARAPL2* | ENSG00000034713.8 | 6.91 | 6.47 | 0.43 | 1.35 | 1.07661E-05 | 0.000227527 |
| 4860 | *PNP* | ENSG00000198805.12 | 5.58 | 5.14 | 0.43 | 1.35 | 0.000448335 | 0.003506546 |
| 372 | *ARCN1* | ENSG00000095139.15 | 8.54 | 8.11 | 0.43 | 1.35 | 1.02804E-05 | 0.000219798 |
| 286451 | *YIPF6* | ENSG00000181704.12 | 6.41 | 5.98 | 0.43 | 1.35 | 0.000128875 | 0.001400646 |
| 9748 | *SLK* | ENSG00000065613.15 | 7.49 | 7.06 | 0.43 | 1.35 | 0.000151778 | 0.001579771 |
| 51399 | *TRAPPC4* | ENSG00000196655.12 | 5.80 | 5.36 | 0.43 | 1.35 | 0.001626388 | 0.009152778 |
| 50808 | *AK3* | ENSG00000147853.17 | 6.30 | 5.86 | 0.43 | 1.35 | 0.000266761 | 0.002406142 |
| 6617 | *SNAPC1* | ENSG00000023608.5 | 4.75 | 4.32 | 0.43 | 1.35 | 0.000917815 | 0.005897038 |
| 3939 | *LDHA* | ENSG00000134333.14 | 8.75 | 8.32 | 0.43 | 1.35 | 0.000426208 | 0.003370364 |
| 5264 | *PHYH* | ENSG00000107537.14 | 5.82 | 5.38 | 0.43 | 1.35 | 3.55942E-05 | 0.000554805 |
| 10063 | *COX17* | ENSG00000138495.7 | 5.96 | 5.53 | 0.43 | 1.35 | 0.000600974 | 0.004346996 |
| 23484 | *LEPROTL1* | ENSG00000104660.19 | 5.64 | 5.21 | 0.43 | 1.35 | 6.67712E-05 | 0.000883746 |
| 7086 | *TKT* | ENSG00000163931.17 | 9.14 | 8.71 | 0.43 | 1.35 | 1.45249E-05 | 0.000284166 |
| 11040 | *PIM2* | ENSG00000102096.9 | 4.80 | 4.37 | 0.43 | 1.35 | 0.000234987 | 0.002179801 |
| 387882 | *C12orf75* | ENSG00000235162.9 | 5.08 | 4.65 | 0.43 | 1.35 | 0.000877817 | 0.005730821 |
| 10509 | *SEMA4B* | ENSG00000185033.15 | 6.37 | 5.94 | 0.43 | 1.35 | 5.52726E-05 | 0.000766008 |
| 6045 | *RNF2* | ENSG00000121481.11 | 5.00 | 4.57 | 0.43 | 1.35 | 0.001748049 | 0.00961395 |
| 5110 | *PCMT1* | ENSG00000120265.19 | 7.34 | 6.91 | 0.43 | 1.35 | 4.23356E-06 | 0.000114843 |
| 167153 | *TENT2* | ENSG00000164329.14 | 6.55 | 6.12 | 0.43 | 1.35 | 0.000297883 | 0.002600887 |
| 3799 | *KIF5B* | ENSG00000170759.11 | 8.94 | 8.52 | 0.43 | 1.35 | 8.48033E-06 | 0.000188845 |
| 8826 | *IQGAP1* | ENSG00000140575.13 | 9.26 | 8.83 | 0.43 | 1.35 | 5.37521E-05 | 0.000751178 |
| 56994 | *CHPT1* | ENSG00000111666.11 | 5.81 | 5.38 | 0.43 | 1.34 | 3.90502E-05 | 0.000596847 |
| 7296 | *TXNRD1* | ENSG00000198431.17 | 9.37 | 8.94 | 0.43 | 1.34 | 4.06992E-06 | 0.000113331 |
| 113402 | *SFT2D1* | ENSG00000198818.10 | 5.43 | 5.00 | 0.43 | 1.34 | 8.89127E-05 | 0.001075335 |
| 133619 | *PRRC1* | ENSG00000164244.21 | 7.15 | 6.73 | 0.43 | 1.34 | 0.000423327 | 0.003358197 |
| 84893 | *FBH1* | ENSG00000134452.20 | 6.56 | 6.14 | 0.43 | 1.34 | 0.000574338 | 0.004209099 |
| 5786 | *PTPRA* | ENSG00000132670.21 | 6.78 | 6.35 | 0.43 | 1.34 | 5.81466E-05 | 0.000797637 |
| 9470 | *EIF4E2* | ENSG00000135930.15 | 5.94 | 5.51 | 0.43 | 1.34 | 0.000313522 | 0.002685246 |
| 7385 | *UQCRC2* | ENSG00000140740.11 | 7.39 | 6.96 | 0.43 | 1.34 | 5.21395E-05 | 0.00073411 |

| | | | | | | | | |
|---|---|---|---|---|---|---|---|---|
| 5261 | *PHKG2* | ENSG00000156873.16 | 4.84 | 4.42 | 0.43 | 1.34 | 0.000488548 | 0.00372592 |
| 25912 | *C1orf43* | ENSG00000143612.21 | 8.46 | 8.03 | 0.43 | 1.34 | 2.19634E-05 | 0.000387366 |
| 85450 | *ITPRIP* | ENSG00000148841.17 | 4.77 | 4.34 | 0.43 | 1.34 | 0.000900054 | 0.00582783 |
| 23158 | *TBC1D9* | ENSG00000109436.8 | 6.50 | 6.07 | 0.43 | 1.34 | 0.00019821 | 0.001915609 |
| 3703 | *STT3A* | ENSG00000134910.14 | 8.37 | 7.94 | 0.42 | 1.34 | 8.88461E-05 | 0.001075335 |
| 51124 | *IER3IP1* | ENSG00000134049.6 | 6.33 | 5.90 | 0.42 | 1.34 | 0.000177367 | 0.001760652 |
| 9656 | *MDC1* | ENSG00000137337.16 | 6.62 | 6.19 | 0.42 | 1.34 | 0.00129158 | 0.007668246 |
| 22872 | *SEC31A* | ENSG00000138674.17 | 7.97 | 7.54 | 0.42 | 1.34 | 0.000320622 | 0.00273111 |
| 5516 | *PPP2CB* | ENSG00000104695.13 | 6.38 | 5.95 | 0.42 | 1.34 | 0.000107395 | 0.001230449 |
| 2319 | *FLOT2* | ENSG00000132589.16 | 6.94 | 6.52 | 0.42 | 1.34 | 0.000134434 | 0.001443291 |
| 23321 | *TRIM2* | ENSG00000109654.16 | 6.72 | 6.30 | 0.42 | 1.34 | 0.000246889 | 0.002274747 |
| 10615 | *SPAG5* | ENSG00000076382.17 | 7.47 | 7.04 | 0.42 | 1.34 | 0.000148202 | 0.001553329 |
| 51272 | *BET1L* | ENSG00000177951.18 | 5.10 | 4.68 | 0.42 | 1.34 | 0.00077858 | 0.005247289 |
| 27333 | *GOLIM4* | ENSG00000173905.9 | 7.06 | 6.64 | 0.42 | 1.34 | 1.52957E-05 | 0.000293571 |
| 90231 | *KIAA2013* | ENSG00000116685.16 | 6.21 | 5.79 | 0.42 | 1.34 | 0.000114902 | 0.001295638 |
| 1183 | *CLCN4* | ENSG00000073464.13 | 6.87 | 6.45 | 0.42 | 1.34 | 4.91499E-05 | 0.000705924 |
| 23353 | *SUN1* | ENSG00000164828.18 | 7.40 | 6.98 | 0.42 | 1.34 | 0.00041614 | 0.003324024 |
| 1198 | *CLK3* | ENSG00000179335.20 | 6.09 | 5.67 | 0.42 | 1.34 | 0.000609375 | 0.004390834 |
| 11072 | *DUSP14* | ENSG00000276023.5 | 5.87 | 5.45 | 0.42 | 1.34 | 4.52671E-05 | 0.000666092 |
| 55857 | *KIZ* | ENSG00000088970.16 | 5.64 | 5.22 | 0.42 | 1.34 | 0.000226077 | 0.002115455 |
| 58505 | *OSTC* | ENSG00000198856.13 | 6.96 | 6.54 | 0.42 | 1.34 | 0.00015889 | 0.001630065 |
| 3638 | *INSIG1* | ENSG00000186480.13 | 7.90 | 7.48 | 0.42 | 1.34 | 5.24298E-05 | 0.000736126 |
| 5045 | *FURIN* | ENSG00000140564.13 | 6.39 | 5.97 | 0.42 | 1.34 | 0.000168175 | 0.001690645 |
| 10613 | *ERLIN1* | ENSG00000107566.14 | 6.87 | 6.45 | 0.42 | 1.34 | 6.93556E-05 | 0.000905181 |
| 404203 | *SPINK6* | ENSG00000178172.7 | 9.89 | 9.47 | 0.42 | 1.34 | 0.000183623 | 0.001806031 |
| 81533 | *ITFG1* | ENSG00000129636.13 | 7.74 | 7.32 | 0.42 | 1.34 | 9.45812E-06 | 0.000204191 |
| 84448 | *ABLIM2* | ENSG00000163995.21 | 5.43 | 5.01 | 0.42 | 1.34 | 0.000302494 | 0.002633123 |
| 57631 | *LRCH2* | ENSG00000130224.15 | 5.51 | 5.10 | 0.42 | 1.33 | 0.000967465 | 0.006124283 |
| 11101 | *ATE1* | ENSG00000107669.19 | 6.19 | 5.78 | 0.41 | 1.33 | 0.000228612 | 0.002129882 |
| 780 | *DDR1* | ENSG00000204580.14 | 9.26 | 8.85 | 0.41 | 1.33 | 4.25385E-05 | 0.0006391 |
| 9585 | *KIF20B* | ENSG00000138182.15 | 6.97 | 6.56 | 0.41 | 1.33 | 0.000200772 | 0.001935108 |
| 21 | *ABCA3* | ENSG00000167972.14 | 6.34 | 5.92 | 0.41 | 1.33 | 4.48613E-05 | 0.000663338 |
| 57459 | *GATAD2B* | ENSG00000143614.10 | 7.09 | 6.68 | 0.41 | 1.33 | 1.39398E-05 | 0.000277116 |
| 90488 | *TMEM263* | ENSG00000151135.10 | 7.53 | 7.12 | 0.41 | 1.33 | 8.8156E-06 | 0.000195152 |
| 143384 | *CACUL1* | ENSG00000151893.15 | 7.18 | 6.77 | 0.41 | 1.33 | 0.000580478 | 0.004239605 |
| 2770 | *GNAI1* | ENSG00000127955.17 | 6.72 | 6.31 | 0.41 | 1.33 | 1.60125E-05 | 0.000304603 |
| 5168 | *ENPP2* | ENSG00000136960.13 | 9.17 | 8.76 | 0.41 | 1.33 | 1.73016E-05 | 0.000322985 |
| 5528 | *PPP2R5D* | ENSG00000112640.16 | 7.06 | 6.65 | 0.41 | 1.33 | 1.52839E-05 | 0.000293571 |
| 401548 | *SNX30* | ENSG00000148158.17 | 6.44 | 6.03 | 0.41 | 1.33 | 3.74417E-05 | 0.000576964 |
| 138050 | *HGSNAT* | ENSG00000165102.15 | 6.40 | 5.99 | 0.41 | 1.33 | 0.000102368 | 0.00118828 |
| 6892 | *TAPBP* | ENSG00000231925.13 | 7.65 | 7.24 | 0.41 | 1.33 | 3.76844E-05 | 0.000580108 |
| 79101 | *TAF1D* | ENSG00000166012.17 | 5.52 | 5.11 | 0.41 | 1.33 | 0.000664401 | 0.004668056 |
| 2288 | *FKBP4* | ENSG00000004478.8 | 8.35 | 7.94 | 0.41 | 1.33 | 2.62858E-05 | 0.000445789 |
| 89958 | *SAPCD2* | ENSG00000186193.9 | 5.67 | 5.26 | 0.41 | 1.33 | 8.54491E-05 | 0.001044385 |
| 5290 | *PIK3CA* | ENSG00000121879.6 | 5.97 | 5.56 | 0.41 | 1.33 | 0.000269114 | 0.002417198 |
| 64951 | *MRPS24* | ENSG00000062582.14 | 6.92 | 6.51 | 0.41 | 1.33 | 2.26777E-05 | 0.000395778 |
| 54918 | *CMTM6* | ENSG00000091317.8 | 7.87 | 7.47 | 0.41 | 1.33 | 1.4902E-05 | 0.000289346 |
| 112 | *ADCY6* | ENSG00000174233.12 | 6.75 | 6.34 | 0.41 | 1.33 | 0.000157059 | 0.001615699 |
| 6868 | *ADAM17* | ENSG00000151694.14 | 6.68 | 6.27 | 0.41 | 1.33 | 0.000215965 | 0.002039349 |
| 224 | *ALDH3A2* | ENSG00000072210.19 | 9.04 | 8.64 | 0.41 | 1.33 | 9.0678E-06 | 0.000198395 |
| 25820 | *ARIH1* | ENSG00000166233.15 | 7.67 | 7.26 | 0.41 | 1.33 | 0.001211129 | 0.007288625 |
| 5214 | *PFKP* | ENSG00000067057.18 | 6.73 | 6.32 | 0.41 | 1.33 | 0.000547513 | 0.004062098 |
| 28998 | *MRPL13* | ENSG00000172172.8 | 6.04 | 5.64 | 0.41 | 1.32 | 0.000255247 | 0.002333415 |
| 211 | *ALAS1* | ENSG00000023330.15 | 7.32 | 6.92 | 0.41 | 1.32 | 1.72237E-05 | 0.000322332 |
| 153339 | *TMEM167A* | ENSG00000174695.10 | 8.09 | 7.69 | 0.41 | 1.32 | 0.000453267 | 0.003534067 |
| 1810 | *DR1* | ENSG00000117505.13 | 7.50 | 7.09 | 0.40 | 1.32 | 1.65143E-05 | 0.000311385 |
| 23338 | *JADE2* | ENSG00000043143.22 | 6.55 | 6.15 | 0.40 | 1.32 | 0.000284368 | 0.002515076 |
| 9950 | *GOLGA5* | ENSG00000066455.13 | 6.30 | 5.90 | 0.40 | 1.32 | 0.000218304 | 0.00205683 |
| 57326 | *PBXIP1* | ENSG00000163346.17 | 8.84 | 8.43 | 0.40 | 1.32 | 4.37944E-06 | 0.000117979 |
| 7423 | *VEGFB* | ENSG00000173511.10 | 6.21 | 5.81 | 0.40 | 1.32 | 0.001519763 | 0.008673051 |
| 83606 | *GUCD1* | ENSG00000138867.17 | 7.17 | 6.77 | 0.40 | 1.32 | 0.000628891 | 0.004484098 |
| 63915 | *BLOC1S5* | ENSG00000188428.20 | 5.58 | 5.18 | 0.40 | 1.32 | 0.000327632 | 0.002775069 |
| 26959 | *HBP1* | ENSG00000105856.14 | 6.96 | 6.55 | 0.40 | 1.32 | 3.10416E-05 | 0.00050345 |
| 196527 | *ANO6* | ENSG00000177119.17 | 7.98 | 7.58 | 0.40 | 1.32 | 1.72136E-05 | 0.000322332 |
| 2935 | *GSPT1* | ENSG00000103342.13 | 8.53 | 8.13 | 0.40 | 1.32 | 4.96007E-06 | 0.000128799 |
| 60313 | *GPBP1L1* | ENSG00000159592.11 | 6.96 | 6.56 | 0.40 | 1.32 | 0.000639583 | 0.004543067 |
| 2879 | *GPX4* | ENSG00000167468.19 | 7.32 | 6.92 | 0.40 | 1.32 | 1.8709E-05 | 0.00034161 |
| 51478 | *HSD17B7* | ENSG00000132196.16 | 5.12 | 4.72 | 0.40 | 1.32 | 0.000957458 | 0.006078884 |
| 80321 | *CEP70* | ENSG00000114107.9 | 5.75 | 5.35 | 0.40 | 1.32 | 0.000144495 | 0.001525122 |
| 4864 | *NPC1* | ENSG00000141458.13 | 6.98 | 6.58 | 0.40 | 1.32 | 0.001210118 | 0.007285461 |
| 729082 | *OIP5-AS1* | ENSG00000247556.6 | 8.34 | 7.94 | 0.40 | 1.32 | 6.34837E-05 | 0.000848486 |
| 10120 | *ACTR1B* | ENSG00000115073.8 | 6.16 | 5.76 | 0.40 | 1.32 | 0.000101057 | 0.001176698 |
| 6785 | *ELOVL4* | ENSG00000118402.6 | 6.14 | 5.75 | 0.40 | 1.32 | 0.000189921 | 0.001852197 |
| 51512 | *GTSE1* | ENSG00000075218.19 | 6.08 | 5.68 | 0.40 | 1.32 | 0.000166782 | 0.001684544 |
| 5236 | *PGM1* | ENSG00000079739.17 | 7.60 | 7.20 | 0.40 | 1.32 | 4.56392E-05 | 0.000670253 |
| 79102 | *RNF26* | ENSG00000173456.5 | 6.65 | 6.25 | 0.40 | 1.32 | 0.000110542 | 0.001255965 |
| 23157 | *SEPTIN6* | ENSG00000125354.24 | 8.00 | 7.61 | 0.40 | 1.32 | 7.16371E-06 | 0.000167313 |
| 27250 | *PDCD4* | ENSG00000150593.18 | 7.32 | 6.93 | 0.40 | 1.32 | 0.000302628 | 0.002633123 |
| 64793 | *CEP85* | ENSG00000130695.16 | 6.05 | 5.65 | 0.40 | 1.32 | 0.000423598 | 0.003358572 |
| 54629 | *MINDY2* | ENSG00000128923.11 | 5.59 | 5.20 | 0.40 | 1.32 | 0.00123734 | 0.007404797 |
| 9040 | *UBE2M* | ENSG00000130725.8 | 6.57 | 6.17 | 0.39 | 1.31 | 0.000166646 | 0.001684306 |
| 29889 | *GNL2* | ENSG00000134697.13 | 6.93 | 6.53 | 0.39 | 1.31 | 6.92489E-05 | 0.000904575 |
| 967 | *CD63* | ENSG00000135404.12 | 9.95 | 9.56 | 0.39 | 1.31 | 3.28701E-05 | 0.000525396 |
| 54386 | *TERF2IP* | ENSG00000166848.7 | 7.12 | 6.73 | 0.39 | 1.31 | 3.24613E-05 | 0.000519415 |
| 3298 | *HSF2* | ENSG00000025156.13 | 6.19 | 5.80 | 0.39 | 1.31 | 0.000466515 | 0.003607332 |
| 10525 | *HYOU1* | ENSG00000149428.20 | 8.74 | 8.34 | 0.39 | 1.31 | 1.14018E-05 | 0.000237021 |
| 64981 | *MRPL34* | ENSG00000130312.7 | 5.17 | 4.78 | 0.39 | 1.31 | 0.000534731 | 0.003988954 |
| 3835 | *KIF22* | ENSG00000079616.14 | 7.06 | 6.66 | 0.39 | 1.31 | 3.55971E-05 | 0.000554805 |
| 7988 | *ZNF212* | ENSG00000170260.9 | 5.29 | 4.90 | 0.39 | 1.31 | 0.000731145 | 0.005006276 |

| | | | | | | | | |
|---|---|---|---|---|---|---|---|---|
| 871 | *SERPINH1* | ENSG00000149257.16 | 7.36 | 6.97 | 0.39 | 1.31 | 8.43769E-05 | 0.001035372 |
| 54926 | *UBE2R2* | ENSG00000107341.5 | 7.19 | 6.80 | 0.39 | 1.31 | 1.39763E-05 | 0.000277474 |
| 9296 | *ATP6V1F* | ENSG00000128524.5 | 6.99 | 6.60 | 0.39 | 1.31 | 7.99825E-05 | 0.00099623 |
| 5376 | *PMP22* | ENSG00000109099.16 | 5.56 | 5.17 | 0.39 | 1.31 | 0.00023223 | 0.002160905 |
| 51776 | *MAP3K20* | ENSG00000091436.17 | 6.60 | 6.21 | 0.39 | 1.31 | 0.000563999 | 0.004141422 |
| 1196 | *CLK2* | ENSG00000176444.19 | 6.00 | 5.61 | 0.39 | 1.31 | 0.000149667 | 0.001562141 |
| 10444 | *ZER1* | ENSG00000160445.11 | 7.67 | 7.28 | 0.39 | 1.31 | 1.29427E-05 | 0.000260531 |
| 22903 | *BTBD3* | ENSG00000132640.15 | 6.42 | 6.03 | 0.39 | 1.31 | 0.000646246 | 0.004568776 |
| 7841 | *MOGS* | ENSG00000115275.15 | 6.43 | 6.05 | 0.39 | 1.31 | 0.001084983 | 0.006682196 |
| 5087 | *PBX1* | ENSG00000185630.19 | 9.07 | 8.68 | 0.39 | 1.31 | 2.11435E-05 | 0.000375998 |
| 55334 | *SLC39A9* | ENSG00000029364.12 | 6.91 | 6.52 | 0.39 | 1.31 | 6.36371E-05 | 0.000849004 |
| 9820 | *CUL7* | ENSG00000044090.13 | 6.33 | 5.94 | 0.39 | 1.31 | 7.90431E-05 | 0.000989456 |
| 170954 | *PPP1R18* | ENSG00000146112.12 | 5.94 | 5.55 | 0.39 | 1.31 | 0.000215381 | 0.002036959 |
| 29927 | *SEC61A1* | ENSG00000058262.10 | 9.45 | 9.06 | 0.39 | 1.31 | 1.72618E-05 | 0.000322644 |
| 4363 | *ABCC1* | ENSG00000103222.20 | 6.73 | 6.34 | 0.39 | 1.31 | 0.000130732 | 0.001412634 |
| 3572 | *IL6ST* | ENSG00000134352.20 | 8.43 | 8.04 | 0.39 | 1.31 | 0.000590726 | 0.00429356 |
| 200933 | *FBXO45* | ENSG00000174013.8 | 6.19 | 5.80 | 0.39 | 1.31 | 0.000141477 | 0.001501223 |
| 27107 | *ZBTB11* | ENSG00000066422.6 | 6.21 | 5.82 | 0.39 | 1.31 | 8.79464E-05 | 0.001067088 |
| 64083 | *GOLPH3* | ENSG00000113384.14 | 7.01 | 6.63 | 0.39 | 1.31 | 4.19995E-05 | 0.000631633 |
| 80223 | *RAB11FIP1* | ENSG00000156675.16 | 6.79 | 6.41 | 0.39 | 1.31 | 0.00089996 | 0.00582783 |
| 79901 | *CYBRD1* | ENSG00000071967.12 | 5.68 | 5.30 | 0.39 | 1.31 | 0.000655014 | 0.004615541 |
| 4061 | *LY6E* | ENSG00000160932.11 | 8.61 | 8.23 | 0.39 | 1.31 | 7.82736E-05 | 0.000982281 |
| 4891 | *SLC11A2* | ENSG00000110911.17 | 6.54 | 6.16 | 0.38 | 1.31 | 0.000375692 | 0.003070983 |
| 51187 | *RSL24D1* | ENSG00000137876.11 | 7.38 | 7.00 | 0.38 | 1.31 | 0.000111989 | 0.001268557 |
| 51006 | *SLC35C2* | ENSG00000080189.15 | 6.16 | 5.78 | 0.38 | 1.30 | 0.001013617 | 0.006354795 |
| 6009 | *RHEB* | ENSG00000106615.10 | 8.50 | 8.11 | 0.38 | 1.30 | 9.30203E-06 | 0.000201755 |
| 7326 | *UBE2G1* | ENSG00000132388.13 | 7.04 | 6.66 | 0.38 | 1.30 | 0.000980728 | 0.00618737 |
| 1997 | *ELF1* | ENSG00000120690.16 | 5.92 | 5.54 | 0.38 | 1.30 | 0.000142441 | 0.001506617 |
| 8886 | *DDX18* | ENSG00000088205.13 | 7.17 | 6.79 | 0.38 | 1.30 | 0.000811641 | 0.005426005 |
| 132 | *ADK* | ENSG00000156110.15 | 5.43 | 5.05 | 0.38 | 1.30 | 0.001235204 | 0.007397859 |
| 7048 | *TGFBR2* | ENSG00000163513.19 | 6.29 | 5.91 | 0.38 | 1.30 | 0.000294465 | 0.002578501 |
| 5518 | *PPP2R1A* | ENSG00000105568.18 | 9.83 | 9.45 | 0.38 | 1.30 | 1.6001E-05 | 0.000304603 |
| 6944 | *VPS72* | ENSG00000163159.15 | 6.53 | 6.15 | 0.38 | 1.30 | 2.39746E-05 | 0.000413521 |
| 9882 | *TBC1D4* | ENSG00000136111.14 | 8.91 | 8.53 | 0.38 | 1.30 | 6.57646E-05 | 0.000871306 |
| 6536 | *SLC6A9* | ENSG00000196517.13 | 6.83 | 6.45 | 0.38 | 1.30 | 0.001769744 | 0.009701276 |
| 55095 | *SAMD4B* | ENSG00000179134.16 | 7.09 | 6.71 | 0.38 | 1.30 | 0.000373939 | 0.00306524 |
| 9353 | *SLIT2* | ENSG00000145147.20 | 5.60 | 5.22 | 0.38 | 1.30 | 0.001288447 | 0.007658733 |
| 332 | *BIRC5* | ENSG00000089685.15 | 7.24 | 6.86 | 0.38 | 1.30 | 0.001067381 | 0.006610734 |
| 29093 | *MRPL22* | ENSG00000082515.18 | 4.96 | 4.58 | 0.38 | 1.30 | 0.000875542 | 0.005724516 |
| 3727 | *JUND* | ENSG00000130522.6 | 6.78 | 6.40 | 0.38 | 1.30 | 0.001553521 | 0.00883212 |
| 10426 | *TUBGCP3* | ENSG00000126216.15 | 6.24 | 5.86 | 0.38 | 1.30 | 0.000459962 | 0.003569584 |
| 54887 | *UHRF1BP1* | ENSG00000065060.18 | 6.42 | 6.04 | 0.38 | 1.30 | 4.50375E-05 | 0.000664015 |
| 10133 | *OPTN* | ENSG00000123240.17 | 7.18 | 6.80 | 0.38 | 1.30 | 0.000123504 | 0.001361993 |
| 23200 | *ATP11B* | ENSG00000058063.16 | 6.60 | 6.23 | 0.38 | 1.30 | 0.000332988 | 0.002810918 |
| 8731 | *RNMT* | ENSG00000101654.18 | 6.60 | 6.23 | 0.38 | 1.30 | 0.000129235 | 0.001403532 |
| 4741 | *NEFM* | ENSG00000104722.14 | 5.70 | 5.32 | 0.38 | 1.30 | 0.000793809 | 0.005326007 |
| 7105 | *TSPAN6* | ENSG00000000003.15 | 6.88 | 6.50 | 0.38 | 1.30 | 0.00172621 | 0.009553351 |
| 54861 | *SNRK* | ENSG00000163788.14 | 5.62 | 5.25 | 0.38 | 1.30 | 0.001126766 | 0.006897761 |
| 51094 | *ADIPOR1* | ENSG00000159346.13 | 7.44 | 7.06 | 0.38 | 1.30 | 3.9886E-05 | 0.000607467 |
| 7040 | *TGFB1* | ENSG00000105329.11 | 6.13 | 5.75 | 0.38 | 1.30 | 0.000364425 | 0.003005306 |
| 9853 | *RUSC2* | ENSG00000198853.12 | 6.49 | 6.11 | 0.38 | 1.30 | 0.000310404 | 0.00267438 |
| 84056 | *KATNAL1* | ENSG00000102781.14 | 5.77 | 5.40 | 0.38 | 1.30 | 0.001535291 | 0.008741723 |
| 9337 | *CNOT8* | ENSG00000155508.14 | 6.69 | 6.31 | 0.38 | 1.30 | 8.91696E-05 | 0.001076707 |
| 64853 | *AIDA* | ENSG00000186063.13 | 7.30 | 6.92 | 0.38 | 1.30 | 2.28998E-05 | 0.000398265 |
| 7745 | *ZKSCAN8* | ENSG00000198315.11 | 6.70 | 6.32 | 0.38 | 1.30 | 0.000362536 | 0.002994661 |
| 9659 | *PDE4DIP* | ENSG00000178104.19 | 6.39 | 6.01 | 0.38 | 1.30 | 0.000449803 | 0.003514366 |
| 3688 | *ITGB1* | ENSG00000150093.20 | 9.54 | 9.16 | 0.38 | 1.30 | 7.06892E-06 | 0.000166037 |
| 7417 | *VDAC2* | ENSG00000165637.14 | 7.34 | 6.96 | 0.38 | 1.30 | 9.79136E-05 | 0.00115445 |
| 4217 | *MAP3K5* | ENSG00000197442.10 | 6.52 | 6.15 | 0.37 | 1.30 | 0.000487685 | 0.003721229 |
| 7386 | *UQCRFS1* | ENSG00000169021.6 | 7.41 | 7.04 | 0.37 | 1.30 | 8.03258E-05 | 0.000998848 |
| 9144 | *SYNGR2* | ENSG00000108639.8 | 5.94 | 5.57 | 0.37 | 1.30 | 0.000365656 | 0.003013798 |
| 4286 | *MITF* | ENSG00000187098.18 | 6.59 | 6.22 | 0.37 | 1.30 | 0.000192013 | 0.001867039 |
| 84515 | *MCM8* | ENSG00000125885.13 | 5.77 | 5.39 | 0.37 | 1.30 | 0.000582209 | 0.004248117 |
| 84268 | *RPAIN* | ENSG00000129197.15 | 4.98 | 4.60 | 0.37 | 1.30 | 0.000930074 | 0.005960495 |
| 54492 | *NEURL1B* | ENSG00000214357.9 | 7.90 | 7.53 | 0.37 | 1.30 | 3.16406E-05 | 0.000510022 |
| 7994 | *KAT6A* | ENSG00000083168.11 | 7.13 | 6.76 | 0.37 | 1.29 | 0.000307757 | 0.002662566 |
| 51727 | *CMPK1* | ENSG00000162368.14 | 8.09 | 7.72 | 0.37 | 1.29 | 4.68843E-05 | 0.000683854 |
| 81857 | *MED25* | ENSG00000104973.19 | 7.64 | 7.26 | 0.37 | 1.29 | 8.438E-05 | 0.001035372 |
| 92259 | *MRPS36* | ENSG00000134056.12 | 5.76 | 5.38 | 0.37 | 1.29 | 0.000455071 | 0.003537187 |
| 112483 | *SAT2* | ENSG00000141504.12 | 5.87 | 5.50 | 0.37 | 1.29 | 0.00030998 | 0.002672305 |
| 541578 | *EOLA2* | ENSG00000197021.9 | 5.31 | 4.94 | 0.37 | 1.29 | 0.000132703 | 0.001427812 |
| 84148 | *KAT8* | ENSG00000103510.20 | 4.66 | 4.29 | 0.37 | 1.29 | 0.00149615 | 0.008570881 |
| 54704 | *PDP1* | ENSG00000164951.16 | 6.48 | 6.10 | 0.37 | 1.29 | 0.000672071 | 0.004704822 |
| 103910 | *MYL12B* | ENSG00000118680.14 | 8.65 | 8.28 | 0.37 | 1.29 | 2.02793E-05 | 0.000364518 |
| 10424 | *PGRMC2* | ENSG00000164040.18 | 6.80 | 6.42 | 0.37 | 1.29 | 0.000923307 | 0.005927252 |
| 3752 | *KCND3* | ENSG00000171385.10 | 6.39 | 6.02 | 0.37 | 1.29 | 0.000124245 | 0.001366147 |
| 9867 | *PJA2* | ENSG00000198961.10 | 8.12 | 7.75 | 0.37 | 1.29 | 2.4343E-05 | 0.00041803 |
| 2132 | *EXT2* | ENSG00000151348.16 | 8.00 | 7.63 | 0.37 | 1.29 | 0.000281467 | 0.002497588 |
| 439921 | *MXRA7* | ENSG00000182534.14 | 6.56 | 6.19 | 0.37 | 1.29 | 0.000424805 | 0.003366363 |
| 55605 | *KIF21A* | ENSG00000139116.19 | 6.49 | 6.12 | 0.37 | 1.29 | 0.000936703 | 0.005990188 |
| 80830 | *APOL6* | ENSG00000221963.6 | 6.15 | 5.78 | 0.37 | 1.29 | 0.000438262 | 0.00344391 |
| 10746 | *MAP3K2* | ENSG00000169967.17 | 7.08 | 6.71 | 0.37 | 1.29 | 0.000418038 | 0.003330324 |
| 10163 | *WASF2* | ENSG00000158195.11 | 7.62 | 7.25 | 0.37 | 1.29 | 0.000477838 | 0.003661124 |
| 5584 | *PRKCI* | ENSG00000163558.13 | 7.10 | 6.74 | 0.37 | 1.29 | 7.07198E-05 | 0.00091899 |
| 25797 | *QPCT* | ENSG00000115828.17 | 10.16 | 9.80 | 0.37 | 1.29 | 7.95809E-05 | 0.000992876 |
| 286410 | *ATP11C* | ENSG00000101974.15 | 6.08 | 5.71 | 0.37 | 1.29 | 0.000861431 | 0.005665591 |
| 2339 | *FNTA* | ENSG00000168522.13 | 5.91 | 5.55 | 0.37 | 1.29 | 0.000798336 | 0.005344437 |
| 79589 | *RNF128* | ENSG00000133135.14 | 7.24 | 6.87 | 0.36 | 1.29 | 7.76894E-05 | 0.000977402 |
| 57617 | *VPS18* | ENSG00000104142.11 | 6.24 | 5.88 | 0.36 | 1.29 | 0.001427789 | 0.008270814 |

| | | | | | | | | |
|---|---|---|---|---|---|---|---|---|
| 80790 | *CMIP* | ENSG00000153815.18 | 6.47 | 6.10 | 0.36 | 1.29 | 0.000349628 | 0.002922817 |
| 10159 | *ATP6AP2* | ENSG00000182220.15 | 7.81 | 7.44 | 0.36 | 1.29 | 7.34924E-05 | 0.000940271 |
| 523 | *ATP6V1A* | ENSG00000114573.10 | 8.48 | 8.11 | 0.36 | 1.29 | 2.51722E-05 | 0.000430308 |
| 3842 | *TNPO1* | ENSG00000083312.19 | 8.53 | 8.17 | 0.36 | 1.29 | 5.07067E-05 | 0.00072138 |
| 10550 | *ARL6IP5* | ENSG00000144746.7 | 8.15 | 7.79 | 0.36 | 1.29 | 1.65734E-05 | 0.000312108 |
| 10116 | *FEM1B* | ENSG00000169018.6 | 8.36 | 8.00 | 0.36 | 1.29 | 5.81926E-05 | 0.000797637 |
| 25994 | *HIGD1A* | ENSG00000181061.14 | 7.41 | 7.05 | 0.36 | 1.29 | 0.000759647 | 0.005154402 |
| 55352 | *COPRS* | ENSG00000172301.11 | 6.11 | 5.75 | 0.36 | 1.28 | 0.000120071 | 0.001341882 |
| 10567 | *RABAC1* | ENSG00000105404.11 | 6.12 | 5.76 | 0.36 | 1.28 | 0.001055005 | 0.006545918 |
| 10329 | *RXYLT1* | ENSG00000118600.13 | 5.94 | 5.58 | 0.36 | 1.28 | 0.001046538 | 0.006509528 |
| 9444 | *QKI* | ENSG00000112531.17 | 8.30 | 7.94 | 0.36 | 1.28 | 0.00038152 | 0.00309861 |
| 3608 | *ILF2* | ENSG00000143621.17 | 9.12 | 8.76 | 0.36 | 1.28 | 1.04515E-05 | 0.000221564 |
| 5978 | *REST* | ENSG00000084093.19 | 6.52 | 6.16 | 0.36 | 1.28 | 0.000811967 | 0.005426005 |
| 7046 | *TGFBR1* | ENSG00000106799.13 | 8.15 | 7.79 | 0.36 | 1.28 | 0.000463541 | 0.003591681 |
| 54890 | *ALKBH5* | ENSG00000091542.9 | 7.71 | 7.35 | 0.36 | 1.28 | 0.000110087 | 0.001252688 |
| 11041 | *B4GAT1* | ENSG00000174684.7 | 6.23 | 5.87 | 0.36 | 1.28 | 0.000206299 | 0.001975966 |
| 7873 | *MANF* | ENSG00000145050.19 | 6.21 | 5.85 | 0.36 | 1.28 | 0.000189797 | 0.00185219 |
| 55787 | *TXLNG* | ENSG00000086712.13 | 5.75 | 5.39 | 0.36 | 1.28 | 0.000939199 | 0.005995238 |
| 57515 | *SERINC1* | ENSG00000111897.7 | 9.11 | 8.76 | 0.36 | 1.28 | 8.36721E-05 | 0.00103059 |
| 219743 | *TYSND1* | ENSG00000156521.14 | 4.55 | 4.19 | 0.36 | 1.28 | 0.000550068 | 0.004075012 |
| 81608 | *FIP1L1* | ENSG00000145216.16 | 6.33 | 5.97 | 0.36 | 1.28 | 0.000131165 | 0.001414578 |
| 26470 | *SEZ6L2* | ENSG00000174938.15 | 7.28 | 6.92 | 0.36 | 1.28 | 7.62653E-05 | 0.000966779 |
| 55788 | *LMBRD1* | ENSG00000168216.13 | 6.40 | 6.04 | 0.36 | 1.28 | 0.000466519 | 0.003607332 |
| 54928 | *BPNT2* | ENSG00000104331.9 | 8.53 | 8.18 | 0.36 | 1.28 | 5.57212E-05 | 0.000770091 |
| 2889 | *RAPGEF1* | ENSG00000107263.19 | 7.47 | 7.11 | 0.36 | 1.28 | 0.000310641 | 0.00267438 |
| 1889 | *ECE1* | ENSG00000117298.16 | 8.03 | 7.67 | 0.36 | 1.28 | 0.000220715 | 0.002074333 |
| 3091 | *HIF1A* | ENSG00000100644.17 | 9.68 | 9.32 | 0.36 | 1.28 | 2.18121E-05 | 0.000385151 |
| 2787 | *GNG5* | ENSG00000174021.12 | 6.29 | 5.94 | 0.36 | 1.28 | 9.03841E-05 | 0.001084817 |
| 11321 | *GPN1* | ENSG00000198522.14 | 5.78 | 5.42 | 0.36 | 1.28 | 0.000114485 | 0.001291963 |
| 55041 | *PLEKHB2* | ENSG00000115762.17 | 8.10 | 7.75 | 0.35 | 1.28 | 3.16703E-05 | 0.000510022 |
| 22974 | *TPX2* | ENSG00000088325.16 | 8.72 | 8.36 | 0.35 | 1.28 | 1.06531E-05 | 0.000225519 |
| 54732 | *TMED9* | ENSG00000184840.12 | 7.49 | 7.13 | 0.35 | 1.28 | 0.000417469 | 0.003327555 |
| 5861 | *RAB1A* | ENSG00000138069.18 | 7.44 | 7.09 | 0.35 | 1.28 | 5.68647E-05 | 0.000781578 |
| 154007 | *SNRNP48* | ENSG00000168566.13 | 6.54 | 6.19 | 0.35 | 1.28 | 0.000337263 | 0.002834252 |
| 1894 | *ECT2* | ENSG00000114346.14 | 8.42 | 8.07 | 0.35 | 1.28 | 9.89184E-05 | 0.001159896 |
| 25829 | *TMEM184B* | ENSG00000198792.13 | 6.57 | 6.21 | 0.35 | 1.28 | 0.000153423 | 0.001592657 |
| 400 | *ARL1* | ENSG00000120805.14 | 7.60 | 7.25 | 0.35 | 1.28 | 0.001046356 | 0.006509528 |
| 50854 | *SNHG32* | ENSG00000204387.14 | 6.51 | 6.16 | 0.35 | 1.28 | 0.000198818 | 0.001920243 |
| 28996 | *HIPK2* | ENSG00000064393.16 | 8.01 | 7.66 | 0.35 | 1.27 | 0.000149014 | 0.001558568 |
| 23019 | *CNOT1* | ENSG00000125107.19 | 8.92 | 8.57 | 0.35 | 1.27 | 0.001521928 | 0.008682106 |
| 54885 | *TBC1D8B* | ENSG00000133138.20 | 5.74 | 5.39 | 0.35 | 1.27 | 0.000257681 | 0.002351082 |
| 85403 | *EAF1* | ENSG00000144597.14 | 6.57 | 6.22 | 0.35 | 1.27 | 0.000259688 | 0.002362645 |
| 1054 | *CEBPG* | ENSG00000153879.9 | 6.83 | 6.48 | 0.35 | 1.27 | 0.000160354 | 0.001640098 |
| 51290 | *ERGIC2* | ENSG00000087502.19 | 7.41 | 7.06 | 0.35 | 1.27 | 0.00076135 | 0.005157052 |
| 5869 | *RAB5B* | ENSG00000111540.16 | 8.72 | 8.37 | 0.35 | 1.27 | 0.000194694 | 0.001886485 |
| 1832 | *DSP* | ENSG00000096696.15 | 7.74 | 7.39 | 0.35 | 1.27 | 0.000552287 | 0.0040854 |
| 10137 | *RBM12* | ENSG00000244462.8 | 7.64 | 7.29 | 0.35 | 1.27 | 0.000646202 | 0.004568776 |
| 83540 | *NUF2* | ENSG00000143228.13 | 6.51 | 6.17 | 0.35 | 1.27 | 0.000159581 | 0.001634912 |
| 6480 | *ST6GAL1* | ENSG00000073849.16 | 6.93 | 6.59 | 0.35 | 1.27 | 0.000523552 | 0.003928996 |
| 53838 | *C11orf24* | ENSG00000171067.11 | 5.70 | 5.35 | 0.35 | 1.27 | 0.000349518 | 0.002922817 |
| 9493 | *KIF23* | ENSG00000137807.16 | 8.29 | 7.94 | 0.35 | 1.27 | 3.56524E-05 | 0.000555091 |
| 10403 | *NDC80* | ENSG00000080986.13 | 6.32 | 5.98 | 0.35 | 1.27 | 0.001035812 | 0.006456186 |
| 22931 | *RAB18* | ENSG00000099246.18 | 7.87 | 7.52 | 0.35 | 1.27 | 0.000138246 | 0.001475771 |
| 23210 | *JMJD6* | ENSG00000070495.15 | 5.22 | 4.87 | 0.35 | 1.27 | 0.000491934 | 0.003746037 |
| 7110 | *TMF1* | ENSG00000144747.17 | 6.62 | 6.28 | 0.35 | 1.27 | 0.000366873 | 0.003020388 |
| 57462 | *MYORG* | ENSG00000164976.9 | 6.45 | 6.11 | 0.35 | 1.27 | 0.001623569 | 0.009140341 |
| 8727 | *CTNNAL1* | ENSG00000119326.15 | 9.25 | 8.91 | 0.35 | 1.27 | 3.54512E-05 | 0.000554257 |
| 5347 | *PLK1* | ENSG00000166851.15 | 7.24 | 6.90 | 0.35 | 1.27 | 0.000173819 | 0.00173345 |
| 994 | *CDC25B* | ENSG00000101224.18 | 6.82 | 6.47 | 0.34 | 1.27 | 0.000390523 | 0.003152966 |
| 65056 | *GPBP1* | ENSG00000062194.16 | 7.76 | 7.42 | 0.34 | 1.27 | 6.45539E-05 | 0.000860471 |
| 144455 | *E2F7* | ENSG00000165891.16 | 5.89 | 5.55 | 0.34 | 1.27 | 0.001135657 | 0.006934533 |
| 50999 | *TMED5* | ENSG00000117500.13 | 8.58 | 8.24 | 0.34 | 1.27 | 0.000190662 | 0.001856692 |
| 84687 | *PPP1R9B* | ENSG00000108819.11 | 6.45 | 6.11 | 0.34 | 1.27 | 0.000411454 | 0.003295364 |
| 51107 | *APH1A* | ENSG00000117362.13 | 8.56 | 8.22 | 0.34 | 1.27 | 0.000115078 | 0.001295733 |
| 55839 | *CENPN* | ENSG00000166451.14 | 7.23 | 6.88 | 0.34 | 1.27 | 7.78854E-05 | 0.000979047 |
| 114882 | *OSBPL8* | ENSG00000091039.17 | 7.55 | 7.21 | 0.34 | 1.27 | 9.84423E-05 | 0.001156906 |
| 53354 | *PANK1* | ENSG00000152782.18 | 5.21 | 4.87 | 0.34 | 1.27 | 0.000760066 | 0.005154917 |
| 146223 | *CMTM4* | ENSG00000183723.13 | 7.50 | 7.16 | 0.34 | 1.27 | 5.06972E-05 | 0.00072138 |
| 27043 | *PELP1* | ENSG00000141456.16 | 6.97 | 6.63 | 0.34 | 1.27 | 0.000211438 | 0.002012346 |
| 54205 | *CYCS* | ENSG00000172115.9 | 8.27 | 7.93 | 0.34 | 1.27 | 2.79047E-05 | 0.000465357 |
| 5654 | *HTRA1* | ENSG00000166033.13 | 5.96 | 5.62 | 0.34 | 1.27 | 0.000840892 | 0.005564791 |
| 10787 | *NCKAP1* | ENSG00000061676.15 | 8.63 | 8.29 | 0.34 | 1.27 | 6.30648E-05 | 0.000845125 |
| 347902 | *AMIGO2* | ENSG00000139211.7 | 5.95 | 5.61 | 0.34 | 1.27 | 0.0009614 | 0.006098752 |
| 64781 | *CERK* | ENSG00000100422.14 | 4.80 | 4.46 | 0.34 | 1.27 | 0.001734787 | 0.009576773 |
| 4240 | *MFGE8* | ENSG00000140545.15 | 7.32 | 6.98 | 0.34 | 1.27 | 4.95578E-05 | 0.000709162 |
| 55770 | *EXOC2* | ENSG00000112685.14 | 6.46 | 6.12 | 0.34 | 1.27 | 7.21736E-05 | 0.000929034 |
| 8237 | *USP11* | ENSG00000102226.10 | 7.67 | 7.33 | 0.34 | 1.27 | 0.000907991 | 0.005864595 |
| 55165 | *CEP55* | ENSG00000138180.16 | 6.71 | 6.37 | 0.34 | 1.27 | 6.92058E-05 | 0.000904575 |
| 4698 | *NDUFA5* | ENSG00000128609.16 | 8.38 | 8.04 | 0.34 | 1.26 | 9.40693E-05 | 0.001119656 |
| 4247 | *MGAT2* | ENSG00000168282.6 | 7.00 | 6.66 | 0.34 | 1.26 | 8.73571E-05 | 0.001061654 |
| 6416 | *MAP2K4* | ENSG00000065559.15 | 6.30 | 5.96 | 0.34 | 1.26 | 0.000581271 | 0.004243335 |
| 5048 | *PAFAH1B1* | ENSG00000007168.14 | 8.24 | 7.90 | 0.34 | 1.26 | 0.001407282 | 0.008196312 |
| 2931 | *GSK3A* | ENSG00000105723.13 | 7.11 | 6.78 | 0.34 | 1.26 | 0.000269725 | 0.002419784 |
| 64282 | *TENT4B* | ENSG00000121274.14 | 6.71 | 6.37 | 0.34 | 1.26 | 0.001760998 | 0.009668126 |
| 2011 | *MARK2* | ENSG00000072518.22 | 6.02 | 5.68 | 0.34 | 1.26 | 0.000477499 | 0.003661124 |
| 1459 | *CSNK2A2* | ENSG00000070770.10 | 7.40 | 7.06 | 0.34 | 1.26 | 0.000714764 | 0.004925571 |
| 9802 | *DAZAP2* | ENSG00000183283.16 | 9.20 | 8.87 | 0.34 | 1.26 | 5.14437E-05 | 0.000727041 |
| 4253 | *MIA2* | ENSG00000150527.18 | 8.99 | 8.66 | 0.34 | 1.26 | 0.000145349 | 0.00153091 |
| 23215 | *PRRC2C* | ENSG00000117523.18 | 8.85 | 8.52 | 0.34 | 1.26 | 0.000212918 | 0.002020981 |

| | | | | | | | | |
|---|---|---|---|---|---|---|---|---|
| 9821 | *RB1CC1* | ENSG00000023287.13 | 7.00 | 6.66 | 0.34 | 1.26 | 0.000689251 | 0.004793777 |
| 10491 | *CRTAP* | ENSG00000170275.15 | 8.47 | 8.13 | 0.34 | 1.26 | 0.000155521 | 0.001609801 |
| 9230 | *RAB11B* | ENSG00000185236.12 | 6.73 | 6.39 | 0.34 | 1.26 | 0.000227693 | 0.002125275 |
| 6730 | *SRP68* | ENSG00000167881.15 | 7.43 | 7.09 | 0.34 | 1.26 | 0.000162266 | 0.001653392 |
| 6612 | *SUMO3* | ENSG00000184900.16 | 7.81 | 7.47 | 0.34 | 1.26 | 0.000196698 | 0.001903439 |
| 4134 | *MAP4* | ENSG00000047849.22 | 8.04 | 7.70 | 0.34 | 1.26 | 6.98918E-05 | 0.000911387 |
| 345778 | *MTX3* | ENSG00000177034.17 | 6.79 | 6.46 | 0.34 | 1.26 | 0.000331701 | 0.002803207 |
| 5894 | *RAF1* | ENSG00000132155.14 | 7.95 | 7.61 | 0.33 | 1.26 | 5.62574E-05 | 0.000776073 |
| 4154 | *MBNL1* | ENSG00000152601.18 | 7.58 | 7.25 | 0.33 | 1.26 | 0.000114984 | 0.001295638 |
| 90417 | *KNSTRN* | ENSG00000128944.14 | 6.57 | 6.24 | 0.33 | 1.26 | 0.000259735 | 0.002362645 |
| 10983 | *CCNI* | ENSG00000118816.11 | 8.45 | 8.12 | 0.33 | 1.26 | 4.56178E-05 | 0.000670253 |
| 10376 | *TUBA1B* | ENSG00000123416.15 | 11.92 | 11.58 | 0.33 | 1.26 | 0.000132096 | 0.001423285 |
| 51504 | *TRMT112* | ENSG00000173113.7 | 6.91 | 6.58 | 0.33 | 1.26 | 0.000102767 | 0.001191064 |
| 79960 | *JADE1* | ENSG00000077684.16 | 6.81 | 6.48 | 0.33 | 1.26 | 7.72946E-05 | 0.000975706 |
| 23207 | *PLEKHM2* | ENSG00000116786.13 | 6.55 | 6.21 | 0.33 | 1.26 | 0.000266755 | 0.002406142 |
| 51335 | *NGRN* | ENSG00000182768.9 | 7.67 | 7.34 | 0.33 | 1.26 | 9.28434E-05 | 0.001106821 |
| 23198 | *PSME4* | ENSG00000068878.15 | 7.31 | 6.97 | 0.33 | 1.26 | 0.000826788 | 0.00550056 |
| 79646 | *PANK3* | ENSG00000120137.7 | 7.94 | 7.61 | 0.33 | 1.26 | 0.000192386 | 0.001868942 |
| 3069 | *HDLBP* | ENSG00000115677.18 | 8.50 | 8.17 | 0.33 | 1.26 | 0.001126715 | 0.006897761 |
| 38 | *ACAT1* | ENSG00000075239.14 | 7.60 | 7.27 | 0.33 | 1.26 | 0.000100285 | 0.001168739 |
| 51324 | *SPG21* | ENSG00000090487.11 | 6.82 | 6.49 | 0.33 | 1.26 | 0.000180319 | 0.001784054 |
| 55970 | *GNG12* | ENSG00000172380.6 | 8.54 | 8.21 | 0.33 | 1.26 | 2.72592E-05 | 0.000457132 |
| 2720 | *GLB1* | ENSG00000170266.16 | 7.48 | 7.15 | 0.33 | 1.26 | 0.0010837 | 0.006677178 |
| 3915 | *LAMC1* | ENSG00000135862.6 | 9.51 | 9.18 | 0.33 | 1.26 | 0.00124077 | 0.007419406 |
| 152006 | *RNF38* | ENSG00000137075.18 | 7.87 | 7.54 | 0.33 | 1.26 | 0.000243686 | 0.00225076 |
| 1112 | *FOXN3* | ENSG00000053254.16 | 8.16 | 7.83 | 0.33 | 1.26 | 0.000551497 | 0.004083578 |
| 10959 | *TMED2* | ENSG00000086598.11 | 8.52 | 8.19 | 0.33 | 1.26 | 0.000258257 | 0.002354904 |
| 64395 | *GMCL1* | ENSG00000087338.5 | 5.72 | 5.39 | 0.33 | 1.26 | 0.001488062 | 0.008536456 |
| 9741 | *LAPTM4A* | ENSG00000068697.7 | 8.74 | 8.41 | 0.33 | 1.26 | 0.000351934 | 0.002931283 |
| 55830 | *GLT8D1* | ENSG00000016864.19 | 7.51 | 7.18 | 0.33 | 1.26 | 0.000316365 | 0.002705601 |
| 55754 | *TMEM30A* | ENSG00000112697.17 | 8.77 | 8.44 | 0.33 | 1.26 | 0.000179064 | 0.001773978 |
| 51246 | *SHISA5* | ENSG00000164054.17 | 6.69 | 6.36 | 0.33 | 1.26 | 0.000597051 | 0.004327431 |
| 29959 | *NRBP1* | ENSG00000115216.14 | 7.74 | 7.41 | 0.33 | 1.26 | 3.53636E-05 | 0.000553464 |
| 64116 | *SLC39A8* | ENSG00000138821.14 | 7.44 | 7.11 | 0.33 | 1.26 | 9.13835E-05 | 0.001092889 |
| 23608 | *MKRN1* | ENSG00000133606.11 | 7.70 | 7.37 | 0.33 | 1.26 | 9.85093E-05 | 0.001156906 |
| 2734 | *GLG1* | ENSG00000090863.12 | 8.73 | 8.40 | 0.33 | 1.26 | 9.57521E-05 | 0.001136082 |
| 2305 | *FOXM1* | ENSG00000111206.13 | 8.53 | 8.21 | 0.33 | 1.25 | 0.000442216 | 0.00347135 |
| 22993 | *HMGXB3* | ENSG00000113716.13 | 6.52 | 6.19 | 0.33 | 1.25 | 0.000219652 | 0.002066929 |
| 84925 | *SLC49A4* | ENSG00000138463.9 | 5.91 | 5.58 | 0.33 | 1.25 | 0.001000153 | 0.006283505 |
| 8099 | *CDK2AP1* | ENSG00000111328.7 | 6.26 | 5.93 | 0.33 | 1.25 | 0.001492162 | 0.008551302 |
| 57410 | *SCYL1* | ENSG00000142186.18 | 6.53 | 6.20 | 0.33 | 1.25 | 0.000425693 | 0.003369794 |
| 51510 | *CHMP5* | ENSG00000086065.14 | 6.60 | 6.28 | 0.32 | 1.25 | 0.000262515 | 0.002382157 |
| 8408 | *ULK1* | ENSG00000177169.10 | 6.86 | 6.53 | 0.32 | 1.25 | 0.000626199 | 0.004469152 |
| 84930 | *MASTL* | ENSG00000120539.15 | 5.95 | 5.63 | 0.32 | 1.25 | 0.00035077 | 0.002928615 |
| 8545 | *CGGBP1* | ENSG00000163320.12 | 7.93 | 7.61 | 0.32 | 1.25 | 0.000389028 | 0.003144279 |
| 79786 | *KLHL36* | ENSG00000135686.13 | 6.96 | 6.63 | 0.32 | 1.25 | 0.000412786 | 0.003304272 |
| 7360 | *UGP2* | ENSG00000169764.16 | 7.59 | 7.26 | 0.32 | 1.25 | 0.000249532 | 0.002293466 |
| 8565 | *YARS1* | ENSG00000134684.12 | 7.12 | 6.80 | 0.32 | 1.25 | 0.000356117 | 0.002954652 |
| 79893 | *GGNBP2* | ENSG00000278311.5 | 6.33 | 6.01 | 0.32 | 1.25 | 0.001437848 | 0.008316244 |
| 1456 | *CSNK1G3* | ENSG00000151292.17 | 5.83 | 5.51 | 0.32 | 1.25 | 0.001830613 | 0.009972564 |
| 9275 | *BCL7B* | ENSG00000106635.8 | 6.13 | 5.81 | 0.32 | 1.25 | 0.000121336 | 0.001345991 |
| 9787 | *DLGAP5* | ENSG00000126787.13 | 7.26 | 6.93 | 0.32 | 1.25 | 0.000355605 | 0.002953665 |
| 57693 | *ZNF317* | ENSG00000130803.15 | 6.11 | 5.79 | 0.32 | 1.25 | 0.000587584 | 0.004279011 |
| 79784 | *MYH14* | ENSG00000105357.20 | 5.60 | 5.28 | 0.32 | 1.25 | 0.001649651 | 0.009247667 |
| 1362 | *CPD* | ENSG00000108582.12 | 7.81 | 7.49 | 0.32 | 1.25 | 0.000325426 | 0.002759501 |
| 23174 | *ZCCHC14* | ENSG00000140948.13 | 7.63 | 7.31 | 0.32 | 1.25 | 0.001298799 | 0.007684019 |
| 9928 | *KIF14* | ENSG00000118193.12 | 5.97 | 5.65 | 0.32 | 1.25 | 0.001371443 | 0.008024946 |
| 79080 | *CCDC86* | ENSG00000110104.13 | 6.19 | 5.87 | 0.32 | 1.25 | 0.000204743 | 0.001966087 |
| 2631 | *NIPSNAP2* | ENSG00000146729.10 | 7.23 | 6.91 | 0.32 | 1.25 | 8.98311E-05 | 0.001080349 |
| 10483 | *SEC23B* | ENSG00000101310.17 | 7.48 | 7.16 | 0.32 | 1.25 | 0.000285216 | 0.002519605 |
| 23582 | *CCNDBP1* | ENSG00000166946.14 | 5.74 | 5.42 | 0.32 | 1.25 | 0.001192101 | 0.007206968 |
| 7159 | *TP53BP2* | ENSG00000143514.17 | 6.15 | 5.84 | 0.32 | 1.25 | 0.001476276 | 0.008479733 |
| 6655 | *SOS2* | ENSG00000100485.12 | 6.97 | 6.65 | 0.32 | 1.25 | 0.000130536 | 0.001412561 |
| 63935 | *PCIF1* | ENSG00000100982.12 | 5.94 | 5.62 | 0.32 | 1.25 | 0.001442504 | 0.008335877 |
| 1725 | *DHPS* | ENSG00000095059.17 | 5.85 | 5.53 | 0.32 | 1.25 | 0.000894282 | 0.005802971 |
| 56889 | *TM9SF3* | ENSG00000077147.16 | 8.91 | 8.59 | 0.32 | 1.25 | 0.000165969 | 0.00168086 |
| 11021 | *RAB35* | ENSG00000111737.12 | 6.37 | 6.05 | 0.32 | 1.25 | 0.000589376 | 0.004285823 |
| 9213 | *XPR1* | ENSG00000143324.14 | 8.33 | 8.01 | 0.32 | 1.25 | 0.000762215 | 0.005160162 |
| 23137 | *SMC5* | ENSG00000198887.9 | 7.86 | 7.54 | 0.32 | 1.25 | 0.001139044 | 0.006938678 |
| 4478 | *MSN* | ENSG00000147065.17 | 8.20 | 7.88 | 0.32 | 1.25 | 0.000570535 | 0.004185317 |
| 4190 | *MDH1* | ENSG00000014641.21 | 8.31 | 8.00 | 0.32 | 1.25 | 0.000202065 | 0.001943997 |
| 4105 | *MAGEA6* | ENSG00000197172.11 | 7.53 | 7.22 | 0.32 | 1.25 | 0.000360557 | 0.002984103 |
| 5792 | *PTPRF* | ENSG00000142949.17 | 7.87 | 7.55 | 0.32 | 1.25 | 0.001740727 | 0.009587735 |
| 9445 | *ITM2B* | ENSG00000136156.15 | 8.69 | 8.38 | 0.32 | 1.24 | 6.27931E-05 | 0.000842988 |
| 5250 | *SLC25A3* | ENSG00000075415.15 | 9.19 | 8.87 | 0.32 | 1.24 | 0.000285815 | 0.002521926 |
| 4170 | *MCL1* | ENSG00000143384.14 | 8.84 | 8.53 | 0.32 | 1.24 | 3.46981E-05 | 0.000547617 |
| 51669 | *SARAF* | ENSG00000133872.14 | 7.75 | 7.43 | 0.31 | 1.24 | 9.89977E-05 | 0.001159919 |
| 348801 | *LNP1* | ENSG00000206535.8 | 4.62 | 4.30 | 0.31 | 1.24 | 0.001350078 | 0.007937061 |
| 23376 | *UFL1* | ENSG00000014123.10 | 6.72 | 6.40 | 0.31 | 1.24 | 0.000217144 | 0.002047184 |
| 51096 | *UTP18* | ENSG00000011260.14 | 5.94 | 5.63 | 0.31 | 1.24 | 0.000225204 | 0.002109916 |
| 60592 | *SCOC* | ENSG00000153130.18 | 8.23 | 7.91 | 0.31 | 1.24 | 0.000141585 | 0.001501223 |
| 57142 | *RTN4* | ENSG00000115310.19 | 8.18 | 7.86 | 0.31 | 1.24 | 9.94068E-05 | 0.001162897 |
| 5885 | *RAD21* | ENSG00000164754.16 | 9.44 | 9.13 | 0.31 | 1.24 | 8.32303E-05 | 0.001026461 |
| 10594 | *PRPF8* | ENSG00000174231.17 | 9.77 | 9.45 | 0.31 | 1.24 | 0.001239908 | 0.007417208 |
| 26224 | *FBXL3* | ENSG00000005812.11 | 5.93 | 5.62 | 0.31 | 1.24 | 0.000183526 | 0.001806031 |
| 1027 | *CDKN1B* | ENSG00000111276.12 | 7.27 | 6.95 | 0.31 | 1.24 | 0.000320963 | 0.002731711 |
| 56681 | *SAR1A* | ENSG00000079332.15 | 7.48 | 7.17 | 0.31 | 1.24 | 0.000160524 | 0.001640098 |
| 26097 | *CHTOP* | ENSG00000160679.13 | 7.30 | 6.98 | 0.31 | 1.24 | 0.00098389 | 0.006204708 |
| 3021 | *H3-3B* | ENSG00000132475.10 | 8.72 | 8.41 | 0.31 | 1.24 | 0.000394288 | 0.003174822 |

| | | | | | | | | |
|---|---|---|---|---|---|---|---|---|
| 25940 | *FAM98A* | ENSG00000119812.20 | 6.71 | 6.40 | 0.31 | 1.24 | 0.000431268 | 0.00340142 |
| 83658 | *DYNLRB1* | ENSG00000125971.17 | 6.90 | 6.59 | 0.31 | 1.24 | 0.000831802 | 0.005517878 |
| 6619 | *SNAPC3* | ENSG00000164975.16 | 6.35 | 6.04 | 0.31 | 1.24 | 0.000188047 | 0.001838692 |
| 10010 | *TANK* | ENSG00000136560.14 | 5.99 | 5.68 | 0.31 | 1.24 | 0.000816645 | 0.005449987 |
| 6745 | *SSR1* | ENSG00000124783.14 | 8.55 | 8.24 | 0.31 | 1.24 | 0.000106604 | 0.001223252 |
| 92370 | *PXYLP1* | ENSG00000155893.13 | 8.23 | 7.92 | 0.31 | 1.24 | 0.000141017 | 0.00149895 |
| 3679 | *ITGA7* | ENSG00000135424.19 | 8.58 | 8.27 | 0.31 | 1.24 | 0.000225629 | 0.002112575 |
| 682 | *BSG* | ENSG00000172270.22 | 9.23 | 8.92 | 0.31 | 1.24 | 0.000362116 | 0.002992841 |
| 9663 | *LPIN2* | ENSG00000101577.10 | 6.26 | 5.95 | 0.31 | 1.24 | 0.000894044 | 0.005802971 |
| 10776 | *ARPP19* | ENSG00000128989.11 | 9.06 | 8.75 | 0.31 | 1.24 | 0.000252428 | 0.002311584 |
| 60343 | *FAM3A* | ENSG00000071889.17 | 5.92 | 5.61 | 0.31 | 1.24 | 0.001667553 | 0.009319686 |
| 161176 | *SYNE3* | ENSG00000176438.13 | 6.73 | 6.42 | 0.31 | 1.24 | 0.001133042 | 0.006927016 |
| 26046 | *LTN1* | ENSG00000198862.14 | 6.60 | 6.30 | 0.31 | 1.24 | 0.0005938 | 0.004307563 |
| 55872 | *PBK* | ENSG00000168078.10 | 6.39 | 6.08 | 0.31 | 1.24 | 0.00056101 | 0.004127696 |
| 51569 | *UFM1* | ENSG00000120686.12 | 5.69 | 5.38 | 0.31 | 1.24 | 0.001228109 | 0.007373077 |
| 25949 | *SYF2* | ENSG00000117614.10 | 6.28 | 5.97 | 0.31 | 1.24 | 0.001577423 | 0.00894767 |
| 84661 | *DPY30* | ENSG00000162961.14 | 5.73 | 5.43 | 0.31 | 1.24 | 0.001298905 | 0.007684019 |
| 1314 | *COPA* | ENSG00000122218.16 | 8.43 | 8.12 | 0.31 | 1.24 | 0.001650027 | 0.009247667 |
| 3052 | *HCCS* | ENSG00000004961.15 | 5.76 | 5.46 | 0.31 | 1.24 | 0.001019652 | 0.006376652 |
| 25847 | *ANAPC13* | ENSG00000129055.13 | 6.83 | 6.52 | 0.31 | 1.24 | 0.000694467 | 0.004818885 |
| 57544 | *TXNDC16* | ENSG00000087301.9 | 6.66 | 6.36 | 0.30 | 1.23 | 0.000377599 | 0.003075272 |
| 7533 | *YWHAH* | ENSG00000128245.15 | 8.47 | 8.17 | 0.30 | 1.23 | 0.000267629 | 0.00241084 |
| 51526 | *OSER1* | ENSG00000132823.11 | 6.04 | 5.74 | 0.30 | 1.23 | 0.000484441 | 0.003702125 |
| 81610 | *FAM83D* | ENSG00000101447.15 | 6.89 | 6.59 | 0.30 | 1.23 | 0.001710326 | 0.009483932 |
| 57179 | *KIAA1191* | ENSG00000122203.15 | 7.71 | 7.40 | 0.30 | 1.23 | 7.24344E-05 | 0.000931592 |
| 53827 | *FXYD5* | ENSG00000089327.15 | 8.04 | 7.74 | 0.30 | 1.23 | 0.000472178 | 0.003636186 |
| 55270 | *NUDT15* | ENSG00000136159.4 | 5.70 | 5.39 | 0.30 | 1.23 | 0.000580258 | 0.004239605 |
| 6628 | *SNRPB* | ENSG00000125835.20 | 7.87 | 7.56 | 0.30 | 1.23 | 0.000543676 | 0.004043621 |
| 6166 | *RPL36AL* | ENSG00000165502.7 | 7.11 | 6.81 | 0.30 | 1.23 | 0.000321057 | 0.002731711 |
| 9550 | *ATP6V1G1* | ENSG00000136888.8 | 7.67 | 7.36 | 0.30 | 1.23 | 0.000425911 | 0.003369794 |
| 8976 | *WASL* | ENSG00000106299.8 | 7.65 | 7.35 | 0.30 | 1.23 | 0.000377547 | 0.003075272 |
| 23588 | *KLHDC2* | ENSG00000165516.11 | 6.26 | 5.95 | 0.30 | 1.23 | 0.000913046 | 0.005881506 |
| 58506 | *SCAF1* | ENSG00000126461.15 | 7.08 | 6.78 | 0.30 | 1.23 | 0.001679363 | 0.009363134 |
| 51719 | *CAB39* | ENSG00000135932.12 | 6.40 | 6.10 | 0.30 | 1.23 | 0.000350833 | 0.002928615 |
| 6272 | *SORT1* | ENSG00000134243.12 | 8.20 | 7.90 | 0.30 | 1.23 | 0.000292717 | 0.00256923 |
| 9246 | *UBE2L6* | ENSG00000156587.16 | 6.40 | 6.10 | 0.30 | 1.23 | 0.001241553 | 0.007421134 |
| 7289 | *TULP3* | ENSG00000078246.17 | 6.67 | 6.37 | 0.30 | 1.23 | 0.000901522 | 0.005834818 |
| 8242 | *KDM5C* | ENSG00000126012.13 | 7.35 | 7.05 | 0.30 | 1.23 | 0.000210705 | 0.00200875 |
| 805 | *CALM2* | ENSG00000143933.20 | 9.31 | 9.00 | 0.30 | 1.23 | 0.00010364 | 0.001197484 |
| 23181 | *DIP2A* | ENSG00000160305.18 | 6.27 | 5.97 | 0.30 | 1.23 | 0.001303429 | 0.007701791 |
| 25901 | *CCDC28A* | ENSG00000024862.18 | 5.29 | 4.99 | 0.30 | 1.23 | 0.000993974 | 0.006256565 |
| 24144 | *TFIP11* | ENSG00000100109.17 | 6.33 | 6.02 | 0.30 | 1.23 | 0.001368268 | 0.008022004 |
| 84232 | *MAF1* | ENSG00000179632.10 | 6.00 | 5.70 | 0.30 | 1.23 | 0.000965187 | 0.006112581 |
| 90550 | *MCU* | ENSG00000156026.15 | 5.80 | 5.50 | 0.30 | 1.23 | 0.000759296 | 0.005154353 |
| 24149 | *ZNF318* | ENSG00000171467.16 | 6.71 | 6.41 | 0.30 | 1.23 | 0.000622106 | 0.004452641 |
| 55161 | *TMEM33* | ENSG00000109133.13 | 8.13 | 7.84 | 0.30 | 1.23 | 0.000149227 | 0.001559484 |
| 10527 | *IPO7* | ENSG00000205339.10 | 7.88 | 7.58 | 0.30 | 1.23 | 0.000275569 | 0.002460395 |
| 7009 | *TMBIM6* | ENSG00000139644.13 | 11.20 | 10.90 | 0.30 | 1.23 | 0.00016727 | 0.001685862 |
| 9781 | *RNF144A* | ENSG00000151692.15 | 6.61 | 6.31 | 0.30 | 1.23 | 0.000727082 | 0.004987554 |
| 10440 | *TIMM17A* | ENSG00000134375.11 | 6.81 | 6.51 | 0.30 | 1.23 | 0.000890897 | 0.005788519 |
| 29956 | *CERS2* | ENSG00000143418.20 | 8.35 | 8.06 | 0.30 | 1.23 | 4.1705E-05 | 0.000628465 |
| 26123 | *TCTN3* | ENSG00000119977.22 | 6.47 | 6.17 | 0.30 | 1.23 | 0.0006506 | 0.004595227 |
| 8665 | *EIF3F* | ENSG00000175390.15 | 6.52 | 6.22 | 0.30 | 1.23 | 0.001519208 | 0.008673051 |
| 7469 | *NELFA* | ENSG00000185049.16 | 5.60 | 5.30 | 0.30 | 1.23 | 0.001399322 | 0.008159452 |
| 51203 | *NUSAP1* | ENSG00000137804.14 | 8.80 | 8.50 | 0.30 | 1.23 | 0.000933071 | 0.00597715 |
| 79875 | *THSD4* | ENSG00000187720.15 | 7.60 | 7.31 | 0.30 | 1.23 | 0.001447529 | 0.008351959 |
| 23673 | *STX12* | ENSG00000117758.14 | 6.41 | 6.11 | 0.30 | 1.23 | 0.001203809 | 0.007264968 |
| 5432 | *POLR2C* | ENSG00000102978.13 | 6.77 | 6.47 | 0.30 | 1.23 | 0.000882944 | 0.005754283 |
| 10149 | *ADGRG2* | ENSG00000173698.18 | 7.78 | 7.48 | 0.30 | 1.23 | 0.000626006 | 0.004469152 |
| 221092 | *HNRNPUL2* | ENSG00000214753.4 | 8.26 | 7.96 | 0.30 | 1.23 | 0.000349749 | 0.002922817 |
| 64131 | *XYLT1* | ENSG00000103489.12 | 7.47 | 7.17 | 0.30 | 1.23 | 0.000749719 | 0.005106597 |
| 83636 | *C19orf12* | ENSG00000131943.20 | 5.80 | 5.50 | 0.29 | 1.23 | 0.000950235 | 0.006048379 |
| 84726 | *PRRC2B* | ENSG00000288701.1 | 9.11 | 8.81 | 0.29 | 1.23 | 0.001177091 | 0.007138163 |
| 11143 | *KAT7* | ENSG00000136504.13 | 6.54 | 6.25 | 0.29 | 1.22 | 0.001157696 | 0.007033687 |
| 3313 | *HSPA9* | ENSG00000113013.16 | 9.55 | 9.25 | 0.29 | 1.22 | 9.68163E-05 | 0.001147046 |
| 2040 | *STOM* | ENSG00000148175.13 | 7.94 | 7.65 | 0.29 | 1.22 | 9.57224E-05 | 0.001136082 |
| 7965 | *AIMP2* | ENSG00000106305.10 | 5.26 | 4.96 | 0.29 | 1.22 | 0.001764307 | 0.009675001 |
| 60485 | *SAV1* | ENSG00000151748.15 | 6.45 | 6.16 | 0.29 | 1.22 | 0.000491194 | 0.0037423 |
| 10523 | *CHERP* | ENSG00000085872.15 | 6.66 | 6.37 | 0.29 | 1.22 | 0.001030426 | 0.006425289 |
| 6856 | *SYPL1* | ENSG00000008282.9 | 8.03 | 7.74 | 0.29 | 1.22 | 0.000265652 | 0.002401908 |
| 11057 | *ABHD2* | ENSG00000140526.18 | 9.79 | 9.50 | 0.29 | 1.22 | 0.000213018 | 0.002020981 |
| 348235 | *SKA2* | ENSG00000182628.13 | 7.29 | 6.99 | 0.29 | 1.22 | 0.000502227 | 0.003807036 |
| 593 | *BCKDHA* | ENSG00000248098.12 | 6.58 | 6.29 | 0.29 | 1.22 | 0.000885757 | 0.005762602 |
| 8933 | *RTL8C* | ENSG00000134590.14 | 6.60 | 6.31 | 0.29 | 1.22 | 0.000444267 | 0.003485623 |
| 9117 | *SEC22C* | ENSG00000093183.14 | 7.03 | 6.74 | 0.29 | 1.22 | 0.000366619 | 0.003020078 |
| 3183 | *HNRNPC* | ENSG00000092199.19 | 9.54 | 9.25 | 0.29 | 1.22 | 9.49253E-05 | 0.001128949 |
| 3710 | *ITPR3* | ENSG00000096433.11 | 6.05 | 5.77 | 0.29 | 1.22 | 0.000909819 | 0.005873321 |
| 10672 | *GNA13* | ENSG00000120063.10 | 7.59 | 7.30 | 0.29 | 1.22 | 0.000704906 | 0.004880042 |
| 4249 | *MGAT5* | ENSG00000152127.9 | 7.24 | 6.95 | 0.29 | 1.22 | 0.001487038 | 0.008534974 |
| 26001 | *RNF167* | ENSG00000108523.16 | 7.02 | 6.73 | 0.29 | 1.22 | 0.000667032 | 0.004678265 |
| 79692 | *ZNF322* | ENSG00000181315.11 | 6.45 | 6.16 | 0.29 | 1.22 | 0.001583175 | 0.008963362 |
| 51726 | *DNAJB11* | ENSG00000090520.12 | 6.35 | 6.06 | 0.29 | 1.22 | 0.001018559 | 0.006376652 |
| 51449 | *PCYOX1* | ENSG00000116005.12 | 7.71 | 7.42 | 0.29 | 1.22 | 0.000178546 | 0.001770008 |
| 6059 | *ABCE1* | ENSG00000164163.11 | 7.66 | 7.37 | 0.29 | 1.22 | 0.00047359 | 0.003643321 |
| 8804 | *CREG1* | ENSG00000143162.9 | 6.56 | 6.27 | 0.29 | 1.22 | 0.000545062 | 0.004051928 |
| 7705 | *ZNF146* | ENSG00000167635.12 | 7.60 | 7.31 | 0.29 | 1.22 | 0.000875306 | 0.005724516 |
| 6535 | *SLC6A8* | ENSG00000130821.17 | 6.50 | 6.22 | 0.29 | 1.22 | 0.001181119 | 0.007155151 |
| 567 | *B2M* | ENSG00000166710.22 | 10.48 | 10.20 | 0.29 | 1.22 | 0.00046699 | 0.003607332 |
| 1975 | *EIF4B* | ENSG00000063046.18 | 9.98 | 9.70 | 0.29 | 1.22 | 7.35303E-05 | 0.000940271 |

| | | | | | | | | |
|---|---|---|---|---|---|---|---|---|
| 3098 | *HK1* | ENSG00000156515.24 | 7.28 | 6.99 | 0.29 | 1.22 | 0.001285911 | 0.007649716 |
| 1398 | *CRK* | ENSG00000167193.8 | 6.96 | 6.67 | 0.29 | 1.22 | 0.000416441 | 0.00332466 |
| 3191 | *HNRNPL* | ENSG00000104824.18 | 8.87 | 8.58 | 0.28 | 1.22 | 0.000242388 | 0.002240148 |
| 57092 | *PCNP* | ENSG00000081154.12 | 8.26 | 7.97 | 0.28 | 1.22 | 0.000197457 | 0.001909555 |
| 7763 | *ZFAND5* | ENSG00000107372.13 | 8.74 | 8.45 | 0.28 | 1.22 | 0.000351225 | 0.002928707 |
| 4893 | *NRAS* | ENSG00000213281.5 | 7.59 | 7.31 | 0.28 | 1.22 | 0.000990997 | 0.006244277 |
| 10434 | *LYPLA1* | ENSG00000120992.18 | 6.98 | 6.69 | 0.28 | 1.22 | 0.001374914 | 0.00804212 |
| 283820 | *NOMO2* | ENSG00000185164.15 | 8.56 | 8.27 | 0.28 | 1.22 | 0.000122448 | 0.001354223 |
| 2923 | *PDIA3* | ENSG00000167004.14 | 10.02 | 9.74 | 0.28 | 1.22 | 7.54437E-05 | 0.000957173 |
| 26586 | *CKAP2* | ENSG00000136108.15 | 7.50 | 7.22 | 0.28 | 1.22 | 0.000143541 | 0.001516124 |
| 2664 | *GDI1* | ENSG00000203879.12 | 8.19 | 7.91 | 0.28 | 1.21 | 0.000155796 | 0.001610159 |
| 221937 | *FOXK1* | ENSG00000164916.11 | 6.75 | 6.47 | 0.28 | 1.21 | 0.001775204 | 0.009724102 |
| 5886 | *RAD23A* | ENSG00000179262.10 | 7.31 | 7.03 | 0.28 | 1.21 | 0.000401617 | 0.00322691 |
| 1386 | *ATF2* | ENSG00000115966.18 | 6.91 | 6.63 | 0.28 | 1.21 | 0.000247131 | 0.002275578 |
| 10652 | *YKT6* | ENSG00000106636.9 | 8.59 | 8.31 | 0.28 | 1.21 | 0.000311534 | 0.002676486 |
| 5660 | *PSAP* | ENSG00000197746.15 | 10.95 | 10.67 | 0.28 | 1.21 | 0.000773983 | 0.005223342 |
| 10906 | *TRAFD1* | ENSG00000135148.12 | 6.10 | 5.82 | 0.28 | 1.21 | 0.001429302 | 0.008276385 |
| 9975 | *NR1D2* | ENSG00000174738.13 | 8.25 | 7.98 | 0.28 | 1.21 | 0.00102822 | 0.006419534 |
| 29978 | *UBQLN2* | ENSG00000188021.9 | 6.65 | 6.38 | 0.28 | 1.21 | 0.000400904 | 0.003222908 |
| 28986 | *MAGEH1* | ENSG00000187601.5 | 6.31 | 6.04 | 0.28 | 1.21 | 0.001157988 | 0.007033687 |
| 3423 | *IDS* | ENSG00000010404.18 | 8.68 | 8.41 | 0.27 | 1.21 | 0.00088518 | 0.005761344 |
| 7534 | *YWHAZ* | ENSG00000164924.18 | 9.55 | 9.28 | 0.27 | 1.21 | 0.000164184 | 0.001667276 |
| 6926 | *TBX3* | ENSG00000135111.16 | 9.76 | 9.49 | 0.27 | 1.21 | 0.000169744 | 0.0017002 |
| 7879 | *RAB7A* | ENSG00000075785.14 | 8.29 | 8.02 | 0.27 | 1.21 | 0.000369442 | 0.003035007 |
| 48 | *ACO1* | ENSG00000122729.19 | 6.80 | 6.52 | 0.27 | 1.21 | 0.001632777 | 0.009178408 |
| 9554 | *SEC22B* | ENSG00000265808.4 | 7.36 | 7.09 | 0.27 | 1.21 | 0.0008418 | 0.005568348 |
| 10130 | *PDIA6* | ENSG00000143870.13 | 8.50 | 8.23 | 0.27 | 1.21 | 0.000337143 | 0.002834252 |
| 51604 | *PIGT* | ENSG00000124155.19 | 7.21 | 6.94 | 0.27 | 1.21 | 0.000914654 | 0.00588681 |
| 8887 | *TAX1BP1* | ENSG00000106052.14 | 8.29 | 8.01 | 0.27 | 1.21 | 0.000294248 | 0.002578148 |
| 81624 | *DIAPH3* | ENSG00000139734.19 | 6.27 | 6.00 | 0.27 | 1.21 | 0.001083723 | 0.006677178 |
| 83737 | *ITCH* | ENSG00000078747.16 | 7.22 | 6.95 | 0.27 | 1.21 | 0.0011926 | 0.007206968 |
| 219988 | *PATL1* | ENSG00000166889.14 | 7.26 | 6.99 | 0.27 | 1.21 | 0.000311993 | 0.002678802 |
| 10857 | *PGRMC1* | ENSG00000101856.10 | 8.13 | 7.85 | 0.27 | 1.21 | 0.001029401 | 0.006421856 |
| 537 | *ATP6AP1* | ENSG00000071553.18 | 8.74 | 8.47 | 0.27 | 1.21 | 0.000258434 | 0.002355096 |
| 7171 | *TPM4* | ENSG00000167460.17 | 9.09 | 8.82 | 0.27 | 1.21 | 0.000108119 | 0.001236856 |
| 6096 | *RORB* | ENSG00000198963.11 | 8.87 | 8.60 | 0.27 | 1.21 | 0.000876899 | 0.005727318 |
| 9403 | *SELENOF* | ENSG00000183291.18 | 8.13 | 7.86 | 0.27 | 1.21 | 0.00050921 | 0.003844432 |
| 10212 | *DDX39A* | ENSG00000123136.15 | 5.44 | 5.17 | 0.27 | 1.20 | 0.001489003 | 0.008536456 |
| 4055 | *LTBR* | ENSG00000111321.11 | 6.42 | 6.15 | 0.27 | 1.20 | 0.001819322 | 0.009922312 |
| 583 | *BBS2* | ENSG00000125124.14 | 7.52 | 7.25 | 0.27 | 1.20 | 0.000463767 | 0.003591681 |
| 7295 | *TXN* | ENSG00000136810.13 | 8.22 | 7.95 | 0.27 | 1.20 | 0.000911242 | 0.005874974 |
| 80204 | *FBXO11* | ENSG00000138081.22 | 6.52 | 6.26 | 0.27 | 1.20 | 0.00055846 | 0.004120906 |
| 51247 | *PAIP2* | ENSG00000120727.13 | 7.66 | 7.39 | 0.26 | 1.20 | 0.000514046 | 0.0038751 |
| 5577 | *PRKAR2B* | ENSG00000005249.13 | 8.61 | 8.34 | 0.26 | 1.20 | 0.00040393 | 0.003243761 |
| 10092 | *ARPC5* | ENSG00000162704.16 | 8.74 | 8.47 | 0.26 | 1.20 | 0.000504492 | 0.003816495 |
| 56949 | *XAB2* | ENSG00000076924.12 | 6.03 | 5.77 | 0.26 | 1.20 | 0.00142135 | 0.008253617 |
| 5253 | *PHF2* | ENSG00000197724.11 | 7.12 | 6.85 | 0.26 | 1.20 | 0.000891691 | 0.005791167 |
| 93663 | *ARHGAP18* | ENSG00000146376.11 | 8.08 | 7.82 | 0.26 | 1.20 | 0.00052682 | 0.003947599 |
| 80351 | *TNKS2* | ENSG00000107854.6 | 7.18 | 6.92 | 0.26 | 1.20 | 0.000337137 | 0.002834252 |
| 9528 | *TMEM59* | ENSG00000116209.12 | 8.01 | 7.75 | 0.26 | 1.20 | 0.000578061 | 0.004228127 |
| 6729 | *SRP54* | ENSG00000100883.13 | 7.35 | 7.09 | 0.26 | 1.20 | 0.001379178 | 0.008063921 |
| 1857 | *DVL3* | ENSG00000161202.20 | 7.57 | 7.31 | 0.26 | 1.20 | 0.00074895 | 0.005104901 |
| 6638 | *SNRPN* | ENSG00000128739.23 | 6.78 | 6.52 | 0.26 | 1.20 | 0.00158301 | 0.008963362 |
| 154807 | *VKORC1L1* | ENSG00000196715.7 | 7.18 | 6.92 | 0.26 | 1.20 | 0.001303901 | 0.007701791 |
| 8087 | *FXR1* | ENSG00000114416.18 | 7.83 | 7.57 | 0.26 | 1.20 | 0.000500184 | 0.003795382 |
| 3685 | *ITGAV* | ENSG00000138448.12 | 7.73 | 7.47 | 0.26 | 1.20 | 0.00058488 | 0.004261697 |
| 23568 | *ARL2BP* | ENSG00000102931.8 | 6.97 | 6.71 | 0.26 | 1.20 | 0.000645756 | 0.004568776 |
| 53339 | *BTBD1* | ENSG00000064726.10 | 7.15 | 6.89 | 0.26 | 1.20 | 0.000602765 | 0.004357854 |
| 8803 | *SUCLA2* | ENSG00000136143.16 | 6.19 | 5.93 | 0.26 | 1.20 | 0.000870767 | 0.005706625 |
| 84193 | *SETD3* | ENSG00000183576.13 | 6.20 | 5.95 | 0.26 | 1.20 | 0.001436558 | 0.00831199 |
| 351 | *APP* | ENSG00000142192.21 | 8.90 | 8.64 | 0.26 | 1.20 | 0.000240537 | 0.002225784 |
| 7088 | *TLE1* | ENSG00000196781.16 | 7.89 | 7.63 | 0.26 | 1.20 | 0.000606324 | 0.004375152 |
| 4102 | *MAGEA3* | ENSG00000221867.9 | 8.67 | 8.41 | 0.26 | 1.20 | 0.000902981 | 0.00584174 |
| 23291 | *FBXW11* | ENSG00000072803.18 | 7.14 | 6.88 | 0.26 | 1.20 | 0.001735909 | 0.009576773 |
| 3927 | *LASP1* | ENSG00000002834.18 | 7.66 | 7.40 | 0.26 | 1.20 | 0.001290951 | 0.007668246 |
| 4952 | *OCRL* | ENSG00000122126.18 | 7.10 | 6.84 | 0.26 | 1.20 | 0.001053847 | 0.006541437 |
| 8848 | *TSC22D1* | ENSG00000102804.15 | 6.60 | 6.35 | 0.26 | 1.19 | 0.001308336 | 0.007718871 |
| 4666 | *NACA* | ENSG00000196531.14 | 9.06 | 8.80 | 0.26 | 1.19 | 0.000285156 | 0.002519605 |
| 3607 | *FOXK2* | ENSG00000141568.21 | 7.07 | 6.81 | 0.26 | 1.19 | 0.001758909 | 0.009666591 |
| 5781 | *PTPN11* | ENSG00000179295.19 | 8.19 | 7.94 | 0.26 | 1.19 | 0.000470533 | 0.003627233 |
| 10605 | *PAIP1* | ENSG00000172239.14 | 7.66 | 7.40 | 0.26 | 1.19 | 0.00068681 | 0.004781227 |
| 10099 | *TSPAN3* | ENSG00000140391.15 | 8.59 | 8.34 | 0.25 | 1.19 | 0.000630937 | 0.004495746 |
| 444 | *ASPH* | ENSG00000198363.18 | 9.20 | 8.94 | 0.25 | 1.19 | 0.000416715 | 0.003325077 |
| 51142 | *CHCHD2* | ENSG00000106153.13 | 8.76 | 8.50 | 0.25 | 1.19 | 0.001345054 | 0.007910677 |
| 28972 | *SPCS1* | ENSG00000114902.14 | 7.43 | 7.17 | 0.25 | 1.19 | 0.001786586 | 0.009772185 |
| 26092 | *TOR1AIP1* | ENSG00000143337.19 | 8.01 | 7.75 | 0.25 | 1.19 | 0.001357179 | 0.00796633 |
| 10009 | *ZBTB33* | ENSG00000177485.7 | 6.53 | 6.28 | 0.25 | 1.19 | 0.00176223 | 0.009668126 |
| 91369 | *ANKRD40* | ENSG00000154945.7 | 6.55 | 6.29 | 0.25 | 1.19 | 0.000871968 | 0.005710018 |
| 9908 | *G3BP2* | ENSG00000138757.15 | 8.52 | 8.26 | 0.25 | 1.19 | 0.001683787 | 0.009377352 |
| 116150 | *NUS1* | ENSG00000153989.8 | 7.10 | 6.85 | 0.25 | 1.19 | 0.000624693 | 0.004464769 |
| 5547 | *PRCP* | ENSG00000137509.12 | 8.31 | 8.05 | 0.25 | 1.19 | 0.001255722 | 0.007496872 |
| 1736 | *DKC1* | ENSG00000130826.18 | 7.84 | 7.59 | 0.25 | 1.19 | 0.000256042 | 0.002337551 |
| 25793 | *FBXO7* | ENSG00000100225.18 | 7.58 | 7.33 | 0.25 | 1.19 | 0.000835266 | 0.00553733 |
| 10238 | *DCAF7* | ENSG00000136485.16 | 8.85 | 8.61 | 0.25 | 1.19 | 0.001525907 | 0.008696589 |
| 476 | *ATP1A1* | ENSG00000163399.16 | 9.38 | 9.13 | 0.25 | 1.19 | 0.001345063 | 0.007910677 |
| 3329 | *HSPD1* | ENSG00000144381.18 | 11.52 | 11.28 | 0.25 | 1.19 | 0.000205295 | 0.001970121 |
| 382 | *ARF6* | ENSG00000165527.8 | 7.63 | 7.39 | 0.25 | 1.19 | 0.000820337 | 0.005469762 |
| 387 | *RHOA* | ENSG00000067560.14 | 9.50 | 9.25 | 0.25 | 1.19 | 0.000405517 | 0.003251913 |
| 334 | *APLP2* | ENSG00000084234.18 | 10.04 | 9.80 | 0.24 | 1.18 | 0.000546263 | 0.00405683 |

| | | | | | | | | |
|---|---|---|---|---|---|---|---|---|
| 10899 | *JTB* | ENSG00000143543.15 | 6.53 | 6.28 | 0.24 | 1.18 | 0.001526205 | 0.008696589 |
| 1390 | *CREM* | ENSG00000095794.20 | 7.22 | 6.98 | 0.24 | 1.18 | 0.00170904 | 0.009483932 |
| 23028 | *KDM1A* | ENSG00000004487.18 | 7.62 | 7.38 | 0.24 | 1.18 | 0.000939287 | 0.005995238 |
| 5034 | *P4HB* | ENSG00000185624.16 | 9.65 | 9.41 | 0.24 | 1.18 | 0.000763655 | 0.005167585 |
| 4678 | *NASP* | ENSG00000132780.17 | 7.52 | 7.28 | 0.24 | 1.18 | 0.001709773 | 0.009483932 |
| 23071 | *ERP44* | ENSG00000023318.9 | 7.22 | 6.98 | 0.24 | 1.18 | 0.000987958 | 0.006227745 |
| 2632 | *GBE1* | ENSG00000114480.13 | 7.88 | 7.64 | 0.24 | 1.18 | 0.001337167 | 0.007870405 |
| 3777 | *KCNK3* | ENSG00000171303.8 | 9.63 | 9.39 | 0.24 | 1.18 | 0.000643672 | 0.004561986 |
| 2547 | *XRCC6* | ENSG00000196419.13 | 8.40 | 8.16 | 0.24 | 1.18 | 0.001216581 | 0.007315569 |
| 51150 | *SDF4* | ENSG00000078808.19 | 6.38 | 6.14 | 0.24 | 1.18 | 0.001293066 | 0.007669926 |
| 51552 | *RAB14* | ENSG00000119396.11 | 8.11 | 7.88 | 0.24 | 1.18 | 0.001664737 | 0.009309253 |
| 10140 | *TOB1* | ENSG00000141232.5 | 7.72 | 7.48 | 0.24 | 1.18 | 0.000745149 | 0.005085921 |
| 2052 | *EPHX1* | ENSG00000143819.13 | 9.68 | 9.44 | 0.23 | 1.18 | 0.001286775 | 0.007651826 |
| 10342 | *TFG* | ENSG00000114354.15 | 7.82 | 7.58 | 0.23 | 1.18 | 0.00050301 | 0.003811045 |
| 27173 | *SLC39A1* | ENSG00000143570.19 | 7.12 | 6.89 | 0.23 | 1.17 | 0.000996043 | 0.00626555 |
| 10955 | *SERINC3* | ENSG00000132824.14 | 8.39 | 8.16 | 0.23 | 1.17 | 0.001078456 | 0.006655649 |
| 9295 | *SRSF11* | ENSG00000116754.14 | 7.75 | 7.52 | 0.23 | 1.17 | 0.001283296 | 0.007640215 |
| 94239 | *H2AZ2* | ENSG00000105968.19 | 9.84 | 9.61 | 0.23 | 1.17 | 0.000846766 | 0.00559626 |
| 130074 | *FAM168B* | ENSG00000152102.18 | 8.30 | 8.07 | 0.23 | 1.17 | 0.000805574 | 0.005390484 |
| 11171 | *STRAP* | ENSG00000023734.11 | 8.01 | 7.78 | 0.23 | 1.17 | 0.001007657 | 0.00632271 |
| 27069 | *GHITM* | ENSG00000165678.22 | 8.83 | 8.60 | 0.23 | 1.17 | 0.001404471 | 0.008186295 |
| 7514 | *XPO1* | ENSG00000082898.19 | 8.31 | 8.08 | 0.23 | 1.17 | 0.001233063 | 0.00739298 |
| 5355 | *PLP2* | ENSG00000102007.11 | 7.41 | 7.18 | 0.23 | 1.17 | 0.00171226 | 0.009486643 |
| 113201 | *GOLM2* | ENSG00000166734.20 | 8.79 | 8.56 | 0.22 | 1.17 | 0.001229375 | 0.007377724 |
| 226 | *ALDOA* | ENSG00000149925.22 | 9.42 | 9.19 | 0.22 | 1.17 | 0.001736259 | 0.009576773 |
| 7416 | *VDAC1* | ENSG00000213585.11 | 8.17 | 7.95 | 0.22 | 1.17 | 0.001667844 | 0.009319686 |
| 506 | *ATP5F1B* | ENSG00000110955.9 | 10.52 | 10.29 | 0.22 | 1.17 | 0.00094475 | 0.006021128 |
| 26035 | *GLCE* | ENSG00000138604.10 | 7.57 | 7.34 | 0.22 | 1.17 | 0.001407213 | 0.008196312 |
| 647979 | *NORAD* | ENSG00000260032.2 | 9.51 | 9.29 | 0.22 | 1.17 | 0.001721954 | 0.009533314 |
| 10313 | *RTN3* | ENSG00000133318.14 | 8.14 | 7.92 | 0.22 | 1.17 | 0.001688602 | 0.009393709 |
| 10618 | *TGOLN2* | ENSG00000152291.15 | 8.55 | 8.34 | 0.22 | 1.16 | 0.001810299 | 0.009887473 |
| 133686 | *NADK2* | ENSG00000152620.13 | 7.73 | 7.52 | 0.22 | 1.16 | 0.001818381 | 0.009922312 |
| 9500 | *MAGED1* | ENSG00000179222.18 | 8.54 | 8.32 | 0.21 | 1.16 | 0.000858862 | 0.005658762 |
| 2778 | *GNAS* | ENSG00000087460.29 | 10.03 | 9.82 | 0.21 | 1.16 | 0.001112933 | 0.006826344 |
| 10971 | *YWHAQ* | ENSG00000134308.14 | 9.39 | 9.18 | 0.21 | 1.16 | 0.000993354 | 0.006256507 |
| 80829 | *ZFP91* | ENSG00000186660.15 | 8.19 | 7.98 | 0.21 | 1.16 | 0.001606302 | 0.009060124 |
| 5298 | *PI4KB* | ENSG00000143393.17 | 7.48 | 7.27 | 0.21 | 1.16 | 0.001787923 | 0.009775932 |
| 10953 | *TOMM34* | ENSG00000025772.8 | 7.52 | 7.32 | 0.21 | 1.15 | 0.001736823 | 0.009576773 |
| 26112 | *CCDC69* | ENSG00000198624.13 | 8.51 | 8.31 | 0.20 | 1.15 | 0.001797758 | 0.009822552 |
| 23466 | *CBX6* | ENSG00000183741.12 | 6.93 | 7.14 | -0.21 | 0.87 | 0.001762409 | 0.009668126 |
| 9375 | *TM9SF2* | ENSG00000125304.10 | 9.56 | 9.78 | -0.21 | 0.86 | 0.001204985 | 0.00726914 |
| 23511 | *NUP188* | ENSG00000095319.14 | 7.56 | 7.78 | -0.22 | 0.86 | 0.001078106 | 0.006655649 |
| 3178 | *HNRNPA1* | ENSG00000135486.19 | 10.59 | 10.81 | -0.22 | 0.86 | 0.001235687 | 0.007397859 |
| 1434 | *CSE1L* | ENSG00000124207.17 | 7.53 | 7.76 | -0.23 | 0.85 | 0.001504001 | 0.008606003 |
| 23500 | *DAAM2* | ENSG00000146122.17 | 8.02 | 8.25 | -0.23 | 0.85 | 0.001773512 | 0.009718381 |
| 140735 | *DYNLL2* | ENSG00000264364.3 | 8.62 | 8.85 | -0.23 | 0.85 | 0.001386529 | 0.008091139 |
| 9782 | *MATR3* | ENSG00000015479.20 | 8.76 | 9.00 | -0.24 | 0.85 | 0.000695375 | 0.004821034 |
| 5095 | *PCCA* | ENSG00000175198.17 | 6.85 | 7.09 | -0.24 | 0.85 | 0.001264045 | 0.007543559 |
| 6602 | *SMARCD1* | ENSG00000066117.15 | 7.47 | 7.71 | -0.24 | 0.85 | 0.001027196 | 0.006415808 |
| 51666 | *ASB4* | ENSG00000005981.13 | 8.98 | 9.22 | -0.24 | 0.85 | 0.000289234 | 0.002547218 |
| 51701 | *NLK* | ENSG00000087095.13 | 7.41 | 7.65 | -0.24 | 0.85 | 0.001539063 | 0.008759877 |
| 1031 | *CDKN2C* | ENSG00000123080.12 | 7.52 | 7.76 | -0.24 | 0.85 | 0.000535176 | 0.003990288 |
| 23270 | *TSPYL4* | ENSG00000187189.11 | 7.11 | 7.35 | -0.24 | 0.84 | 0.000798087 | 0.005344437 |
| 11180 | *WDR6* | ENSG00000178252.19 | 7.29 | 7.54 | -0.24 | 0.84 | 0.000501933 | 0.003806728 |
| 23607 | *CD2AP* | ENSG00000198087.7 | 6.74 | 6.99 | -0.25 | 0.84 | 0.001380012 | 0.008065654 |
| 118 | *ADD1* | ENSG00000087274.19 | 7.61 | 7.86 | -0.25 | 0.84 | 0.001458044 | 0.008397458 |
| 149603 | *RNF187* | ENSG00000168159.15 | 7.73 | 7.98 | -0.25 | 0.84 | 0.000898629 | 0.005823626 |
| 26985 | *AP3M1* | ENSG00000185009.13 | 6.71 | 6.96 | -0.25 | 0.84 | 0.001422185 | 0.008253617 |
| 178 | *AGL* | ENSG00000162688.17 | 6.14 | 6.39 | -0.25 | 0.84 | 0.000897037 | 0.005818336 |
| 8125 | *ANP32A* | ENSG00000140350.15 | 7.50 | 7.75 | -0.25 | 0.84 | 0.001589495 | 0.008982205 |
| 1933 | *EEF1B2* | ENSG00000114942.14 | 7.03 | 7.28 | -0.25 | 0.84 | 0.001500738 | 0.008593885 |
| 9352 | *TXNL1* | ENSG00000091164.13 | 6.17 | 6.42 | -0.25 | 0.84 | 0.00093949 | 0.005995238 |
| 7343 | *UBTF* | ENSG00000108312.15 | 6.93 | 7.18 | -0.25 | 0.84 | 0.000643995 | 0.004561986 |
| 824 | *CAPN2* | ENSG00000162909.18 | 6.90 | 7.15 | -0.25 | 0.84 | 0.001551828 | 0.008825837 |
| 10220 | *GDF11* | ENSG00000135414.10 | 7.70 | 7.96 | -0.25 | 0.84 | 0.001280376 | 0.00762586 |
| 7529 | *YWHAB* | ENSG00000166913.13 | 8.28 | 8.54 | -0.26 | 0.84 | 0.000505881 | 0.003823148 |
| 91775 | *NXPE3* | ENSG00000144815.17 | 6.89 | 7.15 | -0.26 | 0.84 | 0.00113851 | 0.006938678 |
| 11130 | *ZWINT* | ENSG00000122952.17 | 6.43 | 6.69 | -0.26 | 0.84 | 0.000708242 | 0.004898623 |
| 140885 | *SIRPA* | ENSG00000198053.12 | 6.60 | 6.86 | -0.26 | 0.84 | 0.000554227 | 0.004097729 |
| 5962 | *RDX* | ENSG00000137710.17 | 9.27 | 9.53 | -0.26 | 0.83 | 0.000208821 | 0.001996299 |
| 6950 | *TCP1* | ENSG00000120438.12 | 7.67 | 7.93 | -0.26 | 0.83 | 0.000306703 | 0.002657796 |
| 3151 | *HMGN2* | ENSG00000198830.11 | 9.12 | 9.39 | -0.26 | 0.83 | 0.000636406 | 0.004523765 |
| 29 | *ABR* | ENSG00000159842.16 | 7.50 | 7.77 | -0.26 | 0.83 | 0.000812927 | 0.005430002 |
| 4199 | *ME1* | ENSG00000065833.9 | 6.64 | 6.91 | -0.26 | 0.83 | 0.00148873 | 0.008536456 |
| 4257 | *MGST1* | ENSG00000008394.13 | 6.09 | 6.36 | -0.27 | 0.83 | 0.001474329 | 0.008475777 |
| 9791 | *PTDSS1* | ENSG00000156471.13 | 6.53 | 6.79 | -0.27 | 0.83 | 0.001601818 | 0.00903823 |
| 39 | *ACAT2* | ENSG00000120437.9 | 6.48 | 6.75 | -0.27 | 0.83 | 0.001137168 | 0.00693828 |
| 5705 | *PSMC5* | ENSG00000087191.13 | 6.40 | 6.67 | -0.27 | 0.83 | 0.000739433 | 0.0050538 |
| 55243 | *KIRREL1* | ENSG00000183853.18 | 6.73 | 7.00 | -0.27 | 0.83 | 0.000890621 | 0.005788519 |
| 646719 | *NIPBL-DT* | ENSG00000285967.1 | 5.48 | 5.75 | -0.27 | 0.83 | 0.001147315 | 0.00698582 |
| 220988 | *HNRNPA3* | ENSG00000170144.21 | 9.37 | 9.64 | -0.27 | 0.83 | 0.000680135 | 0.004742573 |
| 89891 | *DYNC2I2* | ENSG00000119333.12 | 5.59 | 5.86 | -0.27 | 0.83 | 0.001635377 | 0.009189468 |
| 3434 | *IFIT1* | ENSG00000185745.10 | 4.77 | 5.04 | -0.27 | 0.83 | 0.001683517 | 0.009377352 |
| 3192 | *HNRNPU* | ENSG00000153187.20 | 9.23 | 9.50 | -0.27 | 0.83 | 0.000267764 | 0.00241084 |
| 2058 | *EPRS1* | ENSG00000136628.18 | 11.19 | 11.46 | -0.27 | 0.83 | 0.000108912 | 0.001243092 |
| 10745 | *PHTF1* | ENSG00000116793.16 | 5.53 | 5.80 | -0.27 | 0.83 | 0.001293396 | 0.007669926 |
| 6421 | *SFPQ* | ENSG00000116560.11 | 8.38 | 8.66 | -0.27 | 0.83 | 0.000245539 | 0.002263987 |
| 9371 | *KIF3B* | ENSG00000101350.8 | 6.71 | 6.99 | -0.28 | 0.83 | 0.001354597 | 0.007954285 |
| 1639 | *DCTN1* | ENSG00000204843.13 | 7.51 | 7.78 | -0.28 | 0.83 | 0.000331575 | 0.002803207 |

| | | | | | | | | |
|---|---|---|---|---|---|---|---|---|
| 23339 | *VPS39* | ENSG00000166887.16 | 6.84 | 7.12 | -0.28 | 0.82 | 0.000539199 | 0.004016299 |
| 6434 | *TRA2B* | ENSG00000136527.19 | 7.26 | 7.54 | -0.28 | 0.82 | 0.000266145 | 0.002404923 |
| 25929 | *GEMIN5* | ENSG00000082516.9 | 5.57 | 5.85 | -0.28 | 0.82 | 0.00118201 | 0.007156431 |
| 79142 | *PHF23* | ENSG00000040633.13 | 5.95 | 6.23 | -0.28 | 0.82 | 0.001327408 | 0.007816035 |
| 59349 | *KLHL12* | ENSG00000117153.16 | 5.53 | 5.81 | -0.28 | 0.82 | 0.000618717 | 0.004432609 |
| 10928 | *RALBP1* | ENSG00000017797.13 | 7.87 | 8.15 | -0.29 | 0.82 | 0.000332686 | 0.002809955 |
| 4715 | *NDUFB9* | ENSG00000147684.10 | 7.08 | 7.37 | -0.29 | 0.82 | 0.000445101 | 0.003488523 |
| 27350 | *APOBEC3C* | ENSG00000244509.4 | 5.72 | 6.01 | -0.29 | 0.82 | 0.001669935 | 0.009327898 |
| 50512 | *PODXL2* | ENSG00000114631.11 | 5.37 | 5.66 | -0.29 | 0.82 | 0.000725235 | 0.004981718 |
| 55173 | *MRPS10* | ENSG00000048544.6 | 5.33 | 5.62 | -0.29 | 0.82 | 0.000884681 | 0.005760598 |
| 5865 | *RAB3B* | ENSG00000169213.7 | 7.65 | 7.94 | -0.29 | 0.82 | 0.000232931 | 0.002166085 |
| 10488 | *CREB3* | ENSG00000107175.12 | 6.70 | 6.99 | -0.29 | 0.82 | 0.00028408 | 0.002514403 |
| 8220 | *ESS2* | ENSG00000100056.12 | 4.98 | 5.27 | -0.29 | 0.82 | 0.001272224 | 0.007583327 |
| 124222 | *PAQR4* | ENSG00000162073.14 | 6.13 | 6.42 | -0.29 | 0.82 | 0.000193406 | 0.001877641 |
| 9373 | *PLAA* | ENSG00000137055.15 | 5.88 | 6.17 | -0.29 | 0.82 | 0.000444861 | 0.003488464 |
| 9779 | *TBC1D5* | ENSG00000131374.14 | 6.77 | 7.06 | -0.29 | 0.82 | 0.000211726 | 0.00201381 |
| 7077 | *TIMP2* | ENSG00000035862.12 | 6.80 | 7.09 | -0.30 | 0.81 | 0.000675388 | 0.004721422 |
| 6611 | *SMS* | ENSG00000102172.16 | 9.05 | 9.35 | -0.30 | 0.81 | 0.000167362 | 0.001685862 |
| 7389 | *UROD* | ENSG00000126088.14 | 6.35 | 6.65 | -0.30 | 0.81 | 0.001639699 | 0.009202527 |
| 29960 | *MRM2* | ENSG00000122687.19 | 5.37 | 5.67 | -0.30 | 0.81 | 0.00088656 | 0.005765329 |
| 25946 | *ZNF385A* | ENSG00000161642.18 | 7.47 | 7.77 | -0.30 | 0.81 | 0.000132707 | 0.001427812 |
| 26227 | *PHGDH* | ENSG00000092621.13 | 7.47 | 7.76 | -0.30 | 0.81 | 0.00074683 | 0.005092762 |
| 22858 | *CILK1* | ENSG00000112144.16 | 6.26 | 6.56 | -0.30 | 0.81 | 0.000325933 | 0.002762239 |
| 1432 | *MAPK14* | ENSG00000112062.11 | 7.19 | 7.49 | -0.30 | 0.81 | 0.001295437 | 0.007675962 |
| 51535 | *PPHLN1* | ENSG00000134283.18 | 6.97 | 7.27 | -0.30 | 0.81 | 0.000576635 | 0.004221813 |
| 78988 | *MRPL57* | ENSG00000173141.5 | 5.39 | 5.69 | -0.30 | 0.81 | 0.000522466 | 0.003924768 |
| 6414 | *SELENOP* | ENSG00000250722.6 | 5.95 | 6.25 | -0.30 | 0.81 | 0.001066613 | 0.006609745 |
| 90701 | *SEC11C* | ENSG00000166562.9 | 5.33 | 5.63 | -0.30 | 0.81 | 0.001029448 | 0.006421856 |
| 79590 | *MRPL24* | ENSG00000143314.12 | 5.91 | 6.21 | -0.30 | 0.81 | 0.000976011 | 0.006165213 |
| 5306 | *PITPNA* | ENSG00000174238.16 | 6.74 | 7.04 | -0.30 | 0.81 | 0.000604972 | 0.004369598 |
| 4171 | *MCM2* | ENSG00000073111.14 | 7.07 | 7.37 | -0.30 | 0.81 | 0.00013015 | 0.001410398 |
| 27115 | *PDE7B* | ENSG00000171408.14 | 5.32 | 5.63 | -0.30 | 0.81 | 0.000965209 | 0.006112581 |
| 9063 | *PIAS2* | ENSG00000078043.16 | 5.51 | 5.82 | -0.31 | 0.81 | 0.001187508 | 0.007183921 |
| 11078 | *TRIOBP* | ENSG00000100106.22 | 5.67 | 5.97 | -0.31 | 0.81 | 0.000711242 | 0.004914011 |
| 1499 | *CTNNB1* | ENSG00000168036.18 | 9.03 | 9.34 | -0.31 | 0.81 | 4.63167E-05 | 0.000677551 |
| 50650 | *ARHGEF3* | ENSG00000163947.12 | 6.62 | 6.93 | -0.31 | 0.81 | 0.00016395 | 0.001667158 |
| 5325 | *PLAGL1* | ENSG00000118495.21 | 7.62 | 7.93 | -0.31 | 0.81 | 0.001038318 | 0.006469121 |
| 7913 | *DEK* | ENSG00000124795.17 | 8.04 | 8.35 | -0.31 | 0.81 | 0.000140161 | 0.001494042 |
| 7818 | *DAP3* | ENSG00000132676.16 | 6.75 | 7.06 | -0.31 | 0.81 | 0.000382128 | 0.003101874 |
| 222553 | *SLC35F1* | ENSG00000196376.11 | 5.86 | 6.16 | -0.31 | 0.81 | 0.000302229 | 0.00263271 |
| 8323 | *FZD6* | ENSG00000164930.12 | 5.90 | 6.21 | -0.31 | 0.81 | 0.001817132 | 0.009921182 |
| 55238 | *SLC38A7* | ENSG00000103042.9 | 5.57 | 5.88 | -0.31 | 0.81 | 0.000999092 | 0.006279467 |
| 10969 | *EBNA1BP2* | ENSG00000117395.13 | 5.87 | 6.18 | -0.31 | 0.81 | 0.0008815 | 0.005749859 |
| 23654 | *PLXNB2* | ENSG00000196576.16 | 6.09 | 6.41 | -0.31 | 0.80 | 0.001474466 | 0.008475777 |
| 8805 | *TRIM24* | ENSG00000122779.18 | 7.43 | 7.75 | -0.31 | 0.80 | 8.15312E-05 | 0.001010488 |
| 2647 | *BLOC1S1* | ENSG00000135441.8 | 6.03 | 6.34 | -0.31 | 0.80 | 0.00074643 | 0.005092349 |
| 85458 | *DIXDC1* | ENSG00000150764.14 | 5.54 | 5.85 | -0.31 | 0.80 | 0.000505311 | 0.003820759 |
| 3337 | *DNAJB1* | ENSG00000132002.9 | 7.18 | 7.49 | -0.31 | 0.80 | 0.000955098 | 0.006074179 |
| 11112 | *HIBADH* | ENSG00000106049.9 | 6.75 | 7.07 | -0.32 | 0.80 | 0.000195669 | 0.001894704 |
| 7027 | *TFDP1* | ENSG00000198176.13 | 8.01 | 8.32 | -0.32 | 0.80 | 4.90278E-05 | 0.000704845 |
| 11331 | *PHB2* | ENSG00000215021.10 | 7.06 | 7.37 | -0.32 | 0.80 | 0.000249878 | 0.002295237 |
| 11338 | *U2AF2* | ENSG00000063244.13 | 7.41 | 7.73 | -0.32 | 0.80 | 0.000281559 | 0.002497588 |
| 84298 | *LLPH* | ENSG00000139233.7 | 6.10 | 6.42 | -0.32 | 0.80 | 0.000794322 | 0.005327068 |
| 2067 | *ERCC1* | ENSG00000012061.16 | 5.30 | 5.63 | -0.32 | 0.80 | 0.000879786 | 0.005741177 |
| 10382 | *TUBB4A* | ENSG00000104833.12 | 6.37 | 6.69 | -0.32 | 0.80 | 0.000607571 | 0.004382043 |
| 1836 | *SLC26A2* | ENSG00000155850.9 | 5.72 | 6.04 | -0.32 | 0.80 | 0.000492449 | 0.003748059 |
| 140465 | *MYL6B* | ENSG00000196465.10 | 5.25 | 5.58 | -0.32 | 0.80 | 0.001518041 | 0.008669814 |
| 55588 | *MED29* | ENSG00000063322.15 | 6.09 | 6.41 | -0.32 | 0.80 | 0.000956512 | 0.006076651 |
| 100505876 | *CEBPZOS* | ENSG00000218739.10 | 5.27 | 5.59 | -0.32 | 0.80 | 0.001381229 | 0.008069628 |
| 54888 | *NSUN2* | ENSG00000037474.15 | 6.00 | 6.33 | -0.32 | 0.80 | 0.0001041 | 0.001200024 |
| 9797 | *TATDN2* | ENSG00000157014.11 | 6.51 | 6.83 | -0.32 | 0.80 | 0.000475384 | 0.003652825 |
| 55000 | *TUG1* | ENSG00000253352.10 | 8.20 | 8.53 | -0.33 | 0.80 | 0.0001829 | 0.001801278 |
| 1466 | *CSRP2* | ENSG00000175183.10 | 5.20 | 5.52 | -0.33 | 0.80 | 0.000757594 | 0.005147455 |
| 2260 | *FGFR1* | ENSG00000077782.22 | 7.18 | 7.51 | -0.33 | 0.80 | 0.001561208 | 0.008869101 |
| 8078 | *USP5* | ENSG00000111667.14 | 7.99 | 8.32 | -0.33 | 0.80 | 0.00012502 | 0.001371089 |
| 9937 | *DCLRE1A* | ENSG00000198924.8 | 4.66 | 4.98 | -0.33 | 0.80 | 0.000616997 | 0.004425886 |
| 162494 | *RHBDL3* | ENSG00000141314.13 | 6.37 | 6.70 | -0.33 | 0.80 | 4.76914E-05 | 0.000691184 |
| 115265 | *DDIT4L* | ENSG00000145358.6 | 6.71 | 7.04 | -0.33 | 0.80 | 7.28629E-05 | 0.000936301 |
| 85406 | *DNAJC14* | ENSG00000135392.19 | 6.47 | 6.80 | -0.33 | 0.80 | 0.000122517 | 0.001354223 |
| 9343 | *EFTUD2* | ENSG00000108883.13 | 7.53 | 7.87 | -0.33 | 0.80 | 0.000377754 | 0.003075272 |
| 80851 | *SH3BP5L* | ENSG00000175137.11 | 5.41 | 5.75 | -0.33 | 0.80 | 0.000176864 | 0.001757985 |
| 2516 | *NR5A1* | ENSG00000136931.10 | 8.45 | 8.78 | -0.33 | 0.79 | 0.000267392 | 0.002410381 |
| 9765 | *ZFYVE16* | ENSG00000039319.17 | 6.65 | 6.98 | -0.33 | 0.79 | 4.94293E-05 | 0.000709162 |
| 54859 | *ELP6* | ENSG00000163832.16 | 5.19 | 5.52 | -0.33 | 0.79 | 0.001081455 | 0.00667142 |
| 54946 | *SLC41A3* | ENSG00000114544.17 | 5.97 | 6.30 | -0.33 | 0.79 | 0.001698698 | 0.009442871 |
| 7559 | *ZNF12* | ENSG00000164631.19 | 5.18 | 5.51 | -0.33 | 0.79 | 0.000531611 | 0.003971605 |
| 57542 | *KLHL42* | ENSG00000087448.11 | 6.59 | 6.93 | -0.33 | 0.79 | 5.43417E-05 | 0.000755899 |
| 231 | *AKR1B1* | ENSG00000085662.14 | 6.35 | 6.68 | -0.34 | 0.79 | 0.000277492 | 0.002471739 |
| 57226 | *LYRM2* | ENSG00000083099.11 | 5.71 | 6.04 | -0.34 | 0.79 | 0.000526057 | 0.003945794 |
| 4605 | *MYBL2* | ENSG00000101057.16 | 6.15 | 6.48 | -0.34 | 0.79 | 0.00015581 | 0.001610159 |
| 10154 | *PLXNC1* | ENSG00000136040.9 | 6.69 | 7.03 | -0.34 | 0.79 | 0.001269399 | 0.007569494 |
| 1376 | *CPT2* | ENSG00000157184.7 | 5.49 | 5.83 | -0.34 | 0.79 | 0.000420863 | 0.003342186 |
| 9631 | *NUP155* | ENSG00000113569.16 | 6.85 | 7.19 | -0.34 | 0.79 | 0.000177549 | 0.001761287 |
| 93622 | *LOC93622* | ENSG00000170846.17 | 4.42 | 4.77 | -0.34 | 0.79 | 0.001581836 | 0.008963362 |
| 51435 | *SCARA3* | ENSG00000168077.14 | 4.86 | 5.21 | -0.34 | 0.79 | 0.000729163 | 0.004997264 |
| 781 | *CACNA2D1* | ENSG00000153956.16 | 7.02 | 7.36 | -0.34 | 0.79 | 0.001090584 | 0.00670843 |
| 79710 | *MORC4* | ENSG00000133131.15 | 4.96 | 5.30 | -0.35 | 0.79 | 0.000513615 | 0.003873792 |
| 835 | *CASP2* | ENSG00000106144.20 | 6.81 | 7.15 | -0.35 | 0.79 | 0.000452999 | 0.003533816 |
| 51300 | *TIMMDC1* | ENSG00000113845.10 | 5.39 | 5.74 | -0.35 | 0.79 | 0.001562318 | 0.008872053 |

| | | | | | | | | |
|---|---|---|---|---|---|---|---|---|
| 4940 | *OAS3* | ENSG00000111331.14 | 6.01 | 6.36 | -0.35 | 0.79 | 0.000130687 | 0.001412634 |
| 9315 | *NREP* | ENSG00000134986.14 | 8.09 | 8.44 | -0.35 | 0.79 | 0.000688396 | 0.004790047 |
| 9324 | *HMGN3* | ENSG00000118418.14 | 6.29 | 6.64 | -0.35 | 0.78 | 3.71862E-05 | 0.000574797 |
| 9459 | *ARHGEF6* | ENSG00000129675.16 | 6.97 | 7.32 | -0.35 | 0.78 | 2.27495E-05 | 0.000396165 |
| 11169 | *WDHD1* | ENSG00000198554.12 | 5.37 | 5.72 | -0.35 | 0.78 | 0.000608018 | 0.004383164 |
| 63979 | *FIGNL1* | ENSG00000132436.12 | 6.57 | 6.93 | -0.35 | 0.78 | 0.000146126 | 0.00153693 |
| 57325 | *KAT14* | ENSG00000149474.15 | 5.09 | 5.45 | -0.35 | 0.78 | 0.00077194 | 0.005211897 |
| 2029 | *ENSA* | ENSG00000143420.19 | 7.45 | 7.81 | -0.36 | 0.78 | 5.42093E-05 | 0.000754819 |
| 8540 | *AGPS* | ENSG00000018510.18 | 7.39 | 7.75 | -0.36 | 0.78 | 8.45047E-05 | 0.001035372 |
| 7425 | *VGF* | ENSG00000128564.8 | 5.81 | 6.17 | -0.36 | 0.78 | 0.001393369 | 0.008127895 |
| 55827 | *DCAF6* | ENSG00000143164.16 | 5.70 | 6.06 | -0.36 | 0.78 | 0.000335215 | 0.002824952 |
| 9830 | *TRIM14* | ENSG00000106785.15 | 5.40 | 5.76 | -0.36 | 0.78 | 0.000187566 | 0.001835191 |
| 8899 | *PRPF4B* | ENSG00000112739.17 | 6.28 | 6.64 | -0.36 | 0.78 | 0.000161377 | 0.001646574 |
| 10528 | *NOP56* | ENSG00000101361.17 | 6.25 | 6.61 | -0.36 | 0.78 | 0.000303298 | 0.002637426 |
| 57153 | *SLC44A2* | ENSG00000129353.15 | 6.41 | 6.77 | -0.36 | 0.78 | 0.000882369 | 0.005753034 |
| 23461 | *ABCA5* | ENSG00000154265.16 | 5.53 | 5.90 | -0.36 | 0.78 | 0.000719629 | 0.004952274 |
| 1522 | *CTSZ* | ENSG00000101160.15 | 5.33 | 5.69 | -0.36 | 0.78 | 0.00113276 | 0.006927016 |
| 90324 | *CCDC97* | ENSG00000142039.4 | 4.78 | 5.14 | -0.36 | 0.78 | 0.000275727 | 0.002460395 |
| 7779 | *SLC30A1* | ENSG00000170385.10 | 11.39 | 11.76 | -0.36 | 0.78 | 0.000153489 | 0.001592657 |
| 4841 | *NONO* | ENSG00000147140.17 | 8.67 | 9.03 | -0.37 | 0.78 | 2.83021E-05 | 0.000468611 |
| 92399 | *MRRF* | ENSG00000148187.18 | 4.70 | 5.07 | -0.37 | 0.78 | 0.000941392 | 0.006004822 |
| 2941 | *GSTA4* | ENSG00000170899.11 | 6.32 | 6.69 | -0.37 | 0.78 | 0.000119674 | 0.001338711 |
| 29085 | *PHPT1* | ENSG00000054148.18 | 5.22 | 5.59 | -0.37 | 0.78 | 0.000289881 | 0.002550305 |
| 65082 | *VPS33A* | ENSG00000139719.11 | 5.08 | 5.45 | -0.37 | 0.77 | 0.001352981 | 0.007951016 |
| 100506668 | *NRAV* | ENSG00000248008.4 | 3.41 | 3.78 | -0.37 | 0.77 | 0.001458702 | 0.008398026 |
| 7174 | *TPP2* | ENSG00000134900.12 | 6.27 | 6.64 | -0.37 | 0.77 | 0.000176187 | 0.00175241 |
| 8482 | *SEMA7A* | ENSG00000138623.10 | 5.89 | 6.26 | -0.37 | 0.77 | 0.000124113 | 0.001365695 |
| 221154 | *MICU2* | ENSG00000165487.14 | 5.55 | 5.92 | -0.37 | 0.77 | 0.000418313 | 0.003330751 |
| 388969 | *C2orf68* | ENSG00000168887.11 | 4.91 | 5.28 | -0.37 | 0.77 | 0.00113911 | 0.006938678 |
| 284716 | *RIMKLA* | ENSG00000177181.15 | 5.14 | 5.51 | -0.37 | 0.77 | 0.001411829 | 0.008211821 |
| 1967 | *EIF2B1* | ENSG00000111361.13 | 5.62 | 5.99 | -0.37 | 0.77 | 0.000164163 | 0.001667276 |
| 493 | *ATP2B4* | ENSG00000058668.15 | 7.35 | 7.72 | -0.37 | 0.77 | 0.000654264 | 0.004612921 |
| 5445 | *PON2* | ENSG00000105854.13 | 6.79 | 7.17 | -0.37 | 0.77 | 5.22934E-05 | 0.000734898 |
| 25879 | *DCAF13* | ENSG00000164934.14 | 4.80 | 5.17 | -0.37 | 0.77 | 0.001619318 | 0.009123252 |
| 57643 | *ZSWIM5* | ENSG00000162415.7 | 5.33 | 5.70 | -0.38 | 0.77 | 7.76388E-05 | 0.000977402 |
| 4858 | *NOVA2* | ENSG00000104967.8 | 4.95 | 5.33 | -0.38 | 0.77 | 0.000770415 | 0.005203945 |
| 10109 | *ARPC2* | ENSG00000163466.16 | 6.31 | 6.69 | -0.38 | 0.77 | 0.000117064 | 0.001315134 |
| 51315 | *KRCC1* | ENSG00000172086.9 | 3.87 | 4.24 | -0.38 | 0.77 | 0.000697866 | 0.004833535 |
| 9962 | *SLC23A2* | ENSG00000089057.15 | 8.59 | 8.97 | -0.38 | 0.77 | 9.37285E-06 | 0.000202705 |
| 388021 | *TMEM179* | ENSG00000258986.7 | 6.24 | 6.62 | -0.38 | 0.77 | 0.000600891 | 0.004346996 |
| 55086 | *RADX* | ENSG00000147231.14 | 5.60 | 5.98 | -0.38 | 0.77 | 0.000972372 | 0.006150164 |
| 5970 | *RELA* | ENSG00000173039.20 | 10.32 | 10.70 | -0.38 | 0.77 | 1.37031E-05 | 0.000272771 |
| 10885 | *WDR3* | ENSG00000065183.16 | 6.19 | 6.57 | -0.38 | 0.77 | 0.000219217 | 0.002064136 |
| 25828 | *TXN2* | ENSG00000100348.10 | 5.79 | 6.17 | -0.38 | 0.77 | 0.000756693 | 0.005143662 |
| 10040 | *TOM1L1* | ENSG00000141198.16 | 4.66 | 5.04 | -0.38 | 0.77 | 0.000665266 | 0.004668056 |
| 64759 | *TNS3* | ENSG00000136205.17 | 7.05 | 7.43 | -0.38 | 0.77 | 0.00014842 | 0.001554074 |
| 2110 | *ETFDH* | ENSG00000171503.13 | 5.26 | 5.64 | -0.38 | 0.77 | 0.000547144 | 0.00406137 |
| 26750 | *RPS6KC1* | ENSG00000136643.12 | 5.82 | 6.21 | -0.38 | 0.77 | 0.000268247 | 0.002410853 |
| 27032 | *ATP2C1* | ENSG00000017260.20 | 7.23 | 7.61 | -0.38 | 0.77 | 0.000393159 | 0.00316743 |
| 26523 | *AGO1* | ENSG00000092847.13 | 7.77 | 8.15 | -0.38 | 0.77 | 1.35887E-05 | 0.000271213 |
| 6742 | *SSBP1* | ENSG00000106028.11 | 6.81 | 7.19 | -0.38 | 0.77 | 7.30042E-05 | 0.000937313 |
| 55794 | *DDX28* | ENSG00000182810.7 | 3.85 | 4.23 | -0.38 | 0.77 | 0.000839592 | 0.005558638 |
| 11140 | *CDC37* | ENSG00000105401.9 | 6.45 | 6.83 | -0.38 | 0.77 | 2.41179E-05 | 0.00041512 |
| 25936 | *NSL1* | ENSG00000117697.15 | 4.68 | 5.06 | -0.38 | 0.77 | 0.001503799 | 0.008606003 |
| 26019 | *UPF2* | ENSG00000151461.20 | 5.19 | 5.57 | -0.38 | 0.77 | 0.000476654 | 0.003659383 |
| 79665 | *DHX40* | ENSG00000108406.10 | 6.57 | 6.96 | -0.38 | 0.77 | 4.81548E-05 | 0.000695626 |
| 3140 | *MR1* | ENSG00000153029.16 | 5.18 | 5.57 | -0.38 | 0.77 | 0.000163625 | 0.001664982 |
| 90861 | *JPT2* | ENSG00000206053.13 | 7.04 | 7.43 | -0.39 | 0.77 | 0.000958594 | 0.006083525 |
| 3028 | *HSD17B10* | ENSG00000072506.14 | 5.67 | 6.05 | -0.39 | 0.77 | 0.001246172 | 0.007442815 |
| 79954 | *NOL10* | ENSG00000115761.16 | 5.19 | 5.58 | -0.39 | 0.76 | 8.94629E-05 | 0.001077648 |
| 55142 | *HAUS2* | ENSG00000137814.12 | 4.95 | 5.33 | -0.39 | 0.76 | 0.000450056 | 0.003514515 |
| 55055 | *ZWILCH* | ENSG00000174442.12 | 6.13 | 6.52 | -0.39 | 0.76 | 0.000778991 | 0.005247703 |
| 84629 | *TNRC18* | ENSG00000182095.15 | 5.61 | 6.00 | -0.39 | 0.76 | 0.001137742 | 0.006938678 |
| 225 | *ABCD2* | ENSG00000173208.4 | 4.40 | 4.79 | -0.39 | 0.76 | 0.000559759 | 0.004125864 |
| 54552 | *GNL3L* | ENSG00000130119.17 | 4.66 | 5.04 | -0.39 | 0.76 | 0.001098865 | 0.006753837 |
| 125988 | *MICOS13* | ENSG00000174917.9 | 4.08 | 4.47 | -0.39 | 0.76 | 0.001831194 | 0.009972564 |
| 64219 | *PJA1* | ENSG00000181191.12 | 6.19 | 6.58 | -0.39 | 0.76 | 0.000265359 | 0.002401908 |
| 7447 | *VSNL1* | ENSG00000163032.12 | 5.54 | 5.93 | -0.39 | 0.76 | 0.000664891 | 0.004668056 |
| 79738 | *BBS10* | ENSG00000179941.9 | 4.54 | 4.93 | -0.39 | 0.76 | 0.00026632 | 0.002405056 |
| 6241 | *RRM2* | ENSG00000171848.16 | 7.65 | 8.04 | -0.39 | 0.76 | 4.40641E-05 | 0.000655459 |
| 112858 | *TP53RK* | ENSG00000172315.6 | 4.70 | 5.09 | -0.39 | 0.76 | 0.0009241 | 0.005929809 |
| 81889 | *FAHD1* | ENSG00000180185.12 | 4.54 | 4.93 | -0.39 | 0.76 | 0.000295574 | 0.002583735 |
| 79903 | *NAA60* | ENSG00000122390.19 | 5.73 | 6.12 | -0.39 | 0.76 | 0.000724215 | 0.004976985 |
| 7756 | *ZNF207* | ENSG00000010244.19 | 7.48 | 7.87 | -0.39 | 0.76 | 6.09018E-06 | 0.000149115 |
| 114805 | *GALNT13* | ENSG00000144278.15 | 5.05 | 5.44 | -0.39 | 0.76 | 0.00110797 | 0.006803377 |
| 286077 | *FAM83H* | ENSG00000180921.7 | 4.37 | 4.77 | -0.40 | 0.76 | 0.000299791 | 0.002615981 |
| 22906 | *TRAK1* | ENSG00000182606.17 | 5.57 | 5.96 | -0.40 | 0.76 | 0.000455081 | 0.003537187 |
| 4082 | *MARCKS* | ENSG00000277443.3 | 6.53 | 6.93 | -0.40 | 0.76 | 4.12255E-05 | 0.000623743 |
| 55296 | *TBC1D19* | ENSG00000109680.11 | 4.63 | 5.02 | -0.40 | 0.76 | 0.001632504 | 0.009178408 |
| 10587 | *TXNRD2* | ENSG00000184470.21 | 5.58 | 5.98 | -0.40 | 0.76 | 0.000311017 | 0.00267438 |
| 152007 | *GLIPR2* | ENSG00000122694.16 | 6.94 | 7.33 | -0.40 | 0.76 | 9.06193E-06 | 0.000198395 |
| 80823 | *BHLHB9* | ENSG00000198908.12 | 3.42 | 3.82 | -0.40 | 0.76 | 0.001231801 | 0.007389332 |
| 51652 | *CHMP3* | ENSG00000115561.16 | 6.28 | 6.68 | -0.40 | 0.76 | 0.000884101 | 0.005759322 |
| 9577 | *BABAM2* | ENSG00000158019.21 | 4.24 | 4.64 | -0.40 | 0.76 | 0.000754339 | 0.005132312 |
| 57461 | *ISY1* | ENSG00000240682.10 | 4.95 | 5.35 | -0.40 | 0.76 | 0.001414876 | 0.008221397 |
| 51520 | *LARS1* | ENSG00000133706.19 | 6.73 | 7.13 | -0.40 | 0.76 | 0.000121757 | 0.001348678 |
| 57494 | *RIMKLB* | ENSG00000166532.16 | 5.15 | 5.55 | -0.40 | 0.76 | 0.000437055 | 0.003438027 |
| 339290 | *LINC00667* | ENSG00000263753.9 | 4.54 | 4.94 | -0.40 | 0.76 | 0.000928019 | 0.005949868 |
| 55572 | *FOXRED1* | ENSG00000110074.12 | 4.12 | 4.53 | -0.40 | 0.76 | 0.00015694 | 0.00161558 |

| | | | | | | | | |
|---|---|---|---|---|---|---|---|---|
| 6573 | *SLC19A1* | ENSG00000173638.19 | 4.74 | 5.15 | -0.41 | 0.75 | 0.000874663 | 0.005724516 |
| 27090 | *ST6GALNAC4* | ENSG00000136840.19 | 3.40 | 3.80 | -0.41 | 0.75 | 0.001454545 | 0.008386965 |
| 3184 | *HNRNPD* | ENSG00000138668.19 | 8.20 | 8.60 | -0.41 | 0.75 | 3.69311E-06 | 0.00010518 |
| 9054 | *NFS1* | ENSG00000244005.13 | 5.48 | 5.89 | -0.41 | 0.75 | 0.000422374 | 0.003352412 |
| 4598 | *MVK* | ENSG00000110921.14 | 5.40 | 5.81 | -0.41 | 0.75 | 0.000600203 | 0.004345609 |
| 5073 | *PARN* | ENSG00000140694.17 | 6.02 | 6.43 | -0.41 | 0.75 | 5.14333E-05 | 0.000727041 |
| 159 | *ADSS2* | ENSG00000035687.10 | 5.81 | 6.22 | -0.41 | 0.75 | 2.55513E-05 | 0.000435715 |
| 84851 | *TRIM52* | ENSG00000183718.6 | 4.05 | 4.46 | -0.41 | 0.75 | 0.000998446 | 0.00627804 |
| 10144 | *FAM13A* | ENSG00000138640.15 | 4.69 | 5.10 | -0.41 | 0.75 | 0.000203682 | 0.001957142 |
| 23557 | *SNAPIN* | ENSG00000143553.11 | 4.96 | 5.37 | -0.41 | 0.75 | 0.000354317 | 0.002946231 |
| 55269 | *PSPC1* | ENSG00000121390.19 | 5.27 | 5.68 | -0.41 | 0.75 | 0.000129894 | 0.001408658 |
| 57446 | *NDRG3* | ENSG00000101079.21 | 5.89 | 6.30 | -0.41 | 0.75 | 2.57237E-05 | 0.00043774 |
| 56886 | *UGGT1* | ENSG00000136731.13 | 7.00 | 7.41 | -0.41 | 0.75 | 0.0007265 | 0.004987134 |
| 23089 | *PEG10* | ENSG00000242265.6 | 10.27 | 10.68 | -0.41 | 0.75 | 0.000233416 | 0.002169249 |
| 3762 | *KCNJ5* | ENSG00000120457.12 | 7.68 | 8.10 | -0.42 | 0.75 | 4.52319E-05 | 0.000666092 |
| 3312 | *HSPA8* | ENSG00000109971.14 | 9.78 | 10.20 | -0.42 | 0.75 | 2.81641E-06 | 8.64449E-05 |
| 55631 | *LRRC40* | ENSG00000066557.6 | 4.68 | 5.09 | -0.42 | 0.75 | 0.000153652 | 0.001592657 |
| 10274 | *STAG1* | ENSG00000118007.13 | 5.91 | 6.33 | -0.42 | 0.75 | 1.8135E-05 | 0.000334384 |
| 26121 | *PRPF31* | ENSG00000105618.14 | 5.55 | 5.97 | -0.42 | 0.75 | 0.000126351 | 0.001379722 |
| 116224 | *PABIR1* | ENSG00000187866.10 | 5.95 | 6.37 | -0.42 | 0.75 | 2.89171E-05 | 0.000475375 |
| 65108 | *MARCKSL1* | ENSG00000175130.7 | 6.74 | 7.16 | -0.42 | 0.75 | 1.43201E-05 | 0.000282118 |
| 6920 | *TCEA3* | ENSG00000204219.11 | 5.60 | 6.02 | -0.42 | 0.75 | 0.00027338 | 0.002445267 |
| 6941 | *TCF19* | ENSG00000137310.12 | 6.44 | 6.86 | -0.42 | 0.75 | 1.54056E-05 | 0.000294852 |
| 51616 | *TAF9B* | ENSG00000187325.5 | 5.15 | 5.57 | -0.42 | 0.75 | 8.19988E-05 | 0.001014608 |
| 3704 | *ITPA* | ENSG00000125877.13 | 4.05 | 4.47 | -0.42 | 0.75 | 0.001587103 | 0.008974198 |
| 1020 | *CDK5* | ENSG00000164885.13 | 5.21 | 5.63 | -0.42 | 0.75 | 0.000268247 | 0.002410853 |
| 9987 | *HNRNPDL* | ENSG00000152795.18 | 7.58 | 8.00 | -0.42 | 0.75 | 6.35851E-06 | 0.000154425 |
| 10922 | *FASTK* | ENSG00000164896.21 | 5.34 | 5.77 | -0.42 | 0.75 | 0.000130844 | 0.001412832 |
| 5859 | *QARS1* | ENSG00000172053.18 | 6.36 | 6.79 | -0.42 | 0.75 | 0.001657769 | 0.009282495 |
| 28978 | *TMEM14A* | ENSG00000096092.6 | 5.48 | 5.90 | -0.42 | 0.74 | 0.00018597 | 0.001822422 |
| 9134 | *CCNE2* | ENSG00000175305.18 | 5.48 | 5.91 | -0.43 | 0.74 | 3.09474E-05 | 0.000502952 |
| 5358 | *PLS3* | ENSG00000102024.19 | 7.42 | 7.85 | -0.43 | 0.74 | 0.000367843 | 0.003025179 |
| 987 | *LRBA* | ENSG00000198589.14 | 6.05 | 6.47 | -0.43 | 0.74 | 0.001761385 | 0.009668126 |
| 54931 | *TRMT10C* | ENSG00000174173.7 | 5.71 | 6.14 | -0.43 | 0.74 | 0.000427739 | 0.003378916 |
| 60492 | *CCDC90B* | ENSG00000137500.10 | 5.31 | 5.73 | -0.43 | 0.74 | 0.000355423 | 0.002953665 |
| 64785 | *GINS3* | ENSG00000181938.14 | 5.75 | 6.18 | -0.43 | 0.74 | 1.07783E-05 | 0.000227527 |
| 54884 | *RETSAT* | ENSG00000042445.14 | 6.04 | 6.47 | -0.43 | 0.74 | 3.1061E-05 | 0.00050345 |
| 81605 | *URM1* | ENSG00000167118.11 | 5.84 | 6.27 | -0.43 | 0.74 | 0.000118632 | 0.001328771 |
| 4810 | *NHS* | ENSG00000188158.17 | 4.60 | 5.03 | -0.43 | 0.74 | 0.00053221 | 0.003974101 |
| 50944 | *SHANK1* | ENSG00000161681.17 | 6.20 | 6.63 | -0.43 | 0.74 | 0.000675775 | 0.004721933 |
| 27072 | *VPS41* | ENSG00000006715.16 | 5.82 | 6.25 | -0.43 | 0.74 | 0.001067654 | 0.006610734 |
| 64327 | *LMBR1* | ENSG00000105983.23 | 7.00 | 7.43 | -0.43 | 0.74 | 0.00025906 | 0.002359364 |
| 5889 | *RAD51C* | ENSG00000108384.15 | 4.19 | 4.62 | -0.43 | 0.74 | 0.000758925 | 0.005154164 |
| 1983 | *EIF5* | ENSG00000100664.11 | 6.91 | 7.35 | -0.43 | 0.74 | 0.000389979 | 0.00315027 |
| 3187 | *HNRNPH1* | ENSG00000169045.17 | 8.62 | 9.05 | -0.43 | 0.74 | 6.6984E-06 | 0.000160345 |
| 5239 | *PGM5* | ENSG00000154330.13 | 6.85 | 7.29 | -0.44 | 0.74 | 7.39544E-05 | 0.000943862 |
| 2629 | *GBA* | ENSG00000177628.16 | 6.77 | 7.21 | -0.44 | 0.74 | 4.757E-05 | 0.000690501 |
| 124637 | *CYB5D1* | ENSG00000182224.12 | 4.14 | 4.58 | -0.44 | 0.74 | 0.000120919 | 0.001345942 |
| 3837 | *KPNB1* | ENSG00000108424.11 | 8.28 | 8.72 | -0.44 | 0.74 | 1.98401E-05 | 0.000357479 |
| 90333 | *ZNF468* | ENSG00000204604.13 | 4.88 | 5.32 | -0.44 | 0.74 | 0.001291433 | 0.007668246 |
| 5827 | *PXMP2* | ENSG00000176894.10 | 4.03 | 4.47 | -0.44 | 0.74 | 0.000994197 | 0.006256565 |
| 79035 | *NABP2* | ENSG00000139579.13 | 5.26 | 5.69 | -0.44 | 0.74 | 0.000976395 | 0.006165213 |
| 56926 | *NCLN* | ENSG00000125912.11 | 6.18 | 6.62 | -0.44 | 0.74 | 5.49216E-05 | 0.000763258 |
| 2653 | *GCSH* | ENSG00000140905.11 | 4.82 | 5.26 | -0.44 | 0.74 | 0.000819116 | 0.005464047 |
| 54839 | *LRRC49* | ENSG00000137821.12 | 6.69 | 7.13 | -0.44 | 0.74 | 1.02401E-05 | 0.000219249 |
| 375748 | *ERCC6L2* | ENSG00000182150.19 | 5.43 | 5.87 | -0.44 | 0.74 | 0.00037637 | 0.003073414 |
| 168455 | *CCDC71L* | ENSG00000253276.5 | 4.61 | 5.05 | -0.44 | 0.74 | 0.001082574 | 0.006675574 |
| 3437 | *IFIT3* | ENSG00000119917.15 | 3.92 | 4.36 | -0.44 | 0.74 | 0.000308693 | 0.002665809 |
| 56160 | *NSMCE3* | ENSG00000185115.6 | 5.33 | 5.77 | -0.44 | 0.74 | 0.000167589 | 0.001687018 |
| 373156 | *GSTK1* | ENSG00000197448.14 | 6.40 | 6.84 | -0.44 | 0.74 | 8.96118E-06 | 0.000196923 |
| 1496 | *CTNNA2* | ENSG00000066032.19 | 6.27 | 6.72 | -0.44 | 0.73 | 5.93662E-05 | 0.000809289 |
| 54809 | *SAMD9* | ENSG00000205413.8 | 6.05 | 6.50 | -0.44 | 0.73 | 2.61567E-05 | 0.000444102 |
| 80742 | *PRR3* | ENSG00000204576.12 | 4.86 | 5.31 | -0.45 | 0.73 | 0.000118095 | 0.001324729 |
| 23065 | *EMC1* | ENSG00000127463.16 | 6.64 | 7.09 | -0.45 | 0.73 | 0.000102883 | 0.001191486 |
| 1075 | *CTSC* | ENSG00000109861.17 | 7.07 | 7.52 | -0.45 | 0.73 | 2.88042E-05 | 0.000474597 |
| 27143 | *PALD1* | ENSG00000107719.9 | 3.30 | 3.75 | -0.45 | 0.73 | 0.001528634 | 0.008707124 |
| 1355 | *COX15* | ENSG00000014919.13 | 5.63 | 6.08 | -0.45 | 0.73 | 0.001622521 | 0.00913787 |
| 1793 | *DOCK1* | ENSG00000150760.13 | 5.32 | 5.77 | -0.45 | 0.73 | 5.08574E-05 | 0.000721473 |
| 65110 | *UPF3A* | ENSG00000169062.15 | 6.75 | 7.20 | -0.45 | 0.73 | 1.85258E-05 | 0.000338677 |
| 1114 | *CHGB* | ENSG00000089199.10 | 6.14 | 6.59 | -0.45 | 0.73 | 1.52608E-05 | 0.000293571 |
| 54939 | *COMMD4* | ENSG00000140365.16 | 4.96 | 5.41 | -0.45 | 0.73 | 2.37263E-05 | 0.000410263 |
| 7324 | *UBE2E1* | ENSG00000170142.12 | 5.29 | 5.74 | -0.45 | 0.73 | 0.000438057 | 0.00344391 |
| 9836 | *LCMT2* | ENSG00000168806.8 | 4.18 | 4.63 | -0.45 | 0.73 | 0.000392966 | 0.00316743 |
| 2800 | *GOLGA1* | ENSG00000136935.14 | 4.97 | 5.43 | -0.46 | 0.73 | 0.000898105 | 0.005822744 |
| 79968 | *WDR76* | ENSG00000092470.12 | 5.70 | 6.16 | -0.46 | 0.73 | 0.000363291 | 0.002999246 |
| 11098 | *PRSS23* | ENSG00000150687.12 | 6.75 | 7.20 | -0.46 | 0.73 | 0.000574313 | 0.004209099 |
| 8437 | *RASAL1* | ENSG00000111344.12 | 4.77 | 5.23 | -0.46 | 0.73 | 0.000263429 | 0.002387141 |
| 1945 | *EFNA4* | ENSG00000243364.8 | 4.90 | 5.36 | -0.46 | 0.73 | 0.000142373 | 0.001506617 |
| 85476 | *GFM1* | ENSG00000168827.15 | 6.18 | 6.63 | -0.46 | 0.73 | 0.000168873 | 0.001695658 |
| 286333 | *FAM225A* | ENSG00000231528.3 | 3.94 | 4.40 | -0.46 | 0.73 | 0.000468198 | 0.003614806 |
| 57827 | *C6orf47* | ENSG00000204439.4 | 4.91 | 5.37 | -0.46 | 0.73 | 0.000380606 | 0.00309286 |
| 10421 | *CD2BP2* | ENSG00000169217.9 | 6.39 | 6.85 | -0.46 | 0.73 | 8.78446E-05 | 0.001066714 |
| 83759 | *RBM4B* | ENSG00000173914.12 | 4.38 | 4.84 | -0.46 | 0.73 | 0.000205993 | 0.001974295 |
| 63926 | *ANKEF1* | ENSG00000132623.16 | 4.24 | 4.70 | -0.46 | 0.73 | 0.001018923 | 0.006376652 |
| 4707 | *NDUFB1* | ENSG00000183648.11 | 4.11 | 4.57 | -0.46 | 0.73 | 0.000829744 | 0.005515338 |
| 7353 | *UFD1* | ENSG00000070010.19 | 6.46 | 6.92 | -0.46 | 0.73 | 7.09392E-05 | 0.000920633 |
| 203522 | *INTS6L* | ENSG00000165359.16 | 3.66 | 4.12 | -0.46 | 0.73 | 0.001209133 | 0.007282458 |
| 3988 | *LIPA* | ENSG00000107798.18 | 5.30 | 5.76 | -0.46 | 0.73 | 4.69929E-05 | 0.000684109 |
| 5303 | *PIN4* | ENSG00000102309.15 | 3.52 | 3.98 | -0.46 | 0.73 | 0.001243094 | 0.007427386 |

| | | | | | | | | |
|---|---|---|---|---|---|---|---|---|
| 161823 | *ADAL* | ENSG00000168803.16 | 4.04 | 4.50 | -0.46 | 0.73 | 0.001716885 | 0.009508754 |
| 57181 | *SLC39A10* | ENSG00000196950.14 | 6.81 | 7.28 | -0.46 | 0.73 | 2.76782E-05 | 0.000462456 |
| 4686 | *NCBP1* | ENSG00000136937.13 | 6.87 | 7.34 | -0.46 | 0.72 | 4.16494E-06 | 0.000113721 |
| 51367 | *POP5* | ENSG00000167272.11 | 3.91 | 4.38 | -0.47 | 0.72 | 0.000711777 | 0.004914011 |
| 11164 | *NUDT5* | ENSG00000165609.13 | 5.13 | 5.60 | -0.47 | 0.72 | 6.81657E-05 | 0.000898089 |
| 2946 | *GSTM2* | ENSG00000213366.13 | 4.83 | 5.29 | -0.47 | 0.72 | 0.000795775 | 0.005334426 |
| 83594 | *NUDT12* | ENSG00000112874.10 | 4.30 | 4.76 | -0.47 | 0.72 | 0.00031914 | 0.002723125 |
| 84305 | *PYM1* | ENSG00000170473.17 | 4.64 | 5.10 | -0.47 | 0.72 | 0.000318136 | 0.002717643 |
| 51659 | *GINS2* | ENSG00000131153.9 | 5.34 | 5.81 | -0.47 | 0.72 | 9.79017E-05 | 0.00115445 |
| 55220 | *KLHDC8A* | ENSG00000162873.15 | 4.53 | 5.00 | -0.47 | 0.72 | 0.001510748 | 0.008634733 |
| 5588 | *PRKCQ* | ENSG00000065675.16 | 3.32 | 3.79 | -0.47 | 0.72 | 0.000454433 | 0.003535814 |
| 1159 | *CKMT1B* | ENSG00000237289.10 | 4.64 | 5.11 | -0.47 | 0.72 | 0.001009314 | 0.006330464 |
| 114785 | *MBD6* | ENSG00000166987.15 | 5.63 | 6.11 | -0.47 | 0.72 | 6.85066E-05 | 0.000900364 |
| 9380 | *GRHPR* | ENSG00000137106.18 | 6.81 | 7.29 | -0.47 | 0.72 | 5.78772E-06 | 0.000144059 |
| 203 | *AK1* | ENSG00000106992.19 | 5.16 | 5.63 | -0.48 | 0.72 | 0.000489586 | 0.003731943 |
| 5702 | *PSMC3* | ENSG00000165916.9 | 7.20 | 7.68 | -0.48 | 0.72 | 0.000306675 | 0.002657796 |
| 54621 | *VSIG10* | ENSG00000176834.14 | 4.79 | 5.27 | -0.48 | 0.72 | 0.000616007 | 0.004423755 |
| 301 | *ANXA1* | ENSG00000135046.14 | 4.61 | 5.08 | -0.48 | 0.72 | 0.00012804 | 0.001392574 |
| 56950 | *SMYD2* | ENSG00000143499.14 | 4.25 | 4.73 | -0.48 | 0.72 | 0.001207001 | 0.007275452 |
| 9910 | *RABGAP1L* | ENSG00000152061.24 | 5.32 | 5.80 | -0.48 | 0.72 | 0.001108285 | 0.006803377 |
| 53 | *ACP2* | ENSG00000134575.13 | 5.52 | 6.00 | -0.48 | 0.72 | 0.000323354 | 0.002748102 |
| 83982 | *IFI27L2* | ENSG00000119632.4 | 3.86 | 4.34 | -0.48 | 0.72 | 0.000367059 | 0.003020388 |
| 83543 | *AIF1L* | ENSG00000126878.13 | 7.49 | 7.98 | -0.48 | 0.72 | 4.99363E-05 | 0.000712446 |
| 79600 | *TCTN1* | ENSG00000204852.17 | 4.40 | 4.87 | -0.48 | 0.72 | 0.000477856 | 0.003661124 |
| 210 | *ALAD* | ENSG00000148218.16 | 5.81 | 6.29 | -0.48 | 0.72 | 1.74572E-05 | 0.000325081 |
| 6904 | *TBCD* | ENSG00000141556.22 | 5.65 | 6.13 | -0.48 | 0.72 | 7.33127E-05 | 0.000939667 |
| 55971 | *BAIAP2L1* | ENSG00000006453.14 | 5.62 | 6.10 | -0.48 | 0.72 | 6.68374E-05 | 0.000883843 |
| 25941 | *TPGS2* | ENSG00000134779.15 | 6.11 | 6.59 | -0.48 | 0.72 | 5.97757E-05 | 0.000814091 |
| 23169 | *SLC35D1* | ENSG00000116704.8 | 6.51 | 6.99 | -0.48 | 0.72 | 3.37288E-05 | 0.000536835 |
| 716 | *C1S* | ENSG00000182326.16 | 4.86 | 5.35 | -0.48 | 0.71 | 0.000244137 | 0.002253534 |
| 2940 | *GSTA3* | ENSG00000174156.15 | 5.61 | 6.09 | -0.48 | 0.71 | 9.1362E-05 | 0.001092889 |
| 271 | *AMPD2* | ENSG00000116337.20 | 5.87 | 6.35 | -0.48 | 0.71 | 9.87463E-05 | 0.001158783 |
| 643988 | *FNDC10* | ENSG00000228594.4 | 4.61 | 5.10 | -0.48 | 0.71 | 9.15047E-05 | 0.001093467 |
| 119032 | *BORCS7* | ENSG00000166275.16 | 5.75 | 6.23 | -0.49 | 0.71 | 0.000391368 | 0.003158084 |
| 55793 | *MINDY1* | ENSG00000143409.17 | 3.85 | 4.33 | -0.49 | 0.71 | 0.001646392 | 0.009234194 |
| 26133 | *TRPC4AP* | ENSG00000100991.12 | 6.05 | 6.54 | -0.49 | 0.71 | 1.33911E-06 | 5.44324E-05 |
| 192683 | *SCAMP5* | ENSG00000198794.12 | 5.57 | 6.06 | -0.49 | 0.71 | 0.000615665 | 0.004423755 |
| 9555 | *MACROH2A1* | ENSG00000113648.17 | 7.39 | 7.88 | -0.49 | 0.71 | 4.66596E-06 | 0.000123295 |
| 3428 | *IFI16* | ENSG00000163565.20 | 5.18 | 5.67 | -0.49 | 0.71 | 9.68291E-05 | 0.001147046 |
| 11010 | *GLIPR1* | ENSG00000139278.10 | 5.37 | 5.86 | -0.49 | 0.71 | 0.000673683 | 0.004711701 |
| 84861 | *KLHL22* | ENSG00000099910.17 | 5.39 | 5.88 | -0.49 | 0.71 | 0.000122529 | 0.001354223 |
| 1588 | *CYP19A1* | ENSG00000137869.16 | 5.42 | 5.92 | -0.49 | 0.71 | 5.06449E-05 | 0.00072138 |
| 5356 | *PLRG1* | ENSG00000171566.12 | 5.47 | 5.96 | -0.49 | 0.71 | 0.000236475 | 0.002192247 |
| 9774 | *BCLAF1* | ENSG00000029363.17 | 7.97 | 8.46 | -0.49 | 0.71 | 7.73779E-06 | 0.000177579 |
| 126792 | *B3GALT6* | ENSG00000176022.7 | 3.85 | 4.35 | -0.49 | 0.71 | 0.000962466 | 0.006102938 |
| 116461 | *TSEN15* | ENSG00000198860.14 | 4.40 | 4.89 | -0.49 | 0.71 | 0.000430744 | 0.003399074 |
| 5046 | *PCSK6* | ENSG00000140479.18 | 5.03 | 5.53 | -0.49 | 0.71 | 0.000177232 | 0.00176047 |
| 79912 | *PYROXD1* | ENSG00000121350.16 | 4.51 | 5.01 | -0.50 | 0.71 | 0.000937631 | 0.005993573 |
| 1968 | *EIF2S3* | ENSG00000130741.11 | 6.58 | 7.08 | -0.50 | 0.71 | 7.47912E-06 | 0.000172965 |
| 93210 | *PGAP3* | ENSG00000161395.14 | 4.40 | 4.89 | -0.50 | 0.71 | 0.000463392 | 0.003591681 |
| 9158 | *FIBP* | ENSG00000172500.13 | 4.89 | 5.39 | -0.50 | 0.71 | 0.000305096 | 0.002648455 |
| 9931 | *HELZ* | ENSG00000198265.12 | 6.29 | 6.79 | -0.50 | 0.71 | 0.000341858 | 0.002866453 |
| 84964 | *ALKBH6* | ENSG00000239382.11 | 2.96 | 3.45 | -0.50 | 0.71 | 0.001088612 | 0.006699045 |
| 113675 | *SDSL* | ENSG00000139410.15 | 3.03 | 3.53 | -0.50 | 0.71 | 0.000787829 | 0.005290612 |
| 54799 | *MBTD1* | ENSG00000011258.16 | 4.47 | 4.97 | -0.50 | 0.71 | 0.000127102 | 0.001384384 |
| 1660 | *DHX9* | ENSG00000135829.17 | 8.17 | 8.67 | -0.50 | 0.71 | 1.03648E-06 | 4.62182E-05 |
| 90957 | *DHX57* | ENSG00000163214.21 | 5.38 | 5.88 | -0.50 | 0.71 | 9.9742E-05 | 0.001165002 |
| 6646 | *SOAT1* | ENSG00000057252.13 | 9.47 | 9.97 | -0.50 | 0.71 | 8.7972E-07 | 4.10053E-05 |
| 5106 | *PCK2* | ENSG00000100889.12 | 3.94 | 4.44 | -0.50 | 0.71 | 0.000556982 | 0.00411405 |
| 1719 | *DHFR* | ENSG00000228716.7 | 7.48 | 7.98 | -0.50 | 0.71 | 7.04821E-06 | 0.000165971 |
| 26512 | *INTS6* | ENSG00000102786.15 | 4.82 | 5.32 | -0.50 | 0.71 | 7.66923E-05 | 0.000971371 |
| 9829 | *DNAJC6* | ENSG00000116675.16 | 4.87 | 5.37 | -0.50 | 0.71 | 0.001233395 | 0.00739298 |
| 9899 | *SV2B* | ENSG00000185518.12 | 5.97 | 6.47 | -0.50 | 0.71 | 1.23813E-05 | 0.000250784 |
| 219285 | *SAMD9L* | ENSG00000177409.12 | 4.49 | 4.99 | -0.50 | 0.71 | 0.001305985 | 0.007708034 |
| 2235 | *FECH* | ENSG00000066926.13 | 5.84 | 6.34 | -0.50 | 0.71 | 0.000141233 | 0.00150019 |
| 776 | *CACNA1D* | ENSG00000157388.20 | 6.36 | 6.86 | -0.50 | 0.71 | 0.000126785 | 0.001382934 |
| 728661 | *SLC35E2B* | ENSG00000189339.12 | 4.86 | 5.36 | -0.50 | 0.71 | 0.000210792 | 0.00200875 |
| 79587 | *CARS2* | ENSG00000134905.17 | 4.76 | 5.26 | -0.50 | 0.71 | 7.87828E-05 | 0.000987021 |
| 3376 | *IARS1* | ENSG00000196305.19 | 8.15 | 8.66 | -0.50 | 0.71 | 1.65259E-06 | 6.15274E-05 |
| 7629 | *ZNF76* | ENSG00000065029.15 | 4.42 | 4.93 | -0.50 | 0.71 | 0.000167813 | 0.001688143 |
| 9076 | *CLDN1* | ENSG00000163347.6 | 8.47 | 8.97 | -0.51 | 0.70 | 1.48814E-06 | 5.80141E-05 |
| 148156 | *ZNF558* | ENSG00000167785.9 | 3.25 | 3.75 | -0.51 | 0.70 | 0.000853183 | 0.005626286 |
| 285362 | *SUMF1* | ENSG00000144455.14 | 3.47 | 3.97 | -0.51 | 0.70 | 0.000617096 | 0.004425886 |
| 51161 | *C3orf18* | ENSG00000088543.15 | 4.22 | 4.73 | -0.51 | 0.70 | 0.001077226 | 0.006655093 |
| 7145 | *TNS1* | ENSG00000079308.20 | 6.22 | 6.73 | -0.51 | 0.70 | 0.000227304 | 0.002122967 |
| 1718 | *DHCR24* | ENSG00000116133.13 | 8.87 | 9.38 | -0.51 | 0.70 | 1.08039E-06 | 4.6866E-05 |
| 23506 | *BICRAL* | ENSG00000112624.13 | 4.49 | 5.00 | -0.51 | 0.70 | 0.00043367 | 0.003413192 |
| 9568 | *GABBR2* | ENSG00000136928.7 | 7.07 | 7.59 | -0.51 | 0.70 | 3.22862E-05 | 0.000517165 |
| 2766 | *GMPR* | ENSG00000137198.10 | 3.28 | 3.79 | -0.51 | 0.70 | 0.000945803 | 0.006024636 |
| 11346 | *SYNPO* | ENSG00000171992.13 | 6.82 | 7.33 | -0.51 | 0.70 | 2.75587E-06 | 8.54897E-05 |
| 3092 | *HIP1* | ENSG00000127946.17 | 5.52 | 6.04 | -0.51 | 0.70 | 0.000405595 | 0.003251913 |
| 145957 | *NRG4* | ENSG00000169752.17 | 3.60 | 4.12 | -0.52 | 0.70 | 0.000850956 | 0.005616532 |
| 9844 | *ELMO1* | ENSG00000155849.16 | 4.04 | 4.55 | -0.52 | 0.70 | 0.000789343 | 0.00529841 |
| 374618 | *TEX9* | ENSG00000151575.14 | 3.07 | 3.59 | -0.52 | 0.70 | 0.000651067 | 0.00459636 |
| 503637 | *DUXAP8* | ENSG00000206195.11 | 5.63 | 6.15 | -0.52 | 0.70 | 4.36776E-05 | 0.000651648 |
| 5591 | *PRKDC* | ENSG00000253729.8 | 8.05 | 8.57 | -0.52 | 0.70 | 0.000227262 | 0.002122967 |
| 83990 | *BRIP1* | ENSG00000136492.10 | 5.08 | 5.60 | -0.52 | 0.70 | 6.57589E-05 | 0.000871306 |
| 4507 | *MTAP* | ENSG00000099810.21 | 5.67 | 6.19 | -0.52 | 0.70 | 0.000377827 | 0.003075272 |
| 1601 | *DAB2* | ENSG00000153071.15 | 9.28 | 9.80 | -0.52 | 0.70 | 1.364E-06 | 5.47387E-05 |

| | | | | | | | | |
|---|---|---|---|---|---|---|---|---|
| 58155 | *PTBP2* | ENSG00000117569.19 | 4.61 | 5.13 | -0.52 | 0.70 | 0.000664239 | 0.004668056 |
| 56654 | *NPDC1* | ENSG00000107281.10 | 3.82 | 4.33 | -0.52 | 0.70 | 0.000170587 | 0.001706891 |
| 78991 | *PCYOX1L* | ENSG00000145882.11 | 4.15 | 4.67 | -0.52 | 0.70 | 0.000375177 | 0.003068684 |
| 55005 | *RMND1* | ENSG00000155906.20 | 3.97 | 4.50 | -0.52 | 0.70 | 0.000496411 | 0.003772473 |
| 93183 | *PIGM* | ENSG00000143315.8 | 5.70 | 6.23 | -0.52 | 0.70 | 1.23457E-05 | 0.000250566 |
| 6603 | *SMARCD2* | ENSG00000108604.17 | 5.07 | 5.59 | -0.52 | 0.70 | 0.000162534 | 0.001655 |
| 9897 | *WASHC5* | ENSG00000164961.16 | 6.29 | 6.82 | -0.52 | 0.70 | 4.54994E-06 | 0.000120867 |
| 51209 | *RAB9B* | ENSG00000123570.4 | 3.30 | 3.83 | -0.53 | 0.69 | 0.001074638 | 0.006648493 |
| 8881 | *CDC16* | ENSG00000130177.16 | 7.10 | 7.62 | -0.53 | 0.69 | 1.96676E-06 | 6.94566E-05 |
| 340526 | *RTL5* | ENSG00000242732.4 | 4.95 | 5.48 | -0.53 | 0.69 | 7.05207E-05 | 0.000917992 |
| 1944 | *EFNA3* | ENSG00000143590.14 | 5.29 | 5.82 | -0.53 | 0.69 | 2.83013E-06 | 8.66887E-05 |
| 53407 | *STX18* | ENSG00000168818.10 | 4.81 | 5.33 | -0.53 | 0.69 | 0.000641975 | 0.004553595 |
| 435 | *ASL* | ENSG00000126522.17 | 3.43 | 3.96 | -0.53 | 0.69 | 0.001776358 | 0.00972687 |
| 7321 | *UBE2D1* | ENSG00000072401.15 | 4.41 | 4.94 | -0.53 | 0.69 | 0.000938112 | 0.005994093 |
| 83666 | *PARP9* | ENSG00000138496.17 | 5.35 | 5.88 | -0.53 | 0.69 | 7.35478E-05 | 0.000940271 |
| 3070 | *HELLS* | ENSG00000119969.15 | 5.49 | 6.02 | -0.53 | 0.69 | 1.10912E-05 | 0.000232132 |
| 57628 | *DPP10* | ENSG00000175497.17 | 6.43 | 6.96 | -0.53 | 0.69 | 1.50369E-05 | 0.000290684 |
| 57610 | *RANBP10* | ENSG00000141084.12 | 4.13 | 4.66 | -0.53 | 0.69 | 3.72869E-05 | 0.000575761 |
| 5164 | *PDK2* | ENSG00000005882.12 | 5.08 | 5.62 | -0.53 | 0.69 | 0.001585048 | 0.008970583 |
| 131540 | *ZDHHC19* | ENSG00000163958.14 | 4.10 | 4.63 | -0.53 | 0.69 | 0.000353887 | 0.002944282 |
| 100289097 | *FRG1CP* | ENSG00000282826.3 | 3.58 | 4.11 | -0.53 | 0.69 | 0.000166513 | 0.001684088 |
| 79624 | *ARMT1* | ENSG00000146476.11 | 6.55 | 7.08 | -0.53 | 0.69 | 2.45126E-06 | 7.9462E-05 |
| 54985 | *HCFC1R1* | ENSG00000103145.11 | 3.31 | 3.85 | -0.53 | 0.69 | 0.001137196 | 0.00693828 |
| 4144 | *MAT2A* | ENSG00000168906.13 | 6.62 | 7.15 | -0.53 | 0.69 | 0.000113894 | 0.001287221 |
| 10801 | *SEPTIN9* | ENSG00000184640.20 | 7.05 | 7.58 | -0.54 | 0.69 | 1.74491E-05 | 0.000325081 |
| 24148 | *PRPF6* | ENSG00000101161.8 | 5.25 | 5.78 | -0.54 | 0.69 | 3.26611E-06 | 9.7527E-05 |
| 27304 | *MOCS3* | ENSG00000124217.5 | 3.36 | 3.90 | -0.54 | 0.69 | 0.000294632 | 0.002578501 |
| 51157 | *ZNF580* | ENSG00000213015.9 | 3.36 | 3.89 | -0.54 | 0.69 | 0.001048886 | 0.006521431 |
| 55869 | *HDAC8* | ENSG00000147099.21 | 3.98 | 4.52 | -0.54 | 0.69 | 0.000787063 | 0.005287838 |
| 6822 | *SULT2A1* | ENSG00000105398.4 | 7.07 | 7.61 | -0.54 | 0.69 | 1.44214E-06 | 5.68113E-05 |
| 27291 | *R3HCC1L* | ENSG00000166024.14 | 3.73 | 4.27 | -0.54 | 0.69 | 0.000293424 | 0.002573935 |
| 100506054 | *RNASEH1-DT* | ENSG00000234171.4 | 2.67 | 3.20 | -0.54 | 0.69 | 0.001658096 | 0.009282495 |
| 402665 | *IGLON5* | ENSG00000142549.10 | 5.28 | 5.82 | -0.54 | 0.69 | 2.80088E-06 | 8.61442E-05 |
| 6426 | *SRSF1* | ENSG00000136450.13 | 7.94 | 8.48 | -0.54 | 0.69 | 7.09885E-07 | 3.55095E-05 |
| 112487 | *DTD2* | ENSG00000129480.13 | 3.74 | 4.28 | -0.54 | 0.69 | 0.000584922 | 0.004261697 |
| 80017 | *DGLUCY* | ENSG00000133943.21 | 5.38 | 5.93 | -0.54 | 0.69 | 0.000478918 | 0.003665516 |
| 715 | *C1R* | ENSG00000159403.18 | 5.14 | 5.68 | -0.54 | 0.69 | 2.99176E-05 | 0.000489143 |
| 23078 | *VWA8* | ENSG00000102763.18 | 3.36 | 3.91 | -0.54 | 0.69 | 0.000353735 | 0.002944282 |
| 23595 | *ORC3* | ENSG00000135336.16 | 5.34 | 5.89 | -0.54 | 0.69 | 0.000355906 | 0.002954528 |
| 79184 | *BRCC3* | ENSG00000185515.16 | 5.55 | 6.10 | -0.55 | 0.69 | 1.19155E-05 | 0.000246153 |
| 7626 | *ZNF75D* | ENSG00000186376.15 | 5.44 | 5.99 | -0.55 | 0.69 | 3.35419E-05 | 0.000534427 |
| 4311 | *MME* | ENSG00000196549.13 | 8.73 | 9.27 | -0.55 | 0.68 | 4.2442E-06 | 0.000114843 |
| 64792 | *IFT22* | ENSG00000128581.17 | 5.55 | 6.10 | -0.55 | 0.68 | 5.21105E-05 | 0.00073411 |
| 11160 | *ERLIN2* | ENSG00000147475.17 | 5.73 | 6.28 | -0.55 | 0.68 | 0.001363982 | 0.008000002 |
| 4200 | *ME2* | ENSG00000082212.13 | 4.80 | 5.34 | -0.55 | 0.68 | 2.6965E-05 | 0.00045372 |
| 56062 | *KLHL4* | ENSG00000102271.14 | 4.94 | 5.49 | -0.55 | 0.68 | 6.89064E-06 | 0.000164161 |
| 113419 | *TEX261* | ENSG00000144043.12 | 5.92 | 6.47 | -0.55 | 0.68 | 8.89899E-05 | 0.001075403 |
| 9344 | *TAOK2* | ENSG00000149930.18 | 5.90 | 6.45 | -0.55 | 0.68 | 2.88936E-05 | 0.000475375 |
| 57476 | *GRAMD1B* | ENSG00000023171.20 | 8.10 | 8.65 | -0.55 | 0.68 | 9.23687E-06 | 0.000200922 |
| 23233 | *EXOC6B* | ENSG00000144036.16 | 4.50 | 5.05 | -0.55 | 0.68 | 0.000478821 | 0.003665516 |
| 9989 | *PPP4R1* | ENSG00000154845.16 | 5.72 | 6.27 | -0.55 | 0.68 | 1.79092E-05 | 0.000331441 |
| 116442 | *RAB39B* | ENSG00000155961.5 | 4.36 | 4.91 | -0.55 | 0.68 | 4.8837E-05 | 0.000703449 |
| 57567 | *ZNF319* | ENSG00000166188.3 | 4.03 | 4.58 | -0.55 | 0.68 | 0.001596132 | 0.009016312 |
| 27086 | *FOXP1* | ENSG00000114861.24 | 3.47 | 4.02 | -0.55 | 0.68 | 0.001041269 | 0.006484814 |
| 108783654 | *HSD17B1-AS* | ENSG00000266962.2 | 3.17 | 3.72 | -0.55 | 0.68 | 0.00082279 | 0.005478817 |
| 91975 | *ZNF300* | ENSG00000145908.13 | 3.10 | 3.65 | -0.55 | 0.68 | 0.00010501 | 0.001206811 |
| 26999 | *CYFIP2* | ENSG00000055163.20 | 5.74 | 6.29 | -0.55 | 0.68 | 0.000194644 | 0.001886485 |
| 90139 | *TSPAN18* | ENSG00000157570.12 | 3.58 | 4.13 | -0.55 | 0.68 | 0.001299204 | 0.007684019 |
| 29082 | *CHMP4A* | ENSG00000254505.11 | 3.29 | 3.85 | -0.55 | 0.68 | 0.000530503 | 0.003967277 |
| 6319 | *SCD* | ENSG00000099194.6 | 9.20 | 9.76 | -0.55 | 0.68 | 9.77742E-07 | 4.44695E-05 |
| 56154 | *TEX15* | ENSG00000133863.9 | 3.25 | 3.80 | -0.55 | 0.68 | 0.000632311 | 0.004501425 |
| 2521 | *FUS* | ENSG00000089280.19 | 8.64 | 9.19 | -0.55 | 0.68 | 1.34533E-06 | 5.44324E-05 |
| 645323 | *LINC00461* | ENSG00000245526.12 | 5.49 | 6.04 | -0.55 | 0.68 | 0.000312161 | 0.002678802 |
| 2317 | *FLNB* | ENSG00000136068.16 | 6.42 | 6.97 | -0.55 | 0.68 | 0.00062433 | 0.004464318 |
| 2582 | *GALE* | ENSG00000117308.15 | 2.86 | 3.41 | -0.56 | 0.68 | 0.001505722 | 0.008612568 |
| 114815 | *SORCS1* | ENSG00000108018.16 | 6.28 | 6.83 | -0.56 | 0.68 | 1.87625E-06 | 6.70597E-05 |
| 10165 | *SLC25A13* | ENSG00000004864.14 | 5.53 | 6.09 | -0.56 | 0.68 | 2.82303E-05 | 0.000468611 |
| 64976 | *MRPL40* | ENSG00000185608.9 | 4.88 | 5.43 | -0.56 | 0.68 | 3.64233E-05 | 0.000565333 |
| 440465 | *BAIAP2-DT* | ENSG00000226137.5 | 4.58 | 5.14 | -0.56 | 0.68 | 6.57732E-05 | 0.000871306 |
| 2121 | *EVC* | ENSG00000072840.13 | 4.28 | 4.83 | -0.56 | 0.68 | 0.000740219 | 0.005056333 |
| 116228 | *COX20* | ENSG00000203667.10 | 4.83 | 5.38 | -0.56 | 0.68 | 0.001183201 | 0.007160754 |
| 27077 | *B9D1* | ENSG00000108641.20 | 3.99 | 4.55 | -0.56 | 0.68 | 0.000342239 | 0.002868044 |
| 10432 | *RBM14* | ENSG00000239306.5 | 5.85 | 6.41 | -0.56 | 0.68 | 0.00015291 | 0.001589358 |
| 64848 | *YTHDC2* | ENSG00000047188.16 | 4.35 | 4.91 | -0.56 | 0.68 | 0.001278557 | 0.007618049 |
| 22994 | *CEP131* | ENSG00000141577.14 | 3.84 | 4.40 | -0.56 | 0.68 | 0.000740477 | 0.005056333 |
| 8913 | *CACNA1G* | ENSG00000006283.18 | 3.51 | 4.07 | -0.56 | 0.68 | 6.69298E-05 | 0.000884287 |
| 27340 | *UTP20* | ENSG00000120800.5 | 5.47 | 6.04 | -0.56 | 0.68 | 0.000294233 | 0.002578148 |
| 2176 | *FANCC* | ENSG00000158169.13 | 4.72 | 5.28 | -0.56 | 0.68 | 0.001443464 | 0.00833588 |
| 88745 | *RRP36* | ENSG00000124541.7 | 4.76 | 5.33 | -0.57 | 0.68 | 1.21339E-05 | 0.000248254 |
| 139285 | *AMER1* | ENSG00000184675.11 | 5.44 | 6.01 | -0.57 | 0.67 | 5.25612E-06 | 0.000134853 |
| 4886 | *NPY1R* | ENSG00000164128.7 | 3.22 | 3.79 | -0.57 | 0.67 | 0.000831716 | 0.005517878 |
| 338761 | *C1QL4* | ENSG00000186897.5 | 3.77 | 4.34 | -0.57 | 0.67 | 0.000200873 | 0.001935108 |
| 55508 | *SLC35E3* | ENSG00000175782.11 | 4.90 | 5.47 | -0.57 | 0.67 | 6.86822E-05 | 0.00090067 |
| 57447 | *NDRG2* | ENSG00000165795.25 | 6.03 | 6.59 | -0.57 | 0.67 | 3.7629E-06 | 0.000106361 |
| 3551 | *IKBKB* | ENSG00000104365.16 | 4.56 | 5.14 | -0.57 | 0.67 | 5.53164E-06 | 0.000139069 |
| 2937 | *GSS* | ENSG00000100983.12 | 5.09 | 5.67 | -0.57 | 0.67 | 1.67936E-05 | 0.000315069 |
| 10752 | *CHL1* | ENSG00000134121.10 | 4.91 | 5.48 | -0.58 | 0.67 | 0.001597583 | 0.009021116 |
| 9851 | *KIAA0753* | ENSG00000198920.11 | 3.57 | 4.15 | -0.58 | 0.67 | 0.001115721 | 0.006840213 |
| 55701 | *ARHGEF40* | ENSG00000165801.10 | 5.78 | 6.36 | -0.58 | 0.67 | 9.84689E-05 | 0.001156906 |

| | | | | | | | | |
|---|---|---|---|---|---|---|---|---|
| 55027 | *HEATR3* | ENSG00000155393.14 | 4.47 | 5.05 | -0.58 | 0.67 | 0.000427411 | 0.003378103 |
| 11282 | *MGAT4B* | ENSG00000161013.17 | 5.49 | 6.07 | -0.58 | 0.67 | 0.000134145 | 0.001441216 |
| 9479 | *MAPK8IP1* | ENSG00000121653.11 | 5.10 | 5.68 | -0.58 | 0.67 | 7.25247E-07 | 3.58067E-05 |
| 5229 | *PGGT1B* | ENSG00000164219.10 | 5.00 | 5.58 | -0.58 | 0.67 | 0.001044971 | 0.006505173 |
| 348262 | *MCRIP1* | ENSG00000225663.8 | 4.60 | 5.18 | -0.58 | 0.67 | 9.65952E-06 | 0.00020741 |
| 65123 | *INTS3* | ENSG00000143624.14 | 6.00 | 6.59 | -0.58 | 0.67 | 2.51624E-06 | 8.07355E-05 |
| 10400 | *PEMT* | ENSG00000133027.18 | 3.01 | 3.59 | -0.59 | 0.67 | 0.000732068 | 0.005009947 |
| 138162 | *C9orf116* | ENSG00000160345.13 | 3.27 | 3.86 | -0.59 | 0.67 | 0.000155877 | 0.001610159 |
| 10277 | *UBE4B* | ENSG00000130939.20 | 5.40 | 5.99 | -0.59 | 0.67 | 0.000216698 | 0.00204426 |
| 79084 | *WDR77* | ENSG00000116455.14 | 5.14 | 5.73 | -0.59 | 0.67 | 6.08755E-05 | 0.000823877 |
| 6604 | *SMARCD3* | ENSG00000082014.17 | 5.32 | 5.91 | -0.59 | 0.67 | 1.56588E-06 | 5.96504E-05 |
| 159371 | *SLC35G1* | ENSG00000176273.15 | 3.35 | 3.94 | -0.59 | 0.67 | 0.000636564 | 0.004523765 |
| 222183 | *SRRM3* | ENSG00000177679.16 | 5.46 | 6.05 | -0.59 | 0.66 | 7.05506E-06 | 0.000165971 |
| 5422 | *POLA1* | ENSG00000101868.13 | 4.80 | 5.39 | -0.59 | 0.66 | 3.5254E-05 | 0.000552325 |
| 200895 | *DHFR2* | ENSG00000178700.9 | 3.47 | 4.06 | -0.59 | 0.66 | 0.000135822 | 0.001454034 |
| 4306 | *NR3C2* | ENSG00000151623.15 | 3.13 | 3.72 | -0.59 | 0.66 | 0.000209831 | 0.002003404 |
| 9681 | *DEPDC5* | ENSG00000100150.20 | 4.51 | 5.10 | -0.59 | 0.66 | 0.000767857 | 0.005191337 |
| 84263 | *HSDL2* | ENSG00000119471.15 | 7.16 | 7.75 | -0.59 | 0.66 | 4.63343E-07 | 2.67474E-05 |
| 79989 | *TTC26* | ENSG00000105948.13 | 5.36 | 5.96 | -0.60 | 0.66 | 0.000152694 | 0.001588211 |
| 55709 | *KBTBD4* | ENSG00000123444.14 | 4.70 | 5.30 | -0.60 | 0.66 | 5.25256E-05 | 0.000736782 |
| 55384 | *MEG3* | ENSG00000214548.18 | 7.08 | 7.68 | -0.60 | 0.66 | 0.000829279 | 0.005514684 |
| 29882 | *ANAPC2* | ENSG00000176248.9 | 4.67 | 5.27 | -0.60 | 0.66 | 7.74451E-05 | 0.000976784 |
| 1910 | *EDNRB* | ENSG00000136160.17 | 3.89 | 4.49 | -0.60 | 0.66 | 0.001476876 | 0.00847989 |
| 114132 | *SIGLEC11* | ENSG00000161640.15 | 5.98 | 6.58 | -0.60 | 0.66 | 1.11753E-05 | 0.000232958 |
| 92105 | *INTS4* | ENSG00000149262.17 | 4.30 | 4.90 | -0.60 | 0.66 | 0.000906495 | 0.00586195 |
| 9202 | *ZMYM4* | ENSG00000146463.12 | 6.48 | 7.08 | -0.60 | 0.66 | 1.45974E-05 | 0.000284166 |
| 55076 | *TMEM45A* | ENSG00000181458.10 | 3.75 | 4.36 | -0.61 | 0.66 | 0.000446067 | 0.003493176 |
| 23474 | *ETHE1* | ENSG00000105755.8 | 2.25 | 2.86 | -0.61 | 0.66 | 0.000943232 | 0.006014006 |
| 23228 | *PLCL2* | ENSG00000154822.18 | 7.26 | 7.87 | -0.61 | 0.66 | 1.06754E-07 | 9.89059E-06 |
| 348110 | *ARPIN* | ENSG00000242498.8 | 5.08 | 5.70 | -0.61 | 0.66 | 0.000603481 | 0.004360931 |
| 56848 | *SPHK2* | ENSG00000063176.16 | 4.25 | 4.86 | -0.61 | 0.66 | 1.42687E-05 | 0.000281788 |
| 80310 | *PDGFD* | ENSG00000170962.13 | 6.24 | 6.85 | -0.61 | 0.65 | 1.48029E-06 | 5.78587E-05 |
| 10964 | *IFI44L* | ENSG00000137959.17 | 3.54 | 4.15 | -0.61 | 0.65 | 0.000261993 | 0.002380303 |
| 4670 | *HNRNPM* | ENSG00000099783.12 | 7.76 | 8.38 | -0.61 | 0.65 | 6.78411E-07 | 3.45161E-05 |
| 51092 | *SIDT2* | ENSG00000149577.16 | 4.12 | 4.73 | -0.61 | 0.65 | 0.000270829 | 0.00242824 |
| 9208 | *LRRFIP1* | ENSG00000124831.19 | 3.82 | 4.43 | -0.62 | 0.65 | 5.27584E-05 | 0.000738667 |
| 339229 | *OXLD1* | ENSG00000204237.5 | 3.07 | 3.68 | -0.62 | 0.65 | 0.000121171 | 0.001345942 |
| 80199 | *FUZ* | ENSG00000010361.14 | 3.61 | 4.22 | -0.62 | 0.65 | 0.00064881 | 0.004584742 |
| 9852 | *EPM2AIP1* | ENSG00000178567.8 | 6.01 | 6.63 | -0.62 | 0.65 | 6.20599E-05 | 0.000836137 |
| 129831 | *RBM45* | ENSG00000155636.15 | 3.04 | 3.66 | -0.62 | 0.65 | 0.001382016 | 0.00807108 |
| 92806 | *CENPBD1P* | ENSG00000177946.7 | 3.06 | 3.68 | -0.62 | 0.65 | 0.000504105 | 0.003815856 |
| 1468 | *SLC25A10* | ENSG00000183048.12 | 4.14 | 4.76 | -0.62 | 0.65 | 6.87939E-05 | 0.000900984 |
| 23229 | *ARHGEF9* | ENSG00000131089.17 | 4.25 | 4.87 | -0.62 | 0.65 | 0.000320155 | 0.002728682 |
| 9957 | *HS3ST1* | ENSG00000002587.10 | 2.91 | 3.53 | -0.62 | 0.65 | 0.001427143 | 0.008270269 |
| 2934 | *GSN* | ENSG00000148180.21 | 5.10 | 5.73 | -0.62 | 0.65 | 1.54213E-05 | 0.000294852 |
| 219854 | *TMEM218* | ENSG00000150433.10 | 3.20 | 3.82 | -0.62 | 0.65 | 0.00048611 | 0.00371299 |
| 51062 | *ATL1* | ENSG00000198513.14 | 4.14 | 4.77 | -0.63 | 0.65 | 0.000145542 | 0.001531863 |
| 55154 | *MSTO1* | ENSG00000125459.17 | 4.55 | 5.18 | -0.63 | 0.65 | 0.000137472 | 0.00146855 |
| 2938 | *GSTA1* | ENSG00000243955.6 | 8.85 | 9.48 | -0.63 | 0.65 | 1.06339E-06 | 4.63967E-05 |
| 6522 | *SLC4A2* | ENSG00000164889.15 | 6.78 | 7.40 | -0.63 | 0.65 | 4.52986E-06 | 0.000120547 |
| 168400 | *DDX53* | ENSG00000184735.7 | 2.73 | 3.35 | -0.63 | 0.65 | 0.000459614 | 0.003568722 |
| 55532 | *SLC30A10* | ENSG00000196660.11 | 7.37 | 8.00 | -0.63 | 0.65 | 9.95184E-07 | 4.5069E-05 |
| 4651 | *MYO10* | ENSG00000145555.15 | 7.51 | 8.14 | -0.63 | 0.65 | 2.63514E-06 | 8.29157E-05 |
| 5092 | *PCBD1* | ENSG00000166228.9 | 3.49 | 4.12 | -0.63 | 0.65 | 0.000164392 | 0.001668262 |
| 26149 | *ZNF658* | ENSG00000274349.5 | 3.29 | 3.92 | -0.63 | 0.65 | 0.000307963 | 0.002662566 |
| 412 | *STS* | ENSG00000101846.9 | 3.57 | 4.20 | -0.63 | 0.65 | 0.00015358 | 0.001592657 |
| 55860 | *ACTR10* | ENSG00000131966.14 | 5.47 | 6.10 | -0.63 | 0.64 | 1.10794E-05 | 0.000232132 |
| 4065 | *LY75* | ENSG00000054219.11 | 4.76 | 5.39 | -0.63 | 0.64 | 7.13221E-05 | 0.000921456 |
| 29903 | *CCDC106* | ENSG00000173581.8 | 3.32 | 3.96 | -0.63 | 0.64 | 0.000351812 | 0.002931283 |
| 23532 | *PRAME* | ENSG00000185686.18 | 4.71 | 5.34 | -0.64 | 0.64 | 4.41674E-05 | 0.000655696 |
| 7390 | *UROS* | ENSG00000188690.15 | 4.79 | 5.43 | -0.64 | 0.64 | 7.67372E-06 | 0.000176378 |
| 133121 | *ENPP6* | ENSG00000164303.11 | 2.56 | 3.19 | -0.64 | 0.64 | 0.000842541 | 0.00557079 |
| 57727 | *NCOA5* | ENSG00000124160.12 | 5.74 | 6.38 | -0.64 | 0.64 | 8.34176E-06 | 0.000186868 |
| 9609 | *RAB36* | ENSG00000100228.14 | 5.65 | 6.28 | -0.64 | 0.64 | 1.27441E-05 | 0.000257092 |
| 11022 | *TDRKH* | ENSG00000182134.17 | 3.51 | 4.15 | -0.64 | 0.64 | 0.000419186 | 0.003334165 |
| 92840 | *REEP6* | ENSG00000115255.12 | 4.47 | 5.11 | -0.64 | 0.64 | 3.57052E-06 | 0.00010286 |
| 5557 | *PRIM1* | ENSG00000198056.15 | 3.90 | 4.54 | -0.64 | 0.64 | 0.001200802 | 0.007252648 |
| 55644 | *OSGEP* | ENSG00000092094.11 | 3.38 | 4.01 | -0.64 | 0.64 | 0.000611874 | 0.004404609 |
| 55160 | *ARHGEF10L* | ENSG00000074964.17 | 3.17 | 3.81 | -0.64 | 0.64 | 0.000521148 | 0.00391683 |
| 10810 | *WASF3* | ENSG00000132970.14 | 5.54 | 6.18 | -0.64 | 0.64 | 0.000955849 | 0.006076385 |
| 65987 | *KCTD14* | ENSG00000151364.17 | 2.94 | 3.58 | -0.64 | 0.64 | 8.63046E-05 | 0.001051417 |
| 10778 | *ZNF271P* | ENSG00000257267.4 | 4.53 | 5.17 | -0.64 | 0.64 | 7.16784E-06 | 0.000167313 |
| 25925 | *ZNF521* | ENSG00000198795.11 | 5.60 | 6.24 | -0.64 | 0.64 | 0.000397343 | 0.003195991 |
| 138428 | *PTRH1* | ENSG00000187024.15 | 2.53 | 3.17 | -0.64 | 0.64 | 0.001072426 | 0.006637542 |
| 56911 | *MAP3K7CL* | ENSG00000156265.16 | 3.23 | 3.87 | -0.64 | 0.64 | 0.00101954 | 0.006376652 |
| 130535 | *KCTD18* | ENSG00000155729.13 | 2.85 | 3.50 | -0.64 | 0.64 | 0.000280251 | 0.002491552 |
| 400713 | *ZNF880* | ENSG00000221923.9 | 3.45 | 4.09 | -0.64 | 0.64 | 0.000186018 | 0.001822422 |
| 3065 | *HDAC1* | ENSG00000116478.12 | 6.31 | 6.96 | -0.65 | 0.64 | 1.74406E-06 | 6.38453E-05 |
| 10046 | *MAMLD1* | ENSG00000013619.15 | 2.85 | 3.49 | -0.65 | 0.64 | 0.001673383 | 0.009336731 |
| 29940 | *DSE* | ENSG00000111817.19 | 3.18 | 3.82 | -0.65 | 0.64 | 0.001320357 | 0.007780618 |
| 56896 | *DPYSL5* | ENSG00000157851.17 | 4.88 | 5.53 | -0.65 | 0.64 | 0.001368941 | 0.008022818 |
| 221883 | *HOXA11-AS* | ENSG00000240990.10 | 4.59 | 5.24 | -0.65 | 0.64 | 2.128E-05 | 0.000377532 |
| 55364 | *IMPACT* | ENSG00000154059.11 | 5.49 | 6.14 | -0.65 | 0.64 | 0.001075717 | 0.006649683 |
| 100129461 | *LYRM4-AS1* | ENSG00000272142.4 | 2.42 | 3.07 | -0.65 | 0.64 | 0.000254056 | 0.002325074 |
| 5836 | *PYGL* | ENSG00000100504.17 | 4.53 | 5.18 | -0.65 | 0.64 | 1.8678E-06 | 6.70597E-05 |
| 7733 | *ZNF180* | ENSG00000167384.11 | 2.91 | 3.56 | -0.65 | 0.64 | 0.00015765 | 0.001620661 |
| 3482 | *IGF2R* | ENSG00000197081.16 | 7.63 | 8.28 | -0.65 | 0.64 | 0.000102039 | 0.001186288 |
| 9308 | *CD83* | ENSG00000112149.10 | 7.32 | 7.97 | -0.65 | 0.64 | 5.71157E-08 | 6.12322E-06 |
| 80208 | *SPG11* | ENSG00000104133.16 | 5.72 | 6.37 | -0.66 | 0.64 | 1.10186E-05 | 0.000231298 |

| | | | | | | | | |
|---|---|---|---|---|---|---|---|---|
| 128853 | *DUSP15* | ENSG00000149599.16 | 3.17 | 3.83 | -0.66 | 0.63 | 0.000549478 | 0.004072648 |
| 27148 | *STK36* | ENSG00000163482.12 | 3.23 | 3.89 | -0.66 | 0.63 | 0.000693114 | 0.004811727 |
| 10533 | *ATG7* | ENSG00000197548.13 | 3.72 | 4.39 | -0.66 | 0.63 | 0.001152741 | 0.00701033 |
| 79183 | *TTPAL* | ENSG00000124120.11 | 4.53 | 5.19 | -0.66 | 0.63 | 4.02656E-05 | 0.000611068 |
| 388341 | *LRRC75A* | ENSG00000181350.12 | 4.19 | 4.86 | -0.67 | 0.63 | 2.8204E-05 | 0.000468611 |
| 150726 | *FBXO41* | ENSG00000163013.12 | 3.74 | 4.40 | -0.67 | 0.63 | 0.000415429 | 0.003320114 |
| 54872 | *PIGG* | ENSG00000174227.16 | 5.87 | 6.54 | -0.67 | 0.63 | 1.99168E-06 | 6.98526E-05 |
| 79622 | *SNRNP25* | ENSG00000161981.11 | 4.98 | 5.65 | -0.67 | 0.63 | 1.79019E-06 | 6.51661E-05 |
| 90806 | *ANGEL2* | ENSG00000174606.14 | 4.89 | 5.55 | -0.67 | 0.63 | 2.39268E-05 | 0.000413253 |
| 25903 | *OLFML2B* | ENSG00000162745.11 | 2.44 | 3.11 | -0.67 | 0.63 | 0.000150241 | 0.001565944 |
| 93643 | *TJAP1* | ENSG00000137221.14 | 4.16 | 4.84 | -0.67 | 0.63 | 0.000755409 | 0.005137261 |
| 2135 | *EXTL2* | ENSG00000162694.14 | 5.64 | 6.31 | -0.67 | 0.63 | 2.09163E-06 | 7.2003E-05 |
| 10518 | *CIB2* | ENSG00000136425.14 | 2.95 | 3.62 | -0.67 | 0.63 | 0.000135439 | 0.001452004 |
| 378708 | *CENPS* | ENSG00000175279.22 | 2.11 | 2.78 | -0.67 | 0.63 | 0.000578724 | 0.004230916 |
| 5001 | *ORC5* | ENSG00000164815.11 | 4.59 | 5.26 | -0.67 | 0.63 | 1.87655E-06 | 6.70597E-05 |
| 63920 | *ZBED8* | ENSG00000221886.4 | 3.17 | 3.85 | -0.67 | 0.63 | 6.79769E-05 | 0.000896543 |
| 64789 | *EXO5* | ENSG00000164002.12 | 2.99 | 3.66 | -0.68 | 0.63 | 0.000135684 | 0.001453591 |
| 23542 | *MAPK8IP2* | ENSG00000008735.14 | 3.13 | 3.81 | -0.68 | 0.63 | 0.000183927 | 0.001807649 |
| 84981 | *MIR22HG* | ENSG00000186594.15 | 2.50 | 3.18 | -0.68 | 0.62 | 0.001202096 | 0.007257545 |
| 7507 | *XPA* | ENSG00000136936.11 | 3.30 | 3.98 | -0.68 | 0.62 | 0.000156579 | 0.001614071 |
| 101927111 | *SUCLG2-DT* | ENSG00000241316.8 | 1.46 | 2.14 | -0.68 | 0.62 | 0.000860514 | 0.005664675 |
| 5257 | *PHKB* | ENSG00000102893.16 | 6.78 | 7.46 | -0.68 | 0.62 | 2.11355E-06 | 7.20414E-05 |
| 4820 | *NKTR* | ENSG00000114857.19 | 5.18 | 5.86 | -0.68 | 0.62 | 2.06863E-05 | 0.000369619 |
| 6660 | *SOX5* | ENSG00000134532.19 | 2.95 | 3.63 | -0.68 | 0.62 | 0.001701201 | 0.009453283 |
| 81790 | *RNF170* | ENSG00000120925.16 | 3.51 | 4.19 | -0.68 | 0.62 | 0.000729559 | 0.004997699 |
| 728819 | *C1GALT1C1L* | ENSG00000223658.8 | 2.85 | 3.54 | -0.68 | 0.62 | 7.09689E-05 | 0.000920633 |
| 8850 | *KAT2B* | ENSG00000114166.8 | 3.68 | 4.36 | -0.68 | 0.62 | 0.000154244 | 0.001597691 |
| 10785 | *WDR4* | ENSG00000160193.12 | 2.55 | 3.24 | -0.69 | 0.62 | 0.0007139 | 0.004921876 |
| 2184 | *FAH* | ENSG00000103876.14 | 3.79 | 4.47 | -0.69 | 0.62 | 0.000120449 | 0.0013451 |
| 60529 | *ALX4* | ENSG00000052850.8 | 2.29 | 2.98 | -0.69 | 0.62 | 0.000112568 | 0.001274161 |
| 54556 | *ING3* | ENSG00000071243.16 | 4.40 | 5.09 | -0.69 | 0.62 | 2.76169E-06 | 8.54897E-05 |
| 11170 | *FAM107A* | ENSG00000168309.18 | 4.79 | 5.48 | -0.69 | 0.62 | 2.06674E-06 | 7.14739E-05 |
| 23187 | *PHLDB1* | ENSG00000019144.20 | 5.61 | 6.30 | -0.69 | 0.62 | 2.43243E-05 | 0.00041803 |
| 3321 | *IGSF3* | ENSG00000143061.18 | 4.94 | 5.64 | -0.69 | 0.62 | 1.32349E-05 | 0.000264856 |
| 2639 | *GCDH* | ENSG00000105607.13 | 4.32 | 5.01 | -0.69 | 0.62 | 9.04918E-05 | 0.001084817 |
| 57118 | *CAMK1D* | ENSG00000183049.14 | 2.53 | 3.22 | -0.69 | 0.62 | 0.001153356 | 0.007011226 |
| 5575 | *PRKAR1B* | ENSG00000188191.15 | 4.13 | 4.82 | -0.69 | 0.62 | 3.07777E-06 | 9.2946E-05 |
| 23586 | *DDX58* | ENSG00000107201.11 | 1.83 | 2.53 | -0.69 | 0.62 | 0.001746504 | 0.009608973 |
| 2972 | *BRF1* | ENSG00000185024.18 | 4.58 | 5.28 | -0.69 | 0.62 | 0.000911209 | 0.005874974 |
| 4292 | *MLH1* | ENSG00000076242.16 | 4.73 | 5.43 | -0.70 | 0.62 | 6.93374E-06 | 0.000164405 |
| 11267 | *SNF8* | ENSG00000159210.10 | 5.22 | 5.91 | -0.70 | 0.62 | 5.32793E-06 | 0.000135793 |
| 25927 | *CNRIP1* | ENSG00000119865.9 | 2.23 | 2.93 | -0.70 | 0.62 | 0.001506632 | 0.008614492 |
| 23428 | *SLC7A8* | ENSG00000092068.21 | 6.53 | 7.22 | -0.70 | 0.62 | 2.90969E-07 | 1.93958E-05 |
| 360200 | *TMPRSS9* | ENSG00000178297.14 | 3.32 | 4.01 | -0.70 | 0.62 | 0.000588059 | 0.00428028 |
| 64925 | *CCDC71* | ENSG00000177352.10 | 3.56 | 4.25 | -0.70 | 0.62 | 1.12124E-05 | 0.000233408 |
| 11178 | *LZTS1* | ENSG00000061337.15 | 2.47 | 3.17 | -0.70 | 0.62 | 0.000476937 | 0.003659685 |
| 6988 | *TCTA* | ENSG00000145022.5 | 4.58 | 5.29 | -0.70 | 0.62 | 7.91591E-06 | 0.000180837 |
| 10439 | *OLFM1* | ENSG00000130558.20 | 4.37 | 5.07 | -0.70 | 0.61 | 1.47021E-06 | 5.76145E-05 |
| 2874 | *GPS2* | ENSG00000132522.16 | 4.76 | 5.46 | -0.70 | 0.61 | 7.40725E-06 | 0.000171567 |
| 344595 | *DUBR* | ENSG00000243701.8 | 5.05 | 5.75 | -0.70 | 0.61 | 0.000145215 | 0.00153057 |
| 2264 | *FGFR4* | ENSG00000160867.15 | 4.31 | 5.02 | -0.70 | 0.61 | 5.52129E-05 | 0.000765888 |
| 84152 | *PPP1R1B* | ENSG00000131771.14 | 2.75 | 3.46 | -0.70 | 0.61 | 0.000189487 | 0.001850369 |
| 60490 | *PPCDC* | ENSG00000138621.12 | 2.85 | 3.56 | -0.71 | 0.61 | 0.000659613 | 0.004641412 |
| 6431 | *SRSF6* | ENSG00000124193.16 | 5.93 | 6.64 | -0.71 | 0.61 | 6.54525E-06 | 0.000157938 |
| 25999 | *CLIP3* | ENSG00000105270.15 | 5.57 | 6.27 | -0.71 | 0.61 | 0.000250263 | 0.002295969 |
| 2798 | *GNRHR* | ENSG00000109163.7 | 4.95 | 5.66 | -0.71 | 0.61 | 5.03863E-07 | 2.85793E-05 |
| 83401 | *ELOVL3* | ENSG00000119915.5 | 3.09 | 3.80 | -0.71 | 0.61 | 7.66061E-06 | 0.000176347 |
| 100289274 | *DNAJC3-DT* | ENSG00000247400.5 | 3.08 | 3.79 | -0.71 | 0.61 | 0.000122902 | 0.001357352 |
| 29099 | *COMMD9* | ENSG00000110442.12 | 3.81 | 4.52 | -0.71 | 0.61 | 0.001181322 | 0.007155151 |
| 388566 | *ZNF470* | ENSG00000197016.12 | 3.29 | 4.01 | -0.71 | 0.61 | 0.000251637 | 0.002306526 |
| 730101 | *LOC730101* | ENSG00000216775.4 | 4.71 | 5.43 | -0.71 | 0.61 | 3.19659E-06 | 9.61476E-05 |
| 9617 | *MTRF1* | ENSG00000120662.16 | 1.74 | 2.46 | -0.71 | 0.61 | 0.001134088 | 0.006930009 |
| 7753 | *ZNF202* | ENSG00000166261.11 | 3.81 | 4.53 | -0.71 | 0.61 | 6.35376E-05 | 0.000848486 |
| 54962 | *TIPIN* | ENSG00000075131.10 | 3.23 | 3.94 | -0.71 | 0.61 | 3.82747E-05 | 0.000586788 |
| 6932 | *TCF7* | ENSG00000081059.20 | 4.39 | 5.10 | -0.71 | 0.61 | 3.41308E-06 | 0.000100642 |
| 389023 | *DPP10-AS1* | ENSG00000235026.7 | 3.28 | 4.00 | -0.71 | 0.61 | 5.19621E-05 | 0.000732988 |
| 9227 | *LRAT* | ENSG00000121207.12 | 3.74 | 4.46 | -0.72 | 0.61 | 0.000136061 | 0.001455555 |
| 79643 | *CHMP6* | ENSG00000176108.9 | 3.73 | 4.45 | -0.72 | 0.61 | 2.04242E-05 | 0.000366244 |
| 29945 | *ANAPC4* | ENSG00000053900.11 | 3.88 | 4.60 | -0.72 | 0.61 | 2.11719E-05 | 0.000376058 |
| 1666 | *DECR1* | ENSG00000104325.7 | 5.41 | 6.13 | -0.72 | 0.61 | 1.13194E-06 | 4.82652E-05 |
| 285613 | *RELL2* | ENSG00000164620.9 | 2.92 | 3.64 | -0.72 | 0.61 | 0.000295529 | 0.002583735 |
| 84456 | *L3MBTL3* | ENSG00000198945.8 | 5.33 | 6.05 | -0.72 | 0.61 | 1.40976E-06 | 5.59764E-05 |
| 6936 | *GCFC2* | ENSG00000005436.14 | 2.01 | 2.73 | -0.72 | 0.61 | 0.000184691 | 0.001811778 |
| 5152 | *PDE9A* | ENSG00000160191.18 | 4.03 | 4.75 | -0.72 | 0.61 | 2.93341E-05 | 0.000481702 |
| 119392 | *SFR1* | ENSG00000156384.15 | 1.72 | 2.44 | -0.72 | 0.61 | 0.000615003 | 0.004422893 |
| 11221 | *DUSP10* | ENSG00000143507.18 | 2.07 | 2.79 | -0.72 | 0.61 | 0.001088081 | 0.006698526 |
| 26521 | *TIMM8B* | ENSG00000150779.12 | 4.87 | 5.60 | -0.72 | 0.61 | 1.54567E-06 | 5.91808E-05 |
| 118738 | *ZNF488* | ENSG00000265763.4 | 3.20 | 3.93 | -0.73 | 0.60 | 0.000124372 | 0.001366547 |
| 57102 | *C12orf4* | ENSG00000047621.12 | 3.45 | 4.18 | -0.73 | 0.60 | 0.000680309 | 0.004742573 |
| 253039 | *CUTALP* | ENSG00000226752.10 | 4.93 | 5.66 | -0.73 | 0.60 | 9.32378E-06 | 0.000201935 |
| 23613 | *ZMYND8* | ENSG00000101040.20 | 6.32 | 7.05 | -0.73 | 0.60 | 1.75267E-05 | 0.000325568 |
| 116987 | *AGAP1* | ENSG00000157985.19 | 3.77 | 4.51 | -0.73 | 0.60 | 0.001311556 | 0.007731793 |
| 284695 | *ZNF326* | ENSG00000162664.17 | 5.26 | 5.99 | -0.73 | 0.60 | 1.84397E-06 | 6.66896E-05 |
| 90203 | *SNX21* | ENSG00000124104.19 | 3.94 | 4.68 | -0.73 | 0.60 | 6.08135E-06 | 0.000149115 |
| 105370333 | *PCCA-DT* | ENSG00000274605.3 | 2.45 | 3.19 | -0.73 | 0.60 | 0.000453997 | 0.003535104 |
| 6695 | *SPOCK1* | ENSG00000152377.14 | 6.38 | 7.12 | -0.74 | 0.60 | 3.63306E-07 | 2.23478E-05 |
| 88 | *ACTN2* | ENSG00000077522.15 | 4.71 | 5.45 | -0.74 | 0.60 | 0.000265614 | 0.002401908 |
| 2648 | *KAT2A* | ENSG00000108773.11 | 3.96 | 4.70 | -0.74 | 0.60 | 4.41565E-05 | 0.000655696 |
| 379025 | *PSMA3-AS1* | ENSG00000257621.9 | 3.93 | 4.66 | -0.74 | 0.60 | 0.001744986 | 0.009604141 |

| | | | | | | | | |
|---|---|---|---|---|---|---|---|---|
| 348094 | *ANKDD1A* | ENSG00000166839.17 | 2.80 | 3.54 | -0.74 | 0.60 | 0.000831965 | 0.005517878 |
| 92104 | *TTC30A* | ENSG00000197557.7 | 4.55 | 5.29 | -0.74 | 0.60 | 1.19346E-06 | 5.00352E-05 |
| 196740 | *VSTM4* | ENSG00000165633.13 | 5.87 | 6.61 | -0.74 | 0.60 | 8.4508E-08 | 8.23623E-06 |
| 146198 | *ZFP90* | ENSG00000184939.16 | 3.92 | 4.66 | -0.74 | 0.60 | 0.001744476 | 0.009604141 |
| 9704 | *DHX34* | ENSG00000134815.19 | 3.15 | 3.90 | -0.75 | 0.60 | 9.30619E-05 | 0.001108545 |
| 9557 | *CHD1L* | ENSG00000131778.20 | 4.95 | 5.70 | -0.75 | 0.59 | 1.11071E-06 | 4.77671E-05 |
| 23211 | *ZC3H4* | ENSG00000130749.10 | 5.11 | 5.86 | -0.75 | 0.59 | 2.46174E-06 | 7.96299E-05 |
| 54361 | *WNT4* | ENSG00000162552.15 | 5.59 | 6.34 | -0.75 | 0.59 | 1.61562E-06 | 6.10802E-05 |
| 374393 | *FAM111B* | ENSG00000189057.11 | 5.16 | 5.91 | -0.75 | 0.59 | 6.10342E-06 | 0.000149196 |
| 56122 | *PCDHB14* | ENSG00000120327.7 | 2.36 | 3.11 | -0.75 | 0.59 | 0.001412647 | 0.008211821 |
| 10628 | *TXNIP* | ENSG00000265972.6 | 7.50 | 8.25 | -0.75 | 0.59 | 6.86577E-08 | 7.03548E-06 |
| 51313 | *GASK1B* | ENSG00000164125.16 | 5.66 | 6.41 | -0.75 | 0.59 | 6.35641E-06 | 0.000154425 |
| 79810 | *PTCD2* | ENSG00000049883.15 | 4.04 | 4.80 | -0.75 | 0.59 | 3.4225E-05 | 0.000541686 |
| 79695 | *GALNT12* | ENSG00000119514.7 | 2.12 | 2.87 | -0.75 | 0.59 | 0.001650667 | 0.0092478 |
| 23649 | *POLA2* | ENSG00000014138.9 | 5.32 | 6.08 | -0.76 | 0.59 | 0.000300928 | 0.002622894 |
| 147949 | *ZNF583* | ENSG00000198440.9 | 2.21 | 2.96 | -0.76 | 0.59 | 0.001370727 | 0.008024314 |
| 83989 | *FAM172A* | ENSG00000113391.19 | 4.01 | 4.77 | -0.76 | 0.59 | 9.52622E-05 | 0.001132058 |
| 66008 | *TRAK2* | ENSG00000115993.13 | 5.59 | 6.35 | -0.76 | 0.59 | 5.28047E-06 | 0.000135247 |
| 64943 | *NT5DC2* | ENSG00000168268.11 | 5.30 | 6.06 | -0.76 | 0.59 | 1.03555E-05 | 0.000220501 |
| 101930085 | *HERPUD2-AS* | ENSG00000271122.1 | 3.11 | 3.87 | -0.76 | 0.59 | 1.19564E-05 | 0.000246163 |
| 4163 | *MCC* | ENSG00000171444.18 | 2.50 | 3.26 | -0.76 | 0.59 | 0.001359691 | 0.007977954 |
| 548596 | *CKMT1A* | ENSG00000223572.10 | 3.46 | 4.22 | -0.77 | 0.59 | 0.000672443 | 0.004705219 |
| 54948 | *MRPL16* | ENSG00000166902.5 | 3.96 | 4.73 | -0.77 | 0.59 | 1.43789E-05 | 0.000282848 |
| 169981 | *SPIN3* | ENSG00000204271.13 | 3.24 | 4.00 | -0.77 | 0.59 | 1.29493E-05 | 0.000260531 |
| 26267 | *FBXO10* | ENSG00000147912.13 | 6.24 | 7.01 | -0.77 | 0.59 | 3.70577E-05 | 0.000573992 |
| 5241 | *PGR* | ENSG00000082175.16 | 5.18 | 5.96 | -0.77 | 0.59 | 0.000541019 | 0.004027857 |
| 124930 | *ANKRD13B* | ENSG00000198720.13 | 2.75 | 3.52 | -0.77 | 0.59 | 7.43389E-05 | 0.000946355 |
| 56922 | *MCCC1* | ENSG00000078070.14 | 4.52 | 5.29 | -0.77 | 0.58 | 7.91445E-07 | 3.77105E-05 |
| 8677 | *STX10* | ENSG00000104915.15 | 3.20 | 3.97 | -0.78 | 0.58 | 7.06904E-05 | 0.00091899 |
| 1174 | *AP1S1* | ENSG00000106367.15 | 7.24 | 8.02 | -0.78 | 0.58 | 3.1539E-08 | 4.21588E-06 |
| 151613 | *TTC14* | ENSG00000163728.11 | 3.97 | 4.75 | -0.78 | 0.58 | 2.9408E-05 | 0.000482388 |
| 8504 | *PEX3* | ENSG00000034693.15 | 4.13 | 4.90 | -0.78 | 0.58 | 5.7876E-07 | 3.17519E-05 |
| 93611 | *FBXO44* | ENSG00000132879.14 | 3.25 | 4.02 | -0.78 | 0.58 | 8.87709E-05 | 0.001075335 |
| 10940 | *POP1* | ENSG00000104356.11 | 4.21 | 4.99 | -0.78 | 0.58 | 1.44006E-05 | 0.000282903 |
| 56935 | *SMCO4* | ENSG00000166002.7 | 2.42 | 3.20 | -0.78 | 0.58 | 0.000416985 | 0.003325463 |
| 585 | *BBS4* | ENSG00000140463.14 | 4.43 | 5.22 | -0.78 | 0.58 | 3.36263E-06 | 9.95458E-05 |
| 23208 | *SYT11* | ENSG00000132718.9 | 2.19 | 2.98 | -0.78 | 0.58 | 0.000348266 | 0.002915293 |
| 729991 | *BORCS8* | ENSG00000254901.8 | 2.73 | 3.51 | -0.78 | 0.58 | 0.00047196 | 0.003636186 |
| 79736 | *TEFM* | ENSG00000172171.11 | 2.27 | 3.05 | -0.78 | 0.58 | 0.00012761 | 0.001388907 |
| 84674 | *CARD6* | ENSG00000132357.14 | 4.50 | 5.29 | -0.79 | 0.58 | 3.46681E-06 | 0.000101394 |
| 150 | *ADRA2A* | ENSG00000150594.7 | 2.51 | 3.30 | -0.79 | 0.58 | 0.001134455 | 0.006930009 |
| 2845 | *GPR22* | ENSG00000172209.6 | 4.54 | 5.32 | -0.79 | 0.58 | 5.12148E-05 | 0.000725857 |
| 4900 | *NRGN* | ENSG00000154146.13 | 2.04 | 2.82 | -0.79 | 0.58 | 0.000159016 | 0.001630241 |
| 23657 | *SLC7A11* | ENSG00000151012.13 | 5.11 | 5.90 | -0.79 | 0.58 | 1.21415E-05 | 0.000248254 |
| 255928 | *SYT14* | ENSG00000143469.20 | 2.67 | 3.46 | -0.79 | 0.58 | 0.000669318 | 0.004689916 |
| 112268269 | *LOC1122682ϵ* | ENSG00000268858.2 | 2.00 | 2.79 | -0.79 | 0.58 | 0.000156793 | 0.001615175 |
| 60401 | *EDA2R* | ENSG00000131080.15 | 2.70 | 3.50 | -0.79 | 0.58 | 6.27657E-05 | 0.000842988 |
| 79979 | *TRMT2B* | ENSG00000188917.15 | 3.73 | 4.53 | -0.80 | 0.58 | 8.44808E-05 | 0.001035372 |
| 3216 | *HOXB6* | ENSG00000108511.10 | 2.85 | 3.64 | -0.80 | 0.58 | 0.000290606 | 0.002553692 |
| 51196 | *PLCE1* | ENSG00000138193.17 | 2.78 | 3.58 | -0.80 | 0.58 | 0.001192758 | 0.007206968 |
| 56892 | *TCIM* | ENSG00000176907.5 | 3.67 | 4.47 | -0.80 | 0.58 | 6.91183E-06 | 0.000164341 |
| 5287 | *PIK3C2B* | ENSG00000133056.14 | 1.81 | 2.61 | -0.80 | 0.58 | 0.000782249 | 0.005267282 |
| 55157 | *DARS2* | ENSG00000117593.12 | 5.84 | 6.64 | -0.80 | 0.58 | 3.32238E-08 | 4.33614E-06 |
| 10795 | *ZNF268* | ENSG00000090612.22 | 4.46 | 5.27 | -0.80 | 0.57 | 1.82742E-05 | 0.000336536 |
| 55622 | *TTC27* | ENSG00000018699.13 | 3.17 | 3.98 | -0.81 | 0.57 | 0.000558991 | 0.0041228 |
| 340348 | *TSPAN33* | ENSG00000158457.6 | 3.08 | 3.89 | -0.81 | 0.57 | 0.000250236 | 0.002295969 |
| 3936 | *LCP1* | ENSG00000136167.15 | 4.85 | 5.66 | -0.81 | 0.57 | 3.53223E-06 | 0.000102149 |
| 9580 | *SOX13* | ENSG00000143842.15 | 4.50 | 5.31 | -0.81 | 0.57 | 3.79881E-07 | 2.29905E-05 |
| 149233 | *IL23R* | ENSG00000162594.16 | 2.23 | 3.04 | -0.81 | 0.57 | 0.000640208 | 0.004545354 |
| 259230 | *SGMS1* | ENSG00000198964.14 | 4.73 | 5.54 | -0.81 | 0.57 | 0.001299069 | 0.007684019 |
| 26268 | *FBXO9* | ENSG00000112146.17 | 5.60 | 6.41 | -0.81 | 0.57 | 7.08571E-07 | 3.55095E-05 |
| 404093 | *CUEDC1* | ENSG00000180891.13 | 3.57 | 4.38 | -0.81 | 0.57 | 0.000308558 | 0.002665809 |
| 10654 | *PMVK* | ENSG00000163344.6 | 3.90 | 4.71 | -0.82 | 0.57 | 7.32955E-06 | 0.00017003 |
| 6103 | *RPGR* | ENSG00000156313.15 | 2.36 | 3.18 | -0.82 | 0.57 | 0.000640513 | 0.004545369 |
| 83879 | *CDCA7* | ENSG00000144354.14 | 3.65 | 4.47 | -0.82 | 0.57 | 2.36228E-06 | 7.75832E-05 |
| 3623 | *INHA* | ENSG00000123999.5 | 2.06 | 2.87 | -0.82 | 0.57 | 0.000221965 | 0.002083476 |
| 594 | *BCKDHB* | ENSG00000083123.15 | 3.37 | 4.18 | -0.82 | 0.57 | 0.000161369 | 0.001646574 |
| 50863 | *NTM* | ENSG00000182667.15 | 3.17 | 3.99 | -0.82 | 0.57 | 0.000157765 | 0.001620735 |
| 7587 | *ZNF37A* | ENSG00000075407.19 | 4.26 | 5.08 | -0.82 | 0.57 | 1.42498E-06 | 5.64313E-05 |
| 114801 | *TMEM200A* | ENSG00000164484.12 | 5.18 | 6.00 | -0.82 | 0.57 | 2.9191E-07 | 1.93958E-05 |
| 8908 | *GYG2* | ENSG00000056998.20 | 2.97 | 3.79 | -0.82 | 0.57 | 3.49986E-06 | 0.000101801 |
| 29104 | *N6AMT1* | ENSG00000156239.12 | 3.25 | 4.07 | -0.82 | 0.57 | 0.000807796 | 0.005402944 |
| 2868 | *GRK4* | ENSG00000125388.20 | 2.56 | 3.39 | -0.82 | 0.57 | 0.000124992 | 0.001371089 |
| 9941 | *EXOG* | ENSG00000157036.13 | 3.77 | 4.59 | -0.82 | 0.56 | 0.000669142 | 0.004689916 |
| 463 | *ZFHX3* | ENSG00000140836.17 | 3.92 | 4.74 | -0.83 | 0.56 | 0.000392563 | 0.003166025 |
| 100329135 | *TRPC5OS* | ENSG00000204025.8 | 4.21 | 5.03 | -0.83 | 0.56 | 7.22321E-06 | 0.000167822 |
| 10481 | *HOXB13* | ENSG00000159184.8 | 3.29 | 4.12 | -0.83 | 0.56 | 1.66942E-05 | 0.000313988 |
| 220963 | *SLC16A9* | ENSG00000165449.12 | 1.05 | 1.88 | -0.83 | 0.56 | 0.000562952 | 0.004137778 |
| 285382 | *C3orf70* | ENSG00000187068.3 | 2.31 | 3.14 | -0.83 | 0.56 | 0.000124108 | 0.001365695 |
| 92369 | *SPSB4* | ENSG00000175093.5 | 1.85 | 2.68 | -0.83 | 0.56 | 0.000125151 | 0.001371089 |
| 4684 | *NCAM1* | ENSG00000149294.17 | 6.01 | 6.84 | -0.83 | 0.56 | 1.49922E-06 | 5.82618E-05 |
| 90673 | *PPP1R3E* | ENSG00000235194.9 | 3.45 | 4.28 | -0.83 | 0.56 | 0.000593102 | 0.004304577 |
| 582 | *BBS1* | ENSG00000174483.20 | 4.20 | 5.03 | -0.83 | 0.56 | 2.21161E-06 | 7.44261E-05 |
| 54841 | *BIVM* | ENSG00000134897.14 | 3.96 | 4.79 | -0.83 | 0.56 | 1.40453E-05 | 0.000278475 |
| 56952 | *PRTFDC1* | ENSG00000099256.19 | 3.40 | 4.23 | -0.83 | 0.56 | 4.47061E-05 | 0.00066173 |
| 2898 | *GRIK2* | ENSG00000164418.22 | 4.73 | 5.56 | -0.84 | 0.56 | 6.64802E-06 | 0.000159904 |
| 26022 | *TMEM98* | ENSG00000006042.12 | 3.76 | 4.59 | -0.84 | 0.56 | 7.80673E-05 | 0.000980512 |
| 84277 | *DNAJC30* | ENSG00000176410.8 | 3.43 | 4.27 | -0.84 | 0.56 | 2.20737E-05 | 0.000388059 |
| 5198 | *PFAS* | ENSG00000178921.14 | 4.09 | 4.93 | -0.84 | 0.56 | 0.000334796 | 0.00282301 |

| | | | | | | | | |
|---|---|---|---|---|---|---|---|---|
| 81033 | *KCNH6* | ENSG00000173826.15 | 1.53 | 2.37 | -0.85 | 0.56 | 0.000732349 | 0.005009947 |
| 79641 | *ROGDI* | ENSG00000067836.13 | 3.30 | 4.15 | -0.85 | 0.56 | 7.33026E-05 | 0.000939667 |
| 3669 | *ISG20* | ENSG00000172183.16 | 1.62 | 2.47 | -0.85 | 0.56 | 0.000482815 | 0.003691576 |
| 55170 | *PRMT6* | ENSG00000198890.9 | 4.45 | 5.30 | -0.85 | 0.56 | 8.4928E-07 | 3.99587E-05 |
| 85440 | *DOCK7* | ENSG00000116641.18 | 5.72 | 6.56 | -0.85 | 0.56 | 0.000215954 | 0.002039349 |
| 125113 | *KRT222* | ENSG00000213424.9 | 3.12 | 3.97 | -0.85 | 0.56 | 4.0341E-05 | 0.000611594 |
| 4773 | *NFATC2* | ENSG00000101096.20 | 2.20 | 3.05 | -0.85 | 0.56 | 0.000211306 | 0.002012346 |
| 286101 | *ZNF252P* | ENSG00000196922.11 | 3.80 | 4.65 | -0.85 | 0.55 | 8.14796E-05 | 0.001010488 |
| 10813 | *UTP14A* | ENSG00000156697.13 | 3.69 | 4.54 | -0.85 | 0.55 | 1.92313E-06 | 6.82372E-05 |
| 2053 | *EPHX2* | ENSG00000120915.14 | 2.80 | 3.65 | -0.85 | 0.55 | 0.000433499 | 0.003413192 |
| 134 | *ADORA1* | ENSG00000163485.17 | 2.34 | 3.20 | -0.85 | 0.55 | 0.000396567 | 0.003191462 |
| 101928601 | *MEI4* | ENSG00000269964.4 | 3.54 | 4.40 | -0.85 | 0.55 | 3.21164E-05 | 0.000514995 |
| 5565 | *PRKAB2* | ENSG00000131791.8 | 4.29 | 5.15 | -0.85 | 0.55 | 0.000130243 | 0.001410398 |
| 11092 | *SPACA9* | ENSG00000165698.16 | 2.90 | 3.76 | -0.86 | 0.55 | 2.70715E-05 | 0.000454492 |
| 64425 | *POLR1E* | ENSG00000137054.16 | 5.61 | 6.46 | -0.86 | 0.55 | 4.83408E-09 | 1.31918E-06 |
| 59269 | *HIVEP3* | ENSG00000127124.16 | 2.65 | 3.51 | -0.86 | 0.55 | 3.52067E-05 | 0.000552325 |
| 83478 | *ARHGAP24* | ENSG00000138639.18 | 3.17 | 4.03 | -0.86 | 0.55 | 0.000497164 | 0.003775165 |
| 27034 | *ACAD8* | ENSG00000151498.12 | 3.80 | 4.66 | -0.87 | 0.55 | 7.21135E-06 | 0.000167806 |
| 1589 | *CYP21A2* | ENSG00000231852.9 | 7.28 | 8.15 | -0.87 | 0.55 | 8.40837E-08 | 8.23623E-06 |
| 114787 | *GPRIN1* | ENSG00000169258.7 | 4.13 | 5.00 | -0.87 | 0.55 | 2.61224E-06 | 8.23678E-05 |
| 128710 | *SLX4IP* | ENSG00000149346.15 | 2.77 | 3.63 | -0.87 | 0.55 | 0.000109067 | 0.001243911 |
| 91947 | *ARRDC4* | ENSG00000140450.9 | 4.72 | 5.59 | -0.87 | 0.55 | 1.67085E-07 | 1.34827E-05 |
| 6427 | *SRSF2* | ENSG00000161547.17 | 6.82 | 7.69 | -0.87 | 0.55 | 7.18254E-09 | 1.63338E-06 |
| 162282 | *ANKFN1* | ENSG00000153930.13 | 6.49 | 7.36 | -0.87 | 0.55 | 5.68765E-09 | 1.47183E-06 |
| 79634 | *SCRN3* | ENSG00000144306.15 | 3.50 | 4.38 | -0.87 | 0.55 | 3.40697E-05 | 0.000540542 |
| 102724532 | *SP2-DT* | ENSG00000264920.3 | 1.18 | 2.06 | -0.87 | 0.55 | 0.001457676 | 0.008397458 |
| 2048 | *EPHB2* | ENSG00000133216.17 | 4.85 | 5.73 | -0.88 | 0.55 | 1.39857E-06 | 5.58276E-05 |
| 5311 | *PKD2* | ENSG00000118762.8 | 4.49 | 5.36 | -0.88 | 0.54 | 1.63749E-06 | 6.14428E-05 |
| 8941 | *CDK5R2* | ENSG00000171450.6 | 1.03 | 1.91 | -0.88 | 0.54 | 0.000448856 | 0.003508794 |
| 23154 | *NCDN* | ENSG00000020129.16 | 4.26 | 5.14 | -0.88 | 0.54 | 3.47911E-06 | 0.000101394 |
| 79640 | *C22orf46* | ENSG00000184208.12 | 3.09 | 3.97 | -0.88 | 0.54 | 9.14165E-06 | 0.000199241 |
| 2150 | *F2RL1* | ENSG00000164251.5 | 2.21 | 3.09 | -0.88 | 0.54 | 0.000330336 | 0.002795513 |
| 10069 | *RWDD2B* | ENSG00000156253.7 | 3.16 | 4.04 | -0.88 | 0.54 | 5.21949E-05 | 0.000734201 |
| 81704 | *DOCK8* | ENSG00000107099.18 | 6.88 | 7.77 | -0.88 | 0.54 | 5.92861E-09 | 1.48304E-06 |
| 285440 | *CYP4V2* | ENSG00000145476.16 | 3.14 | 4.03 | -0.89 | 0.54 | 0.001310818 | 0.007730479 |
| 65094 | *JMJD4* | ENSG00000081692.13 | 3.07 | 3.96 | -0.89 | 0.54 | 2.24377E-05 | 0.000392502 |
| 399726 | *MIR1915HG* | ENSG00000204682.8 | 2.11 | 3.00 | -0.89 | 0.54 | 0.000175376 | 0.001745506 |
| 8507 | *ENC1* | ENSG00000171617.15 | 2.77 | 3.65 | -0.89 | 0.54 | 0.000777226 | 0.005240512 |
| 10999 | *SLC27A4* | ENSG00000167114.13 | 5.83 | 6.72 | -0.89 | 0.54 | 9.31533E-09 | 1.93127E-06 |
| 158572 | *USP27X-DT* | ENSG00000234390.4 | 2.34 | 3.23 | -0.89 | 0.54 | 7.42927E-05 | 0.000946355 |
| 100507266 | *STX18-AS1* | ENSG00000247708.8 | 2.14 | 3.04 | -0.89 | 0.54 | 0.000334275 | 0.002820202 |
| 84993 | *UBL7* | ENSG00000138629.16 | 5.31 | 6.21 | -0.89 | 0.54 | 1.15971E-06 | 4.88934E-05 |
| 157273 | *LOC157273* | ENSG00000248538.9 | 1.99 | 2.88 | -0.90 | 0.54 | 0.000184352 | 0.001809639 |
| 51276 | *ZNF571* | ENSG00000180479.14 | 2.58 | 3.48 | -0.90 | 0.54 | 0.000202184 | 0.001943997 |
| 4053 | *LTBP2* | ENSG00000119681.12 | 2.11 | 3.01 | -0.90 | 0.54 | 0.000245571 | 0.002263987 |
| 875 | *CBS* | ENSG00000160200.18 | 3.39 | 4.30 | -0.90 | 0.54 | 1.09755E-05 | 0.000230716 |
| 1780 | *DYNC1I1* | ENSG00000158560.14 | 3.45 | 4.36 | -0.90 | 0.53 | 0.0001466 | 0.00154084 |
| 150967 | *LINC01963* | ENSG00000260804.3 | 1.27 | 2.18 | -0.90 | 0.53 | 0.001734462 | 0.009576773 |
| 124152 | *IQCK* | ENSG00000174628.16 | 1.75 | 2.66 | -0.90 | 0.53 | 0.000167306 | 0.001685862 |
| 2939 | *GSTA2* | ENSG00000244067.3 | 3.03 | 3.94 | -0.90 | 0.53 | 1.75775E-05 | 0.000326107 |
| 670 | *BPHL* | ENSG00000137274.13 | 2.98 | 3.89 | -0.91 | 0.53 | 4.35553E-05 | 0.00065047 |
| 8100 | *IFT88* | ENSG00000032742.18 | 3.44 | 4.35 | -0.91 | 0.53 | 4.79211E-05 | 0.000692917 |
| 18 | *ABAT* | ENSG00000183044.12 | 2.90 | 3.81 | -0.91 | 0.53 | 2.6521E-05 | 0.00044927 |
| 140738 | *TMEM37* | ENSG00000171227.7 | 1.67 | 2.57 | -0.91 | 0.53 | 0.000275905 | 0.002460518 |
| 100129518 | *SOD2-OT1* | ENSG00000285427.1 | 0.87 | 1.78 | -0.91 | 0.53 | 0.001540596 | 0.00876528 |
| 114987 | *WDR31* | ENSG00000148225.16 | 2.79 | 3.70 | -0.91 | 0.53 | 1.88061E-05 | 0.000342965 |
| 4968 | *OGG1* | ENSG00000114026.22 | 3.24 | 4.15 | -0.91 | 0.53 | 1.37849E-06 | 5.51726E-05 |
| 10799 | *RPP40* | ENSG00000124787.14 | 1.88 | 2.79 | -0.91 | 0.53 | 0.000123629 | 0.001362371 |
| 7771 | *ZNF112* | ENSG00000062370.17 | 2.49 | 3.40 | -0.91 | 0.53 | 0.000107751 | 0.001233592 |
| 6720 | *SREBF1* | ENSG00000072310.18 | 6.42 | 7.33 | -0.92 | 0.53 | 1.54995E-06 | 5.91941E-05 |
| 9619 | *ABCG1* | ENSG00000160179.19 | 1.68 | 2.60 | -0.92 | 0.53 | 0.000454106 | 0.003535104 |
| 2395 | *FXN* | ENSG00000165060.15 | 3.54 | 4.46 | -0.92 | 0.53 | 2.20803E-05 | 0.000388059 |
| 100302692 | *FTX* | ENSG00000230590.12 | 2.77 | 3.68 | -0.92 | 0.53 | 0.000533077 | 0.003978594 |
| 1153 | *CIRBP* | ENSG00000099622.14 | 6.65 | 7.57 | -0.92 | 0.53 | 9.96928E-07 | 4.5069E-05 |
| 64881 | *PCDH20* | ENSG00000280165.1 | 3.06 | 3.98 | -0.92 | 0.53 | 2.20111E-06 | 7.44066E-05 |
| 9515 | *STXBP5L* | ENSG00000145087.13 | 4.42 | 5.34 | -0.92 | 0.53 | 3.39776E-05 | 0.000539898 |
| 3992 | *FADS1* | ENSG00000149485.19 | 5.01 | 5.94 | -0.92 | 0.53 | 3.41073E-07 | 2.13299E-05 |
| 28988 | *DBNL* | ENSG00000136279.21 | 6.75 | 7.68 | -0.92 | 0.53 | 8.17212E-08 | 8.12287E-06 |
| 10402 | *ST3GAL6* | ENSG00000064225.13 | 2.04 | 2.95 | -0.92 | 0.53 | 0.000559956 | 0.004125864 |
| 9079 | *LDB2* | ENSG00000169744.13 | 4.24 | 5.16 | -0.93 | 0.53 | 1.45688E-05 | 0.000284166 |
| 23541 | *SEC14L2* | ENSG00000100003.18 | 1.56 | 2.49 | -0.93 | 0.53 | 0.000655778 | 0.004618759 |
| 440104 | *TMEM198B* | ENSG00000182796.15 | 3.72 | 4.65 | -0.93 | 0.53 | 7.69507E-05 | 0.00097227 |
| 150737 | *TTC30B* | ENSG00000196659.10 | 3.93 | 4.86 | -0.93 | 0.52 | 2.20262E-07 | 1.59706E-05 |
| 126526 | *C19orf47* | ENSG00000160392.14 | 3.45 | 4.37 | -0.93 | 0.52 | 0.000935006 | 0.005986991 |
| 54487 | *DGCR8* | ENSG00000128191.16 | 5.05 | 5.98 | -0.93 | 0.52 | 0.000205522 | 0.001971042 |
| 4919 | *ROR1* | ENSG00000185483.13 | 2.83 | 3.77 | -0.94 | 0.52 | 4.85466E-05 | 0.000699938 |
| 8187 | *ZNF239* | ENSG00000196793.14 | 0.78 | 1.71 | -0.94 | 0.52 | 0.000970621 | 0.006141676 |
| 144571 | *A2M-AS1* | ENSG00000245105.4 | 1.95 | 2.89 | -0.94 | 0.52 | 3.07093E-05 | 0.000499909 |
| 58499 | *ZNF462* | ENSG00000148143.13 | 5.92 | 6.87 | -0.95 | 0.52 | 0.000172678 | 0.001725519 |
| 84253 | *GARNL3* | ENSG00000136895.19 | 2.21 | 3.17 | -0.95 | 0.52 | 0.000597116 | 0.004327431 |
| 1543 | *CYP1A1* | ENSG00000140465.15 | 3.33 | 4.28 | -0.95 | 0.52 | 0.000425112 | 0.003367019 |
| 1010 | *CDH12* | ENSG00000154162.15 | 3.99 | 4.94 | -0.95 | 0.52 | 1.84631E-05 | 0.000337942 |
| 80298 | *MTERF2* | ENSG00000120832.10 | 3.40 | 4.36 | -0.95 | 0.52 | 2.39974E-05 | 0.000413521 |
| 57821 | *CCDC181* | ENSG00000117477.13 | 1.27 | 2.21 | -0.95 | 0.52 | 0.000935995 | 0.005988215 |
| 80206 | *FHOD3* | ENSG00000134775.16 | 3.80 | 4.76 | -0.96 | 0.52 | 2.36215E-05 | 0.000408922 |
| 54982 | *CLN6* | ENSG00000128973.13 | 5.23 | 6.19 | -0.96 | 0.52 | 3.20375E-06 | 9.61703E-05 |
| 440173 | *LINC02893* | ENSG00000269994.3 | 3.34 | 4.29 | -0.96 | 0.51 | 3.74489E-06 | 0.000106251 |
| 55930 | *MYO5C* | ENSG00000128833.13 | 3.84 | 4.81 | -0.97 | 0.51 | 0.000108535 | 0.001240677 |
| 57545 | *CC2D2A* | ENSG00000048342.18 | 3.26 | 4.23 | -0.97 | 0.51 | 4.77092E-05 | 0.000691184 |

| | | | | | | | | |
|---|---|---|---|---|---|---|---|---|
| 7412 | *VCAM1* | ENSG00000162692.12 | 6.76 | 7.73 | -0.98 | 0.51 | 8.00092E-09 | 1.74037E-06 |
| 134145 | *ATPSCKMT* | ENSG00000150756.14 | 1.88 | 2.86 | -0.98 | 0.51 | 0.000216041 | 0.002039349 |
| 19 | *ABCA1* | ENSG00000165029.17 | 3.79 | 4.77 | -0.98 | 0.51 | 1.5039E-05 | 0.000290684 |
| 55332 | *DRAM1* | ENSG00000136048.14 | 2.25 | 3.23 | -0.98 | 0.51 | 0.000248668 | 0.002288327 |
| 745 | *MYRF* | ENSG00000124920.14 | 2.41 | 3.39 | -0.98 | 0.51 | 0.000159741 | 0.001635438 |
| 163589 | *TDRD5* | ENSG00000162782.16 | 1.29 | 2.28 | -0.98 | 0.51 | 0.000946103 | 0.006024636 |
| 389840 | *MAP3K15* | ENSG00000180815.15 | 2.66 | 3.65 | -0.98 | 0.51 | 1.52709E-06 | 5.86194E-05 |
| 80320 | *SP6* | ENSG00000189120.5 | 4.14 | 5.13 | -0.98 | 0.51 | 4.349E-08 | 5.18389E-06 |
| 169834 | *ZNF883* | ENSG00000228623.7 | 2.76 | 3.74 | -0.99 | 0.51 | 0.000181309 | 0.001790305 |
| 29075 | *LINC00652* | ENSG00000179935.10 | 0.53 | 1.51 | -0.99 | 0.50 | 0.001205792 | 0.007271087 |
| 50617 | *ATP6V0A4* | ENSG00000105929.16 | 0.48 | 1.47 | -0.99 | 0.50 | 0.000119698 | 0.001338711 |
| 1187 | *CLCNKA* | ENSG00000186510.12 | 1.39 | 2.37 | -0.99 | 0.50 | 0.0013708 | 0.008024314 |
| 101927761 | *TH2LCRR* | ENSG00000223442.1 | 4.01 | 5.01 | -0.99 | 0.50 | 1.01886E-07 | 9.61765E-06 |
| 79818 | *ZNF552* | ENSG00000178935.5 | 1.67 | 2.67 | -0.99 | 0.50 | 6.14611E-05 | 0.000829558 |
| 286016 | *TPI1P2* | ENSG00000230359.5 | 0.68 | 1.67 | -0.99 | 0.50 | 0.00056103 | 0.004127696 |
| 64220 | *STRA6* | ENSG00000137868.19 | 2.06 | 3.05 | -0.99 | 0.50 | 5.40308E-05 | 0.000753669 |
| 54843 | *SYTL2* | ENSG00000137501.18 | 7.92 | 8.92 | -0.99 | 0.50 | 1.07738E-09 | 5.41117E-07 |
| 2304 | *FOXE1* | ENSG00000178919.9 | 1.86 | 2.86 | -1.00 | 0.50 | 0.000109315 | 0.001245791 |
| 100009676 | *ZBTB11-AS1* | ENSG00000256628.3 | 1.85 | 2.85 | -1.00 | 0.50 | 5.54572E-06 | 0.00013919 |
| 728130 | *NUTM2D* | ENSG00000214562.15 | 2.62 | 3.61 | -1.00 | 0.50 | 2.1337E-05 | 0.000378096 |
| 136895 | *C7orf31* | ENSG00000153790.12 | 2.44 | 3.43 | -1.00 | 0.50 | 0.001432879 | 0.008293896 |
| 26220 | *DGCR5* | ENSG00000273032.3 | 3.38 | 4.38 | -1.00 | 0.50 | 2.3805E-06 | 7.78557E-05 |
| 339201 | *ASB16-AS1* | ENSG00000267080.7 | 2.04 | 3.04 | -1.00 | 0.50 | 0.001112936 | 0.006826344 |
| 10900 | *RUNDC3A* | ENSG00000108309.14 | 1.83 | 2.84 | -1.01 | 0.50 | 3.09632E-05 | 0.000502952 |
| 36 | *ACADSB* | ENSG00000196177.13 | 4.79 | 5.80 | -1.01 | 0.50 | 6.27524E-08 | 6.54063E-06 |
| 2925 | *GRPR* | ENSG00000126010.6 | 1.04 | 2.06 | -1.01 | 0.50 | 0.001586255 | 0.008974031 |
| 92170 | *MTG1* | ENSG00000148824.19 | 3.54 | 4.54 | -1.01 | 0.50 | 0.000464015 | 0.003591748 |
| 8745 | *ADAM23* | ENSG00000114948.13 | 3.29 | 4.30 | -1.02 | 0.49 | 1.60924E-05 | 0.000305037 |
| 256764 | *WDR72* | ENSG00000166415.15 | 1.80 | 2.82 | -1.02 | 0.49 | 0.000719224 | 0.004951762 |
| 79159 | *NOL12* | ENSG00000273899.5 | 2.54 | 3.55 | -1.02 | 0.49 | 0.000147854 | 0.001551843 |
| 50861 | *STMN3* | ENSG00000197457.10 | 4.51 | 5.53 | -1.02 | 0.49 | 2.60951E-08 | 3.62649E-06 |
| 284618 | *RUSC1-AS1* | ENSG00000225855.7 | 0.35 | 1.36 | -1.02 | 0.49 | 0.000935614 | 0.005988215 |
| 64429 | *ZDHHC6* | ENSG00000023041.12 | 4.58 | 5.61 | -1.03 | 0.49 | 1.32193E-07 | 1.12732E-05 |
| 285600 | *KIAA0825* | ENSG00000185261.15 | 0.30 | 1.32 | -1.03 | 0.49 | 0.001018224 | 0.006376652 |
| 29896 | *TRA2A* | ENSG00000164548.12 | 5.03 | 6.05 | -1.03 | 0.49 | 1.73756E-07 | 1.37258E-05 |
| 197342 | *EME2* | ENSG00000197774.14 | 1.28 | 2.31 | -1.03 | 0.49 | 0.000631124 | 0.004495746 |
| 84457 | *PHYHIPL* | ENSG00000165443.12 | 3.73 | 4.76 | -1.03 | 0.49 | 8.86809E-07 | 4.12078E-05 |
| 164633 | *CABP7* | ENSG00000100314.4 | 0.85 | 1.89 | -1.03 | 0.49 | 0.000507849 | 0.003836088 |
| 285368 | *PRRT3* | ENSG00000163704.12 | 3.85 | 4.88 | -1.03 | 0.49 | 3.69029E-05 | 0.000572186 |
| 101929780 | *NBPF25P* | ENSG00000272150.5 | 0.18 | 1.22 | -1.03 | 0.49 | 0.00170931 | 0.009483932 |
| 79414 | *LRFN3* | ENSG00000126243.9 | 4.43 | 5.47 | -1.04 | 0.49 | 9.1598E-05 | 0.001093711 |
| 9068 | *ANGPTL1* | ENSG00000116194.13 | 2.79 | 3.83 | -1.04 | 0.49 | 0.000284125 | 0.002514403 |
| 115207 | *KCTD12* | ENSG00000178695.6 | 2.17 | 3.22 | -1.04 | 0.49 | 7.6149E-06 | 0.000175564 |
| 729013 | *ZBED5-AS1* | ENSG00000247271.8 | 0.91 | 1.95 | -1.04 | 0.49 | 0.000121056 | 0.001345942 |
| 5608 | *MAP2K6* | ENSG00000108984.15 | 3.09 | 4.14 | -1.04 | 0.49 | 4.50329E-05 | 0.000664015 |
| 282679 | *AQP11* | ENSG00000178301.4 | 2.33 | 3.38 | -1.04 | 0.49 | 5.97771E-06 | 0.000147565 |
| 25939 | *SAMHD1* | ENSG00000101347.11 | 6.53 | 7.58 | -1.05 | 0.48 | 1.41341E-08 | 2.38358E-06 |
| 55359 | *STYK1* | ENSG00000060140.9 | 2.72 | 3.78 | -1.05 | 0.48 | 6.08246E-06 | 0.000149115 |
| 1122 | *CHML* | ENSG00000203668.3 | 4.74 | 5.80 | -1.05 | 0.48 | 9.38904E-07 | 4.29634E-05 |
| 85376 | *RIMBP3* | ENSG00000275793.1 | 1.49 | 2.55 | -1.05 | 0.48 | 0.00037458 | 0.003068684 |
| 9187 | *SLC24A1* | ENSG00000074621.14 | 2.65 | 3.72 | -1.06 | 0.48 | 0.000283694 | 0.002513559 |
| 55329 | *MNS1* | ENSG00000138587.6 | 1.53 | 2.59 | -1.06 | 0.48 | 4.39562E-05 | 0.000655152 |
| 8536 | *CAMK1* | ENSG00000134072.11 | 3.29 | 4.35 | -1.06 | 0.48 | 6.57492E-06 | 0.0001584 |
| 135932 | *TMEM139* | ENSG00000178826.11 | 0.13 | 1.20 | -1.06 | 0.48 | 0.000223187 | 0.002093633 |
| 148213 | *ZNF681* | ENSG00000196172.9 | 1.44 | 2.50 | -1.06 | 0.48 | 0.000626695 | 0.004470565 |
| 255082 | *CASC2* | ENSG00000177640.16 | 1.93 | 3.00 | -1.07 | 0.48 | 6.47514E-05 | 0.000862337 |
| 1462 | *VCAN* | ENSG00000038427.16 | 5.65 | 6.72 | -1.07 | 0.48 | 0.000691613 | 0.004807978 |
| 170712 | *COX7B2* | ENSG00000170516.17 | 0.51 | 1.58 | -1.07 | 0.48 | 0.000227204 | 0.002122967 |
| 4674 | *NAP1L2* | ENSG00000186462.9 | 3.34 | 4.42 | -1.07 | 0.48 | 9.48301E-06 | 0.000204204 |
| 151647 | *TAFA4* | ENSG00000163377.16 | 6.63 | 7.70 | -1.07 | 0.47 | 9.70003E-09 | 1.93127E-06 |
| 390927 | *ZNF793* | ENSG00000188227.14 | 1.63 | 2.71 | -1.08 | 0.47 | 0.000379486 | 0.003085431 |
| 56124 | *PCDHB12* | ENSG00000120328.6 | 0.65 | 1.73 | -1.08 | 0.47 | 0.000721014 | 0.004959532 |
| 100507516 | *LOC10050751* | ENSG00000237807.4 | 1.08 | 2.16 | -1.08 | 0.47 | 8.94434E-05 | 0.001077648 |
| 23245 | *ASTN2* | ENSG00000148219.18 | 1.56 | 2.63 | -1.08 | 0.47 | 7.13222E-05 | 0.000921456 |
| 51725 | *FBXO40* | ENSG00000163833.8 | 3.84 | 4.93 | -1.09 | 0.47 | 1.02846E-07 | 9.64763E-06 |
| 653689 | *GSTT2B* | ENSG00000133433.11 | 5.82 | 6.91 | -1.09 | 0.47 | 1.43076E-08 | 2.38602E-06 |
| 100101467 | *ZSCAN30* | ENSG00000186814.14 | 3.30 | 4.40 | -1.09 | 0.47 | 8.12701E-06 | 0.000184536 |
| 442213 | *PTCHD4* | ENSG00000244694.8 | 3.01 | 4.10 | -1.09 | 0.47 | 6.00689E-07 | 3.19953E-05 |
| 404281 | *YY2* | ENSG00000230797.3 | 1.26 | 2.36 | -1.10 | 0.47 | 9.04499E-05 | 0.001084817 |
| 51286 | *CEND1* | ENSG00000184524.6 | -0.35 | 0.75 | -1.10 | 0.47 | 0.000752273 | 0.005120576 |
| 55511 | *SAGE1* | ENSG00000181433.11 | 1.38 | 2.48 | -1.10 | 0.47 | 5.07615E-05 | 0.000721473 |
| 83451 | *ABHD11* | ENSG00000106077.19 | 1.79 | 2.89 | -1.10 | 0.47 | 0.000563898 | 0.004141422 |
| 202243 | *CCDC125* | ENSG00000183323.13 | 2.56 | 3.67 | -1.10 | 0.47 | 0.000565806 | 0.004152655 |
| 1577 | *CYP3A5* | ENSG00000106258.15 | 2.74 | 3.84 | -1.10 | 0.47 | 0.000101927 | 0.001185906 |
| 285908 | *LINC00174* | ENSG00000179406.9 | 2.30 | 3.40 | -1.11 | 0.46 | 8.47068E-06 | 0.000188845 |
| 245812 | *CNPY4* | ENSG00000166997.8 | 2.97 | 4.08 | -1.11 | 0.46 | 6.75473E-07 | 3.45161E-05 |
| 51308 | *REEP2* | ENSG00000132563.17 | 1.63 | 2.74 | -1.12 | 0.46 | 0.001424086 | 0.008258929 |
| 27300 | *ZNF544* | ENSG00000198131.15 | 2.13 | 3.24 | -1.12 | 0.46 | 0.000466802 | 0.003607332 |
| 55026 | *TMEM255A* | ENSG00000125355.16 | 0.80 | 1.92 | -1.12 | 0.46 | 0.001836079 | 0.009991919 |
| 51167 | *CYB5R4* | ENSG00000065615.14 | 2.74 | 3.86 | -1.12 | 0.46 | 0.001005786 | 0.006316254 |
| 55672 | *NBPF1* | ENSG00000219481.11 | 6.94 | 8.06 | -1.12 | 0.46 | 1.76844E-06 | 6.45804E-05 |
| 203238 | *CCDC171* | ENSG00000164989.17 | 0.32 | 1.45 | -1.12 | 0.46 | 0.000860054 | 0.005664131 |
| 400591 | *TMEM132E-D* | ENSG00000197322.4 | -0.35 | 0.77 | -1.12 | 0.46 | 0.001056622 | 0.006553237 |
| 157983 | *DOCK8-AS1* | ENSG00000183784.7 | -1.01 | 0.11 | -1.12 | 0.46 | 0.001426576 | 0.008270174 |
| 100289635 | *ZNF605* | ENSG00000196458.11 | 3.09 | 4.21 | -1.13 | 0.46 | 0.000678562 | 0.004736405 |
| 256643 | *BCLAF3* | ENSG00000173681.17 | 2.02 | 3.15 | -1.13 | 0.46 | 1.79672E-05 | 0.000332105 |
| 51306 | *FAM13B* | ENSG00000031003.11 | 5.93 | 7.06 | -1.13 | 0.46 | 1.77237E-07 | 1.38667E-05 |
| 11182 | *SLC2A6* | ENSG00000160326.14 | 1.29 | 2.43 | -1.14 | 0.45 | 0.000497268 | 0.003775165 |
| 26052 | *DNM3* | ENSG00000197959.15 | 5.95 | 7.09 | -1.14 | 0.45 | 3.41038E-08 | 4.41261E-06 |

| | | | | | | | | |
|---|---|---|---|---|---|---|---|---|
| 6855 | *SYP* | ENSG00000102003.12 | 4.64 | 5.78 | -1.14 | 0.45 | 3.87017E-08 | 4.76126E-06 |
| 3909 | *LAMA3* | ENSG00000053747.17 | 2.33 | 3.47 | -1.14 | 0.45 | 1.23538E-05 | 0.000250566 |
| 147495 | *APCDD1* | ENSG00000154856.13 | 6.77 | 7.91 | -1.14 | 0.45 | 1.06168E-07 | 9.89059E-06 |
| 56967 | *C14orf132* | ENSG00000227051.7 | 1.91 | 3.07 | -1.15 | 0.45 | 0.001560108 | 0.008866208 |
| 9540 | *TP53I3* | ENSG00000115129.14 | 1.41 | 2.56 | -1.15 | 0.45 | 0.000338054 | 0.002837726 |
| 100303728 | *SLC25A5-AS1* | ENSG00000224281.5 | 0.13 | 1.28 | -1.15 | 0.45 | 0.000316732 | 0.002707197 |
| 387640 | *SKIDA1* | ENSG00000180592.17 | 1.31 | 2.48 | -1.16 | 0.45 | 0.000126895 | 0.001383123 |
| 55959 | *SULF2* | ENSG00000196562.15 | 3.81 | 4.97 | -1.16 | 0.45 | 4.95685E-07 | 2.8396E-05 |
| 93517 | *SDR42E1* | ENSG00000184860.10 | 2.52 | 3.68 | -1.16 | 0.45 | 0.000337133 | 0.002834252 |
| 1947 | *EFNB1* | ENSG00000090776.6 | 2.49 | 3.65 | -1.16 | 0.45 | 1.0627E-06 | 4.63967E-05 |
| 22885 | *ABLIM3* | ENSG00000173210.20 | 2.55 | 3.71 | -1.16 | 0.45 | 7.66856E-07 | 3.71282E-05 |
| 84532 | *ACSS1* | ENSG00000154930.15 | 6.16 | 7.32 | -1.16 | 0.45 | 1.36959E-09 | 6.22915E-07 |
| 185 | *AGTR1* | ENSG00000144891.19 | 5.18 | 6.34 | -1.16 | 0.45 | 1.11764E-09 | 5.41117E-07 |
| 728723 | *ZBED3-AS1* | ENSG00000250802.7 | 0.80 | 1.95 | -1.16 | 0.45 | 0.000455474 | 0.003538414 |
| 10332 | *CLEC4M* | ENSG00000104938.18 | 0.51 | 1.68 | -1.18 | 0.44 | 0.000131194 | 0.001414578 |
| 4744 | *NEFH* | ENSG00000100285.10 | 1.96 | 3.14 | -1.18 | 0.44 | 4.05765E-05 | 0.000614544 |
| 399948 | *COLCA1* | ENSG00000196167.10 | 2.92 | 4.11 | -1.18 | 0.44 | 1.00672E-06 | 4.5375E-05 |
| 100288842 | *B3GALT9* | ENSG00000214654.9 | 1.77 | 2.96 | -1.18 | 0.44 | 1.27143E-05 | 0.000256836 |
| 100506930 | *LINC00665* | ENSG00000232677.10 | 4.10 | 5.28 | -1.18 | 0.44 | 2.98567E-08 | 4.07381E-06 |
| 5935 | *RBM3* | ENSG00000102317.18 | 5.64 | 6.82 | -1.18 | 0.44 | 9.58261E-08 | 9.16086E-06 |
| 1837 | *DTNA* | ENSG00000134769.23 | 4.18 | 5.36 | -1.19 | 0.44 | 1.27581E-06 | 5.2606E-05 |
| 219931 | *TPCN2* | ENSG00000162341.18 | 1.72 | 2.91 | -1.20 | 0.44 | 0.00010351 | 0.001196904 |
| 503569 | *RGMB-AS1* | ENSG00000246763.7 | -1.17 | 0.03 | -1.20 | 0.44 | 0.001409717 | 0.008204124 |
| 283601 | *LINC00523* | ENSG00000196273.8 | 0.53 | 1.73 | -1.20 | 0.44 | 0.0005833 | 0.004254005 |
| 4856 | *CCN3* | ENSG00000136999.5 | 7.18 | 8.38 | -1.20 | 0.44 | 9.10729E-11 | 1.29661E-07 |
| 10202 | *DHRS2* | ENSG00000100867.15 | 0.43 | 1.64 | -1.21 | 0.43 | 0.000181873 | 0.001794692 |
| 285848 | *PNPLA1* | ENSG00000180316.13 | 2.25 | 3.46 | -1.21 | 0.43 | 0.000201759 | 0.001942402 |
| 374378 | *GALNT18* | ENSG00000110328.6 | 1.12 | 2.34 | -1.22 | 0.43 | 4.16727E-06 | 0.000113721 |
| 444882 | *IGFL4* | ENSG00000204869.9 | 0.27 | 1.49 | -1.22 | 0.43 | 0.000588328 | 0.00428028 |
| 83873 | *GPR61* | ENSG00000156097.13 | 2.66 | 3.88 | -1.22 | 0.43 | 2.80383E-05 | 0.000466548 |
| 100507437 | *LOC1005074[illegible]* | ENSG00000251602.7 | 1.38 | 2.61 | -1.22 | 0.43 | 3.80103E-05 | 0.00058333 |
| 1795 | *DOCK3* | ENSG00000088538.13 | 2.19 | 3.41 | -1.23 | 0.43 | 6.68493E-06 | 0.000160278 |
| 163131 | *ZNF780B* | ENSG00000128000.17 | 2.69 | 3.91 | -1.23 | 0.43 | 0.000913738 | 0.005883441 |
| 57573 | *ZNF471* | ENSG00000196263.8 | 2.35 | 3.59 | -1.23 | 0.43 | 0.000865067 | 0.00567858 |
| 113 | *ADCY7* | ENSG00000121281.13 | 0.90 | 2.14 | -1.24 | 0.42 | 4.85156E-05 | 0.000699938 |
| 441046 | *GUSBP5* | ENSG00000236296.8 | 0.88 | 2.12 | -1.24 | 0.42 | 5.1372E-05 | 0.000727041 |
| 389799 | *CFAP77* | ENSG00000188523.9 | 0.41 | 1.65 | -1.24 | 0.42 | 0.000273943 | 0.002448258 |
| 729262 | *NUTM2B* | ENSG00000188199.11 | 0.81 | 2.05 | -1.24 | 0.42 | 2.58516E-05 | 0.000439419 |
| 114898 | *C1QTNF2* | ENSG00000145861.9 | 0.03 | 1.27 | -1.24 | 0.42 | 0.000383935 | 0.003113174 |
| 100507433 | *ZNF571-AS1* | ENSG00000267470.6 | 2.04 | 3.30 | -1.25 | 0.42 | 0.000911681 | 0.005875236 |
| 388389 | *CCDC103* | ENSG00000167131.17 | 1.91 | 3.16 | -1.25 | 0.42 | 0.00091723 | 0.005895808 |
| 283481 | *FGF14-AS2* | ENSG00000272143.1 | 0.73 | 1.99 | -1.26 | 0.42 | 0.000228572 | 0.002129882 |
| 4168 | *MCF2* | ENSG00000101977.21 | 3.53 | 4.79 | -1.27 | 0.41 | 8.67744E-07 | 4.05731E-05 |
| 176 | *ACAN* | ENSG00000157766.19 | 1.93 | 3.22 | -1.28 | 0.41 | 3.17152E-06 | 9.55849E-05 |
| 79173 | *BRME1* | ENSG00000132016.12 | 1.72 | 3.01 | -1.29 | 0.41 | 0.000865255 | 0.00567858 |
| 283234 | *CCDC88B* | ENSG00000168071.22 | -0.12 | 1.17 | -1.29 | 0.41 | 0.001052737 | 0.006539952 |
| 10544 | *PROCR* | ENSG00000101000.6 | 1.65 | 2.94 | -1.30 | 0.41 | 7.01434E-05 | 0.000913873 |
| 643276 | *NOC2LP1* | ENSG00000213225.7 | 1.99 | 3.29 | -1.30 | 0.41 | 3.23423E-07 | 2.04821E-05 |
| 793 | *CALB1* | ENSG00000104327.7 | 4.43 | 5.73 | -1.30 | 0.41 | 1.16707E-08 | 2.1423E-06 |
| 57002 | *YAE1* | ENSG00000241127.8 | 1.64 | 2.93 | -1.31 | 0.40 | 1.74882E-05 | 0.000325255 |
| 101927417 | *FHAD1-AS1* | ENSG00000233485.1 | 2.89 | 4.21 | -1.31 | 0.40 | 9.30756E-07 | 4.27208E-05 |
| 83758 | *RBP5* | ENSG00000139194.8 | 0.70 | 2.01 | -1.31 | 0.40 | 0.000169783 | 0.0017002 |
| 3284 | *HSD3B2* | ENSG00000203859.10 | 3.80 | 5.11 | -1.31 | 0.40 | 2.01824E-08 | 2.99918E-06 |
| 55277 | *FGGY* | ENSG00000172456.18 | 1.00 | 2.33 | -1.32 | 0.40 | 0.000963984 | 0.006109983 |
| 256933 | *NPB* | ENSG00000183979.8 | -0.64 | 0.68 | -1.32 | 0.40 | 0.001353853 | 0.007953022 |
| 84622 | *ZNF594* | ENSG00000180626.10 | 1.88 | 3.22 | -1.33 | 0.40 | 6.47094E-06 | 0.000156649 |
| 158401 | *SHOC1* | ENSG00000165181.17 | 3.75 | 5.08 | -1.33 | 0.40 | 5.35501E-06 | 0.000135851 |
| 440078 | *FAM66C* | ENSG00000226711.8 | 0.59 | 1.93 | -1.33 | 0.40 | 0.001215192 | 0.007310147 |
| 120379 | *PIH1D2* | ENSG00000150773.12 | 0.05 | 1.40 | -1.33 | 0.40 | 0.001097058 | 0.006745494 |
| 54332 | *GDAP1* | ENSG00000104381.14 | 4.27 | 5.62 | -1.34 | 0.39 | 9.26169E-09 | 1.93127E-06 |
| 55520 | *ELAC1* | ENSG00000141642.9 | 1.77 | 3.12 | -1.35 | 0.39 | 4.58579E-05 | 0.000672149 |
| 201625 | *DNAH12* | ENSG00000174844.15 | 1.77 | 3.11 | -1.35 | 0.39 | 0.000126399 | 0.001379722 |
| 101927671 | *MIR202HG* | ENSG00000166917.11 | -1.02 | 0.32 | -1.35 | 0.39 | 0.001007382 | 0.00632271 |
| 105369364 | *LINC02701* | ENSG00000250508.1 | 3.30 | 4.66 | -1.35 | 0.39 | 5.66542E-08 | 6.12322E-06 |
| 137209 | *ZNF572* | ENSG00000180938.6 | 0.87 | 2.24 | -1.36 | 0.39 | 0.000251722 | 0.002306526 |
| 283008 | *NUTM2E* | ENSG00000228570.8 | 0.57 | 1.94 | -1.37 | 0.39 | 5.28777E-05 | 0.000739647 |
| 23284 | *ADGRL3* | ENSG00000150471.17 | 2.72 | 4.09 | -1.37 | 0.39 | 1.40589E-07 | 1.1594E-05 |
| 284443 | *ZNF493* | ENSG00000196268.12 | 1.20 | 2.56 | -1.37 | 0.39 | 0.001464132 | 0.008422826 |
| 3747 | *KCNC2* | ENSG00000166006.14 | 1.58 | 2.96 | -1.38 | 0.38 | 1.34549E-06 | 5.44324E-05 |
| 283726 | *SAXO2* | ENSG00000188659.10 | -0.60 | 0.78 | -1.38 | 0.38 | 0.001014617 | 0.006358408 |
| 3780 | *KCNN1* | ENSG00000105642.16 | 1.87 | 3.25 | -1.39 | 0.38 | 2.66469E-05 | 0.000450664 |
| 167681 | *PRSS35* | ENSG00000146250.7 | -0.03 | 1.35 | -1.39 | 0.38 | 0.000272196 | 0.00243904 |
| 3995 | *FADS3* | ENSG00000221968.9 | 3.91 | 5.31 | -1.40 | 0.38 | 6.20817E-08 | 6.51597E-06 |
| 51385 | *ZNF589* | ENSG00000164048.14 | 2.13 | 3.54 | -1.40 | 0.38 | 3.52267E-05 | 0.000552325 |
| 91409 | *CCDC74B* | ENSG00000152076.19 | 0.33 | 1.73 | -1.40 | 0.38 | 0.000824751 | 0.005489442 |
| 1268 | *CNR1* | ENSG00000118432.13 | 3.14 | 4.54 | -1.40 | 0.38 | 2.04254E-06 | 7.09641E-05 |
| 6004 | *RGS16* | ENSG00000143333.7 | 0.13 | 1.54 | -1.42 | 0.37 | 0.000618608 | 0.004432609 |
| 140706 | *CCM2L* | ENSG00000101331.17 | 0.34 | 1.76 | -1.43 | 0.37 | 0.000212322 | 0.002016922 |
| 340533 | *NEXMIF* | ENSG00000050030.16 | 0.97 | 2.42 | -1.44 | 0.37 | 6.03144E-05 | 0.000819239 |
| 84894 | *LINGO1* | ENSG00000169783.13 | 2.57 | 4.02 | -1.44 | 0.37 | 2.74269E-05 | 0.00045892 |
| 55231 | *CCDC87* | ENSG00000182791.5 | -1.13 | 0.32 | -1.45 | 0.37 | 0.000861786 | 0.005665591 |
| 79187 | *FSD1* | ENSG00000105255.11 | 1.52 | 2.96 | -1.45 | 0.37 | 8.27334E-06 | 0.00018643 |
| 121643 | *FOXN4* | ENSG00000139445.18 | 0.03 | 1.50 | -1.46 | 0.36 | 0.001830222 | 0.009972564 |
| 389602 | *LOC389602* | ENSG00000204876.5 | 1.81 | 3.27 | -1.46 | 0.36 | 6.51922E-07 | 3.37403E-05 |
| 92960 | *PEX11G* | ENSG00000104883.8 | -0.71 | 0.76 | -1.47 | 0.36 | 0.000360661 | 0.002984103 |
| 492 | *ATP2B3* | ENSG00000067842.18 | 3.69 | 5.15 | -1.47 | 0.36 | 4.15872E-08 | 5.03373E-06 |
| 84276 | *NICN1* | ENSG00000145029.14 | 0.52 | 1.99 | -1.47 | 0.36 | 2.70209E-05 | 0.000454151 |
| 89846 | *FGD3* | ENSG00000127084.19 | -0.29 | 1.19 | -1.48 | 0.36 | 0.000336053 | 0.002830428 |
| 140862 | *ISM1* | ENSG00000101230.6 | 4.02 | 5.50 | -1.48 | 0.36 | 6.96419E-10 | 4.18102E-07 |

| | | | | | | | | |
|---|---|---|---|---|---|---|---|---|
| 100996307 | *LIPE-AS1* | ENSG00000213904.10 | 0.86 | 2.34 | -1.48 | 0.36 | 0.000371728 | 0.003050445 |
| 3910 | *LAMA4* | ENSG00000112769.20 | 0.51 | 2.00 | -1.49 | 0.35 | 0.000515302 | 0.003878718 |
| 112268389 | *LOC11226838* | ENSG00000255052.5 | -0.92 | 0.56 | -1.50 | 0.35 | 0.000605985 | 0.004374809 |
| 126755 | *LRRC38* | ENSG00000162494.6 | -1.09 | 0.42 | -1.50 | 0.35 | 0.00035825 | 0.002970706 |
| 7433 | *VIPR1* | ENSG00000114812.13 | 2.13 | 3.64 | -1.51 | 0.35 | 6.34257E-07 | 3.32852E-05 |
| 401207 | *C5orf63* | ENSG00000164241.14 | 0.35 | 1.87 | -1.53 | 0.35 | 0.000493558 | 0.003754596 |
| 283130 | *SLC25A45* | ENSG00000162241.13 | 0.68 | 2.22 | -1.54 | 0.34 | 0.000233583 | 0.002169462 |
| 5138 | *PDE2A* | ENSG00000186642.16 | 5.35 | 6.89 | -1.54 | 0.34 | 3.46541E-08 | 4.4455E-06 |
| 60680 | *CELF5* | ENSG00000161082.13 | 1.33 | 2.88 | -1.54 | 0.34 | 4.78963E-05 | 0.000692917 |
| 129049 | *SGSM1* | ENSG00000167037.19 | 1.30 | 2.87 | -1.55 | 0.34 | 0.000447586 | 0.003502514 |
| 102723775 | *LINC01315* | ENSG00000229891.3 | -1.73 | -0.18 | -1.56 | 0.34 | 0.000749879 | 0.005106597 |
| 101929074 | *PIK3CD-AS2* | ENSG00000231789.3 | -1.69 | -0.12 | -1.57 | 0.34 | 0.001409168 | 0.00820411 |
| 103724390 | *LINC01287* | ENSG00000234722.5 | -1.99 | -0.41 | -1.57 | 0.34 | 0.001711232 | 0.009484448 |
| 4067 | *LYN* | ENSG00000254087.8 | 0.37 | 1.95 | -1.58 | 0.34 | 0.001304866 | 0.00770446 |
| 136306 | *SVOPL* | ENSG00000157703.16 | 0.45 | 2.04 | -1.58 | 0.33 | 0.000313628 | 0.002685246 |
| 105369147 | *LOC10536914* | ENSG00000247081.8 | 0.50 | 2.07 | -1.58 | 0.33 | 6.10124E-05 | 0.000824987 |
| 84536 | *LINC01547* | ENSG00000183250.13 | 1.28 | 2.86 | -1.59 | 0.33 | 7.20547E-06 | 0.000167806 |
| 114769 | *CARD16* | ENSG00000204397.10 | -1.16 | 0.44 | -1.59 | 0.33 | 0.000830239 | 0.005516183 |
| 728411 | *GUSBP1* | ENSG00000215190.9 | 0.52 | 2.13 | -1.60 | 0.33 | 0.000182691 | 0.001800669 |
| 4128 | *MAOA* | ENSG00000189221.11 | -0.54 | 1.06 | -1.62 | 0.33 | 0.000351234 | 0.002928707 |
| 29951 | *PDZRN4* | ENSG00000165966.16 | -0.41 | 1.20 | -1.62 | 0.33 | 0.000296099 | 0.002586815 |
| 643783 | *USP46-DT* | ENSG00000248866.1 | -0.94 | 0.69 | -1.63 | 0.32 | 0.00046996 | 0.003624679 |
| 3291 | *HSD11B2* | ENSG00000176387.7 | -0.79 | 0.83 | -1.64 | 0.32 | 0.000863462 | 0.005674125 |
| 5820 | *PVT1* | ENSG00000249859.13 | -0.75 | 0.88 | -1.64 | 0.32 | 6.82783E-05 | 0.00089815 |
| 133022 | *TRAM1L1* | ENSG00000174599.5 | 0.23 | 1.88 | -1.64 | 0.32 | 0.000324101 | 0.002751377 |
| 100506810 | *LINC01132* | ENSG00000227630.4 | 0.16 | 1.81 | -1.66 | 0.32 | 9.95607E-05 | 0.00116379 |
| 56135 | *PCDHAC1* | ENSG00000248383.5 | -1.03 | 0.62 | -1.66 | 0.32 | 0.000364257 | 0.003005306 |
| 347442 | *DCAF8L2* | ENSG00000189186.11 | 0.85 | 2.53 | -1.68 | 0.31 | 0.000286967 | 0.002529119 |
| 257396 | *MOCS2-DT* | ENSG00000247796.3 | 1.63 | 3.31 | -1.68 | 0.31 | 3.68195E-07 | 2.25561E-05 |
| 283050 | *ZMIZ1-AS1* | ENSG00000224596.9 | -0.57 | 1.12 | -1.70 | 0.31 | 9.17294E-05 | 0.001094409 |
| 100506622 | *LINC00540* | ENSG00000276476.4 | 0.04 | 1.74 | -1.70 | 0.31 | 5.58625E-05 | 0.000771334 |
| 375704 | *ENHO* | ENSG00000168913.7 | -0.37 | 1.34 | -1.71 | 0.31 | 3.20987E-05 | 0.000514995 |
| 101927021 | *ZFAND2A-DT* | ENSG00000229043.3 | 0.03 | 1.74 | -1.71 | 0.31 | 1.20734E-05 | 0.000247555 |
| 5288 | *PIK3C2G* | ENSG00000139144.11 | 0.40 | 2.12 | -1.72 | 0.30 | 0.000157986 | 0.001621899 |
| 57127 | *RHBG* | ENSG00000132677.13 | 0.84 | 2.56 | -1.72 | 0.30 | 2.78095E-07 | 1.91464E-05 |
| 91646 | *TDRD12* | ENSG00000173809.18 | 0.20 | 1.95 | -1.74 | 0.30 | 0.000237763 | 0.002201471 |
| 153657 | *TTC23L* | ENSG00000205838.14 | -0.14 | 1.61 | -1.75 | 0.30 | 0.001301294 | 0.007692449 |
| 200931 | *SLC51A* | ENSG00000163959.10 | -0.83 | 0.91 | -1.76 | 0.30 | 0.00021425 | 0.002028816 |
| 54959 | *ODAM* | ENSG00000109205.17 | -1.21 | 0.57 | -1.77 | 0.29 | 0.000377345 | 0.003075272 |
| 23416 | *KCNH3* | ENSG00000135519.8 | 1.20 | 2.96 | -1.77 | 0.29 | 1.93046E-05 | 0.00034874 |
| 94 | *ACVRL1* | ENSG00000139567.13 | 0.83 | 2.64 | -1.81 | 0.29 | 1.66671E-06 | 6.17669E-05 |
| 653427 | *FOXD4L5* | ENSG00000204779.2 | -1.93 | -0.12 | -1.81 | 0.28 | 0.001728038 | 0.009559944 |
| 55016 | *MARCHF1* | ENSG00000145416.14 | -1.21 | 0.60 | -1.82 | 0.28 | 0.001264787 | 0.007544989 |
| 2900 | *GRIK4* | ENSG00000149403.13 | 2.03 | 3.86 | -1.83 | 0.28 | 1.8826E-08 | 2.92753E-06 |
| 160728 | *SLC5A8* | ENSG00000256870.3 | -2.52 | -0.66 | -1.84 | 0.28 | 0.001758453 | 0.009666591 |
| 7694 | *ZNF135* | ENSG00000176293.20 | -1.26 | 0.57 | -1.85 | 0.28 | 0.000359272 | 0.002977537 |
| 115677 | *NOSTRIN* | ENSG00000163072.16 | 0.75 | 2.58 | -1.85 | 0.28 | 4.11092E-06 | 0.000113457 |
| 339929 | *LPP-AS2* | ENSG00000270959.1 | -0.79 | 1.09 | -1.87 | 0.27 | 1.91002E-05 | 0.000346226 |
| 140711 | *TLDC2* | ENSG00000101342.10 | -1.97 | -0.09 | -1.87 | 0.27 | 0.001459436 | 0.008399032 |
| 100128025 | *WWTR1-AS1* | ENSG00000241313.3 | -2.70 | -0.83 | -1.88 | 0.27 | 0.000517662 | 0.003892578 |
| 174 | *AFP* | ENSG00000081051.8 | 0.44 | 2.31 | -1.89 | 0.27 | 7.9244E-05 | 0.000990319 |
| 25854 | *FAM149A* | ENSG00000109794.14 | -0.85 | 1.05 | -1.89 | 0.27 | 0.000211984 | 0.002014987 |
| 5457 | *POU4F1* | ENSG00000152192.8 | -0.14 | 1.78 | -1.92 | 0.26 | 5.51318E-05 | 0.000765471 |
| 2298 | *FOXD4* | ENSG00000170122.6 | -2.42 | -0.50 | -1.93 | 0.26 | 0.001587481 | 0.008974198 |
| 105371708 | *LINC01992* | ENSG00000260019.2 | 0.54 | 2.48 | -1.94 | 0.26 | 1.03101E-06 | 4.61924E-05 |
| 8911 | *CACNA1I* | ENSG00000100346.18 | -1.58 | 0.40 | -1.99 | 0.25 | 1.83643E-05 | 0.000336955 |
| 375791 | *CYSRT1* | ENSG00000197191.6 | -1.82 | 0.18 | -1.99 | 0.25 | 0.00061981 | 0.004438322 |
| 22844 | *FRMPD1* | ENSG00000070601.10 | -1.39 | 0.64 | -2.04 | 0.24 | 0.001148181 | 0.006988263 |
| 340895 | *MALRD1* | ENSG00000204740.11 | -1.97 | 0.08 | -2.05 | 0.24 | 8.876E-05 | 0.001075335 |
| 9914 | *ATP2C2* | ENSG00000064270.13 | -1.62 | 0.46 | -2.08 | 0.24 | 0.000361096 | 0.002986053 |
| 92340 | *PRR29* | ENSG00000224383.8 | -0.24 | 1.85 | -2.09 | 0.24 | 0.000140857 | 0.001498318 |
| 2953 | *GSTT2* | ENSG00000099984.11 | -0.94 | 1.14 | -2.09 | 0.24 | 2.33078E-05 | 0.000404424 |
| 112267871 | *LOC11226787* | ENSG00000228549.5 | -2.23 | -0.14 | -2.09 | 0.23 | 0.001635969 | 0.009189468 |
| 729747 | *ZNF878* | ENSG00000257446.4 | -2.67 | -0.57 | -2.10 | 0.23 | 0.000868021 | 0.005691627 |
| 729467 | *HSD52* | ENSG00000224609.8 | -2.67 | -0.58 | -2.10 | 0.23 | 0.000140298 | 0.001494042 |
| 105274304 | *LOC1052743* | ENSG00000261335.1 | -1.73 | 0.38 | -2.12 | 0.23 | 0.001619049 | 0.009123252 |
| 196872 | *LINC00638* | ENSG00000258701.1 | -0.04 | 2.13 | -2.16 | 0.22 | 5.49802E-07 | 3.05629E-05 |
| 9971 | *NR1H4* | ENSG00000012504.15 | 0.53 | 2.70 | -2.18 | 0.22 | 0.000108633 | 0.001240845 |
| 7092 | *TLL1* | ENSG00000038295.8 | -1.93 | 0.28 | -2.20 | 0.22 | 0.000556151 | 0.004109932 |
| 221938 | *MMD2* | ENSG00000136297.15 | -1.35 | 0.83 | -2.20 | 0.22 | 0.000229769 | 0.002139335 |
| 100131193 | *CCDC183-AS* | ENSG00000228544.1 | -1.00 | 1.21 | -2.20 | 0.22 | 5.93303E-06 | 0.000146945 |
| 101928728 | *LOC10192872* | ENSG00000260063.1 | -0.82 | 1.39 | -2.23 | 0.21 | 4.0566E-06 | 0.00011317 |
| 728039 | *SSR4P1* | ENSG00000235374.2 | -0.77 | 1.52 | -2.27 | 0.21 | 0.000624332 | 0.004464318 |
| 5148 | *PDE6G* | ENSG00000185527.12 | -2.58 | -0.28 | -2.30 | 0.20 | 1.46747E-05 | 0.000285301 |
| 158056 | *MAMDC4* | ENSG00000177943.14 | -1.68 | 0.64 | -2.33 | 0.20 | 0.00014848 | 0.001554074 |
| 100507462 | *CCDC28A-AS* | ENSG00000279968.3 | -2.97 | -0.65 | -2.33 | 0.20 | 0.000654335 | 0.004612921 |
| 256536 | *TCERG1L* | ENSG00000176769.9 | -0.50 | 1.91 | -2.41 | 0.19 | 4.40417E-05 | 0.000655459 |
| 221458 | *KIF6* | ENSG00000164627.18 | -1.01 | 1.39 | -2.41 | 0.19 | 0.001292132 | 0.007668491 |
| 154075 | *SAMD3* | ENSG00000164483.17 | -0.29 | 2.13 | -2.43 | 0.19 | 0.000678793 | 0.004736405 |
| 101929567 | *LINC01814* | ENSG00000236008.4 | 0.78 | 3.22 | -2.44 | 0.18 | 2.22123E-09 | 7.5769E-07 |
| 654780 | *LOC654780* | ENSG00000250685.7 | -2.92 | -0.34 | -2.56 | 0.17 | 0.000319333 | 0.002723224 |
| 2169 | *FABP2* | ENSG00000145384.4 | -0.62 | 1.96 | -2.57 | 0.17 | 5.36028E-07 | 3.00196E-05 |
| 105376845 | *LOC10537684* | ENSG00000285873.1 | -2.90 | -0.28 | -2.61 | 0.16 | 6.35418E-05 | 0.000848486 |
| 101928489 | *CD109-AS1* | ENSG00000231652.2 | -3.41 | -0.69 | -2.72 | 0.15 | 8.93544E-05 | 0.001077648 |
| 56137 | *PCDHA12* | ENSG00000251664.5 | -2.15 | 0.59 | -2.74 | 0.15 | 0.000123129 | 0.00135885 |
| 8796 | *SCEL* | ENSG00000136155.17 | -0.97 | 1.79 | -2.75 | 0.15 | 0.000168899 | 0.001695658 |
| 100129781 | *MIR193BHG* | ENSG00000262454.4 | -2.61 | 0.19 | -2.83 | 0.14 | 0.000413221 | 0.003305991 |
| 28 | *ABO* | ENSG00000175164.16 | -2.49 | 0.50 | -2.99 | 0.13 | 0.000359873 | 0.002980866 |
| 8328 | *GFI1B* | ENSG00000165702.15 | -1.96 | 1.14 | -3.10 | 0.12 | 9.10373E-07 | 4.21722E-05 |

| | | | | | | | | |
|---|---|---|---|---|---|---|---|---|
| 114794 | *ELFN2* | ENSG00000166897.16 | -2.50 | 0.63 | -3.15 | 0.11 | 0.001447918 | 0.008351959 |
| 146712 | *B3GNTL1* | ENSG00000175711.9 | -2.07 | 1.11 | -3.18 | 0.11 | 2.17653E-05 | 0.00038509 |
| 100506776 | *TRG-AS1* | ENSG00000281103.3 | -3.81 | -0.57 | -3.26 | 0.10 | 0.000839448 | 0.005558638 |
| 101927973 | *LINC01695* | ENSG00000236532.6 | -2.47 | 0.99 | -3.48 | 0.09 | 0.000338054 | 0.002837726 |
| 2563 | *GABRD* | ENSG00000187730.9 | -3.90 | -0.05 | -3.86 | 0.07 | 0.000125746 | 0.001376602 |
| 285220 | *EPHA6* | ENSG00000080224.18 | -3.62 | 0.41 | -4.03 | 0.06 | 0.000272375 | 0.002439188 |
